# pLM representations unlock metagenomic space beyond homology

**DOI:** 10.64898/2026.07.28.739874

**Authors:** Lola Le Breton, David Heurtel-Depeiges, Douglas C. Millar, Lara E. Zetzsche, Robert M. Vernon, Christopher James Langmead, Sarath Chandar, Quentin Fournier

## Abstract

Metagenomic sequencing has uncovered billions of proteins from uncultured microorganisms, vastly expanding the known protein space. Yet most remain functionally inaccessible because existing annotation methods depend on close homologs or accurate structure predictions. Here, we show that protein language models (pLMs) can unlock this diversity only when their training data are appropriately curated. We introduce Residue Embedding Diversity (RED), a metric for protein quality assessment orders of magnitude cheaper than likelihood, and a calibration task that measures model alignment with natural evolutionary distributions. We discover a fundamental trade-off between evolutionary calibration and structural modeling, establishing training data composition as a primary determinant of pLM behavior. Finally, we successfully retrieve diverse enzyme candidates from billions of metagenomic sequences and validate their expression *in vivo*.

## 1. Introduction

Metagenomic sequencing has cataloged billions of protein sequences from uncultured microorganisms, the majority of which have no close homolog in any annotated database (*1*). Standard tools such as homology search and structure prediction are only effective when a query protein has characterized relatives or folds into a confident structural model. As a result, functional characterization and effective retrieval in this space remain fragmentary. The standard response has been to set these sequences aside, missing a vast reservoir of functional diversity generated by the intense evolutionary pressure of the microbial world.

Protein language models (pLMs), trained to recover masked amino acids from unannotated sequences, learn representations that encode co-evolutionary statistics and aspects of biophysical stability without structural supervision (*2*). They successfully infer structural contacts from primary sequences (*3*), predict mutational fitness without homologous alignments (*4*), and generate candidate protein binders (*5*). Because pLMs learn from sequence statistics alone, they can in principle be trained on any corpus of protein sequences, including the billions of unannotated metagenomic assemblies. But like any statistical model, a pLM is shaped by its training distribution. Whether metagenomic data can be incorporated productively and how the composition of pretraining data shapes the resulting representations remains poorly characterized. Moreover, recent work shows diminishing and even negative returns from scaling model parameters (*6, 7, 8*), suggesting that the bottleneck has shifted from model capacity to training data. Understanding how data composition shapes protein representations is therefore a prerequisite for modeling the full natural protein landscape, including metagenomic spaces.

Two obstacles make this difficult in practice. The first is noise: metagenomic assemblies contain frameshifts, fragments, and pseudogenes that dilute the functional signal (*9, 10*), and naively adding them to pretraining degrades downstream performance. Current filtering heuristics rely on length thresholds or identity cutoffs (*11,12*), which cannot capture the long-range dependencies that distinguish functional sequences from genomic artifacts, and no scalable method can assess individual sequence quality across billions of entries. The second is distributional: curated databases such as UniProtKB (*13, 14*) provide dense evolutionary signal but impose anthropocentric biases and remain limited in scale, whereas metagenomic databases provide breadth across the tree of life but sample a fundamentally different region of sequence space. How mixing these distributions affects what a model learns and how these effects impact downstream tasks is not well studied.

Here we address both obstacles. We first show that pLM representations carry a functional signal suitable for candidate selection. Perplexity (PPL), a model-derived score inversely proportional to sequence likelihood normalized by length, aligns with expert-quality annotations: a sequence with higher perplexity is more surprising, or less likely, according to the model, than one with lower perplexity. However, perplexity is computationally expensive because it requires one forward pass per residue. To overcome this limitation, we introduce Residue Embedding Diversity (RED), a scalable metric that requires a single forward pass per sequence. We derive RED from a residue-level representation collapse phenomenon in low-quality sequences. RED reduces scoring costs by three orders of magnitude and provides an unsupervised filtering criterion applicable to massive databases.

Using RED-enabled curation and more than 100 controlled pretraining ablations, we establish how training data composition shapes protein representations. In addition to a broad set of in silico evaluations, we introduce a calibration task that isolates positions where the evolutionary consensus residue differs from the residue found in the training proteome, providing a direct measure of alignment to the natural evolutionary distribution. We uncover a fundamental trade-off: dense homologous coverage favors evolutionary calibration and fitness prediction, whereas sparse and diverse data favor structural modeling. Guided by this discovery, we develop AMPLIFY-B and AMPLIFY-C, specialized models that sit on opposite ends of the trade-off. We validate AMPLIFY-C in a wet-lab assay of Fe/αKG-dependent dioxygenases and show that pLM-guided selection outperforms a control pipeline in which no pLM is used. With pLM embedding retrieval, we recover enzyme candidates from metagenomic databases that share low sequence identity with the annotated enzyme class and experimentally confirm measurable expression for 31.3% of retrieved candidates.

Collectively, these results establish training data composition as a primary determinant of protein language model abilities. We discover a fundamental trade-off between evolutionary calibration and structural modeling, introduce RED as a scalable metric for metagenomic protein curation, and establish calibration as a new task for evaluating pLMs. We further demonstrate that, with appropriate curation, metagenomic data can be transformed from a source of noise into a valuable extension of the accessible protein sequence space.

## 2. Results

### The diversity of residue-level pLM representations is a scalable proxy for perplexity

Protein language models are trained on nothing but evolutionary sequence co-occurrence, and yet their perplexity, or measure of how unlikely a sequence is, aligns with expert annotations. This is tested *in silico* by computing the perplexity under AMPLIFY 350M (*6*) of a broad set of curated proteins from UniProtKB (Figure 1A, Table S17). Well-characterized human proteins (mean PPL = 3.57) and curated intrinsically disordered proteins from DisProt (*15*) (mean PPL = 4.21) both receive low PPL, corresponding to a high likelihood according to the model, despite disordered proteins being structurally atypical and poorly modeled by structure prediction tools. In contrast, shuffled human sequences that preserve amino acid composition but destroy positional syntax receive high perplexity (mean PPL = 17.88). In an uncertainty zone, among sequences with the weakest evidence level in UniProtKB (PE = 5, protein uncertain), we differentiate sequences by annotation count. We find that mean perplexity decreases monotonically with the level of annotation, from 17.00 at annotation level 1 to 4.70 at level 4. The trend reverses at the highest annotation level (mean PPL = 7.14): this set is bimodal, and inspection of the high-perplexity cluster reveals that all of the sequences carry a UniProtKB *Caution* flag identifying it as a likely pseudogene. For instance, YG21B YEAST contains a stop codon disrupting the open reading frame (ORF) coding for protein TY21, as well as a frameshift which disrupts the ORF coding for protein TY2B. From sequence statistics alone, the model recovers a quality assessment that curators reached through independent evidence and scientific publication. That this signal differentiates disordered proteins from ambiguous sequences confirms that pLMs capture evolutionary plausibility in a sense orthogonal to structural foldability.

**Figure 1:**
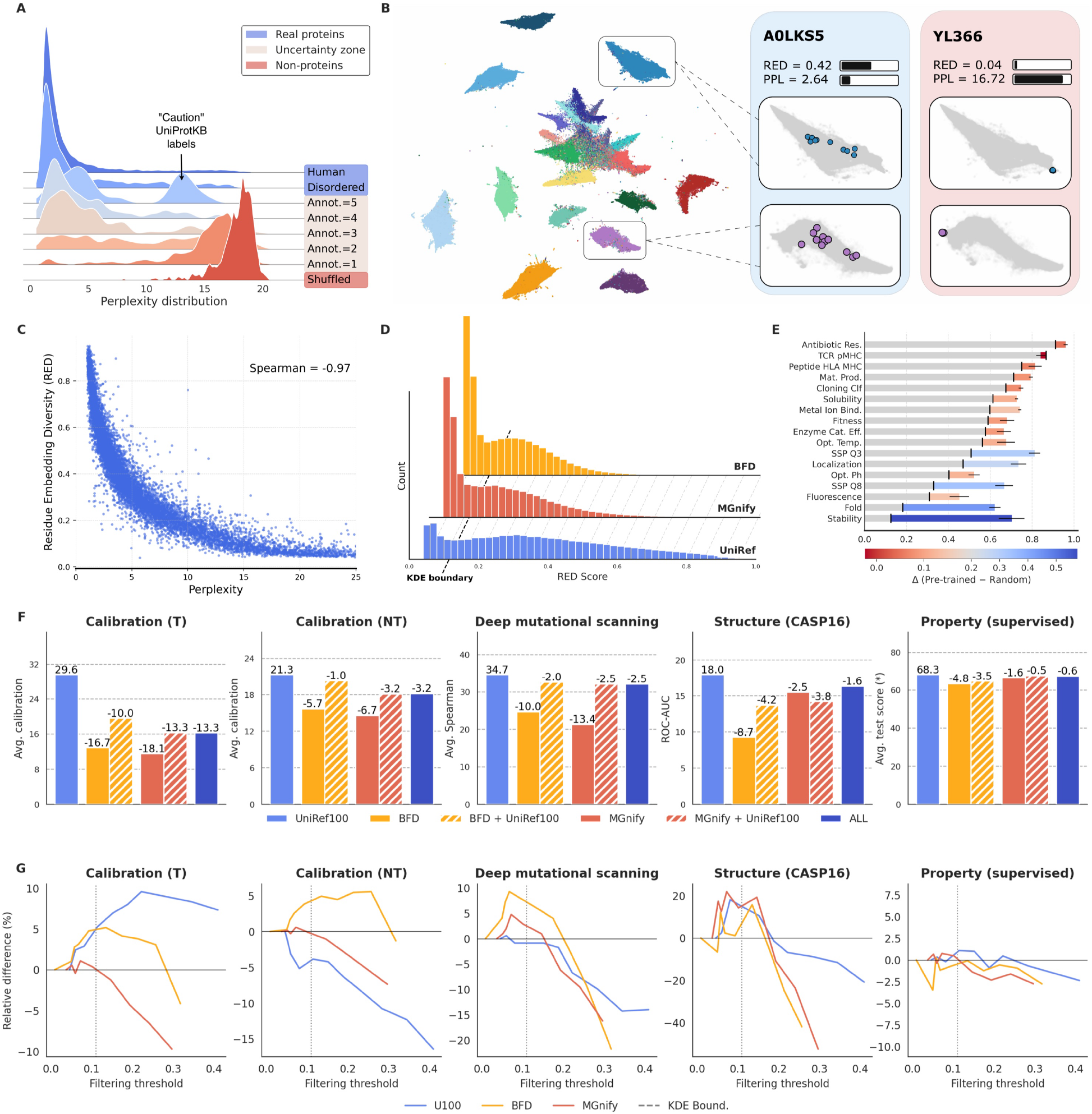
Residue-level representation collapse is a scalable proxy for perplexity and enables filtering of metagenomic databases. (**A**) Perplexity distributions for human, disordered, weak-evidence (PE=5), and shuffled sequences. In the PE=5 high-annotation set, all high-PPL sequences are flagged as unlikely to encode a functional protein in UniProtKB. (**B**) 2D projection (UMAP) of residue-level pLM embeddings colored by identity. A low-PPL sequence shows context-dependent intra-cluster diversity (left); a high-PPL sequence of similar length shows collapse to a single point per amino acid type (right). (**C**) RED and PPL correlation on a UniProtKB validation set (Spearman *r* = −0.97, 10,000 samples). (**D**) RED distribution for UniRef100, BFD, and MGnify. KDE-identified boundaries between the modes (vertical lines) are computed independently per dataset and converge to similar values. (**E**) Per-task test scores on property prediction for randomly initialized versus pretrained models. The largest pretraining gains occur on structural and stability-related tasks. (**F**) Relative scores of models pretrained on different datasets compared to the UniRef100-only baseline. Pretraining on unfiltered metagenomic data consistently degrades scores; including UniRef100 mitigates the effect. (**G**) Relative scores as a function of data filtering with RED. Scores improve up to a per-dataset optimum; the KDE boundary (vertical line) aligns with the empirical peaks.

This motivates the use of pLMs as quality filters for large, noisy metagenomic databases such as MGnify (*16*) and BFD (Big Fantastic Database) (*17*). However, computing pLM scores at scale is not straightforward as perplexity requires one forward pass per residue; scoring the billions of entries in metagenomic databases would take an estimated 90 H100 GPU-years. Investigating the geometry of residue-level pLM embeddings reveals a phenomenon that allows us to bypass this prohibitive cost. Perplexity is directly linked to the diversity of AMPLIFY 350M’s residue embeddings, with a sharp collapse for low-likelihood sequences. Within this embedding space, points cluster by amino acid type (Figure 1B). For a well-modeled, low-perplexity sequence, residues of the same type are diverse and modulated by their local context within the cluster: each occurrence of a given amino acid occupies a distinct position shaped by its sequence context. For a high-perplexity sequence of similar length, this intra-cluster diversity disappears. Every residue of a given amino acid type collapses to a single identical point, regardless of its sequential context; the representation is determined entirely by amino acid identity rather than by the sequence. In other words, while global chemical and structural contexts modulate the residue representations in well-modeled sequences, the model encodes no information beyond residue identity for poorly modeled sequences.

Residue Embedding Diversity (RED) is defined as one minus the mean pairwise cosine similarity of residue embeddings within a sequence: context-sensitive representations yield high RED; collapsed representations yield low RED. RED correlates near-perfectly with perplexity on our validation UniProtKB subset (10,000 sequences, Spearman *r* = −0.97; Figure 1C). Although this relationship saturates at high perplexity values (PPL > 15), such cases correspond to a regime where sequences are considered noise and where changes in perplexity do not lead to qualitative changes in likelihood (e.g., < 10^−6^ or < 10^−9^). RED’s sensitivity is therefore concentrated across the informative range of the model. Crucially, RED requires only a single forward pass per sequence rather than one per residue, reducing the estimated compute cost of scoring the 2.0B sequences in BFD and 2.5B sequences in MGnify from approximately 90 years to 25 days on a single H100 GPU.

### Unfiltered metagenomic pretraining degrades downstream performance but mitigates overfitting

Applying RED scores as a quality filter for pretraining datasets introduces the risk of inheriting the internal biases of the scoring pLM. Therefore, orthogonal evaluations are necessary to determine how data curation strategies influence pLM representations. We start by establishing baselines with models pretrained on unfiltered datasets. In zero-shot tasks, which require no additional training or supervision, models pretrained on metagenomic data consistently underperform compared to a UniRef100-only baseline (*18*). Specifically, there are absolute drops of 10.0% (BFD) and 13.4% (MGnify) on post-mutation fitness prediction (deep mutational scanning; DMS) (*4*), and 8.7% and 2.5% on contact prediction of CASP16 targets (Structure) (*3*). Incorporating UniRef100 into the pretraining mixture mitigates this effect, reducing the decline to 2.0% (BFD) and 2.5% (MGnify) on DMS (Figure 1F). These results confirm that expanding pretraining datasets with uncurated metagenomic sequences degrades the quality of signals learned during pretraining.

Supervised tasks, in which probes are trained to predict sequence or residue properties from model embeddings, exhibit variable sensitivity to pretraining data mixtures. Across 17 supervised property prediction tasks, aggregate test scores (F1 or Matthews coefficient) differ by less than 5% among all baselines (Figure 1F). Notably, we find that a shallow linear probe trained on random representations can recover most of the performance on several tasks (Figure 1E). Since scores are derived from classification or regression heads trained on frozen pLM embeddings, they conflate information transferred from pretraining with information learned during the second phase of supervised training. To separate these contributions, the difference in scores between random representations and pretrained embeddings can be computed. Structure-related tasks benefit substantially from pretraining, with stability prediction (+57.7%), fold classification (+44.0%), and secondary structure prediction (+30.4% Q3, +33.7% Q8) showing significant improvements over random baselines. Other functional tasks show minimal differences, with TCR-pMHC prediction even exhibiting a 2.5% average decline. Among these tasks, some are saturated, with random embeddings already achieving high accuracy, while others depend on features that masked language modeling captures only weakly. Thus, variation in this property prediction benchmark is driven primarily by models’ ability to predict stability and structure.

In a novel zero-shot task assessing model calibration to the evolutionary landscape (see section 4), metagenomic models perform worse than the UniRef100 baseline, but exhibit reduced signs of overfitting. In multiple sequence alignments, at positions where the target residue matches the evolutionary consensus (T), both memorization and generalization predict the same residue. At positions where it differs (NT), only a model that has memorized the sequence favors the target variant over the more stable consensus, making the T–NT gap indicative of overfitting to the training proteomes. In this task, unfiltered metagenomic models underperform the UniRef100 baseline at T positions (−16.7% BFD, −18.1% MGnify) but show much smaller declines at NT positions (−5.7% BFD, −6.7% MGnify; Figure 1F), suggesting reduced memorization despite a weaker overall fit to the natural protein distribution.

### RED-based filtering consistently improves representations and increasing training steps reveals divergent scaling dynamics

Although metagenomic pretrained models underperform the UniRef100 baseline overall, we hypothesize that this can be mitigated by filtering noise using pLM quality scores. RED scores under AMPLIFY 350M exhibit a consistent bimodal distribution across UniRef100 and metagenomic databases (Figure 1D): a sharp primary mode of low-RED, collapsed sequences and a broader secondary mode of high-RED, well-modeled sequences. Decision boundaries between the two modes, identified by kernel density estimation (KDE), fall at similar RED values (∈ [0.11, 0.13]) when computed independently on the three datasets. The convergence of these boundaries across sets of very different composition suggests that the threshold reflects the scoring model’s representation mechanisms rather than any particular dataset’s properties.

To determine whether RED scores provide a quality signal transferable to metagenomic datasets, we pretrain models on each source, filtered at progressively stricter RED thresholds, and evaluate them on the tasks established in the previous section (Figure 1G). Downstream performance follows a consistent trend: filtering improves scores up to a per-dataset optimum, beyond which the removal of well-modeled sequences degrades them. Metagenomic databases, which contain the highest proportion of low-RED sequences, show the largest absolute gains. The empirical optima align with the KDE-identified boundary between the two modes of each distribution, suggesting that this unsupervised threshold separates sequences with downstream utility from those without. However, filtering metagenomic data does not close the gap with UniRef100: even after removing low-RED sequences, metagenomic models retain a performance deficit relative to models trained on curated data.

We hypothesize that metagenomic models are undertrained: the size and diversity of metagenomic sequence space shifts the compute-optimal frontier toward longer training, as suggested by standard scaling laws (*19*). Increasing the training budget from 200B to 2T tokens reveals dataset-dependent scaling dynamics on the zero-shot benchmarks (Figure 2A-B). UniRef100-trained models achieve strong absolute scores early but gain little from additional training. Models trained on metagenomic data start at a lower level but show larger relative improvements with more steps. On structural modeling, as measured by the categorical Jacobian on CASP16 targets, metagenomic models match or surpass UniRef100 models at scale (MGnify AUC 24.1 vs. UniRef100 22.8; CASP16 zero-shot contact prediction). Moreover, while models pretrained on UniRef100 consistently improve with longer training at T positions, their calibration degrades at NT positions (mean drop of 14.1 points), suggesting progressive memorization of the training proteomes. By contrast, models pretrained on diverse metagenomic datasets do not exhibit the same degradation (11.2 and 3.1 average improvement for BFD and MGnify models). These distinct scaling behaviors motivate finer-grained strategies of data composition to harness the full potential of metagenomic sources.

**Figure 2:**
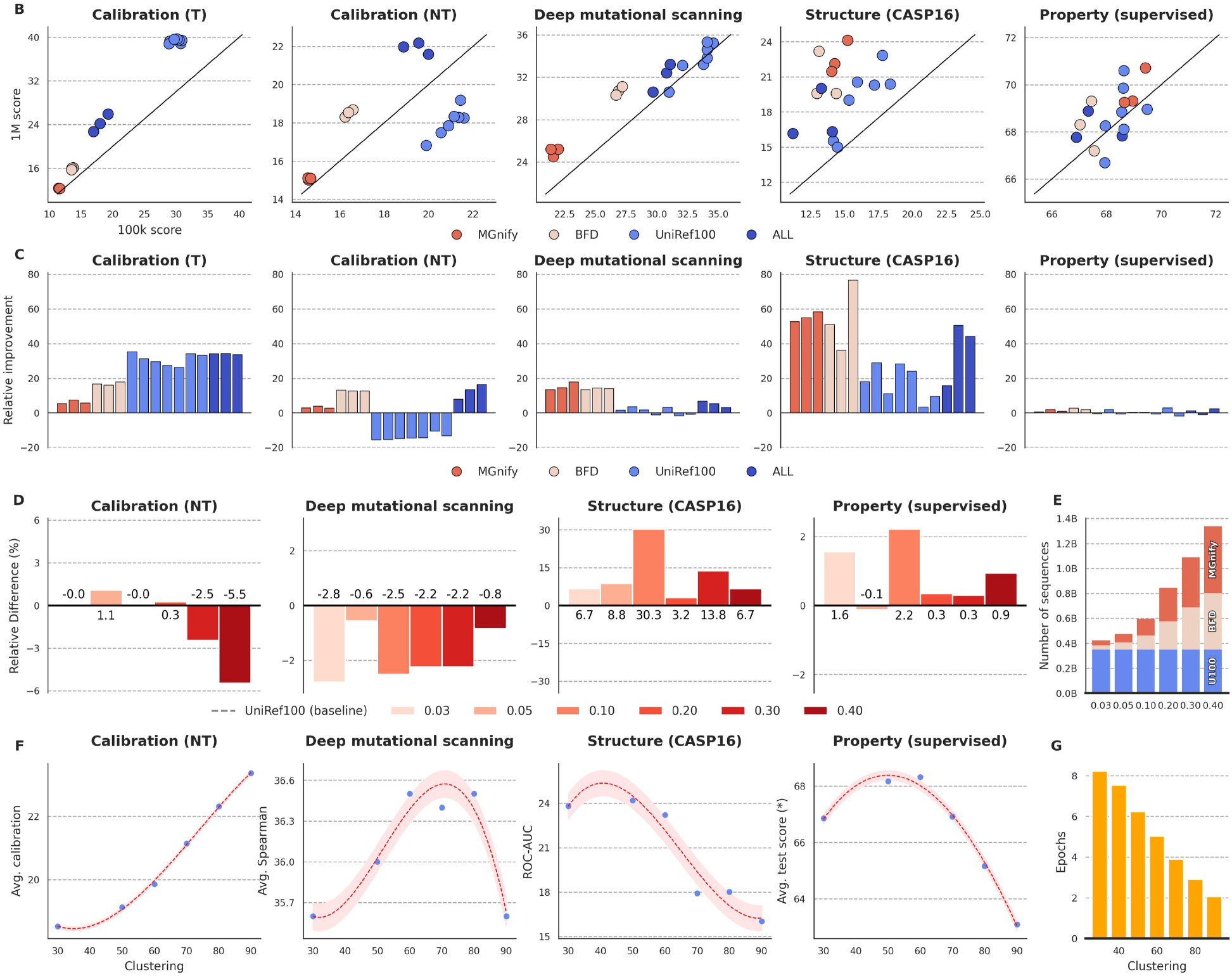
Pretraining data strategies lead to trade-offs in downstream task performance. (**A**) Scores at 200B versus 2T pretraining tokens; colored by dataset. Longer pretraining improves all tasks regardless of the data quality, except for calibration on non-target positions for UniRef100 models. This is a direct signature of overfitting to the training data distribution. (**B**) Relative score change from 200B to 2T tokens. Metagenomic models show the largest gains, particularly on DMS and CASP16 contact prediction. (**C**) Relative score change compared to the UniRef100-only baseline as a function of metagenomic subsampling proportion. Structural and property prediction peak at 0.10, at which point calibration recovers to baseline levels; DMS spearman decreases systematically with metagenomic datasets. (**D**) Composition of pretraining mixtures as metagenomic subsampling increases from 0.03 to 0.4. (**E**) Scores as a function of UniRef100 clustering threshold (30% to 90% identity). Structural tasks improve with aggressive clustering; calibration and DMS degrade. (**F**) Training epochs at each clustering level; lower thresholds produce smaller datasets and more epochs for the same step budget.

### Heterogeneous data mixtures and sequence clustering expose a memorization–generalization trade-off

Careful construction of heterogeneous pretraining data mixtures reveals trade-offs in downstream task performance, with structural modeling and fitness prediction responding in opposite directions. After RED filtering, metagenomic sequences are dynamically subsampled at rates ranging from 0.03 to 0.4 and mixed with UniRef100 (Figure 2C-D). Incorporating this metagenomic data improves performance on structural tasks, peaking at a subsampling rate of 10%, which incidentally corresponds to a calibration score (NT) that matches the UniRef100 baseline. Further increases in metagenomic data degrade performance. The average Spearman correlation for DMS, however, consistently drops relative to the UniRef100-only baseline. The dense evolutionary signal required to resolve the functional impact of single-amino-acid substitutions appears to be diluted by the addition of metagenomic data, even though this diversity broadens the structural patterns accessible to the model.

Sequence clustering produces a consistent pattern from a different angle. Clustering UniRef100 at identity thresholds from 90% to 30% collapses local evolutionary neighborhoods into single representatives, progressively removing the redundancy that distinguishes close homologs. Under a fixed training budget of 200B tokens (Figure 2E-F), aggressive clustering improves structural tasks, with maximum scores achieved at a 50% identity threshold, but degrades calibration and DMS. Both strategies, adding metagenomic diversity and clustering, thin the local homologous context around any given sequence, and both shift downstream performance toward structural modeling and away from evolutionary precision.

These two axes point to a common mechanism. In dense, unclustered training data, each sequence is surrounded by close homologs with minor variations. Memorizing individual sequences in this context is impractical; the abundance of local variation likely forces the model to learn the stability rules inherent to the underlying distribution rather than encoding specific sequences. This regime favors tasks that depend on robust generalization over local evolutionary landscapes, such as calibration (NT) and DMS. Aggressive clustering or the addition of diverse metagenomic sequences sparsifies the training landscape, retaining only distinct representatives and removing the small variations between them. Under a fixed number of clusters, this reduction lowers the cost of memorization, enabling the model to efficiently encode structural patterns from individual sequences, consistent with recent work showing that residue contacts can be recovered by matching memorized patterns (*3*). No single training distribution simultaneously optimizes both regimes. These findings motivate training two 350M-parameter models on complementary data mixtures: AMPLIFY-B for structural modeling and AMPLIFY-C for calibration to the evolutionary landscape (section 4). Because neither the architecture nor the training strategy was modified between these models, performance differences directly reflect the impact of the training data mixture. Compared to AMPLIFY 350M, both models improve across the three zero-shot evaluation tasks, with the largest gains aligning with their intended specialization. AMPLIFY-B achieves the greatest structural improvement, increasing performance on the CASP16 categorical Jacobian task by 59.3%, while also improving calibration (NT) by 88.6% and DMS by 7.7%. AMPLIFY-C achieves the largest calibration gain, improving calibration (NT) by 120.9%, while also improving the CASP16 categorical Jacobian task by 18.3% and DMS by 5.2%. Data mixture therefore provides a controllable axis for specializing pLM capabilities: AMPLIFY-B and AMPLIFY-C prioritize different aspects of the memorization–generalization trade-off, while surpassing the UniRef100-only baseline in structural modeling, evolutionary calibration, and DMS.

### pLMs capture sequence plausibility and are predictors of wet-lab expression

The preceding analyses rely on the assumption that pLM scores and representations encode protein viability. We validate this assumption *in vivo* with a wet-lab discovery campaign within the EC 1.14.11 enzyme class (Fe/αKG-dependent dioxygenases). Fe/αKG-dependent enzymes have emerged as valuable biocatalysts for challenging C-H oxidation reactions owing to their exquisite selectivity. After pre-filtering, candidate sequences are selected from a set of 34k enzymes, with selected candidates constrained to share less than 50% sequence identity. If perplexity captures viability, then guiding candidate selection with pLM scores should improve experimental outcomes. Here, we use expression and purification yields as proxies for protein solubility and stable folding, i.e., biophysical viability. 288 candidates, split into three 96-well plates, are selected using three strategies with AMPLIFY-C: quality-focused pLM-guided selection (Plate A), embedding diversity-focused pLM-guided selection (Plate B), and a sequence diversity-only control without any pLM score (Plate C). The selection strategies, detailed in section 4, rely on the same quality filters for all three plates but are augmented with pLM scores for plates A and B. Resulting band volumes, as determined by SDS-PAGE analysis, demonstrate that perplexity acts as a requirement for reaching minimum expression levels: the convex hull of expression versus perplexity forms a right-triangular envelope whose hypotenuse slopes downward (Figure 3C), indicating that sequences must fall below a perplexity threshold to achieve a given expression level, although low perplexity alone does not guarantee expressibility in E. coli (purified band Pearson *r* = −0.15, *p* = 9.10 × 10^−3^, lysate band Pearson *r* = −0.14, *p* = −1.90 × 10^−2^). For lysate expression, Plates A and B achieve comparable total volume (normalized band volumes of 0.16 and 0.17, respectively), while Plate C, the diversity-only control, reaches roughly half that level (0.08). For purified expression, total volume decreases monotonically with average plate perplexity (Plate A highest, Plate C lowest; Figure 3B). In both assays, incorporating pLM likelihood into the selection criterion improves wet-lab yield relative to a sequence-diversity-only strategy.

**Figure 3:**
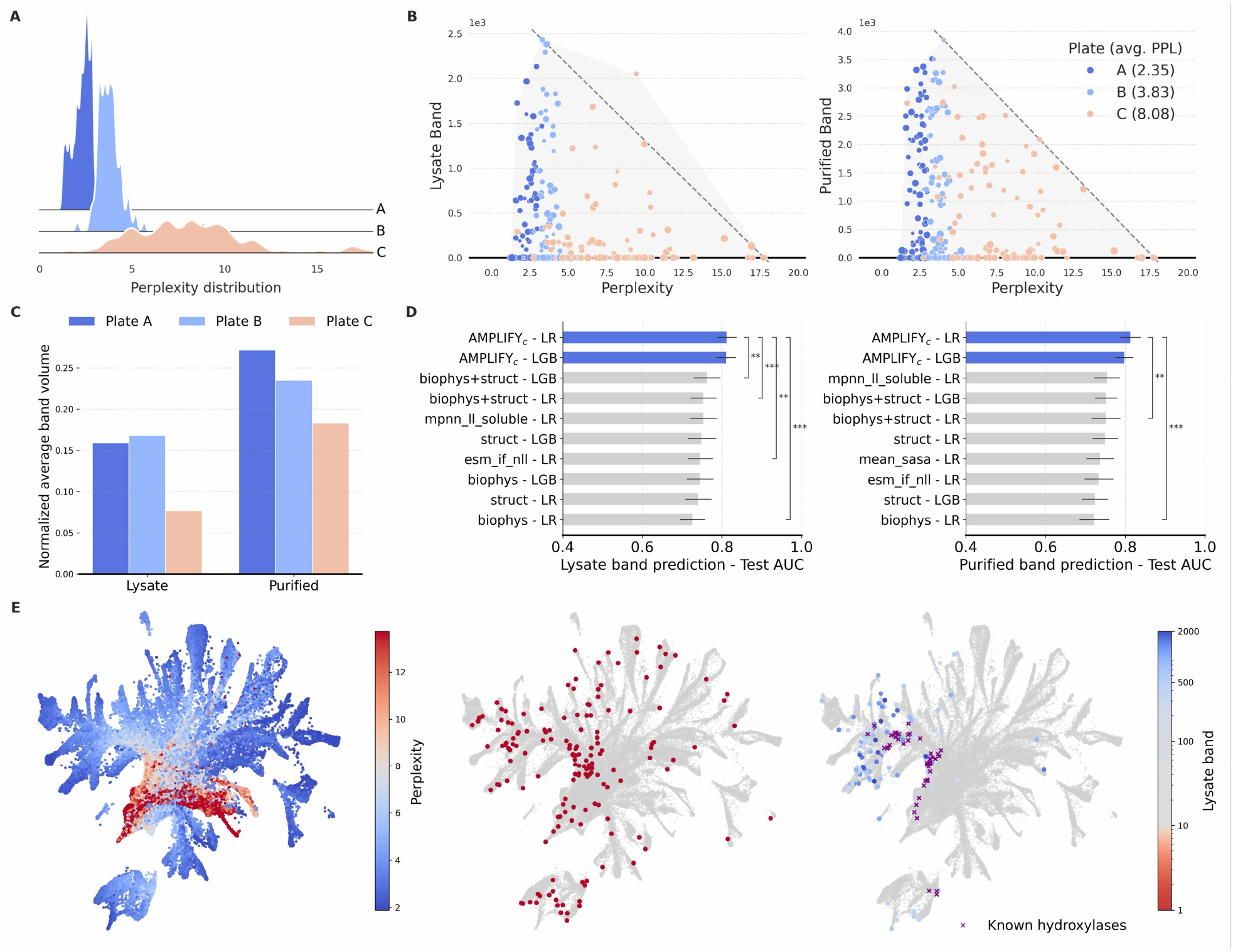
pLM representations encode expressibility. (**A**) Per-plate perplexity distributions. Plates use similar selection pipelines, with varying reliance on the pLM. Plate A emphasizes PPL-based quality; Plate B balances quality and embedding diversity; Plate C uses sequence diversity only, without any pLM score. (**B**) Average normalized band volumes per plate. (**C**) Band expression as a function of perplexity for synthesized candidates. (**D**) Test set ROC-AUC for binary expression classifiers with LightGBM (LGB) and logistic regression (LR). pLM embedding-based classifiers outperform all structural and biophysical feature sets. (**E**) UMAP of pLM sequence embeddings, colored by perplexity (left), purified band expression (< 10 center; > 10 right). Purple crosses indicate characterized Fe/αKG-dependent hydroxylases from the literature.

Moreover, biophysical viability is not only linked to a scalar perplexity score but is also topologically organized within the pLM embedding space. A 2D projection with UMAP (*20*) of sequence embeddings shows that strongly expressed candidates, as identified by lysate and purified band assays, concentrate in overlapping spatial regions (Figure 3E). This organization is predictively stronger than the conventional toolkit: binary classifiers trained on pLM embeddings significantly outperform classifiers trained on combinations of features, including AlphaFold mean pLDDT (*17*), MPNN soluble log-likelihood (*21*), ESM inverse folding negative log-likelihood (*22*), or mean solvent-accessible surface area on both prediction tasks (paired permutation test, *p* < 0.011 for all comparisons, 8 of 12 at *p* < 0.001; Figure 3D). The embedding, derived from sequence alone without any structural computation, is therefore a reliable discriminator of wet-lab expression in this experiment.

Taken together, these results establish complementary signals. Perplexity ranks and filters candidates in alignment with curator-level quality assessments, thereby improving experimental success rates. The embedding encodes a topology that is a direct predictor of expression. Both signals arise from evolutionary sequence statistics alone, with no structural supervision.

### pLM-guided retrieval recovers highly expressed metagenomic proteins inaccessible to homology search and structure prediction tools

Beyond selecting UniProtKB-annotated enzymes, we evaluate AMPLIFY-C’s ability to recover viable proteins from the metagenomic space. From the billions of sequences in BFD and MGnify, candidates are retrieved through embedding similarity to the sequences selected from enzyme class 1.14.11, after RED filtering, kNN search, and MMR optimization (see Methods). Homology-based retrieval would not return any of these candidates, as none of the 96 selected sequences have detectable sequence similarity to members of the enzyme class 1.14.11 under MMseqs2 (Figure 4A). Despite this, 30 candidates show detectable expression (band > 10), 24 of these reaching high expression levels (purified or lysate band values over 500).

**Figure 4:**
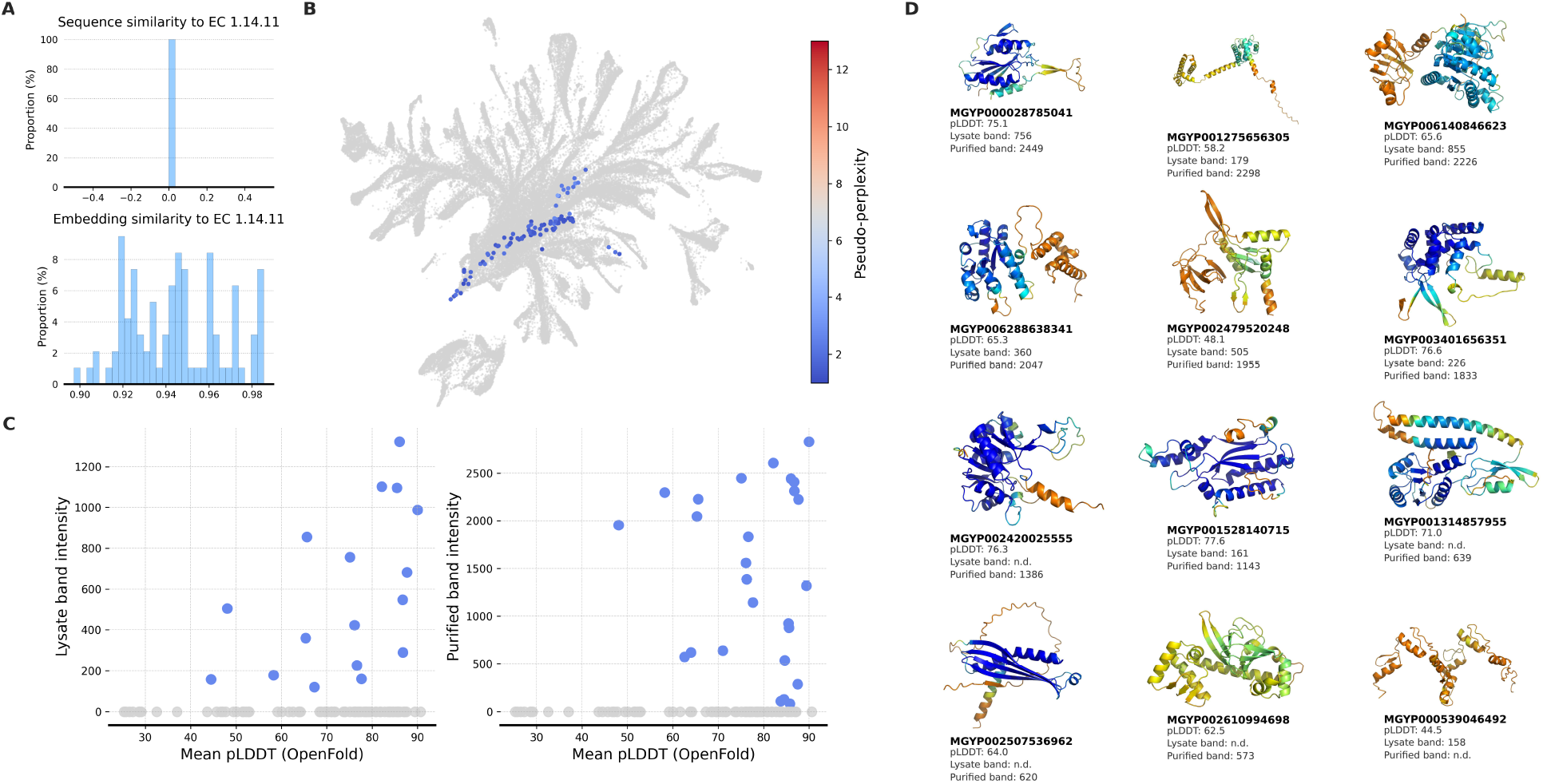
pLM representations allow retrieval of viable sequences from metagenomic space. (**A**) Sequence and embedding similarities between metagenomic candidates and annotated members of the EC 1.14.11 enzyme class. None of the candidates has detectable sequence similarity to any annotated member under MMseqs2 (*23*), and all have high embedding cosine similarity under AMPLIFY-C. (**B**) Projection of metagenomic candidates on the UMAP of wet-lab candidates from Figure 3E, colored by perplexity under AMPLIFY-C. (**C**) Lysate and purified band volume from recombinant expression against pLDDT predicted by OpenFold for metagenomic candidates. A number of highly expressed proteins receive low structural confidence, indicating that structure prediction alone would not have prioritized them. (**D**) Structures predicted by OpenFold for a subset of highly expressed metagenomic candidates with low predicted structural confidence.

Notably, seven of the most highly expressed candidates receive low predicted structural confidence from OpenFold (*24*) (mean pLDDT < 70; Figure 4C, D). Structure prediction alone would not have prioritized these sequences: they lack both the sequence identity required for homology search and the fold confidence that would flag them as plausible candidates. That they nonetheless express as soluble proteins shows that the pLM embedding carries information about expressibility that predicted structural confidence does not: it ranked these sequences as viable where pLDDT would have discarded them.

Expression rates are lower than for the curated selections (Plates A and B), as expected: there, the pLM prioritizes sequences within a pool of annotated enzymes already known to belong to the target class. In the unannotated metagenomic space, it must first identify which sequences are plausible enzymes before judging their expressibility. The lower yield reflects the difficulty of selecting candidates from unannotated space. Nevertheless, pLM representations can organize metagenomic sequences well enough to retrieve expressible proteins in regions inaccessible to conventional tools. pLM scores can therefore serve as starting points for experimental groups to probe metagenomic space in a computationally tractable way. That metagenomic sequences with no detectable homology can be retrieved and expressed as soluble proteins demonstrates that a significant portion of the functional protein universe remains accessible only through the learned representations of a sequence model trained on the right data.

## 3. Conclusion

Protein language models can recover viable proteins from regions of sequence space beyond the reach of homology search and structure prediction. From billions of metagenomic sequences, we retrieved 96 candidate dioxygenases using likelihood and embedding similarity alone, none sharing detectable homology with any annotated member of the enzyme class. Of these candidates, 30 expressed as soluble proteins and 24 at high levels, including seven highly expressed proteins with low predicted structural confidence. These results establish that protein language model representations carry a signal complementary to classical sequence alignment and structure prediction.

Across more than 100 controlled pretraining runs, we establish that data composition, rather than scale, determines what protein language models learn. We discover that dense homologous coverage favors evolutionary calibration and fitness prediction, whereas sparse, diverse data favor structural modeling. The fundamental trade-off depends on the density of homologs around each sequence. We introduce calibration to the evolutionary distribution as a task that makes this trade-off measurable and show that longer training on curated data exacerbates memorization. Because structural modeling gains came at the cost of evolutionary calibration across all mixtures examined, we trained and released AMPLIFY-B and AMPLIFY-C as specialized models that sit on different ends of the trade-off.

Working at metagenomic scale required a sequence-quality measure that could be computed cheaply. We establish that perplexity, a sequence’s likelihood under the model, aligns with expert curation, separating disordered but functional proteins from pseudogenes and frameshifted artifacts that structural confidence cannot distinguish. Because perplexity costs one forward pass per residue, we introduce Residue Embedding Diversity (RED), a single-pass proxy correlated with perplexity at Spearman’s *r* = −0.97 that lowers the cost of scoring billions of sequences from an estimated 90 GPU-years to 25 days. We further show that embeddings predict experimental expression more accurately than structural and biophysical features.

Together, these results transform metagenomic sequences into a tractable extension of the accessible protein universe. We introduce RED to enable billion-scale metagenomic curation, establish calibration as a diagnostic for evaluating protein language models, and discover that data composition controls the trade-off between evolutionary calibration and structural modeling. Guided by the model’s own measure of quality, sequence diversity that current pipelines ignore becomes raw material for discovery, and learned representations turn that diversity into a searchable space. While much of the functional protein world still lies beyond the reach of homology and structure, our results show that it is within reach of representations trained on the right data.

## Supporting information

Code bundle

## 4. Acknowledgment

We thank Amgen’s Alan Russell, Marti Head, Carolyn Ch’ing, Matt Foster, and Karen Chou; Mila’s Almer van der Sloot, Shawn Whitfield, and Robert Farias. We also thank Amgen, Mila - Quebec AI Institute, Lambda AI, Compute Canada, and the Royal Military College of Canada for providing compute resources.

## AI tools

Generative AI tools were used for code review and manuscript editing.

## Data and material availability

All result details are available in the main text or in supplementary material. Code is available on GitHub at https://github.com/flair-bio. Models and datasets are available on HuggingFace at https://huggingface.co/flair-bio.

## Material and Methods

### Data acquisition and preprocessing

#### Pretraining databases

Three open-source databases are used. UniRef (UniProtKB Reference Clusters, 2024_05 release) provides high-quality, clustered protein sequences and annotations. BFD (Big Fantastic Database) is used as a source of metagenomic diversity; we preprocess it by filtering out artificial consensus sequences and entries from TremBL/Swiss-Prot, capitalizing residues, and removing alignment gaps. MGnify (EBI-Metagenomics, 2024_04 release) is used to maximize coverage of the microbial world, retaining all sequences regardless of full-length indicators. For the final specialized models (AMPLIFY-B and AMPLIFY-C), we additionally include all paired antibody sequences from the Observed Antibody Space (OAS) (*25*). Each paired entry consists of a heavy chain and a light chain; we augment the data by concatenating both orderings (heavy|light and light|heavy) as separate training sequences, exposing the model to both chain arrangements during pretraining. A validation set was built following the AMPLIFY protocol (*6*). All training sets were deduplicated against this validation set using a 90% sequence identity threshold. While a 90% threshold is too permissive for measuring generalization, we use this deduplicated validation set to assess the models’ language modeling abilities on the training landscape, and we rely on our extensive suite of downstream tasks to evaluate the models’ utility and generalization to novel functions.

#### Ambiguous Amino Acids

Open-source databases frequently contain non-standard characters (B, O, U, X, Z, J) resulting from sequencing ambiguity or the presence of non-canonical amino acids. These characters carry varying degrees of biological meaning: O (pyrrolysine) is a genuine, but rare, non-canonical amino acid; Z denotes ambiguity between glutamic acid (E) and glutamine (Q); while others, such as X, represent wholly unresolved reads. These characters reflect noise or real biochemical uncertainty and therefore are not tokens the model should be trained to predict. Discarding the corresponding sequences, however, wastes otherwise valid data. To avoid this, we adopt a masking strategy where ambiguous residues are removed from the input token sequence but accounted for in the positional indexing. By shifting the Rotary Position Embeddings (RoPE) (*26*) of subsequent residues to account for the gap, we preserve correct positional information without forcing the model to predict tokens outside the canonical amino acid vocabulary.

### Data mixture strategies

#### Scoring

The likelihood of a sequence S of length L decomposes as a product of conditional probabilities, 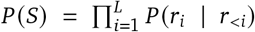 which means likelihood decreases with sequence length. Researchers therefore commonly rely on perplexity, which provides a length-normalized measure of a model’s predictive confidence.

For a masked language model, however, the true perplexity cannot be computed exactly and must be approximated. One approach, termed pseudo-perplexity, masks a random subset of positions and estimates the full perplexity from that single conditional probability. This estimate is fast but stochastic and noisy. A more principled approximation is to mask each residue individually and aggregate the resulting conditionals:

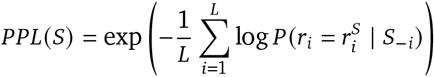

This quantity is deterministic and well-defined, but requires one forward pass per residue. This is the definition of perplexity we adopt in this work. However, computing it for the billions of sequences in metagenomic databases is computationally intractable.

To address this, we introduce **Residue Embedding Diversity (RED)**, a computationally efficient proxy computable in a single forward pass per sequence. RED measures the diversity of residue-level embeddings via average pairwise cosine similarity:

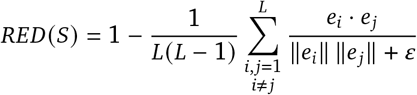

where *e*_*i*_ is the embedding vector of the *i*-th residue. RED is motivated by the observation that low-likelihood sequences cause representation collapse for pLMs, where residue embeddings degenerate to static representations of residue identity rather than reflecting sequence context.

#### Filtering

We score all datasets using RED computed by AMPLIFY 350M (*6*). This model is used exclusively for RED scoring and filtering of the pretraining datasets and is not the subject of the ablation or wet lab experiments. We filter datasets independently by removing sequences in the bottom *n*% quantile, *n* varying from 0% to 90%. We train and evaluate models on resulting sets in identical settings to determine the optimal experimental filtering points on downstream benchmarks.

#### Subsampling

Mixtures are constructed by filtering out sequences below a RED threshold of 0.115. This threshold is an empirical aggregate of the optimal filtering points for downstream tasks and the boundary separating the two modes of the distribution of RED scores across all datasets. Subsequently, metagenomic sources (BFD, MGnify) are dynamically subsampled to a proportion *p* at each new epoch, thereby modulating the distribution of our pretraining data to balance curated and uncurated domains.

#### Clustering

We cluster UniRef100 at sequence identity thresholds ranging from 90% to 30% and retain only cluster representatives. Clustering is performed with MMseqs2 `easy-linclust` (*27*). We apply an iterative strategy, similar to UniRef: the dataset is first clustered at 90% identity, and each subsequent threshold (80%, 70%, 60%, 50%, 40%, 30%) is applied to the representative sequences of the previous level rather than to the full database, reducing computational cost while preserving cluster hierarchy. At all levels, we require a minimum alignment coverage of 80% of the target sequence (`–cov-mode 1, -c 0.8`).

### Model architecture and training

#### Model architecture

All models are transformer-based encoder pLMs following the AMPLIFY architecture (*6*), a BERT-like masked language model with pre-layer normalization, Rotary Position Embedding (RoPE), and a SwiGLU activation. Unless stated otherwise, ablation models use 24 layers with a hidden size of 640, resulting in 120M parameters.

#### Training protocol

We implemented a codebase that makes use of state-of-the-art libraries and training accelerations. In particular, we use the packed version of flash-attention (2.7.1) (*28*), compile the model, and rely on HuggingFace’s accelerate (1.4.0) (*29*) and DeepSpeed ZeRO 2 (0.14.4) (*30*) with PyTorch (2.5.0). We train all models with a warmup followed by a cosine decay schedule, with a peak learning rate of 1*e* − 3. We use the AdamW (*31*) optimizer with weight decay of 0.01, betas of (0.9, 0.95), and eps 1*e* − 8.

#### Small-scale ablations

To bring the training time of our ablations under a day on 8 H100 GPUs, we combine this fast implementation with a smaller training scale of 100,000 steps, with a maximum sequence length of 512 and a batch size of 4,096 (approximately 200B tokens).

#### Specialized models

The ablation experiments identify two data regimes with complementary strengths: dense homologous coverage for evolutionary precision and filtered metagenomic diversity for structural modeling. We translate these findings into two specialized 350M-parameter models (32 layers, hidden size 960), each trained in two stages following standard practice: 1M steps on sequences of length 512, followed by 50,000 steps on sequences of length 2,048 for context extension, for a total of approximately 2.1T tokens.

**AMPLIFY-benchmark (AMPLIFY-B)** targets structural modeling and broad benchmark performance. It is trained on UniRef100 at a 0.05 quantile filtering threshold on RED scores with 50% identity clustering; BFD at a 0.2 quantile threshold with 0.03 subsampling; and MGnify at a 0.2 quantile threshold with 0.03 subsampling.

**AMPLIFY-consensus (AMPLIFY-C)** targets evolutionary precision and functional viability scoring. It is trained on UniRef100 at a 0.1 quantile filtering threshold on RED scores with 90% identity clustering; BFD at a 0.5 quantile threshold with 0.05 subsampling; and MGnify at a 0.2 quantile threshold with 0.05 subsampling.

Both models use the same architecture and optimizer settings as the ablation models. AMPLIFY-C is used for all wet-lab candidate selection, embedding-based retrieval, and expression analysis reported in this study.

### In silico evaluations

#### Perplexity distributions

We score five curated protein sets with perplexity (PPL) under the public model AMPLIFY 350M. (i) The human reference proteome (UniProtKB Proteome ID UP000005640; 2025/07/10; 20,541 sequences).(ii) Reviewed intrinsically disordered proteins from the DisProt database (*15*) (2025/07/09; 1,742 entries). These serve as a positive control for evolutionary plausibility in the absence of a stable fold. (iii) Shuffled human sequences, generated by randomly permuting the residue order of each human protein while preserving amino acid composition; 692 entries. These serve as a negative control, destroying positional syntax while retaining global compositional statistics. (iv) All UniProtKB entries with protein existence level 5 (uncertain; 2025/07/10; 1,738 entries), stratified by annotation count (1 through 5) as a proxy for curation confidence. (v) A subset of the PE = 5 sequences at the highest annotation level (*n* = 23), analyzed separately because of their bimodal perplexity distribution; high-perplexity members of this group were cross-referenced against UniProtKB *Caution* annotations to identify likely pseudogenes. Full per-set statistics are reported in Table S17.

#### Calibration task

This task measures whether a model has learned the evolutionary consensus at conserved positions or has instead memorized the specific proteome sequences seen during training. We use the human proteome from UniProtKB (Proteome ID UP000005640) and align each human protein to the BFD database with HHblits (*32*), requiring a minimum sequence identity and coverage of 50% and a maximum pairwise sequence identity of 99%. Proteins with fewer than 10 homologs are discarded. Aligning the entire human proteome against BFD with HHblits is a computationally intensive step, as runtime scales with proteome size and database depth, but is performed only once as a preprocessing step.

For each retained protein, we compute the positional amino acid frequency across all homologs and identify positions where the consensus residue exceeds a conservation threshold (> *n*% for different values of *n*). These conserved positions are split into two categories. **Target (T)** positions are those where the human residue matches the evolutionary consensus: a model that has learned either the consensus or the human sequence will predict the correct residue, so high accuracy is expected from any well-trained model. **Non-target (NT)** positions are those where the human residue differs from the consensus: here, the more stable prediction is likely the consensus residue, but a model that has memorized the human proteome will instead predict the human variant. Accuracy on NT positions therefore isolates memorization from generalization: a model that improves on T positions while stagnating or degrading on NT positions is progressively overfitting to the training proteome.

Beyond top-1 accuracy, we quantify how well the full predicted distribution matches the evolutionary distribution with a calibration score:

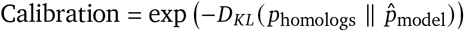

where 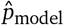 is the softmax distribution over amino acids at the masked posi tion and *P*_homologs_ is the empirical frequency distribution from the multiple sequence alignment. A score of 1 indicates perfect agreement between the model’s output distribution and the evolutionary distribution; scores decrease as the two distributions diverge. This scalar summarizes, in a single number per position, whether the model has internalized the broader evolutionary landscape or has collapsed onto the target proteome.

#### Validation of consensus mutations via in silico mutagenesis

To assess whether consensus residues identified by multiple sequence alignment confer greater thermodynamic stability than the corresponding human residues, we performed systematic *in silico* mutagenesis using the Rosetta macromolecular modeling suite (version 2025.51) (*33*). We focused on positions in the human proteome where the evolutionary consensus residue exceeded 70% frequency across homologs yet differed from the human sequence (3,302 positions across 1,905 proteins). For each position, we retrieved experimental three-dimensional structures by querying the SIFTS (Structure Integration with Function, Taxonomy and Sequences) database (*34*) to map UniProtKB accessions to PDB entries (*35*). When multiple chains covered a given site, we selected the chain with the best crystallographic resolution and highest sequence coverage. We restricted analysis to X-ray structures, excluding NMR models. A subset of 454 positions was sampled uniformly across proteins after filtering to sites covered by a PDB chain.

Each selected PDB chain was extracted, stripped of non-protein atoms, and subjected to Cartesian-space energy minimization using the Rosetta FastRelax protocol (*36*) with the ref2015_cart scoring function. This relaxation step is critical to resolve steric clashes inherent in deposited coordinates before computing energy differences. The free energy change upon mutation (ΔΔ*G*) was then estimated using the Rosetta cartesian_ddg application, running three independent iterations per mutation with a backbone neighbor shell of one residue. ΔΔ*G* was computed as the difference in mean Rosetta energy units (REU) between the consensus mutant and the human wild-type: ΔΔ*G* = *E* (*consensus*) - *E* (*human*). A negative ΔΔ*G* indicates that the consensus residue is predicted to be more stable in the context of the human protein backbone.

We note that this approach evaluates stability changes on a fixed backbone and does not capture conformational rearrangements that may accompany the substitution in a natural evolutionary context. Nevertheless, Cartesian ΔΔ*G* with ref2015_cart has been shown to correlate well with experimentally measured stability changes for single-point mutations (*37*), making it a suitable proxy for the biophysical signal we seek to quantify.

#### Deep Mutational Scanning (DMS)

We evaluate zero-shot fitness prediction on the substitutions split of the ProteinGym benchmark (September 2025 release; 217 assays, 2.4M mutated sequences) (*4*). Mutations are scored with the Masked-Marginal method: each mutated position is masked and the log-likelihood ratio of the mutant versus wild-type amino acid is computed from the wild-type sequence context; multi-site mutations are scored by summing per-position ratios. Performance is the Spearman correlation between model scores and experimental fitness, aggregated by a weighted average over UniProtKB sequences and then functional categories, following the ProteinGym aggregation protocol (*4*).

All ProteinGym wild-type sequences are present in UniProt, creating a potential data-leakage advantage for models pretrained on these sequences. We follow community practice (*2, 12*) and do not deduplicate against ProteinGym wild-types. However, our validation-set deduplication (90% identity) may inadvertently remove evolutionary context around ProteinGym targets, potentially suppressing reported scores. Moreover, we note that fitness in ProteinGym spans diverse biophysical properties (enzymatic activity, binding affinity, thermostability), and there is no guarantee that evolutionary sequence likelihoods correlate uniformly with all of them in a zero-shot setting.

#### Unsupervised structure prediction

As a zero-shot probe of the structural information encoded in pLM representations, we compute the categorical Jacobian introduced previously (*3*). For a sequence of length *L*, the Jacobian entry *J* [*i, a, j, b*] measures how substituting residue *b* at position *j* changes the model’s log-probability of observing amino acid *a* at position *i*:

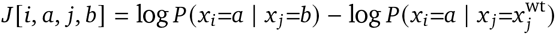

Where 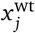 is the wild-type residue at position *j*. Computing the full *L* × 20 × *L* × 20 tensor requires *L* × 20 forward passes (one per single-site substitution). Predicted contact maps are derived by four-pass sequential centering of *J* over each axis, symmetrization, Frobenius norm over the amino acid dimensions, and average product correction (APC) to remove phylogenetic background. This procedure requires no supervised training head and extracts inter-residue coupling directly from the masked language model.

We evaluate on CASP16 (Critical Assessment of protein Structure Prediction) (*38*), excluding sequences exceeding 1,024 residues. Contact prediction quality is assessed on long-range contacts (sequence separation ≥ 6) using precision at *L* (P@L), P@L/2, P@L/5, mean binned precision (AUC), and ROC-AUC, following the previous evaluation protocol (*39*). We report AUC as the primary metric (**Categorical Jacobian (CASP16)** in all figures.

#### Supervised property and structure prediction

We evaluate models on a suite of 17 classification and regression tasks from the xTrimo benchmark suite (*40*), excluding *temperature stability* due to its large size. We evaluate under two paradigms: feature extraction with a frozen backbone and a shallow task-specific head, and full fine-tuning where all parameters are updated after the first epoch. Residue-level embeddings are extracted from the last layer and mean-pooled for sequence-level tasks. Test performance on all 17 tasks are reported under an averaged *Property prediction* metric, based on the F1 score for classification tasks and the Matthews correlation coefficient for regression tasks.

Performance on these tasks is sensitive to hyperparameter choices. We therefore evaluate across multiple random seeds with a systematic grid search: a standard grid for intermediary ablations (REST.) and an extended grid for final models (EXT.). For full task descriptions and data splits, we refer the reader to the original benchmark description (*40*).

**Transfer learning evaluation**. Transfer is quantified as the difference in score between prediction heads trained on top of embeddings derived from pretrained models and randomly re-initialized model weights (Figure 1E). To ensure that the transfer learning results reported in Figure 1E reflect general properties of pretraining data rather than architecture or scale-specific effects, we measure transfer across five models spanning two families: AMPLIFY 120M and 350M (*6*), and ESM2 35M, 150M, and 650M (*2*). All transfer scores reported for the randomized-versus-pretrained comparison are means ± standard deviations across these five models.

### Experimental validation (Enzyme Discovery)

#### Candidate selection (96 sequences x 4 plates)

##### Target selection and filtering

We process UniProtKB entries annotated as belonging to the EC 1.14.11 enzyme class. Given the large number of sequences annotated within this class, a series of straightforward filters was applied to retain sequences that a practitioner would typically consider well-characterized members of this family.

Sequences shorter than 200 or longer than 450 amino acids were removed, as well as any sequences containing ambiguous amino acids (X, O, U, B, J, Z). Sequences lacking the *HXD/E* motif were also removed: this conserved pattern of histidine, any amino acid, and an aspartate or glutamate is responsible for chelating the catalytic iron in the active site and is a defining feature of Fe/αKG-dependent enzymes. Transmembrane proteins were excluded from this study, as they are incompatible with our validation assay for soluble protein expression.

A second set of distribution-based filters was then applied to remove outlier sequences relative to the broader dataset. The bottom 5% of sequences by mean pLDDT score from AlphaFold2 (*17*) structure predictions were removed, as persistently low confidence across a full sequence is indicative of intrinsically disordered or poorly folded proteins unlikely to harbor a well-defined active site, and more broadly suggestive of misannotation or pseudogenes. Additionally, the top 5% of sequences by KL divergence from the expected amino acid composition were discarded, as an atypical composition signals deviation from natural evolutionary constraints. The expected amino acid frequencies were taken from King and Jukes (*41*). Together, these filters reduced the candidate pool from 95,165 to 34,539 sequences.

##### Maximal marginal relevance

A diverse set of high-quality targets was selected from the ~34,000 UniProtKB candidates using the Maximal Marginal Relevance (MMR) algorithm. MMR is an iterative greedy selection procedure that balances two competing objectives: the intrinsic quality of each candidate and its diversity relative to the sequences already selected. At each iteration, the candidate *c* maximizing:

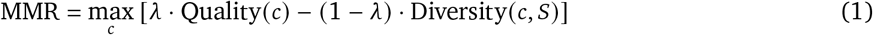

is added to the selected set *S*. Quality is measured as the normalized perplexity of the candidate under AMPLIFY-C,

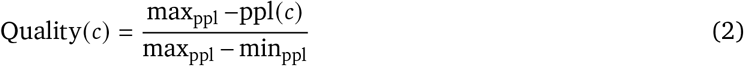

so that lower perplexity, reflecting higher model confidence in the sequence, yields a higher quality score. Diversity is computed by taking the maximum cosine similarity between the candidate embedding and any sequence embedding already in *S*,

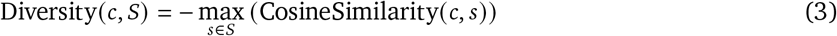

ensuring that sequences too similar to already selected candidates are deprioritized. The trade-off between the two objectives is controlled by *λ*: a value close to 1 favors quality, while a value close to 0 favors diversity.

##### pLM-guided retrieval from EC 1.14.11: Plate A, quality-focused

This experiment prioritizes selecting the highest-confidence enzymes with a relaxed diversity requirement (*λ* = 0.9). An additional hard constraint is enforced so that candidates have at most 50% sequence identity to any other candidate within Plate 1.

##### pLM-guided retrieval from EC 1.14.11: Plate B, diversity-focused

This experiment prioritizes covering the embedding space by selecting candidates distinct from each other and from Plate 1 (*λ* = 0.1). The hard constraint is that candidates must have ≤ 50% sequence identity to any candidate currently selected for Plate 2 and those already selected in Plate 1.

##### pLM-agnostic retrieval from EC 1.14.11: Plate C, control

Candidate sequences were selected from a starting pool identical to experiments 1 and 2. However, the selection was agnostic to pLM scores and based solely on sequence diversity. Concretely, the lambda parameter was set to zero in the MMR score function, and the diversity term was computed with BLAST scores (*42*) rather than embedding similarity. We kept the hard constraint that candidates have at most 50% sequence identity to any other candidates within this plate.

##### pLM-guided retrieval from BFD and MGnify: Plate D, metagenomic

Candidates were retrieved from metagenomic datasets (*BFD* and *MGnify*). The databases were first pre-processed by removing the bottom 50% of sequences based on their RED score, relative to AMPLIFY 350M. Given the computational cost of scoring the entire datasets, pre-computed scores from AMPLIFY 350M were reused rather than generating new scores with AMPLIFY-C. Length (200-450) and H.D/E motif filters were additionally applied, reducing the dataset from 4.5 billion to ~500 million sequences.

Embeddings were then computed for the remaining sequences using AMPLIFY-C, and a k-nearest neighbors search was performed to retrieve the 1,000 closest sequences for each of the UniProtKB enzyme targets from previous plates using cosine similarity distance. Sequences sharing more than 80% sequence identity with any selected UniProtKB target were subsequently removed, as were fragment sequences not starting with methionine, yielding 44,005 unique sequences. A larger number of candidate sequences could be obtained by increasing the number of selected nearest neighbors. This was capped at 1,000 to keep only the closest targets to the selected candidates from other plates. Final candidate selection was performed by running the MMR optimization algorithm, as in plates 1 and 2, with *λ* = 0.9 and a hard threshold of 50% on the maximum pairwise sequence identity among selected sequences. Incidentally, none of the selected metagenomic sequences show any detectable similarity to EC 1.14.11 sequences under MMseqs.

#### Wet lab protocol

##### Recombinant expression in E. coli

The genes encoding the selected sequences were codon-optimized for recombinant expression in Escherichia coli. The codon-optimized DNA sequence was synthesized as an insert within the pET28a(+) vector between the BamHI and HindIII restriction sites. Based on this design, all recombinant proteins were produced in BL21(DE3) E. coli cells with an N-terminal polyhistidine tag. Chemically competent BL21(DE3) E. coli cells were transformed with each plasmid, and the resulting cells were grown in 96-well plates containing 800-µL cultures of Luria–Bertani broth containing 50 µg/mL kanamycin. The cultures were incubated overnight at 37 °C and 80% humidity, with shaking at 1000 rpm. A portion of the overnight cultures (30 µL) was used to inoculate 400-µL cultures of Terrific Broth containing 50 µg/mL kanamycin. In plate format, the cultures were incubated overnight at 37 °C and 80% humidity, with shaking at 1000 rpm. After the desired cell density was reached (OD600 of 0.8–1.5), the expression was induced with the addition of isopropyl *β*-D-1-thiogalactopyranoside to a final concentration of 0.5 mM. The expression cultures were incubated overnight at 18 °C and 80% humidity, with shaking at 1000 rpm. After 20 hours, the plates were centrifuged at 3500 rpm for 5 minutes. The supernatant was discarded, and the plates containing the remaining cell pellets containing recombinant protein were stored at –80 °C.

##### Protein preparation

Prior to lysing the cells, 96-well plates containing cell pellets were thawed at room temperature. The thawed cells were resuspended in lysis buffer (100 mM potassium phosphate, pH 7.5, 2 mg/mL lysozyme, 10 U/mL benzonase, 0.1 mM phenylmethylsulfonyl fluoride) and incubated at 22 °C with shaking at 1000 rpm. After 2 hours, the plates were centrifuged at 3500 rpm for 5 minutes. A portion of the supernatant (20 µL) was saved for SDS-PAGE analysis. To capture the His-tagged recombinant proteins, 50 µL of equilibrated Ni-NTA resin was added to the cleared lysate, and the plates were incubated with shaking for 30 minutes. The resulting slurries were transferred to 96-well filter plates and washed with buffer to remove unspecific interactions. Finally, the His-tagged proteins were eluted with buffer containing 250 mM imidazole. The resulting semi-purified protein samples were analyzed by SDS-PAGE.

##### SDS-PAGE analysis

All collected lysate and semi-purified protein samples were diluted 1:1 in LDS dye containing reducing agent (10 mM DTT) and heated at 80 °C for 10 minutes. Each sample (10 µL) was loaded onto a 4-12% Bis-Tris MOPS gel and run for 45 minutes. Gels were rinsed in water and stained with Imperial Stain for 1 hour, followed by a 2-day destain in water. Recombinant expression success was quantified based on the band volume in the lysate gels at the expected molecular weight for the recombinant protein. The presence of the band in gels following Ni-NTA enrichment was used to validate the successful recombinant expression of the target His-tagged sequence.

#### Wet lab analysis

##### Embedding projections

Two-dimensional projections of sequence embeddings (Figure 3E, Figure 4B) were computed with UMAP (*20*) on mean-pooled AMPLIFY-C embeddings (960 dimensions). We used 200 nearest neighbors, a minimum distance of 0.5, 2 components, and 1,000 optimization epochs. The UMAP model was fit on the entire enzyme class 1.14.11 and reused to project metagenomic candidates into the same coordinate space.

##### In silico expression prediction

We predicted enzyme expression as a binary outcome for 287 sequences from Plates 1, 2, and 4 (UniProtKB). Four feature families were compared: (i) AMPLIFY-C embeddings (960-d) mean-pooled over residues, (ii) 22 sequence-derived biophysical descriptors, (iii) four structural descriptors from AlphaFold-predicted structures (mean pLDDT, mean SASA, ESM-IF and soluble-ProteinMPNN log-likelihoods), and (iv) their concatenation. All features were standardized on training-split statistics.

Two classifiers were compared: *L*_2_-regularized logistic regression and LightGBM (*43*). Hyperparameters were selected per (feature × model × seed) combination by stratified 5-fold cross-validation scored on ROC-AUC, using Bayesian optimization with Optuna (*44*) (50 trials) for LightGBM, and grid search over *C* ∈ {10^−4^, …, 10^2^} for logistic regression. Each configuration was evaluated over 20 independent 85%/15% stratified splits. Pairwise differences were assessed by paired permutation tests (10,000 permutations) with Benjamini–Hochberg correction at 5% false discovery rate.

##### On activity

The selection combines two signals: perplexity, which captures whether a sequence is a viable protein and is an effective filter for expression, and embedding proximity to characterized enzymes, which goes one step further. The first signal is confirmed: pLM-augmented retrieval candidates have higher expression levels. Retrieval rests on the stronger hypothesis that nearby embeddings encode functionally related proteins, which raises a natural question: are the retrieved metagenomic candidates catalytically active? While characterizing catalysis with systematic activity screening is hardly feasible due to enzymes in this class being highly substrate-specific, so no subset of substrates can be rationally screened to conclude whether the overexpressed enzymes are active or not, we nonetheless assayed a small set of substrates and observed conversion to product for several enzymes, showing that at least some of the retrieved proteins are catalytically competent. Some of these substrates deviate from the classic amino-acid hydroxylation, suggesting that the panel may unlock activity not observed in the well-studied enzymes of this family.

