## Supplementary information for "pLM representations unlock metagenomic space beyond homology"

**Lola Le Breton[1,2], David Heurtel-Depeiges[1,2], Douglas C. Millar[3], Lara E. Zetzsche[3], Robert M. Vernon[3], Christopher James Langmead[4,†], Sarath Chandar[1,2,5,6], and Quentin Fournier[1,5]**
[1]Mila – Quebec AI Institute, [2]Polytechnique Montréal, [3]Amgen, [4]Danaher Corporation, [5]University of Montréal, [6]Canada CIFAR AI Chair

**This file includes:**

- Supplementary text and consolidated Materials and Methods (Sections S1–S12)
- Figs. S1 to S24
- Tables S1 to S24

**Other supplementary materials (separate files):**

- `in_silico_results.csv` — per-model in silico scores across evaluation axes (§S6)
- `property_prediction_results.csv` — per-model, per-task supervised property prediction scores (§S6.1)
- `categorical_jacobian_results.csv` — per-model CASP16 contact metrics (§S6.5)
- `human_proteome.csv` — human proteome query set for the calibration task (§S7.1)
- `human_consensus_positions.csv` — per-position homolog amino-acid frequency profiles (§S7.1)
- `ddg_results_random.csv` — per-mutation Rosetta $\Delta\Delta G$ for the random-substitution control (§S7.4)
- `ddg_results.csv` — per-mutation Rosetta $\Delta\Delta G$ for consensus substitutions (§S7.4)
- `kde_boundaries.csv` — per-dataset RED filtering thresholds (§S5)
- `all_plates_features.parquet` — per-candidate in silico features for all four plates (§S9)
- `pairwise_cosine_similarity.parquet` — AMPLIFY-C embedding cosine-similarity matrix across candidates (§S8.4)
- `in_vivo_results.csv` — wet-lab expression measurements (lysate/purified band) and synthesized-construct details for all four plates (§S9)
- `metagenomic_additional_features.csv` — per-candidate Plate D OpenFold pLDDT and MMseqs2 identity to EC 1.14.11 (§S11)
- `expression_prediction_results.csv` — per-split expression-classifier ROC-AUC (§S10)
- `literature_hydroxylases.csv` — hydroxylases characterized in the literature

### S1. Reading guide and notation

This document provides the extended methods, supporting data, and supplementary figures for the main text. The main text is organized around six result blocks. The table below maps each key claim to the SI section where supporting material can be found.

#### Result-to-SI mapping

***R1 — RED: a scalable proxy for perplexity***

- Mean PPL: human proteome (3.57), DisProt (4.21), shuffled (17.88); PE = 5 by annotation level (17.00 → 4.70 → 7.14) §S4
- PPL across AMPLIFY 350M and ESM-2 (35M, 150M, 650M) §S4
- Scoring cost: ~90 GPU-years (PPL) vs. 25 days (RED) on BFD+MGnify §S3, §S4
- RED–PPL Spearman $r = -0.97$ (10k sequences) §S4
- KDE boundaries converge to RED $\in [0.11, 0.13]$ across datasets §S5

***R2 — Unfiltered metagenomic pretraining degrades downstream performance***

- DMS drops: −10.0% (BFD), −13.4% (MGnify); mitigated to −2.0%/−2.5% with UniRef100 §S6
- CASP16 drops: −8.7% (BFD), −2.5% (MGnify) §S6
- Property prediction: 17-task aggregate < 5% variation §S6
- Transfer: stability +57.7%, fold +44.0%, Q3 +30.4%, Q8 +33.7%; TCR pMHC −2.5% §S6.3
- Calibration baselines: T/NT gaps for BFD (−16.7%/−5.7%) and MGnify (−18.1%/−6.7%) §S7
- Rosetta validation: 35.1% stabilizing (consensus) vs. 18.9% (random); $n$ = 454/3,302/1,905 §S7

***R3 — RED filtering and training-scale dynamics***

- Filtering optima align with KDE boundary; gap with UniRef100 persists after filtering §S5, §S6
- Scaling 200B → 2T: UniRef100 plateaus; MGnify ROC-AUC 24.1 vs. UniRef100 22.8 (CASP16) §S6
- UniRef100 NT calibration −14.1 pts (memorization); BFD +11.2, MGnify +3.1 pts §S7, §S6

***R4 — Memorization–generalization trade-off***

- Subsampling 0.03–0.4: structural tasks peak at 10%; DMS drops monotonically §S6
- Clustering 90%–30%: structural tasks peak at 50%; calibration and DMS degrade §S6
- AMPLIFY-B and AMPLIFY-C final model specifications §S3.4

***R5 — pLMs predict wet-lab expression (curated selection)***

- Candidate funnel: 95,165 → 34,539 (HXD/E motif, pLDDT, composition KL); MMR per plate §S8
- SDS-PAGE gels (lysate and Ni-NTA), all four plates §S9
- PPL–expression envelope; Pearson $r = -0.15$ (purified), −0.14 (lysate) §S8.5, §S9.2
- Band volumes; per-sequence data §S9.2
- Codon-optimized sequences, all four plates §S9.2
- Embedding classifiers vs. structural/biophysical features: ROC-AUC, $p < 0.011$ §S10

***R6 — Metagenomic retrieval and validation***

- kNN retrieval funnel: 4.5B → 500M → 44,005 → 96 candidates §S8
- Zero MMseqs2 similarity to EC 1.14.11 (settings, full distribution) §S8.4
- 30 expressed, 24 at high levels; 7 highly expressed with mean pLDDT < 70 §S11
- Predicted structures for expressed candidates §S11

***Additional Methodology***

- Dataset versions, licenses, sequence-accounting table (raw → final token counts) §S2
- Ambiguous amino acid masking (B, O, U, X, Z, J) and RoPE-shift §S2
- Model registry: all 100 ablations (dataset, quantile, clustering, subsampling, tokens) §S3
- HP grids: REST. 108 configs/task; EXT. 2,880 configs/task; rank stability §S6
- Software versions, random seeds, compute budget; statistical methods §S12

##### Notation

| Symbol / abbreviation | Meaning |
|---|---|
| pLM | Protein language model. |
| PPL | Perplexity: $\exp\left(-\frac{1}{L}\sum_{i=1}^{L}\log P(r_i \mid S_{-i})\right)$. One forward pass per residue. |
| pseudo-PPL | Single-pass approximation of PPL using one randomly drawn mask per sequence. |
| RED | Residue Embedding Diversity: $1 - \frac{1}{L(L-1)}\sum_{i\neq j}\frac{e_i\cdot e_j}{\|e_i\|\|e_j\|+\varepsilon}$. One forward pass per sequence. |
| MCS | Mean Cosine Similarity: 1 – RED. Used in cosine-similarity histogram figures. |
| T position | Target: conserved MSA position where the human residue matches the evolutionary consensus. |
| NT position | Non-target: conserved MSA position where the human residue *differs* from the consensus. |
| $\Delta\Delta G$ | Rosetta-estimated free-energy change upon mutation (REU); negative = stabilizing. |
| MMR | Maximal Marginal Relevance: greedy selection balancing quality and diversity with weight $\lambda$. |
| $\lambda$ | MMR trade-off parameter: $\lambda \approx 1$ favors quality, $\lambda \approx 0$ favors diversity. |
| P@L | Precision at $L$ contacts: fraction of top-$L$ predicted long-range contacts that are true contacts. |
| ROC-AUC | Area under the receiver operating characteristic curve for contact prediction. |
| REST. | Restricted hyperparameter grid (108 configs/task; used for subsampling ablations). |
| EXT. | Extended hyperparameter grid (2,880 configs/task; used for all other models). |
| BH | Benjamini–Hochberg false-discovery-rate correction at 5%. |

Table S1: Notation and abbreviations used throughout the Supplementary Materials.

#### S2. Datasets and preprocessing

##### S2.1. Databases and versions

Four databases are used for pretraining.

**UniRef100** (UniProtKB Reference Clusters, release 2024_05) provides high-quality, clustered protein sequences and annotations. Download: https://ftp.uniprot.org/pub/databases/uniprot/uniref/uniref100/. License: Creative Commons Attribution 4.0 International (CC BY 4.0).

**BFD** (Big Fantastic Database) is used as a source of metagenomic diversity. Preprocessing removes artificial consensus sequences and entries originating from TrEMBL/Swiss-Prot, capitalizes all residues, and strips alignment gaps. Download: https://bfd.mmseqs.com/. License: Creative Commons Attribution 4.0 International (CC BY 4.0).

**MGnify** (EBI-Metagenomics, release 2024_04) maximizes coverage of the microbial world. All sequences are retained regardless of full-length indicators. Download: https://www.ebi.ac.uk/metagenomics/. License: Creative Commons Zero 1.0 Universal (CC0 1.0).

**OAS** (Observed Antibody Space (*1*)) paired antibody sequences are included for the two final specialized models (AMPLIFY-B and AMPLIFY-C) only. Each paired entry consists of a heavy chain and a light chain; both chain orderings (heavy|light and light|heavy) are added as separate training sequences. Download: https://opig.stats.ox.ac.uk/webapps/oas/. License: Creative Commons Attribution 4.0 International (CC BY 4.0).

The **Validation set** is constructed following the protocol described in AMPLIFY (*2*). The validation set contains 10k sequences sampled from UniProt reference proteomes with high completeness (> 80% BUSCO score), covering all three domains of the phylogenetic tree of life, and excluding proteins lacking experimental evidence (PE = 1 or 2). In addition, it contains 10k paired OAS sequences (light and heavy chains) randomly sampled from representatives of 90% similarity clusters. SCOP sequences were not included in the validation set. All training databases are deduplicated against this validation set using a 90% sequence identity threshold, except for OAS which uses a 99% threshold.

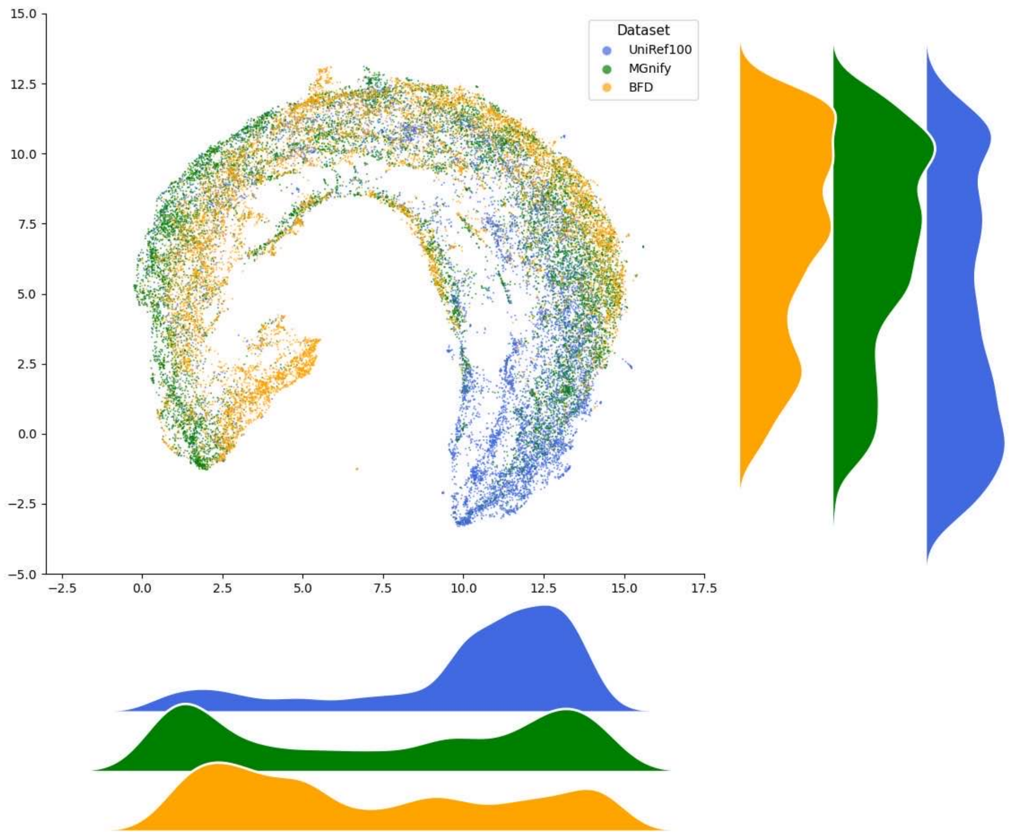

**Figure** S1: **UMAP density of sequence embeddings** for UniRef100, BFD, MGnify (10k sequences each) under AMPLIFY 350M.

| Database | Raw seqs | Train seqs | Avg length | Min length | Max length | Seqs w/ ambig. AA |
|---|---|---|---|---|---|---|
| UniRef100 | 420,305,229 | 418,829,208 | 383.9 | 2 | 49,499 | 3,778,047 |
| BFD | 2,032,538,970 | 2,032,203,748 | 684.6 | 49 | 37,324 | 718,353,100 |
| MGnify | 2,455,939,992 | 2,455,358,433 | 189.7 | 20 | 57,795 | 34,390,412 |
| OAS | 3,774,974 | 1,971,900 | 115.6 | 99 | 164 | — |

Table S2: Sequence statistics per pretraining database. *Raw seqs*: sequences in the downloaded database after basic preprocessing. *Train seqs*: sequences remaining after deduplication against the validation set (90% identity threshold); this is the pool from which RED filtering and subsampling are applied. *Avg / Min / Max length*: computed on train sequences, in amino acids. *Seqs w/ ambig. AA*: number of train sequences containing at least one character from {B, O, U, X, Z, J} (these sequences are retained with the ambiguous positions masked).

#### S2.2. Ambiguous amino acids and RoPE-shift masking

Open-source databases frequently contain non-standard amino acid characters resulting from sequencing ambiguity or the presence of non-canonical residues. The characters encountered and their biological interpretation are listed in Table S3.

| Character | Meaning |
|---|---|
| O | Pyrrolysine |
| U | Selenocysteine |
| B | Aspartate or asparagine (ambiguous) |
| Z | Glutamate or glutamine (ambiguous) |
| J | Leucine or isoleucine (ambiguous) |
| X | Wholly unresolved read |

Table S3: Ambiguous or non-canonical amino acid characters and their meaning. All six characters are removed from the input token sequence. The corresponding positions are accounted for in the positional index via RoPE-shift (see text).

Rather than discarding sequences that contain these characters, we retain the surrounding valid residues and mask only the ambiguous positions. Rotary Position Embeddings (RoPE) (*3*) are shifted for all residues following a masked position so that the model sees correct positional distances between the remaining canonical residues, without ever being asked to predict a token outside the standard 20-amino-acid vocabulary.

The following example is drawn from sequence scaffold1532777_1 in FD_GraSoiStandDraft_30_1057271. The crop spans positions 122–160. The 24-residue run of X characters (positions 129–152) represents wholly unresolved reads.

*Original sequence:*

| | | | | | | | | | | | | | | |
|---|---|---|---|---|---|---|---|---|---|---|---|---|---|---|
| D | T | D | S | T | F | I | X | X | ··· | X | X | E | I | R |
| 122 | 123 | 124 | 125 | 126 | 127 | 128 | 129 | 130 | | 151 | 152 | 153 | 154 | 155 |

*After masking: X residues removed, RoPE indices preserved:*

| | | | | | | | | | | | | | | |
|---|---|---|---|---|---|---|---|---|---|---|---|---|---|---|
| D | T | D | S | T | F | I | E | I | R | K | Q | L | I | A |
| 122 | 123 | 124 | 125 | 126 | 127 | 128 | 153 | 154 | 155 | 156 | 157 | 158 | 159 | 160 |

The 24 X tokens are dropped from the input, but the RoPE indices of the downstream residues are preserved at their original values (153, 154, . . . ) rather than being renumbered from 129 onward. The model therefore encodes the positional distance between I (128) and E (153) as 25 steps, preserving correct positional relationships among the retained canonical residues.

#### S2.3. Tokenization and input representation

The AMPLIFY tokenizer uses a vocabulary of 32 tokens listed in Table S4. Five are special tokens; one (|) is a chain separator used for paired antibody sequences in OAS; six are ambiguous amino acid characters that are masked during preprocessing (section S2); and the remaining 20 are the canonical amino acids.

Sequences are encoded as a flat token sequence with no added <bos> or <eos> tokens. The maximum sequence length is 512 tokens for ablation models and 2,048 tokens for the context-extension stage of the final models. Sequences exceeding the maximum length are randomly truncated.

During training, 15% of positions are selected for masking. All selected positions are replaced with the <mask> token (i.e., a pure masking scheme, without the 80/10/10 random-replacement variant used in the original BERT).

| Index | Token | Type | Note |
|---|---|---|---|
| 0 | <pad> | Special | Padding |
| 1 | <unk> | Special | Unknown |
| 2 | <mask> | Special | Masked position (MLM target) |
| 3 | <bos> | Special | Beginning of sequence |
| 4 | <eos> | Special | End of sequence |
| 5 | \| | Special | Chain separator (OAS paired sequences) |
| 6 | X | Ambiguous AA | Unresolved read |
| 7 | B | Ambiguous AA | Asp or Asn |
| 8 | O | Ambiguous AA | Pyrrolysine |
| 9 | U | Ambiguous AA | Selenocysteine |
| 10 | Z | Ambiguous AA | Glu or Gln |
| 11 | J | Ambiguous AA | Leu or Ile |
| 12 | L | Canonical AA | Leucine |
| 13 | A | Canonical AA | Alanine |
| 14 | G | Canonical AA | Glycine |
| 15 | V | Canonical AA | Valine |
| 16 | S | Canonical AA | Serine |
| 17 | E | Canonical AA | Glutamic acid |
| 18 | R | Canonical AA | Arginine |
| 19 | T | Canonical AA | Threonine |
| 20 | I | Canonical AA | Isoleucine |
| 21 | D | Canonical AA | Aspartic acid |
| 22 | P | Canonical AA | Proline |
| 23 | K | Canonical AA | Lysine |
| 24 | Q | Canonical AA | Glutamine |
| 25 | N | Canonical AA | Asparagine |
| 26 | F | Canonical AA | Phenylalanine |
| 27 | Y | Canonical AA | Tyrosine |
| 28 | M | Canonical AA | Methionine |
| 29 | H | Canonical AA | Histidine |
| 30 | W | Canonical AA | Tryptophan |
| 31 | C | Canonical AA | Cysteine |

Table S4: AMPLIFY tokenizer vocabulary (32 tokens). The model is trained to predict only the 20 canonical amino acids; ambiguous characters are masked out of the input prior to training (§S2).

### S3. Model architecture, training, and compute

#### S3.1. Architectures

All AMPLIFY models are transformer encoder pLMs using pre-layer normalization, Rotary Position Embeddings (RoPE, $\theta = 10{,}000$), and SwiGLU activations. Each encoder block comprises: a fused QKV projection (no bias), an output projection $W_O$, and a two-gate SwiGLU FFN whose hidden dimension is $\lfloor 2/3 \cdot d_{\mathrm{FFN}} \rfloor$ rounded to the nearest multiple of 8. Layer normalization uses RMSNorm ($\varepsilon = 10^{-5}$) applied before attention and before the FFN. A final RMSNorm precedes the output decoder.

All ablation experiments use the 120M configuration (24 layers, hidden 640). The two specialized final models (AMPLIFY-B and AMPLIFY-C) use the 350M configuration.

Transfer-learning experiments additionally use ESM2 checkpoints `esm2_t12_35M_UR50D` (35M), `esm2_t30_150M_UR50D` (150M), and `esm2_t33_650M_UR50D` (650M), and AMPLIFY checkpoints `chandar-lab/AMPLIFY_120M` and `chandar-lab/AMPLIFY_350M` from Hugging Face (2). The performance improvements of AMPLIFY-B and AMPLIFY-C are also computed relative to `chandar-lab/AMPLIFY_350M`, a model identical in architecture.

| Hyperparameter | AMPLIFY 120M | AMPLIFY 350M |
|---|---|---|
| Layers ($L$) | 24 | 32 |
| Hidden size ($d$) | 640 | 960 |
| Attention heads ($H$) | 10 | 15 |
| Head dimension ($d/H$) | 64 | 64 |
| FFN intermediate ($d_{\text{FFN}}$) | 2,560 | 3,840 |
| SwiGLU actual hidden | 1,712 | 2,560 |
| Vocabulary size | 32 | 32 |
| Context length (Stage 1) | 512 | 512 |
| Context length (Stage 2) | — | 2,048 |
| RoPE $\theta$ | 10,000 | 10,000 |
| Parameters (approx.) | 118M | 354M |

Table S5: Architecture hyperparameters for the two AMPLIFY model sizes.

##### S3.2. Training protocol and optimizer

**Optimizer..** All models are trained with AdamW (*4*) with peak learning rate $10^{-3}$, betas (0.9, 0.95), $\varepsilon = 10^{-8}$, and weight decay 0.01. Gradient clipping is applied at norm 1.

**Learning rate schedule..** A cosine decay schedule is used with 1,000 linear warmup steps, followed by cosine decay of 90,000 or 900,000 steps to a final learning rate of $10^{-4}$ (0.1× the peak) for the last 10% of training.

**Masking..** 15% of token positions are selected uniformly at random and replaced with `<mask>` (pure masking; no random-replacement or keep-original variant). Ambiguous amino acid positions are excluded from both input and loss (§S2).

**Batch size and sequence packing..** All models are trained with the flash-attention packed variant (flash-attn 2.8.3.post1 (*5*)), which eliminates padding overhead. Training uses HuggingFace Accelerate 1.4.0 (*6*) with DeepSpeed ZeRO-2 (0.14.4) (*7*) and PyTorch 2.5.0, with `bfloat16` mixed precision and `tf32` enabled.

**Two-stage training for specialized models..** AMPLIFY-B and AMPLIFY-C are each trained in two stages:

1. **Stage 1:** 1,000,000 steps at a maximum sequence length of 512.
2. **Stage 2:** 50,000 steps at a maximum sequence length of 2,048 (context extension).

The combined token budget is approximately 2.1T tokens.

##### S3.3. Model registry (all ablations)

All ablation models use the 120M architecture, a single seed, and bfloat16 mixed precision. All are trained for 100k steps with a complete schedule (at a batch size of 2M tokens) unless the model name carries the `_1m` suffix, in which case they were trained for 1M steps.

###### *Reference models*

| Model | UniRef100 | BFD | MGnify | Seq w. Ambig. AA excluded | Train seqs. |
|---|---|---|---|---|---|
| MILA_BFD | × | ✓ | × | No | 2,032,203,748 |
| MILA_MGnify | × | × | ✓ | No | 2,455,358,433 |
| MILA_BFD_no_ambig | × | ✓ | × | Yes | 1,313,850,648 |
| MILA_MGnify_no_ambig | × | × | ✓ | Yes | 2,420,968,021 |
| MILA_UR100_BFD_no_ambig | ✓ | ✓ | × | Yes | 1,730,377,830 |
| MILA_UR100_MGnify_no_ambig | ✓ | × | ✓ | Yes | 2,836,019,182 |
| MILA_UR100_BFD_MGnify_no_ambig | ✓ | ✓ | ✓ | Yes | 4,149,869,830 |

Table S6: Baseline models trained on unfiltered mixtures of the three train datasets. No filtering, subsampling, or clustering applied.

| Model | Steps | RED quantile | Train seqs (UniRef100) |
|---|---|---|---|
| MILA_U100_0.00 | 100k | 0.00 | 418,829,208 |
| MILA_U100_0.01 | 100k | 0.01 | 414,640,916 |
| MILA_U100_0.02 | 100k | 0.02 | 410,452,624 |
| MILA_U100_0.03 | 100k | 0.03 | 406,264,332 |
| MILA_U100_0.05 | 100k | 0.05 | 397,887,748 |
| MILA_U100_0.05_1m | 1M | 0.05 | 397,887,748 |
| MILA_U100_0.10 | 100k | 0.10 | 376,946,287 |
| MILA_U100_0.10_1m | 1M | 0.10 | 376,946,287 |
| MILA_U100_0.15 | 100k | 0.15 | 356,004,827 |
| MILA_U100_0.15_1m | 1M | 0.15 | 356,004,827 |
| MILA_U100_0.20 | 100k | 0.20 | 335,063,366 |
| MILA_U100_0.20_1m | 1M | 0.20 | 335,063,366 |
| MILA_U100_0.25 | 100k | 0.25 | 314,121,906 |
| MILA_U100_0.25_1m | 1M | 0.25 | 314,121,906 |
| MILA_U100_0.30 | 100k | 0.30 | 293,180,446 |
| MILA_U100_0.30_1m | 1M | 0.30 | 293,180,446 |
| MILA_U100_0.35 | 100k | 0.35 | 272,238,985 |
| MILA_U100_0.35_1m | 1M | 0.35 | 272,238,985 |
| MILA_U100_0.40 | 100k | 0.40 | 251,297,525 |
| MILA_U100_0.40_1m | 1M | 0.40 | 251,297,525 |
| MILA_U100_0.50 | 100k | 0.50 | 209,414,604 |
| MILA_U100_0.60 | 100k | 0.60 | 167,531,683 |
| MILA_U100_0.65 | 100k | 0.65 | 146,590,223 |
| MILA_U100_0.75 | 100k | 0.75 | 104,707,302 |

Table S7: RED filtering ablations on UniRef100 only. The RED quantile represents the fraction of lowest-scoring sequences removed.

| Model | Steps | RED quantile | Train seqs (BFD) |
|---|---|---|---|
| MILA_BFD_0.10 | 100k | 0.10 | 1,828,983,373 |
| MILA_BFD_0.20 | 100k | 0.20 | 1,625,762,998 |
| MILA_BFD_0.20_1m | 1M | 0.20 | 1,625,762,998 |
| MILA_BFD_0.30 | 100k | 0.30 | 1,422,542,624 |
| MILA_BFD_0.30_1m | 1M | 0.30 | 1,422,542,624 |
| MILA_BFD_0.40 | 100k | 0.40 | 1,219,322,249 |
| MILA_BFD_0.40_1m | 1M | 0.40 | 1,219,322,249 |
| MILA_BFD_0.50 | 100k | 0.50 | 1,016,101,874 |
| MILA_BFD_0.50_1m | 1M | 0.50 | 1,016,101,874 |
| MILA_BFD_0.60 | 100k | 0.60 | 812,881,499 |
| MILA_BFD_0.60_1m | 1M | 0.60 | 812,881,499 |
| MILA_BFD_0.70 | 100k | 0.70 | 609,661,124 |
| MILA_BFD_0.70_1m | 1M | 0.70 | 609,661,124 |
| MILA_BFD_0.80 | 100k | 0.80 | 406,440,750 |
| MILA_BFD_0.80_1m | 1M | 0.80 | 406,440,750 |

Table S8: RED filtering ablation on BFD only.

| Model | Steps | RED quantile | Train seqs (MGnify) |
|---|---|---|---|
| MILA_MGnify_0.10 | 100k | 0.10 | 2,209,822,590 |
| MILA_MGnify_0.20 | 100k | 0.20 | 1,964,286,746 |
| MILA_MGnify_0.20_1m | 1M | 0.20 | 1,964,286,746 |
| MILA_MGnify_0.30 | 100k | 0.30 | 1,718,750,903 |
| MILA_MGnify_0.30_1m | 1M | 0.30 | 1,718,750,903 |
| MILA_MGnify_0.40 | 100k | 0.40 | 1,473,215,060 |
| MILA_MGnify_0.40_1m | 1M | 0.40 | 1,473,215,060 |
| MILA_MGnify_0.50 | 100k | 0.50 | 1,227,679,216 |
| MILA_MGnify_0.60 | 100k | 0.60 | 982,143,373 |
| MILA_MGnify_0.70 | 100k | 0.70 | 736,607,530 |
| MILA_MGnify_0.80 | 100k | 0.80 | 491,071,687 |
| MILA_MGnify_0.90 | 100k | 0.90 | 245,535,843 |

Table S9: RED filtering ablation on MGnify only.

| Model | Steps | RED $q$ | Seqs (UR100) | Seqs (BFD) | Seqs (MGnify) |
|---|---|---|---|---|---|
| MILA_ALL_0.30 | 100k | 0.30 | 293,180,446 | 1,422,542,624 | 1,718,750,903 |
| MILA_ALL_0.40 | 100k | 0.40 | 251,297,525 | 1,219,322,249 | 1,473,215,060 |
| MILA_ALL_0.50 | 100k | 0.50 | 209,414,604 | 1,016,101,874 | 1,227,679,216 |
| MILA_ALL_0.50_1m | 1M | 0.50 | 209,414,604 | 1,016,101,874 | 1,227,679,216 |
| MILA_ALL_0.60 | 100k | 0.60 | 167,531,683 | 812,881,499 | 982,143,373 |
| MILA_ALL_0.60_1m | 1M | 0.60 | 167,531,683 | 812,881,499 | 982,143,373 |
| MILA_ALL_0.70 | 100k | 0.70 | 125,648,762 | 609,661,124 | 736,607,530 |
| MILA_ALL_0.70_1m | 1M | 0.70 | 125,648,762 | 609,661,124 | 736,607,530 |
| MILA_ALL_0.80 | 100k | 0.80 | 83,765,842 | 406,440,750 | 491,071,687 |
| MILA_ALL_0.80_1m | 1M | 0.80 | 83,765,842 | 406,440,750 | 491,071,687 |
| MILA_ALL_0.90 | 100k | 0.90 | 41,882,921 | 203,220,375 | 245,535,843 |

Table S10: RED filtering applied uniformly across all three pretraining datasets simultaneously.

| Model | Steps | $s$ | Seqs (UR100) | Seqs (BFD) | Seqs (MGnify) |
|---|---|---|---|---|---|
| MILA_OPT_0.10_sub10 | 100k | 0.10 | 335,063,366 | 40,644,075 | 171,875,090 |
| MILA_OPT_0.10_sub10_1m | 1M | 0.10 | 335,063,366 | 40,644,075 | 171,875,090 |
| MILA_OPT_0.20_sub20 | 100k | 0.20 | 335,063,366 | 81,288,150 | 343,750,181 |
| MILA_OPT_0.20_sub20_1m | 1M | 0.20 | 335,063,366 | 81,288,150 | 343,750,181 |
| MILA_OPT_0.30_sub30_1m | 1M | 0.30 | 335,063,366 | 121,932,225 | 515,625,271 |
| MILA_OPT_0.40_sub40_1m | 1M | 0.40 | 335,063,366 | 162,576,300 | 687,500,361 |
| MILA_OPT_0.50_sub50 | 100k | 0.50 | 335,063,366 | 203,220,375 | 859,375,452 |
| MILA_OPT_0.60_sub60 | 100k | 0.60 | 335,063,366 | 243,864,450 | 1,031,250,542 |

Table S11: All datasets filtered at fixed per-dataset RED quantiles (UniRef100: $q = 0.20$; BFD: $q = 0.80$; MGnify: $q = 0.30$); metagenomic datasets (BFD and MGnify) further subsampled at rate $s$ after filtering. UniRef100 is not subsampled.

| Model | Steps | $s$ | RED thresh. | Seqs (UR100) | Seqs (BFD) | Seqs (MGnify) |
|---|---|---|---|---|---|---|
| MILA_RED_0.03_0.88 | 100k | 0.03 | 0.115 | 351,816,535 | 32,921,701 | 40,513,414 |
| MILA_RED_0.05_0.88 | 100k | 0.05 | 0.115 | 351,816,535 | 54,869,501 | 67,522,357 |
| MILA_RED_0.10_0.88 | 100k | 0.10 | 0.115 | 351,816,535 | 109,739,002 | 135,044,714 |
| MILA_RED_0.20_0.88 | 100k | 0.20 | 0.115 | 351,816,535 | 219,478,005 | 270,089,428 |
| MILA_RED_0.30_0.88 | 100k | 0.30 | 0.115 | 351,816,535 | 329,217,007 | 405,134,141 |
| MILA_RED_0.40_0.88 | 100k | 0.40 | 0.115 | 351,816,535 | 438,956,010 | 540,178,855 |

Table S12: All datasets filtered with a common RED threshold of 0.115; metagenomic datasets (BFD and MGnify) further subsampled at rate $s$ after filtering.

| Model | Steps | Clustering identity | Fixed epochs | Train seqs (UniRef100) |
|---|---|---|---|---|
| MILA_U100_clust90 | 100k | 90% | No | 199,268,886 |
| MILA_U100_clust90_fixed_epochs | 2 ep. | 90% | Yes | 199,268,886 |
| MILA_U100_clust80 | 100k | 80% | No | 141,071,243 |
| MILA_U100_clust80_fixed_epochs | 2 ep. | 80% | Yes | 141,071,243 |
| MILA_U100_clust70 | 100k | 70% | No | 104,962,993 |
| MILA_U100_clust70_fixed_epochs | 2 ep. | 70% | Yes | 104,962,993 |
| MILA_U100_clust60 | 100k | 60% | No | 81,461,435 |
| MILA_U100_clust60_fixed_epochs | 2 ep. | 60% | Yes | 81,461,435 |
| MILA_U100_clust50 | 100k | 50% | No | 65,847,724 |
| MILA_U100_clust50_fixed_epochs | 2 ep. | 50% | Yes | 65,847,724 |
| MILA_U100_clust40 | 100k | 40% | No | 54,339,116 |
| MILA_U100_clust40_fixed_epochs | 2 ep. | 40% | Yes | 54,339,116 |
| MILA_U100_clust30 | 100k | 30% | No | 49,769,155 |
| MILA_U100_clust30_fixed_epochs | 2 ep. | 30% | Yes | 49,769,155 |

Table S13: Group 8: clustering ablation on UniRef100. Sequences are clustered at the given identity threshold before training. *Fixed epochs* models run for 2 epochs over the clustered set rather than a fixed step count.

| Model | Description |
|---|---|
| AMPLIFY_120M | Public AMPLIFY 120M checkpoint (*2*) (chandar-lab/AMPLIFY_120M) |
| AMPLIFY_350M | Public AMPLIFY 350M checkpoint (*2*) (chandar-lab/AMPLIFY_350M) |
| AMPLIFY_benchmark | AMPLIFY-B (350M), final specialized model (§S3) |
| AMPLIFY_consensus | AMPLIFY-C (350M), final specialized model (§S3) |

Table S14: Reference models included in the in silico evaluation results for comparison.

#### S3.4. Specialized models: AMPLIFY-B and AMPLIFY-C

| Property | AMPLIFY-B | AMPLIFY-C |
|---|---|---|
| Architecture | 350M (32L, $d$=960) | 350M (32L, $d$=960) |
| Primary target | Structural modeling | Evolutionary calibration |
| *UniRef100* | | |
| RED quantile | 0.05 | 0.10 |
| Clustering | 50% identity | 90% identity |
| Subsampling | — | — |
| *BFD* | | |
| RED quantile | 0.20 | 0.50 |
| Subsampling | 0.03 | 0.05 |
| *MGnify* | | |
| RED quantile | 0.20 | 0.20 |
| Subsampling | 0.03 | 0.05 |
| *OAS (paired antibodies)* | | |
| Included | Yes | Yes |
| Training stages | 2 (512 + 2,048 ctx) | 2 (512 + 2,048 ctx) |
| Total tokens (approx.) | 2.1T | 2.1T |

Table S15: Data mixture and training configuration for the two specialized final models. RED quantile refers to the fraction of lowest-scoring sequences removed per dataset. AMPLIFY-C is used for all wet-lab candidate selection and metagenomic retrieval experiments.

#### S3.5. Compute budget

All models were trained on NVIDIA H100 GPUs. We account for compute using the standard approximations $C_{\text{train}} \approx 6ND$ for training and $C_{\text{score}} \approx 2ND$ for inference, where $N$ is the number of non-embedding parameters and $D$ the number of tokens processed. These FLOP counts are hardware and implementation-independent. We convert them to GPU-hours through the model FLOP utilization (MFU) measured on our hardware.

To calibrate the MFU, we benchmarked a 350M-parameter run that processed 2T tokens in 118 wall-clock hours on 32 H100 GPUs (3,776 GPU-hours). This corresponds to a sustained $3 \times 10^{14}$ FLOP/s per GPU, or an MFU of $\approx$30% relative to the $10^{15}$ FLOP/s BF16 peak.

Pretraining comprised 72 models (120M parameters, 100k steps), 26 models (120M, 1M steps), and 2 models (350M, 1M steps), all at $\approx$2M tokens per step. Scoring covers a single forward pass over $\approx$5B metagenomic sequences (mean length $\approx$400 tokens) with a 350M model. The resulting budget is summarized in Table S16. All numbers are rounded to one significant figure to reflect estimation uncertainty. In total, GPU work amounts to $\approx 8 \times 10^4$ H100 GPU-hours ($\approx 6 \times 10^{22}$ FLOP, equivalently $\approx$9 GPU-years). The GPU-hour figures for the 120M models are scaled from the 350M MFU.

Homology search was performed separately on CPU: homologs for the ~20,000 human-proteome sequences were identified with HHblits against BFD. At an estimated $\approx$6 hours per query (8 core), this amounts to $\approx 10^6$ CPU-hours.

**RED versus perplexity scoring cost..** The RED score requires a single forward pass per sequence, so scoring the full 5B-sequence pool costs $\approx 2 \times 10^3$ GPU-hours. Computing perplexity over the same pool would instead require one masked forward pass per residue, inflating the cost by a factor in the order of the mean sequence length ($\approx$400×) to $\approx 8 \times 10^5$ GPU-hours ($\approx$90 GPU-years), which is 10 times more than the total compute spent for the entire project. This gap is the practical motivation for RED: it makes metagenomic-scale quality scoring tractable.

| Job | Models | FLOP | GPU-hours |
|---|---|---|---|
| Pretraining — 120M, 100k steps | 72 | $1.0 \times 10^{22}$ | 14,000 |
| Pretraining — 120M, 1M steps | 26 | $3.7 \times 10^{22}$ | 50,000 |
| Pretraining — 350M, 1M steps | 2 | $8.4 \times 10^{21}$ | 11,000 |
| Scoring (RED) — 350M | — | $1.4 \times 10^{21}$ | 1,900 |
| Total | | $5.8 \times 10^{22}$ | ≈80,000 |

Table S16: Compute budget on NVIDIA H100 GPUs. FLOP counts use $6ND$ (training) and $2ND$ (inference) and are codebase-independent. GPU-hours are reported for the experiment codebase (≈31% MFU); the 350M row is anchored by a direct wall-clock measurement, the remaining rows are scaled at the measured MFU.

### S4. Sequence scoring: perplexity and RED

#### S4.1. Perplexity, pseudo-perplexity, and Residue Embedding Diversity (RED)

**Perplexity..** The likelihood of a sequence $S = (r_1, \ldots, r_L)$ of length $L$ decomposes as a product of conditional probabilities, $P(S) = \prod_{i=1}^{L} P(r_i \mid r_{<i})$. Because this quantity decreases with sequence length, researchers commonly use perplexity, a length-normalized measure of predictive confidence. For a masked language model the true perplexity cannot be computed exactly: the model does not have access to a causal factorization. One approximation, *pseudo-perplexity*, masks a random subset of positions and estimates the full perplexity from those conditionals in a single forward pass; this is fast but stochastic and noisy. A more principled approximation masks each residue individually and aggregates the resulting conditionals:

$$\mathrm{PPL}(S) = \exp\left(-\frac{1}{L}\sum_{i=1}^{L} \log P\left(r_i = r_i^S \mid S_{-i}\right)\right), \tag{S1}$$

where $S_{-i}$ denotes $S$ with position $i$ masked. This quantity is deterministic and well-defined, but requires *one forward pass per residue*: scoring a sequence of length $L$ costs $L$ forward passes instead of one. At metagenomic scale this is prohibitive (see §S3.5).

**Residue Embedding Diversity (RED)..** RED is a computationally efficient proxy for perplexity computable in a single forward pass per sequence. It quantifies the diversity of residue-level embeddings: diverse representations (low perplexity) yield high RED; collapsed representations (high perplexity) yield low RED. Formally,

$$\mathrm{RED}(S) = 1 - \frac{1}{L(L-1)} \sum_{\substack{i,j=1 \\ i \neq j}}^{L} \frac{e_i \cdot e_j}{\|e_i\|\,\|e_j\| + \varepsilon}, \tag{S2}$$

where $e_i \in \mathbb{R}^d$ is the embedding vector of the $i$-th residue output by the final encoder layer, and $\varepsilon$ is a small numerical stability constant. RED is motivated by a representation-collapse phenomenon: for low-likelihood sequences, every residue of a given amino acid type collapses to a single identical point in embedding space regardless of its sequential context, so the mean pairwise cosine similarity approaches 1 and RED approaches 0. For well-modeled sequences, the same-type residues are modulated by their local context, yielding a larger spread and a RED value closer to 1. RED requires no gradient computation and no per-residue masking, reducing the cost of scoring a sequence from L forward passes to one. On a single H100 GPU, this lowers the estimated cost of scoring the 2B BFD sequences and 2.5B MGnify sequences from ≈ 90 GPU-years to ≈ 25 GPU-days. RED is computed using AMPLIFY 350M (*2*) as the scoring model throughout this work. This model is used exclusively for scoring and filtering of pretraining datasets; it is not the subject of any ablation or wet-lab experiment.

#### S4.2. Perplexity across curated control sets

| Set | $n$ | Mean PPL |
|---|---|---|
| Human proteome (UP000005640) | 20,541 | 3.57 |
| DisProt (reviewed IDPs) | 1,742 | 4.21 |
| Shuffled human controls | 692 | 17.88 |
| PE = 5 (all) | 1,738 | 13.04 |
| Annotation level 1 | 852 | 17.00 |
| Annotation level 2 | 599 | 10.97 |
| Annotation level 3 | 220 | 5.61 |
| Annotation level 4 | 44 | 4.70 |
| Annotation level 5 | 23 | 7.14 |

Table S17: Perplexity statistics for curated protein sets scored by AMPLIFY 350M.

##### S4.3. Residue embedding geometry (RED motivation)

RED tracks perplexity closely. Because perplexity requires one masked forward pass per residue, it was only computed on a random subset of 10,000 UniRef100 sequences drawn from the training set. On this subset, RED and pseudo-perplexity are correlated at Spearman $r = -0.97$ ($p < 10^{-3}$). This strong rank correlation motivates the use of RED as a single-forward-pass proxy for the per-residue perplexity computation.

#### S5. RED-based filtering

##### S5.1. RED distributions and KDE boundaries

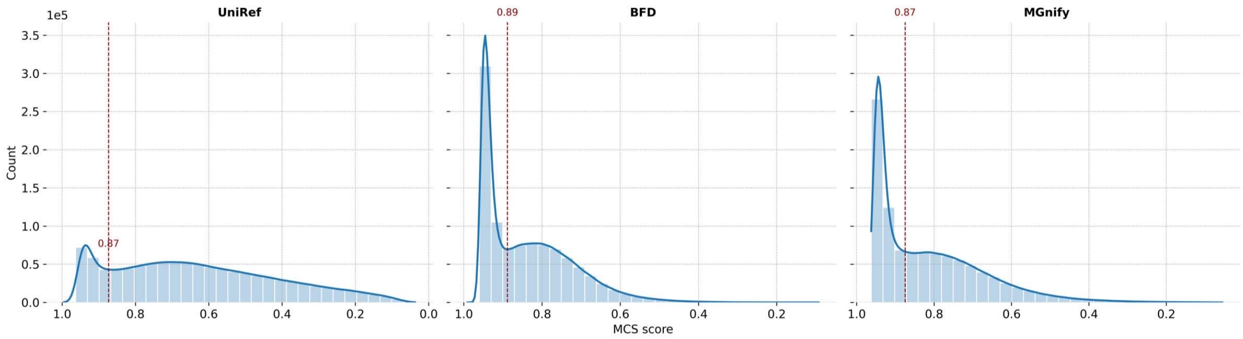

**Figure** S2: Intra-sequence mean pairwise cosine similarity (MCS = 1−RED) and KDE boundaries, all datasets.

The RED score distribution is bimodal, separating a low-quality mode from a high-quality mode. Per dataset, the boundary between the two modes was located from a kernel density estimate (KDE) of the RED scores and used as the filtering threshold. The boundaries are consistent across datasets. RED = 0.112 (BFD), 0.125 (MGnify), and 0.127 (UniRef100) fall at the 44.4th, 45.3th, and 16.7th percentile of each dataset, respectively. The per-dataset boundaries are provided in `kde_boundaries.csv`.

##### S5.2. Effect of filtering on sequence distribution

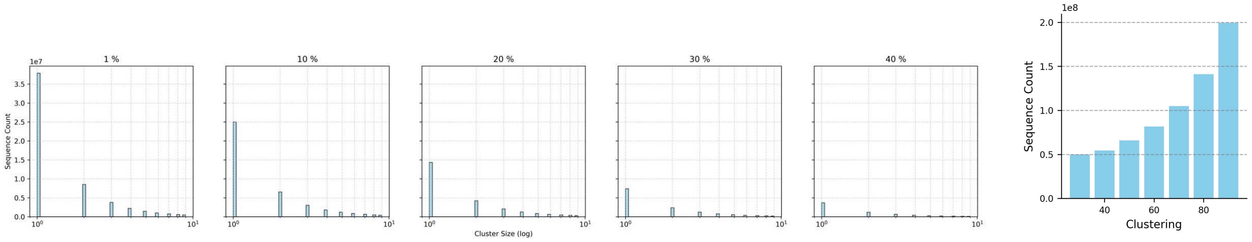

**Figure** S3: Cluster-size distribution after filtering (UniRef100, 30% identity clustering) (left). Sequence count vs. clustering (UniRef100) (right).

#### S6. Downstream evaluation suite

Complete numerical results for all ablation models across all evaluation axes are provided in `in_silico_results.csv` (100 model rows; three identifier fields plus seven evaluation columns). Each row identifies a model by three fields: `model_name` (matches the registry in §S3), `1m` (boolean; `True` if trained for 1M steps, `False` for 100k), and `filtering` (the RED quantile applied, or `0.0` for unfiltered models). The seven evaluation columns are: *KL homologs (mean)*, *Calibration (T)*, *Calibration (NT)*, *Categorical Jacobian (CASP16)*, *Deep mutational scanning*, *Property (FE, ext.)* and *Property (FE, rest.)*. The two *Property* columns are aggregates over the 17 supervised property-prediction tasks listed in Table S18, computed under extended (EXT., 2,880 hyperparameter configurations) and restricted (REST., 108 configurations) grids, respectively. Per-task scores for all models (each task under both grids) are provided in `property_prediction_results.csv` (see §S6.1). All data-composition results discussed in the main text are contained in this file and are not duplicated as separate tables: filtering ablations and metagenomic subsampling are encoded by the `filtering` field and the `model_name` prefixes (e.g. `MILA_BFD_0.50`, subsampling proportions), UniRef100 clustering by the `MILA_U100_clust{30...90}` prefixes, and the 200B-versus-2T-token scaling comparison by the `1m` flag. Each result can thus be reproduced by selecting the corresponding rows.

##### S6.1. Supervised property-prediction tasks

| Task | Type | Approx. Size | Description |
|---|---|---|---|
| antibiotic resistance | multiclass (19) | 3,420 | resistance class; CARD |
| cloning CLF | binary | 28,200 | clonability |
| enzyme catalytic efficiency | regression | 16,800 | $k_{\text{cat}}/K_M$ from sequence |
| fitness prediction | regression | 8,730 | scalar variant fitness |
| fluorescence prediction | regression | 54,000 | GFP-mutant intensity |
| localization prediction | multiclass | 8,460 | subcellular localization |
| material production | binary | 28,100 | functional-material production |
| metal ion binding | binary | 7,330 | metal-ion binding |
| optimal pH | regression | 9,860 | optimal pH |
| optimal temperature | regression | N/A | optimal temperature |
| TCR pMHC affinity | binary | N/A | TCR–pMHC binding |
| stability prediction | regression | N/A | $\Delta\Delta G$ stability |
| solubility prediction | binary | N/A | soluble expression |
| peptide HLA/MHC affinity | binary | 72,770 | peptide–HLA/MHC binding |
| secondary structure (Q3) | multiclass | 10,900 | 3-class per-residue SS |
| secondary structure (Q8) | multiclass | 10,900 | 8-class per-residue SS |
| fold | multiclass | 12,400 | 1,195 folds |

Table S18: Downstream tasks from the xTrimoPGLM benchmark.

##### S6.2. Hyperparameter grids and sensitivity

| Hyperparameter | Values |
|---|---|
| Hidden size | 1024, 2048 |
| Learning rate | $10^{-3}$, $5{\times}10^{-4}$, $10^{-4}$ |
| Epochs | 10, 20, 40 |
| Weight decay | $10^{-4}$, $10^{-3}$ |
| Batch size | 32, 64, 128 |
| Random seeds | 42, 43, 44 |

Table S19: Restricted grid (REST.); 108 configs/task.

| Hyperparameter | Values |
|---|---|
| Hidden size | 512, 1024, 2048, 4096 |
| Learning rate | $5{\times}10^{-3}$, $10^{-3}$, $5{\times}10^{-4}$, $10^{-4}$, $5{\times}10^{-5}$, $10^{-5}$ |
| Epochs | 3, 5, 10, 20, 40, 80 |
| Weight decay | 0, $10^{-4}$, $10^{-3}$, $10^{-2}$ |
| Batch size | 16, 32, 64, 128, 256 |
| Random seeds | 42, 43, 44 |

Table S20: Extended grid (EXT.); 2,880 configs/task.

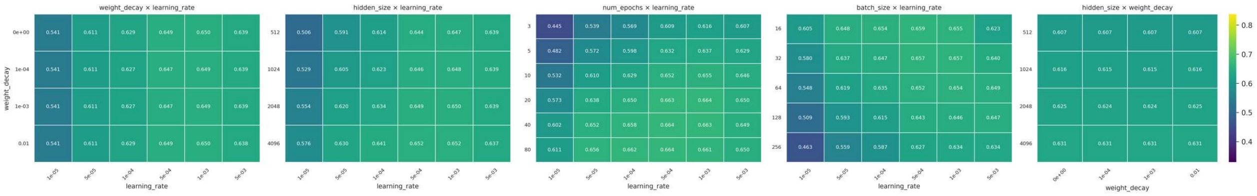

(a) 2D performance landscapes.

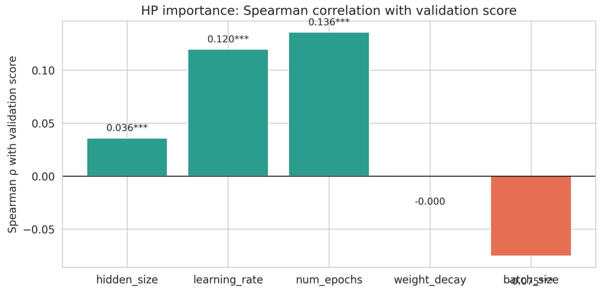

(b) HP–score correlation.

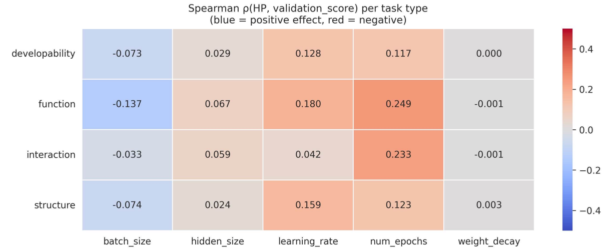

(c) Sensitivity by task type.

**Figure** S4: **Hyperparameter search: landscape and sensitivity.**

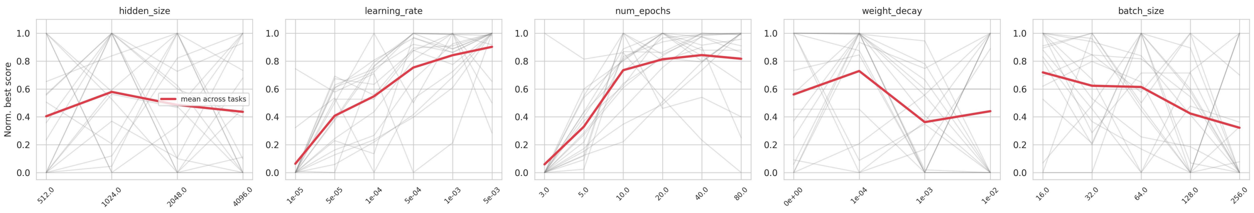

(a) Sensitivity curves.

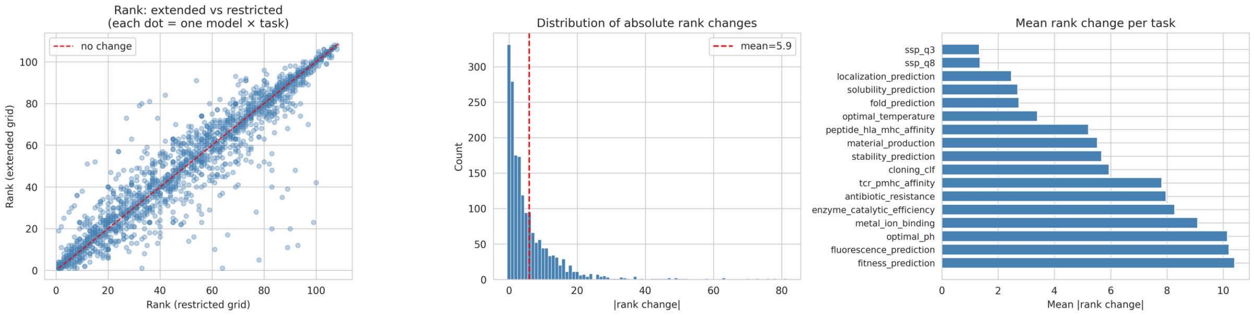

(b) Rank stability across grids (going from the restricted to the extended grid search).

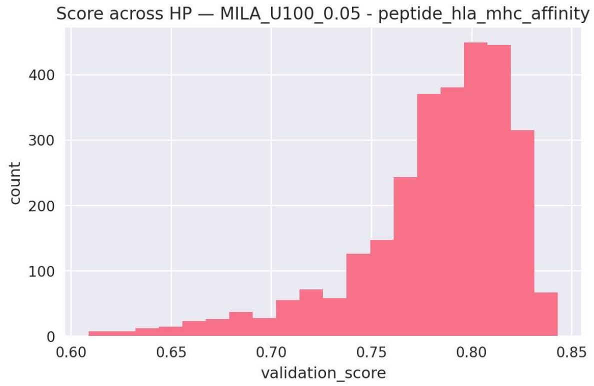

(c) Validation-score distribution for a single model (peptide HLA/MHC).

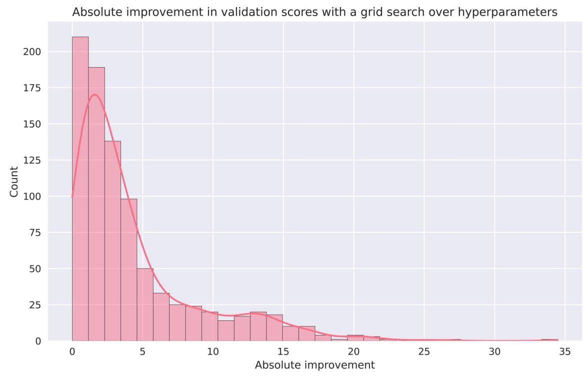

(d) Distribution of val. score improvements over default HPs.

**Figure** S5: **Hyperparameter search: stability and improvement.** These figures demonstrate the necessity of running HP grid searches to fairly compare models.

##### S6.3. Transfer learning (random vs pretrained)

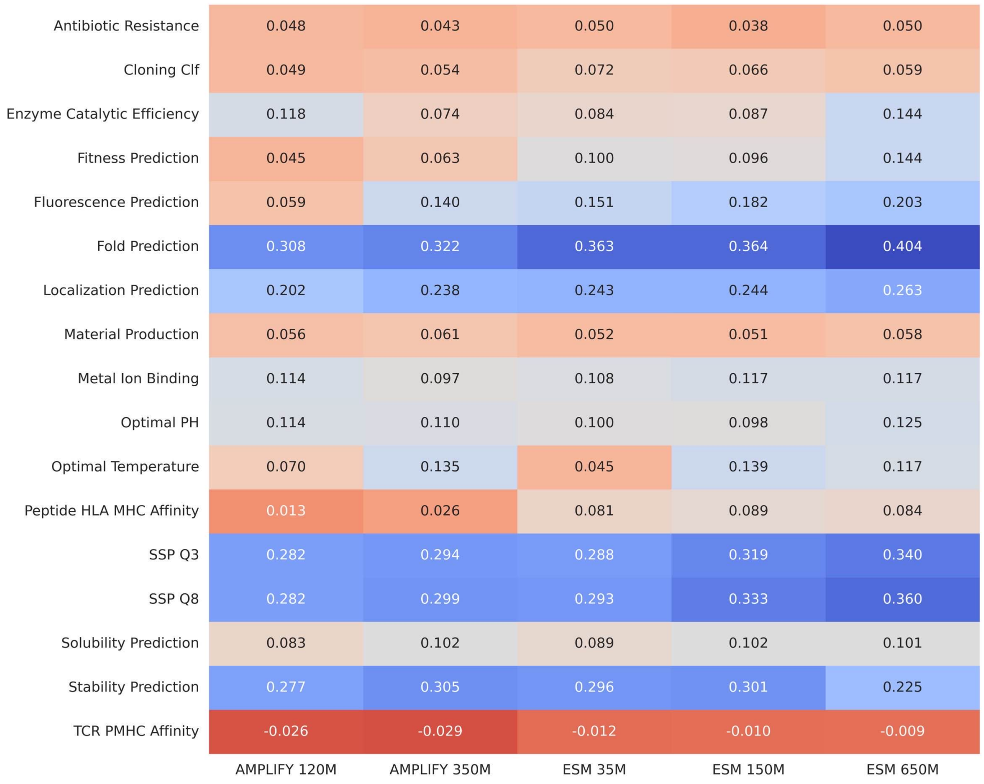

**Figure** S6: **Transfer evaluation across model families and scales.** The differences are computed on the test scores corresponding to the best validation performance, between heads trained on pre-trained models and randomly re-initialized models of the same architecture.

##### S6.4. Deep mutational scanning (DMS)

**Protocol..** We evaluate zero-shot fitness prediction on the substitution split of the ProteinGym benchmark (September 2025 release; 217 assays, 2.4M mutated sequences) (*8*). Each variant is scored by the masked-marginal method: the mutated position is masked and the log-likelihood ratio of the mutant versus wild-type residue is read from the wild-type sequence context. Multi-site mutants are scored by summing the per-position ratios. The reported score is the Spearman correlation between model scores and experimental fitness, aggregated by a weighted average first over UniProtKB sequences and then over functional categories, following the ProteinGym aggregation protocol (*8*). The aggregate per-model DMS Spearman is the *Deep mutational scanning* column of `in_silico_results.csv`.

All ProteinGym wild-type sequences are present in UniProt, a potential data-leakage advantage for UniProt-trained models. Following community practice (*9, 10*), we do not deduplicate against ProteinGym wild-types, though our 90%-identity validation-set deduplication may remove evolutionary context around these targets and suppress reported scores. Moreover, ProteinGym fitness spans heterogeneous biophysical properties (enzymatic activity, binding affinity, thermostability), not all of which are expected to track evolutionary sequence likelihood in a zero-shot setting.

##### S6.5. Unsupervised structure prediction (categorical Jacobian, CASP16)

As a zero-shot probe of the structural information encoded in pLM representations, we compute the categorical Jacobian (*11*). The Jacobian measures how a single substitution at one position perturbs the model's predicted distribution at every other position. Coupled positions are those whose predictions are most sensitive to each other's identity, and these couplings recover residue–residue contacts without any supervised training or structural labels.

**Jacobian construction..** Let $\sigma = (x_1, \ldots, x_L)$ be a sequence of length $L$ and let $\mathcal{A}$ be the alphabet of the 20 standard amino acids. For a masked-language-model head producing logits over the vocabulary, write $\log P(x_i{=}a \mid \sigma)$ for the log-probability of amino acid $a$ at position $i$ given input $\sigma$, restricted to $\mathcal{A}$. The categorical Jacobian entry is the change in this log-probability when position $j$ is set to amino acid $b$:

$$J[i, a, j, b] \;=\; \log P\Big(x_i{=}a \mid \sigma^{j\to b}\Big) \;-\; \log P(x_i{=}a \mid \sigma)\,, \tag{S3}$$

where $\sigma^{j\to b}$ is $\sigma$ with position $j$ substituted by $b$. We compute it as follows:

1. **Reference pass.** Run a single forward pass on the wild-type sequence and store the reference log-probabilities $\log P(x_i{=}a \mid \sigma)$ for all $i \in \{1, \ldots, L\}$, $a \in \mathcal{A}$. Tokenizer special tokens (e.g. a prepended `BOS`/`CLS`) are detected and offset so that sequence position $i$ maps to the correct token index.
2. **Mutant passes.** For each of the $L \times 20$ single substitutions $(j, b)$, build the mutated sequence $\sigma^{j\to b}$ and run a forward pass (batched), recording $\log P(x_i{=}a \mid \sigma^{j\to b})$ for all $i, a$.
3. Assemble $J \in \mathbb{R}^{L\times20\times L\times20}$ via Equation S3.

**Contact extraction..** The full Jacobian is reduced to an $L \times L$ contact-score map by the centering/APC pipeline used in coevolution analysis:

1. **Four-pass centering.** Sequentially subtract the mean along each of the four axes of $J$ (positions $i$, amino acids $a$, positions $j$, amino acids $b$, in that order), removing per-axis offsets.
2. **Symmetrization.** Replace $J$ by $\frac{1}{2}(J + J^\top)$, where $\top$ exchanges the index pairs $(i, a) \leftrightarrow (j, b)$.
3. **Frobenius norm.** Collapse the two amino-acid axes into a single coupling strength per position pair, $C_{ij} = \sqrt{\sum_{a,b} J[i, a, j, b]^2}$, and set the diagonal to zero.
4. **Average product correction (APC).** Apply $C_{ij} \leftarrow C_{ij} - \dfrac{C_{i\cdot}\, C_{\cdot j}}{C_{\cdot\cdot}}$, where $C_{i\cdot}$, $C_{\cdot j}$ are row/column sums and $C_{\cdot\cdot}$ is the total sum; set the diagonal to zero again. The resulting $C$ is the predicted contact-score map.

**Evaluation protocol and metrics..** We evaluate on CASP16 targets (*12*), excluding sequences longer than 1,024 residues. Following the ESM contact-evaluation protocol (*13*), metrics are restricted to long-range pairs with sequence separation $|i - j| \geq 6$, computed over the upper triangle; pairs with missing ground-truth labels are masked out. Let the predicted scores for valid pairs be ranked in decreasing order. We report:

- **P@L, P@L/2, P@L/5** — precision among the top $L$, $L/2$, and $L/5$ ranked pairs;
- **AUC** — the mean of the binned precisions evaluated at the top $0.1L, 0.2L, \ldots, 1.0L$ pairs;
- **ROC-AUC** — the area under the ROC curve over all valid long-range pairs.

AUC (mean binned precision) is the primary metric reported as **Categorical Jacobian (CASP16)** throughout (and notably in `in_silico_results.csv`). The remaining metrics are computed per sequence and averaged across targets. All five metrics (P@L, P@L/2, P@L/5, AUC, and ROC-AUC, each as the mean and standard deviation over the CASP16 targets) are provided for every model in `categorical_jacobian_results.csv`.

### S7. Calibration task and mutation stability

#### S7.1. Definition, consensus computation, and the T/NT split

The calibration task measures whether a model has learned the *evolutionary consensus* at conserved positions, or has instead memorized the specific proteome sequences seen during training. It is a single-forward-pass, zero-shot diagnostic that requires no labels beyond a multiple-sequence alignment per query protein.

**Alignments and homolog profiles..** We use the human proteome from UniProtKB (Proteome ID UP000005640; `human_proteome.csv`, 205,155 sequences) as the query set, and align each protein to the BFD database with

HHblits (*14*), requiring a minimum sequence identity and coverage of 50% and a maximum pairwise sequence identity of 99%. Proteins with fewer than 10 homologs are discarded. Aligning the full proteome against BFD is the only computationally heavy step and is performed once as preprocessing. From each resulting `a3m` alignment we build a per-position amino-acid frequency profile: insertion columns (lowercase in `a3m`) are stripped, and at each query (non-gap) column we count, over the query and all homologs, only the 20 standard residues (gaps and non-standard symbols excluded), normalizing the counts to a distribution $p_{\text{homologs}}(\cdot)$ over $\mathcal{A}$. The aggregated profiles are released as `human_consensus_positions.csv` (columns `sequence_name`, `position`, `wt_aa`, and the 20 per-residue frequencies).

**Consensus positions and conservation thresholds..** A position is a *consensus* (conserved) position at threshold $\tau$ if some amino acid attains frequency $\geq \tau$ in its homolog profile. The consensus residue is $a^\star = \arg\max_{a\in\mathcal{A}} p_{\text{homologs}}(a)$. We report results at conservation thresholds $\tau \in \{0.70, 0.90\,0.96\}$ for the calibration score (figs. S8 and S9) and use $\tau = 0.70$ to define the consensus mutations reported in the calibration task and explored with Rosetta (§S7.4).

**Target (T) and non-target (NT) split..** Consensus positions are partitioned by comparing the human (wild-type) residue $x_i$ to the consensus residue $a^\star$:

**Target (T): $x_i = a^\star$.** The human residue coincides with the evolutionary consensus. A model that has learned the consensus *or* memorized the human sequence predicts the same residue, so any well-trained model is expected to score high.

**Non-target (NT): $x_i \neq a^\star$.** The human residue departs from the consensus. The evolutionary landscape converges to the consensus residue $a^\star$, but a model that has memorized the human proteome will instead favor the human variant $x_i$. Accuracy at NT positions therefore isolates memorization from generalization: a model that improves at T positions while stagnating or degrading at NT positions is progressively overfitting to its training proteomes.

**Scoring procedure..** For each consensus position we mask the query residue and run a single forward pass. Sequences longer than 512 residues are truncated to a 512-residue window centered on the masked position. The output logits are restricted to the 20 standard amino acids and renormalized by a softmax to give the model distribution $p_{\text{model}}(\cdot)$ over $\mathcal{A}$; the top-1 prediction is $\hat{a} = \arg\max_a p_{\text{model}}(a)$. Two complementary quantities are recorded per position.

First, a *distributional* agreement between the model and the homolog profile, via the Kullback–Leibler divergence:

$$D_{\text{KL}}\big(\mathbf{p}_{\text{homologs}} \,\big\|\, \mathbf{p}_{\text{model}}\big) \;=\; \sum_{a\in\mathcal{A}} p_{\text{homologs}}(a)\,\log\frac{p_{\text{homologs}}(a)}{p_{\text{model}}(a)}, \tag{S4}$$

with both distributions clipped to $[\varepsilon, 1]$ ($\varepsilon = 10^{-10}$) for numerical stability. The per-class **calibration score** is obtained by averaging over positions within a class (T or NT) and mapping to $[0, 1]$,

$$\text{Calibration} \;=\; \exp\Big(-\overline{D_{\text{KL}}}\Big), \tag{S5}$$

reported as a percentage (higher is better). Per-model T and NT calibration scores for all 100 registry models are in `in_silico_results.csv` (columns `Calibration (T)` and `Calibration (NT)`; the column `KL homologs (mean)` reports $\overline{D_{\text{KL}}}$ pooled over all consensus positions).

Second, a *top-1* breakdown of the predicted residue $\hat{a}$ against the wild-type $x_i$ and consensus $a^\star$: at T positions we record whether $\hat{a} = x_i\ (= a^\star)$; at NT positions we distinguish the three outcomes $\hat{a} = x_i$ (human variant, the memorization signature), $\hat{a} = a^\star$ (consensus, the generalization signature), and $\hat{a} \notin \{x_i, a^\star\}$ (other). These prediction-type frequencies underlie the figures in §S7.5.

#### S7.2. Calibration by filtering level and dataset

Figure S8 reports *sequence recovery* as a function of the RED filtering quantile, separately for target (T) and non-target (NT) positions. Sequence recovery is the top-1 recovery rate: the fraction of masked conserved positions at which the model's most likely amino acid ($\hat{a} = \arg\max_a p_{\text{model}}(a)$, §S7.1) equals the reference residue. At T positions the reference is the wild-type residue (which coincides with the consensus); at NT positions, recovery of the human wild-type versus the consensus is the memorization-versus-generalization signal dissected in §S7.5. This top-1 recovery rate is an alternative, more interpretable view used only in this figure (and Figure S9); every other calibration result in this work uses the calibration score of (S5).

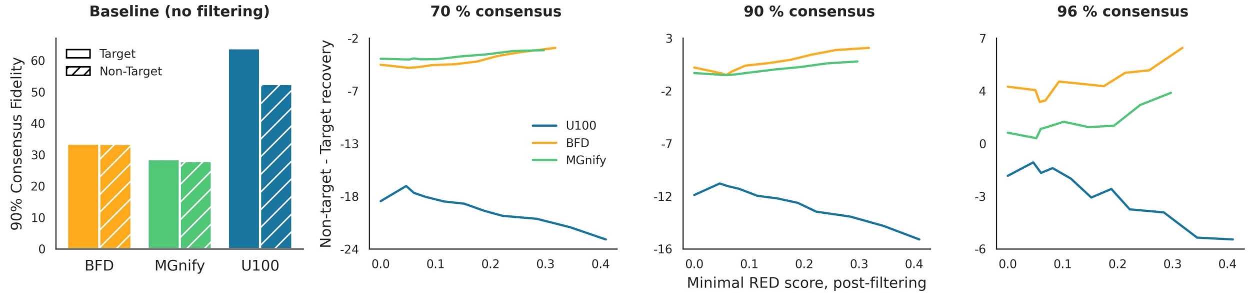

**Figure** S7: Absolute calibration scores (90%; left) and absolute calibration differences between NT and T vs filtering (at different thresholds; right). Increased filtering improves memorization for models trained on UniRef100 but reduces it for models trained on metagenomic datasets.

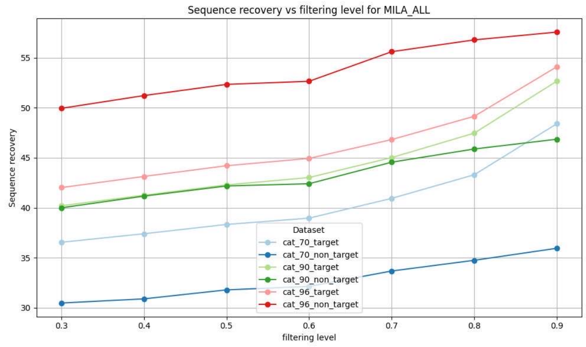

(a) All datasets.

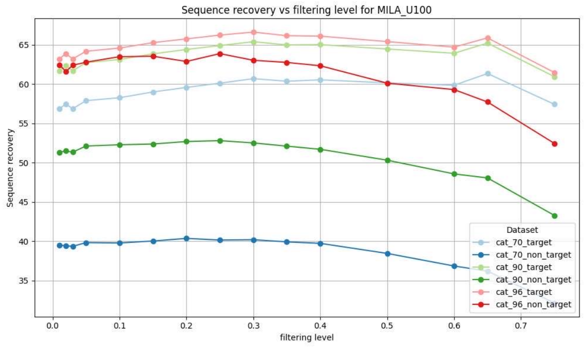

(b) UniRef100.

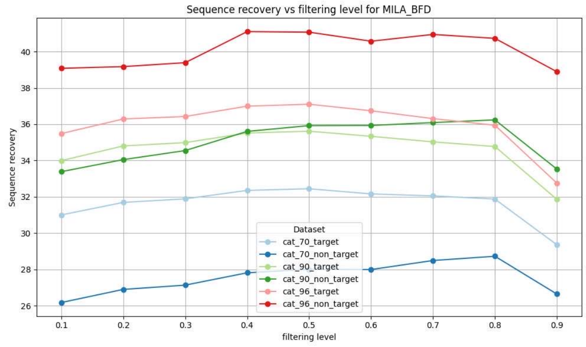

(c) BFD.

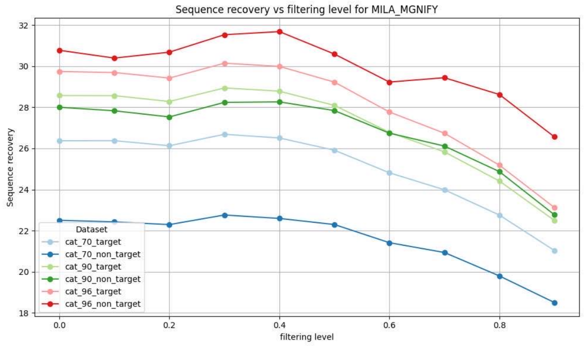

(d) MGnify.

**Figure** S8: Consensus (top1 accuracy) by filtering level at T and NT positions, for different consensus thresholds.

#### S7.3. Calibration vs training step (memorization signature)

Extending the training budget from 100k to 1M steps ($\approx$ 200B to 2T tokens) exposes a dataset-dependent signature of memorization that the T/NT split is designed to isolate. At T positions, where the wild-type residue coincides with the consensus, calibration improves with longer training across all datasets. This is consistent

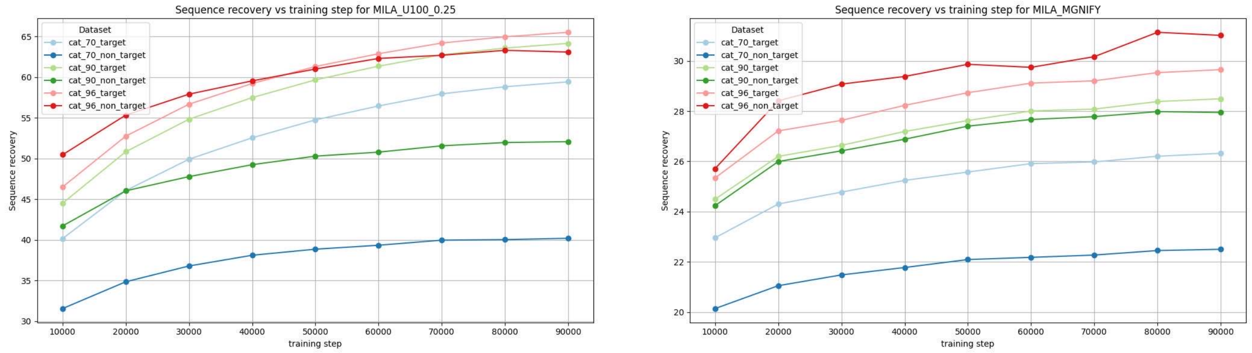

(a) UniRef100 (0.25 filtering quantile). (b) MGnify (no filtering).

**Figure** S9: **Consensus recovery (top1 accuracy) vs training step** at T and NT positions.

with a better overall fit to the natural sequence distribution. The diagnostic behavior is at NT positions, where models pretrained on UniRef100 *degrade* with longer training (mean drop of 14.1 points across filtering levels), indicating that additional steps increase memorization of training variants $x_i$ over the more stable consensus $a^\star$. Models pretrained on diverse metagenomic data show no such degradation: BFD and MGnify models instead *improve* at NT positions with longer training (+11.2 and +3.1 points on average, respectively). Per-model T and NT calibration scores at both step counts are available in `in_silico_results.csv`: the 100k-step and 1M-step runs are distinguished by the boolean `1m` column, the dataset and filtering quantile by the `model_name` and `filtering` fields, so the full dataset × filtering × step breakdown behind figs. S8 and S9 can be reconstructed directly.

#### S7.4. Rosetta in silico mutagenesis ($\Delta\Delta G$)

This assay tests whether the consensus residues identified by the calibration task (§S7.1) are thermodynamically stabilizing under the Rosetta cartesian ddG protocol.

**Position selection..** We selected human-proteome positions where the homolog consensus residue differs from the human residue at a consensus frequency ≥ 70%. Experimental structures were obtained by mapping UniProtKB accessions to PDB entries through the SIFTS database (*15, 16*). When several chains covered a site, we kept the one with the best crystallographic resolution and highest coverage, restricting to X-ray structures (NMR models excluded). The consensus substitution (human→consensus) at each PDB-covered position was carried through the $\Delta\Delta G$ computation after relaxation, for 983 consensus substitutions across 917 proteins. As a control, 4,682 random non-consensus substitutions at the same positions were scored identically.

**$\Delta\Delta G$ computation..**

1. Extract the selected PDB chain and strip non-protein atoms.
2. Apply Cartesian-space energy minimization with the Rosetta FastRelax protocol (*17*) under the `ref2015_cart` scoring function to resolve clashes in the deposited coordinates before energy differences are computed.
3. Estimate $\Delta\Delta G$ with the Rosetta `cartesian_ddg` application (Rosetta v2025.51 (*18*)), running three independent iterations per mutation with a backbone neighbor shell of one residue.
4. Define $\Delta\Delta G = E(\text{consensus}) - E(\text{human})$ in Rosetta energy units. $\Delta\Delta G < 0$ indicates the consensus residue is more stable on the human backbone.

This evaluates stability on a fixed backbone and does not capture conformational rearrangement; Cartesian $\Delta\Delta G$ with `ref2015_cart` nonetheless correlates well with measured single-point stability changes (*19*).

**Result..** Consensus substitutions are markedly enriched for stabilizing changes ($\Delta\Delta G < 0$) relative to random substitutions (Figure S10): 35.1% of the 983 consensus substitutions are stabilizing, versus 18.9% of the 4,682 random substitutions, and the consensus set has a lower mean $\Delta\Delta G$ (1.21 versus 3.69 Rosetta energy units).

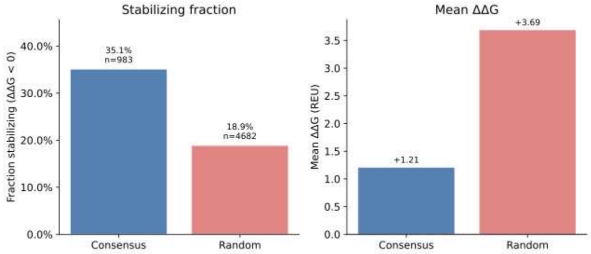

(a) Stabilizing fraction: consensus vs random.

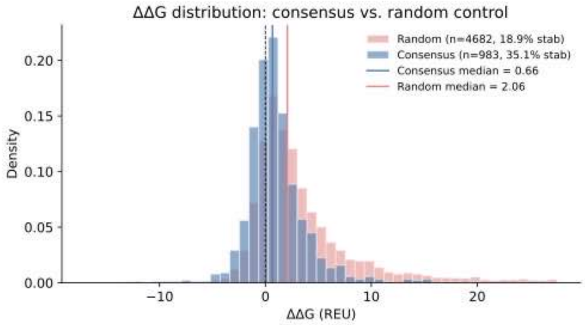

(b) $\Delta\Delta G$ distribution.

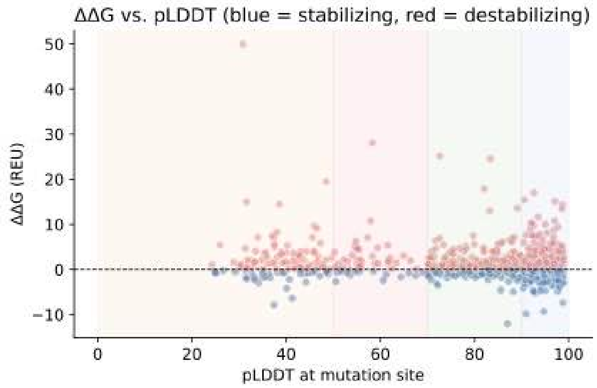

(c) $\Delta\Delta G$ vs pLDDT.

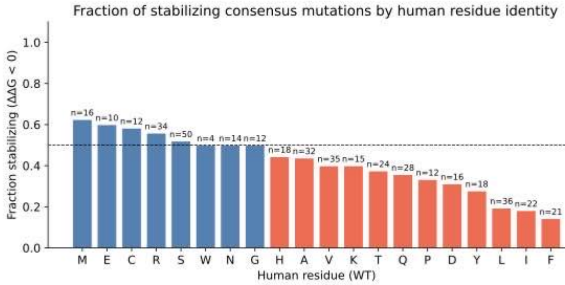

(d) Stabilizing fraction by AA.

**Figure** S10: **Rosetta in silico mutagenesis validation.**

Per-mutation $\Delta\Delta G$ values are provided in `ddg_results.csv` (consensus) and `ddg_results_random.csv` (random control).

#### S7.5. Prediction-error structure (KL analyses)

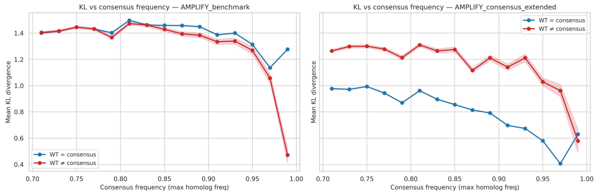

(a) KL vs consensus frequency.

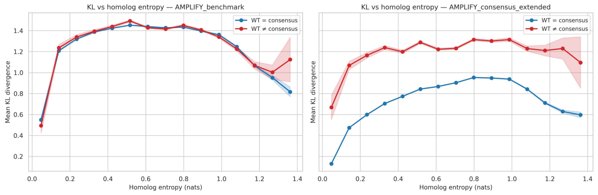

(b) KL vs positional entropy.

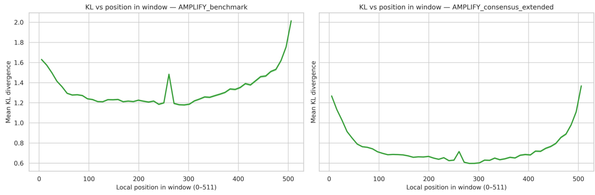

(c) KL vs local position.

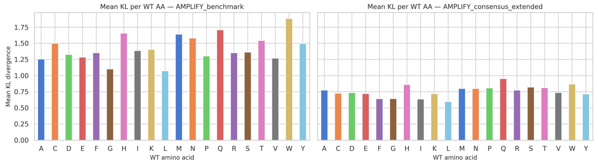

(d) KL vs wild-type AA.

**Figure S11: Prediction error (KL) vs consensus frequency, homolog entropy, position and AA type.**

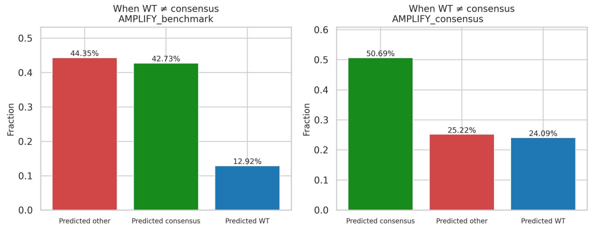

(a) Prediction types when WT ≠ consensus.

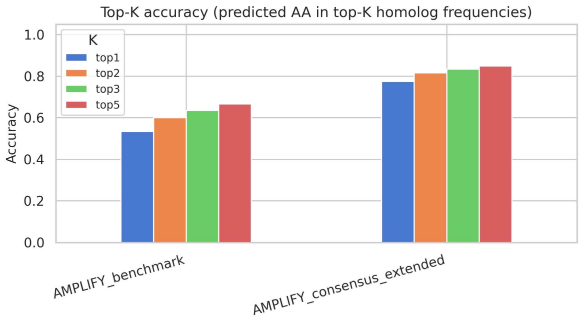

(b) Top-$k$ accuracy by consensus status.

**Figure S12: Calibration prediction-type breakdown, NT positions.**

### S8. Enzyme discovery: candidate selection

#### S8.1. Target filtering pipeline (EC 1.14.11)

The curated candidate pool was drawn from all UniProtKB entries annotated to the EC 1.14.11 enzyme class (Fe/$\alpha$KG-dependent dioxygenases). Two successive filter stages were applied, designed to retain sequences a practitioner would consider plausible members of the family.

##### Stage 1 — hard filters..

(1) **Length**: sequences shorter than 200 or longer than 450 residues were removed.
(2) **Ambiguous residues**: sequences containing any of `X, O, U, B, J, Z` were removed.
(3) **Catalytic motif**: sequences lacking the *HXD/E* motif (a histidine, any residue, then an aspartate or glutamate) were removed. This motif chelates the catalytic iron and is a defining feature of the enzyme class.
(4) **Transmembrane**: predicted transmembrane proteins (DeepTMHMM) were excluded, as they are incompatible with the soluble-expression validation assay.

##### Stage 2 — distribution-based outlier removal..

(5) **Structural confidence**: the bottom 5% of sequences by mean AlphaFold2 pLDDT (*20*) were removed.

(6) **Compositional typicality**: the top 5% of sequences by Kullback–Leibler divergence between their amino-acid composition and the expected composition of King and Jukes (*21*) were removed, as an atypical composition signals deviation from natural evolutionary constraints.

Together, these filters reduced the pool from 95,165 to 34,539 sequences (most of the reduction coming from length constraints), which served as the common starting set for the curated plates (A, B, C).

##### S8.2. Maximal Marginal Relevance (MMR)

All plates were assembled with the same greedy Maximal Marginal Relevance (MMR) procedure, which balances candidate *quality* against *diversity* relative to the already-selected set $S$. At each step it adds the candidate $c$ maximizing

$$\mathrm{MMR}(c) \;=\; \lambda\,\mathrm{Quality}(c) \;-\; (1-\lambda)\,\max_{s\in S}\mathrm{sim}(c,s), \tag{S6}$$

where quality is the min–max-normalized AMPLIFY-C perplexity,

$$\mathrm{Quality}(c) = \frac{\mathrm{ppl}_{\max} - \mathrm{ppl}(c)}{\mathrm{ppl}_{\max} - \mathrm{ppl}_{\min}}, \tag{S7}$$

so that lower perplexity yields higher quality, and $\mathrm{sim}(c,s)$ is the diversity term (cosine similarity between AMPLIFY-C embeddings, or, for the control plate, a BLAST identity). The hyperparameter $\lambda \in [0,1]$ trades quality ($\lambda \to 1$) against diversity ($\lambda \to 0$).

**Algorithm..** Given quality scores, a pairwise similarity matrix, a pairwise identity matrix, a target count $N$, weight $\lambda$, and an identity threshold $\tau$ (optionally seeded with the selections from earlier plates):

1. Initialize $S$ with the highest-quality candidate (unless a preselected set is supplied).
2. For each remaining candidate, compute its maximum similarity to $S$ and its MMR score (Equation S6); rank candidates by MMR descending.
3. Walk down the ranked list and add the first candidate whose pairwise identity to *every* member of $S$ is $\leq \tau$ (the hard diversity constraint).
4. If no candidate satisfies the constraint, relax $\tau$ by 5 percentage points and retry.
5. Repeat steps 2–4 until $|S| = N$.

The identity threshold was $\tau = 50\%$ for all plates, enforced both within a plate and against candidates already chosen for earlier plates.

**Per-plate settings..** The three curated plates draw from the same 34,539-sequence pool (§S8.1); each plate holds 96 candidates.

- **Plate A (quality-focused)**: $\lambda = 0.9$; embedding-cosine diversity; $\leq 50\%$ identity within the plate.
- **Plate B (diversity-focused)**: $\lambda = 0.1$; embedding-cosine diversity; $\leq 50\%$ identity to Plate B and to Plate A.
- **Plate C (pLM-agnostic control)**: $\lambda = 0$ (quality term inactive); diversity computed from BLAST identities (*22*) rather than embeddings, so no pLM score informs the selection; $\leq 50\%$ identity within the plate.
- **Plate D (metagenomic)**: $\lambda = 0.9$ applied to the retrieved metagenomic pool (§S8.3); $\leq 50\%$ identity within the plate.

##### S8.3. kNN retrieval from metagenomic space (Plate D)

Plate D candidates were retrieved directly from the metagenomic databases (BFD and MGnify) by embedding similarity to the curated EC 1.14.11 targets, with no homology or annotation input.

**Prefiltering..** The combined databases ($\approx$4.5 billion sequences) were reduced by (i) removing the bottom 50% of sequences by RED score (reusing the precomputed AMPLIFY-350M scores rather than rescoring with AMPLIFY-C, given the cost of scoring billions of sequences) and (ii) applying the same length (200–450) and *HXD/E* motif filters as the curated pipeline. This left $\approx$500 million sequences.

**Embedding and nearest-neighbor search..** The surviving sequences were embedded with AMPLIFY-C (mean-pooled, 960-d). Using the curated UniProtKB targets from Plates A and B as queries, we performed an exact $k$-nearest-neighbor search under cosine similarity: query and database embeddings were $L_2$-normalized and scored by inner product, streamed over the sharded embedding files with a running top-$k$ merge. We retrieved the $k = 1{,}000$ closest metagenomic sequences per target. The arbitrary cap of 1,000 keeps only the closest neighbors (a larger candidate set could be obtained by increasing $k$).

**Deduplication..** From the pooled neighbors we removed (i) any sequence sharing more than 80% identity with any UniProtKB target and (ii) fragment sequences not beginning with methionine, yielding 44,005 unique metagenomic candidates. Final selection applied the MMR procedure (§S8.2) with $\lambda = 0.9$ and the $\leq 50\%$ pairwise-identity constraint, producing the 96 Plate D candidates.

##### S8.4. Set similarities

**Within and between-plate similarity..** The MMR hard constraint (§S8.2) guarantees that no two selected sequences within a plate exceed 50% pairwise identity, and that Plates B and D additionally satisfy this bound against sequences selected for earlier plates. The full AMPLIFY-C embedding cosine-similarity matrix across all 384 selected candidates is provided in `pairwise_cosine_similarity.parquet`.

**Plate D versus EC 1.14.11 (zero homology)..** To test whether the retrieved metagenomic candidates share detectable homology with the annotated enzyme class, we computed, for every Plate D sequence, its maximum percent identity to any EC 1.14.11 member with MMseqs2 (*23*). Searches used maximum sensitivity (`-s 7.0`), a target coverage of 0.8 (`-cov-mode 0`), and reported percent identity over aligned pairs. None of the 96 Plate D candidates produced a detectable MMseqs2 hit against EC 1.14.11, i.e. a maximum identity of 0 across the plate, confirming that homology search would not have surfaced any of them.

**Characterized hydroxylases..** For context, a set of experimentally characterized Fe/$\alpha$KG hydroxylases from the literature were embedded with AMPLIFY-C. These reference enzymes are shown as overlays in the embedding projections (main-text Fig. 3E and Fig. 4B) and available in `literature_hydroxylases.csv`.

##### S8.5. Per-sequence expression data

Per-candidate in silico features for all four plates (384 sequences) are provided in `all_plates_features.parquet`: `id`, `plate`, `length`, AMPLIFY-C RED and pseudo-perplexity, the 22 biophysical descriptors, and the structural descriptors (mean pLDDT, mean SASA, ESM-IF NLL, soluble-ProteinMPNN log-likelihood).

##### S8.6. Additional property distributions of selected candidates

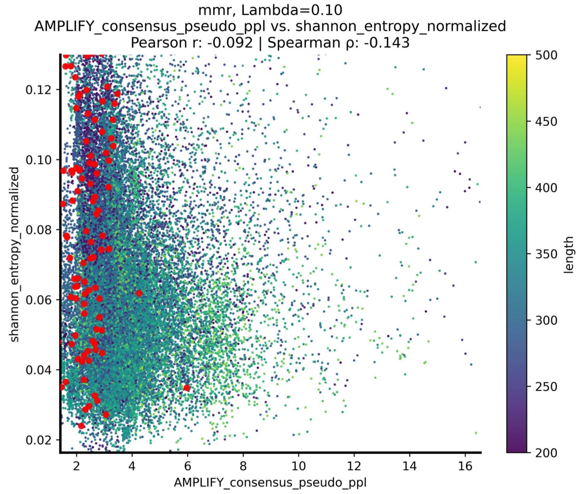

(a) PPL vs entropy ($\lambda$=0.10).

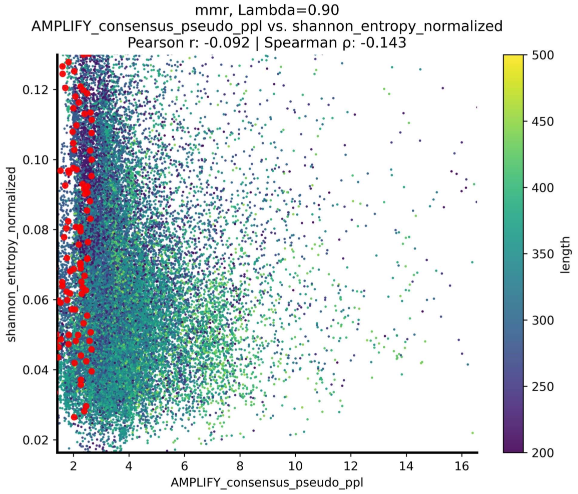

(b) PPL vs entropy ($\lambda$=0.90).

**Figure S13: PPL vs sequence entropy under different MMR diversity weights.**

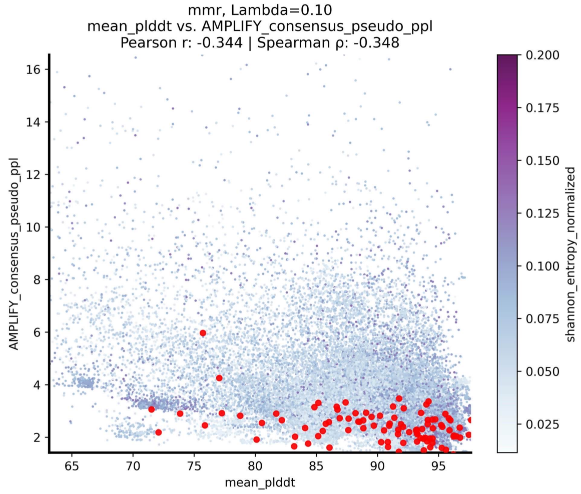

(a) Mean pLDDT ($\lambda$=0.10).

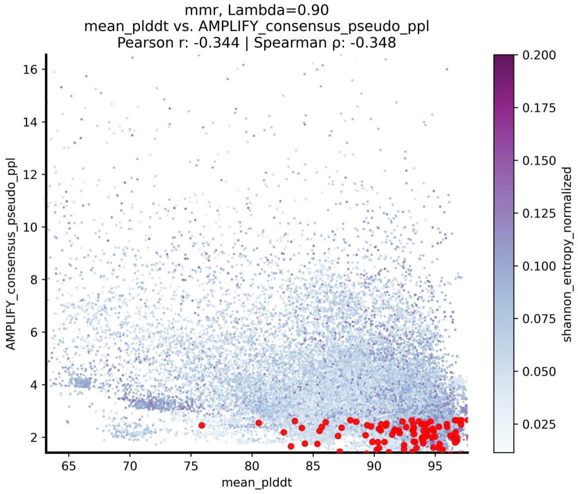

(b) Mean pLDDT ($\lambda$=0.90).

**Figure S14: Mean pLDDT of selected candidates by MMR diversity weight.**

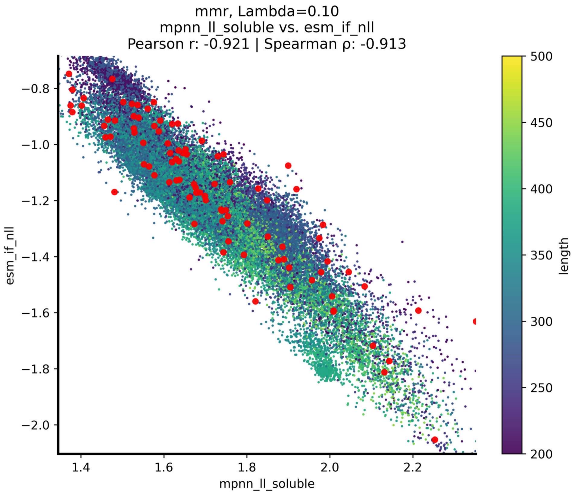

(a) MPNN / ESM-IF ($\lambda$=0.10).

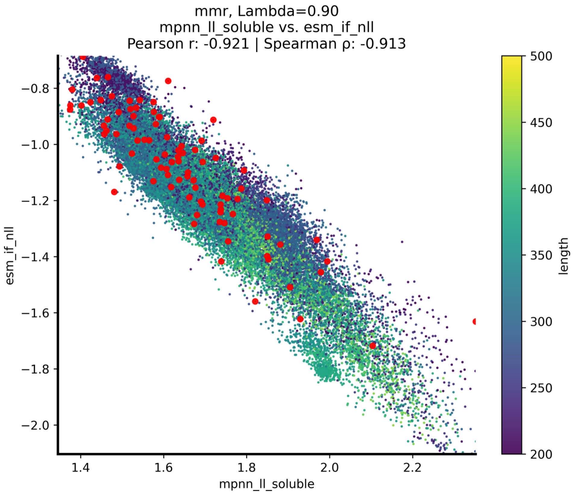

(b) MPNN / ESM-IF ($\lambda$=0.90).

**Figure** S15: **ProteinMPNN and ESM-IF scores by MMR diversity weight.**

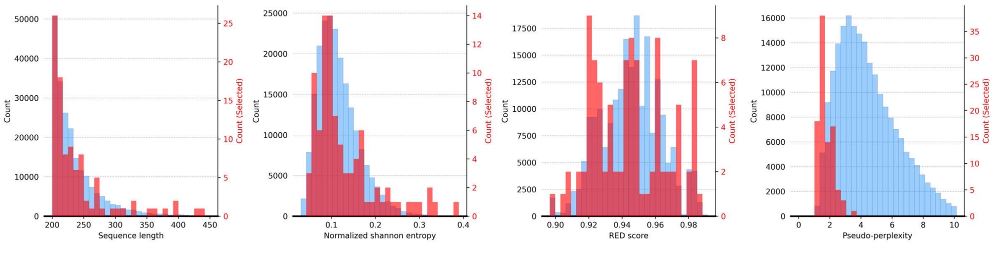

**Figure** S16: Biophysical properties of Plate D selections vs full library. In blue is the distribution of the full retrieved pool (the 44,005 sequences), and in red the distribution of the selected metagenomic candidates (plate D).

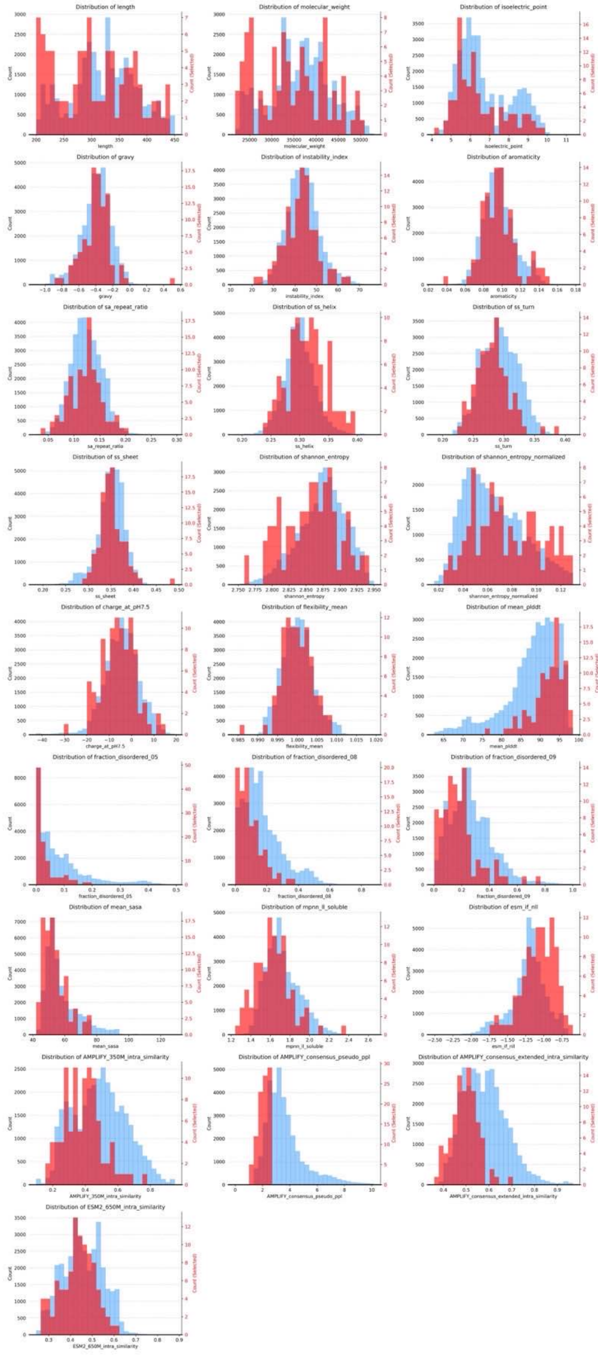

(a) Plate A properties ($\lambda$=0.90).

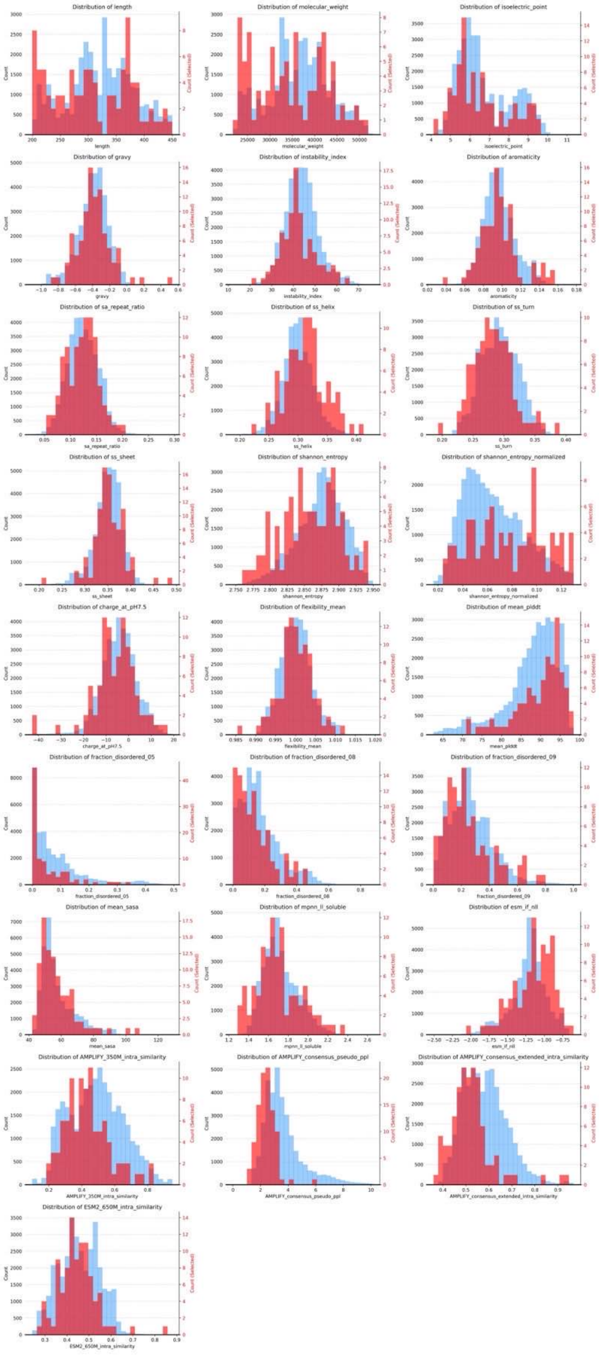

(b) Plate B properties ($\lambda$=0.10).

**Figure** S17: Biophysical property distributions of selected candidates by MMR diversity weight (plate A vs. B). In blue is the distribution of the full filtered pool (the 34,539 sequences surviving the initial target filters; §S8.1), and in red the distribution of the selected candidates for the quality-focused ($\lambda$ = 0.90) and diversity-focused ($\lambda$ = 0.10) plates.

### S9. Enzyme discovery: wet-lab protocols and data

#### S9.1. Recombinant expression, purification, and SDS-PAGE

**Recombinant expression in *E. coli*..** The genes encoding the selected sequences were codon-optimized for recombinant expression in *Escherichia coli* and synthesized as inserts in the `pET28a(+)` vector between the BamHI and HindIII restriction sites, giving an N-terminal polyhistidine tag. Chemically competent BL21(DE3) cells were transformed with each plasmid and grown in 96-well plates in 800 $\mu$L Luria–Bertani broth with 50 $\mu$g/mL kanamycin (overnight, 37 °C, 80% humidity, 1000 rpm). A 30 $\mu$L aliquot was used to inoculate 400 $\mu$L Terrific Broth with 50 $\mu$g/mL kanamycin; at $OD_{600}$ = 0.8–1.5, expression was induced with 0.5 mM IPTG and cultures were incubated overnight (18 °C, 80% humidity, 1000 rpm). After 20 h, plates were centrifuged (3500 rpm, 5 min), the supernatant discarded, and the cell pellets stored at −80 °C.

**Protein preparation..** Thawed pellets were resuspended in lysis buffer (100 mM potassium phosphate pH 7.5, 2 mg/mL lysozyme, 10 U/mL benzonase, 0.1 mM phenylmethylsulfonyl fluoride) and incubated (22 °C, 1000 rpm, 2 h), then centrifuged (3500 rpm, 5 min). A 20 $\mu$L portion of the cleared lysate was reserved for SDS-PAGE. His-tagged proteins were captured by adding 50 $\mu$L equilibrated Ni-NTA resin (30 min, shaking), transferred to 96-well filter plates, washed, and eluted with buffer containing 250 mM imidazole; the semi-purified eluates were analyzed by SDS-PAGE.

**SDS-PAGE and band quantification..** Lysate and semi-purified samples were diluted 1:1 in LDS dye with reducing agent (10 mM DTT), heated (80 °C, 10 min), and 10 $\mu$L was loaded onto a 4–12% Bis-Tris MOPS gel and run for 45 min. Gels were stained with Imperial Stain (1 h) and destained in water (2 days). Recombinant expression was quantified from the band volume at the expected molecular weight in the lysate gel; the presence of a band after Ni-NTA enrichment validated successful expression of the His-tagged target. The per-plate *normalized* band volume used in the main text (Fig. 3B) is obtained by min–max normalization of the band volumes.

#### S9.2. Per-sequence expression data

Complete expression results are available in `in_vivo_results.csv`, including per-sequence `kDa`, `mg/mL`, `lysate band`, and `purified band` volumes. This file also includes per-sequence `gene index`, `gene name`, `insertion point name`, `vector name`, and `insert sequence`. The plate-level expression summaries and the perplexity–expression correlations are reported in the main text.

#### S9.3. SDS-PAGE gels

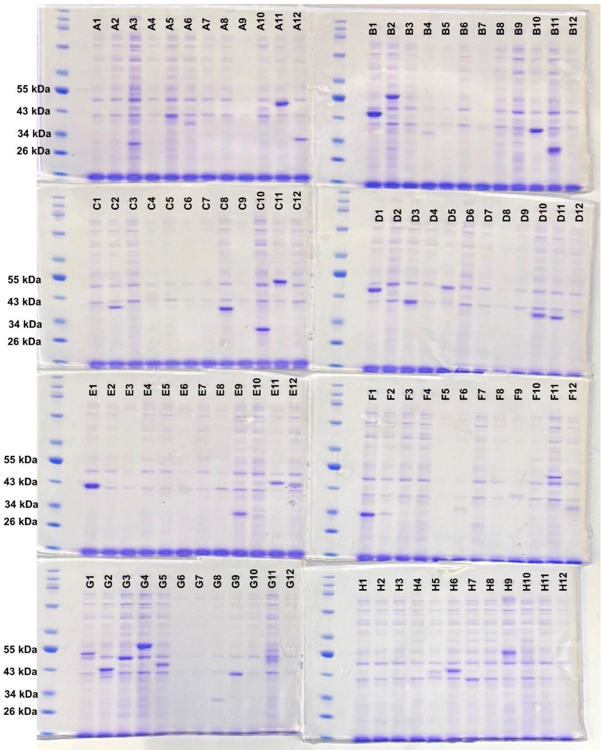

(a) Plate A.

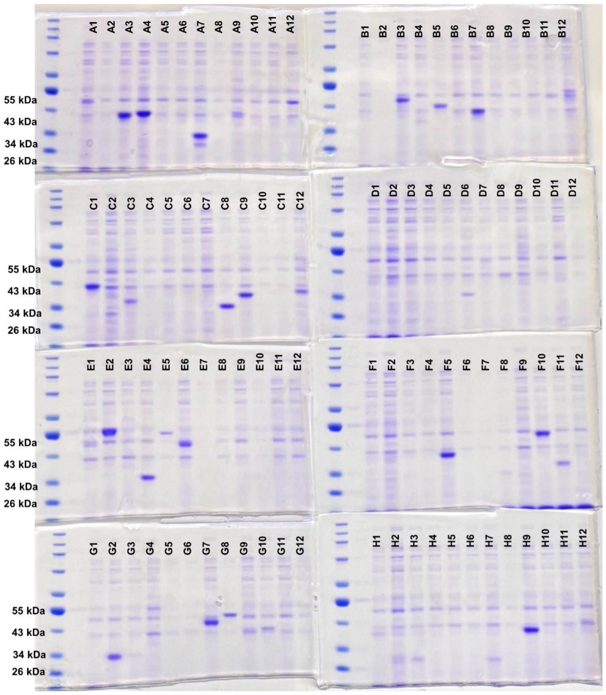

(b) Plate B.

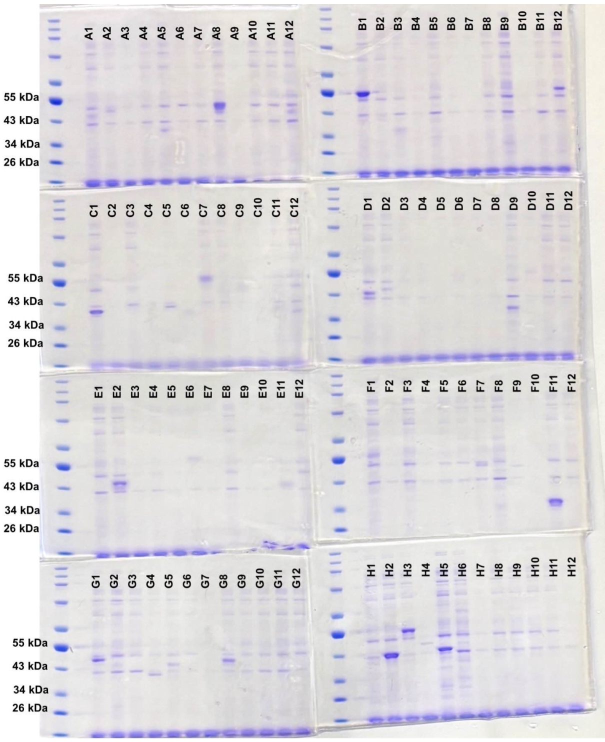

(c) Plate C.

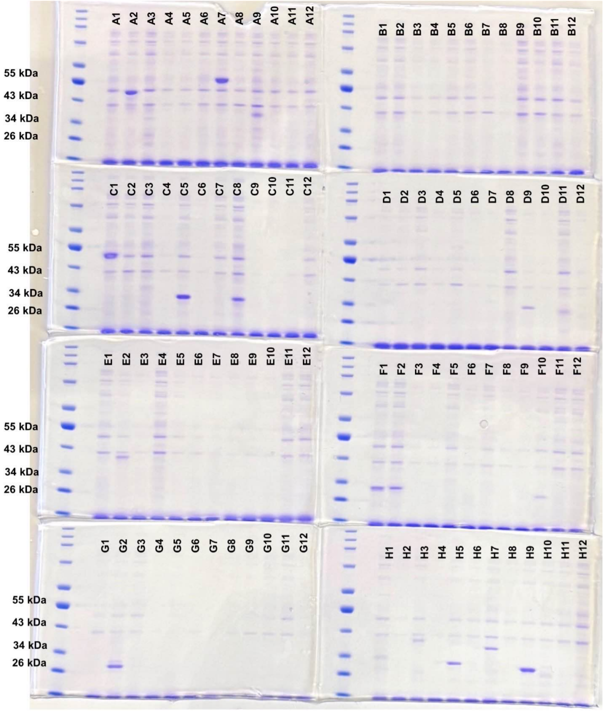

(d) Plate D.

**Figure** S18: **SDS-PAGE of soluble lysate fractions, all four plates.**

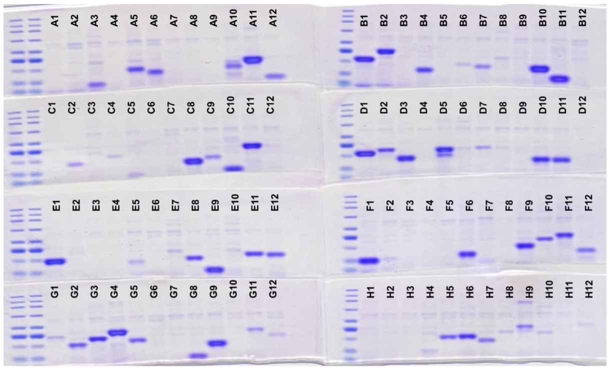

(a) Plate A.

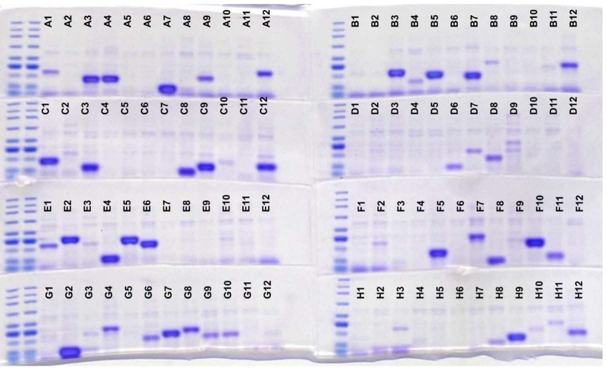

(b) Plate B.

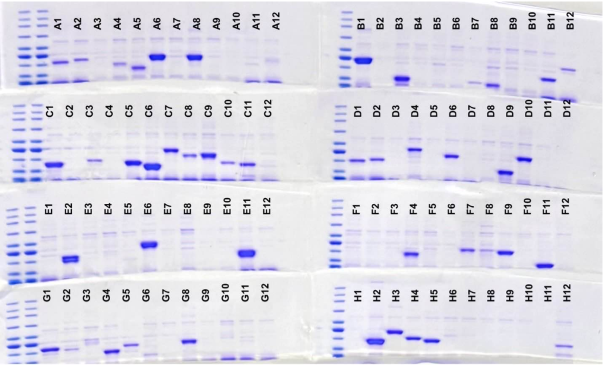

(c) Plate C.

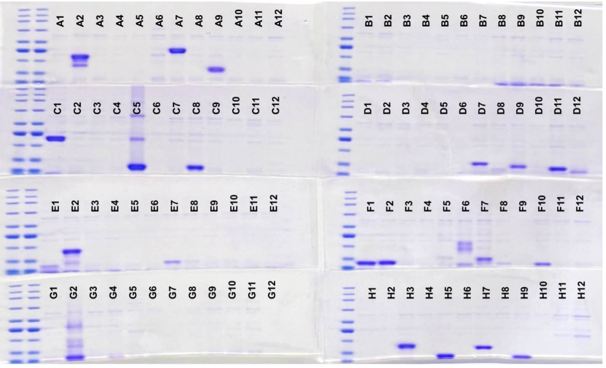

(d) Plate D.

**Figure** S19: **SDS-PAGE of Ni-NTA enriched fractions, all four plates.**

### S10. Expression-prediction models

#### S10.1. Features and classifiers

We tested whether recombinant expression can be predicted from sequence and structure-derived features, and in particular whether AMPLIFY-C embeddings discriminate expression more reliably than the conventional biophysical and structural descriptors. The task is binary classification of expression outcome for the 287 curated UniProtKB enzymes assayed on Plates A, B, and C (§S8). Two readouts are predicted independently: detectable versus undetectable band in the crude *lysate* fraction and in the affinity-*purified* fraction, with band quantification and the detection threshold set at one. The pool consists of remote homologs (pairwise identity ≤ 50% by construction; §S8), precluding train/test leakage through sequence similarity.

**Feature families..** Four feature families were compared:

(i) **AMPLIFY-C embedding** (960-d) — the per-residue final-layer representations of AMPLIFY-C, mean-pooled over the sequence;
(ii) **Biophysical** (22 descriptors) — aromaticity; grand average of hydropathy (GRAVY); isoelectric point; net charge at pH 7.0, 7.5, and 8.0; mean residue flexibility; instability index; molar extinction coefficient (oxidized and reduced cysteines); molecular weight; Shannon entropy of the residue composition (raw and length-normalized); predicted disordered-residue fraction at three thresholds (0.5, 0.8, 0.9); signal-peptide likelihood; predicted secondary-structure content (helix, sheet, turn); single-amino-acid repeat ratio; and non-standard-residue count;
(iii) **Structural** (4 descriptors), computed from the AlphaFold-predicted structure (*20*): mean pLDDT, mean solvent-accessible surface area (SASA), per-residue ESM-IF negative log-likelihood (*24*), and per-residue soluble-ProteinMPNN log-likelihood (*25*);
(iv) **Biophysical** + **Structural** — the concatenation of families (ii) and (iii).

All features were standardized to zero mean and unit variance using statistics fit on the training split only.

**Classifiers and training..** Two classifiers were compared: $L_2$-regularized logistic regression and LightGBM gradient-boosted trees (*26*). For every (feature family × classifier × split) combination, hyperparameters were selected by stratified 5-fold cross-validation on the training split, scored by mean validation ROC-AUC. LightGBM was tuned by Bayesian optimization (Optuna (*27*), 50 trials) and class-reweighted through `scale_pos_weight`; logistic regression was tuned by grid search over the inverse-regularization strength $C \in \{10^{-4}, 10^{-3}, 10^{-2}, 10^{-1}, 1, 10, 10^2\}$. The selected configuration was refit on the full training split before evaluation. Each configuration was evaluated over 20 independent 85%/15% stratified train/test splits.

**Statistical analysis..** Performance is summarized as the mean ± standard deviation of test ROC-AUC across the 20 splits. Differences between feature families were assessed by a one-sided paired permutation test (10,000 sign-flip permutations) on the per-split AUCs, testing whether the embedding family exceeds the comparator. The resulting $p$-values were corrected for multiple comparisons with the Benjamini–Hochberg procedure at a 5% false-discovery rate. All random operations were controlled by explicit seeds.

#### S10.2. Results

Table S21 reports the test-set ROC-AUC of each feature family and classifier on the two readouts. The AMPLIFY-C embedding attains the highest ROC-AUC on both readouts (∼0.80–0.81), well above the biophysical, structural, and combined descriptor sets (∼0.72–0.76). Classifiers trained on the AMPLIFY-C embedding significantly outperform those trained on every non-embedding family, on both readouts and with both classifiers (one-sided paired permutation test with Benjamini–Hochberg correction; $p \leq 0.011$, and 8 of 12 comparisons at $p < 0.001$; Table S22). This establishes that the AMPLIFY-C representation discriminates wet-lab expression more reliably than biophysical and structure-based descriptors in the context of these experiments. The per-split ROC-AUC values underlying both tables are provided in `expression_prediction_results.csv`.

| Feature family | Lysate band LR | Lysate band LGB | Purified band LR | Purified band LGB |
|---|---|---|---|---|
| AMPLIFY-C embedding (960-d) | 0.812±0.056 | 0.811±0.056 | 0.813±0.058 | 0.798±0.051 |
| Biophysical (22) | 0.726±0.071 | 0.745±0.076 | 0.722±0.085 | 0.721±0.075 |
| Structural (4) | 0.741±0.076 | 0.749±0.080 | 0.750±0.073 | 0.724±0.075 |
| Biophysical + Structural | 0.754±0.075 | 0.763±0.076 | 0.752±0.083 | 0.752±0.065 |

Table S21: **Expression-prediction ROC-AUC by feature family and classifier** (mean ± s.d. over 20 stratified 85%/15% splits; 287-protein complete pool). LR: $L_2$-regularized logistic regression; LGB: LightGBM.

| Comparison | Lysate band LR | Lysate band LGB | Purified band LR | Purified band LGB |
|---|---|---|---|---|
| AMPLIFY-C vs Biophysical | < 0.001 | < 0.001 | < 0.001 | < 0.001 |
| AMPLIFY-C vs Structural | < 0.001 | 0.002 | < 0.001 | 0.001 |
| AMPLIFY-C vs Biophysical+Structural | < 0.001 | < 0.001 | 0.001 | 0.011 |

Table S22: **AMPLIFY-C versus non-embedding feature families: one-sided paired-permutation $p$-values** (10,000 sign-flip permutations testing whether the AMPLIFY-C embedding exceeds the comparator; Benjamini–Hochberg-corrected across all 12 comparisons at 5% FDR; 287-protein complete pool).

### S11. Metagenomic retrieval: results and characterization

#### S11.1. Retrieval funnel and per-candidate data

Table S23 traces the metagenomic candidates from the raw databases to the experimentally validated set. Each stage corresponds to a step of the pipeline in §S8.3 (prefiltering, kNN, deduplication, MMR) and the wet-lab assay (§S9). Expression is scored as a detectable band (> 10) in the lysate or purified readout; high expression as a band > 500 in either readout; low structural confidence as an OpenFold (*28*) mean pLDDT < 70.

Table S23: **Metagenomic retrieval funnel**, from the raw BFD+MGnify databases to the validated Plate D candidates.

| Stage | Count | Operation |
|---|---|---|
| Raw BFD + MGnify | $\approx 4.5 \times 10^9$ | input databases |
| After RED, length, HXD/E prefilter | $\approx 5 \times 10^8$ | bottom-50% RED, length 200–450, motif |
| Unique kNN candidates | 44,005 | $k$=1,000 cosine kNN + dedup (>80% to targets, non-Met fragments) |
| Plate D (selected) | 96 | MMR ($\lambda$=0.9, ≤50% identity) |
| Expressed | 30 | lysate or purified band > 10 |
| Highly expressed | 24 | lysate or purified band > 500 |
| Low structural confidence | 7 | highly expressed with OpenFold pLDDT < 70 |

**Per-candidate data..** Per-candidate Plate D structural-confidence and homology features are provided in `metagenomic_additional_features.csv` (one row per sequence): `id`, `sequence`, `length`, source database (`dataset`), `panel`, `openfold_plddt`, and the MMseqs2 maximum identity to EC 1.14.11 (0 for all candidates). The corresponding wet-lab expression measurements (lysate/purified band, yield) are in `in_vivo_results.csv`, and the in silico scores (RED, AMPLIFY-C perplexity, and the biophysical/structural descriptors) for these and the curated-plate candidates are in `all_plates_features.parquet`.

#### S11.2. Predicted structures of expressed candidates

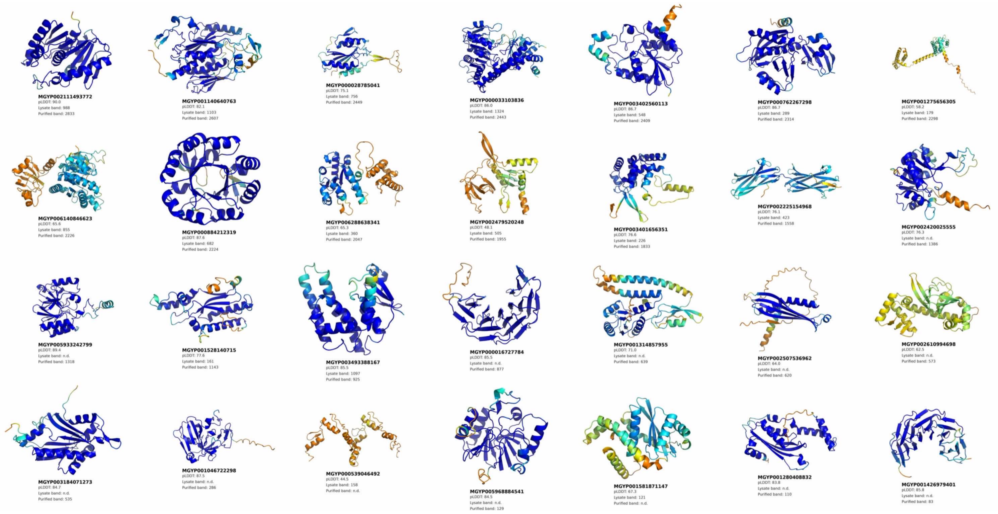

**Figure** S20: Predicted structures with OpenFold for 28 expressed metagenomic sequences.

##### S11.3. Low-pLDDT expression

Of the 24 highly expressed Plate D candidates (Table S23), seven receive low predicted structural confidence from OpenFold (*28*) (mean pLDDT < 70; Table S24). These sequences would have been invisible to the conventional toolkit: they lack the sequence identity required for homology search (§S8.4) and the fold confidence that would flag them as plausible candidates, yet they express as soluble proteins. This demonstrates that the AMPLIFY-C embedding encodes information about expressibility that predicted structural confidence does not: it ranked these sequences as viable where pLDDT would have discarded them. Predicted structures for 28 expressed candidates are shown in Figure S20.

Table S24: **Highly expressed Plate D candidates with low predicted structural confidence** (OpenFold mean pLDDT < 70 and lysate or purified band > 500). Band volumes are in arbitrary SDS-PAGE units. Six are MGnify-derived and one is BFD-derived.

| Sequence ID | OpenFold pLDDT | Lysate band | Purified band |
|---|---|---|---|
| MGYP002479520248 | 48.1 | 505 | 1,955 |
| MGYP001275656305 | 58.2 | 179 | 2,298 |
| BFD_AP12_2_1047962.scaffolds.fasta_ scaffold13649_1 | 60.9 | 283 | 1,104 |
| MGYP002610994698 | 62.5 | 0 | 573 |
| MGYP002507536962 | 64.0 | 0 | 620 |
| MGYP006288638341 | 65.3 | 360 | 2,047 |
| MGYP006140846623 | 65.6 | 855 | 2,226 |

#### S12. Software, reproducibility, and data availability

##### S12.1. Software and library versions

Table S25 lists the principal software used.

| Component | Version | Reference |
|---|---|---|
| *Pretraining and modeling* | | |
| PyTorch | 2.5.0 | — |
| CUDA | 12.6.0 | — |
| FlashAttention | 2.8.3.post1 | (5) |
| DeepSpeed | 0.18.0 | (7) |
| Accelerate | ⟨⟩ | (6) |
| *Structure, homology, and biophysics* | | |
| Rosetta | 2025.51 | (18) |
| AlphaFold2 | ⟨⟩ | (20) |
| OpenFold | ⟨⟩ | (28) |
| ProteinMPNN | ⟨⟩ | (25) |
| ESM-IF | ⟨⟩ | (24) |
| HHblits | ⟨⟩ | (14) |
| BLAST | ⟨⟩ | (22) |
| MMseqs2 | ⟨⟩ | (23) |
| *Analysis and machine learning* | | |
| Optuna | ⟨⟩ | (27) |
| LightGBM | ⟨⟩ | (26) |
| scikit-learn | ⟨⟩ | — |
| UMAP | ⟨⟩ | (29) |

Table S25: **Software and library versions.** Entries marked ⟨⟩ are pending final confirmation.

##### S12.2. Random seeds and hardware

**Random seeds..** All stochastic procedures used explicit seeds. The supervised property-prediction grids (§S6.1) were repeated over seeds $\{42, 43, 44\}$; the expression-prediction benchmark (§S10) was evaluated over 20 stratified train/test splits (seeds 0–19). Permutation tests used 10,000 sign-flip resamples under a fixed seed.

**Hardware..** Pretraining was performed on NVIDIA H100 GPUs. Evaluation and preprocessing ran on a heterogeneous mix of GPUs available on the Mila and Digital Research Alliance of Canada (Compute Canada) clusters, including NVIDIA A100, L40S, and RTX-series cards; because these workloads are not performance-sensitive, the specific device varied by job.

##### S12.3. Data, model, and code availability

Ablation models, final models (AMPLIFY-B and AMPLIFY-C), and datasets with scores are released publicly at `https://huggingface.co/flair-bio`. The pre-processing, training, evaluation, and wet lab candidate selection code is released at `https://github.com/flair-bio`.

**Column dictionaries..** The data files listed in the front matter have the following schemas.

**`in_silico_results.csv`** one row per ablation model. `model_name` (registry id, §S3); `1m` (boolean; `True`=1M steps, `False`=100k); `filtering` (RED quantile applied, `0.0`=unfiltered); *KL homologs (mean)* (mean $D_{\mathrm{KL}}(p_{\mathrm{homologs}} \,\|\, p_{\mathrm{model}})$ over consensus positions); *Calibration (T)*, *Calibration (NT)* (per-class calibration scores, §S7.1); *Categorical Jacobian (CASP16)* (AUC, mean binned precision); *Deep mutational scanning* (ProteinGym weighted Spearman); *Property (FE, ext.)*, *Property (FE, rest.)* (aggregate property-prediction scores under the extended/restricted grids).

**`property_prediction_results.csv`** one row per model (`model_name`, `1m`, `filtering` as above) and one column per task × grid, named `<Task> (FE, Extended)` / `<Task> (FE, Restricted)` for the 17 tasks of Table S18.

**`categorical_jacobian_results.csv`** one row per model. `model_name`; `model` (Hugging Face repo path); `split` (`casp16`); `P@L`, `P@L/2`, `P@L/5`, `AUC`, `ROC-AUC` (means over targets); `n_sequences` (number of CASP16 targets, 49); and the matching `*_std` columns (standard deviation over targets).

**`expression_prediction_results.csv`** one row per (feature, task, classifier, split). `regime` (`complete`); `seed` (0–19); `feature` (feature family); `task` (`lysate`/`purified`); `method` (`lr`/`lgb`); `n_train`, `n_test`; `test_auc`.

**`human_proteome.csv`** `id` (UniProtKB accession); `sequence`.

**`human_consensus_positions.csv`** `sequence_name` (UniProtKB id); `position` (0-based residue index); `wt_aa` (human residue); and 20 columns `A`...`Y` giving the homolog amino-acid frequencies at that position.

**`all_plates_features.parquet`** per-candidate in silico features, all four plates. `id`, `sequence`, `length`, `plate`, `dataset`, `AMPLIFY_consensus_embedding`, `AMPLIFY_consensus_red`, `AMPLIFY_consensus_pseudo_ppl`, the 22 biophysical descriptors, and the structural descriptors (`mean_plddt`, `mean_sasa`, `esm_if_nll`, `mpnn_ll_soluble`).

**`in_vivo_results.csv`** wet-lab measurements and synthesized-construct details, one row per construct (all four plates). `seq ID`, `location`, `sequence`, `kDa`, `lysate band`, `purified band`, `mg/mL`, `plate`, `well`, `gene_index`, `gene_name`, `insertion_point_name`, `vector_name`, `insert_sequence`.

**`metagenomic_additional_features.csv`** Plate D candidates. `id`, `sequence`, `length`, `dataset`, `panel`, `openfold_plddt`, `mmseqs2_ec_1.14.11` (maximum MMseqs2 identity to EC 1.14.11).

**`pairwise_cosine_similarity.parquet`** symmetric $384 \times 384$ matrix of AMPLIFY-C embedding cosine similarities; rows and columns are candidate ids.

**`kde_boundaries.csv`** `dataset`; `red_boundary` (per-dataset RED filtering threshold); `qt` (percentile of that threshold within the dataset).

**`ddg_results.csv`, `ddg_results_random.csv`** one row per scored mutation (consensus and random control respectively). `uniprot_id`, `af_id`, `res_ix`/`res_1based`, `AA` (wild-type), `most_common` (consensus residue), `max_freq`, `mutation_id`, `wt_score`, `mut_score`, `ddg`, `stabilizing` (boolean, $\Delta\Delta G < 0$).

**`literature_hydroxylases.csv`** reference enzymes: `Enzyme`, `UniProt ID`, `Reaction`, `Kingdom`, `Organism`, `Sequence`.

##### S12.4. Statistical methods

**Significance testing..** Pairwise comparisons between feature families (§S10) were assessed by a paired permutation test on the per-split metric (10,000 sign-flip permutations), valid because all families were evaluated on identical splits. Tests were one-sided when the hypothesis was directional (a pLM feature exceeding a comparator). $p$-values were corrected for multiple comparisons with the Benjamini–Hochberg procedure at a 5% false-discovery rate, applied across the family of comparisons reported in the corresponding table.

**Correlations..** Rank correlations (RED versus perplexity, §S4.3; DMS fitness, §S6.4) are Spearman coefficients; perplexity–expression associations in the main text are Pearson coefficients with the $n$ and $p$ stated alongside.
