## Supplementary material for "pLM representations unlock metagenomic space beyond homology": Code bundle: accelerate_ddp.yaml

```
compute_environment: LOCAL_MACHINE
debug: false
distributed_type: MULTI_GPU
downcast_bf16: 'no'
enable_cpu_affinity: false
gpu_ids: all
machine_rank: 0
main_training_function: main
mixed_precision: fp16
num_machines: 1
num_processes: 1
rdzv_backend: static
same_network: true
tpu_env: []
tpu_use_cluster: false
tpu_use_sudo: false
use_cpu: false
```
