## Supplementary material for "pLM representations unlock metagenomic space beyond homology": Code bundle: csv_to_fasta.py

```
import csv
from tqdm import tqdm
from argparse import ArgumentParser


if __name__ == "__main__":
    parser = ArgumentParser()
    parser.add_argument("-i", "--inputs", nargs="+", help="path to the input csv file")
    parser.add_argument("-o", "--output", help="path to the output fasta file")
    args = parser.parse_args()

    writer = open(args.output, "w")

    for input_path in args.inputs:
        infile = open(input_path, "r")
        reader = csv.reader(infile)
        next(reader)
        for row in tqdm(reader):
            if len(row) > 1:
                identifier = row[0].strip()
                sequence = row[1].strip()
                writer.write(f">{identifier}\n{sequence}\n")
            else:
                print(f"Skipping row due to insufficient columns: {row}")
        infile.close()

    writer.close()
```
