## Supplementary material for "pLM representations unlock metagenomic space beyond homology": Code bundle: 03_plm_embeddings.sh

```
#!/bin/bash
#SBATCH --job-name=03_plm_embeddings
#SBATCH --ntasks=1
#SBATCH --cpus-per-task=6
#SBATCH --gpus-per-task=1
#SBATCH --mem=32G
#SBATCH --time=1-00:00:00
#SBATCH --output=$HOME/enzymes/output/%x_%j.out
#SBATCH --error=$HOME/enzymes/output/%x_%j.out
#SBATCH --partition=unkillable

# Change to the project directory
cd $HOME/enzymes

module load python/3.10 cuda/12.6.0

# Activate the environment
source $HOME/AMPLIFY-private/.venv/bin/activate

# Hugging Face organization hosting the released checkpoints.
HF_ORG=${HF_ORG:-flair-bio}

# `model` doubles as the column-name prefix used downstream.
model=AMPLIFY-C
model_repo=${HF_ORG}/${model}

# Get the arguments from CONFIGS based on the SLURM_ARRAY_TASK_ID
python scripts/03_plm_embeddings.py \
    --input_csv $SCRATCH/proseqo/datasets/uniprot/uniprot.csv \
    --data_name uniprot \
    --model_name $model \
    --model_path $model_repo \
    --output_dir $SCRATCH/proseqo/enzymes/output \
    --max_len 512 \
    --batch_size 32 \
    --mask_type boolean \

# # Get the arguments from CONFIGS based on the SLURM_ARRAY_TASK_ID
# python scripts/03_plm_embeddings.py \
#     --input_csv $HOME/enzymes/data/known_hydroxylases.csv \
#     --data_name known_hydroxylases \
#     --model_name $model \
#     --model_path $model_repo \
#     --output_dir $SCRATCH/proseqo/enzymes/output \
#     --max_len 512 \
#     --batch_size 32 \
#     --mask_type boolean \
```
