## Supplementary material for "pLM representations unlock metagenomic space beyond homology": Code bundle: 04_plm_perplexity.sh

```
#!/bin/bash
#SBATCH --job-name=04_plm_perplexity
#SBATCH --ntasks=1
#SBATCH --cpus-per-task=6
#SBATCH --gres=gpu:4
#SBATCH --mem=32G
#SBATCH --time=03:00:00
#SBATCH --output=$HOME/enzymes/output/%x_%j.out
#SBATCH --error=$HOME/enzymes/output/%x_%j.out
#SBATCH --partition=short-unkillable

# Change to the project directory
cd $HOME/enzymes

module load python/3.10 cuda/12.6.0

# Activate the environment
source $HOME/AMPLIFY-private/.venv/bin/activate

# MASTER_PORT=1900
# MASTER_ADDR=cn-n001
# SLURM_NODEID=O
# SLURM_GPUS_ON_NODE=1

export MASTER_PORT=$(expr 10000 + $(echo -n $SLURM_JOBID | tail -c 4))
export MASTER_ADDR=$(scontrol show hostnames $SLURM_JOB_NODELIST | head -n 1)

# Hugging Face organization hosting the released checkpoints.
HF_ORG=${HF_ORG:-flair-bio}

# `model` doubles as the column-name prefix used downstream.
model=AMPLIFY-C
model_repo=${HF_ORG}/${model}

python scripts/04_plm_perplexity.py \
    --model_name $model \
    --model_path $model_repo \
    --output_dir $SCRATCH/proseqo/swissprot_diff \
    --data_name removed_sprot \
    --input_csv $SCRATCH/proseqo/swissprot_diff/removed_sequences.csv \
    --max_len 2048 \
    --packing

# python scripts/04_plm_perplexity.py \
#     --model_name $model \
#     --model_path $model_repo \
#     --output_dir $SCRATCH/proseqo/enzymes/output \
#     --data_name all_selected \
#     --input_csv $HOME/enzymes/data/all_selected.csv \
#     --max_len 450 \
#     --packing

# cmd=(
#     accelerate launch \
#         --config_file=$HOME/enzymes/accelerate_ddp.yaml \
#         --machine_rank=\$SLURM_NODEID \
#         --main_process_ip=$MASTER_ADDR \
#         --main_process_port=$MASTER_PORT \
#         --num_processes=$SLURM_GPUS_ON_NODE \
#         --num_machines=1 \
#     scripts/04_plm_perplexity.py \
#     --model_name $model \
#     --model_path $model_repo \
#     --output_dir $SCRATCH/proseqo/enzymes/output \
#     --data_name all_selected \
#     --input_csv $HOME/enzymes/data/all_selected.csv \
#     --max_len 450 \
#     --packing
# )


srun --kill-on-bad-exit=1 bash -c "$(for a in "${cmd[@]}" ; do echo -n \"$a\" "" ; done)"
```
