## Supplementary material for "pLM representations unlock metagenomic space beyond homology": Code bundle: 08_proteinmpnn.sh

```
#!/bin/bash
#SBATCH --job-name=08_proteinmpnn
#SBATCH --ntasks=1
#SBATCH --cpus-per-task=12
#SBATCH --mem=32G
#SBATCH --time=1-00:00:00
#SBATCH --output=$HOME/enzymes/output/%x_%j.out
#SBATCH --error=$HOME/enzymes/output/%x_%j.out

# Change to the project directory
cd $HOME/enzymes

module load python/3.10

# Activate the environment
source env/bin/activate

# # Get the arguments from CONFIGS based on the SLURM_ARRAY_TASK_ID
# python scripts/08_proteinmpnn.py \
#     --input_csv data/uniprotkb_ec_1_14_11_2025_10_20.csv \
#     --output_dir $SCRATCH/proseqo/enzymes/output \
#     --n_samples 0


# python scripts/08_proteinmpnn.py \
#     --input_csv data/bfd_candidates.csv \
#     --output_dir $SCRATCH/proseqo/enzymes/bfd_output \
#     --model_family soluble \
#     --n_samples 0

python scripts/08_proteinmpnn.py \
    --input_csv data/mgnify_candidates.csv \
    --output_dir $SCRATCH/proseqo/enzymes/mgnify_output \
    --model_family soluble \
    --n_samples 0
```
