## Supplementary material for "pLM representations unlock metagenomic space beyond homology": Code bundle: 10_pairwise_cossim.sh

```
#!/bin/bash
#SBATCH --job-name=10_pairwise_cossim
#SBATCH --ntasks=1
#SBATCH --cpus-per-task=6
#SBATCH --mem=256G
#SBATCH --time=01:00:00
#SBATCH --output=$HOME/enzymes/output/%x_%j.out
#SBATCH --error=$HOME/enzymes/output/%x_%j.out

# Change to the project directory
cd $HOME/enzymes

module load python/3.10

# Activate the environment
source env/bin/activate

file=plm_embeddings_AMPLIFY-C.parquet

# Get the arguments from CONFIGS based on the SLURM_ARRAY_TASK_ID
python scripts/10_pairwise_cossim.py \
    --input_parquet $file \
    --output_dir $SCRATCH/proseqo/enzymes/metagenomic_output \
    --n_samples 0
```
