## Supplementary material for "pLM representations unlock metagenomic space beyond homology": Code bundle: 12_openfold.sh

```
#!/bin/bash
#SBATCH --job-name=12_openfold
#SBATCH --ntasks=1
#SBATCH --cpus-per-task=12
#SBATCH --gres=gpu:1
#SBATCH --mem=32G
#SBATCH --time=1-00:00:00
#SBATCH --output=$HOME/enzymes/output/%x_%j.out
#SBATCH --error=$HOME/enzymes/output/%x_%j.out
#SBATCH --partition=lab-chandar
#SBATCH --array=0-7

# Change to the project directory
cd $HOME/enzymes

module purge
module load anaconda/3

conda init
conda activate openfold

module load cuda/12.6.0

SLURM_ARRAY_TASK_ID=7

# Get the arguments from CONFIGS based on the SLURM_ARRAY_TASK_ID
python scripts/12_openfold.py \
    --input_csv $HOME/enzymes/data/metagenomic_selected/metagenomic_selected.csv \
    --output_dir $SCRATCH/proseqo/enzymes/metagenomic_output/openfold \
    --n_samples 0 \
```
