## Supplementary material for "pLM representations unlock metagenomic space beyond homology": Code bundle: filter.sh

```
#!/bin/bash
#SBATCH --job-name=filter_dataframes
#SBATCH --time=02:00:00
#SBATCH --error=$HOME/enzymes/output/output_%x_%j.txt
#SBATCH --output=$HOME/enzymes/output/output_%x_%j.txt
#SBATCH --cpus-per-task=6            # number of cpus per gpu
#SBATCH --mem=200G                    # memory

cd $HOME/enzymes
module load python/3.10

# Activate the environment
source env/bin/activate

python scripts/filter_dataframes.py \
    --input_dir $SCRATCH/proseqo/enzymes/output \
    --output_dir $SCRATCH/proseqo/enzymes/filtered_outputs \
    --csv_file $HOME/enzymes/data/uniprotkb_ec_1_14_11_2025_10_20.csv \
    --pairwise_similarity_file "$SCRATCH/proseqo/enzymes/filtered_outputs/pairwise_similarity_000.parquet" \
    --pairwise_identity_file "$SCRATCH/proseqo/enzymes/filtered_outputs/pairwise_identity_000.parquet" \
    --pairwise_blast_file "$SCRATCH/proseqo/enzymes/filtered_outputs/pairwise_blast_000.parquet" \
```
