## Supplementary material for "pLM representations unlock metagenomic space beyond homology": Code bundle: filter_bfd.sh

```
#!/bin/bash
#SBATCH --job-name=filter_for_enzymes
#SBATCH --time=1-00:00:00
#SBATCH --error=$HOME/enzymes/output/output_%x_%j.txt
#SBATCH --output=$HOME/enzymes/output/output_%x_%j.txt
#SBATCH --cpus-per-task=6            # number of cpus per gpu
#SBATCH --mem=400G                    # memory


# # Check if input file is provided
# if [ "$#" -ne 2 ]; then
#     echo "Usage: $0 <input.fasta.gz> <output.fasta>"
#     exit 1
# fi

# INPUT_FILE=$1
# OUTPUT_FILE=$2

INPUT_FILE=$SCRATCH/proseqo/datasets/bfd/train/bfd_train_no_ambig.fasta.gz
OUTPUT_FILE=$SCRATCH/proseqo/datasets/bfd/filtered/bfd_potential_enzymes_00.fasta

echo "Processing $INPUT_FILE..."

# Definition of Standard Amino Acids
AAS="ACDEFGHIKLMNPQRSTVWY"

# Combined Regex Logic:
# 1. ^[$AAS]* : Start with any number of standard AAs
# 2. H             : The 'H' in H-X-D/E
# 3. [$AAS]        : The 'X' (must be a standard AA)
# 4. [DE]          : The 'D' or 'E'
# 5. [$AAS]*$      : End with any number of standard AAs
#
# This regex ensures the WHOLE sequence is composed of standard AAs (no X, B, Z, etc.)
# AND that it contains the specific motif.

REGEX="^[$AAS]*H[$AAS][DE][$AAS]*$"

# Pipeline:
# 1. seqkit seq: Fast length filtering (Condition 1)
# 2. seqkit grep: Combined regex for ambiguity and motif (Conditions 2 & 3)
# Note: 'seq' is separate because it is extremely fast at parsing headers/length
# without needing to process the full sequence body like 'grep' does.

seqkit seq -w 0 -m 100 -M 450 "$INPUT_FILE" \
  | seqkit grep -w 0 -s -r -p "$REGEX" \
  > "$OUTPUT_FILE"

echo "Done. Filtered sequences saved to $OUTPUT_FILE"
```
