## Supplementary material for "pLM representations unlock metagenomic space beyond homology": Code bundle: jobs.sh

```
#!/bin/bash
#SBATCH --job-name=enzymes
#SBATCH --ntasks=1
#SBATCH --cpus-per-task=8
#SBATCH --gpus-per-task=a100l:1
#SBATCH --mem=64G
#SBATCH --time=3-00:00:00
#SBATCH --output=$HOME/enzymes/output/%x_%j.out
#SBATCH --error=$HOME/enzymes/output/%x_%j.out

# Define the jobs
JOBS=(
    # "scripts/01_physicochem.py 32"
    # "scripts/02_pairwise_identity.py 32"
    # "scripts/03_plm_embeddings.py AMPLIFY_350M chandar-lab/AMPLIFY_350M additive"
    # "scripts/03_plm_embeddings.py AMPLIFY_350M chandar-lab/AMPLIFY_350M additive"
    # "scripts/04_plm_perplexity.py AMPLIFY-C ${HF_ORG:-flair-bio}/AMPLIFY-C"
    # "scripts/05_structural_properties.py 32"
    # "scripts/06_signalp.py"
    # "scripts/07_deeptmhmm.py"
    # "scripts/08_proteinmpnn.py vanilla"
    # "scripts/08_proteinmpnn.py soluble"
    # "scripts/09_esm_if.py"
)

# Change to the project directory
cd $HOME/enzymes

module load python/3.10

# Activate the environment
source env/bin/activate

IFS=' ' read -r -a ARGS <<< "${JOBS[$SLURM_ARRAY_TASK_ID]}"

# Get the arguments from CONFIGS based on the SLURM_ARRAY_TASK_ID
python ${ARGS[0]} \
    --model_name ${ARGS[1]} \
    --model_path ${ARGS[2]} \
    --output_dir $SCRATCH/proseqo/enzymes/output \
    --max_len 512 \
    --packing
```
