## Supplementary material for "pLM representations unlock metagenomic space beyond homology": Code bundle: mmseqs.sh

```
#!/bin/bash
#SBATCH --job-name=mmseqs
#SBATCH --time=1-00:00:00
#SBATCH --error=$HOME/enzymes/output/output_%x_%j.txt
#SBATCH --output=$HOME/enzymes/output/output_%x_%j.txt
#SBATCH --cpus-per-task=6            # number of cpus per gpu
#SBATCH --mem=1000G                    # memory

source $HOME/AMPLIFY-private/.venv/bin/activate
cd $HOME/enzymes

# python scripts/mmseqs.py \
#   --query_fasta data/metagenomic_selected/active_metagenomic.fasta \
#   --target_fasta $SCRATCH/proseqo/datasets/uniprot/uniprot_trembl.fasta.gz \
#   --output_csv data/metagenomic_selected/active3_uniprot_trembl_mmseqs.csv \
#   --mmseqs_path data/mmseqs/bin/mmseqs \
#   --cores 6 \
#   --tmp_dir /tmp \
#   --m8_dir $SCRATCH/proseqo/enzymes/metagenomic_output/mmseqs_m8

# python scripts/mmseqs.py \
#   --query_fasta data/metagenomic_selected/active_metagenomic.fasta \
#   --target_fasta $SCRATCH/proseqo/datasets/uniprot/uniprot_sprot.fasta.gz \
#   --output_csv data/metagenomic_selected/active3_uniprot_sprot_mmseqs.csv \
#   --mmseqs_path data/mmseqs/bin/mmseqs \
#   --cores 6 \
#   --tmp_dir /tmp \
#   --m8_dir $SCRATCH/proseqo/enzymes/metagenomic_output/mmseqs_m8

# python scripts/mmseqs.py \
#   --query_fasta data/metagenomic_selected/expressed_metagenomic.fasta \
#   --target_fasta $SCRATCH/proseqo/datasets/uniprot/uniprot_sprot.fasta.gz \
#   --output_csv data/metagenomic_selected/expressed3_uniprot_sprot_mmseqs.csv \
#   --mmseqs_path data/mmseqs/bin/mmseqs \
#   --cores 6 \
#   --tmp_dir /tmp \
#   --m8_dir $SCRATCH/proseqo/enzymes/metagenomic_output/mmseqs_m8

python scripts/mmseqs.py \
  --query_fasta data/metagenomic_selected/expressed_metagenomic.fasta \
  --target_fasta $SCRATCH/proseqo/datasets/uniprot/uniprot_trembl.fasta.gz \
  --output_csv data/metagenomic_selected/expressed3_uniprot_trembl_mmseqs.csv \
  --mmseqs_path data/mmseqs/bin/mmseqs \
  --cores 6 \
  --tmp_dir /tmp \
  --m8_dir $SCRATCH/proseqo/enzymes/metagenomic_output/mmseqs_m8
```
