## Supplementary material for "pLM representations unlock metagenomic space beyond homology": Code bundle: run_mmr.sh

```
#!/bin/bash
#SBATCH --job-name=run_mmr_metagenomic
#SBATCH --time=03:00:00
#SBATCH --error=$HOME/enzymes/output/%x_%j.txt
#SBATCH --output=$HOME/enzymes/output/%x_%j.txt
#SBATCH --cpus-per-task=6            # number of cpus per gpu
#SBATCH --mem=64G                    # memory
# #SBATCH --partition=main-cpu

cd $HOME/enzymes/

module load python/3.10
source env/bin/activate

output_dir=$HOME/enzymes/selection_logs/filter_metagenomic_000

mkdir -p $output_dir

python optimization/marginal_relevance.py \
    --input_dir $SCRATCH/proseqo/enzymes/metagenomic_output/dataframes \
    --output_dir $output_dir \
    --version 0 \
    --id_col metagenomic_id \
    --blast_threshold 50 \
    --N_select 96 \
    --lambda_param 0.80 \
```
