## Supplementary material for "pLM representations unlock metagenomic space beyond homology": Code bundle: search_metagenomic.sh

```
#!/bin/bash
#SBATCH --job-name=search_metagenomic
#SBATCH --time=01:00:00
#SBATCH --error=$HOME/enzymes/output/%x_%j_%a.txt
#SBATCH --output=$HOME/enzymes/output/%x_%j_%a.txt
#SBATCH --cpus-per-task=6            # number of cpus per gpu
#SBATCH --mem=100G                    # memory
#SBATCH --array=0-4

cd $HOME/enzymes
module load python/3.10

# Activate the environment
source env/bin/activate

SEARCH_SETS=(
    "bfd_potential_enzymes_0.50"
    "mgy_0_potential_enzymes_0.50"
    "mgy_1_potential_enzymes_0.50"
    "mgy_2_potential_enzymes_0.50"
    "mgy_3_potential_enzymes_0.50"
)

search_set=${SEARCH_SETS[$SLURM_ARRAY_TASK_ID]}
lambda=0.90

echo "Current Job ID: $SLURM_ARRAY_JOB_ID"
echo "Current Task ID: $SLURM_ARRAY_TASK_ID"
echo "Processing search_set: $search_set"

python scripts/metagenomic_search.py \
    --queries_id "$HOME/enzymes/selection_logs/final/selected_uniprot_ids_mmr_lambda_"$lambda"_50.txt" \
    --data_path $SCRATCH/proseqo/enzymes/filtered_outputs/merged_data_004.parquet \
    --parquet_dir $SCRATCH/proseqo/scores/Embedding-scores/$search_set \
    --output_dir $SCRATCH/proseqo/enzymes/metagenomic_search/$search_set/$lambda \
    --k 1000 \
    --query_vector_col AMPLIFY-C_embedding \
    --query_id_col uniprot_id \
    --vector_col embeddings \
    --id_col name \
```
