## Supplementary material for "pLM representations unlock metagenomic space beyond homology": Code bundle: control.ipynb

```
# %% [code]
from pathlib import Path
from functools import reduce

import random
import os
import math
import numpy as np
import pandas as pd
import json
import shutil

from tqdm import tqdm

from scipy.stats import pearsonr, spearmanr
from scipy.spatial.distance import pdist, squareform
from sklearn.metrics.pairwise import cosine_similarity

import matplotlib.pyplot as plt
import seaborn as sns
from umap import UMAP

from utils import merge_dataframes, plot_distribution
%config InlineBackend.figure_format='retina'

# %% [code]
idx=4
KEY_COL = "uniprot_id"

DATA_DIR = Path("$SCRATCH/proseqo/enzymes/filtered_outputs")
LOG_DIR = Path("$HOME/enzymes/selection_logs/control")
LOG_DIR.mkdir(exist_ok=True)

MERGED_FILE = f"merged_data_{idx:03}.parquet"
BLAST_MATRIX_FILE = f"pairwise_blast_{idx:03}.parquet"

# %% [code]
df_main = pd.read_parquet(DATA_DIR / MERGED_FILE)
blast_df = pd.read_parquet(DATA_DIR / BLAST_MATRIX_FILE)

blast_df.sort_index(axis=0, inplace=True)
blast_df.sort_index(axis=1, inplace=True)
df_main.sort_values(KEY_COL, inplace=True)

ids = list(df_main[KEY_COL])

# %% [code]
def select_proteins_mmr(
    blast_matrix,
    n_select=96,
    blast_threshold=50,
    preselected_indices=[],
):
    if preselected_indices == []:
        first_choice = random.choice(range(len(blast_matrix)))
        selected_indices = [first_choice]
        preselected_indices.append(first_choice)
    else:
        selected_indices = []

    n_select -= len(selected_indices)

    # Iterative Selection
    for _ in tqdm(range(n_select)):
        remaining_indices = [i for i in range(len(blast_matrix)) if i not in preselected_indices]
        candidate_to_add = -1

        # Shape: (Num_Remaining, Num_Selected)
        sim_to_selected = blast_matrix[remaining_indices][:, preselected_indices]

        # Calculate max similarity to the selected set for each remaining item
        max_sim_vals = np.max(sim_to_selected, axis=1)

        # Calculate MMR scores for all remaining items
        # mmr = lambda * Quality - (1-lambda) * Max_Sim
        # current_qualities = quality_scores[remaining_indices]
        # mmr_scores = (lambda_param * current_qualities) - ((1 - lambda_param) * max_sim_vals)
        mmr_scores = - max_sim_vals

        # Sort candidates by MMR score descending
        sorted_candidate_args = np.argsort(mmr_scores)[::-1]

        # Apply BLAST Hard Constraint
        # We iterate down the ranked list and pick the first one that passes the BLAST check
        found_next = False
        while not found_next:
            for local_idx in sorted_candidate_args:
                global_idx = remaining_indices[local_idx]

                # Check BLAST against ALL currently selected
                blast_scores = blast_matrix[global_idx, preselected_indices]

                if np.all(blast_scores <= blast_threshold):
                    candidate_to_add = global_idx
                    found_next = True
                    break

            if found_next:
                selected_indices.append(candidate_to_add)
                preselected_indices.append(candidate_to_add)
            else:
                blast_threshold += 5
                print(f"Warning: Could not find enough candidates satisfying BLAST constraint, increasing threshold to {blast_threshold}.")

    return selected_indices

# %% [code]
blast_matrix = blast_df.to_numpy()
blast_threshold = 50

selected_indices = select_proteins_mmr(blast_matrix=blast_matrix, n_select=110)

selected_uniprot_ids = [ids[idx] for idx in selected_indices]

with open("$HOME/enzymes/selection_logs/control/selected_uniprot_ids_50.txt", "w") as f:
    for uid in selected_uniprot_ids:
        f.write(f"{uid}\n")

# %% [code]
output_file = "$HOME/enzymes/selection_logs/control/selected_uniprot_ids_50.txt"
with open(output_file, "r") as file:
    control_ids = [line.strip() for line in file]

# %% [code]
output_file = "$HOME/enzymes/selection_logs/final/selected_uniprot_ids.txt"
with open(output_file, "r") as file:
    selected_uniprot_ids = [line.strip() for line in file]

# %% [code]
tmp = control_ids[:98]

# %% [code]
len(set(selected_uniprot_ids) - set(tmp)) - len(set(selected_uniprot_ids))

# %% [code]
len(set(tmp) - set(selected_uniprot_ids))

# %% [code]
new_ids = list(set(tmp) - set(selected_uniprot_ids))

# %% [code]
with open("$HOME/enzymes/selection_logs/control/all_selected_uniprot_ids.txt", "w") as f:
    for uid in control_ids[:98]:
        f.write(f"{uid}\n")

with open("$HOME/enzymes/selection_logs/control/control_selected_uniprot_ids.txt", "w") as f:
    for uid in new_ids:
        f.write(f"{uid}\n")

# %% [code]
selected_file = "$HOME/enzymes/selection_logs/control/control_selected_uniprot_ids.txt"
with open(selected_file, "r") as f:
    selected_ids = set(line.strip() for line in f if line.strip())

# %% [code]
df_selected = df_main[df_main[KEY_COL].isin(selected_ids)]

# %% [code]
df_selected

# %% [code]
blast_df_selected = blast_df.loc[list(selected_ids)]
blast_df_selected = blast_df_selected[list(selected_ids)]

# %% [code]
blast_df_selected.to_parquet("$HOME/enzymes/selection_logs/control/pairwise_blast_control_plates.parquet")

# %% [code]
df_selected.to_parquet("$HOME/enzymes/selection_logs/control/data_control_plates.parquet")

# %% [code]
df_selected

# %% [code]
fasta_path = "$HOME/enzymes/selection_logs/control/control_selected.fasta"
with open(fasta_path, "w") as f:
    for id, seq in zip(df_selected["uniprot_id"], df_selected["sequence"]):
        f.write(f">{id}\n")
        f.write(f"{seq}\n")

# %% [code]
pdb_files = list(df_selected["pdb_file"].unique())
base_path = "$SCRATCH/proseqo/enzymes"
destination_base_path = "$HOME/enzymes/selection_logs/control/pdb_files"

# %% [code]
for file in pdb_files:
    source_item_path = os.path.join(base_path, file)
    file_name = file.split("/")[-1]
    destination_item_path = os.path.join(destination_base_path, file_name)
    shutil.copy2(source_item_path, destination_item_path)

# %% [code]
```
