## Supplementary material for "pLM representations unlock metagenomic space beyond homology": Code bundle: gbm.ipynb

```
# %% [code]
from pathlib import Path

import numpy as np
import pandas as pd
import warnings
warnings.filterwarnings('ignore')

from scipy.stats import spearmanr
from sklearn.model_selection import train_test_split, StratifiedKFold
from sklearn.preprocessing import StandardScaler
from sklearn.metrics import roc_auc_score, mean_absolute_error, r2_score

import optuna
optuna.logging.set_verbosity(optuna.logging.WARNING)

import matplotlib.pyplot as plt

FIG_DIR = Path('figures')
FIG_DIR.mkdir(exist_ok=True)

%config InlineBackend.figure_format='retina'

# %% [code]
data = pd.read_csv('$HOME/enzymes/notebooks/data.csv')

# %% [markdown]
## Data preprocessing

# %% [code]
def parse_embedding(s):
    return np.fromstring(s.strip('[]'), sep=' ')

X_all = np.stack(data['AMPLIFY-C_embedding'].map(parse_embedding))

targets = {
    'lysate':   data['lysate band'],
    'purified': data['purified band'],
}

datasets = {}
for name, y_raw in targets.items():
    mask = y_raw.notna()
    X = X_all[mask]
    y = y_raw[mask].values.astype(float)
    strat = (y > 0).astype(int)

    X_train, X_test, y_train, y_test = train_test_split(
        X, y, test_size=0.15, random_state=42, stratify=strat
    )
    # Tree methods don't require scaling, but we keep it consistent with the MLP notebook
    scaler = StandardScaler()
    X_train = scaler.fit_transform(X_train)
    X_test  = scaler.transform(X_test)

    datasets[name] = dict(
        X_train=X_train, X_test=X_test,
        y_train=y_train, y_test=y_test,
    )
    pos_tr = (y_train > 0).sum()
    pos_te = (y_test  > 0).sum()
    print(f'{name:10s}  train={len(y_train)} (non-zero={pos_tr})  '
          f'test={len(y_test)} (non-zero={pos_te})')

# %% [markdown]
## Two-stage helper

Same two-stage logic as the MLP notebook:
- **Stage 1** (classifier): predict zero vs. non-zero
- **Stage 2** (regressor): predict magnitude, trained only on positive samples
- **Final prediction**: `P(non-zero) × predicted_value`

Both stages accept any sklearn-compatible estimator, allowing us to swap
XGBoost and LightGBM in and out.

# %% [code]
from sklearn.base import BaseEstimator, RegressorMixin, clone

class TwoStageEstimator(BaseEstimator, RegressorMixin):
    """
    Two-stage wrapper around any pair of sklearn-compatible estimators.
    clf: binary classifier  (0 = no expression, 1 = expression)
    reg: regressor trained only on positive (non-zero) samples
    prediction = P(non-zero) * reg_output.clip(0)
    """
    def __init__(self, clf, reg):
        self.clf = clf
        self.reg = reg

    def fit(self, X, y):
        y_bin = (y > 0).astype(int)
        self.clf_ = clone(self.clf).fit(X, y_bin)
        pos = y > 0
        self.reg_ = clone(self.reg).fit(X[pos], y[pos])
        return self

    def predict(self, X):
        p_pos = self.clf_.predict_proba(X)[:, 1]
        y_reg = self.reg_.predict(X).clip(min=0)
        return p_pos * y_reg

    def clf_proba(self, X):
        return self.clf_.predict_proba(X)[:, 1]

# %% [markdown]
## Baseline 1 — Classifier only (zero vs. non-zero)

# %% [code]
import xgboost as xgb
import lightgbm as lgb

N_TRIALS_BL = 50
N_CV_FOLDS  = 5


def make_clf_objective_xgb(X_tr, y_tr):
    y_bin = (y_tr > 0).astype(int)
    cv = StratifiedKFold(n_splits=N_CV_FOLDS, shuffle=True, random_state=42)
    splits = list(cv.split(X_tr, y_bin))
    scale_pos = (y_bin == 0).sum() / (y_bin == 1).sum()

    def objective(trial):
        params = dict(
            n_estimators      = trial.suggest_int(  'n_estimators',   50,  500),
            max_depth         = trial.suggest_int(  'max_depth',       2,    8),
            learning_rate     = trial.suggest_float('learning_rate', 1e-3, 0.3, log=True),
            subsample         = trial.suggest_float('subsample',      0.5,  1.0),
            colsample_bytree  = trial.suggest_float('colsample_bytree', 0.3, 1.0),
            reg_alpha         = trial.suggest_float('reg_alpha',     1e-5, 10.0, log=True),
            reg_lambda        = trial.suggest_float('reg_lambda',    1e-5, 10.0, log=True),
            scale_pos_weight  = scale_pos,
            use_label_encoder = False,
            eval_metric       = 'logloss',
            random_state      = 0,
            device            = 'cuda',
        )
        aucs = []
        for tr_idx, va_idx in splits:
            m = xgb.XGBClassifier(**params)
            m.fit(X_tr[tr_idx], y_bin[tr_idx])
            p = m.predict_proba(X_tr[va_idx])[:, 1]
            aucs.append(roc_auc_score(y_bin[va_idx], p))
        return float(np.mean(aucs))

    return objective


def make_clf_objective_lgb(X_tr, y_tr):
    y_bin = (y_tr > 0).astype(int)
    cv = StratifiedKFold(n_splits=N_CV_FOLDS, shuffle=True, random_state=42)
    splits = list(cv.split(X_tr, y_bin))
    scale_pos = (y_bin == 0).sum() / (y_bin == 1).sum()

    def objective(trial):
        params = dict(
            n_estimators      = trial.suggest_int(  'n_estimators',   50,  500),
            max_depth         = trial.suggest_int(  'max_depth',       2,    8),
            learning_rate     = trial.suggest_float('learning_rate', 1e-3, 0.3, log=True),
            num_leaves        = trial.suggest_int(  'num_leaves',      8,   64),
            subsample         = trial.suggest_float('subsample',      0.5,  1.0),
            colsample_bytree  = trial.suggest_float('colsample_bytree', 0.3, 1.0),
            reg_alpha         = trial.suggest_float('reg_alpha',     1e-5, 10.0, log=True),
            reg_lambda        = trial.suggest_float('reg_lambda',    1e-5, 10.0, log=True),
            scale_pos_weight  = scale_pos,
            random_state      = 0,
            device            = 'gpu',
            verbosity         = -1,
        )
        aucs = []
        for tr_idx, va_idx in splits:
            m = lgb.LGBMClassifier(**params)
            m.fit(X_tr[tr_idx], y_bin[tr_idx])
            p = m.predict_proba(X_tr[va_idx])[:, 1]
            aucs.append(roc_auc_score(y_bin[va_idx], p))
        return float(np.mean(aucs))

    return objective


clf_only_estimators = {}   # best model per target

print('Classifier-only  —  CV')
print('=' * 60)
for name, ds in datasets.items():
    X_tr, y_tr = ds['X_train'], ds['y_train']
    y_bin = (y_tr > 0).astype(int)

    best_auc, best_model = -np.inf, None
    for label, make_obj, ModelCls, extra in [
        ('XGB', make_clf_objective_xgb, xgb.XGBClassifier,
         dict(use_label_encoder=False, eval_metric='logloss', device='cuda')),
        ('LGB', make_clf_objective_lgb, lgb.LGBMClassifier,
         dict(device='gpu', verbosity=-1)),
    ]:
        study = optuna.create_study(direction='maximize',
                                    sampler=optuna.samplers.TPESampler(seed=42))
        study.optimize(make_obj(X_tr, y_tr), n_trials=N_TRIALS_BL,
                       show_progress_bar=True)
        print(f'  {name} / {label}  best CV AUC = {study.best_value:.4f}')

        if study.best_value > best_auc:
            best_auc = study.best_value
            bp = {k: v for k, v in study.best_params.items()}
            scale_pos = (y_bin == 0).sum() / (y_bin == 1).sum()
            best_model = ModelCls(**bp, scale_pos_weight=scale_pos,
                                  random_state=0, **extra).fit(X_tr, y_bin)

    clf_only_estimators[name] = best_model

print('\nClassifier-only  —  test-set results')
print('=' * 60)
for name, ds in datasets.items():
    X_te, y_te = ds['X_test'], ds['y_test']
    y_bin = (y_te > 0).astype(int)
    p = clf_only_estimators[name].predict_proba(X_te)[:, 1]
    auc = roc_auc_score(y_bin, p)
    rho = spearmanr(y_te, p).statistic
    print(f'  {name:10s}  AUC={auc:.4f}  Spearman(p_pos vs y)={rho:.4f}')

# %% [markdown]
## Baseline 2 — Regressor only (all samples, including zeros)

# %% [code]
def make_reg_objective_xgb(X_tr, y_tr):
    strat = (y_tr > 0).astype(int)
    cv = StratifiedKFold(n_splits=N_CV_FOLDS, shuffle=True, random_state=42)
    splits = list(cv.split(X_tr, strat))

    def objective(trial):
        params = dict(
            n_estimators     = trial.suggest_int(  'n_estimators',   50,  500),
            max_depth        = trial.suggest_int(  'max_depth',       2,    8),
            learning_rate    = trial.suggest_float('learning_rate', 1e-3, 0.3, log=True),
            subsample        = trial.suggest_float('subsample',      0.5,  1.0),
            colsample_bytree = trial.suggest_float('colsample_bytree', 0.3, 1.0),
            reg_alpha        = trial.suggest_float('reg_alpha',     1e-5, 10.0, log=True),
            reg_lambda       = trial.suggest_float('reg_lambda',    1e-5, 10.0, log=True),
            random_state     = 0,
            device           = 'cuda',
        )
        rhos = []
        for tr_idx, va_idx in splits:
            m = xgb.XGBRegressor(**params)
            m.fit(X_tr[tr_idx], y_tr[tr_idx])
            y_pred = m.predict(X_tr[va_idx]).clip(min=0)
            rhos.append(spearmanr(y_tr[va_idx], y_pred).statistic)
        return float(np.mean(rhos))

    return objective


def make_reg_objective_lgb(X_tr, y_tr):
    strat = (y_tr > 0).astype(int)
    cv = StratifiedKFold(n_splits=N_CV_FOLDS, shuffle=True, random_state=42)
    splits = list(cv.split(X_tr, strat))

    def objective(trial):
        params = dict(
            n_estimators     = trial.suggest_int(  'n_estimators',   50,  500),
            max_depth        = trial.suggest_int(  'max_depth',       2,    8),
            learning_rate    = trial.suggest_float('learning_rate', 1e-3, 0.3, log=True),
            num_leaves       = trial.suggest_int(  'num_leaves',      8,   64),
            subsample        = trial.suggest_float('subsample',      0.5,  1.0),
            colsample_bytree = trial.suggest_float('colsample_bytree', 0.3, 1.0),
            reg_alpha        = trial.suggest_float('reg_alpha',     1e-5, 10.0, log=True),
            reg_lambda       = trial.suggest_float('reg_lambda',    1e-5, 10.0, log=True),
            random_state     = 0,
            device           = 'gpu',
            verbosity        = -1,
        )
        rhos = []
        for tr_idx, va_idx in splits:
            m = lgb.LGBMRegressor(**params)
            m.fit(X_tr[tr_idx], y_tr[tr_idx])
            y_pred = m.predict(X_tr[va_idx]).clip(min=0)
            rhos.append(spearmanr(y_tr[va_idx], y_pred).statistic)
        return float(np.mean(rhos))

    return objective


reg_only_estimators = {}

print('Regressor-only  —  CV')
print('=' * 60)
for name, ds in datasets.items():
    X_tr, y_tr = ds['X_train'], ds['y_train']

    best_rho, best_model = -np.inf, None
    for label, make_obj, ModelCls, extra in [
        ('XGB', make_reg_objective_xgb, xgb.XGBRegressor,
         dict(device='cuda')),
        ('LGB', make_reg_objective_lgb, lgb.LGBMRegressor,
         dict(device='gpu', verbosity=-1)),
    ]:
        study = optuna.create_study(direction='maximize',
                                    sampler=optuna.samplers.TPESampler(seed=42))
        study.optimize(make_obj(X_tr, y_tr), n_trials=N_TRIALS_BL,
                       show_progress_bar=True)
        print(f'  {name} / {label}  best CV Spearman = {study.best_value:.4f}')

        if study.best_value > best_rho:
            best_rho = study.best_value
            bp = study.best_params
            best_model = ModelCls(**bp, random_state=0,
                                  **extra).fit(X_tr, y_tr)

    reg_only_estimators[name] = best_model

print('\nRegressor-only  —  test-set results')
print('=' * 60)
for name, ds in datasets.items():
    X_te, y_te = ds['X_test'], ds['y_test']
    y_pred = reg_only_estimators[name].predict(X_te).clip(min=0)
    rho = spearmanr(y_te, y_pred).statistic
    mae = mean_absolute_error(y_te, y_pred)
    r2  = r2_score(y_te, y_pred)
    print(f'  {name:10s}  Spearman={rho:.4f}  MAE={mae:.2f}  R²={r2:.4f}')

# %% [markdown]
## Two-stage model (classifier + regressor on positives)

# %% [code]
N_TRIALS = 100


def make_twostage_objective(X_tr, y_tr, clf_cls, reg_cls, clf_extra, reg_extra):
    strat = (y_tr > 0).astype(int)
    cv = StratifiedKFold(n_splits=N_CV_FOLDS, shuffle=True, random_state=42)
    splits = list(cv.split(X_tr, strat))
    scale_pos = (strat == 0).sum() / (strat == 1).sum()

    def objective(trial):
        shared = dict(
            n_estimators     = trial.suggest_int(  'n_estimators',   50,  500),
            max_depth        = trial.suggest_int(  'max_depth',       2,    8),
            learning_rate    = trial.suggest_float('learning_rate', 1e-3, 0.3, log=True),
            subsample        = trial.suggest_float('subsample',      0.5,  1.0),
            colsample_bytree = trial.suggest_float('colsample_bytree', 0.3, 1.0),
            reg_alpha        = trial.suggest_float('reg_alpha',     1e-5, 10.0, log=True),
            reg_lambda       = trial.suggest_float('reg_lambda',    1e-5, 10.0, log=True),
        )
        clf = clf_cls(**shared, scale_pos_weight=scale_pos, random_state=0, **clf_extra)
        reg = reg_cls(**shared, random_state=0, **reg_extra)

        rhos = []
        for tr_idx, va_idx in splits:
            m = TwoStageEstimator(clf, reg)
            m.fit(X_tr[tr_idx], y_tr[tr_idx])
            y_pred = m.predict(X_tr[va_idx])
            rhos.append(spearmanr(y_tr[va_idx], y_pred).statistic)
        return float(np.mean(rhos))

    return objective


best_estimators = {}

print('Two-stage  —  CV')
print('=' * 60)
for name, ds in datasets.items():
    X_tr, y_tr = ds['X_train'], ds['y_train']
    strat = (y_tr > 0).astype(int)
    scale_pos = (strat == 0).sum() / (strat == 1).sum()

    best_rho, best_model = -np.inf, None
    for label, clf_cls, reg_cls, clf_extra, reg_extra in [
        ('XGB',
         xgb.XGBClassifier, xgb.XGBRegressor,
         dict(use_label_encoder=False, eval_metric='logloss', device='cuda'),
         dict(device='cuda')),
        ('LGB',
         lgb.LGBMClassifier, lgb.LGBMRegressor,
         dict(device='gpu', verbosity=-1),
         dict(device='gpu', verbosity=-1)),
    ]:
        study = optuna.create_study(direction='maximize',
                                    sampler=optuna.samplers.TPESampler(seed=42))
        study.optimize(
            make_twostage_objective(X_tr, y_tr, clf_cls, reg_cls, clf_extra, reg_extra),
            n_trials=N_TRIALS, show_progress_bar=True,
        )
        print(f'  {name} / {label}  best CV Spearman = {study.best_value:.4f}')

        if study.best_value > best_rho:
            best_rho = study.best_value
            bp = study.best_params
            clf = clf_cls(**bp, scale_pos_weight=scale_pos,
                          random_state=0, **clf_extra)
            reg = reg_cls(**bp, random_state=0, **reg_extra)
            best_model = TwoStageEstimator(clf, reg).fit(X_tr, y_tr)

    best_estimators[name] = best_model

# %% [markdown]
## Test-set evaluation — two-stage

# %% [code]
print('Two-stage  —  test-set results')
print('=' * 60)

fig, axes = plt.subplots(1, 2, figsize=(12, 5))

for ax, (name, ds) in zip(axes, datasets.items()):
    X_te, y_te = ds['X_test'], ds['y_test']
    model  = best_estimators[name]
    y_pred = model.predict(X_te)

    rho_all, p_all = spearmanr(y_te, y_pred)
    pos = y_te > 0
    rho_pos = spearmanr(y_te[pos], y_pred[pos]) if pos.sum() > 1 else None
    y_bin   = (y_te > 0).astype(int)
    p_pos_s = model.clf_proba(X_te)
    auc     = roc_auc_score(y_bin, p_pos_s) if y_bin.sum() > 0 else float('nan')
    mae     = mean_absolute_error(y_te, y_pred)
    r2      = r2_score(y_te, y_pred)

    print(f'\nTarget: {name}')
    print(f'  Spearman (all):      {rho_all:.4f}  (p={p_all:.3g})')
    if rho_pos:
        print(f'  Spearman (non-zero): {rho_pos.statistic:.4f}  (p={rho_pos.pvalue:.3g})')
    print(f'  Classifier AUC:      {auc:.4f}')
    print(f'  MAE:                 {mae:.2f}')
    print(f'  R²:                  {r2:.4f}')

    ax.scatter(y_te[~pos], y_pred[~pos], alpha=0.5, label='true=0',  color='gray',      s=30)
    ax.scatter(y_te[pos],  y_pred[pos],  alpha=0.7, label='true>0',  color='steelblue', s=40)
    lim = max(y_te.max(), y_pred.max()) * 1.05
    ax.plot([0, lim], [0, lim], 'r--', lw=1)
    ax.set_xlabel('True value')
    ax.set_ylabel('Predicted value')
    ax.set_title(f'{name} band  (ρ={rho_all:.3f}, AUC={auc:.3f})')
    ax.legend(fontsize=8)

plt.tight_layout()
plt.savefig(FIG_DIR / 'gbm_twostage_test.png', dpi=150, bbox_inches='tight')
plt.show()
```
