## Supplementary material for "pLM representations unlock metagenomic space beyond homology": Code bundle: maximum_marginal_relevance.ipynb

```
# %% [code]
import numpy as np
import matplotlib.pyplot as plt
import scipy
import random
import torch 
import pyarrow.parquet as pq
import pandas as pd
import time
import os

import numpy as np
from tqdm import tqdm

from pathlib import Path

from utils import (
    plot_distribution
)

device = torch.device("cuda" if torch.cuda.is_available() else "cpu")

# %% [code]
# Define the directory where your results are stored
LOG_DIR = Path("$HOME/enzymes/selection_logs/metagenomic_output")
DATA_DIR = Path("$SCRATCH/proseqo/enzymes/metagenomic_output/dataframes")
FIG_DIR = Path("$SCRATCH/proseqo/enzymes/output/figures")
FIG_DIR.mkdir(exist_ok=True)
LOG_DIR.mkdir(exist_ok=True)

# Define the primary key for merging
KEY_COL = "metagenomic_id"

# Data version
idx = 1

# Add all your generated metrics filenames to this list.
MERGED_FILE = f"merged_data_{idx:03}.parquet"
SIMILARITY_MATRIX_FILE = f"pairwise_similarity_{idx:03}.parquet"
BLAST_MATRIX_FILE = f"pairwise_blast_{idx:03}.parquet"

# %% [code]
print("Loading metrics files...")
df_main = pd.read_parquet(DATA_DIR / MERGED_FILE)
df_main.sort_values(KEY_COL, inplace=True)

df_main = df_main[~df_main["AMPLIFY-C_pseudo_ppl"].isna()]
quality = df_main["AMPLIFY-C_pseudo_ppl"]
ids = list(df_main[KEY_COL])

# %% [code]
print(f"Loading pairwise BLAST matrix ({BLAST_MATRIX_FILE})...")
blast_df = pd.read_parquet(DATA_DIR / BLAST_MATRIX_FILE)

print(f"Loading pairwise similarity matrix ({SIMILARITY_MATRIX_FILE})...")
similarity_df = pd.read_parquet(DATA_DIR / SIMILARITY_MATRIX_FILE)

# %% [code]
# to_drop = [col for col in similarity_df.columns if col not in ids]

# blast_df.drop(index=to_drop, columns=to_drop, inplace=True)
# similarity_df.drop(index=to_drop, columns=to_drop, inplace=True)

# idx = 1
# SIMILARITY_MATRIX_FILE = f"pairwise_similarity_{idx:03}.parquet"
# BLAST_MATRIX_FILE = f"pairwise_blast_{idx:03}.parquet"

# blast_df.sort_index(axis=0, inplace=True)
# blast_df.sort_index(axis=1, inplace=True)

# similarity_df.sort_index(axis=0, inplace=True)
# similarity_df.sort_index(axis=1, inplace=True)

# print(f"Writing pairwise BLAST matrix ({BLAST_MATRIX_FILE})...")
# blast_df.to_parquet(DATA_DIR / BLAST_MATRIX_FILE)

# print(f"Writing pairwise similarity matrix ({SIMILARITY_MATRIX_FILE})...")
# similarity_df.to_parquet(DATA_DIR / SIMILARITY_MATRIX_FILE)

# %% [code]
def select_proteins_mmr(
    quality_scores,
    cosine_matrix,
    blast_matrix,
    n_select=96,
    lambda_param=0.5,
    blast_threshold=80,
    preselected_indices=[],
):
    # Initialization: Pick the absolute best quality not in to start
    if preselected_indices == []:
        first_choice = np.argmax(quality_scores)
        selected_indices = [first_choice]
        preselected_indices.append(first_choice)
    else:
        selected_indices = []

    n_select -= len(selected_indices)
    print(n_select)

    # Iterative Selection
    for _ in tqdm(range(n_select)):
        remaining_indices = [i for i in range(len(quality_scores)) if i not in preselected_indices]
        candidate_to_add = -1

        # Shape: (Num_Remaining, Num_Selected)
        sim_to_selected = cosine_matrix[remaining_indices][:, preselected_indices]

        # Calculate max similarity to the selected set for each remaining item
        max_sim_vals = np.max(sim_to_selected, axis=1)

        # Calculate MMR scores for all remaining items
        # mmr = lambda * Quality - (1-lambda) * Max_Sim
        current_qualities = quality_scores[remaining_indices]
        mmr_scores = (lambda_param * current_qualities) - ((1 - lambda_param) * max_sim_vals)

        # Sort candidates by MMR score descending
        sorted_candidate_args = np.argsort(mmr_scores)[::-1]

        # Apply BLAST Hard Constraint
        # We iterate down the ranked list and pick the first one that passes the BLAST check
        found_next = False
        while not found_next:
            for local_idx in sorted_candidate_args:
                global_idx = remaining_indices[local_idx]

                # Check BLAST against ALL currently selected
                blast_scores = blast_matrix[global_idx, preselected_indices]

                if np.all(blast_scores <= blast_threshold):
                    candidate_to_add = global_idx
                    found_next = True
                    break

            if found_next:
                selected_indices.append(candidate_to_add)
                preselected_indices.append(candidate_to_add)
            else:
                blast_threshold += 5
                print(f"Warning: Could not find enough candidates satisfying BLAST constraint, increasing threshold to {blast_threshold}.")

    return selected_indices

# %% [code]
def select_proteins_mmr_fast(
    quality_scores,
    cosine_matrix,
    blast_matrix,
    inter_scores,
    beta=0.1,
    n_select=96,
    lambda_param=0.5,
    blast_threshold=80,
    preselected_indices=None,
):
    N = len(quality_scores)

    is_selected = np.zeros(N, dtype=bool)

    blast_violation_counts = np.zeros(N, dtype=int)
    current_max_sim = np.full(N, -1.0, dtype=np.float32)

    selected_indices_list = []

    # Handle Pre-selection
    if preselected_indices:
        for idx in preselected_indices:
            is_selected[idx] = True
            selected_indices_list.append(idx)
            # Update max sim state
            current_max_sim = np.maximum(current_max_sim, cosine_matrix[idx])
            # Update BLAST blocks
            bad_blast = blast_matrix[idx] > blast_threshold
            blast_violation_counts[bad_blast] += 1

    else:
        first_choice = np.argmax(quality_scores)
        is_selected[first_choice] = True
        selected_indices_list.append(first_choice)

        current_max_sim = np.maximum(current_max_sim, cosine_matrix[first_choice])

        bad_blast = blast_matrix[first_choice] > blast_threshold
        blast_violation_counts[bad_blast] += 1

    n_remaining_to_select = n_select - len(selected_indices_list)
    print(f"Selecting {n_remaining_to_select} more proteins...")

    for _ in tqdm(range(n_remaining_to_select)):
        candidate_found = False

        while not candidate_found:
            valid_candidates_mask = (~is_selected) & (blast_violation_counts == 0)

            if not np.any(valid_candidates_mask):
                blast_threshold += 5
                print(f"Warning: Relaxing BLAST threshold to {blast_threshold}...")

                blast_violation_counts[:] = 0
                for sel_idx in selected_indices_list:
                    bad_blast = blast_matrix[sel_idx] > blast_threshold
                    blast_violation_counts[bad_blast] += 1
                continue

            mmr_scores = (lambda_param * quality_scores) - ((1 - lambda_param) * current_max_sim) + beta * inter_scores
            mmr_scores[~valid_candidates_mask] = -np.inf

            best_idx = np.argmax(mmr_scores)

            selected_indices_list.append(best_idx)
            is_selected[best_idx] = True

            current_max_sim = np.maximum(current_max_sim, cosine_matrix[best_idx])
            bad_blast = blast_matrix[best_idx] > blast_threshold
            blast_violation_counts[bad_blast] += 1

            candidate_found = True

    return selected_indices_list

# %% [code]
l_min = np.min(quality)
l_max = np.max(quality)

# Avoid division by zero if all scores are identical
if l_max - l_min == 0:
    norm_quality = np.ones_like(quality)
else:
    # Linear inversion: (Max - Current) / Range
    norm_quality = (l_max - quality) / (l_max - l_min)

# %% [code]
plt.figure(figsize=(6,4))
plot_distribution(
    plt.gca(),
    {"norm_quality": norm_quality},
    "norm_quality",
)
plt.show()

# %% [code]
inter_scores = df_main["inter_score"].to_numpy()

l_min = np.min(inter_scores)
l_max = np.max(inter_scores)

if l_max - l_min == 0:
    norm_inter_scores = np.ones_like(inter_scores)
else:
    norm_inter_scores = (inter_scores - l_min) / (l_max - l_min)

# %% [code]
plt.figure(figsize=(6, 4))
plot_distribution(
    plt.gca(),
    {"norm_inter_scores": norm_inter_scores},
    "norm_inter_scores",
)
plt.show()

# %% [code]
assert (ids == similarity_df.index).all()
assert (ids == blast_df.index).all()

# %% [code]
cosine_matrix = similarity_df.to_numpy()
blast_matrix = blast_df.to_numpy()
norm_quality = norm_quality.to_numpy()

# %% [code]
blast_threshold = 50

lambda_param = 0.1
# for lambda_param in np.arange(0.1, 1, 0.05):
for beta in np.arange(0.01, 0.11, 0.01):
    selected_indices = select_proteins_mmr_fast(
        norm_quality,
        cosine_matrix=cosine_matrix,
        blast_matrix=blast_matrix,
        inter_scores=norm_inter_scores,
        beta=beta,
        lambda_param=lambda_param,
        blast_threshold=blast_threshold,
    )

    selected_uniprot_ids = [ids[idx] for idx in selected_indices]

    output_file = f"{LOG_DIR}/selected_uniprot_ids_mmr_lambda_{lambda_param:.2f}_{blast_threshold}_beta_{beta:.2f}.txt"

    with open(output_file, "w") as f:
        for uid in selected_uniprot_ids:
            f.write(f"{uid}\n")

# %% [code]
blast_df_selected = blast_df[selected_uniprot_ids]

# %% [code]
blast_df_selected = blast_df_selected.loc[selected_uniprot_ids]

# %% [code]
similarity_df_selected = similarity_df.loc[selected_uniprot_ids]

# %% [code]
similarity_df_selected = similarity_df_selected[selected_uniprot_ids]

# %% [code]
similarity_df_selected

# %% [code]
blast_df_selected

# %% [code]
pairs = [(id1, id2) for id1 in blast_df_selected.index for id2 in blast_df_selected.columns if id1 < id2]

row_indices = [p[0] for p in pairs]
col_indices = [p[1] for p in pairs]
x_values = blast_df.loc[row_indices, col_indices]
x_values = x_values.values.flatten()

# %% [code]
plt.figure(figsize=(6, 4))
ax = plt.gca()

plt.hist(
    x_values,
    bins=50,
    density=True,
    alpha=0.75,
)

ax.set_title(f"Distribution of Pairwise Similarity")
ax.set_xlabel("Pairwise Similarity")
ax.set_ylabel("Probability (%)")
ax.tick_params(left=False)
ax.spines[["top", "right", "left"]].set_visible(False)
ax.spines["bottom"].set_linewidth(2)
ax.yaxis.grid(True, which="major", linestyle=":")
ax.set_axisbelow(True)

plt.tight_layout()
plt.show()

# %% [code]
df_selected

# %% [code]
```
