## Supplementary material for "pLM representations unlock metagenomic space beyond homology": Code bundle: metagenomic.ipynb

```
# %% [code]

from pathlib import Path
from functools import reduce

import random
import os
import math
import numpy as np
import pandas as pd
import json
import polars as pl

from scipy.stats import pearsonr, spearmanr
from scipy.spatial.distance import pdist, squareform
from sklearn.metrics.pairwise import cosine_similarity

import matplotlib.pyplot as plt
import seaborn as sns
from umap import UMAP

from scipy.sparse import csr_matrix, save_npz, load_npz
from pandas.api.types import CategoricalDtype

from utils import merge_dataframes, plot_distribution
%config InlineBackend.figure_format='retina'

# %% [code]
DATA_DIR = Path("$SCRATCH/proseqo/enzymes/metagenomic_output/dataframes")
FIG_DIR = Path("$SCRATCH/proseqo/enzymes/metagenomic_output/figures")
FIG_DIR.mkdir(exist_ok=True)

# Add all your generated metrics filenames to this list.
# MERGED_FILE = None
MERGED_FILE = "merged_data.parquet"
SIMILARITY_MATRIX_FILE = f"pairwise_similarity.parquet"
IDENTITY_MATRIX_FILE = f"pairwise_identity.parquet"
BLAST_MATRIX_FILE = f"pairwise_blast.parquet"

DROP_EMBEDDINGS = False

if MERGED_FILE is None:
    df = pd.read_parquet(DATA_DIR / "metagenomic_candidates.parquet")
    df = df.reset_index(drop=False)

    df2 = pd.read_parquet(DATA_DIR / "physicochem_properties.parquet")
    df2 = df2.rename(columns={"uniprot_id": "metagenomic_id"})

    df3 = pd.read_parquet(DATA_DIR / "plm_perplexity_AMPLIFY-C.parquet")
    df3 = df3.rename(columns={"id": "metagenomic_id"})

    df = df.merge(df2, on="metagenomic_id", how="left")
    df = df.merge(df3, on="metagenomic_id", how="left")

    df = df.rename(columns={"length_x": "length"})
    df = df.drop(columns=["length_y"])
    df.to_parquet(DATA_DIR / "merged_data.parquet")

else:
    df = pd.read_parquet(DATA_DIR / MERGED_FILE)

if DROP_EMBEDDINGS == True:
    print("Dropping embedding columns to reduce dataframe size...")
    df = df.drop(columns=[c for c in df.columns if "embedding" in c])

# %% [code]
tmp = df[df["sequence_metagenomic"].str.startswith("M")]
tmp = tmp[tmp["blast"] < 50]
unique_ids = set(list(tmp["metagenomic_id"].unique()))

# %% [code]
tmp.to_parquet("$SCRATCH/proseqo/enzymes/metagenomic_output/dataframes/merged_data_001.parquet")
# tmp = pd.read_parquet(DATA_DIR / "merged_data_001.parquet")

# %% [code]
unique_ids = set(list(tmp["metagenomic_id"].unique()))

# %% [code]
selected_file = "$HOME/enzymes/selection_logs/metagenomic_output/selected_uniprot_ids_mmr_lambda_0.10_50.txt"

with open(selected_file, "r") as f:
    # Read IDs, strip whitespace, and remove empty lines
    selected_ids = set(line.strip() for line in f if line.strip())

df_selected = df[df["metagenomic_id"].isin(selected_ids)]

# %% [markdown]
### Cosine sim

# %% [code]
emb = tmp[["metagenomic_id", "embedding"]]

# %% [code]
emb.to_parquet(
    "$SCRATCH/proseqo/enzymes/metagenomic_output/dataframes/plm_embeddings_AMPLIFY-C.parquet"
)

# %% [markdown]
### BLAST

# %% [code]
tsv1 = pd.read_csv(
    "$SCRATCH/proseqo/enzymes/metagenomic_output/metagenomic_candidates_blast_pairwise_similarity_1.tsv", sep="\t", header=None
)
tsv2 = pd.read_csv(
    "$SCRATCH/proseqo/enzymes/metagenomic_output/metagenomic_candidates_blast_pairwise_similarity_1.tsv",
    sep="\t",
    header=None,
)
tsv = pd.concat([tsv1, tsv2])
del tsv2, tsv1
tsv.columns = ["id1", "id2", "blast"]
tsv_inverse = tsv.rename(columns={"id1": "id2", "id2": "id1"})
all_tsv = pd.concat([tsv, tsv_inverse], ignore_index=True).drop_duplicates(subset=["id1", "id2"])
del tsv, tsv_inverse

# %% [code]
all_tsv = all_tsv[(all_tsv["id1"].isin(unique_ids)) & (all_tsv["id2"].isin(unique_ids))]

# %% [code]
id_type = CategoricalDtype(categories=unique_ids, ordered=True)
row = all_tsv["id1"].astype(id_type).cat.codes
col = all_tsv["id2"].astype(id_type).cat.codes
sparse_matrix = csr_matrix((all_tsv["blast"], (row, col)), shape=(len(unique_ids), len(unique_ids)))
save_npz("$SCRATCH/proseqo/enzymes/metagenomic_output/dataframes/sparse_blast.npz", sparse_matrix)

# %% [code]
matrix = load_npz("$SCRATCH/proseqo/enzymes/metagenomic_output/dataframes/sparse_blast.npz")

# %% [code]
matrix = matrix.toarray()
blast_df = pd.DataFrame(matrix)
blast_df.columns = list(unique_ids)
blast_df.index = list(unique_ids)

# %% [code]
blast_df = blast_df.astype(np.float16)

# %% [code]
blast_df

# %% [code]
blast_df.to_parquet("$SCRATCH/proseqo/enzymes/metagenomic_output/dataframes/blast.parquet")
# blast_df = pd.read_parquet("$SCRATCH/proseqo/enzymes/metagenomic_output/dataframes/blast.parquet")

# %% [code]
blast_df_sub = blast_df.head(1000)
blast_df_sub = blast_df_sub.iloc[:, :1000]
pairs = [(id1, id2) for id1 in blast_df_sub.index for id2 in blast_df_sub.columns if id1 < id2]

sub_pairs = random.sample(pairs, 10000)
row_indices = [p[0] for p in sub_pairs]
col_indices = [p[1] for p in sub_pairs]
x_values = blast_df.loc[row_indices, col_indices]
x_values = x_values.values.flatten()

# %% [code]
plt.figure(figsize=(6, 4))
ax = plt.gca()

plt.hist(
    x_values,
    bins=50,
    density=True,
    alpha=0.75,
)

ax.set_title(f"Distribution of Pairwise Similarity")
ax.set_xlabel("Pairwise Similarity")
ax.set_ylabel("Probability (%)")
ax.tick_params(left=False)
ax.spines[["top", "right", "left"]].set_visible(False)
ax.spines["bottom"].set_linewidth(2)
ax.yaxis.grid(True, which="major", linestyle=":")
ax.set_axisbelow(True)

plt.tight_layout()
plt.show()

# %% [code]
non_zero = x_values[x_values > 0]
plt.figure(figsize=(6, 4))
ax = plt.gca()

plt.hist(
    non_zero,
    bins=50,
    density=True,
    alpha=0.75,
)

ax.set_title(f"Distribution of Pairwise Identity")
ax.set_xlabel("Pairwise Similarity")
ax.set_ylabel("Probability (%)")
ax.tick_params(left=False)
ax.spines[["top", "right", "left"]].set_visible(False)
ax.spines["bottom"].set_linewidth(2)
ax.yaxis.grid(True, which="major", linestyle=":")
ax.set_axisbelow(True)

plt.tight_layout()
plt.show()

# %% [markdown]
## Plots

# %% [code]
def plot_correlation(
    df,
    metric1,
    metric2,
    color_metric,
    remove_outliers=False,
    min_correlation=None,
    figsize=(7, 6),
    ax=None,
    vmax_color=None,
    cmap="viridis",
    lambda_param=None,
    method=None,
):
    # Check if all required columns exist before plotting
    required_cols = [metric1, metric2, color_metric]
    if all(col in df.columns for col in required_cols):
        plot_data = df[[metric1, metric2, color_metric]].dropna()

        if not plot_data.empty:
            p_corr, _ = pearsonr(plot_data[metric1], plot_data[metric2])
            s_corr, _ = spearmanr(plot_data[metric1], plot_data[metric2])

            # Early exit if correlation is too low
            if min_correlation is not None and (abs(p_corr) < min_correlation and abs(s_corr) < min_correlation):
                return None, None

            if ax is None:
                fig, ax = plt.subplots(figsize=figsize)
                is_standalone = True
                title_fs, label_fs = 12, 10
            else:
                fig = ax.get_figure()  # Get the parent figure of the passed axis
                is_standalone = False
                title_fs, label_fs = 12, 10

            points = ax.scatter(
                plot_data[metric1],
                plot_data[metric2],
                c=plot_data[color_metric].values,
                cmap=cmap,
                s=1,
                alpha=0.9,
                vmax=vmax_color,
            )

            if remove_outliers:
                x_min, x_max = plot_data[metric1].quantile(0.001), plot_data[metric1].quantile(0.999)
                y_min, y_max = plot_data[metric2].quantile(0.001), plot_data[metric2].quantile(0.999)
                ax.set_xlim(x_min, x_max)
                ax.set_ylim(y_min, y_max)

            cbar = fig.colorbar(points, ax=ax)
            cbar.set_label(color_metric, fontsize=label_fs)

            if is_standalone:
                title = f"{metric1} vs. {metric2}\nPearson r: {p_corr:.3f} | Spearman ρ: {s_corr:.3f}"
                if method is not None and lambda_param is not None:
                    title = f"{method}, Lambda={lambda_param:.2f}\n" + title
                ax.set_title(title)

            ax.set_xlabel(metric1, fontsize=label_fs)
            ax.set_ylabel(metric2, fontsize=label_fs)

            ax.tick_params(axis="both", which="major", labelsize=label_fs)

            ax.spines[["top", "right"]].set_visible(False)
            ax.spines[["left", "bottom"]].set_linewidth(2)

            if is_standalone:
                plt.tight_layout()
                if method is not None and lambda_param is not None:
                    plt.savefig(f"figures/{metric1}_{metric2}_{method}_lambda_{lambda_param:.2f}.png", dpi=600)

            return p_corr, s_corr
        else:
            print(f"Skipping: No data for {required_cols}")
    else:
        print(f"Skipping: Missing columns {required_cols}")

# %% [code]
def plot_properties(df_main, df_selected, properties_to_plot, lambda_param=None, method=None, filename=None):

    crop_ranges = [prop for prop in properties_to_plot.keys() if "ppl" in prop or "shannon" in prop or "pH" in prop]

    # Filter list to only properties that exist in the final dataframe
    properties_to_plot = {p: label for p, label in properties_to_plot.items() if p in df_main.columns}

    n_props = len(properties_to_plot)
    n_cols = 4
    n_rows = int(np.ceil(n_props / n_cols))
    num_bins = 30

    fig, axes = plt.subplots(n_rows, n_cols, figsize=(18, n_rows * 4.5), tight_layout=True)

    for i, prop in enumerate(properties_to_plot):
        ax = axes.flat[i]
        data_to_plot = df_main.dropna(subset=[prop])

        crop_range = prop in crop_ranges
        if "ppl" in prop:
            binrange = (0, data_to_plot[prop].quantile(0.99))
        elif "shannon" in prop or "pH" in prop:
            binrange = (data_to_plot[prop].quantile(0.001), data_to_plot[prop].quantile(0.999))
        else:
            binrange = None
        bin_edges = np.histogram_bin_edges(data_to_plot[prop], bins=num_bins, range=binrange)

        sns.histplot(
            data=data_to_plot,
            x=prop,
            alpha=0.4,
            lw=0.1,
            color="#0085ff",
            ax=ax,
            bins=bin_edges,
            stat="count",
        )

        ax2 = ax.twinx()
        ax2.hist(df_selected[prop], bins=bin_edges, color="red", alpha=0.6, label="Selected IDs")

        ax2.spines[["top", "left", "bottom"]].set_visible(False)
        ax2.spines["right"].set_visible(True)
        ax2.set_ylabel("Count (Selected)", color="red")
        ax2.tick_params(axis="y", labelcolor="red")

        # ax.set_title(f"Distribution of {prop}")
        ax.set_xlabel(properties_to_plot[prop])
        ax.set_ylabel("Count")
        ax.tick_params(left=False)
        ax.spines[["top", "right", "left"]].set_visible(False)
        ax.spines["bottom"].set_linewidth(2)
        ax.yaxis.grid(True, which="major", linestyle=":")
        ax.set_axisbelow(True)

    # Hide any unused subplots
    for j in range(i + 1, n_rows * n_cols):
        axes.flat[j].set_visible(False)

    plt.tight_layout()
    if lambda_param is not None and method is not None:
        plt.savefig(f"figures/properties_{method}_lambda_{lambda_param:.2f}.png", dpi=600)
    elif filename is not None:
        plt.savefig(filename, dpi=600)

# %% [code]
# df_selected.to_parquet("$HOME/enzymes/selection_logs/final/data_metagenomic.parquet")
df_selected = pd.read_parquet("$HOME/enzymes/selection_logs/final/data_metagenomic.parquet")

# %% [code]
df_selected.columns

# %% [code]
properties_to_plot = {
    "length": "Sequence length",
    "shannon_entropy_normalized": "Normalized shannon entropy",
    "AMPLIFY-C_pseudo_ppl": "Pseudo-perplexity",
    "intra_score": "RED score",
    "inter_score": "Embedding similarity to closest EC 1.14.11 enzyme",
}

# %% [code]
df["AMPLIFY-C_pseudo_ppl"].isna().sum()

# %% [code]
FIG_DIR

# %% [code]
plot_properties(df, df_selected, properties_to_plot, filename=None) # "figures/plate3_properties_selected_vs_all.png")

# %% [code]
properties_to_plot

# %% [code]
# Filter list to only properties that exist in the final dataframe
properties_to_plot = [
    "length",
    "inter_score",
    "mean_sasa",
    "mpnn_ll_soluble",
    "esm_if_nll",
    "AMPLIFY-C_pseudo_ppl",
]

tmp = df_selected

properties_to_plot = [prop for prop in properties_to_plot if prop in tmp.columns]

n_props = len(properties_to_plot)
n_cols = 3
n_rows = int(np.ceil(n_props / n_cols))

fig, axes = plt.subplots(n_rows, n_cols, figsize=(18, n_rows * 4.5), tight_layout=True)

for i, prop in enumerate(properties_to_plot):
    ax = axes.flat[i]
    data_to_plot = tmp.dropna(subset=[prop])

    sns.histplot(
        data=data_to_plot,
        x=prop,
        bins=30,
        stat="percent",
        alpha=0.4,
        lw=0.1,
        color="#0085ff",
        ax=ax,
    )

    ax.set_title(f"Distribution of {prop}")
    ax.set_xlabel(prop)
    ax.set_ylabel("Probability (%)")
    ax.tick_params(left=False)
    ax.spines[["top", "right", "left"]].set_visible(False)
    ax.spines["bottom"].set_linewidth(2)
    ax.yaxis.grid(True, which="major", linestyle=":")
    ax.set_axisbelow(True)

# Hide any unused subplots
for j in range(i + 1, n_rows * n_cols):
    axes.flat[j].set_visible(False)

plt.tight_layout()
plt.show()

# %% [code]
df_selected["lambda"].value_counts()

# %% [code]
df_selected["dataset"].value_counts()

# %% [code]
# Filter list to only properties that exist in the final dataframe
properties_to_plot = [
    "length",
    "inter_score",
    "blast",
    "molecular_weight",
    "isoelectric_point",
    "gravy",
    "instability_index",
    "aromaticity",
    "sa_repeat_ratio",
    "ss_helix",
    "ss_turn",
    "ss_sheet",
    "shannon_entropy",
    "shannon_entropy_normalized",
    "charge_at_pH7.5",
    "flexibility_mean",
    "mean_sasa",
    "deeptmhmm_sp",
    "deeptmhmm_tmalpha",
    "deeptmhmm_tmbeta",
    "mpnn_ll_vanilla",
    "mpnn_ll_soluble",
    "esm_if_nll",
    "intra_score",
    "AMPLIFY-C_pseudo_ppl",
]

properties_to_plot = [prop for prop in properties_to_plot if prop in df.columns]

n_props = len(properties_to_plot)
n_cols = 3
n_rows = int(np.ceil(n_props / n_cols))

fig, axes = plt.subplots(n_rows, n_cols, figsize=(18, n_rows * 4.5), tight_layout=True)

for i, prop in enumerate(properties_to_plot):
    ax = axes.flat[i]
    data_to_plot = df.dropna(subset=[prop])

    sns.histplot(
        data=data_to_plot,
        x=prop,
        bins=30,
        stat="percent",
        alpha=0.4,
        lw=0.1,
        color="#0085ff",
        ax=ax,
    )

    ax.set_title(f"Distribution of {prop}")
    ax.set_xlabel(prop)
    ax.set_ylabel("Probability (%)")
    ax.tick_params(left=False)
    ax.spines[["top", "right", "left"]].set_visible(False)
    ax.spines["bottom"].set_linewidth(2)
    ax.yaxis.grid(True, which="major", linestyle=":")
    ax.set_axisbelow(True)

# Hide any unused subplots
for j in range(i + 1, n_rows * n_cols):
    axes.flat[j].set_visible(False)

plt.tight_layout()
plt.show()

# %% [code]
df.columns

# %% [code]
# Filter list to only properties that exist in the final dataframe
properties_to_plot = [
    "length",
    "inter_score",
    "blast",
    "molecular_weight",
    "isoelectric_point",
    "gravy",
    "instability_index",
    "aromaticity",
    "sa_repeat_ratio",
    "ss_helix",
    "ss_turn",
    "ss_sheet",
    "shannon_entropy",
    "shannon_entropy_normalized",
    "charge_at_pH7.5",
    "flexibility_mean",
    "mean_sasa",
    "deeptmhmm_sp",
    "deeptmhmm_tmalpha",
    "deeptmhmm_tmbeta",
    "mpnn_ll_vanilla",
    "mpnn_ll_soluble",
    "esm_if_nll",
    "intra_score",
    "AMPLIFY-C_pseudo_ppl",
]

properties_to_plot = [prop for prop in properties_to_plot if prop in tmp.columns]

n_props = len(properties_to_plot)
n_cols = 3
n_rows = int(np.ceil(n_props / n_cols))

fig, axes = plt.subplots(n_rows, n_cols, figsize=(18, n_rows * 4.5), tight_layout=True)

for i, prop in enumerate(properties_to_plot):
    ax = axes.flat[i]
    data_to_plot = tmp.dropna(subset=[prop])

    sns.histplot(
        data=data_to_plot,
        x=prop,
        bins=30,
        stat="percent",
        alpha=0.4,
        lw=0.1,
        color="#0085ff",
        ax=ax,
    )

    ax.set_title(f"Distribution of {prop}")
    ax.set_xlabel(prop)
    ax.set_ylabel("Probability (%)")
    ax.tick_params(left=False)
    ax.spines[["top", "right", "left"]].set_visible(False)
    ax.spines["bottom"].set_linewidth(2)
    ax.yaxis.grid(True, which="major", linestyle=":")
    ax.set_axisbelow(True)

# Hide any unused subplots
for j in range(i + 1, n_rows * n_cols):
    axes.flat[j].set_visible(False)

plt.tight_layout()
plt.show()

# %% [code]
tmp = tmp.sort_values("inter_score", ascending=False)

# %% [code]
tmp.columns

# %% [markdown]
## Properties

# %% [markdown]
### BFD

# %% [code]
DROP_EMBEDDINGS = True

# Define the directory where your results are stored
DATA_DIR = Path("$SCRATCH/proseqo/enzymes/bfd_output/dataframes")
FIG_DIR = Path("$SCRATCH/proseqo/enzymes/bfd_output/figures")
FIG_DIR.mkdir(exist_ok=True)
MERGED_FILE = None
KEY_COL = "uniprot_id"

if MERGED_FILE is None:
    # Add all your generated metrics filenames to this list.
    FILES_TO_MERGE = [
        "physicochem_properties.parquet",
        "structural_properties.parquet",
        # "signalp_scores.parquet",
        # "plm_embeddings_AMPLIFY_350M.parquet",
        # "plm_embeddings_AMPLIFY-C.parquet",
        # "plm_embeddings_AMPLIFY-C.parquet",
        # "plm_perplexity_AMPLIFY-C.parquet",
        # "plm_perplexity_AMPLIFY-C.parquet",
        # "proteinmpnn_scores_soluble.parquet",
        # "proteinmpnn_scores_vanilla.parquet",
        # "esm_if_scores.parquet",
    ]

    # Pairwise identity matrix (handled separately)
    IDENTITY_MATRIX_FILE = "pairwise_identity.parquet"

    print("Loading metrics files...")
    all_dfs = [pd.read_parquet(DATA_DIR / file) for file in FILES_TO_MERGE]

    print(f"Merging {len(all_dfs)} dataframes on key '{KEY_COL}'...")
    df_main = merge_dataframes(all_dfs, KEY_COL)

    # print(f"Loading pairwise identity matrix ({IDENTITY_MATRIX_FILE})...")
    # identity_df = pd.read_parquet(DATA_DIR / IDENTITY_MATRIX_FILE)

else:

    print("Loading metrics files...")
    df_main = pd.read_parquet(DATA_DIR / MERGED_FILE)

if DROP_EMBEDDINGS == True:
    print("Dropping embedding columns to reduce dataframe size...")
    df_main = df_main.drop(columns=[c for c in df_main.columns if "embedding" in c])

# %% [code]
len(df_main)

# %% [code]
crop_ranges = [prop for prop in properties_to_plot if "ppl" in prop or "pH" in prop]

# Filter list to only properties that exist in the final dataframe
properties_to_plot = [p for p in properties_to_plot if p in df_main.columns]

n_props = len(properties_to_plot)
n_cols = 3
n_rows = int(np.ceil(n_props / n_cols))

fig, axes = plt.subplots(n_rows, n_cols, figsize=(18, n_rows * 4.5), tight_layout=True)

for i, prop in enumerate(properties_to_plot):
    ax = axes.flat[i]
    data_to_plot = df_main.dropna(subset=[prop])

    crop_range = prop in crop_ranges
    binrange = (data_to_plot[prop].quantile(0.01), data_to_plot[prop].quantile(0.99)) if crop_range else None

    sns.histplot(
        data=data_to_plot,
        x=prop,
        bins=30,
        stat="percent",
        alpha=0.4,
        lw=0.1,
        color="#0085ff",
        ax=ax,
        binrange=binrange,
    )

    # Plot vertical lines at min and max
    ax.axvline(data_to_plot[prop].min(), color="red", linestyle="--", lw=1)
    ax.axvline(data_to_plot[prop].max(), color="red", linestyle="--", lw=1)

    ax.set_title(f"Distribution of {prop}")
    ax.set_xlabel(prop)
    ax.set_ylabel("Probability (%)")
    ax.tick_params(left=False)
    ax.spines[["top", "right", "left"]].set_visible(False)
    ax.spines["bottom"].set_linewidth(2)
    ax.yaxis.grid(True, which="major", linestyle=":")
    ax.set_axisbelow(True)

# Hide any unused subplots
for j in range(i + 1, n_rows * n_cols):
    axes.flat[j].set_visible(False)

plt.tight_layout()
plt.show()

# %% [markdown]
### MGnify

# %% [code]
DROP_EMBEDDINGS = True

# Define the directory where your results are stored
DATA_DIR = Path("$SCRATCH/proseqo/enzymes/mgnify_output/dataframes")
FIG_DIR = Path("$SCRATCH/proseqo/enzymes/mgnify_output/figures")
FIG_DIR.mkdir(exist_ok=True)
MERGED_FILE = None
KEY_COL = "uniprot_id"

if MERGED_FILE is None:
    # Add all your generated metrics filenames to this list.
    FILES_TO_MERGE = [
        "physicochem_properties.parquet",
        "structural_properties.parquet",
        # "signalp_scores.parquet",
        # "plm_embeddings_AMPLIFY_350M.parquet",
        # "plm_embeddings_AMPLIFY-C.parquet",
        # "plm_embeddings_AMPLIFY-C.parquet",
        # "plm_perplexity_AMPLIFY-C.parquet",
        # "plm_perplexity_AMPLIFY-C.parquet",
        # "proteinmpnn_scores_soluble.parquet",
        # "proteinmpnn_scores_vanilla.parquet",
        # "esm_if_scores.parquet",
    ]

    # Pairwise identity matrix (handled separately)
    IDENTITY_MATRIX_FILE = "pairwise_identity.parquet"

    print("Loading metrics files...")
    all_dfs = [pd.read_parquet(DATA_DIR / file) for file in FILES_TO_MERGE]

    print(f"Merging {len(all_dfs)} dataframes on key '{KEY_COL}'...")
    df_main = merge_dataframes(all_dfs, KEY_COL)

    # print(f"Loading pairwise identity matrix ({IDENTITY_MATRIX_FILE})...")
    # identity_df = pd.read_parquet(DATA_DIR / IDENTITY_MATRIX_FILE)

else:

    print("Loading metrics files...")
    df_main = pd.read_parquet(DATA_DIR / MERGED_FILE)

if DROP_EMBEDDINGS == True:
    print("Dropping embedding columns to reduce dataframe size...")
    df_main = df_main.drop(columns=[c for c in df_main.columns if "embedding" in c])

# %% [code]


crop_ranges = [prop for prop in properties_to_plot if "ppl" in prop or "pH" in prop]

# Filter list to only properties that exist in the final dataframe
properties_to_plot = [p for p in properties_to_plot if p in df_main.columns]

n_props = len(properties_to_plot)
n_cols = 3
n_rows = int(np.ceil(n_props / n_cols))

fig, axes = plt.subplots(n_rows, n_cols, figsize=(18, n_rows * 4.5), tight_layout=True)

for i, prop in enumerate(properties_to_plot):
    ax = axes.flat[i]
    data_to_plot = df_main.dropna(subset=[prop])

    crop_range = prop in crop_ranges
    binrange = (data_to_plot[prop].quantile(0.01), data_to_plot[prop].quantile(0.99)) if crop_range else None

    sns.histplot(
        data=data_to_plot,
        x=prop,
        bins=30,
        stat="percent",
        alpha=0.4,
        lw=0.1,
        color="#0085ff",
        ax=ax,
        binrange=binrange,
    )

    # Plot vertical lines at min and max
    ax.axvline(data_to_plot[prop].min(), color="red", linestyle="--", lw=1)
    ax.axvline(data_to_plot[prop].max(), color="red", linestyle="--", lw=1)

    ax.set_title(f"Distribution of {prop}")
    ax.set_xlabel(prop)
    ax.set_ylabel("Probability (%)")
    ax.tick_params(left=False)
    ax.spines[["top", "right", "left"]].set_visible(False)
    ax.spines["bottom"].set_linewidth(2)
    ax.yaxis.grid(True, which="major", linestyle=":")
    ax.set_axisbelow(True)

# Hide any unused subplots
for j in range(i + 1, n_rows * n_cols):
    axes.flat[j].set_visible(False)

plt.tight_layout()
plt.show()

# %% [code]
len(df_main)

# %% [code]
```
