## Supplementary material for "pLM representations unlock metagenomic space beyond homology": Code bundle: mlp.ipynb

```
# %% [markdown]
# Enzyme expression prediction: classifier benchmark

Binary classification (zero vs. non-zero expression) across:
- **Methods**: Logistic Regression, LightGBM, MLP
- **Features**: AMPLIFY_c embedding, biophysical, structural, biophysical+structural
- **Tasks**: `lysate band`, `purified band`
- **Regimes**: full data (standard), low-data (n=50, 100 × 5 seeds)

# %% [code]
# ── Imports & constants ────────────────────────────────────────────────────────
import json
import shutil
import warnings
from pathlib import Path

import joblib
import lightgbm as lgb
import numpy as np
import optuna
import pandas as pd
import matplotlib.pyplot as plt
import seaborn as sns
import torch
import torch.nn as nn
import torch.optim as optim
from sklearn.linear_model import LogisticRegression
from sklearn.metrics import roc_auc_score, roc_curve
from sklearn.model_selection import StratifiedKFold, train_test_split
from sklearn.preprocessing import StandardScaler
from torch.utils.data import DataLoader, TensorDataset

optuna.logging.set_verbosity(optuna.logging.WARNING)
warnings.filterwarnings('ignore')
%config InlineBackend.figure_format='retina'

# ── Paths ──────────────────────────────────────────────────────────────────────
DATA_PATH = Path('$HOME/enzymes/notebooks/data.csv')
FIG_DIR   = Path('$HOME/enzymes/notebooks/figures')
OUT_DIR   = Path('$SCRATCH/proseqo/enzymes/mlp')
STD_DIR      = OUT_DIR / 'standard'
COMPLETE_DIR = OUT_DIR / 'complete'
LOW_DIR      = OUT_DIR / 'low_data'
for d in [FIG_DIR, STD_DIR, COMPLETE_DIR, LOW_DIR]:
    d.mkdir(parents=True, exist_ok=True)

DEVICE = torch.device('cuda' if torch.cuda.is_available() else 'cpu')
print(f'Device: {DEVICE}')

# ── Feature column definitions ─────────────────────────────────────────────────
BIOPHYS_COLS = [
    'aromaticity', 'charge_at_pH7', 'charge_at_pH7.5', 'charge_at_pH8',
    'flexibility_mean', 'fraction_disordered_05', 'fraction_disordered_08',
    'fraction_disordered_09', 'gravy', 'instability_index', 'isoelectric_point',
    'mol_ext_oxidized', 'mol_ext_reduced', 'molecular_weight',
    'non_standard_aa_count', 'sa_repeat_ratio', 'shannon_entropy',
    'shannon_entropy_normalized', 'sp_likelihood', 'ss_helix', 'ss_sheet', 'ss_turn',
]
STRUCT_COLS         = ['esm_if_nll', 'mpnn_ll_soluble', 'mean_plddt', 'mean_sasa']
BIOPHYS_STRUCT_COLS = BIOPHYS_COLS + STRUCT_COLS

SINGLE_BIOPHYS_FEATURE_SETS = {
    f'biophys:{col}': ('tabular', [col]) for col in BIOPHYS_COLS
}
SINGLE_STRUCT_FEATURE_SETS = {
    f'struct:{col}': ('tabular', [col]) for col in STRUCT_COLS
}
SINGLE_FEATURE_KEYS = set(SINGLE_BIOPHYS_FEATURE_SETS) | set(SINGLE_STRUCT_FEATURE_SETS)

# kind: 'embedding' (stored as string, needs parsing) or 'tabular' (numeric)
FEATURE_SETS = {
    'amplify_c':      ('embedding', 'AMPLIFY-C_embedding'),
    'biophys':        ('tabular',   BIOPHYS_COLS),
    'struct':         ('tabular',   STRUCT_COLS),
    'biophys+struct': ('tabular',   BIOPHYS_STRUCT_COLS),
    **SINGLE_BIOPHYS_FEATURE_SETS,
    **SINGLE_STRUCT_FEATURE_SETS,
}

TASKS       = ['lysate', 'purified']
TARGET_COLS = {'lysate': 'lysate band', 'purified': 'purified band'}

# ── Training hyperparams ───────────────────────────────────────────────────────
N_EPOCHS      = 150
N_CV_FOLDS    = 5
N_TRIALS_EMB  = 50   # MLP on high-dim embeddings
N_TRIALS_TAB  = 50   # MLP / LGB on tabular features
N_TRIALS_LOW  = 20   # fewer trials in low-data regime

EVAL_SEEDS = list(range(20))  # used for all three regimes

# %% [code]
# ── Data loading ───────────────────────────────────────────────────────────────
data = pd.read_csv(DATA_PATH)
print(f'Loaded {len(data)} rows, {len(data.columns)} columns')

# Pre-parse embeddings once (space-separated vs comma-separated)
_emb_cache = {
    'AMPLIFY-C_embedding': np.stack(
        data['AMPLIFY-C_embedding'].map(
            lambda s: np.fromstring(s.strip('[]'), sep=' ').astype(np.float32))),
}
for col, arr in _emb_cache.items():
    print(f'  {col}: {arr.shape}')

# %% [code]
# ── Dataset builder ────────────────────────────────────────────────────────────

# Rows where ALL tabular features are present — defines the 'complete' pool
_COMPLETE_MASK = data[BIOPHYS_STRUCT_COLS].notna().all(axis=1)
print(f"Complete pool: {_COMPLETE_MASK.sum()} rows  |  Full pool: {len(data)} rows")

def build_dataset(feature_key: str, task: str, seed: int = 42,
                  low_data: bool = False, complete_only: bool = False) -> dict:
    """
    Build a stratified train/test split for (feature_key, task).

    Split ratios (applied to the available pool for this feature set)
    ────────────────────────────────────────────────────────────────
    standard / complete  (low_data=False): 85% train / 15% test
    low-data             (low_data=True):  15% train / 85% test

    Both ratios are computed over the full pool, so n_train and n_test
    are determined by the data, not hardcoded.

    Comparability
    ─────────────
    complete_only=True  → all feature sets use the same pool (~287 rows with
      full biophys+struct coverage) → identical indices across feature sets
      for any given (task, seed).
    complete_only=False → embeddings use the full pool; tabular features are
      always restricted to the complete pool.
    """
    kind, spec = FEATURE_SETS[feature_key]
    y_raw = data[TARGET_COLS[task]]

    if kind == 'tabular' or complete_only:
        mask = y_raw.notna() & _COMPLETE_MASK
    else:
        mask = y_raw.notna()

    X = _emb_cache[spec][mask.values] if kind == 'embedding' \
        else data.loc[mask, spec].values.astype(float)
    y     = y_raw[mask].values.astype(float)
    strat = (y > 0).astype(int)

    test_size = 0.85 if low_data else 0.15
    X_tr, X_te, y_tr, y_te = train_test_split(
        X, y, test_size=test_size, random_state=seed, stratify=strat
    )
    scaler = StandardScaler()
    return dict(
        X_train=scaler.fit_transform(X_tr), X_test=scaler.transform(X_te),
        y_train=y_tr, y_test=y_te, in_dim=X_tr.shape[1],
    )


# %% [code]
# ── MLP architecture & training helpers ────────────────────────────────────────
def build_mlp(in_dim, out_dim, hidden_sizes, dropout):
    layers, prev = [], in_dim
    for h in hidden_sizes:
        layers += [nn.Linear(prev, h), nn.LayerNorm(h), nn.ReLU(), nn.Dropout(dropout)]
        prev = h
    layers.append(nn.Linear(prev, out_dim))
    return nn.Sequential(*layers)


@torch.no_grad()
def predict(model, X):
    model.eval()
    return model(torch.tensor(X, dtype=torch.float32).to(DEVICE)).squeeze(-1).cpu().numpy()


def fit_model(model, X, y, criterion, n_epochs=N_EPOCHS, batch_size=64, lr=1e-3, weight_decay=1e-3):
    loader = DataLoader(TensorDataset(torch.tensor(X, dtype=torch.float32),
                                      torch.tensor(y, dtype=torch.float32)),
                        batch_size=batch_size, shuffle=True)
    opt   = optim.Adam(model.parameters(), lr=lr, weight_decay=weight_decay)
    sched = optim.lr_scheduler.CosineAnnealingLR(opt, T_max=n_epochs)
    for _ in range(n_epochs):
        model.train()
        for xb, yb in loader:
            xb, yb = xb.to(DEVICE), yb.to(DEVICE)
            opt.zero_grad(); criterion(model(xb).squeeze(-1), yb).backward(); opt.step()
        sched.step()
    return model


def _mlp_from_params(bp, in_dim):
    hidden = tuple(bp[f'h{i}'] for i in range(bp['n_layers']))
    return build_mlp(in_dim, 1, hidden, bp['dropout']).to(DEVICE)

# %% [code]
# ── Optuna objectives ──────────────────────────────────────────────────────────
def _n_cv_folds(n_train):
    """Adaptive CV: 3 folds when n_train ≤ 50, else 5. Prevents degenerate folds."""
    return 3 if n_train <= 50 else N_CV_FOLDS


def _mlp_clf_objective(trial, X_tr, y_bin, in_dim, splits):
    n_layers = trial.suggest_int('n_layers', 1, 3)
    hidden   = tuple(trial.suggest_categorical(f'h{i}', [64, 128, 256, 512]) for i in range(n_layers))
    dropout      = trial.suggest_float('dropout',      0.0,  0.5, step=0.1)
    lr           = trial.suggest_float('lr',           1e-4, 1e-2, log=True)
    weight_decay = trial.suggest_float('weight_decay', 1e-5, 1e-2, log=True)
    batch_size   = trial.suggest_categorical('batch_size', [32, 64, 128])
    aucs = []
    for tr_i, va_i in splits:
        m = build_mlp(in_dim, 1, hidden, dropout).to(DEVICE)
        fit_model(m, X_tr[tr_i], y_bin[tr_i].astype('float32'), nn.BCEWithLogitsLoss(),
                  n_epochs=N_EPOCHS, batch_size=batch_size, lr=lr, weight_decay=weight_decay)
        p = torch.sigmoid(torch.tensor(predict(m, X_tr[va_i]))).numpy()
        aucs.append(roc_auc_score(y_bin[va_i], p))
    return float(np.mean(aucs))


def _lgb_clf_objective(trial, X_tr, y_bin, splits, scale_pos):
    params = dict(
        n_estimators     = trial.suggest_int(  'n_estimators',    50,  500),
        max_depth        = trial.suggest_int(  'max_depth',        2,    8),
        learning_rate    = trial.suggest_float('learning_rate',  1e-3, 0.3, log=True),
        num_leaves       = trial.suggest_int(  'num_leaves',       8,   64),
        subsample        = trial.suggest_float('subsample',       0.5, 1.0),
        colsample_bytree = trial.suggest_float('colsample_bytree', 0.3, 1.0),
        reg_alpha        = trial.suggest_float('reg_alpha',      1e-5, 10.0, log=True),
        reg_lambda       = trial.suggest_float('reg_lambda',     1e-5, 10.0, log=True),
        scale_pos_weight=scale_pos, random_state=0, verbosity=-1,
    )
    aucs = []
    for tr_i, va_i in splits:
        m = lgb.LGBMClassifier(**params)
        m.fit(X_tr[tr_i], y_bin[tr_i])
        aucs.append(roc_auc_score(y_bin[va_i], m.predict_proba(X_tr[va_i])[:, 1]))
    return float(np.mean(aucs))


def _lr_cv_auc(X_tr, y_bin, C, splits):
    """Cross-validated AUC for logistic regression at a given C."""
    aucs = []
    for tr_i, va_i in splits:
        m = LogisticRegression(C=C, random_state=0, max_iter=1000)
        m.fit(X_tr[tr_i], y_bin[tr_i])
        aucs.append(roc_auc_score(y_bin[va_i], m.predict_proba(X_tr[va_i])[:, 1]))
    return float(np.mean(aucs))

# %% [code]
# ── Cache-aware train/load helpers ────────────────────────────────────────────
def _cache_paths(cache_dir, key, is_mlp):
    ext = 'model.pt' if is_mlp else 'model.joblib'
    return cache_dir / f'{key}_params.json', cache_dir / f'{key}_{ext}'


def _load_mlp(param_path, model_path, in_dim):
    with open(param_path) as f:
        bp = json.load(f)
    model = _mlp_from_params(bp, in_dim)
    model.load_state_dict(torch.load(model_path, map_location=DEVICE))
    return model, bp


def _run_optuna(objective_fn, n_trials):
    study = optuna.create_study(direction='maximize',
                                sampler=optuna.samplers.TPESampler(seed=42))
    study.optimize(objective_fn, n_trials=n_trials, show_progress_bar=True)
    return study


def train_or_load_mlp(cache_dir, key, X_tr, y_bin, in_dim, n_trials):
    pp, mp = _cache_paths(cache_dir, key, is_mlp=True)
    if pp.exists() and mp.exists():
        model, bp = _load_mlp(pp, mp, in_dim)
        return model, bp.get('best_cv_auc'), True
    cache_dir.mkdir(parents=True, exist_ok=True)
    n_folds = _n_cv_folds(len(X_tr))
    cv      = StratifiedKFold(n_splits=n_folds, shuffle=True, random_state=42)
    splits  = list(cv.split(X_tr, y_bin))
    study   = _run_optuna(
        lambda t: _mlp_clf_objective(t, X_tr, y_bin, in_dim, splits), n_trials)
    bp    = study.best_params
    model = _mlp_from_params(bp, in_dim)
    fit_model(model, X_tr, y_bin.astype('float32'), nn.BCEWithLogitsLoss(),
              n_epochs=N_EPOCHS, batch_size=bp['batch_size'],
              lr=bp['lr'], weight_decay=bp['weight_decay'])
    with open(pp, 'w') as f:
        json.dump({**bp, 'best_cv_auc': study.best_value, 'n_cv_folds': n_folds}, f, indent=2)
    torch.save(model.state_dict(), mp)
    return model, study.best_value, False


def train_or_load_lgb(cache_dir, key, X_tr, y_bin, n_trials):
    pp, mp = _cache_paths(cache_dir, key, is_mlp=False)
    if pp.exists() and mp.exists():
        with open(pp) as f: bp = json.load(f)
        return joblib.load(mp), bp.get('best_cv_auc'), True
    cache_dir.mkdir(parents=True, exist_ok=True)
    scale_pos = (y_bin == 0).sum() / max((y_bin == 1).sum(), 1)
    n_folds   = _n_cv_folds(len(X_tr))
    cv        = StratifiedKFold(n_splits=n_folds, shuffle=True, random_state=42)
    splits    = list(cv.split(X_tr, y_bin))
    study     = _run_optuna(
        lambda t: _lgb_clf_objective(t, X_tr, y_bin, splits, scale_pos), n_trials)
    bp    = study.best_params
    model = lgb.LGBMClassifier(**bp, scale_pos_weight=scale_pos, random_state=0, verbosity=-1)
    model.fit(X_tr, y_bin)
    with open(pp, 'w') as f:
        json.dump({**bp, 'scale_pos_weight': scale_pos,
                   'best_cv_auc': study.best_value, 'n_cv_folds': n_folds}, f, indent=2)
    joblib.dump(model, mp)
    return model, study.best_value, False


# C grid: log-spaced, covers under- and over-regularization for both low-dim and high-dim features
_LR_C_GRID = [1e-4, 1e-3, 1e-2, 0.1, 1.0, 10.0, 100.0]

def train_or_load_lr(cache_dir, key, X_tr, y_bin):
    pp, mp = _cache_paths(cache_dir, key, is_mlp=False)
    if pp.exists() and mp.exists():
        with open(pp) as f: bp = json.load(f)
        return joblib.load(mp), bp.get('best_cv_auc'), True
    cache_dir.mkdir(parents=True, exist_ok=True)
    n_folds = _n_cv_folds(len(X_tr))
    cv      = StratifiedKFold(n_splits=n_folds, shuffle=True, random_state=42)
    splits  = list(cv.split(X_tr, y_bin))
    best_C, best_auc = max(
        ((C, _lr_cv_auc(X_tr, y_bin, C, splits)) for C in _LR_C_GRID),
        key=lambda x: x[1]
    )
    model = LogisticRegression(C=best_C, random_state=0, max_iter=1000)
    model.fit(X_tr, y_bin)
    with open(pp, 'w') as f:
        json.dump({'method': 'lr', 'C': best_C,
                   'best_cv_auc': best_auc, 'n_cv_folds': n_folds}, f, indent=2)
    joblib.dump(model, mp)
    return model, best_auc, False


def evaluate(model, method, X_te, y_te):
    y_bin = (y_te > 0).astype(int)
    if method == 'mlp':
        p = torch.sigmoid(torch.tensor(predict(model, X_te))).numpy()
    else:
        p = model.predict_proba(X_te)[:, 1]
    return float(roc_auc_score(y_bin, p))


def bootstrap_auc(model, method, X_te, y_te, n_boot=1000, ci=0.95, seed=0):
    """Percentile bootstrap CI for test AUC. Returns (auc, lower, upper)."""
    rng   = np.random.default_rng(seed)
    y_bin = (y_te > 0).astype(int)
    if method == 'mlp':
        p = torch.sigmoid(torch.tensor(predict(model, X_te))).numpy()
    else:
        p = model.predict_proba(X_te)[:, 1]
    point = float(roc_auc_score(y_bin, p))
    aucs  = []
    for _ in range(n_boot):
        idx = rng.integers(0, len(y_bin), len(y_bin))
        if y_bin[idx].nunique() if hasattr(y_bin[idx], 'nunique') else len(np.unique(y_bin[idx])) < 2:
            continue
        aucs.append(roc_auc_score(y_bin[idx], p[idx]))
    alpha = (1 - ci) / 2
    lo, hi = np.quantile(aucs, [alpha, 1 - alpha])
    return point, float(lo), float(hi)


def applicable_methods(feature_key):
    """LGB is only applied to low-dim tabular features, not high-dim embeddings."""
    if feature_key in SINGLE_FEATURE_KEYS:
        return ['lr']
    return ['lr', 'lgb'] #['lr', 'mlp'] if FEATURE_SETS[feature_key][0] == 'embedding' else ['lr', 'lgb', 'mlp']


# ── Migrate old flat cache files into standard/ ────────────────────────────────
_COMPAT = {
    'mlp_amplify_c_lysate':         ('clf_only_lysate_params.json',               'clf_only_lysate_model.pt'),
    'mlp_amplify_c_purified':       ('clf_only_purified_params.json',             'clf_only_purified_model.pt'),
    'lgb_biophys_lysate':           ('clf_biophys_lysate_params.json',            'clf_biophys_lysate_model.joblib'),
    'lgb_biophys_purified':         ('clf_biophys_purified_params.json',          'clf_biophys_purified_model.joblib'),
    'lgb_struct_lysate':            ('clf_struct_lysate_params.json',             'clf_struct_lysate_model.joblib'),
    'lgb_struct_purified':          ('clf_struct_purified_params.json',           'clf_struct_purified_model.joblib'),
    'lgb_biophys+struct_lysate':    ('clf_biophys+struct_lysate_params.json',     'clf_biophys+struct_lysate_model.joblib'),
    'lgb_biophys+struct_purified':  ('clf_biophys+struct_purified_params.json',   'clf_biophys+struct_purified_model.joblib'),
    'mlp_biophys_lysate':           ('clf_mlp_biophys_lysate_params.json',        'clf_mlp_biophys_lysate_model.pt'),
    'mlp_biophys_purified':         ('clf_mlp_biophys_purified_params.json',      'clf_mlp_biophys_purified_model.pt'),
    'mlp_struct_lysate':            ('clf_mlp_struct_lysate_params.json',         'clf_mlp_struct_lysate_model.pt'),
    'mlp_struct_purified':          ('clf_mlp_struct_purified_params.json',       'clf_mlp_struct_purified_model.pt'),
    'mlp_biophys+struct_lysate':    ('clf_mlp_biophys+struct_lysate_params.json', 'clf_mlp_biophys+struct_lysate_model.pt'),
    'mlp_biophys+struct_purified':  ('clf_mlp_biophys+struct_purified_params.json','clf_mlp_biophys+struct_purified_model.pt'),
}

migrated = 0
for key, (pname, mname) in _COMPAT.items():
    old_p, old_m = OUT_DIR / pname, OUT_DIR / mname
    new_p, new_m = _cache_paths(STD_DIR, key, is_mlp=mname.endswith('.pt'))
    if old_p.exists() and old_m.exists() and not (new_p.exists() and new_m.exists()):
        shutil.copy(old_p, new_p); shutil.copy(old_m, new_m)
        migrated += 1
print(f'Migrated {migrated} old cache entries to standard/.')

# %% [code]
# ── Standard & complete regimes (5 seeds, 85/15 split) ────────────────────────
# standard : embeddings only, full pool, 85% train / 15% test.
# complete : all feature sets, complete pool, 85% train / 15% test.
# Running over EVAL_SEEDS captures variance from both split and model randomness.

REGIMES = {
    # 'standard': dict(cache_dir=STD_DIR,      complete_only=False,
    #                  feat_keys=[k for k, (kind, _) in FEATURE_SETS.items() if kind == 'embedding']),
    'complete': dict(cache_dir=COMPLETE_DIR,  complete_only=True,
                     feat_keys=list(FEATURE_SETS)),
}

results_standard = []

for regime_name, cfg in REGIMES.items():
    print(f"\n{'='*60}\nRegime: {regime_name}  features: {cfg['feat_keys']}\n{'='*60}")
    for seed in EVAL_SEEDS:
        seed_dir = cfg['cache_dir'] / f'seed{seed}'
        for feat_key in cfg['feat_keys']:
            for task in TASKS:
                ds = build_dataset(feat_key, task, seed=seed,
                                   complete_only=cfg['complete_only'])
                X_tr, y_tr = ds['X_train'], ds['y_train']
                X_te, y_te = ds['X_test'],  ds['y_test']
                y_bin, in_dim = (y_tr > 0).astype(int), ds['in_dim']
                n_trials = N_TRIALS_EMB if FEATURE_SETS[feat_key][0] == 'embedding' else N_TRIALS_TAB

                for method in applicable_methods(feat_key):
                    key = f'{method}_{feat_key}_{task}'

                    if method == 'mlp':
                        model, cv_auc, cached = train_or_load_mlp(seed_dir, key, X_tr, y_bin, in_dim, n_trials)
                    elif method == 'lgb':
                        model, cv_auc, cached = train_or_load_lgb(seed_dir, key, X_tr, y_bin, n_trials)
                    else:
                        model, cv_auc, cached = train_or_load_lr(seed_dir, key, X_tr, y_bin)

                    test_auc = evaluate(model, method, X_te, y_te)
                    print(f"  {'[cache]' if cached else '[train]'} s={seed} {key:50s}  "
                          f"n_train={len(y_tr)} n_test={len(y_te)}  AUC={test_auc:.4f}")
                    results_standard.append(dict(
                        regime=regime_name, seed=seed, feature=feat_key,
                        task=task, method=method,
                        n_train=len(y_tr), n_test=len(y_te), test_auc=test_auc,
                    ))

df_std = pd.DataFrame(results_standard)
df_std.to_csv(STD_DIR / 'results.csv', index=False)


# %% [code]
# ── Results tables — mean ± std AUC across seeds ──────────────────────────────
for regime_name in ['standard', 'complete']:
    print(f"\n{chr(9473)*60}\nRegime: {regime_name}")
    sub = df_std[df_std.regime == regime_name]
    agg = (sub.groupby(['feature', 'task', 'method'])['test_auc']
              .agg(['mean', 'std']).reset_index())
    agg['auc_str'] = (agg['mean'].round(3).astype(str)
                      + ' ± ' + agg['std'].round(3).astype(str))
    for task in TASKS:
        pivot = (agg[agg.task == task]
                 .pivot(index='feature', columns='method', values='auc_str'))
        print(f"\n  Task: {task}")
        display(pivot)


# %% [code]
# ── Paired permutation test: pairwise significance within each regime/task ─────
# For each (feature, task, regime), test every pair of methods.
# H0: A and B are equivalent — sign of per-seed diff is arbitrary.
# p-value = fraction of random sign-flips that produce a mean diff >= observed.
# With n=5 seeds, minimum achievable p = 1/2^5 = 0.031.

def paired_permutation_p(a, b, n_permutations=10_000, rng=None):
    """Two-sided paired permutation test. Returns p-value."""
    rng  = np.random.default_rng(rng)
    diff = np.array(a) - np.array(b)
    observed = np.abs(diff.mean())
    signs = rng.choice([-1, 1], size=(n_permutations, len(diff)))
    null  = np.abs((signs * diff).mean(axis=1))
    return (null >= observed).mean()

regime_dfs = {
    'complete':  df_std[(df_std.regime == 'complete')],
    # 'low-data':  df_low,
}

sig_results_by_regime_task = {}

for regime_name, df_regime in regime_dfs.items():
    print(f"\n{'─'*60}")
    print(f"Regime: {regime_name}")
    for task in TASKS:
        print(f"\n  Task: {task}")
        sub = df_regime[df_regime.task == task]
        # pivot: index=seed, columns=(feature, method)
        pivot = (sub.pivot_table(index='seed', columns=['feature', 'method'],
                                 values='test_auc'))
        keys = list(pivot.columns)
        from itertools import combinations
        rows = []
        for (f1, m1), (f2, m2) in combinations(keys, 2):
            a = pivot[(f1, m1)].dropna().values
            b = pivot[(f2, m2)].dropna().values
            if len(a) < 2 or len(a) != len(b):
                continue
            p = paired_permutation_p(a, b)
            rows.append(dict(A=f'{f1}/{m1}', B=f'{f2}/{m2}',
                             feature_1=f1, method_1=m1,
                             feature_2=f2, method_2=m2,
                             mean_A=a.mean(), mean_B=b.mean(),
                             diff=a.mean()-b.mean(), p=p))
        res = (pd.DataFrame(rows)
               .sort_values('p')
               .reset_index(drop=True))
        # BH correction
        from statsmodels.stats.multitest import multipletests
        reject, p_bh, _, _ = multipletests(res['p'], method='fdr_bh')
        res['p_bh']   = p_bh
        res['sig']    = reject
        sig_results_by_regime_task[(regime_name, task)] = res[res['sig']].copy()
        # display(res[res['sig']][['A', 'B', 'mean_A', 'mean_B', 'diff', 'p', 'p_bh']]
        #         .round(4)
        #         .style.set_caption(f'Significant pairs (BH-corrected p<0.05) — {regime_name} / {task}'))


# %% [code]
# ── Bar plots: methods compared within each data regime ───────────────────────
# One figure per regime, with tasks shown side by side.
# Each bar = one (feature, method) combination, ordered by mean AUC descending.
# Color: AMPLIFY = blue, all other feature sets = grey.
# Error bars = 95% CI (mean ± 1.96 × std / √n).
# Brackets mark BH-corrected significant differences against AMPLIFY among visible bars.

def feat_color(feat):
    return sns.color_palette("coolwarm", n_colors=10)[0] if 'amplify' in feat.lower() else 'lightgrey'

def significance_label(p):
    if p < 1e-3:
        return '***'
    if p < 1e-2:
        return '**'
    return '*'

def amplify_only_sig_pairs(sig_pairs, visible_keys):
    if sig_pairs.empty:
        return sig_pairs

    sig_pairs = sig_pairs.copy()
    sig_pairs = sig_pairs[
        sig_pairs.apply(
            lambda r: ((r['feature_1'], r['method_1']) in visible_keys)
            and ((r['feature_2'], r['method_2']) in visible_keys),
            axis=1,
        )
    ]
    sig_pairs = sig_pairs[
        ((sig_pairs['feature_1'] == 'amplify_c') & (sig_pairs['method_1'] == 'lr'))
        | ((sig_pairs['feature_2'] == 'amplify_c') & (sig_pairs['method_2'] == 'lr'))
    ].copy()
    if sig_pairs.empty:
        return sig_pairs

    swap_mask = (sig_pairs['feature_2'] == 'amplify_c') & (sig_pairs['method_2'] == 'lr')
    if swap_mask.any():
        cols_1 = ['feature_1', 'method_1']
        cols_2 = ['feature_2', 'method_2']
        tmp_vals = sig_pairs.loc[swap_mask, cols_1].copy()
        sig_pairs.loc[swap_mask, cols_1] = sig_pairs.loc[swap_mask, cols_2].to_numpy()
        sig_pairs.loc[swap_mask, cols_2] = tmp_vals.to_numpy()
    return sig_pairs

def collapse_amplify_brackets(sig_pairs, xpos):
    if sig_pairs.empty:
        return []

    collapsed = []
    sig_pairs = sig_pairs.copy()
    sig_pairs['label'] = sig_pairs['p_bh'].map(significance_label)
    sig_pairs['x_anchor'] = sig_pairs.apply(lambda r: xpos[(r['feature_1'], r['method_1'])], axis=1)
    sig_pairs['x_target'] = sig_pairs.apply(lambda r: xpos[(r['feature_2'], r['method_2'])], axis=1)

    for (anchor, label), grp in sig_pairs.sort_values('x_target').groupby(['x_anchor', 'label'], sort=False):
        targets = sorted(grp['x_target'].unique())
        run_start = targets[0]
        run_end = targets[0]
        for x in targets[1:]:
            if x == run_end + 1:
                run_end = x
            else:
                collapsed.append((min(anchor, run_start), max(anchor, run_end), label))
                run_start = run_end = x
        collapsed.append((min(anchor, run_start), max(anchor, run_end), label))

    return sorted(set(collapsed), key=lambda t: (t[1] - t[0], t[0], t[2]))

def add_significance_brackets(ax, tmp, sig_pairs):
    if sig_pairs.empty:
        return

    xpos = {(row['feature'], row['method']): i for i, (_, row) in enumerate(tmp.iterrows())}
    y_max = float((tmp['mean'] + tmp['ci95'].fillna(0)).max())
    bracket_height = 0.01
    level_step = 0.035
    text_pad = 0.006
    base_y = y_max + 0.02
    occupied = []
    max_level = 0

    collapsed = collapse_amplify_brackets(sig_pairs, xpos)
    if not collapsed:
        return

    for x1, x2, label in collapsed:
        level = 0
        while any(not (x2 < a or x1 > b) and level == used_level for a, b, used_level in occupied):
            level += 1
        y = base_y + level * level_step
        ax.plot([x1, x1, x2, x2], [y, y + bracket_height, y + bracket_height, y],
                color='#444', lw=1, clip_on=False, zorder=3)
        ax.text((x1 + x2) / 2, y + bracket_height + text_pad, label,
                ha='center', va='bottom', fontsize=8, color='#222', clip_on=False)
        occupied.append((x1, x2, level))
        max_level = max(max_level, level)

    ymin, ymax = ax.get_ylim()
    ax.set_ylim(ymin, max(ymax, base_y + (max_level + 1) * level_step + 0.03))

def add_significance_brackets_horizontal(ax, tmp, sig_pairs, fontsize=14):
    if sig_pairs.empty:
        return

    ypos = {(row['feature'], row['method']): i for i, (_, row) in enumerate(tmp.iterrows())}
    x_max = float((tmp['mean'] + tmp['ci95'].fillna(0)).max())
    bracket_width = 0.01
    level_step = 0.035
    text_pad = 0.006
    base_x = x_max + 0.02
    occupied = []
    max_level = 0

    collapsed = collapse_amplify_brackets(sig_pairs, ypos)
    if not collapsed:
        return

    for y1, y2, label in collapsed:
        level = 0
        while any(not (y2 < a or y1 > b) and level == used_level for a, b, used_level in occupied):
            level += 1
        x = base_x + level * level_step
        ax.plot([x, x + bracket_width, x + bracket_width, x], [y1, y1, y2, y2],
                color='#444', lw=1, clip_on=False, zorder=3)
        ax.text(x + bracket_width + text_pad, (y1 + y2) / 2, label,
                ha='left', va='center', fontsize=fontsize, color='#222', clip_on=False)
        occupied.append((y1, y2, level))
        max_level = max(max_level, level)

    xmin, xmax = ax.get_xlim()
    ax.set_xlim(xmin, max(xmax, base_x + (max_level + 1) * level_step + 0.05))

# %% [code]
regime_dfs = {
    'complete':  df_std[(df_std.regime == 'complete')],
    # 'low-data':  df_low,
}

if 'sig_results_by_regime_task' not in globals():
    raise RuntimeError('Run the paired permutation test cell first to populate sig_results_by_regime_task.')

for regime_name, df_regime in regime_dfs.items():
    n_seeds = df_regime['seed'].nunique() if 'seed' in df_regime.columns else 1
    agg = (df_regime
           .groupby(['feature', 'task', 'method'])['test_auc']
           .agg(['mean', 'std', 'count'])
           .reset_index())
    agg['ci95'] = 1.96 * agg['std'] / np.sqrt(agg['count'])

    task_subsets = [
        (task,
         agg[(agg.task == task)]
         .sort_values('mean', ascending=False)
         .reset_index(drop=True))
        for task in TASKS
    ]
    n_best = 10
    widths = [0.7 * n_best for _, sub in task_subsets]
    barwidth=0.6
    # fig, axes = plt.subplots(
    #     1, len(task_subsets),
    #     figsize=(sum(widths), 6.5),
    #     sharey=True,
    #     gridspec_kw={'width_ratios': widths}
    # )
    # if len(task_subsets) == 1:
    #     axes = [axes]

    # color_handles = [
    #     plt.Rectangle((0, 0), 1, 1, color=feat_color('amplify'), label='AMPLIFY'),
    #     plt.Rectangle((0, 0), 1, 1, color='#aaaaaa', label='baseline'),
    # ]

    # for ax, (task, sub) in zip(axes, task_subsets):
    #     tmp = sub.head(n_best).copy()
    #     for x, row in tmp.iterrows():
    #         ax.bar(x, row['mean'], width=0.8,
    #                color=feat_color(row['feature']),
    #                edgecolor='white', linewidth=0.5,
    #                yerr=row['ci95'] if not np.isnan(row['ci95']) else 0,
    #                capsize=3, error_kw=dict(lw=1, ecolor='#333'), zorder=2)

    #     ax.set_xticks(range(len(tmp)))
    #     ax.set_xticklabels(
    #         [f"{r['feature'].split(':')[-1].replace('_soluble', '').replace('struct', 'Str.').replace('biophys', 'Bio.').replace('amplify_c', r'AMPLIFY$_c$')}\n{r['method'].upper()}" for _, r in tmp.iterrows()],
    #         fontsize=9, ha='center')
    #     ax.set_ylim(0.4, 1.0)
    #     ax.spines[['top', 'right']].set_visible(False)
    #     ax.spines[['bottom']].set_linewidth(2)
    #     ax.grid(True, axis='y', linestyle='--', alpha=0.5, zorder=0)

    #     visible_keys = set(tmp[['feature', 'method']].itertuples(index=False, name=None))
    #     sig_pairs = sig_results_by_regime_task.get((regime_name, task), pd.DataFrame()).copy()
    #     sig_pairs = amplify_only_sig_pairs(sig_pairs, visible_keys)
    #     add_significance_brackets(ax, tmp, sig_pairs)

    # axes[0].set_ylabel('Test AUC', fontsize=11)
    # axes[0].tick_params(axis='y', labelsize=10)
    # if len(axes) > 1:
    #     for ax in axes[1:]:
    #         ax.tick_params(axis='y', labelsize=10)
    # plt.tight_layout(rect=[0, 0, 1, 0.9])
    # for ax, (task, _) in zip(axes, task_subsets):
    #     bb = ax.get_position()
    #     fig.text(
    #         bb.x0 + bb.width / 2,
    #         0.90,
    #         f'{task.title()} band\n(mean ± 95% CI, {n_seeds} seeds)',
    #         ha='center', va='bottom', fontsize=12
    #     )
    # plt.savefig(FIG_DIR / f'bar_methods_{regime_name}.png',dpi=300, bbox_inches='tight')
    # plt.savefig(FIG_DIR / f'bar_methods_{regime_name}.pdf',dpi=300, bbox_inches='tight')
    # plt.show()
    
    fontsize = 18

    fig, axes = plt.subplots(
        1, len(task_subsets),
        figsize=(sum(widths), 5),
        sharex=True
    )
    if len(task_subsets) == 1:
        axes = [axes]

    for ax, (task, sub) in zip(axes, task_subsets):
        tmp = sub.head(n_best).copy().reset_index(drop=True)
        y = np.arange(len(tmp))
        # Set bar width
        ax.barh(y, tmp['mean'], height=barwidth,
                color=[feat_color(f) for f in tmp['feature']],
                edgecolor='white', linewidth=0.5,
                xerr=tmp['ci95'].fillna(0),
                capsize=0, error_kw=dict(lw=1, ecolor='#333'), zorder=2)

        ax.set_yticks(y)
        ax.set_yticklabels(
            [f"{r['feature'].split(':')[-1].replace('amplify_c', r'AMPLIFY$_c$')} - {r['method'].upper()}" for _, r in tmp.iterrows()],
            fontsize=fontsize - 2
        )
        ax.invert_yaxis()
        ax.set_xlim(0.4, 1.0)
        ax.spines[['top', 'right']].set_visible(False)
        ax.spines[['bottom']].set_linewidth(2)
        ax.grid(True, axis='x', linestyle='--', alpha=0.5, zorder=0)
        ax.set_xlabel(f'{task.title()} band prediction - Test AUC', fontsize=fontsize)

        visible_keys = set(tmp[['feature', 'method']].itertuples(index=False, name=None))
        sig_pairs = sig_results_by_regime_task.get((regime_name, task), pd.DataFrame()).copy()
        sig_pairs = amplify_only_sig_pairs(sig_pairs, visible_keys)
        add_significance_brackets_horizontal(ax, tmp, sig_pairs, fontsize=12)

    axes[0].tick_params(axis='x', labelsize=fontsize)
    if len(axes) > 1:
        for ax in axes[1:]:
            ax.tick_params(axis='x', labelsize=fontsize)
    plt.tight_layout(rect=[0, 0, 1, 0.92])
    # for ax, (task, _) in zip(axes, task_subsets):
    #     bb = ax.get_position()
    #     fig.text(
    #         bb.x0 + bb.width / 2,
    #         0.92,
    #         f'{task.title()} band prediction', #\n(mean ± 95% CI, {n_seeds} seeds)
    #         ha='center', va='bottom', fontsize=fontsize + 2,
    #     )
    plt.savefig(f'figures/bar_methods_{regime_name}_horizontal.png',dpi=300, bbox_inches='tight')
    # plt.savefig(f'figures/bar_methods_{regime_name}_horizontal.pdf',dpi=300, bbox_inches='tight')
    plt.show()


# %% [code]
regime_dfs["complete"].to_csv(STD_DIR / 'results_complete.csv', index=False)

# %% [code]
```
