## Supplementary material for "pLM representations unlock metagenomic space beyond homology": Code bundle: mlp_methodology.md

# Methods — Enzyme expression classification benchmark

**Task and dataset.** We predicted recombinant enzyme expression as a binary outcome — detectable vs. undetectable band — at two independent readouts (crude lysate and affinity-purified fraction) for 382 enzymes. Proteins missing a target value were excluded (378 retained per task). The set was curated by the experimental team to consist of remote homologs only, precluding sequence-identity leakage between train and test splits. Band quantification and the detection threshold were defined by the wet-lab pipeline.

**Features.** Four feature families were benchmarked: (i) a 960-dimensional AMPLIFY-C protein-language-model embedding, obtained by averaging per-residue representations; (ii) 22 sequence-derived biophysical descriptors (aromaticity, GRAVY, isoelectric point, charge at pH 7/7.5/8, flexibility, instability index, molar extinction coefficients, molecular weight, Shannon entropy, disorder fractions, signal-peptide likelihood, predicted secondary-structure content, and repeat ratio); (iii) four structural descriptors computed from AlphaFold-predicted structures — mean pLDDT, mean SASA, and per-residue averages of ESM-IF and soluble-ProteinMPNN log-likelihoods; and (iv) their concatenation. Features were standardized using statistics fit on the training split only.

**Models.** Three classifiers were compared: L2-regularized logistic regression, LightGBM gradient-boosted trees, and a multilayer perceptron (MLP). Logistic regression and the MLP were applied to all feature sets; LightGBM was restricted to the low-dimensional biophysical and structural descriptors, where tree ensembles are best suited, and was not run on the high-dimensional language-model embeddings. The MLP comprised 1–3 (Linear → LayerNorm → ReLU → Dropout) blocks and a linear output head, trained for 150 epochs with binary cross-entropy, Adam with L2 weight decay, and cosine-annealed learning rate. LayerNorm was preferred over BatchNorm because of the small sample size.

**Hyperparameter optimization.** For every (feature × method × task × seed) combination, hyperparameters were selected by stratified 5-fold cross-validation on the training split (3-fold when n ≤ 50), scored by mean validation ROC-AUC. The MLP and LightGBM search spaces were explored with Bayesian optimization (Optuna, Tree-structured Parzen Estimator, 50 trials; 20 trials in the low-data regime); logistic regression was tuned by grid search over C ∈ {10⁻⁴, …, 10²}. LightGBM was class-reweighted via `scale_pos_weight`. The selected configuration was refit on the full training split before evaluation.

**Data regimes.** Three evaluation regimes were defined. In the *standard* regime, sequence-embedding models were trained and tested on the full 378-protein pool. In the *complete* regime, all five feature families were evaluated on the shared 287-protein pool with full descriptor coverage, guaranteeing matched sample identity across feature sets. Both used an 85%/15% stratified train/test split. A *low-data* regime inverted the ratio (15%/85%) on the complete pool to probe data efficiency. Each regime was repeated over 20 independent seeds, with stratification on the binary target.

**Statistical analysis.** Performance was summarized as the mean ± standard deviation of test ROC-AUC across the 20 seeds. Pairwise differences between methods within each (feature, task, regime) stratum were assessed by a paired permutation test (10,000 permutations) on the per-seed AUCs, exploiting the fact that all methods saw identical splits; p-values were corrected for multiple comparisons using the Benjamini–Hochberg procedure at a 5% false-discovery rate. A 95% percentile-bootstrap confidence interval (1,000 resamples) around the single-split AUC was additionally computed for each configuration. All random operations were controlled by explicit seeds for reproducibility.
