## Supplementary material for "pLM representations unlock metagenomic space beyond homology": Code bundle: prepare_blast.ipynb

```
# %% [code]
from pathlib import Path
from functools import reduce

import math
import numpy as np
import pandas as pd
import json

from scipy.stats import pearsonr, spearmanr
from scipy.spatial.distance import pdist, squareform
from sklearn.metrics.pairwise import cosine_similarity

import matplotlib.pyplot as plt
import seaborn as sns
from umap import UMAP

from utils import merge_dataframes, plot_distribution
%config InlineBackend.figure_format='retina'

# %% [code]
# Data version
idx = 4

# Define the primary key for merging
KEY_COL = "uniprot_id"
METHOD = "mmr"
FILTER = "final"

DROP_EMBEDDINGS = True

# Define the directory where your results are stored
DATA_DIR = Path("$SCRATCH/proseqo/enzymes/filtered_outputs")
MERGED_FILE = f"merged_data_{idx:03}.parquet"

FIG_DIR = Path("$SCRATCH/proseqo/enzymes/output/figures")
FIG_DIR.mkdir(exist_ok=True)

print("Loading metrics files...")
df_main = pd.read_parquet(DATA_DIR / MERGED_FILE)

if DROP_EMBEDDINGS == True:
    print("Dropping embedding columns to reduce dataframe size...")
    df_main = df_main.drop(columns=[c for c in df_main.columns if "embedding" in c])

# %% [markdown]
## Uniprot candidates

# %% [code]
def get_selected(df_main, log_dir, lambda_param, method):
    selected_file = log_dir / f"selected_uniprot_ids_{method}_lambda_{lambda_param:.2f}_50.txt"
    # print(f"Loading selected IDs from {selected_file}...")
    try:
        with open(selected_file, "r") as f:
            # Read IDs, strip whitespace, and remove empty lines
            selected_ids = set(line.strip() for line in f if line.strip())

        print(f"Found {len(selected_ids)} unique IDs.")

        # Filter the main dataframe to keep only the selected sequences
        # Using KEY_COL which was defined as 'uniprot_id'
        df_selected = df_main[df_main[KEY_COL].isin(selected_ids)]

        print(f"Successfully filtered dataframe. Retained {len(df_selected)} sequences matching the selected IDs.")

    except FileNotFoundError:
        print(f"Error: {selected_file} not found. Please ensure the file is in the notebook directory.")
        df_selected = pd.DataFrame()  # Empty fallback

    return df_selected

# %% [code]
log_dir = Path("$HOME/enzymes/selection_logs/filter_004_mmr_from_lambda_0.90")
lambda_param = 0.10

df_selected = get_selected(df_main, log_dir, lambda_param=lambda_param, method="mmr")

fasta_path = f"$HOME/enzymes/data/uniprot_selected_{lambda_param}.fasta"
with open(fasta_path, "w") as f:
    for id, seq in zip(df_selected["uniprot_id"], df_selected["sequence"]):
        f.write(f">{id}\n")
        f.write(f"{seq}\n")

# %% [code]
log_dir = Path("$HOME/enzymes/selection_logs/final")
lambda_param = 0.90

df_selected = get_selected(df_main, log_dir, lambda_param=lambda_param, method="mmr")

fasta_path = f"$HOME/enzymes/data/uniprot_selected_{lambda_param}.fasta"
with open(fasta_path, "w") as f:
    for id, seq in zip(df_selected["uniprot_id"], df_selected["sequence"]):
        f.write(f">{id}\n")
        f.write(f"{seq}\n")

# %% [markdown]
### BFD / MGY

# %% [code]
base_path = "$SCRATCH/proseqo/enzymes/metagenomic_search"

# %% [code]
mgnify_csvs = [
    "mgnify_0_potential_enzymes",
    "mgnify_1_potential_enzymes",
    "mgnify_2_potential_enzymes",
    "mgnify_3_potential_enzymes",
]
```
