## Supplementary material for "pLM representations unlock metagenomic space beyond homology": Code bundle: results.ipynb

```
# %% [code]
from pathlib import Path
from functools import reduce

import random
import os
import math
import numpy as np
import pandas as pd
import json
import polars as pl

from scipy.stats import pearsonr, spearmanr
from scipy.spatial.distance import pdist, squareform
from sklearn.metrics.pairwise import cosine_similarity

from matplotlib.patches import Patch

import matplotlib.pyplot as plt
import seaborn as sns

import numpy as np
import pandas as pd
from scipy import stats

FIG_DIR = Path("figures")

%config InlineBackend.figure_format='retina'

# %% [code]
def plot_distribution(ax, data, prop, title = None):
    # Plot distribution
    sns.histplot(
        data,
        x=prop,
        bins=30,
        color="#0085ff",
        alpha=0.4,
        lw=0.1,
        ax=ax,
        stat="count",
    )
    if title is None:
        title = f"Distribution of {prop}"
    ax.set_title(title)
    ax.set_xlabel(prop)
    ax.set_ylabel("Count")
    ax.tick_params(left=False)
    ax.spines[["top", "right", "left"]].set_visible(False)
    ax.spines["bottom"].set_linewidth(2)
    ax.yaxis.grid(True, which="major", linestyle=":")
    ax.set_axisbelow(True)
    
def plot_correlation(df, metric1, metric2, p_corr, s_corr, color_metric, remove_outliers=False, min_correlation=None, figsize=(7, 6), ax=None, vmax_color=2048, cmap="viridis"):
    # Check if all required columns exist before plotting
    required_cols = [metric1, metric2, color_metric]
    if all(col in df.columns for col in required_cols):
        plot_data = df[[metric1, metric2]].dropna()

        if not plot_data.empty:
            # Early exit if correlation is too low
            if min_correlation is not None and (abs(p_corr) < min_correlation and abs(s_corr) < min_correlation):
                return None, None

            if ax is None:
                fig, ax = plt.subplots(figsize=figsize)
                is_standalone = True
                title_fs, label_fs = 12, 10
            else:
                fig = ax.get_figure()  # Get the parent figure of the passed axis
                is_standalone = False
                title_fs, label_fs = 12, 10

            points = ax.scatter(
                plot_data[metric1],
                plot_data[metric2],
                c=plot_data[color_metric].values,
                cmap=cmap,
                s=1,
                alpha=0.9,
                vmax=vmax_color,
            )

            if remove_outliers:
                x_min, x_max = plot_data[metric1].quantile(0.001), plot_data[metric1].quantile(0.999)
                y_min, y_max = plot_data[metric2].quantile(0.001), plot_data[metric2].quantile(0.999)
                ax.set_xlim(x_min, x_max)
                ax.set_ylim(y_min, y_max)

            # y_05 = plot_data[metric2].quantile(0.05)
            # y_95 = plot_data[metric2].quantile(0.95)
            # ax.hlines(y_95, 0, 100, colors='red', linestyles='dashed')
            # ax.hlines(y_05, 0, 100, colors="red", linestyles="dashed")

            # x_05 = plot_data[metric1].quantile(0.05)
            # x_95 = plot_data[metric1].quantile(0.95)
            # ax.vlines(x_05, 0, 100, colors="red", linestyles="dashed")
            # ax.vlines(x_95, 0, 100, colors='red', linestyles='dashed')

            cbar = fig.colorbar(points, ax=ax)
            cbar.set_label(color_metric, fontsize=label_fs)

            if is_standalone:
                ax.set_title(f"{metric1} vs. {metric2}\nPearson r: {p_corr:.3f} | Spearman ρ: {s_corr:.3f}")

            ax.set_xlabel(metric1, fontsize=label_fs)
            ax.set_ylabel(metric2, fontsize=label_fs)

            ax.tick_params(axis="both", which="major", labelsize=label_fs)

            ax.spines[["top", "right"]].set_visible(False)
            ax.spines[["left", "bottom"]].set_linewidth(2)

            if is_standalone:
                plt.tight_layout()
                plt.savefig(FIG_DIR / f"{metric1}_{metric2}_correlation.png", dpi=300)
                plt.show()

            return p_corr, s_corr
        else:
            print(f"Skipping: No data for {required_cols}")
    else:
        print(f"Skipping: Missing columns {required_cols}")

# %% [code]
data = pd.read_csv("$HOME/enzymes/notebooks/data.csv")

# %% [code]
activity_col = ['proline (EIC)', 'lysine (EIC)', 'indanone (245 nm)', 'resorcinol (285 nm)']
# Normalize activity columns by zscore across all plates
for col in activity_col:
    data[col + " (zscore)"] = (data[col] - data[col].mean()) / data[col].std()
    
# Normalize activity columns by zscore within each plate
for col in activity_col:
    data[col + " (zscore_plate)"] = data.groupby("plate")[col].transform(lambda x: (x - x.mean()) / x.std())
    
zscore_col = ['proline (EIC) (zscore)', 'lysine (EIC) (zscore)', 'indanone (245 nm) (zscore)', 'resorcinol (285 nm) (zscore)']
data["max_activity_zscore"] = data[zscore_col].max(axis=1)

zscore_col = ['proline (EIC) (zscore_plate)', 'lysine (EIC) (zscore_plate)', 'indanone (245 nm) (zscore_plate)', 'resorcinol (285 nm) (zscore_plate)']
data["max_activity_zscore_plate"] = data[zscore_col].max(axis=1)

data.to_csv("$HOME/enzymes/notebooks/data.csv", index=False)

# %% [code]
## SI file creation

tmp = pd.read_csv("$HOME/enzymes/notebooks/soluble.csv")

id_matching = {
    "BFD_AP12_scaffold13649": "BFD_AP12_2_1047962.scaffolds.fasta_scaffold13649_1",
    "BFD_AP12_scaffold1914741": "BFD_AP12_2_1047962.scaffolds.fasta_scaffold1914741_1",
    "BFD_CryGeyStandDraft_7": "BFD_CryGeyStandDraft_7_1057128.scaffolds.fasta_scaffold320846_1",
    "BFD_SwirhisoilCB3": "BFD_SwirhisoilCB3_FD_contig_31_13352357_length_237_multi_1_in_0_out_0_1",
}

tmp["seq ID"] = tmp["seq ID"].replace(id_matching)

tmp = tmp.merge(data[["id", "sequence"]], left_on = "seq ID", right_on = "id", how="left")
tmp = tmp[['seq ID', 'location', "sequence", 'kDa', 'lysate band', 'purified band',
       'mg/mL', 'plate', 'well']]

missing_seq = {"A0A1Y0CYF1": "MTMCTDDCYPSRLSHTLAPFKRLDPIVYGSQSTGPLTSTELAQYERDGFLVIKDFLDEQQVQDFQQDLQHYQQHPQQLPNELVIKEAQQANIRSIFDIHKHNPRFDELTREPRLLAMVEQLLGSEVYIHQSRINVKPAFYGKGFDWHSDFETWHTEDGMPRMRALSCSIALTPNTQVNGSLMLIPGSHHYFVPCVGPTPTNNYRSSLQKQQVGVPSEQALNNLANQGGISMPTGPSGSLILFECNTLHASNANLSPWPRSNLFFVYNSVHNRLTTPFCGQAPRPEHLANRQHSGALVSSDIAAYA", 
 "MGYP007091382478": "MRNSIREKHTRKPPTYKHIQTTRKNPTYTHIQTTRKNPHIQTHPDNTKKKTHTNTFRQHEKTTNTNTFRQHEKNNKYKHIQTTRKNPHTNTFRQHEKTTHTDTFRQHEKSHIHTHSNNTKRHTHTNTFRQHEKTHTYKHIQTTRKKHPHTNTFRQHEKNNKYKYIQTTRKQPHIQTHSDNTKKTTHTDTFRQHEKTTNTNTSKQHYCNDSTSNV"}

# Complete sequence column for missing sequences
for seq_id, seq in missing_seq.items():
    tmp.loc[tmp['seq ID'] == seq_id, 'sequence'] = seq
    
tmp.to_csv("$HOME/enzymes/SI/in_vivo_results.csv", index=False)

# %% [code]
data["normalized_lysate_band"] = data["lysate band"].transform(lambda x: (x - x.min()) / (x.max() - x.min()))
data["normalized_purified_band"] = data["purified band"].transform(lambda x: (x - x.min()) / (x.max() - x.min()))
data["max_band"] = data[["normalized_lysate_band", "normalized_purified_band"]].max(axis=1)
tmp = data[data["plate"] == 3] #[['id', 'mg/mL', 'lysate band', 'purified band', 'max_activity_zscore', "max_activity_zscore_plate"]]
expressed = tmp[(tmp["lysate band"] > 0) | (tmp["purified band"] > 0)]
overly_expressed = tmp[(tmp["lysate band"] > 500) | (tmp["purified band"] > 500)]

# %% [code]
tmp = data.groupby("plate").sum(numeric_only=True).reset_index().copy()
tmp = tmp[tmp["plate"] != 3]

fontsize = 14

plate_labels = {
    1: "A",
    2: "B",
    4: "C",
}

palette = sns.color_palette("coolwarm", n_colors=10)
palette = [palette[0], palette[2], palette[6]]
plate_colors = dict(zip(plate_labels.values(), palette))

x = np.arange(len(plate_labels))
width = 0.45

fig, ax = plt.subplots(figsize=(6, 5))

# Purified bars (solid)
ax.bar(x - width/2, tmp["purified band"], width, color=list(plate_colors.values()), alpha=1, lw=1, label="Purified", edgecolor="white")

# Lysate bars (hatched)
ax.bar(x + width/2, tmp["lysate band"], width, color=list(plate_colors.values()), alpha=1, lw=1, hatch="//", edgecolor="white", label="Lysate")

# ax.set_xlabel("Plate")
ax.set_ylabel("Total band volume", fontsize=fontsize)
ax.set_xticks(x)
ax.set_xticklabels(plate_labels.values(), fontsize=fontsize)
ax.grid(axis="y", linestyle=":", alpha=0.7)

# Set y ticks in scientific notation
ax.ticklabel_format(axis="y", style="sci", scilimits=(0,0))

ax.spines[["top", "right", "left"]].set_visible(False)
ax.spines["bottom"].set_linewidth(2)

# Legend: solid patch for purified, hatched patch for lysate
legend_elements = [
    Patch(facecolor="white", label="Purified", edgecolor="black"),
    Patch(facecolor="white", hatch="//", edgecolor="black", label="Lysate"),
]
ax.legend(handles=legend_elements, fontsize=fontsize, title_fontsize=fontsize)

plt.tight_layout()
plt.savefig(FIG_DIR / "total_bands_uniprot.png", dpi=300)
plt.show()

# %% [code]
tmp = data.groupby("plate").sum(numeric_only=True).reset_index().copy()
tmp = tmp[tmp["plate"] != 3]

fontsize = 14

plate_labels = {
    1: "A",
    2: "B",
    4: "C",
}

palette = sns.color_palette("coolwarm", n_colors=10)
palette = [palette[0], palette[2], palette[6]]
plate_colors = dict(zip(plate_labels.values(), palette))

tmp["plate_label"] = tmp["plate"].map(plate_labels)

x = np.arange(len(plate_labels))
width = 0.25

fig, ax1 = plt.subplots(figsize=(8, 5))
ax1.spines[["top"]].set_visible(False)
ax1.spines[["left"]].set_visible(False)
ax1.spines[["right"]].set_visible(False)

ax2 = ax1.twinx()
ax2.spines[["top"]].set_visible(False)
ax2.spines[["left"]].set_visible(False)
ax2.spines[["right"]].set_visible(False)

for i, (plate_num, plate_label) in enumerate(plate_labels.items()):
    row = tmp[tmp["plate"] == plate_num].iloc[0]
    offset = (i - 1) * width
    ax1.bar(0 + offset, row["purified band"], width, color=plate_colors[plate_label], alpha=1, lw=1, edgecolor="white")
    ax2.bar(1 + offset, row["lysate band"], width, color=plate_colors[plate_label], alpha=1, lw=1, edgecolor="white") #hatch="//",

ax1.set_ylabel("Purified band volume", fontsize=fontsize)
ax2.set_ylabel("Lysate band volume", fontsize=fontsize)
ax1.set_xticks([0, 1])
ax1.set_xticklabels(["Purified", "Lysate"], fontsize=fontsize)
ax1.grid(axis="y", linestyle=":", alpha=0.7)
ax1.ticklabel_format(axis="y", style="sci", scilimits=(0,0))
ax2.ticklabel_format(axis="y", style="sci", scilimits=(0,0))

# ax1.spines[["top"]].set_visible(False)
# ax2.spines[["top"]].set_visible(False)
ax1.spines["bottom"].set_linewidth(2)
ax2.spines["bottom"].set_linewidth(2)

legend_elements = [
    Patch(facecolor=plate_colors[label], edgecolor="white", label=f"Plate {label}")
    for label in plate_labels.values()
]
ax1.legend(handles=legend_elements, fontsize=fontsize, title_fontsize=fontsize, ncol=3, loc="upper center", bbox_to_anchor=(0.5, 1.15))

plt.tight_layout()
plt.savefig(FIG_DIR / "total_bands_uniprot.png", dpi=300)
plt.show()


# %% [code]
tmp = data.copy()
tmp = tmp[tmp["plate"] != 3]

# Min-max normalize individual values before aggregating
for col in ["purified band", "lysate band"]:
    col_min = tmp[col].min()
    col_max = tmp[col].max()
    tmp[col + " norm"] = (tmp[col] - col_min) / (col_max - col_min)

tmp = tmp.groupby("plate").mean(numeric_only=True).reset_index()

fontsize = 16

plate_labels = {
    1: "A",
    2: "B",
    4: "C",
}

palette = sns.color_palette("coolwarm", n_colors=10)
palette = [palette[0], palette[2], palette[6]]
plate_colors = dict(zip(plate_labels.values(), palette))

x = np.arange(len(plate_labels))
width = 0.25

fig, ax = plt.subplots(figsize=(6, 5))
ax.spines[["top", "left", "right"]].set_visible(False)
ax.spines["bottom"].set_linewidth(2)

for i, (plate_num, plate_label) in enumerate(plate_labels.items()):
    row = tmp[tmp["plate"] == plate_num].iloc[0]
    offset = (i - 1) * width
    ax.bar(0 + offset, row["lysate band norm"], width, color=plate_colors[plate_label], alpha=1, lw=1, edgecolor="white")
    ax.bar(1 + offset, row["purified band norm"], width, color=plate_colors[plate_label], alpha=1, lw=1, edgecolor="white")

ax.set_ylabel("Normalized average band volume", fontsize=fontsize)
ax.set_xticks([0, 1])
ax.set_xticklabels(["Lysate", "Purified"], fontsize=fontsize)
ax.grid(axis="y", linestyle=":", alpha=0.7)

legend_elements = [
    Patch(facecolor=plate_colors[label], edgecolor="white", label=f"Plate {label}")
    for label in plate_labels.values()
]
ax.legend(handles=legend_elements, fontsize=fontsize, title_fontsize=fontsize, ncol=3, loc="upper center", bbox_to_anchor=(0.5, 1.15), frameon=False)

plt.tight_layout()
plt.savefig("figures/total_bands_uniprot_normalized.png", dpi=300)
plt.show()


# %% [code]
import pandas as pd
from scipy.stats import kruskal, mannwhitneyu
from statsmodels.stats.multitest import multipletests

# Assumes data has columns: 'plate' (1, 2, 4), 'lysate_band', 'purified_band'

assays = ['lysate band', 'purified band']
plate_pairs = [(1, 4), (2, 4), (1, 2)]
pair_labels = ['A vs C', 'B vs C', 'A vs B']

for assay in assays:
    print(f"\n{'='*50}")
    print(f"  {assay.upper()}")
    print(f"{'='*50}")

    groups = [data.loc[data['plate'] == p, assay].values for p in [1, 2, 4]]

    # Omnibus test
    H, p_kw = kruskal(*groups)
    print(f"\nKruskal-Wallis: H = {H:.3f}, p = {p_kw:.4g}")

    # Pairwise Mann-Whitney U
    p_vals = []
    stats = []
    for (p1, p2), label in zip(plate_pairs, pair_labels):
        x = data.loc[data['plate'] == p1, assay].values
        y = data.loc[data['plate'] == p2, assay].values
        U, p_mw = mannwhitneyu(x, y, alternative='two-sided')
        p_vals.append(p_mw)
        stats.append(U)

    # Benjamini-Hochberg correction
    reject, p_adj, _, _ = multipletests(p_vals, method='fdr_bh')

    print(f"\nPairwise Mann-Whitney U (BH-corrected):")
    for label, U, p_raw, p_corr, sig in zip(
        pair_labels, stats, p_vals, p_adj, reject
    ):
        print(f"  {label}: U = {U:.1f}, p_raw = {p_raw:.4g}, "
              f"p_adj = {p_corr:.4g} {'*' if sig else 'ns'}")

# %% [code]
x = "AMPLIFY-C_pseudo_ppl"
y = "purified band"

fontsize=16

# Prepare scatter data
tmp = data[data["plate"] != 3].copy()
tmp = tmp[~tmp[x].isna() & ~tmp[y].isna()]
tmp["plate"] = tmp["plate"].map(plate_labels)

# Prepare bar data: total purified band per plate
bar_data = tmp.groupby("plate")[y].sum()
# Bar x-centers: mean pseudo-perplexity per plate
bar_centers = tmp.groupby("plate")[x].mean()

fig, ax = plt.subplots(figsize=(6, 4))

# Scatter on primary y-axis (foreground)
sns.scatterplot(
    tmp,
    x=x,
    y=y,
    hue="plate",
    size="length",
    palette=palette,
    alpha=1,
    ax=ax,
    zorder=2,
)

# Small negative bottom so dots at y=0 aren't clipped
ax.set_ylim(bottom=-ax.get_ylim()[1] * 0.02)
ax.patch.set_visible(False)
ax.ticklabel_format(axis="y", style="sci", scilimits=(0, 0))

# Align y=0 on both axes: give ax2 the same negative fraction below 0
ax1_bot, ax1_top = ax.get_ylim()
frac_below = -ax1_bot / (ax1_top - ax1_bot)

ax.spines[["top", "right", "left", "bottom"]].set_visible(False)
ax.axhline(0, color="black", linewidth=2, zorder=1)

ax.yaxis.grid(True, which="major", linestyle=":")
ax.set_axisbelow(True)
ax.set_xlabel("Pseudo-perplexity", fontsize=fontsize)
ax.set_ylabel("Purified band", fontsize=fontsize)

# Add a triangle convex hull of all points in the plot
from scipy.spatial import ConvexHull
points = tmp[[x, y]].values
hull = ConvexHull(points)
hull_points = points[hull.vertices]
ax.fill(hull_points[:, 0], hull_points[:, 1], color="lightgray", alpha=0.2, zorder=0, edgecolor="none")

# Draw hypotenuse line extended to plot limits
# Find the two key vertices: highest expression, and lowest expression (highest ppl)
hull_pts = points[hull.vertices]
# Vertex with highest y (expression)
v_high = hull_pts[hull_pts[:, 1].argmax()]
# Vertex with lowest y and highest x (lowest expression, highest ppl)
low_y_pts = hull_pts[hull_pts[:, 1] == hull_pts[:, 1].min()]
v_low = low_y_pts[low_y_pts[:, 0].argmax()]

# Compute slope and extend line to plot limits
slope = (v_high[1] - v_low[1]) / (v_high[0] - v_low[0])
intercept = v_high[1] - slope * v_high[0]
xlims = ax.get_xlim()
x_line = np.array(xlims)
y_line = slope * x_line + intercept
ax.plot(x_line, y_line, color="gray", linestyle="--", linewidth=1.5, zorder=1)

# Create custom legend with color (plate) and corresponding ppl mean
legend_elements = [plt.Line2D([0], [0], marker='o', color='w', markerfacecolor=plate_colors[plate], markersize=8, label=f"{plate} ({bar_centers[plate]:.2f})") for plate in plate_labels.values()]
ax.legend(handles=legend_elements, frameon=False, title="Plate (avg. PPL)", loc="upper right", fontsize=fontsize, title_fontsize=fontsize)

x = "AMPLIFY-C_pseudo_ppl"
y = "lysate band"

fontsize=16

# Prepare scatter data
tmp = data[data["plate"] != 3].copy()
tmp = tmp[~tmp[x].isna() & ~tmp[y].isna()]
tmp["plate"] = tmp["plate"].map(plate_labels)

# Prepare bar data: total lysate band per plate
bar_data = tmp.groupby("plate")[y].sum()
# Bar x-centers: mean pseudo-perplexity per plate
bar_centers = tmp.groupby("plate")[x].mean()

fig, ax = plt.subplots(figsize=(6, 4))

# Scatter on primary y-axis (foreground)
sns.scatterplot(
    tmp,
    x=x,
    y=y,
    hue="plate",
    size="length",
    palette=palette,
    alpha=1,
    ax=ax,
    zorder=2,
)

# Small negative bottom so dots at y=0 aren't clipped
ax.set_ylim(bottom=-ax.get_ylim()[1] * 0.02)
ax.patch.set_visible(False)
ax.ticklabel_format(axis="y", style="sci", scilimits=(0, 0))

# Align y=0 on both axes: give ax2 the same negative fraction below 0
ax1_bot, ax1_top = ax.get_ylim()
frac_below = -ax1_bot / (ax1_top - ax1_bot)

ax.spines[["top", "right", "left", "bottom"]].set_visible(False)
ax.axhline(0, color="black", linewidth=2, zorder=1)

ax.yaxis.grid(True, which="major", linestyle=":")
ax.set_axisbelow(True)
ax.set_xlabel("Pseudo-perplexity", fontsize=fontsize)
ax.set_ylabel("lysate band", fontsize=fontsize)

# Add a triangle convex hull of all points in the plot
from scipy.spatial import ConvexHull
points = tmp[[x, y]].values
hull = ConvexHull(points)
hull_points = points[hull.vertices]
ax.fill(hull_points[:, 0], hull_points[:, 1], color="lightgray", alpha=0.2, zorder=0, edgecolor="none")

# Draw hypotenuse line extended to plot limits
# Find the two key vertices: highest expression, and lowest expression (highest ppl)
hull_pts = points[hull.vertices]
# Vertex with highest y (expression)
v_high = hull_pts[hull_pts[:, 1].argmax()]
# Vertex with lowest y and highest x (lowest expression, highest ppl)
low_y_pts = hull_pts[hull_pts[:, 1] == hull_pts[:, 1].min()]
v_low = low_y_pts[low_y_pts[:, 0].argmax()]

# Compute slope and extend line to plot limits
slope = (v_high[1] - v_low[1]) / (v_high[0] - v_low[0])
intercept = v_high[1] - slope * v_high[0]
xlims = ax.get_xlim()
x_line = np.array(xlims)
y_line = slope * x_line + intercept
ax.plot(x_line, y_line, color="gray", linestyle="--", linewidth=1.5, zorder=1)

# Create custom legend with color (plate) and corresponding ppl mean
legend_elements = [plt.Line2D([0], [0], marker='o', color='w', markerfacecolor=plate_colors[plate], markersize=8, label=f"{plate} ({bar_centers[plate]:.2f})") for plate in plate_labels.values()]
ax.legend(handles=legend_elements, frameon=False, title="Plate (avg. PPL)", loc="upper right", fontsize=fontsize, title_fontsize=fontsize)

plt.tight_layout()
plt.savefig("figures/grouped_pseudo_ppl_vs_lysate_band.png", dpi=300)
plt.show()

plt.tight_layout()
plt.savefig("figures/grouped_pseudo_ppl_vs_purified_band.png", dpi=300)
plt.show()

# %% [code]
x = "AMPLIFY-C_pseudo_ppl"
y = "lysate band"

fontsize=16

fig, axes = plt.subplots(figsize=(12, 4.5), ncols=2, sharex=True)

for i, y in enumerate(["lysate band", "purified band"]):
    ax = axes[i]
    # Prepare scatter data
    tmp = data[data["plate"] != 3].copy()
    tmp = tmp[~tmp[x].isna() & ~tmp[y].isna()]
    tmp["plate"] = tmp["plate"].map(plate_labels)

    # Prepare bar data: total lysate band per plate
    bar_data = tmp.groupby("plate")[y].sum()
    # Bar x-centers: mean pseudo-perplexity per plate
    bar_centers = tmp.groupby("plate")[x].mean()

    # Scatter on primary y-axis (foreground)
    sns.scatterplot(
        tmp,
        x=x,
        y=y,
        hue="plate",
        size="length",
        palette=palette,
        alpha=1,
        ax=ax,
        zorder=2,
    )

    # Small negative bottom so dots at y=0 aren't clipped
    ax.set_ylim(bottom=-ax.get_ylim()[1] * 0.02)
    ax.patch.set_visible(False)
    ax.ticklabel_format(axis="y", style="sci", scilimits=(0, 0))

    # Align y=0 on both axes: give ax2 the same negative fraction below 0
    ax1_bot, ax1_top = ax.get_ylim()
    frac_below = -ax1_bot / (ax1_top - ax1_bot)

    ax.spines[["top", "right", "left", "bottom"]].set_visible(False)
    ax.axhline(0, color="black", linewidth=2, zorder=1)

    ax.yaxis.grid(True, which="major", linestyle=":")
    ax.set_axisbelow(True)
    ax.set_xlabel("Perplexity", fontsize=fontsize)
    ax.set_ylabel(y.title(), fontsize=fontsize)

    # Add a triangle convex hull of all points in the plot
    from scipy.spatial import ConvexHull
    points = tmp[[x, y]].values
    hull = ConvexHull(points)
    hull_points = points[hull.vertices]
    ax.fill(hull_points[:, 0], hull_points[:, 1], color="lightgray", alpha=0.2, zorder=0, edgecolor="none")

    # Draw hypotenuse line extended to plot limits
    # Find the two key vertices: highest expression, and lowest expression (highest ppl)
    hull_pts = points[hull.vertices]
    # Vertex with highest y (expression)
    v_high = hull_pts[hull_pts[:, 1].argmax()]
    # Vertex with lowest y and highest x (lowest expression, highest ppl)
    low_y_pts = hull_pts[hull_pts[:, 1] == hull_pts[:, 1].min()]
    v_low = low_y_pts[low_y_pts[:, 0].argmax()]

    # Compute slope and extend line to plot limits
    slope = (v_high[1] - v_low[1]) / (v_high[0] - v_low[0])
    intercept = v_high[1] - slope * v_high[0]
    xlims = ax.get_xlim()
    x_line = np.array(xlims)
    y_line = slope * x_line + intercept
    ax.plot(x_line, y_line, color="gray", linestyle="--", linewidth=1.5, zorder=1)

    # Create custom legend with color (plate) and corresponding ppl mean
    legend_elements = [plt.Line2D([0], [0], marker='o', color='w', markerfacecolor=plate_colors[plate], markersize=8, label=f"{plate} ({bar_centers[plate]:.2f})") for plate in plate_labels.values()]
    if i == 1:
        ax.legend(handles=legend_elements, frameon=False, title="Plate (avg. PPL)", loc="upper right", fontsize=fontsize, title_fontsize=fontsize)
    else:
        ax.legend().remove()

plt.tight_layout()
plt.savefig("figures/grouped_pseudo_ppl_vs_bands.png", dpi=300)
plt.show()

# %% [code]
tmp = data.groupby("plate").sum(numeric_only=True).reset_index().copy()

metrics = ["lysate band", "purified band"]
fig, ax = plt.subplots(figsize=(12, 5), ncols=len(metrics))
for i, metric in enumerate(metrics):
    ax[i].bar(tmp["plate"], tmp[metric], color="#0085ff", alpha=0.4, lw=0.1)
    ax[i].set_title(f"Total {metric} volume")
    ax[i].set_xlabel("Plate")
    ax[i].set_ylabel(f"Total {metric}")
    ax[i].set_xticks(tmp["plate"])
    ax[i].grid(axis="y", linestyle=":", alpha=0.7)
    
    ax[i].spines[["top", "right", "left"]].set_visible(False)
    ax[i].spines["bottom"].set_linewidth(2)
    
plt.tight_layout()
# plt.savefig(FIG_DIR / "total_bands_per_plate.png", dpi=300)
plt.show()

# %% [code]
result_columns = ["lysate band", "purified band", "mg/mL",] #'proline (EIC)', 'lysine (EIC)', 'indanone (245 nm)', 'resorcinol (285 nm)'
properties_to_plot = [
    "AMPLIFY_c_red",
    'AMPLIFY-C_pseudo_ppl',
    'aromaticity',
    'charge_at_pH7',
    'esm_if_nll',
    'flexibility_mean',
    'fraction_disordered_08',
    'gravy',
    'instability_index',
    'mean_plddt',
    'mean_sasa',
    'mol_ext_oxidized',
    'mol_ext_reduced',
    'molecular_weight',
    'mpnn_ll_soluble',
    'sa_repeat_ratio',
    'shannon_entropy',
    'shannon_entropy_normalized',
    'sp_likelihood',
    'ss_helix',
    'ss_sheet',
    'ss_turn',
    "kDa",
 ]

# %% [code]
# Filter Valid Pairs
valid_pairs = []
min_corr_threshold = 0.2

for metric2 in result_columns:
    for metric1 in properties_to_plot:
        # Get data temporarily to check correlation
        temp_df = data[[metric1, metric2]].dropna()

        if not temp_df.empty:
            try:
                p_corr, _ = pearsonr(list(temp_df[metric1].astype(float)), list(temp_df[metric2].astype(float)))
                s_corr, _ = spearmanr(list(temp_df[metric1].astype(float)), list(temp_df[metric2].astype(float)))
            except Exception as e:
                print(e)
                print(f"Skipping correlation for {metric1} vs. {metric2} due to non-numeric data.")
                continue

            # 2. Store only if it meets the criteria
            if abs(s_corr) >= min_corr_threshold:
                valid_pairs.append((metric1, metric2, p_corr, s_corr))

# %% [code]
# Create Grid & Plot
num_plots = len(valid_pairs)

if num_plots > 0:
    cols = 4
    rows = math.ceil(num_plots / cols)

    # Create the figure with tight sizing
    fig, axes = plt.subplots(rows, cols, figsize=(4 * cols, 3.5 * rows))
    axes = axes.flatten()  # Flatten for easy indexing

    print(f"Plotting {num_plots} valid correlations.")

    for i, (metric1, metric2, p_corr, s_corr) in enumerate(valid_pairs):
        ax=axes[i]
        sns.scatterplot(
            data,
            x=metric1,
            y=metric2,
            hue="plate",
            palette="crest",
            alpha=1,
            ax=ax,
            s=10,
        )

        # Write on plot
        ax.text(0.6, 0.90, f"p={p_corr:.2f}", transform=ax.transAxes, fontsize=10, verticalalignment='top')
        ax.text(0.8, 0.90, f"s={s_corr:.2f}", transform=ax.transAxes, fontsize=10, verticalalignment='top')

        ax.tick_params(left=False)
        ax.spines[["top", "right"]].set_visible(False)
        
        ax.set_xlabel(metric1, fontsize=10)
        ax.set_ylabel(metric2, fontsize=10)
        
        # Hide legend
        ax.get_legend().remove()
        

    # Hide any remaining empty axes (if grid count > plot count)
    for j in range(i + 1, len(axes)):
        axes[j].set_visible(False)

    plt.tight_layout()
    plt.show()
else:
    print("No pairs met the correlation threshold.")

# %% [code]
# Extract sequences from plate 3 that have expression
seq3 = data[(data["plate"] == 3) & ((data["lysate band"] > 0) | (data["purified band"] > 0))][["id", "sequence"]]

# %% [code]
ec = pd.read_csv("$HOME/enzymes/data/uniprotkb_ec_1_14_11_2025_10_20.csv")

# Write ec to fasta file
with open("../data/uniprotkb_ec_1_14_11_2025_10_20.fasta", "w") as f:
    for _, row in ec.iterrows():
        f.write(f">{row['name']}\n{row['sequence']}\n")

# %% [code]
# Write seq3 to fasta file
with open("../data/metagenomic_selected/expressed_metagenomic.fasta", "w") as f:
    for _, row in seq3.iterrows():
        f.write(f">{row['id']}\n{row['sequence']}\n")

# %% [code]
rmsd = pd.read_parquet("$HOME/enzymes/selection_logs/TO_SEND/dataframes/pairwise_rmsd_scores.parquet")

# %% [code]
plate3_sprot = pd.read_csv("$HOME/enzymes/data/metagenomic_selected/expressed3_uniprot_sprot_mmseqs.csv")
plate3_trembl = pd.read_csv("$HOME/enzymes/data/metagenomic_selected/expressed3_uniprot_trembl_mmseqs.csv")

plate3_mmseqs = pd.concat([plate3_sprot, plate3_trembl], ignore_index=True)
plate3_mmseqs = plate3_mmseqs.groupby("query_id")["max_pident"].max().sort_values(ascending=False).reset_index()
expressed_plate3 = data.merge(plate3_mmseqs, left_on="id", right_on="query_id", how="inner")

# %% [code]
expressed_plate3 = expressed_plate3[['query_id', 'max_pident', 'mg/mL', 'lysate band', 'purified band',
       'AMPLIFY_c_red',
       'AMPLIFY-C_pseudo_ppl',
       'AMPLIFY_350M_intra_similarity',
       'length',
       'aromaticity',
       'charge_at_pH7', 'charge_at_pH7.5', 'charge_at_pH8',
       'esm_if_nll', 'flexibility_mean', 'fraction_disordered_05',
       'fraction_disordered_08', 'fraction_disordered_09', 'gravy',
       'instability_index', 'isoelectric_point', 'mean_plddt', 'mean_sasa',
       'mol_ext_oxidized', 'mol_ext_reduced', 'molecular_weight',
       'mpnn_ll_soluble', 'non_standard_aa_count',
       'plate', 'sa_repeat_ratio', 'shannon_entropy',
       'shannon_entropy_normalized', 'sp_likelihood', 'ss_helix', 'ss_sheet',
       'ss_turn', 'panel', 'protein band', 'proline (EIC)', 'lysine (EIC)',
       'indanone (245 nm)', 'resorcinol (285 nm)',
       'AMPLIFY_benchmark_pseudo_ppl',
       'kDa',]]

# %% [code]
expressed_plate3["max band"] = expressed_plate3[["lysate band", "purified band"]].max(axis=1)

# %% [code]
flashy = expressed_plate3[(expressed_plate3["max_pident"] < 50) ].sort_values("max band", ascending=False)
flashy = flashy[["query_id", "max_pident", "lysate band", "purified band", "AMPLIFY-C_pseudo_ppl"]].copy()

# %% [code]
flashy

# %% [code]
flashy.to_csv("../data/metagenomic_selected/flashy_plate3.csv", index=False)

# %% [code]
df_selected = pd.read_parquet("$HOME/enzymes/selection_logs/final/data_metagenomic.parquet")

# %% [code]
df_selected = df_selected[["metagenomic_id", "closest_embedding_uniprot_id", "inter_score"]]

plate3 = data[data["plate"] == 3].merge(df_selected, left_on="id", right_on="metagenomic_id", how="left")
plate3 = plate3[['id', 'sequence',
        "closest_embedding_uniprot_id",
        "inter_score",
        "max_band",
       'AMPLIFY-C_embedding',
       'AMPLIFY-C_pseudo_ppl',
       'AMPLIFY_benchmark_pseudo_ppl', 'protein band', 'mg/mL', 'lysate band', 'purified band']]

plate3 = plate3.merge(data[["id", "max_band", "purified band", "lysate band", "mg/mL"]], left_on="closest_embedding_uniprot_id", right_on="id", how="left", suffixes=("", "_closest_uniprot"))

plate3["purified band"].fillna(0, inplace=True)
plate3["lysate band"].fillna(0, inplace=True)
plate3["max_band"].fillna(0, inplace=True)

# %% [code]
# Plot correlation between purified band and closest uniprot purified band
plt.figure(figsize=(6, 5))
sns.scatterplot(
    plate3,
    x="purified band_closest_uniprot",
    y="purified band",
    alpha=1,
    s=10,
)

def weighted_pearson(x, y, w):
    w = w / w.sum()
    mx = np.average(x, weights=w)
    my = np.average(y, weights=w)
    cov = np.sum(w * (x - mx) * (y - my))
    sx = np.sqrt(np.sum(w * (x - mx)**2))
    sy = np.sqrt(np.sum(w * (y - my)**2))
    return cov / (sx * sy)

p = pearsonr(plate3["purified band_closest_uniprot"].dropna(), plate3["purified band"].dropna())
s = spearmanr(plate3["purified band_closest_uniprot"].dropna(), plate3["purified band"].dropna())

plt.text(0.6, 0.95, f"Pearson r: {p[0]:.2f}", transform=plt.gca().transAxes, fontsize=12, verticalalignment='top')
plt.xlabel("Closest Uniprot purified band")
plt.ylabel("Plate 3 purified band")
plt.title("Correlation between Plate 3 purified band and closest Uniprot purified band")
plt.grid(axis="y", linestyle=":", alpha=0.7)
plt.tight_layout()
# plt.savefig(FIG_DIR / "plate3_closest_uniprot_purified_band_correlation.png", dpi=300)
plt.show()

# %% [code]
# Plot correlation between purified band and closest uniprot purified band
plt.figure(figsize=(6, 5))
sns.scatterplot(
    plate3,
    x="lysate band_closest_uniprot",
    y="lysate band",
    alpha=1,
    s=10,
)

def weighted_pearson(x, y, w):
    w = w / w.sum()
    mx = np.average(x, weights=w)
    my = np.average(y, weights=w)
    cov = np.sum(w * (x - mx) * (y - my))
    sx = np.sqrt(np.sum(w * (x - mx)**2))
    sy = np.sqrt(np.sum(w * (y - my)**2))
    return cov / (sx * sy)

p = pearsonr(plate3["lysate band_closest_uniprot"].dropna(), plate3["lysate band"].dropna())
s = spearmanr(plate3["lysate band_closest_uniprot"].dropna(), plate3["lysate band"].dropna())

plt.text(0.6, 0.95, f"Pearson r: {p[0]:.2f}", transform=plt.gca().transAxes, fontsize=12, verticalalignment='top')
plt.xlabel("Closest Uniprot lysate band")
plt.ylabel("Plate 3 lysate band")
plt.title("Correlation between Plate 3 lysate band and closest Uniprot lysate band")
plt.grid(axis="y", linestyle=":", alpha=0.7)
plt.tight_layout()
# plt.savefig(FIG_DIR / "plate3_closest_uniprot_purified_band_correlation.png", dpi=300)
plt.show()

# %% [code]
# Plot correlation between purified band and closest uniprot purified band
plt.figure(figsize=(6, 5))
sns.scatterplot(
    plate3,
    x="max_band_closest_uniprot",
    y="max_band",
    alpha=1,
    s=10,
)

def weighted_pearson(x, y, w):
    w = w / w.sum()
    mx = np.average(x, weights=w)
    my = np.average(y, weights=w)
    cov = np.sum(w * (x - mx) * (y - my))
    sx = np.sqrt(np.sum(w * (x - mx)**2))
    sy = np.sqrt(np.sum(w * (y - my)**2))
    return cov / (sx * sy)

p = pearsonr(plate3["max_band_closest_uniprot"].dropna(), plate3["max_band"].dropna())
s = spearmanr(plate3["max_band_closest_uniprot"].dropna(), plate3["max_band"].dropna())

plt.text(0.6, 0.95, f"Pearson r: {p[0]:.2f}", transform=plt.gca().transAxes, fontsize=12, verticalalignment='top')
plt.xlabel("Closest Uniprot max_band")
plt.ylabel("Plate 3 max_band")
plt.title("Correlation between Plate 3 max_band and closest Uniprot max_band")
plt.grid(axis="y", linestyle=":", alpha=0.7)
plt.tight_layout()
# plt.savefig(FIG_DIR / "plate3_closest_uniprot_purified_band_correlation.png", dpi=300)
plt.show()

# %% [code]
plate3["metagenomic_expressed"] = (plate3["lysate band"] > 0) | (plate3["purified band"] > 0)
plate3["closest_uniprot_expressed"] = (plate3["lysate band_closest_uniprot"] > 0) | (plate3["purified band_closest_uniprot"] > 0)

# %% [markdown]
## Figures

# %% [code]
y = "max_band"
x = "AMPLIFY-C_pseudo_ppl"

tmp = data[data["plate"] == 3].copy()
tmp = tmp[~tmp[x].isna() & ~tmp[y].isna()]

fig, ax = plt.subplots(figsize=(10, 6))
sns.scatterplot(
    tmp,
    x=x,
    y=y,
    hue="plate",
    size="length",
    palette="crest",
    alpha=1,
    ax=ax,
)

ax.plot([tmp[x].min(), tmp[x].max()], [tmp[y].min(), tmp[y].max()], color='red', linestyle='--', alpha=0.5)

# Compute correlation
pearson_corr, pearson_pval = pearsonr(tmp[x], tmp[y])
spearman_corr, spearman_pval = spearmanr(tmp[x], tmp[y])

# Write on plot
# ax.text(0.05, 0.95, f"Pearson r: {pearson_corr:.2f} (p={pearson_pval:.2e})", transform=ax.transAxes, fontsize=12, verticalalignment='top')
ax.text(0.5, 0.95, r"Spearman $\rho$: " + f"{spearman_corr:.2f} (p={spearman_pval:.2e})", transform=ax.transAxes, fontsize=10, verticalalignment='center')

# ax.tick_params(left=False)
ax.legend(title="", fontsize=10, title_fontsize=12, frameon=False)
ax.spines[["top", "right", "left"]].set_visible(False)
ax.spines["bottom"].set_linewidth(2)
ax.yaxis.grid(True, which="major", linestyle=":")
ax.set_axisbelow(True)

# %% [code]
x = "AMPLIFY_benchmark_pseudo_ppl"
y = "AMPLIFY-C_pseudo_ppl"

tmp = data #[data["plate"] != 3].copy()

fig, ax = plt.subplots(figsize=(10, 6))
sns.scatterplot(
    tmp,
    x=x,
    y=y,
    hue="plate",
    size="length",
    palette="crest",
    alpha=1,
    ax=ax,
)

ax.plot([tmp[x].min(), tmp[x].max()], [tmp[y].min(), tmp[y].max()], color='red', linestyle='--', alpha=0.5)

# Compute correlation
pearson_corr, pearson_pval = pearsonr(tmp[x], tmp[y])
spearman_corr, spearman_pval = spearmanr(tmp[x], tmp[y])

# Write on plot
# ax.text(0.05, 0.95, f"Pearson r: {pearson_corr:.2f} (p={pearson_pval:.2e})", transform=ax.transAxes, fontsize=12, verticalalignment='top')
ax.text(0.5, 0.95, r"Spearman $\rho$: " + f"{spearman_corr:.2f} (p={spearman_pval:.2e})", transform=ax.transAxes, fontsize=10, verticalalignment='center')

# ax.tick_params(left=False)
ax.spines[["top", "right", "left"]].set_visible(False)
ax.spines["bottom"].set_linewidth(2)
ax.yaxis.grid(True, which="major", linestyle=":")
ax.set_axisbelow(True)

# %% [code]
x = "AMPLIFY_benchmark_pseudo_ppl"
y = "AMPLIFY-C_pseudo_ppl"

tmp = data[data["plate"] == 3].copy()
tmp = tmp[~tmp[x].isna() & ~tmp[y].isna()]

fig, ax = plt.subplots(figsize=(10, 6))
sns.scatterplot(
    tmp,
    x=x,
    y=y,
    hue="plate",
    size="length",
    palette="crest",
    alpha=1,
    ax=ax,
)

# Plot y = x
ax.plot([tmp[x].min(), tmp[x].max()], [tmp[y].min(), tmp[y].max()], color='red', linestyle='--', alpha=0.5)

# Compute correlation
pearson_corr, pearson_pval = pearsonr(tmp[x], tmp[y])
spearman_corr, spearman_pval = spearmanr(tmp[x], tmp[y])

# Write on plot
# ax.text(0.05, 0.95, f"Pearson r: {pearson_corr:.2f} (p={pearson_pval:.2e})", transform=ax.transAxes, fontsize=12, verticalalignment='top')
ax.text(0.5, 0.95, r"Spearman $\rho$: " + f"{spearman_corr:.2f} (p={spearman_pval:.2e})", transform=ax.transAxes, fontsize=10, verticalalignment='center')

# ax.tick_params(left=False)
ax.spines[["top", "right", "left"]].set_visible(False)
ax.spines["bottom"].set_linewidth(2)
ax.yaxis.grid(True, which="major", linestyle=":")

ax.set_axisbelow(True)

# %% [code]
# Get 4 plate colors from the crest palette
plate_colors = sns.color_palette("crest", n_colors=4)

pairs = []
for activity in ["lysate band", "purified band"]:
    for model in ["AMPLIFY_benchmark_pseudo_ppl", "AMPLIFY-C_pseudo_ppl"]:
        for plate in range(1, 5):
            pairs.append((model, plate, activity))
            
            
# Create Grid & Plot
num_plots = len(pairs)

cols = 4
rows = math.ceil(num_plots / cols)

# Create the figure with tight sizing
fig, axes = plt.subplots(rows, cols, figsize=(4 * cols, 3.5 * rows), sharex=True, sharey=True)
axes = axes.flatten()  # Flatten for easy indexing

print(f"Plotting {num_plots} plots.")

for i, (model, plate, activity) in enumerate(pairs):
    ax=axes[i]
    
    x = model
    y = activity

    tmp = data[data["plate"] == plate].copy()
    tmp = tmp[~tmp[x].isna() & ~tmp[y].isna()]

    sns.scatterplot(
        tmp,
        x=x,
        y=y,
        color=plate_colors[plate - 1],
        # hue="purified band",
        # palette="crest",
        size="length",
        alpha=1,
        ax=ax,
    )

    # Compute correlation
    pearson_corr, pearson_pval = pearsonr(tmp[x], tmp[y])
    spearman_corr, spearman_pval = spearmanr(tmp[x], tmp[y])

    # Write on plot
    # ax.text(0.05, 0.95, f"Pearson r: {pearson_corr:.2f} (p={pearson_pval:.2e})", transform=ax.transAxes, fontsize=12, verticalalignment='top')
    ax.text(0.5, 0.95, r"$\rho$: " + f"{spearman_corr:.2f}", transform=ax.transAxes, fontsize=10, verticalalignment='center')

    # ax.tick_params(left=False)
    ax.spines[["top", "right", "left"]].set_visible(False)
    ax.spines["bottom"].set_linewidth(2)
    ax.yaxis.grid(True, which="major", linestyle=":")
    ax.set_axisbelow(True)
    
    ax.set_ylabel(y + " - " + x.replace("AMPLIFY_", "").replace("_pseudo_ppl", "").replace("_extended", "").replace("_", " "), fontsize=10)
    ax.set_xlabel("")
    
    ax.legend_.remove()  # Remove legend for cleaner look
    

# Hide any remaining empty axes (if grid count > plot count)
for j in range(i + 1, len(axes)):
    axes[j].set_visible(False)

plt.tight_layout()
plt.show()

# %% [code]
x = "purified band"
y = "lysate band"

tmp = data #[data["plate"] == 3].copy()
tmp = tmp[~tmp[x].isna() & ~tmp[y].isna()]

fig, ax = plt.subplots(figsize=(10, 6))
sns.scatterplot(
    tmp,
    x=x,
    y=y,
    hue="AMPLIFY-C_pseudo_ppl",
    size="length",
    palette="crest",
    alpha=1,
    ax=ax,
)

# Plot y = x
ax.plot([tmp[x].min(), tmp[x].max()], [tmp[y].min(), tmp[y].max()], color='red', linestyle='--', alpha=0.5)

# Compute correlation
pearson_corr, pearson_pval = pearsonr(tmp[x], tmp[y])
spearman_corr, spearman_pval = spearmanr(tmp[x], tmp[y])

# Write on plot
# ax.text(0.05, 0.95, f"Pearson r: {pearson_corr:.2f} (p={pearson_pval:.2e})", transform=ax.transAxes, fontsize=12, verticalalignment='top')
ax.text(0.5, 0.95, r"Spearman $\rho$: " + f"{spearman_corr:.2f} (p={spearman_pval:.2e})", transform=ax.transAxes, fontsize=10, verticalalignment='center')

# ax.tick_params(left=False)
ax.spines[["top", "right", "left"]].set_visible(False)
ax.spines["bottom"].set_linewidth(2)
ax.yaxis.grid(True, which="major", linestyle=":")
ax.set_axisbelow(True)

# %% [code]
x = "AMPLIFY_benchmark_pseudo_ppl"
y = "lysate band"

tmp = data[data["plate"] == 3].copy()
tmp = tmp[~tmp[x].isna() & ~tmp[y].isna()]

fig, ax = plt.subplots(figsize=(10, 6))
sns.scatterplot(
    tmp,
    x=x,
    y=y,
    hue="purified band",
    size="length",
    palette="crest",
    alpha=1,
    ax=ax,
)

# Compute correlation
pearson_corr, pearson_pval = pearsonr(tmp[x], tmp[y])
spearman_corr, spearman_pval = spearmanr(tmp[x], tmp[y])

# Write on plot
# ax.text(0.05, 0.95, f"Pearson r: {pearson_corr:.2f} (p={pearson_pval:.2e})", transform=ax.transAxes, fontsize=12, verticalalignment='top')
ax.text(0.5, 0.95, r"Spearman $\rho$: " + f"{spearman_corr:.2f} (p={spearman_pval:.2e})", transform=ax.transAxes, fontsize=10, verticalalignment='center')

# ax.tick_params(left=False)
ax.spines[["top", "right", "left"]].set_visible(False)
ax.spines["bottom"].set_linewidth(2)
ax.yaxis.grid(True, which="major", linestyle=":")
ax.set_axisbelow(True)

# %% [code]
plate_labels

# %% [code]
FIG_DIR

# %% [code]
x = "AMPLIFY-C_pseudo_ppl"
y = "lysate band"

tmp = data[data["plate"] != 3].copy()
tmp = tmp[~tmp[x].isna() & ~tmp[y].isna()]

fig, ax = plt.subplots(figsize=(10, 6))
sns.scatterplot(
    tmp,
    x=x,
    y=y,
    hue="plate",
    size="length",
    palette="crest",
    alpha=1,
    ax=ax,
)

# Compute correlation
pearson_corr, pearson_pval = pearsonr(tmp[x], tmp[y])
spearman_corr, spearman_pval = spearmanr(tmp[x], tmp[y])

# Write on plot
# ax.text(0.05, 0.95, f"Pearson r: {pearson_corr:.2f} (p={pearson_pval:.2e})", transform=ax.transAxes, fontsize=12, verticalalignment='top')
ax.text(0.5, 0.95, r"Spearman $\rho$: " + f"{spearman_corr:.2f} (p={spearman_pval:.2e})", transform=ax.transAxes, fontsize=10, verticalalignment='center')

# ax.tick_params(left=False)
ax.spines[["top", "right", "left"]].set_visible(False)
ax.spines["bottom"].set_linewidth(2)
ax.yaxis.grid(True, which="major", linestyle=":")
ax.set_axisbelow(True)

# %% [code]
x = "AMPLIFY-C_pseudo_ppl"
y = "purified band"

tmp = data[data["plate"] != 3].copy()
tmp = tmp[~tmp[x].isna() & ~tmp[y].isna()]

fig, ax = plt.subplots(figsize=(10, 6))
sns.scatterplot(
    tmp,
    x=x,
    y=y,
    hue="plate",
    size="length",
    palette="crest",
    alpha=1,
    ax=ax,
)

# Compute correlation
pearson_corr, pearson_pval = pearsonr(tmp[x], tmp[y])
spearman_corr, spearman_pval = spearmanr(tmp[x], tmp[y])

# Write on plot
# ax.text(0.05, 0.95, f"Pearson r: {pearson_corr:.2f} (p={pearson_pval:.2e})", transform=ax.transAxes, fontsize=12, verticalalignment='top')
ax.text(0.5, 0.95, r"Spearman $\rho$: " + f"{spearman_corr:.2f} (p={spearman_pval:.2e})", transform=ax.transAxes, fontsize=10, verticalalignment='center')

# ax.tick_params(left=False)
ax.spines[["top", "right", "left"]].set_visible(False)
ax.spines["bottom"].set_linewidth(2)
ax.yaxis.grid(True, which="major", linestyle=":")
ax.set_axisbelow(True)

# %% [code]
x = "AMPLIFY_benchmark_pseudo_ppl"
y = "lysate band"

tmp = data[data["plate"] != 3].copy()

fig, ax = plt.subplots(figsize=(10, 6))
sns.scatterplot(
    tmp,
    x=x,
    y=y,
    hue="plate",
    size="length",
    palette="crest",
    alpha=1,
    ax=ax,
)

# Compute correlation
pearson_corr, pearson_pval = pearsonr(tmp[x], tmp[y])
spearman_corr, spearman_pval = spearmanr(tmp[x], tmp[y])

# Write on plot
# ax.text(0.05, 0.95, f"Pearson r: {pearson_corr:.2f} (p={pearson_pval:.2e})", transform=ax.transAxes, fontsize=12, verticalalignment='top')
ax.text(0.5, 0.95, r"Spearman $\rho$: " + f"{spearman_corr:.2f} (p={spearman_pval:.2e})", transform=ax.transAxes, fontsize=10, verticalalignment='center')

# ax.tick_params(left=False)
ax.spines[["top", "right", "left"]].set_visible(False)
ax.spines["bottom"].set_linewidth(2)
ax.yaxis.grid(True, which="major", linestyle=":")
ax.set_axisbelow(True)

# %% [code]
fig, ax = plt.subplots(figsize=(10, 6))
sns.scatterplot(
    data,
    x="AMPLIFY-C_pseudo_ppl",
    y="fraction_disordered_09",
    hue = "protein band",
    size="length",
    palette="crest",
    alpha=1,
    ax=ax,
)

# Compute correlation
pearson_corr, pearson_pval = pearsonr(data["AMPLIFY-C_pseudo_ppl"], data["fraction_disordered_09"])
spearman_corr, spearman_pval = spearmanr(data["AMPLIFY-C_pseudo_ppl"], data["fraction_disordered_09"])

# Write on plot
# ax.text(0.05, 0.95, f"Pearson r: {pearson_corr:.2f} (p={pearson_pval:.2e})", transform=ax.transAxes, fontsize=12, verticalalignment='top')
ax.text(0.5, 0.95, r"Spearman $\rho$: " + f"{spearman_corr:.2f} (p={spearman_pval:.2e})", transform=ax.transAxes, fontsize=10, verticalalignment='center')

# ax.tick_params(left=False)
ax.spines[["top", "right", "left"]].set_visible(False)
ax.spines["bottom"].set_linewidth(2)
ax.yaxis.grid(True, which="major", linestyle=":")
ax.set_axisbelow(True)

# %% [code]
def plot_kde_by_metric(data, metrics_to_plot, x_metric="AMPLIFY-C_pseudo_ppl", title="", figname = None):
    # Create Grid & Plot
    num_plots = len(metrics_to_plot)

    rows = 2
    cols = math.ceil(num_plots / rows)

    # Create the figure with tight sizing
    fig, axes = plt.subplots(rows, cols, figsize=(4 * cols, 3.5 * rows), sharey=True)
    axes = axes.flatten()  # Flatten for easy indexing

    for i, metric in enumerate(metrics_to_plot):
        ax=axes[i]
        ax2 = ax.twinx()  # Create a secondary y-axis for the histogram
        sns.kdeplot(
            data=data,
            x=x_metric,
            color="grey",
            fill=True,
            alpha=0.1,
            ax=ax2,
            common_norm=True,
            bw_adjust=0.5,
            warn_singular=False,
        )
        sns.kdeplot(
            data=data[data[metric]],
            x=x_metric,
            color="blue",
            fill=True,
            alpha=0.6,
            ax=ax,
            common_norm=True,
            bw_adjust=0.1,
            warn_singular=False,
        )
        # Plot count instead of density on the y-axis
        # sns.histplot(
        #     data=data[data[metric]],
        #     x=x_metric,
        #     bins=50,
        #     color="blue",
        #     alpha=0.4,
        #     lw=0.1,
        #     ax=ax,
        #     stat="count",
        #     binrange=(0, 20),
        # )

        # Write on plot
        ax.tick_params(left=False)
        ax.spines[["top", "right", "left"]].set_visible(False)
        
        ax2.tick_params(left=False)
        ax.set_yticks([])  # Hide y-ticks for the histogram
        ax2.set_yticks([])  # Hide y-ticks for the histogram
        ax2.spines[["top", "right", "left"]].set_visible(False)
        ax2.set_ylabel("")
        
        ax.set_xlabel(metric, fontsize=10)
        ax.set_ylabel("")
        
    # Hide any remaining empty axes (if grid count > plot count)
    for j in range(i + 1, len(axes)):
        axes[j].set_visible(False)
        
    plt.suptitle(title, fontsize=12)

    plt.tight_layout()
    if figname:
        plt.savefig(f"{figname}.pdf", dpi=300)
        plt.savefig(f"{figname}.png", dpi=300)
    plt.show()

# %% [code]
metrics_to_plot = []
thresholds = [0, 10, 1000, 2000, 3000]
for activity in ["lysate band", "purified band"]:
    for i in range(len(thresholds)):
        threshold = thresholds[i]
        threshold2 = thresholds[i+1] if i < len(thresholds)-1 else None
        if threshold2 == threshold:
            data[f"{activity}: {threshold}"] = (data[activity] == threshold)
            metrics_to_plot.append(f"{activity}: {threshold}")
        elif threshold2:
            data[f"{activity}: {threshold} - {threshold2}"] = (data[activity] >= threshold) & (data[activity] < threshold2)
            metrics_to_plot.append(f"{activity}: {threshold} - {threshold2}")
        else:
            data[f"{activity}: {threshold}"] = data[activity] >= threshold
            metrics_to_plot.append(f"{activity}: {threshold}")


# %% [code]
plot_kde_by_metric(data[data.plate != 3].copy(), metrics_to_plot, title="", figname="figures/ppl_by_activity_range") #Pseudo-perplexity (plates A, B, C)

# %% [code]
plot_kde_by_metric(data[data.plate != 3].copy(), metrics_to_plot, title=" (plates 1, 2, 4)")
plot_kde_by_metric(data[data.plate == 3].copy(), metrics_to_plot, title=" (plate 3)")
plot_kde_by_metric(data[data.plate == 4].copy(), metrics_to_plot, title=" (plate 4)")

# %% [code]
plot_kde_by_metric(data[data.plate != 3].copy(), metrics_to_plot, title=" (plates 1, 2, 4)", x_metric="AMPLIFY_benchmark_pseudo_ppl")
plot_kde_by_metric(data[data.plate == 3].copy(), metrics_to_plot, title=" (plate 3)", x_metric="AMPLIFY_benchmark_pseudo_ppl")
plot_kde_by_metric(data[data.plate == 4].copy(), metrics_to_plot, title=" (plate 4)", x_metric="AMPLIFY_benchmark_pseudo_ppl")

# %% [code]
fig, ax = plt.subplots(nrows=2, ncols=2, figsize=(12, 8), sharex=True, sharey=True)

ax = ax.flatten()
for i, plate in enumerate(range(1, 5)):
    plot_distribution(ax[i], data[data["plate"] == plate], "AMPLIFY-C_pseudo_ppl", title = f"PPL plate {i + 1}")

# %% [code]
data

# %% [code]
plate_colors

# %% [code]
sns.set_style("white", rc={"axes.facecolor": (0, 0, 0, 0)})
plt.rcParams["savefig.dpi"] = 300  # Set save DPI
plt.rcParams["figure.dpi"] = 150  # Set display DPI

tmp = data[data["plate"] != 3].copy()

plate_to_label = {1: "A", 2: "B", 4: "C"}

tmp["plate_label"] = tmp["plate"].map(plate_to_label)

fontsize = 8

plates = sorted(tmp["plate_label"].unique())
# 2. Compute averages and setup palette
averages = tmp.groupby("plate_label")["AMPLIFY-C_pseudo_ppl"].mean().to_dict()
# ordered_plates = sorted(averages, key=averages.get)  # low avg PPL on top
palette = plate_colors

# 4. Plot using FacetGrid
plt.figure(figsize=(6, 6))
sns.set_style("white", rc={"axes.facecolor": (0, 0, 0, 0)})

g = sns.FacetGrid(
    tmp,
    row="plate_label",
    row_order=plates,
    hue="plate_label",
    aspect=3.5,
    height=1,
    palette=palette,
)


g.set(xlim=(0, 18))
g.map(sns.kdeplot, "AMPLIFY-C_pseudo_ppl", clip_on=False, fill=True, alpha=1, linewidth=0.3, bw_adjust=0.3, edgecolor=None)
g.map(sns.kdeplot, "AMPLIFY-C_pseudo_ppl", clip_on=False, color="w", lw=1, bw_adjust=0.3)

# Add horizontal colored lines at y=0
def horiz_line(x, color, label):
    ax = plt.gca()
    ax.axhline(y=0, lw=0.3, color=color, alpha=0.5, clip_on=False)

g.map(horiz_line, "AMPLIFY-C_pseudo_ppl")

# Labeling each subplot with average PPL
def label(x, color, label):
    ax = plt.gca()
    avg = averages.get(label, 0)
    ax.text(1.01, 0, label, fontsize=fontsize, color="black", ha="left", va="center", transform=ax.transAxes)


g.map(label, "AMPLIFY-C_pseudo_ppl")

g.figure.subplots_adjust(hspace=-0.9)
g.set_titles("")
g.set(yticks=[], ylabel="")
g.despine(left=True)
# Set xtick font size and set the ticks every 5 units
for ax in g.axes[:, 0]:
    ax.set_xticks(range(0, 19, 5))
    ax.tick_params(axis='x', labelsize=fontsize - 2)

for ax in g.axes[:, 0]:
    ax.spines["bottom"].set_linewidth(0.5)

plt.xlabel("Perplexity distribution", fontsize=fontsize)
plt.savefig(f"figures/ridgeplot_ppl_UP_plates.png", dpi=300, bbox_inches="tight")
plt.show()

# %% [code]
plate_colors

# %% [code]
```
