## Supplementary material for "pLM representations unlock metagenomic space beyond homology": Code bundle: structure_labels.ipynb

```
# %% [markdown]
# Structure Label Figures for Paper

Generates two sets of figures:
1. **Standalone text labels** — white background panels with sequence name, pLDDT, and band activity (lysate/purified). Crop and place next to structure images.
2. **Annotated structure panels** — text overlaid directly on the structure images, ready for a figure grid.

# %% [code]
import os
import csv
import json
from pathlib import Path
import pandas as pd
import matplotlib.pyplot as plt
import matplotlib.image as mpimg
from PIL import Image
from PIL import ImageDraw, ImageFont
import numpy as np
from IPython.display import Image as IPImage, display

import seaborn as sns
# --- Paths ---
CSV_PATH = "$HOME/enzymes/data/metagenomic_selected/flashy_plate3.csv"
CIF_DIR = Path("$SCRATCH/proseqo/enzymes/metagenomic_output/openfold/dataframes")
IMG_DIR = Path("$SCRATCH/proseqo/enzymes/metagenomic_output/openfold/structure_images")
OUTPUT_DIR = Path("$SCRATCH/proseqo/enzymes/metagenomic_output/openfold/structure_labels")
OUTPUT_DIR.mkdir(parents=True, exist_ok=True)


# %% [code]
data = pd.read_csv("$HOME/enzymes/notebooks/data.csv")
data = data[data["plate"] == 3].copy()

# %% [code]
# --- Load CSV data ---
with open(CSV_PATH) as f:
    reader = csv.DictReader(f)
    rows = list(reader)

# --- Get pLDDT from confidence files (reuse logic from visualize_structures) ---
def get_best_plddt(seq_id: str) -> float | None:
    seq_dir = CIF_DIR / seq_id
    if not seq_dir.is_dir():
        return None
    best_score = -1
    for conf_file in seq_dir.rglob("*_confidences_aggregated.json"):
        with open(conf_file) as f:
            conf = json.load(f)
        if conf["avg_plddt"] > best_score:
            best_score = conf["avg_plddt"]
    return best_score if best_score > 0 else None

data["openfold_plddt"] = data["id"].apply(lambda x: get_best_plddt(x))

# %% [code]
fasta_path = "plate4_expressed.fasta"
with open(fasta_path, "a") as f:
    for id in data[((data["lysate band"] > 100) | (data["purified band"] > 100)) & (data["openfold_plddt"] < 80)]["id"]:
        print(id)
        # Write id, seq to fasta file
        seq = data[data["id"] == id]["sequence"].values[0]
        f.write(f">{id}\n{seq}\n")

# %% [code]
data[((data["lysate band"] > 100) | (data["purified band"] > 100)) & (data["openfold_plddt"] < 80)]

# %% [code]
data.columns

# %% [code]
df_selected = pd.read_parquet("$HOME/enzymes/selection_logs/final/data_metagenomic.parquet")

# %% [code]
data = data.merge(df_selected[["metagenomic_id", "inter_score"]], left_on="id", right_on="metagenomic_id", how="left")

# %% [code]


# %% [code]
# Filter list to only properties that exist in the final dataframe
data["Sequence similarity to EC 1.14.11"] = 0
properties_to_plot = [
    "Sequence similarity to EC 1.14.11",
    "inter_score",
    # "openfold_plddt",
    # "AMPLIFY-C_pseudo_ppl",
]

prop_labels = {
    "Sequence similarity to EC 1.14.11": "Sequence similarity to EC 1.14.11",
    "inter_score": "Embedding similarity to EC 1.14.11",
    "openfold_plddt": "Mean pLDDT (OpenFold)",
    "AMPLIFY-C_pseudo_ppl": "Pseudo-perplexity",
}

tmp = data

properties_to_plot = [prop for prop in properties_to_plot if prop in tmp.columns]

n_props = len(properties_to_plot)
n_cols = 1
n_rows = int(np.ceil(n_props / n_cols))

fig, axes = plt.subplots(n_rows, n_cols, figsize=(4, n_rows * 3), tight_layout=True)

for i, prop in enumerate(properties_to_plot):
    ax = axes.flat[i]
    data_to_plot = tmp.dropna(subset=[prop])

    sns.histplot(
        data=data_to_plot,
        x=prop,
        bins=30,
        stat="percent",
        alpha=0.4,
        lw=0.1,
        color="#0085ff",
        ax=ax,
    )

    ax.set_title(f"{prop_labels.get(prop, prop)}", fontsize=14)
    ax.set_xlabel("")
    ax.set_ylabel("")
    ax.set_ylabel("Proportion (%)", fontsize=12)
    ax.tick_params(left=False)
    ax.spines[["top", "right", "left"]].set_visible(False)
    ax.spines["bottom"].set_linewidth(2)
    ax.yaxis.grid(True, which="major", linestyle=":")
    ax.set_axisbelow(True)

# Hide any unused subplots
for j in range(i + 1, n_rows * n_cols):
    axes.flat[j].set_visible(False)

plt.tight_layout()
plt.savefig("figures/metagenomic similarities.png", dpi=300)
plt.show()

# %% [code]
color = (0.4044212904941176, 0.5346434904470588, 0.9320019126352941)

fontsize = 14

# Plot OpenFold pLDDT vs lysate/purified bands
plt.figure(figsize=(12, 5))
plt.subplot(1, 2, 1)
tmp = data[(data["lysate band"] > 0)]
tmpz = data[(data["lysate band"] == 0)]
plt.scatter(tmp["openfold_plddt"], tmp["lysate band"], color=color, alpha=1, s=80, edgecolors=None)
plt.scatter(tmpz["openfold_plddt"], tmpz["lysate band"], color="lightgray", alpha=0.8, s=80, edgecolors=None)
plt.xlabel("Mean pLDDT (OpenFold)", fontsize=fontsize)
plt.ylabel("Lysate band intensity", fontsize=fontsize)

plt.subplot(1, 2, 2)
tmp = data[(data["purified band"] > 0)]
tmpz = data[(data["purified band"] == 0)]
plt.scatter(tmp["openfold_plddt"], tmp["purified band"],  color=color, alpha=1, s=80, edgecolors=None)
plt.scatter(tmpz["openfold_plddt"], tmpz["purified band"], color="lightgray", alpha=0.8, s=80, edgecolors=None)
plt.xlabel("Mean pLDDT (OpenFold)", fontsize=fontsize)
plt.ylabel("Purified band intensity", fontsize=fontsize)

# Remove top and right spines
for ax in plt.gcf().axes:
    ax.spines["top"].set_visible(False)
    ax.spines["right"].set_visible(False)
    ax.spines["left"].set_visible(False)
    # ax.spines["left"].set_linewidth(0.5)
    ax.spines["bottom"].set_linewidth(2)
    ax.grid(True, which="both", linestyle="--", linewidth=0.5, alpha=0.7)

plt.savefig("figures/openfold_plddt_vs_lab.png", dpi=300)

plt.tight_layout()
plt.show()

# %% [code]
# Plot OpenFold pLDDT vs lysate/purified bands
plt.figure(figsize=(6, 5))
plt.scatter(data["openfold_plddt"], data["max_activity_zscore"], color=(0.4044212904941176, 0.5346434904470588, 0.9320019126352941), alpha=0.7, s=100,)
plt.xlabel("Mean pLDDT (OpenFold)")
plt.ylabel("Maximum activity (z-score)")

# Remove top and right spines
for ax in plt.gcf().axes:
    ax.spines["top"].set_visible(False)
    ax.spines["right"].set_visible(False)
    ax.spines["left"].set_visible(False)
    # ax.spines["left"].set_linewidth(0.5)
    ax.spines["bottom"].set_linewidth(2)
    ax.grid(True, which="both", linestyle="--", linewidth=0.5, alpha=0.7)
    
plt.tight_layout()
plt.show()

# %% [code]
data.columns

# %% [code]
tmp = data[['id', 'sequence', 'length','dataset',
       'panel', 'openfold_plddt', 'Sequence similarity to EC 1.14.11']]

tmp.rename(columns={
    'Sequence similarity to EC 1.14.11': 'mmseqs2_ec_1.14.11',
}, inplace=True)

# %% [code]
tmp.to_csv("$HOME/enzymes/SI/metagenomic_data.csv", index=False)

# %% [markdown]
## Annotated structure panels
Structure image with text overlay in the bottom-left. One clean image per protein, ready for a figure grid.

# %% [code]
def find_content_bottom(img):
    """Find the y-coordinate of the last row that has non-transparent pixels."""
    arr = np.array(img)
    if arr.shape[2] == 4:
        non_empty_rows = np.where(arr[:, :, 3] > 0)[0]
    else:
        non_empty_rows = np.where(arr.min(axis=(1, 2)) < 250)[0]
    if len(non_empty_rows) == 0:
        return img.height
    return int(non_empty_rows[-1]) + 1


def annotate_structure(d, output_path, annotate_expression=True, annotate_activity=True, annotate_pident=False):
    """Place text labels below the structure content, horizontally centered, left-aligned."""
    img = Image.open(d["img_path"]).convert("RGBA")
    draw = ImageDraw.Draw(img)

    font_size_title = max(48, img.width // 20)
    font_size_body = max(46, img.width // 22)
    try:
        font_title = ImageFont.truetype("/usr/share/fonts/truetype/dejavu/DejaVuSans-Bold.ttf", font_size_title)
        font_body = ImageFont.truetype("/usr/share/fonts/truetype/dejavu/DejaVuSans.ttf", font_size_body)
    except OSError:
        font_title = ImageFont.load_default()
        font_body = ImageFont.load_default()

    plddt = d["plddt"]

    lysate = d.get("lysate_band")
    purified = d.get("purified_band")
    lysate_str = f"{lysate:.0f}" if (lysate is not None and lysate > 0) else "n.d."
    purified_str = f"{purified:.0f}" if (purified is not None and purified > 0) else "n.d."

    black = (0, 0, 0, 255)
    dark_gray = (51, 51, 51, 255)
    lines = [
        (d["seq_id"], font_title, black),
        (f"pLDDT: {plddt:.1f}", font_body, dark_gray),
    ]
    if annotate_expression:
        lines.append((f"Lysate band: {lysate_str}", font_body, dark_gray))
        lines.append((f"Purified band: {purified_str}", font_body, dark_gray))
    if annotate_activity:
        lines.append((f"Max activity (z-score): {d['max_activity_zscore']:.2f}", font_body, dark_gray))
    if annotate_pident:
        max_pident = d.get("max_pident")
        pident_str = f"{max_pident:.1f}" if max_pident is not None and not np.isnan(max_pident) else "n.d."
        lines.append((f"Max UniProtKB identity: {pident_str}", font_body, dark_gray))

    padding = 20
    line_spacing = 26
    gap = 50
    line_heights = []
    line_widths = []
    for text, font, _ in lines:
        bbox = draw.textbbox((0, 0), text, font=font)
        line_heights.append(bbox[3] - bbox[1])
        line_widths.append(bbox[2] - bbox[0])
    text_block_height = sum(line_heights) + line_spacing * (len(lines) - 1) + padding
    max_width = max(line_widths)

    content_bottom = find_content_bottom(img)

    new_height = content_bottom + gap + text_block_height
    new_img = Image.new("RGBA", (img.width, new_height), (0, 0, 0, 0))
    new_img.paste(img.crop((0, 0, img.width, content_bottom)), (0, 0))
    draw = ImageDraw.Draw(new_img)

    block_x0 = (img.width - max_width) // 2
    y = content_bottom + gap
    for (text, font, color), lh in zip(lines, line_heights):
        draw.text((block_x0, y), text, fill=color, font=font)
        y += lh + line_spacing

    new_img.save(output_path, dpi=(300, 300))
    return output_path

def plot_sequences(seq_to_plot, output_dir, annotate_expression=True, annotate_activity=True, annotate_pident=False, df=None):
    """Generate annotated structure images.
    
    Parameters
    ----------
    df : DataFrame, optional
        DataFrame to look up per-sequence metadata (must have an "id" column).
        Defaults to the global `data` DataFrame.
    """
    if df is None:
        df = data

    openfold_records = []
    for seq_id in seq_to_plot:
        seq_dir = CIF_DIR / seq_id
        if not seq_dir.is_dir():
            print(f"Skipping {seq_id}: directory not found in {CIF_DIR}")
            continue

        best_plddt = -1.0
        for conf_file in seq_dir.rglob("*_confidences_aggregated.json"):
            with open(conf_file) as f:
                conf = json.load(f)
            if conf["avg_plddt"] > best_plddt:
                best_plddt = conf["avg_plddt"]
        if best_plddt < 0:
            print(f"Skipping {seq_id}: no confidence file found")
            continue

        img_candidates = list(IMG_DIR.glob(f"{seq_id}*.png"))
        if not img_candidates:
            print(f"Skipping {seq_id}: no image found in {IMG_DIR}")
            continue
        img_path = img_candidates[0]

        row = df.loc[df["id"] == seq_id]
        record = {
            "seq_id": seq_id,
            "plddt": best_plddt,
            "img_path": str(img_path),
            "lysate_band": row["lysate band"].values[0],
            "purified_band": row["purified band"].values[0],
            "max_activity_zscore": row["max_activity_zscore"].values[0],
        }
        if "max_pident" in df.columns:
            record["max_pident"] = row["max_pident"].values[0]

        openfold_records.append(record)

    os.makedirs(output_dir, exist_ok=True)
    for d in openfold_records:
        out = output_dir / f"{d['seq_id']}_annotated.png"
        annotate_structure(d, out, annotate_expression=annotate_expression,
                          annotate_activity=annotate_activity, annotate_pident=annotate_pident)
        print(f"Saved: {out.name}")
        display(IPImage(filename=str(out), width=500))

# %% [code]
import math

def make_grid(image_dir, output_path, ncol=4, order=None):
    """Assemble all annotated PNGs in image_dir into a grid with ncol columns.
    
    Each panel is resized to a uniform width; row heights adapt to the
    tallest image in each row.  White background.
    
    Parameters
    ----------
    order : dict, optional
        Mapping of seq_id -> sort value. Images are sorted by descending value.
        seq_id is extracted from the filename (everything before "_annotated.png").
    """
    paths = sorted(Path(image_dir).glob("*_annotated.png"))
    if not paths:
        print(f"No annotated images found in {image_dir}")
        return

    if order is not None:
        # Filter to only images that have an order value, and sort by that value
        paths = [p for p in paths if p.stem.replace("_annotated", "") in order]
        paths = sorted(paths, key=lambda p: order.get(p.stem.replace("_annotated", ""), 0), reverse=True)

    imgs = [Image.open(p).convert("RGBA") for p in paths]
    n = len(imgs)
    nrow = math.ceil(n / ncol)

    # Uniform cell width; scale each image proportionally
    cell_w = max(im.width for im in imgs)
    scaled = []
    for im in imgs:
        ratio = cell_w / im.width
        new_h = int(im.height * ratio)
        scaled.append(im.resize((cell_w, new_h), Image.LANCZOS))

    # Pad the last row with None so the loop is uniform
    while len(scaled) % ncol != 0:
        scaled.append(None)

    # Compute row heights (tallest image in each row)
    row_heights = []
    for r in range(nrow):
        row_imgs = scaled[r * ncol : (r + 1) * ncol]
        row_heights.append(max(im.height for im in row_imgs if im is not None))

    padding = 40  # space between cells
    grid_w = ncol * cell_w + (ncol - 1) * padding
    grid_h = sum(row_heights) + (nrow - 1) * padding

    grid = Image.new("RGBA", (grid_w, grid_h), (255, 255, 255, 255))

    y = 0
    for r in range(nrow):
        x = 0
        for c in range(ncol):
            im = scaled[r * ncol + c]
            if im is not None:
                grid.paste(im, (x, y), im)
            x += cell_w + padding
        y += row_heights[r] + padding

    grid.save(str(output_path), dpi=(300, 300))
    print(f"Saved grid ({n} panels, {nrow}x{ncol}): {output_path}")
    display(IPImage(filename=str(output_path), width=1200))

# %% [code]
seq_to_plot = list(data[(data["lysate band"] > 0) | (data["purified band"] > 0)]["id"].values)
print(f"Plotting {len(seq_to_plot)} sequences with expression bands...")
plot_sequences(seq_to_plot, OUTPUT_DIR / "expressed", annotate_activity=False, annotate_expression=True)

# %% [code]
plate3_sprot = pd.read_csv("$HOME/enzymes/data/metagenomic_selected/expressed3_uniprot_sprot_mmseqs.csv")
plate3_trembl = pd.read_csv("$HOME/enzymes/data/metagenomic_selected/expressed3_uniprot_trembl_mmseqs.csv")

plate3_mmseqs = pd.concat([plate3_sprot, plate3_trembl], ignore_index=True)
plate3_mmseqs = plate3_mmseqs.groupby("query_id")["max_pident"].max().sort_values(ascending=False).reset_index()
expressed_plate3 = data.merge(plate3_mmseqs, left_on="id", right_on="query_id", how="inner")

# %% [code]
seq_to_plot = list(data[(data["lysate band"] > 0) | (data["purified band"] > 0)]["id"].values)
print(f"Plotting {len(seq_to_plot)} sequences with expression bands...")
plot_sequences(seq_to_plot, OUTPUT_DIR / "expressed", annotate_activity=False, annotate_expression=True, annotate_pident=False, df=expressed_plate3)

# %% [code]
expressed_plate3[expressed_plate3["max_pident"] < 60]

# %% [code]
# Order by max expression (lysate or purified band, whichever is higher)
expression_order = data.set_index("id")[["lysate band", "purified band"]].max(axis=1).to_dict()
make_grid(OUTPUT_DIR / "expressed", OUTPUT_DIR / "expressed_grid.png", ncol=7, order=expression_order)

# %% [code]
tmp = expressed_plate3[expressed_plate3["max_pident"] < 60]

# Order by max expression (lysate or purified band, whichever is higher)
expression_order = tmp.set_index("id")[["lysate band", "purified band"]].max(axis=1).to_dict()
make_grid(OUTPUT_DIR / "expressed_mmseqs", OUTPUT_DIR / "expressed_grid_low_pident.png", ncol=5, order=expression_order)

# %% [code]
tmp = expressed_plate3[expressed_plate3["openfold_plddt"] < 80].head(12)

# Order by max expression (lysate or purified band, whichever is higher)
expression_order = tmp.set_index("id")[["lysate band", "purified band"]].max(axis=1).to_dict()
make_grid(OUTPUT_DIR / "expressed", OUTPUT_DIR / "expressed_grid_low_plddt.png", ncol=3, order=expression_order)

# %% [markdown]
## Closest hits and activity

# %% [code]
results = pd.read_csv("$HOME/enzymes/notebooks/data.csv")

# %% [code]
activity_col = ['proline (EIC)', 'lysine (EIC)', 'indanone (245 nm)', 'resorcinol (285 nm)']
# Normalize activity columns by zscore across all plates
for col in activity_col:
    results[col + " (zscore)"] = (results[col] - results[col].mean()) / results[col].std()
    
# Normalize activity columns by zscore within each plate
for col in activity_col:
    results[col + " (zscore_plate)"] = results.groupby("plate")[col].transform(lambda x: (x - x.mean()) / x.std())

# %% [code]
flashy = results[results["id"].isin(flashy_ids)]

# %% [code]
flashy.columns

# %% [code]
flashy[["id", 'AMPLIFY_c_red', 'protein band', 'proline (EIC) (zscore_plate)', 'lysine (EIC) (zscore_plate)',
       'indanone (245 nm) (zscore_plate)',
       'resorcinol (285 nm) (zscore_plate)', 'proline (EIC) (zscore)', 'lysine (EIC) (zscore)',
       'indanone (245 nm) (zscore)', 'resorcinol (285 nm) (zscore)', 'proline (EIC)', 'lysine (EIC)',
       'indanone (245 nm)', 'resorcinol (285 nm)', 'max activity',
       'mg/mL', 'lysate band', 'purified band']]

# %% [code]
zscore_col = ['proline (EIC) (zscore)', 'lysine (EIC) (zscore)', 'indanone (245 nm) (zscore)', 'resorcinol (285 nm) (zscore)']
results["max_activity_zscore"] = results[zscore_col].max(axis=1)

# %% [code]
results.columns

# %% [code]
blast = pd.read_parquet("$HOME/enzymes/selection_logs/TO_SEND/dataframes/pairwise_blast.parquet")
blast = blast[(~blast["sseqid"].str.contains("MGY")) & (~blast["sseqid"].str.contains("BFD"))]
# Find max match for each sequence in seq3 (get full row where pident is maximal)
max_match = blast[blast["qseqid"].isin(tmp["id"])].groupby("qseqid").apply(lambda x: x.loc[x["pident"].idxmax()]).reset_index(drop=True)

# %% [code]
blast.sort_values(by="pident", ascending=False).head(20)

# %% [code]
tmp = results[results["plate"] == 3][["id", 'proline (EIC) (zscore)', 'lysine (EIC) (zscore)',
       'indanone (245 nm) (zscore)', 'resorcinol (285 nm) (zscore)', "purified band", "lysate band", "max_activity_zscore", 'sequence', "AMPLIFY-C_pseudo_ppl"]].sort_values(by='max_activity_zscore', ascending=False)
tmp = tmp.merge(max_match[["qseqid", "pident", "sseqid", "length"]], left_on="id", right_on="qseqid").drop(columns=["qseqid"])


# %% [code]
tmp["length"].describe()

# %% [code]
tmp.rename(columns={"pident": "blast_to_uniprot_plates", "sseqid": "closest_uniprot"}, inplace=True)

# %% [code]
tmp.to_csv("$HOME/enzymes/data/plate3_activity_zscore.csv", index=False)

# %% [code]
tmp.columns

# %% [code]
```
