## Supplementary material for "pLM representations unlock metagenomic space beyond homology": Code bundle: umaps.ipynb

```
# %% [code]
from pathlib import Path
from functools import reduce

import numpy as np
import pandas as pd

from scipy.stats import pearsonr, spearmanr
from scipy.spatial.distance import pdist, squareform
import os
import matplotlib.pyplot as plt
import seaborn as sns
from umap import UMAP
import joblib

from matplotlib.colors import LogNorm, ListedColormap
from matplotlib.colors import TwoSlopeNorm
from matplotlib.cm import ScalarMappable
from matplotlib.colors import Normalize, ListedColormap


from utils import merge_dataframes
%config InlineBackend.figure_format='retina'

# %% [code]
# Define the directory where your results are stored
DATA_DIR = Path("$SCRATCH/proseqo/enzymes/output/dataframes")
FIG_DIR = Path("$SCRATCH/proseqo/enzymes/output/figures")
FIG_DIR.mkdir(exist_ok=True)

# Define the primary key for merging
KEY_COL = "uniprot_id"

# Add all your generated metrics filenames to this list.
FILES_TO_MERGE = [
    "physicochem_properties.parquet",
    "structural_properties.parquet",
    "signalp_scores.parquet",
    "plm_embeddings_AMPLIFY_350M.parquet",
    "plm_embeddings_AMPLIFY-C.parquet",
    "proteinmpnn_scores_soluble.parquet",
    "esm_if_scores.parquet",
    "plm_perplexity_AMPLIFY-C_sequences_filtered_000.parquet",
]

# Pairwise identity matrix (handled separately)
IDENTITY_MATRIX_FILE = "pairwise_identity.parquet"

# %% [code]
print("Loading metrics files...")
all_dfs = [pd.read_parquet(DATA_DIR / file) for file in FILES_TO_MERGE]

print(f"Merging {len(all_dfs)} dataframes on key '{KEY_COL}'...")
df_main = merge_dataframes(all_dfs, KEY_COL)

# %% [code]
df_main.columns

# %% [code]
df_main["AMPLIFY-C_pseudo_ppl"].describe()

# %% [code]
df_main["AMPLIFY-C_red"] = 1 - df_main["AMPLIFY-C_intra_similarity"]

# Define the UMAP parameters
UMAP_N_NEIGHBORS = 200
UMAP_MIN_DIST = 0.5

# UMAP on AMPLIFY-C embeddings colored by pseudo perplexity
emb_col = "AMPLIFY-C_embedding"
color_col = "AMPLIFY-C_pseudo_ppl"
title = "UMAP of AMPLIFY consensus embeddings"

# Filter out NaNs in embedding or color columns
plot_df = df_main[[emb_col, color_col]].dropna()

# %% [code]
# Stack all embeddings into a single 2D NumPy array
if isinstance(plot_df[emb_col].iloc[0], str):
    features = np.array(plot_df[emb_col].apply(lambda x: np.fromstring(x.strip("[]"), sep=",")).tolist())
else:
    features = np.stack(plot_df[emb_col].values)

# Initialize UMAP and fit-transform
reducer = UMAP(
    n_neighbors=UMAP_N_NEIGHBORS,
    min_dist=UMAP_MIN_DIST,
    n_components=2,
    n_epochs=1000,
    n_jobs=-1,
    verbose=True,
)
# Load model if file exists, otherwise fit and save
if os.path.exists(FIG_DIR / f"{title}_umap_model.joblib"):
    print("Loading existing UMAP model...")
    reducer = joblib.load(FIG_DIR / f"{title}_umap_model.joblib")
    projection = reducer.transform(features)
else:
    projection = reducer.fit_transform(features)
    # Save the umap model for future use
    joblib.dump(reducer, FIG_DIR / f"{title}_umap_model.joblib")

# %% [code]
data = pd.read_csv("$HOME/enzymes/notebooks/data.csv")

# %% [code]
import ast

def interpret_embedding(embedding_str):
    return ast.literal_eval(embedding_str.replace("\n", "").replace("[ ", "[").replace("  ", " ").replace(" ", ","))

# %% [code]
df_metagenomic = data[data["plate"] == 3].copy()
df_uniprot = data[data["plate"] != 3].copy()

metagenomic_projection = reducer.transform(np.stack(df_metagenomic["AMPLIFY-C_embedding"].apply(interpret_embedding).values))
uniprot_projection = reducer.transform(np.stack(df_uniprot["AMPLIFY-C_embedding"].apply(interpret_embedding).values))


# %% [code]
# Plot the projection
fig, axes = plt.subplots(figsize=(20, 6), ncols=3)

vmin, vmax = plot_df[color_col].quantile(0.01), plot_df[color_col].quantile(0.95)

ax = axes[0]
points = ax.scatter(
    projection[:, 0],
    projection[:, 1],
    c=plot_df[color_col].values,
    cmap="viridis",
    s=5,
    alpha=1,
    vmin=vmin,
    vmax=vmax,
)

# Add colorbar and labels
cbar = fig.colorbar(points, ax=ax, shrink=0.8, label="Pseudo-perplexity")
ax.set_title(title)
ax.spines[["top", "right", "left", "bottom"]].set_visible(False)
ax.set_xticks([])
ax.set_yticks([])

for i, col in enumerate(["lysate band", "purified band"]):
    ax = axes[i + 1]
    ax.scatter(
        projection[:, 0],
        projection[:, 1],
        c="lightgray",
        s=1,
        alpha=0.1,
        vmin=0,
    )

    # Set midpoint for color normalization with two slopes
    norm = TwoSlopeNorm(vmin=0, vcenter=10, vmax=3500)

    points = ax.scatter(
        uniprot_projection[:, 0],
        uniprot_projection[:, 1],
        c=df_uniprot[col].values,
        cmap="coolwarm_r",
        s=10,
        alpha=1,
        norm=norm
    )
    
    cmap = plt.get_cmap("coolwarm_r")

    N = 512
    vmin, vmax = 1, min(df_uniprot[col].max(), 3500)
    data_grid = np.geomspace(vmin, vmax, N)
    true_colors = cmap(norm(data_grid))        # norm is still the TwoSlopeNorm
    log_cmap = ListedColormap(true_colors)
    

    sm = ScalarMappable(norm=LogNorm(vmin=vmin, vmax=vmax), cmap=log_cmap)
    cbar = fig.colorbar(sm, ax=ax, shrink=0.8, label=col)
    cbar.set_ticks([1, 10, 100, 1000, vmax])
    cbar.set_ticklabels(["1", "10", "100", "1000", str(vmax)])
    
    # Add colorbar and labels
    # cbar = fig.colorbar(points, ax=ax, shrink=0.8, label=col)
    # cbar_ticks = [0, 10, 500, 1000, 2000, 3500]
    # cbar.set_ticks(cbar_ticks)
    # cbar.set_ticklabels([str(tick) for tick in cbar_ticks])
    
    ax.set_title(title)
    ax.spines[["top", "right", "left", "bottom"]].set_visible(False)
    ax.set_xticks([])
    ax.set_yticks([])

# Save and show the figure
plt.tight_layout()
# plt.savefig(FIG_DIR / f"{title}.png", dpi=300)
plt.show()

# %% [code]
known_hydroxylases = pd.read_parquet("$SCRATCH/proseqo/enzymes/output/dataframes/plm_embeddings_known_hydroxylases_AMPLIFY-C.parquet")
knwon_projection = reducer.transform(np.stack(known_hydroxylases["AMPLIFY-C_embedding"].values))

# %% [code]
from mpl_toolkits.axes_grid1 import make_axes_locatable

fontsize = 16

# Plot the projection
fig, axes = plt.subplots(figsize=(20, 6), ncols=3, gridspec_kw={"width_ratios": [1, 0.85, 1]})

color_col = "AMPLIFY-C_pseudo_ppl"
vmin, vmax = plot_df[color_col].quantile(0.05), plot_df[color_col].quantile(0.95)

ax = axes[0]
points = ax.scatter(
    projection[:, 0],
    projection[:, 1],
    c=plot_df[color_col].values,
    cmap="coolwarm",
    s=4,
    alpha=1,
    vmin=vmin,
    vmax=vmax,
)

# Add colorbar and labels
cbar = fig.colorbar(points, ax=ax, shrink=0.8)
cbar.set_label("Perplexity", size=fontsize)
ax.spines[["top", "right", "left", "bottom"]].set_visible(False)
ax.set_xticks([])
ax.set_yticks([])
col = "lysate band"
values = df_uniprot[col].values
masks = [
    ("lysate band < 10", values < 10),
    ("lysate band ≥ 10", values >= 10),
]

for i, (label, mask) in enumerate(masks):
    ax = axes[i + 1]
    ax.scatter(
        projection[:, 0],
        projection[:, 1],
        c="lightgray",
        s=1,
        alpha=0.4,
        vmin=0,
    )

    norm = TwoSlopeNorm(vmin=0, vcenter=10, vmax=2000)

    points = ax.scatter(
        uniprot_projection[mask, 0],
        uniprot_projection[mask, 1],
        c=values[mask],
        cmap="coolwarm_r",
        s=20,
        alpha=1,
        norm=norm,
        # zorder=5,
    )

    if i == 1:
        cmap = plt.get_cmap("coolwarm_r")

        N = 512
        vmin, vmax = 1, min(df_uniprot[col].max(), 2000)
        data_grid = np.geomspace(vmin, vmax, N)
        true_colors = cmap(norm(data_grid))
        log_cmap = ListedColormap(true_colors)

        sm = ScalarMappable(norm=LogNorm(vmin=vmin, vmax=vmax), cmap=log_cmap)
        cbar = fig.colorbar(sm, ax=ax, shrink=0.8, label="lysate band")
        cbar.set_label("Lysate band", size=fontsize)
        cbar.set_ticks([1, 10, 100, 500, vmax])
        cbar.set_ticklabels(["1", "10", "100", "500", str(vmax)])
        
        points = ax.scatter(
            knwon_projection[:, 0],
            knwon_projection[:, 1],
            c="purple",
            marker="x",
            s=20,
            alpha=1,
        )
        
        handle = plt.Line2D([0], [0], marker='x', color='purple', label='Known hydroxylases', linestyle='None', markerfacecolor='purple', markersize=5)
        ax.legend(handles=[handle], loc='lower center', fontsize=fontsize, frameon=False, bbox_to_anchor=(0.7, -0.0))

    ax.spines[["top", "right", "left", "bottom"]].set_visible(False)
    ax.set_xticks([])
    ax.set_yticks([])

# Save and show the figure
plt.tight_layout()
plt.savefig("figures/umap_ppl_uniprot_wetlabs.png", dpi=300)
plt.show()

# %% [code]
from mpl_toolkits.axes_grid1 import make_axes_locatable

# Plot the projection
fig, axes = plt.subplots(figsize=(14, 6), ncols=2, gridspec_kw={"width_ratios": [1, 0.85]})

color_col = "AMPLIFY-C_pseudo_ppl"
vmin, vmax = plot_df[color_col].quantile(0.05), plot_df[color_col].quantile(0.95)

ax = axes[0]
points = ax.scatter(
    projection[:, 0],
    projection[:, 1],
    c=plot_df[color_col].values,
    cmap="coolwarm",
    s=4,
    alpha=1,
    vmin=vmin,
    vmax=vmax,
)

# Add colorbar and labels
cbar = fig.colorbar(points, ax=ax, shrink=0.8)
cbar.set_label("Pseudo-perplexity", size=14)
ax.spines[["top", "right", "left", "bottom"]].set_visible(False)
ax.set_xticks([])
ax.set_yticks([])

ax = axes[1]
ax.scatter(
    projection[:, 0],
    projection[:, 1],
    c="lightgray",
    s=1,
    alpha=0.4,
    vmin=0,
)

norm = TwoSlopeNorm(vmin=0, vcenter=10, vmax=3500)

points = ax.scatter(
    knwon_projection[:, 0],
    knwon_projection[:, 1],
    c="purple",
    s=20,
    alpha=1,
    norm=norm,
)

ax.spines[["top", "right", "left", "bottom"]].set_visible(False)
ax.set_xticks([])
ax.set_yticks([])

# Save and show the figure
plt.tight_layout()
# plt.savefig("figures/umap_ppl_known_wetlabs.png", dpi=300)
plt.show()

# %% [code]
from mpl_toolkits.axes_grid1 import make_axes_locatable

# Plot the projection
fig, axes = plt.subplots(figsize=(20, 6), ncols=3, gridspec_kw={"width_ratios": [1, 0.85, 1]})

color_col = "AMPLIFY-C_pseudo_ppl"
vmin, vmax = plot_df[color_col].quantile(0.05), plot_df[color_col].quantile(0.95)

ax = axes[0]
points = ax.scatter(
    projection[:, 0],
    projection[:, 1],
    c=plot_df[color_col].values,
    cmap="coolwarm",
    s=4,
    alpha=1,
    vmin=vmin,
    vmax=vmax,
)

# Add colorbar and labels
cbar = fig.colorbar(points, ax=ax, shrink=0.8)
cbar.set_label("Pseudo-perplexity", size=14)
ax.spines[["top", "right", "left", "bottom"]].set_visible(False)
ax.set_xticks([])
ax.set_yticks([])
col = "purified band"
values = df_metagenomic[col].values
masks = [
    ("purified band < 10", values < 10),
    ("purified band ≥ 10", values >= 10),
]

for i, (label, mask) in enumerate(masks):
    ax = axes[i + 1]
    ax.scatter(
        projection[:, 0],
        projection[:, 1],
        c="lightgray",
        s=1,
        alpha=0.4,
        vmin=0,
    )

    norm = TwoSlopeNorm(vmin=0, vcenter=10, vmax=3500)

    points = ax.scatter(
        metagenomic_projection[mask, 0],
        metagenomic_projection[mask, 1],
        c=values[mask],
        cmap="coolwarm_r",
        s=20,
        alpha=1,
        norm=norm,
    )

    if i == 1:
        cmap = plt.get_cmap("coolwarm_r")

        N = 512
        vmin, vmax = 1, min(df_metagenomic[col].max(), 3500)
        data_grid = np.geomspace(vmin, vmax, N)
        true_colors = cmap(norm(data_grid))
        log_cmap = ListedColormap(true_colors)

        sm = ScalarMappable(norm=LogNorm(vmin=vmin, vmax=vmax), cmap=log_cmap)
        cbar = fig.colorbar(sm, ax=ax, shrink=0.8, label="Purified band")
        cbar.set_label("Purified band", size=14)
        cbar.set_ticks([1, 10, 100, 1000, vmax])
        cbar.set_ticklabels(["1", "10", "100", "1000", str(vmax)])

    ax.spines[["top", "right", "left", "bottom"]].set_visible(False)
    ax.set_xticks([])
    ax.set_yticks([])

# Save and show the figure
plt.tight_layout()
# plt.savefig("figures/umap_ppl_uniprot_wetlabs.png", dpi=300)
plt.show()

# %% [code]
# Plot the projection
fig, ax = plt.subplots(figsize=(8, 6))
ax.scatter(
    projection[:, 0],
    projection[:, 1],
    c="lightgray",
    s=1,
    alpha=0.4,
)

# norm = TwoSlopeNorm(vmin=0, vcenter=10, vmax=3500)
    
points = ax.scatter(
    metagenomic_projection[:, 0],
    metagenomic_projection[:, 1],
    c=df_metagenomic["AMPLIFY-C_pseudo_ppl"].values,
    cmap="coolwarm",
    s=10,
    alpha=1,
    vmin=1,
    vmax=13,
)

cbar = fig.colorbar(points, ax=ax, shrink=0.8)
cbar.set_label("Pseudo-perplexity", size=14)

ax.spines[["top", "right", "left", "bottom"]].set_visible(False)
ax.set_xticks([])
ax.set_yticks([])

# Save and show the figure
plt.tight_layout()
plt.savefig("figures/umap_ppl_metagenomic.png", dpi=300)
plt.show()

# %% [code]
df_metagenomic["purified band"].values

# %% [code]
# Plot the projection
fig, ax = plt.subplots(figsize=(8, 6))
ax.scatter(
    projection[:, 0],
    projection[:, 1],
    c="lightgray",
    s=1,
    alpha=0.4,
)
points = ax.scatter(
    metagenomic_projection[:, 0],
    metagenomic_projection[:, 1],
    c=df_metagenomic["purified band"].values,
    cmap="coolwarm",
    s=10,
    alpha=1,
    vmin=0,
    vmax=3000,
)

# Add colorbar and labels
cbar = fig.colorbar(points, ax=ax, shrink=0.8, label="Pseudo-perplexity")
cbar.set_label(color_col)
ax.set_title(title)
ax.spines[["top", "right", "left", "bottom"]].set_visible(False)
ax.set_xticks([])
ax.set_yticks([])

# Save and show the figure
plt.tight_layout()
# plt.savefig(FIG_DIR / f"{title}.png", dpi=300)
plt.show()

# %% [code]
# Plot the projection
fig, ax = plt.subplots(figsize=(8, 6))
ax.scatter(
    projection[:, 0],
    projection[:, 1],
    c="lightgray",
    s=1,
    alpha=0.4,
    vmin=0,
)
points = ax.scatter(
    uniprot_projection[:, 0],
    uniprot_projection[:, 1],
    c=df_uniprot["purified band"].values,
    cmap="viridis_r",
    s=10,
    alpha=1,
    vmin=0,
)

# Add colorbar and labels
cbar = fig.colorbar(points, ax=ax, shrink=0.8, label="RED Score")
cbar.set_label(color_col)
ax.set_title(title)
ax.spines[["top", "right", "left", "bottom"]].set_visible(False)
ax.set_xticks([])
ax.set_yticks([])

# Save and show the figure
plt.tight_layout()
# plt.savefig(FIG_DIR / f"{title}.png", dpi=300)
plt.show()

# %% [code]
# Plot the projection
fig, ax = plt.subplots(figsize=(8, 6))
ax.scatter(
    projection[:, 0],
    projection[:, 1],
    c="lightgray",
    s=1,
    alpha=0.4,
    vmin=0,
)
points = ax.scatter(
    metagenomic_projection[:, 0],
    metagenomic_projection[:, 1],
    c=df_metagenomic["intra_score"].values,
    cmap="viridis_r",
    s=10,
    alpha=1,
    vmin=0,
)

# Add colorbar and labels
cbar = fig.colorbar(points, ax=ax, shrink=0.8, label="RED Score")
cbar.set_label(color_col)
ax.set_title(title)
ax.spines[["top", "right", "left", "bottom"]].set_visible(False)
ax.set_xticks([])
ax.set_yticks([])

# Save and show the figure
plt.tight_layout()
# plt.savefig(FIG_DIR / f"{title}.png", dpi=300)
plt.show()

# %% [code]
data = pd.read_csv("$HOME/enzymes/notebooks/data.csv")

# %% [code]
candidates = reducer.transform(np.stack(data[emb_col].apply(lambda x: np.fromstring(x.strip("[]"), sep=",")).tolist()))
data["umap"] = list(candidates)

# %% [code]
data.columns

# %% [code]
# Plot the projection
fig, ax = plt.subplots(figsize=(8, 6))
points = ax.scatter(
    projection[:, 0],
    projection[:, 1],
    c="grey",
    s=1,
    alpha=0.1,
)

points = ax.scatter(
    data["umap"].apply(lambda x: x[0]),
    data["umap"].apply(lambda x: x[1]),
    c=data["AMPLIFY_c_red"].values,
    cmap="viridis_r",
    s=10,
    alpha=1,
    vmin=0,
)

# Add colorbar and labels
cbar = fig.colorbar(points, ax=ax, shrink=0.8, label="RED Score")
cbar.set_label(color_col)
ax.set_title(title)
ax.spines[["top", "right", "left", "bottom"]].set_visible(False)
ax.set_xticks([])
ax.set_yticks([])

# Save and show the figure
plt.tight_layout()
# plt.savefig(FIG_DIR / f"{title}.png", dpi=300)
plt.show()

# %% [code]
from matplotlib.colors import TwoSlopeNorm

# %% [code]
# Plot the projection
fig, ax = plt.subplots(figsize=(8, 6))
points = ax.scatter(
    projection[:, 0],
    projection[:, 1],
    c="grey",
    s=1,
    alpha=0.1,
)

midpoint = 500
norm = TwoSlopeNorm(vmin=data["lysate band"].min(), vcenter=midpoint, vmax=data["lysate band"].max())

points = ax.scatter(
    data["umap"].apply(lambda x: x[0]),
    data["umap"].apply(lambda x: x[1]),
    c=data["lysate band"].values,
    cmap="coolwarm_r",
    s=10,
    alpha=1,
    norm=norm,
)

# Add colorbar and labels
cbar = fig.colorbar(points, ax=ax, shrink=0.8, label="RED Score")
cbar.set_label(color_col)
ax.set_title(title)
ax.spines[["top", "right", "left", "bottom"]].set_visible(False)
ax.set_xticks([])
ax.set_yticks([])

# Save and show the figure
plt.tight_layout()
# plt.savefig(FIG_DIR / f"{title}.png", dpi=300)
plt.show()

# %% [code]
# Plot the projection
fig, ax = plt.subplots(figsize=(8, 6))
points = ax.scatter(
    projection[:, 0],
    projection[:, 1],
    c="grey",
    s=1,
    alpha=0.1,
)

midpoint = 500
norm = TwoSlopeNorm(vmin=data["purified band"].min(), vcenter=midpoint, vmax=data["purified band"].max())

points = ax.scatter(
    data["umap"].apply(lambda x: x[0]),
    data["umap"].apply(lambda x: x[1]),
    c=data["purified band"].values,
    cmap="coolwarm_r",
    s=10,
    alpha=1,
    norm=norm,
)

# Add colorbar and labels
cbar = fig.colorbar(points, ax=ax, shrink=0.8, label="RED Score")
cbar.set_label(color_col)
ax.set_title(title)
ax.spines[["top", "right", "left", "bottom"]].set_visible(False)
ax.set_xticks([])
ax.set_yticks([])

# Save and show the figure
plt.tight_layout()
# plt.savefig(FIG_DIR / f"{title}.png", dpi=300)
plt.show()

# %% [code]
```
