## Supplementary material for "pLM representations unlock metagenomic space beyond homology": Code bundle: utils.py

```
from functools import reduce
import pandas as pd

import seaborn as sns


def plot_distribution(ax, data, prop):
    # Plot distribution
    sns.histplot(
        data,
        x=prop,
        bins=30,
        color="#0085ff",
        alpha=0.4,
        lw=0.1,
        ax=ax,
        stat="count",
    )
    ax.set_title(f"Distribution of {prop}")
    ax.set_xlabel(prop)
    ax.set_ylabel("Count")
    ax.tick_params(left=False)
    ax.spines[["top", "right", "left"]].set_visible(False)
    ax.spines["bottom"].set_linewidth(2)
    ax.yaxis.grid(True, which="major", linestyle=":")
    ax.set_axisbelow(True)


def merge_dataframes(dataframes, key_col):
    """
    Merges a list of DataFrames on a common key column, handling
    duplicate metadata columns.
    """
    # Load the first dataframe completely
    base_df = dataframes[0]

    # Iteratively merge the rest
    if len(dataframes) > 1:
        other_dfs = []
        base_cols = set(base_df.columns)

        for df in dataframes[1:]:
            # Find unique columns in this df
            unique_cols = [col for col in df.columns if col not in base_cols]

            # Add to base_cols so they aren't re-added from another file
            base_cols.update(unique_cols)

            # Only load key + unique columns
            cols_to_load = [key_col] + unique_cols

            # Ensure we don't try to load columns that don't exist
            valid_cols_to_load = [col for col in cols_to_load if col in df.columns]

            other_dfs.append(df[valid_cols_to_load])

        # Now merge all the small dataframes into the base
        final_df = reduce(lambda left, right: pd.merge(left, right, on=key_col, how="outer"), [base_df] + other_dfs)
        return final_df

    return base_df
```
