## Supplementary material for "pLM representations unlock metagenomic space beyond homology": Code bundle: visualize_structures.ipynb

```
# %% [markdown]
# OpenFold Structure Visualization

Renders CIF structures as high-resolution PNGs colored by pLDDT confidence,
for all sequences with predictions found in `CIF_DIR`.

# %% [code]
import json
import os
from pathlib import Path
import pandas as pd

import pymol
from pymol import cmd
pymol.finish_launching(['pymol', '-qc'])

from IPython.display import display, Image as IPImage
from PIL import Image

# %% [code]
CSV_PATH = "$HOME/enzymes/data/metagenomic_selected/flashy_plate3.csv"
CIF_DIR = Path("$SCRATCH/proseqo/enzymes/metagenomic_output/openfold/dataframes")
OUTPUT_DIR = Path("$SCRATCH/proseqo/enzymes/metagenomic_output/openfold/structure_images")
OUTPUT_DIR.mkdir(parents=True, exist_ok=True)


# Image settings
IMG_WIDTH = 1600
IMG_HEIGHT = 1200

# %% [code]
def get_best_sample(seq_dir: Path) -> tuple[Path, dict] | None:
    """Select the sample with the highest pLDDT from a sequence directory,
    skipping CIF files that PyMOL cannot parse."""
    candidates = []

    for conf_file in seq_dir.rglob("*_confidences_aggregated.json"):
        with open(conf_file) as f:
            conf = json.load(f)
        cif_path = conf_file.parent / conf_file.name.replace(
            "_confidences_aggregated.json", "_model.cif"
        )
        if cif_path.exists():
            candidates.append((conf["avg_plddt"], cif_path, conf))

    # Try candidates from highest to lowest pLDDT
    for _, cif_path, conf in sorted(candidates, reverse=True):
        try:
            cmd.reinitialize()
            cmd.load(str(cif_path), "_test")
            cmd.delete("_test")
            return cif_path, conf
        except Exception:
            print(f"    [WARN] corrupt CIF, skipping: {cif_path.name}")
            continue

    return None

# %% [code]
def render_cif_to_png(cif_path: Path, output_path: Path,
                      width=IMG_WIDTH, height=IMG_HEIGHT):
    """Render a CIF structure to PNG as cartoon colored by pLDDT (B-factor)."""
    cmd.reinitialize()
    cmd.load(str(cif_path), "struct")
    cmd.hide("everything")
    cmd.show("cartoon")
    # pLDDT coloring via B-factor
    cmd.spectrum("b", "orange_yellow_cyan_blue", minimum=50, maximum=90)
    cmd.bg_color("white")
    cmd.set("ray_opaque_background", 0)
    cmd.orient()
    cmd.ray(width, height)
    cmd.png(str(output_path), dpi=300)

    img = Image.open(str(output_path))
    return img

# %% [code]
# Discover all sequence directories in CIF_DIR
seq_ids = [d.name for d in sorted(CIF_DIR.iterdir()) if d.is_dir()]
print(f"Sequence directories found in CIF_DIR: {len(seq_ids)}")

# Find which ones have CIF files
sequences = []
for seq_id in seq_ids:
    seq_dir = CIF_DIR / seq_id
    result = get_best_sample(seq_dir)
    if result:
        sequences.append((seq_id, *result))
        print(f"  [OK]   {seq_id} — pLDDT={result[1]['avg_plddt']:.1f}")
    else:
        print(f"  [SKIP] {seq_id} — no CIF files found")


# %% [code]
# Render and save each structure
for seq_id, cif_path, conf in sequences:
    out_path = OUTPUT_DIR / f"{seq_id}.png"
    if out_path.exists():
        # print(f"  [SKIP] {seq_id} — image already exists")
        continue
    
    print(f"Rendering {seq_id} (pLDDT={conf['avg_plddt']:.1f}, pTM={conf['ptm']:.3f}) ...")

    img = render_cif_to_png(cif_path, out_path)
    print(f"  Saved: {out_path} ({img.size[0]}x{img.size[1]})")
    display(IPImage(filename=str(out_path), width=400))
    print()

# %% [markdown]
# 

# %% [code]
```
