## Supplementary material for "pLM representations unlock metagenomic space beyond homology": Code bundle: marginal_relevance.py

```
import numpy as np
from tqdm import tqdm
import pandas as pd
import argparse
from pathlib import Path


def select_proteins_mmr(
    quality_scores,
    cosine_matrix,
    blast_matrix,
    n_select=96,
    lambda_param=0.5,
    blast_threshold=80,
    preselected_indices=[],
):
    # Initialization: Pick the absolute best quality not in to start
    if preselected_indices == []:
        first_choice = np.argmax(quality_scores)
        selected_indices = [first_choice]
        preselected_indices.append(first_choice)
    else:
        selected_indices = []

    n_select -= len(selected_indices)

    # Iterative Selection
    for _ in tqdm(range(n_select)):
        remaining_indices = [i for i in range(len(quality_scores)) if i not in preselected_indices]
        candidate_to_add = -1

        # Shape: (Num_Remaining, Num_Selected)
        sim_to_selected = cosine_matrix[remaining_indices][:, preselected_indices]

        # Calculate max similarity to the selected set for each remaining item
        max_sim_vals = np.max(sim_to_selected, axis=1)

        # Calculate MMR scores for all remaining items
        # mmr = lambda * Quality - (1-lambda) * Max_Sim
        current_qualities = quality_scores[remaining_indices]
        mmr_scores = (lambda_param * current_qualities) - ((1 - lambda_param) * max_sim_vals)

        # Sort candidates by MMR score descending
        sorted_candidate_args = np.argsort(mmr_scores)[::-1]

        # Apply BLAST Hard Constraint
        # We iterate down the ranked list and pick the first one that passes the BLAST check
        found_next = False
        while not found_next:
            for local_idx in sorted_candidate_args:
                global_idx = remaining_indices[local_idx]

                # Check BLAST against ALL currently selected
                # blast_matrix[global_idx, preselected_indices] returns vector of blast scores
                blast_scores = blast_matrix[global_idx, preselected_indices]

                if np.all(blast_scores <= blast_threshold):
                    candidate_to_add = global_idx
                    found_next = True
                    break

            if found_next:
                selected_indices.append(candidate_to_add)
                preselected_indices.append(candidate_to_add)
            else:
                blast_threshold += 5
                print(f"Warning: Could not find enough candidates satisfying BLAST constraint, increasing threshold to {blast_threshold}.")

    return selected_indices


if __name__ == "__main__":

    parser = argparse.ArgumentParser(description="Calculates basic physicochemical properties for protein sequences.")
    parser.add_argument(
        "--input_dir",
        type=str,
        default="input",
        help="Path to the input data directory.",
    )
    parser.add_argument(
        "--output_dir",
        type=str,
        default="selection_logs",
        help="Main directory to store all outputs.",
    )
    parser.add_argument(
        "--version",
        type=int,
        default="selection_logs",
        help="Version of input (filtered) data.",
    )
    parser.add_argument(
        "--id_col",
        type=str,
        default="uniprot_id",
        help="Name of id column.",
    )
    parser.add_argument(
        "--N_select",
        type=int,
        default=96,
        help="Number of candidates to select.",
    )
    parser.add_argument(
        "--lambda_param",
        type=float,
        default=0.5,
        help="Lambda parameter for MMR scoring function.",
    )
    parser.add_argument(
        "--blast_threshold",
        type=int,
        default=50,
        help="Maximum sequence identity between selected candidates.",
    )
    parser.add_argument(
        "--preselected_ids",
        type=str,
        default=None,
        help="Path to list of preselected indices.",
    )
    args = parser.parse_args()

    # Add all your generated metrics filenames to this list.
    data_dir = Path(args.input_dir)

    print(f"Loading metrics files (merged_data_{args.version:03}.parquet).")
    df_main = pd.read_parquet(data_dir / f"merged_data_{args.version:03}.parquet")
    df_main.sort_values(args.id_col, inplace=True)
    quality = df_main["AMPLIFY-C_pseudo_ppl"]
    ids = list(df_main[args.id_col])

    print(f"Loading pairwise BLAST matrix (pairwise_blast_{args.version:03}.parquet).")
    blast_df = pd.read_parquet(data_dir / f"pairwise_blast_{args.version:03}.parquet")
    blast_df.sort_index(axis=0, inplace=True)
    blast_df.sort_index(axis=1, inplace=True)

    print(f"Loading pairwise similarity matrix (pairwise_similarity_{args.version:03}.parquet).")
    similarity_df = pd.read_parquet(data_dir / f"pairwise_similarity_{args.version:03}.parquet")
    similarity_df.sort_index(axis=0, inplace=True)
    similarity_df.sort_index(axis=1, inplace=True)

    # Normalize quality scores
    l_min = np.min(quality)
    l_max = np.max(quality)
    if l_max - l_min == 0:
        norm_quality = np.ones_like(quality)
    else:
        norm_quality = (l_max - quality) / (l_max - l_min)

    print("Converting pairwise matrices to numpy")
    cosine_matrix = similarity_df.to_numpy(dtype=np.float16)
    blast_matrix = blast_df.to_numpy(dtype=np.float16)
    norm_quality = norm_quality.to_numpy()

    del similarity_df, blast_df

    # for lambda_param in np.arange(0.90, 1, 0.05):
    print(f"Running Maximum Marginal Relevance algorithm with lambda={args.lambda_param}, blast_threshold={args.blast_threshold}.")
    if args.preselected_ids is not None:
        with open(args.preselected_ids, "r") as f:
            selected_ids = set(line.strip() for line in f if line.strip())
        preselected_indices = [idx for idx, id in enumerate(ids) if id in selected_ids]
        print(f"Starting from {len(preselected_indices)} preselected sequences.")
    else:
        preselected_indices = []

    selected_indices = select_proteins_mmr(
        norm_quality,
        cosine_matrix=cosine_matrix,
        n_select=args.N_select,
        blast_matrix=blast_matrix,
        lambda_param=args.lambda_param,
        blast_threshold=args.blast_threshold,
        preselected_indices=preselected_indices,
    )

    selected_uniprot_ids = [ids[idx] for idx in selected_indices]
    output_file = f"{args.output_dir}/selected_uniprot_ids_mmr_lambda_{args.lambda_param:.2f}_{args.blast_threshold}.txt"
    with open(output_file, "w") as f:
        for uid in selected_uniprot_ids:
            f.write(f"{uid}\n")

    print(f"Selected ids saved in file {output_file}.")
```
