## Supplementary material for "pLM representations unlock metagenomic space beyond homology": Code bundle: pyproject.toml

```
[build-system]
requires = ["setuptools >= 61.0"]
build-backend = "setuptools.build_meta"

[project]
name = "enzyme-discovery"
version = "0.0.0"
description = "pLM-guided candidate selection and metagenomic retrieval for the Fe/aKG-dependent dioxygenase (EC 1.14.11) wet-lab campaign"
requires-python = ">= 3.10"
dependencies = [
    "numpy<2",
    "pandas",
    "polars",
    "pyarrow",
    "scipy",
    "tqdm",
    "ipykernel",
    "biopython",
    "biotite==0.41.2",
    "matplotlib",
    "seaborn",
    "umap-learn",
    "scikit-learn",
    "lightgbm",
    "optuna",
    "pybiolib",
    "torch",
    "transformers",
    "accelerate",
    "xformers==0.0.24.*",
    "torch-scatter",
    "torch-sparse",
    "torch-cluster",
    "torch-spline-conv",
    "torch-geometric",
    "pymol-open-source-whl",
]

# Not installable from PyPI, and therefore not listed above (see the README):
#   - ESM-IF: `pip install git+https://github.com/facebookresearch/esm.git`
#   - SignalP 6.0: manual download and install from DTU Health Tech.
#   - mmseqs2, ProteinMPNN and hh-suite are external binaries/repositories.
#   - OpenFold is used for structure prediction of retrieved candidates.
#   - AMPLIFY model weights are loaded from the Hugging Face Hub at run time;
#     see the "Models" section of the README.

authors = [
    { name = "Lola Le Breton", email = "" },
    { name = "Quentin Fournier", email = "" },
]
maintainers = [
    { name = "Lola Le Breton", email = "" },
]
license = { text = "MIT License" }
keywords = []

[project.optional-dependencies]
dev = ["black", "pytest"]

# enzymes/ is a collection of standalone scripts and notebooks rather than an
# importable library, so there is nothing to discover. Without this, setuptools
# flat-layout autodiscovery fails on the jobs/, data/, notebooks/ and
# optimization/ directories. Installing the project therefore only provides the
# dependencies; run the scripts directly from this directory.
[tool.setuptools]
py-modules = []
```
