## Supplementary material for "pLM representations unlock metagenomic space beyond homology": Code bundle: 01_physicochem.py

```
"""
Calculates basic physicochemical properties for each sequence.
"""

import argparse
import re
from concurrent.futures import ProcessPoolExecutor
from pathlib import Path

import numpy as np
import pandas as pd
from Bio.SeqUtils.ProtParam import ProteinAnalysis
from tqdm import tqdm
from utils import load_preprocess_data
from scipy.stats import entropy


AMINO_ACIDS = ["A", "C", "D", "E", "F", "G", "H", "I", "K", "L", "M", "N", "P", "Q", "R", "S", "T", "V", "W", "Y"]
EXPECTED_FREQS = [7.4, 3.3, 5.9, 5.8, 4.0, 7.4, 2.9, 3.8, 7.2, 7.6, 1.8, 4.4, 5.0, 3.7, 4.2, 8.1, 6.2, 6.8, 1.3, 3.3]


def calculate_sa_repeat_ratio(seq):
    """Calculates the fraction of the sequence composed of single amino acid repeats (e.g., AAAA, GGG)."""
    n_repeat_res = sum(len(m.group(0)) for m in re.finditer(r"(.)\1+", seq))
    if len(seq) == 0:
        return 0.0
    return n_repeat_res / len(seq)


def compute_protein_properties(sequence):
    """Remove non-standard AAs (U, O, Z, B, J, X, *) for most Biopython analyses."""
    cleaned_seq = re.sub("[UOZBJX*|]", "", str(sequence).upper())

    analysis = ProteinAnalysis(cleaned_seq)
    ss_helix, ss_turn, ss_sheet = analysis.secondary_structure_fraction()
    mol_ext_reduced, mol_ext_oxidized = analysis.molar_extinction_coefficient()

    # Count Observed Frequencies
    observed_count = [sequence.count(aa) for aa in AMINO_ACIDS]

    # Entropy Calculation (will be normilized by default)
    shannon_entropy = entropy(observed_count)
    shannon_entropy_normalized = entropy(observed_count, qk=EXPECTED_FREQS)

    properties = {
        "non_standard_aa_count": sum(1 for char in str(sequence).upper() if char in "UOZBJX*|"),
        "length": len(sequence),
        "molecular_weight": analysis.molecular_weight(),
        "aromaticity": analysis.aromaticity(),
        "instability_index": analysis.instability_index(),
        "isoelectric_point": analysis.isoelectric_point(),
        "flexibility_mean": np.mean(analysis.flexibility()) if analysis.flexibility() else np.nan,
        "charge_at_pH7": analysis.charge_at_pH(7.0),
        "charge_at_pH7.5": analysis.charge_at_pH(7.5),
        "charge_at_pH8": analysis.charge_at_pH(8),
        "gravy": analysis.gravy(),
        "ss_helix": ss_helix,
        "ss_turn": ss_turn,
        "ss_sheet": ss_sheet,
        "mol_ext_reduced": mol_ext_reduced,
        "mol_ext_oxidized": mol_ext_oxidized,
        "sa_repeat_ratio": calculate_sa_repeat_ratio(cleaned_seq),
        "shannon_entropy": shannon_entropy,
        "shannon_entropy_normalized": shannon_entropy_normalized,
    }
    return pd.Series(properties)


if __name__ == "__main__":

    parser = argparse.ArgumentParser(description="Calculates basic physicochemical properties for protein sequences.")
    parser.add_argument(
        "--input_csv",
        type=str,
        default="data/uniprotkb_ec_1_14_11_2025_10_20.csv",
        help="Path to the input CSV file containing sequences.",
    )
    parser.add_argument(
        "--output_dir",
        type=str,
        default="output",
        help="Main directory to store all outputs.",
    )
    parser.add_argument(
        "--n_samples",
        type=int,
        default=0,
        help="Number of sequences to sample. Set to 0 to disable.",
    )
    parser.add_argument(
        "--cores",
        type=int,
        default=8,
        help="Number of CPU cores to use for parallel processing (if applicable).",
    )
    args = parser.parse_args()

    output_dir = Path(args.output_dir)
    df_dir = output_dir / "dataframes"
    output_dir.mkdir(exist_ok=True)
    df_dir.mkdir(exist_ok=True)

    df_main = load_preprocess_data(args.input_csv, args.n_samples)

    print(f"Computing basic physicochemical properties using {args.cores} cores...")
    sequences_to_process = df_main["sequence"].tolist()

    with ProcessPoolExecutor(max_workers=args.cores) as executor:
        results = list(
            tqdm(
                executor.map(compute_protein_properties, sequences_to_process),
                total=len(sequences_to_process),
                desc="Computing properties",
            )
        )

    properties_df = pd.DataFrame(results)
    df_out = df_main[["uniprot_id", "sequence"]].join(properties_df)
    output_path = df_dir / "physicochem_properties.parquet"
    df_out.to_parquet(output_path, index=False)
    print(f"Saved physicochemical properties to {output_path}")
```
