## Supplementary material for "pLM representations unlock metagenomic space beyond homology": Code bundle: 02_pairwise_identity.py

```
"""
Calculates the pairwise sequence identity matrix for all sequences
using the high-speed 'mmseqs' tool.
"""

import argparse
import subprocess
from pathlib import Path

import numpy as np
import pandas as pd
from tqdm import tqdm
from utils import load_preprocess_data


if __name__ == "__main__":

    parser = argparse.ArgumentParser(description="Calculates a pairwise sequence identity matrix using MMseqs.")
    parser.add_argument(
        "--input_csv",
        type=str,
        default="data/uniprotkb_ec_1_14_11_2025_10_20.csv",
        help="Path to the input CSV file containing sequences.",
    )
    parser.add_argument(
        "--output_dir",
        type=str,
        default="output",
        help="Main directory to store all outputs.",
    )
    parser.add_argument(
        "--n_samples",
        type=int,
        default=0,
        help="Number of sequences to sample. Set to 0 to disable.",
    )
    parser.add_argument(
        "--cores",
        type=int,
        default=8,
        help="Number of CPU cores/threads to use for MMseqs.",
    )
    parser.add_argument(
        "--mmseqs_path",
        type=str,
        default="data/mmseqs/bin/mmseqs",
        help="Path to the mmseqs binary.",
    )
    args = parser.parse_args()

    # 1. Setup Paths
    output_dir = Path(args.output_dir)
    df_dir = output_dir / "dataframes"
    tmp_dir = output_dir / "mmseqs_tmp"
    output_dir.mkdir(exist_ok=True)
    df_dir.mkdir(exist_ok=True)
    tmp_dir.mkdir(exist_ok=True)

    mmseqs_bin = Path(args.mmseqs_path)
    if not mmseqs_bin.is_file():
        print(f"Error: MMseqs binary not found at: {mmseqs_bin}")
        print("Please check the --mmseqs_path argument.")
        exit(1)

    # Define file paths
    fasta_file = tmp_dir / "input.fasta"
    db_path = tmp_dir / "protein_db"
    search_res = tmp_dir / "search_res"
    results_m8 = tmp_dir / "results.m8"

    # 2. Load Data and Create FASTA
    print(f"Loading data from {args.input_csv}...")
    df_main = load_preprocess_data(args.input_csv, args.n_samples)

    if args.n_samples > 0:
        print(f"Using a sample of {args.n_samples} sequences.")
    else:
        print(f"Using all {len(df_main)} sequences.")

    sequences = df_main["sequence"].tolist()
    uniprot_ids = df_main["uniprot_id"].tolist()
    n = len(uniprot_ids)

    print(f"Creating FASTA file at {fasta_file}...")
    with open(fasta_file, "w") as f:
        for _, row in df_main.iterrows():
            f.write(f">{row['uniprot_id']}\n")
            f.write(f"{row['sequence']}\n")

    # Create database
    print("Creating MMseqs database")
    cmd_createdb = [str(mmseqs_bin), "createdb", str(fasta_file), str(db_path)]
    subprocess.run(cmd_createdb, check=True, text=True, capture_output=True)

    # Run all-vs-all search
    print("Running all-vs-all search")
    cmd_search = [
        str(mmseqs_bin),
        "search",
        str(db_path),
        str(db_path),
        str(search_res),
        str(tmp_dir),
        "-s",
        "7.0",
        "--threads",
        str(args.cores),
    ]
    subprocess.run(cmd_search, check=True, text=True, capture_output=True)

    # Convert results to BLAST-like .m8 table
    print("Converting results to BLAST-like .m8 format")
    cmd_convert = [
        str(mmseqs_bin),
        "convertalis",
        str(db_path),
        str(db_path),
        str(search_res),
        str(results_m8),
        "--format-output",
        "query,target,pident",
        "--threads",
        str(args.cores),
    ]
    subprocess.run(cmd_convert, check=True, text=True, capture_output=True)

    # Parse Results and Build Matrix
    print(f"\nParsing results from {results_m8}...")

    # Use a dictionary to map string IDs to integer indices
    id_to_index = {uniprot_id: i for i, uniprot_id in enumerate(uniprot_ids)}

    # Initialize a dense identity matrix (1.0 on diagonal, 0.0 elsewhere)
    identity_matrix = np.identity(n)

    # Read the sparse results
    df_results = pd.read_csv(results_m8, sep="\t", header=None, names=["query", "target", "pident"])

    print("Building dense identity matrix...")
    for _, row in tqdm(df_results.iterrows(), total=len(df_results), desc="Populating matrix"):
        # Skip self-hits (already set to 1.0)
        if row["query"] == row["target"]:
            continue

        try:
            i = id_to_index[row["query"]]
            j = id_to_index[row["target"]]

            # pident is 0-1, not 0-100
            identity = row["pident"]

            identity_matrix[i, j] = identity
            identity_matrix[j, i] = identity
        except KeyError:
            # This can happen if an ID in the .m8 file isn't in our list
            print(f"Warning: Skipping unknown ID pair ({row['query']}, {row['target']})")
            pass

    # Save Final DataFrame
    identity_df = pd.DataFrame(identity_matrix, columns=uniprot_ids, index=uniprot_ids)
    output_path = df_dir / "pairwise_identity.parquet"
    identity_df.to_parquet(output_path, index=False)
    print(f"Saved pairwise distance to {output_path}")
```
