## Supplementary material for "pLM representations unlock metagenomic space beyond homology": Code bundle: 03_plm_embeddings.py

```
"""
Calculates pLM embeddings and intra-sequence cosine similarity
for a single pLM model specified via command-line arguments.
"""

import argparse
from functools import partial
from pathlib import Path

import torch
import torch.nn.functional as F
from torch.nn.utils.rnn import pad_sequence
from torch.utils.data import DataLoader
from tqdm import tqdm
from transformers import AutoModel, AutoTokenizer
from utils import load_preprocess_data


def collate_fn_singles(batch, tokenizer):
    """Pads sequences for a single-input model."""
    inputs = pad_sequence(batch, batch_first=True, padding_value=tokenizer.pad_token_id)
    # Pad to a multiple of 8 for tensor core efficiency
    pad = (8 - inputs.size(1) % 8) % 8
    if pad > 0:
        inputs = F.pad(inputs, (0, pad), value=tokenizer.pad_token_id)
    return inputs, (inputs != tokenizer.pad_token_id)


def calculate_intra_sequence_similarity(hidden_states, mask):
    """Calculates the mean pairwise cosine similarity of non-padded token embeddings."""
    similarities = []
    for j in range(hidden_states.shape[0]):
        real_embeddings = hidden_states[j][mask[j]]
        norm_embeddings = F.normalize(real_embeddings, p=2, dim=1)
        cosine_sim_matrix = torch.mm(norm_embeddings, norm_embeddings.T)
        n = cosine_sim_matrix.shape[0]
        mean_sim = (cosine_sim_matrix.sum() - n) / (n * (n - 1))
        similarities.append(mean_sim.item())
    return similarities


if __name__ == "__main__":

    parser = argparse.ArgumentParser(description="Calculates pLM embeddings and intra-sequence similarity for a specified model.")
    parser.add_argument(
        "--input_csv",
        type=str,
        default="data/uniprotkb_ec_1_14_11_2025_10_20.csv",
        help="Path to the input CSV file containing sequences.",
    )
    parser.add_argument(
        "--output_dir",
        type=str,
        default="output",
        help="Main directory to store all outputs.",
    )
    parser.add_argument(
        "--n_samples",
        type=int,
        default=0,
        help="Number of sequences to sample. Set to 0 to disable.",
    )
    parser.add_argument(
        "--batch_size",
        type=int,
        default=128,
        help="Batch size for pLM inference.",
    )
    parser.add_argument(
        "--max_len",
        type=int,
        default=1024,
        help="Maximum sequence length.",
    )
    parser.add_argument(
        "--data_name",
        type=str,
        default="enzymes",
        help="Name of the data (used for output file naming).",
    )
    parser.add_argument(
        "--model_name",
        type=str,
        default="AMPLIFY_120M",
        help="Name of the pLM model (used for output file naming).",
    )
    parser.add_argument(
        "--model_path",
        type=str,
        default="chandar-lab/AMPLIFY_120M",
        help="Path or HuggingFace name of the pLM model to load.",
    )
    parser.add_argument(
        "--token",
        type=str,
        required=False,
        default=None,
        help="HuggingFace token.",
    )
    parser.add_argument(
        "--mask_type",
        type=str,
        default="additive",
        choices=["additive", "boolean"],
        help="Type of attention mask required by the model.",
    )
    parser.add_argument(
        "--device",
        type=str,
        default="cuda:0",
        help="Specify the GPU device to use (e.g., 'cuda:0', 'cuda:1').",
    )
    args = parser.parse_args()

    output_dir = Path(args.output_dir)
    df_dir = output_dir / "dataframes"
    output_dir.mkdir(exist_ok=True)
    df_dir.mkdir(exist_ok=True)

    df_main = load_preprocess_data(args.input_csv, args.n_samples)

    df_out = df_main[["uniprot_id", "sequence"]].copy()

    print(f"Loading {args.model_name} model ({args.model_path}) on {args.device}...")
    model = AutoModel.from_pretrained(args.model_path, trust_remote_code=True).to(args.device).eval()
    tokenizer = AutoTokenizer.from_pretrained(args.model_path, trust_remote_code=True)

    print("Tokenizing sequences...")
    tokenized = [
        tokenizer.encode(str(s), max_length=args.max_len, truncation=True, return_tensors="pt").squeeze(0)
        for s in tqdm(df_main.sequence, desc=f"Tokenizing for {args.model_name}")
    ]

    current_collate_fn = partial(collate_fn_singles, tokenizer=tokenizer)
    dataloader = DataLoader(tokenized, batch_size=args.batch_size, collate_fn=current_collate_fn)

    final_layer_embeddings = []
    final_layer_similarities = []

    print(f"Generating final layer embeddings and similarities for {args.model_name}...")
    with torch.no_grad(), torch.autocast(device_type="cuda", dtype=torch.float16):
        for ids, mask in tqdm(dataloader, desc=f"Generating Embeddings ({args.model_name})"):
            ids, mask = ids.to(args.device), mask.to(args.device)

            if args.mask_type == "additive":
                attention_mask = torch.where(mask, 0.0, float("-inf")).to(device=args.device, dtype=torch.float16)
            else:
                attention_mask = mask.to(device=args.device, dtype=torch.int)

            model_output = model(ids, attention_mask=attention_mask, output_hidden_states=True)
            hidden_states = model_output.hidden_states[-1]
            
            pooled_embedding = (hidden_states * mask.unsqueeze(-1)).sum(1) / torch.clamp(mask.sum(1, keepdim=True), min=1)
            final_layer_embeddings.extend(pooled_embedding.cpu().numpy())

            similarities = calculate_intra_sequence_similarity(hidden_states, mask)
            final_layer_similarities.extend(similarities)

    df_out[f"{args.model_name}_embedding"] = [emb.tolist() for emb in final_layer_embeddings]
    df_out[f"{args.model_name}_intra_similarity"] = final_layer_similarities

    output_path = df_dir / f"plm_embeddings_{args.data_name}_{args.model_name}.parquet"
    df_out.to_parquet(output_path, index=False)
    print(f"Saved embeddings to {output_path}")
```
