## Supplementary material for "pLM representations unlock metagenomic space beyond homology": Code bundle: 04_plm_perplexity.py

```
import os
import torch
import argparse
from pathlib import Path
import numpy as np

# HuggingFace
from transformers import AutoTokenizer, AutoModel

# Ours
from utils import PseudoPPLEvaluator


if __name__ == "__main__":

    parser = argparse.ArgumentParser(description="Calculates pseudo-perplexity (PPL) for a specified model.")
    parser.add_argument(
        "--input_csv",
        type=str,
        default="data/uniprotkb_ec_1_14_11_2025_10_20.csv",
        help="Path to the input CSV file containing sequences.",
    )
    parser.add_argument(
        "--data_name",
        type=str,
        default="uniprotkb_ec_1_14_11_2025_10_20",
        help="Name of the dataset.",
    )
    parser.add_argument(
        "--output_dir",
        type=str,
        default="output",
        help="Main directory to store all outputs.",
    )
    parser.add_argument(
        "--n_samples",
        type=int,
        default=0,
        help="Number of sequences to sample. Set to 0 to disable.",
    )
    parser.add_argument(
        "--max_len",
        type=int,
        default=2048,
        help="Maximum sequence length.",
    )
    parser.add_argument(
        "--batch_size",
        type=int,
        default=128,
        help="Batch size for pLM inference.",
    )
    parser.add_argument(
        "--save_steps",
        type=int,
        default=10000,
        help="Interval between checkpointing.",
    )
    parser.add_argument(
        "--model_name",
        type=str,
        default="AMPLIFY_120M",
        help="Name of the pLM model (used for output file naming).",
    )
    parser.add_argument(
        "--model_path",
        type=str,
        default="chandar-lab/AMPLIFY_120M",
        help="Path or HuggingFace name of the pLM model to load.",
    )
    parser.add_argument(
        "--token",
        type=str,
        default=None,
        help="HuggingFace token.",
    )
    parser.add_argument(
        "--packing",
        action="store_true",
        help="Use the packing implementation of flash attention.",
    )
    parser.add_argument("--compile", action="store_true", help="Enable torch.compile for the model (requires PyTorch 2.0+).")
    args = parser.parse_args()

    output_dir = Path(args.output_dir) / "pseudo_ppl" / args.model_name / args.data_name
    os.makedirs(output_dir, exist_ok=True)
    
    evaluator = PseudoPPLEvaluator(batch_size=args.batch_size, packing=args.packing)

    if "AMPLIFY" in args.model_name:
        model = AutoModel.from_pretrained(args.model_path, trust_remote_code=True, token=args.token)
    else:
        print(f"Error: Model {args.model_name} not supported for PPL. Add logic to this script.")
        exit(1)
    tokenizer = AutoTokenizer.from_pretrained(args.model_path, trust_remote_code=True)

    device = torch.device("cuda" if torch.cuda.is_available() else "cpu")
    model.to(device)

    if args.compile:
        model = torch.compile(model)

    evaluator(
        model=model,
        tokenizer=tokenizer,
        input_csv=args.input_csv,
        max_len=args.max_len,
        model_name=args.model_name,
        data_name=args.data_name,
        output_dir=output_dir,
    )

    ppl = evaluator._compute_metric(
        model_name=args.model_name,
        output_dir=output_dir,
    )

    ppl.to_parquet(output_dir / f"plm_perplexity_{args.model_name}_{args.data_name}.parquet", index=False)
```
