## Supplementary material for "pLM representations unlock metagenomic space beyond homology": Code bundle: 05_structural_properties.py

```
"""
Downloads AlphaFold PDB files (v6) and computes structural properties:
- Mean pLDDT (from B-factor)
- Mean Solvent Accessible Surface Area (SASA)
"""

import argparse
from collections import Counter
from concurrent.futures import ProcessPoolExecutor, as_completed
from pathlib import Path

import numpy as np
import pandas as pd
from Bio.PDB import PDBParser
from Bio.PDB.SASA import ShrakeRupley
from tqdm import tqdm
from utils import download_structure, load_preprocess_data


def compute_structure_properties(pdb_file_path):
    """
    Parses a PDB file to compute mean pLDDT and mean solvent accessibility (SASA).
    Assumes B-factor column stores pLDDT (common for AlphaFold).
    """
    if not Path(pdb_file_path).exists():
        return pd.Series({"mean_plddt": np.nan, "mean_sasa": np.nan})

    # try:
    parser = PDBParser(QUIET=True)
    structure = parser.get_structure("protein", pdb_file_path)
    model = structure[0]

    plddt_scores = np.array([atom.get_bfactor() for atom in model.get_atoms() if atom.get_id() == "CA"], dtype=float).flatten()
    mean_plddt = np.mean(plddt_scores)

    # Compute disorder fractions based on pLDDT thresholds
    fraction_disordered_05 = np.mean(plddt_scores < 50)
    fraction_disordered_08 = np.mean(plddt_scores < 80)
    fraction_disordered_09 = np.mean(plddt_scores < 90)

    sr = ShrakeRupley()
    sr.compute(structure, level="R")

    residue_sasa = [res.sasa for res in model.get_residues() if hasattr(res, "sasa") and res.sasa is not None]
    mean_sasa = np.mean(residue_sasa) if residue_sasa else np.nan

    return pd.Series(
        {
            "mean_plddt": mean_plddt,
            "mean_sasa": mean_sasa,
            "fraction_disordered_05": fraction_disordered_05,
            "fraction_disordered_08": fraction_disordered_08,
            "fraction_disordered_09": fraction_disordered_09,
        }
    )


if __name__ == "__main__":

    parser = argparse.ArgumentParser(description="Downloads PDBs and computes pLDDT and SASA.")
    parser.add_argument(
        "--input_csv",
        type=str,
        default="data/uniprotkb_ec_1_14_11_2025_10_20.csv",
        help="Path to the input CSV file containing sequences.",
    )
    parser.add_argument(
        "--output_dir",
        type=str,
        default="output",
        help="Main directory to store all outputs.",
    )
    parser.add_argument(
        "--n_samples",
        type=int,
        default=0,
        help="Number of sequences to sample. Set to 0 to disable.",
    )
    parser.add_argument(
        "--cores",
        type=int,
        default=8,
        help="Number of CPU cores to use for parallel processing.",
    )
    args = parser.parse_args()

    output_dir = Path(args.output_dir)
    df_dir = output_dir / "dataframes"
    pdb_dir = output_dir / "pdb_files"
    output_dir.mkdir(exist_ok=True)
    df_dir.mkdir(exist_ok=True)
    pdb_dir.mkdir(exist_ok=True)

    # Use shared function to load and preprocess data
    df_main = load_preprocess_data(args.input_csv, args.n_samples)

    print("Starting parallel download of AlphaFold structures...")
    ids_to_download = df_main["uniprot_id"].unique()
    counts = Counter()

    with ProcessPoolExecutor(max_workers=args.cores) as executor:
        futures = {executor.submit(download_structure, uid, pdb_dir): uid for uid in ids_to_download}
        for future in tqdm(as_completed(futures), total=len(futures), desc="Downloading PDBs"):
            uniprot_id, status = future.result()
            counts[status] += 1
    print(f"Download complete. Summary: {counts}")

    df_main["pdb_file"] = df_main["uniprot_id"].apply(lambda x: pdb_dir / f"AF-{x}-F1-model_v6.pdb")
    df_main["pdb_exists"] = df_main["pdb_file"].apply(lambda x: x.exists())

    print("Computing structural properties from PDB files (in parallel)...")
    with ProcessPoolExecutor(max_workers=args.cores) as executor:
        pdb_files_to_process = df_main["pdb_file"].tolist()

        results = list(
            tqdm(
                executor.map(compute_structure_properties, pdb_files_to_process),
                total=len(pdb_files_to_process),
                desc="Computing properties",
            )
        )

    df_main["pdb_file"] = df_main["pdb_file"].apply(lambda x: str(x))

    structure_properties_df = pd.DataFrame(results)
    df_out = df_main[["uniprot_id", "sequence", "pdb_file", "pdb_exists"]].join(structure_properties_df)
    output_path = df_dir / "structural_properties.parquet"
    df_out.to_parquet(output_path, index=False)
    print(f"Saved structural properties to {output_path}")
```
