## Supplementary material for "pLM representations unlock metagenomic space beyond homology": Code bundle: 06_signalp.py

```
"""
Runs SignalP-6.0 to predict signal peptide likelihood.
"""

import argparse
import json
import subprocess
from pathlib import Path

import numpy as np
import pandas as pd

from utils import load_preprocess_data

if __name__ == "__main__":

    parser = argparse.ArgumentParser(description="Runs SignalP 6.0 on a set of sequences.")
    parser.add_argument(
        "--input_csv",
        type=str,
        default="data/uniprotkb_ec_1_14_11_2025_10_20.csv",
        help="Path to the input CSV file containing sequences.",
    )
    parser.add_argument(
        "--output_dir",
        type=str,
        default="output",
        help="Main directory to store all outputs.",
    )
    parser.add_argument(
        "--n_samples",
        type=int,
        default=0,
        help="Number of sequences to sample. Set to 0 to disable.",
    )

    args = parser.parse_args()

    output_dir = Path(args.output_dir)
    df_dir = output_dir / "dataframes"
    tmp_dir = output_dir / "tmp_files"
    output_dir.mkdir(exist_ok=True)
    df_dir.mkdir(exist_ok=True)
    tmp_dir.mkdir(exist_ok=True)

    df_main = load_preprocess_data(args.input_csv, args.n_samples)

    fasta_file_path = tmp_dir / "temp_signalp_input.fasta"
    print(f"Writing {len(df_main)} sequences to {fasta_file_path}...")
    with open(fasta_file_path, "w") as f:
        for _, row in df_main.iterrows():
            f.write(f">{row['uniprot_id']}\n{row['sequence']}\n")

    df_out = df_main[["uniprot_id", "sequence"]].copy()

    local_output_dir = tmp_dir / "signalp_local_out"
    local_output_dir.mkdir(exist_ok=True)

    # Call the local function
    print("Running SignalP 6.0...")
    try:
        cmd = [
            "signalp6",
            "--fastafile",
            str(fasta_file_path),
            "--output_dir",
            str(local_output_dir),
            "--mode",
            "fast",
            "--bsize",
            "64",
        ]
        subprocess.run(cmd, check=True, capture_output=True, text=True, encoding="utf-8")
    except subprocess.CalledProcessError as e:
        print(f"Error running SignalP 6.0:")
        print(e.stderr)
        exit(1)

    print("SignalP run complete. Processing results...")
    with open(local_output_dir / "output.json", "r") as f:
        signalp_results_json = json.load(f)

    sequences_data = signalp_results_json["SEQUENCES"]
    df_signalp = pd.DataFrame.from_dict(sequences_data, orient="index")

    df_signalp["sp_likelihood"] = df_signalp["Likelihood"].apply(
        lambda x: x[1] if isinstance(x, list) and len(x) > 1 else np.nan
    )
    df_out = df_out.merge(
        df_signalp[["sp_likelihood"]],
        left_on="uniprot_id",
        right_index=True,
        how="left",
    )

    output_path = df_dir / "signalp_scores.parquet"
    df_out.to_parquet(output_path, index=False)
    print(f"Saved SignalP scores to {output_path}")
```
