## Supplementary material for "pLM representations unlock metagenomic space beyond homology": Code bundle: 07_deeptmhmm.py

```
#!/usr/bin/env python

"""
Runs DeepTMHMM via biolib to predict transmembrane topology.

This script is refactored to use argparse and conventions from
the 08_proteinmpnn.py script.
"""

import argparse
import sys
from collections import Counter
from pathlib import Path

import biolib
import numpy as np
import pandas as pd
from tqdm import tqdm
from utils import load_preprocess_data


if __name__ == "__main__":
    parser = argparse.ArgumentParser(description="Runs DeepTMHMM via biolib to predict topology.")
    parser.add_argument(
        "--input_csv",
        type=str,
        default="data/uniprotkb_ec_1_14_11_2025_10_20.csv",
        help="Path to the input CSV file containing sequences.",
    )
    parser.add_argument(
        "--output_dir",
        type=str,
        default="output",
        help="Main directory to store all outputs.",
    )
    parser.add_argument(
        "--n_samples",
        type=int,
        default=0,
        help="Number of sequences to sample. Set to 0 to disable.",
    )

    args = parser.parse_args()

    output_dir = Path(args.output_dir)
    df_dir = output_dir / "dataframes"
    tmp_dir = output_dir / "tmp_files"

    output_dir.mkdir(exist_ok=True)
    df_dir.mkdir(exist_ok=True)
    tmp_dir.mkdir(exist_ok=True)

    # Load and preprocess data using the shared function
    print("Loading base dataset...")
    df_main = load_preprocess_data(args.input_csv, args.n_samples)

    # Write sequences to a temporary FASTA file
    fasta_file_path = tmp_dir / "temp_deeptmhmm_input.fasta"
    print(f"Writing {len(df_main)} sequences to {fasta_file_path}...")
    with open(fasta_file_path, "w") as f:
        for _, row in df_main.iterrows():
            f.write(f">{row['uniprot_id']}\n{row['sequence']}\n")

    print("Loading DeepTMHMM model via biolib...")
    deeptmhmm = biolib.load("DTU/DeepTMHMM")

    print("Running DeepTMHMM analysis...")
    deeptmhmm_results = deeptmhmm.run(fasta=str(fasta_file_path))
    deeptmhmm_results.save_files(output_dir=tmp_dir, overwrite=True)

    print("DeepTMHMM run complete. Processing results...")
    deep_file_path = tmp_dir / "predicted_topologies.3line"

    print(f"Processing region details from {deep_file_path}...")

    parsed_data = []
    with open(deep_file_path, "r") as f:
        lines = f.readlines()

    for i in range(0, len(lines), 3):
        if i + 2 >= len(lines):
            continue
        header_line = lines[i].strip()
        # line i+1 is sequence, line i+2 is topology
        topology_string = lines[i + 2].strip()

        # Parse header, e.g., >sp|P01111|RAS_HUMAN | Type=Globular
        header_content, protein_type = header_line.lstrip(">").split(" | ")
        entry_id = header_content.strip().split("|")[1]  # Get Uniprot ID
        protein_type = protein_type.strip()
        region_counts = Counter(topology_string)

        parsed_data.append(
            {
                "uniprot_id": entry_id,
                "deeptmhmm_type": protein_type,
                "deeptmhmm_sp": region_counts.get("S", 0),
                "deeptmhmm_tmalpha": region_counts.get("M", 0),
                "deeptmhmm_tmbeta": region_counts.get("B", 0),
                "deeptmhmm_periplasm": region_counts.get("P", 0),
                "deeptmhmm_inside": region_counts.get("I", 0),
                "deeptmhmm_outside": region_counts.get("O", 0),
            },
        )

    df_deep = pd.DataFrame(parsed_data)

    # Merge results into the DataFrame
    df_main = df_main.merge(df_deep, on="uniprot_id", how="left")

    # Save the updated dataframe as Parquet
    print("Saving final dataframe with all DeepTMHMM scores...")
    output_path = df_dir / "deeptmhmm_scores.parquet"
    df_main.to_parquet(output_path, index=False)
    print(f"Successfully saved DeepTMHMM scores to {output_path}")
```
