## Supplementary material for "pLM representations unlock metagenomic space beyond homology": Code bundle: 08_proteinmpnn.py

```
"""
Downloads AlphaFold PDB files (v6) and then runs ProteinMPNN
(vanilla or soluble models) to compute native sequence log-likelihood.
"""

import argparse
import sys
from collections import Counter
from concurrent.futures import ProcessPoolExecutor, as_completed
from pathlib import Path

import numpy as np
import pandas as pd
from tqdm import tqdm

from utils import download_structure, load_preprocess_data

if __name__ == "__main__":
    parser = argparse.ArgumentParser(description="Downloads PDBs and scores them with ProteinMPNN.")
    parser.add_argument(
        "--input_csv",
        type=str,
        default="data/uniprotkb_ec_1_14_11_2025_10_20.csv",
        help="Path to the input CSV file containing sequences.",
    )
    parser.add_argument(
        "--output_dir",
        type=str,
        default="output",
        help="Main directory to store all outputs.",
    )
    parser.add_argument(
        "--n_samples",
        type=int,
        default=0,
        help="Number of sequences to sample. Set to 0 to disable.",
    )
    parser.add_argument(
        "--cores",
        type=int,
        default=8,
        help="Number of CPU cores to use for parallel PDB download.",
    )
    parser.add_argument(
        "--proteinmpnn_dir",
        type=str,
        default="data/ProteinMPNN",
        help="Path to the cloned ProteinMPNN repository.",
    )
    parser.add_argument(
        "--model_version",
        type=str,
        default="v_48_020",
        help="ProteinMPNN model version to use.",
    )
    parser.add_argument(
        "--model_family",
        type=str,
        default="vanilla",
        choices=["vanilla", "soluble"],
        help="ProteinMPNN model family to use.",
    )

    args = parser.parse_args()

    output_dir = Path(args.output_dir)
    df_dir = output_dir / "dataframes"
    pdb_dir = output_dir / "pdb_files"
    tmp_dir = output_dir / "tmp_files"

    output_dir.mkdir(exist_ok=True)
    df_dir.mkdir(exist_ok=True)
    pdb_dir.mkdir(exist_ok=True)
    tmp_dir.mkdir(exist_ok=True)

    # Import ProteinMPNN from specified directory
    proteinmpnn_path = Path(args.proteinmpnn_dir)
    sys.path.append(args.proteinmpnn_dir)
    import protein_mpnn_run

    df_main = load_preprocess_data(args.input_csv, args.n_samples)

    print("Starting parallel download of AlphaFold structures...")
    ids_to_download = df_main["uniprot_id"].unique()
    counts = Counter()

    with ProcessPoolExecutor(max_workers=args.cores) as executor:
        futures = {executor.submit(download_structure, uid, pdb_dir): uid for uid in ids_to_download}
        for future in tqdm(as_completed(futures), total=len(futures), desc="Downloading PDBs"):
            uniprot_id, status = future.result()
            counts[status] += 1
    print(f"Download complete. Summary: {counts}")

    df_main["pdb_file"] = df_main["uniprot_id"].apply(lambda x: pdb_dir / f"AF-{x}-F1-model_v6.pdb")
    df_main["pdb_exists"] = df_main["pdb_file"].apply(lambda x: x.exists())
    df_main["pdb_file"] = df_main["pdb_file"].astype(str)

    df_out = df_main[["uniprot_id", "sequence", "pdb_file", "pdb_exists"]].copy()

    family = args.model_family
    print(f"Running ProteinMPNN scoring for '{family}' model...")

    model_weights_path = proteinmpnn_path / f"{family}_model_weights"
    if not model_weights_path.is_dir():
        print(f"Error: Model weights not found at {model_weights_path}")
        print(f"Please ensure the '{family}' model weights are downloaded.")
        exit(1)

    base_args = argparse.Namespace(
        path_to_model_weights=str(model_weights_path),
        model_name=args.model_version,
        score_only=1,
        out_folder=str(tmp_dir),
        suppress_print=1,
        use_soluble_model=(family == "soluble"),
        seed=0,
        ca_only=False,
        save_score=0,
        path_to_fasta="",
        save_probs=0,
        conditional_probs_only=0,
        conditional_probs_only_backbone=0,
        unconditional_probs_only=0,
        backbone_noise=0.00,
        num_seq_per_target=1,
        batch_size=1,
        max_length=200000,
        sampling_temp="0.1",
        pdb_path="",
        pdb_path_chains="",
        jsonl_path="",
        chain_id_jsonl="",
        fixed_positions_jsonl="",
        omit_AAs="X",
        bias_AA_jsonl="",
        bias_by_res_jsonl="",
        omit_AA_jsonl="",
        pssm_jsonl="",
        pssm_multi=0.0,
        pssm_threshold=0.0,
        pssm_log_odds_flag=0,
        pssm_bias_flag=0,
        tied_positions_jsonl="",
    )

    mpnn_data = []

    for index, row in tqdm(df_main.iterrows(), total=len(df_main), desc=f"Scoring with {family} MPNN"):
        current_score = np.nan
        if row.get("pdb_exists", False) and pd.notna(row.get("pdb_file")):

            # Define the expected output file path
            pdb_path = Path(row["pdb_file"])
            expected_output_file = tmp_dir / "score_only" / f"{pdb_path.stem}_pdb.npz"

            # Clean up stale file from previous (or failed) runs
            if expected_output_file.exists():
                expected_output_file.unlink()

            # Create a copy of the args for this specific run
            run_args = argparse.Namespace(**vars(base_args))
            run_args.pdb_path = str(pdb_path)

            # Run the main function
            protein_mpnn_run.main(run_args)

            # Read the score from the file
            data = np.load(expected_output_file, allow_pickle=True)
            current_score = data["score"][0]

        mpnn_data.append({"uniprot_id": row.get("uniprot_id"), f"mpnn_ll_{family}": current_score})

    mpnn_df = pd.DataFrame(mpnn_data)
    df_out = df_out.merge(mpnn_df, on="uniprot_id", how="left")

    output_path = df_dir / f"proteinmpnn_scores_{family}.parquet"
    df_out.to_parquet(output_path, index=False)
    print(f"Saved ProteinMPNN scores to {output_path}")
```
