## Supplementary material for "pLM representations unlock metagenomic space beyond homology": Code bundle: 09_esm_if.py

```
"""
Downloads AlphaFold PDB files (v6) and then runs ESM-IF
to compute native sequence negative log-likelihood (NLL).
"""

import argparse
from collections import Counter
from concurrent.futures import ProcessPoolExecutor, as_completed
from pathlib import Path

import esm
import numpy as np
import pandas as pd
from esm.inverse_folding.util import extract_coords_from_structure, load_structure, score_sequence
from tqdm import tqdm
from utils import download_structure, load_preprocess_data

if __name__ == "__main__":
    parser = argparse.ArgumentParser(description="Downloads PDBs and scores them with ESM-IF.")
    parser.add_argument(
        "--input_csv",
        type=str,
        default="data/uniprotkb_ec_1_14_11_2025_10_20.csv",
        help="Path to the input CSV file containing sequences.",
    )
    parser.add_argument(
        "--output_dir",
        type=str,
        default="output",
        help="Main directory to store all outputs.",
    )
    parser.add_argument(
        "--n_samples",
        type=int,
        default=0,
        help="Number of sequences to sample. Set to 0 to disable.",
    )
    parser.add_argument(
        "--cores",
        type=int,
        default=8,
        help="Number of CPU cores to use for parallel PDB download.",
    )
    parser.add_argument(
        "--device",
        type=str,
        default="cuda:0",
        help="Specify the GPU device to use (e.g., 'cuda:0').",
    )

    args = parser.parse_args()

    output_dir = Path(args.output_dir)
    df_dir = output_dir / "dataframes"
    pdb_dir = output_dir / "pdb_files"

    output_dir.mkdir(exist_ok=True)
    df_dir.mkdir(exist_ok=True)
    pdb_dir.mkdir(exist_ok=True)

    # Use shared function to load and preprocess data
    df_main = load_preprocess_data(args.input_csv, args.n_samples)

    print("Starting parallel download of AlphaFold structures...")
    ids_to_download = df_main["uniprot_id"].unique()
    counts = Counter()

    with ProcessPoolExecutor(max_workers=args.cores) as executor:
        futures = {executor.submit(download_structure, uid, pdb_dir): uid for uid in ids_to_download}
        for future in tqdm(as_completed(futures), total=len(futures), desc="Downloading PDBs"):
            uniprot_id, status = future.result()
            counts[status] += 1
    print(f"Download complete. Summary: {counts}")

    df_main["pdb_file"] = df_main["uniprot_id"].apply(lambda x: pdb_dir / f"AF-{x}-F1-model_v6.pdb")
    df_main["pdb_exists"] = df_main["pdb_file"].apply(lambda x: x.exists())

    df_out = df_main[["uniprot_id", "sequence", "pdb_file", "pdb_exists"]].copy()

    print(f"Loading ESM-IF model to {args.device}...")
    model_if, alphabet_if = esm.pretrained.esm_if1_gvp4_t16_142M_UR50()
    model_if = model_if.eval().to(args.device)

    esm_if_data = []
    for index, row in tqdm(df_main.iterrows(), total=len(df_main), desc="Scoring with ESM-IF"):
        current_score = np.nan
        if row.get("pdb_exists", False) and pd.notna(row.get("pdb_file")):
            pdb_path = str(row["pdb_file"])
            try:
                structure = load_structure(pdb_path)
                coords, native_seq_from_pdb = extract_coords_from_structure(structure)

                # Compare PDB sequence to UniProt sequence
                if native_seq_from_pdb != row["sequence"]:
                    print(f"Warning: PDB sequence mismatch for {row['uniprot_id']}. Scoring PDB sequence.")

                current_score, _ = score_sequence(model_if, alphabet_if, coords, native_seq_from_pdb)
                current_score = current_score.item()
            except Exception as e:
                print(f"Warning: ESM-IF failed for {row['uniprot_id']}. Error: {e}")

        esm_if_data.append({"uniprot_id": row.get("uniprot_id"), "esm_if_nll": current_score})

    esm_if_df = pd.DataFrame(esm_if_data)
    df_out = df_out.merge(esm_if_df, on="uniprot_id", how="left")

    output_path = df_dir / "esm_if_scores.parquet"
    df_out.to_parquet(output_path, index=False)
    print(f"Saved ESM-IF scores to {output_path}")
```
