## Supplementary material for "pLM representations unlock metagenomic space beyond homology": Code bundle: 10_pairwise_cossim.py

```
"""
Calculates the pairwise sequence identity matrix for all sequences
using the high-speed 'mmseqs' tool.
"""

import ast
import argparse
from pathlib import Path

import numpy as np
import pandas as pd
from sklearn.preprocessing import normalize


if __name__ == "__main__":

    parser = argparse.ArgumentParser(description="Calculates a pairwise sequence cosine similarity matrix using PLM embeddings.")
    parser.add_argument(
        "--input_parquet",
        type=str,
        default="dataframes/plm_embeddings_AMPLIFY-C.parquet",
        help="Path to the input parquet file containing embeddings.",
    )
    parser.add_argument(
        "--output_dir",
        type=str,
        default="output",
        help="Main directory to store all outputs.",
    )
    parser.add_argument(
        "--n_samples",
        type=int,
        default=0,
        help="Number of sequences to sample. Set to 0 to disable.",
    )
    parser.add_argument(
        "--embedding_col",
        type=str,
        default="AMPLIFY-C_embedding",
    )
    args = parser.parse_args()

    # 1. Setup Paths
    output_dir = Path(args.output_dir)
    df_dir = output_dir / "dataframes"
    output_dir.mkdir(exist_ok=True)
    df_dir.mkdir(exist_ok=True)

    # 2. Load Data
    print(f"Loading data from {args.input_parquet}.")
    df = pd.read_parquet(df_dir / args.input_parquet)

    if args.n_samples > 0:
        print(f"Using a sample of {args.n_samples} sequences.")
        df = df.sample(n=args.n_samples, random_state=42).reset_index(drop=True)
    else:
        print(f"Using all {len(df)} sequences.")

    col = next((c for c in df.columns if "embedding" in c), None)
    id_col = next((c for c in df.columns if "id" in c), None)

    if col is None:
        raise ValueError("No embedding column found in the input parquet file.")

    def safe_parse_embedding(x):
        if isinstance(x, (list, np.ndarray)):
            return x
        if isinstance(x, str):
            x = x.replace("array(", "").replace(")", "")
            return ast.literal_eval(x)
        return x

    embeddings = df[col].apply(safe_parse_embedding)

    # 3. Compute pairwise cosine similarity: result = M . M^T
    print("Normalizing embeddings")
    matrix = np.vstack(embeddings.tolist()).astype(np.float32)
    norm_matrix = normalize(matrix, axis=1, norm="l2")

    print("Computing cosine similarity matrix")
    similarity_matrix = np.dot(norm_matrix, norm_matrix.T)

    # Zero out lower triangle and diagonal (k=1) for compression efficiency
    # Diagonal is known to be 1.0, Lower Triangle is symmetric to Upper.
    similarity_matrix = np.triu(similarity_matrix, k=1)

    # 4. Create DataFrame with IDs as index and columns
    print("Creating dataframe")
    ids = df[id_col].values
    sim_df = pd.DataFrame(similarity_matrix, index=ids, columns=ids)

    # 5. Save to Parquet
    print("Saving dataframe to parquet")
    output_path = df_dir / f"pairwise_cossim_{col}.parquet"
    sim_df.to_parquet(output_path, engine="pyarrow", compression="snappy")

    print(f"Saved pairwise cosine similarities to {output_path}")
```
