## Supplementary material for "pLM representations unlock metagenomic space beyond homology": Code bundle: 12_openfold.py

```
"""Calculates pairwise structure similarity (RMSD) for a list of sequences using PyMOL and multiprocessing."""

import json
import argparse
import subprocess
from concurrent.futures import ProcessPoolExecutor
from pathlib import Path
from functools import partial

import pandas as pd
from tqdm import tqdm


def load_preprocess_data(input_csv, n_samples=0):
    """
    Loads the base dataset from a CSV, samples if needed,
    and preprocesses columns (name -> entry_name, extracts uniprot_id).
    """
    print(f"Loading base dataset from {input_csv}...")
    df_main = pd.read_csv(input_csv)

    if n_samples > 0:
        print(f"Using a subset of {n_samples} sequences.")
        df_main = df_main.sample(n=n_samples, random_state=42).reset_index(drop=True)

    if "name" in df_main.columns:
        df_main = df_main.rename(columns={"name": "entry_name"})

    # Ensure required columns exist
    if "entry_name" not in df_main.columns or "sequence" not in df_main.columns:
        print("Error: Input CSV must contain 'entry_name' (or 'name') and 'sequence' columns.")
        exit(1)

    try:
        df_main["uniprot_id"] = df_main["entry_name"].apply(lambda x: x.split("|")[1])
    except IndexError:
        df_main["uniprot_id"] = df_main["entry_name"]
    df_main["sequence"] = df_main["sequence"].astype(str)
    df_main["uniprot_id"] = df_main["uniprot_id"].astype(str)

    print(f"Loaded and preprocessed {len(df_main)} sequences.")
    return df_main


def run_openfold_prediction(output_dir, query_path):
    openfold_output_dir = output_dir / f"{str(query_path).split('/')[-1].split('_query')[0]}"
    
    # Check if prediction was already done for this query to avoid redundant computation
    if (openfold_output_dir).exists():
        print(f"Prediction already exists for {query_path}, skipping...")
        return {
            "query_path": str(query_path),
            "status": "skipped",
            "error": None,
        }
        
    cmd = ["run_openfold", "predict", f"--query_json={query_path}", f"--output_dir={output_dir}"]
    
    print(cmd)

    try:
        process = subprocess.run(cmd, capture_output=True, text=True, check=True)

        return {
            "query_path": str(query_path),
            "status": "success",
            "error": None,
        }

    except subprocess.CalledProcessError as e:
        return {
            "query_path": str(query_path),
            "status": "failed",
            "stderr": f"Command failed: {e.stderr}",
            "error": str(e),
        }

    except Exception as e:
        return {
            "query_path": str(query_path) if "query_path" in locals() else None,
            "status": "error",
            "error": str(e),
        }


if __name__ == "__main__":

    parser = argparse.ArgumentParser(description="Runs pairwise RMSD calculations on a set of sequences.")
    parser.add_argument(
        "--input_csv",
        type=str,
        default="data/uniprotkb_ec_1_14_11_2025_10_20.csv",
        help="Path to the input CSV file containing sequences.",
    )
    parser.add_argument(
        "--output_dir",
        type=str,
        default="output",
        help="Main directory to store all outputs.",
    )
    parser.add_argument(
        "--n_samples",
        type=int,
        default=0,
        help="Number of sequences to sample. Set to 0 to disable.",
    )

    args = parser.parse_args()

    output_dir = Path(args.output_dir)
    df_dir = output_dir / "dataframes"
    tmp_dir = output_dir / "tmp"

    output_dir.mkdir(exist_ok=True)
    df_dir.mkdir(exist_ok=True)
    tmp_dir.mkdir(exist_ok=True)

    # Use shared function to load and preprocess data
    df_main = load_preprocess_data(args.input_csv, args.n_samples)

    # Create query json files
    queries = []
    for id, sequence in zip(df_main["uniprot_id"], df_main["sequence"]):
        query = {"queries": {id: {"chains": [{"molecule_type": "protein", "chain_ids": ["A"], "sequence": sequence}]}}}
        query_path = tmp_dir / f"{id}_query.json"
        with open(query_path, "w") as f:
            json.dump(query, f)

        queries.append(query_path)

    print(f"Running OpenFold predictions for {len(queries)} sequences...")

    results = []
    for query_path in tqdm(queries, "Computing Structures"):
        print("df_dir", df_dir)
        results.append(run_openfold_prediction(df_dir, query_path=query_path))

    print("Predictions complete. Processing results...")
    df_out = pd.DataFrame(results)

    # Calculate simple stats for reporting
    status_counts = df_out["status"].value_counts().to_dict()
    print(f"Prediction Summary: {status_counts}")

    output_path = df_dir / f"openfold_status_{args.input_csv.split('/')[-1].split('.')[0]}.parquet"
    df_out.to_parquet(output_path, index=False)
    print(f"Saved prediction status logs to {output_path}")
```
