## Supplementary material for "pLM representations unlock metagenomic space beyond homology": Code bundle: create_pairwise_matrix.py

```
import pandas as pd
import numpy as np


def generate_matrix(tsv_paths, ids_path, output_path):
    print("Loading ID list...")
    target_ids = pd.read_csv(ids_path, header=None)[0].unique()
    target_ids.sort()

    id_to_idx = {id_: i for i, id_ in enumerate(target_ids)}
    valid_set = set(target_ids)
    n = len(target_ids)
    print(f"Processing {n} target IDs.")

    # 2. Read TSVs in chunks to avoid OOM
    # Lists to store coordinate data
    rows, cols, values = [], [], []
    chunk_size = 1_000_000

    for tsv_file in tsv_paths:
        print(f"Reading {tsv_file} and filtering...")

        # Assuming no header in TSV, columns: Query, Target, Value
        reader = pd.read_csv(tsv_file, sep="\t", chunksize=chunk_size, header=None, names=["q", "t", "v"])

        for i, chunk in enumerate(reader):
            if i % 10 == 0:
                print(f"Processed {i * chunk_size} rows in current file...", end="\r")

            # Boolean mask: Keep row only if BOTH ids are in our list
            mask = chunk["q"].isin(valid_set) & chunk["t"].isin(valid_set)

            if not mask.any():
                continue

            filtered = chunk.loc[mask]

            # Map string IDs to integer indices [0, n]
            r = filtered["q"].map(id_to_idx).values
            c = filtered["t"].map(id_to_idx).values
            v = filtered["v"].values

            rows.append(r)
            cols.append(c)
            values.append(v)
        print()  # Newline for clean output

    # 3. Concatenate chunks
    print("\nConsolidating data...")
    if not values:
        print("Warning: No matching interactions found.")
        return

    all_rows = np.concatenate(rows)
    all_cols = np.concatenate(cols)
    all_vals = np.concatenate(values)

    # 4. Deduplicate
    # If (ID1, ID2) appears multiple times, we take the MAX value.
    # We use a temporary DF for fast groupby aggregation.
    temp_df = pd.DataFrame({"r": all_rows, "c": all_cols, "v": all_vals})
    temp_df = temp_df.groupby(["r", "c"])["v"].max().reset_index()

    # 5. Build Sparse Matrix -> Dense DataFrame
    # Using float32 to halve memory usage (44k^2 * 4 bytes ~= 7.7 GB)
    print("Constructing dense matrix...")
    matrix_sparse = coo_matrix((temp_df["v"].astype(np.float32), (temp_df["r"], temp_df["c"])), shape=(n, n))

    # Convert to CSR then Dense for speed
    dense_arr = matrix_sparse.tocsr().toarray()

    df_out = pd.DataFrame(dense_arr, index=target_ids, columns=target_ids)

    # 6. Save
    print("Saving to Parquet...")
    df_out.to_parquet(output_path)
    print("Done.")


if __name__ == "__main__":
    # Update filenames as needed
    # Pass a list of input files now
    generate_matrix(["data_part1.tsv", "data_part2.tsv"], "ids.txt", "output.parquet")
```
