## Supplementary material for "pLM representations unlock metagenomic space beyond homology": Code bundle: embed_known_hydroxylases.py

```
"""
Parses ids and sequences from the known hydroxylases xlsx, writes them to a
CSV, and computes per-sequence pLM embeddings stored as a parquet file.
"""

import argparse
from functools import partial
from pathlib import Path

import pandas as pd
import torch
import torch.nn.functional as F
from torch.nn.utils.rnn import pad_sequence
from torch.utils.data import DataLoader
from tqdm import tqdm
from transformers import AutoModel, AutoTokenizer


def parse_xlsx(xlsx_path: Path) -> pd.DataFrame:
    """Find the id and sequence columns across sheets and return a clean DataFrame."""
    sheets = pd.read_excel(xlsx_path, sheet_name=None)

    id_candidates = {"id", "uniprot_id", "uniprot", "accession", "name", "protein_id", "entry"}
    seq_candidates = {"sequence", "seq", "protein_sequence", "aa_sequence"}

    def norm(s: str) -> str:
        return str(s).strip().lower().replace(" ", "_")

    frames = []
    for sheet_name, df in sheets.items():
        cols_lower = {c: norm(c) for c in df.columns}
        id_col = next((c for c, lc in cols_lower.items() if lc in id_candidates), None)
        seq_col = next((c for c, lc in cols_lower.items() if lc in seq_candidates), None)
        if id_col is None or seq_col is None:
            print(f"[skip] sheet {sheet_name!r}: no id/sequence columns (found {list(df.columns)})")
            continue
        sub = df[[id_col, seq_col]].rename(columns={id_col: "id", seq_col: "sequence"})
        sub["sheet"] = sheet_name
        frames.append(sub)

    if not frames:
        raise ValueError(f"No sheet in {xlsx_path} had recognizable id/sequence columns.")

    out = pd.concat(frames, ignore_index=True)
    out = out.dropna(subset=["id", "sequence"])
    out["id"] = out["id"].astype(str).str.strip()
    out["sequence"] = out["sequence"].astype(str).str.strip().str.replace(r"\s+", "", regex=True).str.upper()
    out = out[out["sequence"].str.len() > 0]
    out = out.drop_duplicates(subset=["id"]).reset_index(drop=True)
    return out


def collate_fn(batch, tokenizer):
    inputs = pad_sequence(batch, batch_first=True, padding_value=tokenizer.pad_token_id)
    pad = (8 - inputs.size(1) % 8) % 8
    if pad > 0:
        inputs = F.pad(inputs, (0, pad), value=tokenizer.pad_token_id)
    return inputs, (inputs != tokenizer.pad_token_id)


if __name__ == "__main__":
    parser = argparse.ArgumentParser(description=__doc__)
    parser.add_argument("--xlsx", type=str, default="data/known hydroxylases with characterized activity.xlsx")
    parser.add_argument("--output_csv", type=str, default="data/known_hydroxylases.csv")
    parser.add_argument("--output_dir", type=str, default="output")
    parser.add_argument("--data_name", type=str, default="known_hydroxylases")
    parser.add_argument("--model_name", type=str, default="AMPLIFY_120M")
    parser.add_argument("--model_path", type=str, default="chandar-lab/AMPLIFY_120M")
    parser.add_argument("--mask_type", type=str, default="additive", choices=["additive", "boolean"])
    parser.add_argument("--batch_size", type=int, default=32)
    parser.add_argument("--max_len", type=int, default=1024)
    parser.add_argument("--device", type=str, default="cuda:0" if torch.cuda.is_available() else "cpu")
    args = parser.parse_args()

    xlsx_path = Path(args.xlsx)
    output_csv = Path(args.output_csv)
    output_dir = Path(args.output_dir)
    df_dir = output_dir / "dataframes"
    output_csv.parent.mkdir(parents=True, exist_ok=True)
    df_dir.mkdir(parents=True, exist_ok=True)

    print(f"Parsing {xlsx_path}...")
    df = parse_xlsx(xlsx_path)
    print(f"Parsed {len(df)} unique sequences.")
    df.to_csv(output_csv, index=False)
    print(f"Wrote {output_csv}")

    print(f"Loading {args.model_name} ({args.model_path}) on {args.device}...")
    model = AutoModel.from_pretrained(args.model_path, trust_remote_code=True).to(args.device).eval()
    tokenizer = AutoTokenizer.from_pretrained(args.model_path, trust_remote_code=True)

    tokenized = [
        tokenizer.encode(s, max_length=args.max_len, truncation=True, return_tensors="pt").squeeze(0)
        for s in tqdm(df.sequence, desc="Tokenizing")
    ]
    dataloader = DataLoader(tokenized, batch_size=args.batch_size, collate_fn=partial(collate_fn, tokenizer=tokenizer))

    pooled_embeddings = []
    use_amp = args.device.startswith("cuda")
    autocast_ctx = torch.autocast(device_type="cuda", dtype=torch.float16) if use_amp else torch.autocast(device_type="cpu", enabled=False)

    with torch.no_grad(), autocast_ctx:
        for ids, mask in tqdm(dataloader, desc="Embedding"):
            ids = ids.to(args.device)
            mask = mask.to(args.device)
            if args.mask_type == "additive":
                attn = torch.where(mask, 0.0, float("-inf")).to(device=args.device, dtype=torch.float16 if use_amp else torch.float32)
            else:
                attn = mask.to(device=args.device, dtype=torch.int)

            out = model(ids, attention_mask=attn, output_hidden_states=True)
            hidden = out.hidden_states[-1]
            pooled = (hidden * mask.unsqueeze(-1)).sum(1) / torch.clamp(mask.sum(1, keepdim=True), min=1)
            pooled_embeddings.extend(pooled.float().cpu().numpy())

    df_out = df[["id", "sequence"]].copy()
    df_out[f"{args.model_name}_embedding"] = [e.tolist() for e in pooled_embeddings]

    output_path = df_dir / f"plm_embeddings_{args.data_name}_{args.model_name}.parquet"
    df_out.to_parquet(output_path, index=False)
    print(f"Saved embeddings to {output_path}")
```
