## Supplementary material for "pLM representations unlock metagenomic space beyond homology": Code bundle: filter_dataframes.py

```
import os
import argparse
from pathlib import Path
from functools import reduce

import numpy as np
import pandas as pd

import pyarrow.parquet as pq
import pyarrow.dataset as ds


def merge_dataframes(dataframes, key_col):
    """
    Merges a list of DataFrames on a common key column, handling
    duplicate metadata columns.
    """
    # Load the first dataframe completely
    base_df = dataframes[0]

    # Iteratively merge the rest
    if len(dataframes) > 1:
        other_dfs = []
        base_cols = set(base_df.columns)

        for df in dataframes[1:]:
            # Find unique columns in this df
            unique_cols = [col for col in df.columns if col not in base_cols]

            # Add to base_cols so they aren't re-added from another file
            base_cols.update(unique_cols)

            # Only load key + unique columns
            cols_to_load = [key_col] + unique_cols

            # Ensure we don't try to load columns that don't exist
            valid_cols_to_load = [col for col in cols_to_load if col in df.columns]

            other_dfs.append(df[valid_cols_to_load])

        # Now merge all the small dataframes into the base
        final_df = reduce(lambda left, right: pd.merge(left, right, on=key_col, how="outer"), [base_df] + other_dfs)
        return final_df

    return base_df


if __name__ == "__main__":
    parser = argparse.ArgumentParser(description="Downloads PDBs and scores them with ESM-IF.")
    parser.add_argument(
        "--input_dir",
        type=str,
        default="input",
        help="Main directory where data is stored.",
    )
    parser.add_argument(
        "--output_dir",
        type=str,
        default="filtered_output",
        help="Main directory where outputs are stored.",
    )
    parser.add_argument(
        "--csv_file",
        type=str,
        default=None,
    )
    parser.add_argument(
        "--pairwise_similarity_file",
        type=str,
        default=None,
    )
    parser.add_argument(
        "--pairwise_identity_file",
        type=str,
        default=None,
    )
    parser.add_argument(
        "--pairwise_blast_file",
        type=str,
        default=None,
    )
    args = parser.parse_args()

    # Define the directory where your results are stored
    input_dir = Path(args.input_dir)
    DATA_DIR = input_dir / "dataframes"

    output_dir = Path(args.output_dir)
    output_dir.mkdir(exist_ok=True)

    # Define the primary key for merging
    KEY_COL = "uniprot_id"

    # Add all your generated metrics filenames to this list.
    FILES_TO_MERGE = [
        "physicochem_properties.parquet",
        "structural_properties_old.parquet",
        "signalp_scores.parquet",
        "plm_embeddings_AMPLIFY_350M.parquet",
        "plm_embeddings_AMPLIFY-C.parquet",
        "plm_perplexity_AMPLIFY-C.parquet",
        "proteinmpnn_scores_soluble.parquet",
        "esm_if_scores.parquet",
    ]

    print("Loading metrics files...")
    all_dfs = [pd.read_parquet(DATA_DIR / file) for file in FILES_TO_MERGE]

    print(f"Merging {len(all_dfs)} dataframes on key '{KEY_COL}'...")
    df_main = merge_dataframes(all_dfs, KEY_COL)

    filters = {
        "length": (200, 450),
        "sp_likelihood": (0.0, 0.05),
        "HXD/E_motif": True,
    }

    cut_lowest_5_properties = [
        "mean_plddt",
        # "esm_if_nll",
        # "aromaticity",
    ]

    cut_highest_5_properties = [
        "shannon_entropy_normalized",
        # "mpnn_ll_soluble",
    ]

    df_main_filtered = df_main.copy()
    df_main_filtered["HXD/E_motif"] = df_main_filtered["sequence"].str.contains("H.D|H.E", regex=True)

    for col in cut_lowest_5_properties:
        threshold = df_main_filtered[col].quantile(0.05)
        filters[col] = (threshold, df_main_filtered[col].max())

    for col in cut_highest_5_properties:
        threshold = df_main_filtered[col].quantile(0.95)
        filters[col] = (df_main_filtered[col].min(), threshold)

    for col, filter in filters.items():
        if col in df_main_filtered.columns:
            if isinstance(filter, bool):
                if filter:
                    df_main_filtered = df_main_filtered[df_main_filtered[col] == True]
                    print(f"Filtered {col} for True values. New size: {len(df_main_filtered)}")
                else:
                    df_main_filtered = df_main_filtered[df_main_filtered[col] == False]
                    print(f"Filtered {col} for False values. New size: {len(df_main_filtered)}")
                print(f"Filtered {col} for {filter} values. New size: {len(df_main_filtered)}")
            else:
                min_val, max_val = filter
                df_main_filtered = df_main_filtered[(df_main_filtered[col] >= min_val) & (df_main_filtered[col] <= max_val)]
                print(f"Filtered {col} to range [{min_val}, {max_val}]. New size: {len(df_main_filtered)}")
        else:
            print(f"Warning: Column '{col}' not found in dataframe. Skipping this filter.")

    # Get unique index for this filtering step
    existing_metadata_files = list(output_dir.glob("filtering_metadata_*.txt"))
    existing_indices = [int(f.stem.split("_")[-1]) for f in existing_metadata_files if f.stem.split("_")[-1].isdigit()]
    new_idx = max(existing_indices) + 1 if existing_indices else 0

    df_main_filtered.to_parquet(output_dir / f"merged_data_{new_idx:03}.parquet", index=False)
    metadata = {
        "total_sequences_before_filtering": len(df_main),
        "total_sequences_after_filtering": len(df_main_filtered),
        "filters_applied": filters,
    }
    with open(output_dir / f"filtering_metadata_{new_idx:03}.txt", "w") as f:
        for key, value in metadata.items():
            f.write(f"{key}: {value}\n")

    df_main_filtered_ids = set(df_main_filtered["uniprot_id"])

    if args.csv_file is not None:
        raw_df = pd.read_csv(args.csv_file)
        raw_df["uniprot_id"] = raw_df["name"].apply(lambda x: x.split("|")[1] if "|" in x else x)
        raw_df_filtered = raw_df[raw_df["uniprot_id"].isin(df_main_filtered_ids)]
        csv_dir = Path(os.path.dirname(args.csv_file))
        raw_df_filtered.to_csv(csv_dir / f"sequences_filtered_{new_idx:03}.csv", index=False)

    if args.pairwise_similarity_file is not None:
        print("Filtering pairwise similarity dataframe...")
        df = pd.read_parquet(args.pairwise_similarity_file, columns=df_main_filtered_ids)
        df_filtered = df[(df.index.isin(df_main_filtered_ids))]
        df_filtered.to_parquet(output_dir / f"pairwise_similarity_{new_idx:03}.parquet", index=True, compression="snappy")

    if args.pairwise_identity_file is not None:
        print("Filtering pairwise identity dataframe...")
        dataset = ds.dataset(args.pairwise_identity_file, format="parquet")

        row_filter = ds.field("__index_level_0__").isin(list(df_main_filtered_ids))
        columns_to_read = list(df_main_filtered_ids) + ["__index_level_0__"]

        filtered_table = dataset.to_table(filter=row_filter, columns=columns_to_read)
        df = filtered_table.to_pandas()
        df.to_parquet(output_dir / f"pairwise_identity_{new_idx:03}.parquet", index=True)

    if args.pairwise_blast_file is not None:
        print("Filtering pairwise blast dataframe...")
        dataset = ds.dataset(args.pairwise_blast_file, format="parquet")

        row_filter = ds.field("__index_level_0__").isin(list(df_main_filtered_ids))
        columns_to_read = list(df_main_filtered_ids) + ["__index_level_0__"]

        filtered_table = dataset.to_table(filter=row_filter, columns=columns_to_read)
        df = filtered_table.to_pandas()
        df.to_parquet(output_dir / f"pairwise_blast_{new_idx:03}.parquet", index=True)
```
