## Supplementary material for "pLM representations unlock metagenomic space beyond homology": Code bundle: match_sequences.py

```
import pandas as pd
import pyarrow as pa
import pyarrow.csv as csv
import pyarrow.parquet as pq
import pyarrow.compute as pc
from pathlib import Path


def match_sequences(base_path, base_search_path, lambda_param, ds_names, csv_files):
    base_path = Path(base_path)

    for ds_name in ds_names:
        print(f"Processing: {ds_name}")

        search_path = base_search_path / f"{ds_name}/{lambda_param:.2f}/search_results.parquet"
        if not search_path.exists():
            print(f"Skipping {ds_name}, file not found.")
            continue

        search_table = pq.read_table(search_path)
        unique_ids = pc.unique(search_table.column("id"))
        filtered_tables = []

        for csv_filename in csv_files:
            print(f"Reading and filtering: {csv_filename}")
            csv_path = base_path / csv_filename

            try:
                seq_table = csv.read_csv(csv_path)
            except Exception as e:
                print(f"Error reading {csv_filename}: {e}")
                continue

            is_in_ids = pc.is_in(seq_table.column("name"), value_set=unique_ids)
            filtered_chunk = seq_table.filter(is_in_ids)

            if len(filtered_chunk) > 0:
                filtered_tables.append(filtered_chunk)

            del seq_table

        if filtered_tables:
            combined_seq_table = pa.concat_tables(filtered_tables)
            sequence_df_filtered = combined_seq_table.to_pandas()
            df = search_table.to_pandas()

            merged_df = df.merge(sequence_df_filtered, left_on="id", right_on="name", how="left")

            output_path = search_path.parent / "output.parquet"
            merged_df.to_parquet(output_path, engine="pyarrow")

            print(f"Saved to {output_path}")

            del df, sequence_df_filtered, merged_df, combined_seq_table
        else:
            print(f"No matching sequences found for {ds_name}")


if __name__ == "__main__":
    mgnify_base = "$SCRATCH/proseqo/datasets/mgnify/"
    mgnify_datasets = [
        "mgy_0_potential_enzymes_0.50",
        "mgy_1_potential_enzymes_0.50",
        "mgy_2_potential_enzymes_0.50",
        "mgy_3_potential_enzymes_0.50",
    ]
    mgnify_csvs = [
        "mgnify_0_potential_enzymes.csv",
        "mgnify_1_potential_enzymes.csv",
        "mgnify_2_potential_enzymes.csv",
        "mgnify_3_potential_enzymes.csv",
    ]

    base_search_path = Path("$SCRATCH/proseqo/enzymes/metagenomic_search")
    lambda_param = 0.90

    bfd_base = "$SCRATCH/proseqo/datasets/bfd/"
    bfd_datasets = [
        "bfd_potential_enzymes_0.50",
    ]
    bfd_csvs = [
        "bfd_potential_enzymes.csv",
    ]

    match_sequences(mgnify_base, base_search_path, lambda_param, mgnify_datasets, mgnify_csvs)
    match_sequences(bfd_base, base_search_path, lambda_param, bfd_datasets, bfd_csvs)
```
