## Supplementary material for "pLM representations unlock metagenomic space beyond homology": Code bundle: metagenomic_search.py

```
import numpy as np
import pandas as pd
import glob
import os
import argparse
import sys
from tqdm import tqdm


def normalize_vectors(vectors):
    """
    Normalizes vectors to unit length (L2 norm).
    """
    norm = np.linalg.norm(vectors, axis=1, keepdims=True)
    norm[norm == 0] = 1
    return vectors / norm


import os
import glob
import numpy as np
import pandas as pd
from tqdm import tqdm


def find_top_k_in_parquet(
    query_vectors, parquet_dir, k=5, vector_col="embedding", id_col="id", intra_score_col="mean_cosine_sim", file_pattern="*.parquet"
):
    """
    Iterates over parquet files to find top-k nearest neighbors.
    Returns IDs, Scores, and the actual Vectors of the neighbors.
    """

    # 1. Pre-process Query Vectors
    query_vectors = normalize_vectors(query_vectors)
    n_queries, dim = query_vectors.shape

    # Initialize global storage in a dictionary to avoid managing multiple variables
    # Keys match the data we want to track
    state = {
        "scores": np.full((n_queries, k), -1.0, dtype=np.float32),
        "ids": np.empty((n_queries, k), dtype=object),
        "vectors": np.zeros((n_queries, k, dim), dtype=np.float32),
        "intra_score": np.full((n_queries, k), -1.0, dtype=np.float32),
    }

    files = glob.glob(os.path.join(parquet_dir, file_pattern))
    if not files:
        print(f"No files found in {parquet_dir} matching {file_pattern}")
        return None, None, None, None

    for filepath in tqdm(files, total=len(files), desc="Searching k-nn in parquet files."):
        try:
            # Ensure we read all necessary columns
            df = pd.read_parquet(filepath, columns=[id_col, vector_col, intra_score_col])
        except Exception as e:
            print(f"Skipping {filepath}: {e}")
            continue

        if len(df) == 0:
            continue

        # Extract Raw Data
        db_vectors_raw = np.stack(df[vector_col].values)

        # 2. Compute Similarity
        db_vectors_norm = normalize_vectors(db_vectors_raw.astype(np.float32))
        batch_scores = np.dot(query_vectors, db_vectors_norm.T)

        # 3. Find 'Local' Top-K Candidates (Optimization)
        # If batch is larger than k, filter down to top-k locally to save memory before merging
        if batch_scores.shape[1] >= k:
            top_indices = np.argpartition(batch_scores, -k, axis=1)[:, -k:]
        else:
            top_indices = np.tile(np.arange(batch_scores.shape[1]), (n_queries, 1))

        row_idx = np.arange(n_queries)[:, None]

        # Organize local batch data into a matching dictionary structure
        local_data = {
            "scores": batch_scores[row_idx, top_indices],
            "ids": df[id_col].values[top_indices],
            "vectors": db_vectors_raw[top_indices],
            "intra_score": df[intra_score_col].values[top_indices],
        }

        # 4. Merge with Global Top-K
        # We determine the sort order based on scores, then apply it to all fields
        combined_scores = np.concatenate((state["scores"], local_data["scores"]), axis=1)

        # Determine indices of the top-k in the combined batch
        sorted_indices = np.argsort(-combined_scores, axis=1)[:, :k]

        # Update all state fields using the same indices
        for key in state:
            combined_field = np.concatenate((state[key], local_data[key]), axis=1)
            state[key] = combined_field[row_idx, sorted_indices]

    return state["ids"], state["scores"], state["vectors"], state["intra_score"]


def load_queries(path, data_path, vector_col, id_col=None):
    """
    Loads queries and returns (vectors, ids).
    If id_col is None or not found, generates numeric indices.
    """
    if path.endswith(".npy"):
        vectors = np.load(path)
        ids = np.arange(vectors.shape[0])

    elif path.endswith(".parquet"):
        cols_to_load = [vector_col]
        if id_col:
            cols_to_load.append(id_col)

        df = pd.read_parquet(path, columns=cols_to_load)
        vectors = np.stack(df[vector_col].values)

        if id_col and id_col in df.columns:
            ids = df[id_col].values
        else:
            ids = np.arange(len(df))

    elif path.endswith(".txt"):
        with open(path, "r") as f:
            ids = list(set(line.strip() for line in f if line.strip()))

        df = pd.read_parquet(data_path)
        df = df[df[id_col].isin(ids)]

        vectors = np.stack(df[vector_col].values)

    else:
        raise ValueError("Query file must be .npy or .parquet")

    return vectors, ids


if __name__ == "__main__":
    parser = argparse.ArgumentParser(description="Parquet KNN Search")
    parser.add_argument("--queries_id", required=True, help="Path to query ids (.txt)")
    parser.add_argument("--data_path", required=True, help="Path to full uniprot dataframe (.parquet)")
    parser.add_argument("--parquet_dir", required=True, help="Directory containing DB parquet files")
    parser.add_argument("--output_dir", required=True, help="Directory to save results")
    parser.add_argument("--k", type=int, default=5, help="Number of neighbors")
    parser.add_argument("--vector_col", default="embeddings", help="Vector column name")
    parser.add_argument("--id_col", default="uniprot_id", help="DB ID column name")
    parser.add_argument("--query_vector_col", default="AMPLIFY_embedding", help="Query vector column name")
    parser.add_argument("--query_id_col", default=None, help="Query ID column name (optional)")
    parser.add_argument("--file_pattern", default="*.parquet", help="File pattern for DB files")

    args = parser.parse_args()

    # Load Queries
    try:
        queries, query_ids = load_queries(args.queries_id, args.data_path, args.query_vector_col, args.query_id_col)
        print(f"Loaded {len(queries)} queries.")
    except Exception as e:
        sys.exit(f"Error loading queries: {e}")

    # Create output dir
    os.makedirs(args.output_dir, exist_ok=True)

    # Run Search
    ids, scores, vectors, intra_scores = find_top_k_in_parquet(
        queries, args.parquet_dir, k=args.k, vector_col=args.vector_col, id_col=args.id_col, file_pattern=args.file_pattern
    )

    if ids is not None:
        # Save Results
        print("Saving results...")

        # Prepare data for DataFrame (Long Format)
        n_queries, k_neighbors = ids.shape
        dim = vectors.shape[2]

        # 1. Repeat Query IDs to match flattened results: [q1, q1, q1, q2, q2, q2...]
        flat_query_ids = np.repeat(query_ids, k_neighbors, axis=0)

        # 2. Flatten the retrieved results: [match1, match2, match3...]
        flat_retrieved_ids = ids.flatten()
        flat_scores = scores.flatten()
        flat_intra_scores = intra_scores.flatten()

        # 3. Flatten vectors into a list of arrays (N*K, Dim) -> List of (Dim,) arrays
        # This structure is compatible with Parquet and maintains the vector data
        flat_vectors = list(vectors.reshape(-1, dim))

        # Create DataFrame
        df_results = pd.DataFrame(
            {
                "query_id": flat_query_ids,
                "id": flat_retrieved_ids,
                "inter_score": flat_scores,
                "embedding": flat_vectors,
                "intra_score": flat_intra_scores,
            }
        )

        # Save to Parquet
        output_file = os.path.join(args.output_dir, "search_results.parquet")
        df_results.to_parquet(output_file, index=False)

        print(f"Done. Saved results to {output_file}")
    else:
        sys.exit("Search failed or no files found.")
```
