## Supplementary material for "pLM representations unlock metagenomic space beyond homology": Code bundle: metagenomic_select.py

```
import numpy as np
import pandas as pd
import os


tmp_list = []
for ds in ["mgy_0_potential_enzymes_0.50", "mgy_1_potential_enzymes_0.50", "mgy_2_potential_enzymes_0.50", "mgy_3_potential_enzymes_0.50"]:
    df = pd.read_parquet(f"$SCRATCH/proseqo/enzymes/metagenomic_search/{ds}/0.10/output.parquet")
    df["lambda"] = 0.10
    df["dataset"] = "mgnify"
    tmp_list.append(df)

    df = pd.read_parquet(f"$SCRATCH/proseqo/enzymes/metagenomic_search/{ds}/0.90/output.parquet")
    df["lambda"] = 0.90
    df["dataset"] = "mgnify"
    tmp_list.append(df)

df = pd.read_parquet("$SCRATCH/proseqo/enzymes/metagenomic_search/bfd_potential_enzymes_0.50/0.10/output.parquet")
df["lambda"] = 0.10
df["dataset"] = "bfd"
tmp_list.append(df)

df = pd.read_parquet("$SCRATCH/proseqo/enzymes/metagenomic_search/bfd_potential_enzymes_0.50/0.90/output.parquet")
df["lambda"] = 0.90
df["dataset"] = "bfd"
tmp_list.append(df)

df = pd.concat(tmp_list)
df = df.sort_values("inter_score", ascending=False)
df = df.drop_duplicates(subset="id").reset_index(drop=True)
df["length"] = df["sequence"].apply(len)

# Save to csv
seq = df[["name", "sequence"]]
seq = seq.drop_duplicates()
seq.to_csv("$HOME/enzymes/data/metagenomic_candidates.csv", index=False)

# Save to fasta
fasta_path = f"$HOME/enzymes/data/metagenomic_candidates.fasta"
with open(fasta_path, "w") as f:
    for id, sequence in zip(seq["name"], seq["sequence"]):
        f.write(f">{id}\n")
        f.write(f"{sequence}\n")

ds = pd.read_csv(
    "$SCRATCH/proseqo/enzymes/metagenomic_search/metagenomic_candidates_blast.tsv", sep="\t", header=None
)
ds.columns = ["metagenomic_id", "uniprot_id", "blast"]
ds = ds.sort_values("blast", ascending=False)
ds = ds.drop_duplicates(subset="metagenomic_id", keep="first")
ds = ds.reset_index(drop=True)
df = df.merge(ds, left_on="id", right_on="metagenomic_id", how="left")

uniprot = pd.read_csv("$HOME/enzymes/data/uniprotkb_ec_1_14_11_2025_10_20.csv")
uniprot["uniprot_id"] = uniprot["name"].apply(lambda x: x.split("|")[1] if len(x.split("|")) > 1 else x)
uniprot.drop(columns="name", inplace=True)
df = df.merge(uniprot, on="uniprot_id", how="left")
df.rename(columns={"sequence_x": "sequence_metagenomic", "sequence_y": "sequence_uniprot"}, inplace=True)
df["sequence_uniprot"] = np.where(df["sequence_uniprot"].isna(), None, df["sequence_uniprot"])
df["uniprot_length"] = df["sequence_uniprot"].apply(lambda x: len(x) if x is not None else -1)
df["length"] = df["sequence_metagenomic"].apply(lambda x: len(x) if x is not None else -1)

df.rename(
    columns={
        "uniprot_id": "closest_blast_uniprot_id",
        "sequence_uniprot": "closest_blast_sequence_uniprot",
        "uniprot_length": "closest_blast_uniprot_length",
        "query_id": "closest_embedding_uniprot_id",
    },
    inplace=True,
)
df.drop(columns=["id", "name"], inplace=True)
df.set_index("metagenomic_id", inplace=True)
```
