## Supplementary material for "pLM representations unlock metagenomic space beyond homology": Code bundle: mmseqs.py

```
"""
Computes the maximum sequence similarity of every sequence in a query set
to a target set using MMseqs2.

Inputs: two FASTA files (plain or gzipped). One sequence ID per record (>id).
Outputs a CSV with columns: query_id, max_pident (0-100 scale).
"""

import argparse
import gzip
import subprocess
from pathlib import Path

import pandas as pd


def parse_fasta(path: Path) -> dict[str, str]:
    """Parse a plain or gzipped FASTA file into {id: sequence}."""
    opener = gzip.open if str(path).endswith(".gz") else open
    sequences = {}
    current_id = None
    current_seq = []

    with opener(path, "rt") as f:
        for line in f:
            line = line.strip()
            if line.startswith(">"):
                if current_id is not None:
                    sequences[current_id] = "".join(current_seq)
                current_id = line[1:].split()[0]  # use first token as ID
                current_seq = []
            else:
                current_seq.append(line)

    if current_id is not None:
        sequences[current_id] = "".join(current_seq)

    return sequences


if __name__ == "__main__":

    parser = argparse.ArgumentParser(
        description="Get max sequence similarity of every query to a target set using MMseqs."
    )
    parser.add_argument("--query_fasta", type=str, required=True, help="Query FASTA file (plain or .gz).")
    parser.add_argument("--target_fasta", type=str, required=True, help="Target FASTA file (plain or .gz).")
    parser.add_argument(
        "--output_csv",
        type=str,
        default="output/max_similarity.csv",
        help="Path for the output CSV.",
    )
    parser.add_argument(
        "--tmp_dir",
        type=str,
        default="output/mmseqs_tmp_sim",
        help="Temporary directory for MMseqs intermediate files.",
    )
    parser.add_argument(
        "--m8_dir",
        type=str,
        default="output/mmseqs_m8",
        help="Directory for MMseqs m8 files to keep.",
    )
    parser.add_argument("--cores", type=int, default=8, help="Number of CPU threads for MMseqs.")
    parser.add_argument(
        "--mmseqs_path",
        type=str,
        default="data/mmseqs/bin/mmseqs",
        help="Path to the mmseqs binary.",
    )
    parser.add_argument(
        "--sensitivity",
        type=float,
        default=7.0,
        help="MMseqs search sensitivity (1-7.5, higher = more sensitive but slower).",
    )
    args = parser.parse_args()

    mmseqs_bin = Path(args.mmseqs_path)
    if not mmseqs_bin.is_file():
        print(f"Error: MMseqs binary not found at: {mmseqs_bin}")
        exit(1)

    tmp_dir = Path(args.tmp_dir)
    tmp_dir.mkdir(parents=True, exist_ok=True)
    m8_dir = Path(args.m8_dir)
    m8_dir.mkdir(parents=True, exist_ok=True)
    Path(args.output_csv).parent.mkdir(parents=True, exist_ok=True)

    # --- Parse input FASTAs ---
    print("Parsing query FASTA...")
    query_seqs = parse_fasta(Path(args.query_fasta))
    print("Parsing target FASTA...")
    target_seqs = parse_fasta(Path(args.target_fasta))
    print(f"Query: {len(query_seqs)} sequences | Target: {len(target_seqs)} sequences")

    # If input is gzipped, write decompressed copies for MMseqs
    query_fasta_plain = tmp_dir / "query.fasta"
    target_fasta_plain = tmp_dir / "target.fasta"

    for seqs, out_path in [(query_seqs, query_fasta_plain), (target_seqs, target_fasta_plain)]:
        with open(out_path, "w") as f:
            for seq_id, seq in seqs.items():
                f.write(f">{seq_id}\n{seq}\n")

    # --- File paths ---
    query_db = tmp_dir / "query_db"
    target_db = tmp_dir / "target_db"
    search_res = tmp_dir / "search_res"
    results_m8 = m8_dir / f"{Path(args.output_csv).stem}.m8"

    # --- Build MMseqs databases ---
    print("Creating MMseqs databases...")
    subprocess.run([str(mmseqs_bin), "createdb", str(query_fasta_plain), str(query_db)], check=True, capture_output=True)
    subprocess.run([str(mmseqs_bin), "createdb", str(target_fasta_plain), str(target_db)], check=True, capture_output=True)

    # --- Search ---
    print("Running query-vs-target search...")
    subprocess.run(
        [
            str(mmseqs_bin), "search",
            str(query_db), str(target_db),
            str(search_res), str(tmp_dir),
            "-s", str(args.sensitivity),
            "--threads", str(args.cores),
            "-c", "0.8", "--cov-mode", "0",
        ],
        check=True,
        capture_output=True,
    )

    # --- Convert to tabular format ---
    print("Converting results to m8 format...")
    subprocess.run(
        [
            str(mmseqs_bin), "convertalis",
            str(query_db), str(target_db),
            str(search_res), str(results_m8),
            "--format-output", "query,target,pident",
            "--threads", str(args.cores),
        ],
        check=True,
        capture_output=True,
    )

    # --- Compute max similarity per query ---
    print("Computing max similarity per query...")
    df_results = pd.read_csv(results_m8, sep="\t", header=None, names=["query_id", "target_id", "pident"])

    max_sim = df_results.groupby("query_id")["pident"].max().reset_index()
    max_sim.columns = ["query_id", "max_pident"]

    # Queries with no hits get 0
    max_sim = (
        pd.DataFrame({"query_id": list(query_seqs.keys())})
        .merge(max_sim, on="query_id", how="left")
        .fillna(0.0)
    )

    max_sim.to_csv(args.output_csv, index=False)
    print(f"Saved max similarities to {args.output_csv}")
    print(max_sim.describe())
```
