## Supplementary material for "pLM representations unlock metagenomic space beyond homology": Code bundle: utils.py

```
"""
Utility functions for the protein engineering pipeline.
Includes data loading/preprocessing and PDB downloading.
"""

from pathlib import Path
import pandas as pd
import requests

import glob
import shutil
import csv
import os
import re
import numpy as np
from tqdm import tqdm
import torch
import logging
import pandas as pd
import polars as pl

from typing import Dict, Any

from accelerate import Accelerator
from accelerate.utils import ProjectConfiguration

from torch.utils.data import DataLoader

# HuggingFace
from datasets import load_dataset
from transformers import PreTrainedTokenizerFast, DefaultDataCollator


def create_masked_dataset(input_file, output_dir, max_len: int = None):
    """
    Preprocesses a dataset for computing the pseudo-perplexity. It creates a csv file with replicated sequences with a mask at every position.
    """
    if max_len is not None:
        output_file = os.path.basename(input_file).replace(".csv", f"_max_{max_len}_masked.csv")
    else:
        output_file = os.path.basename(input_file).replace(".csv", f"_masked.csv")

    os.makedirs(output_dir, exist_ok=True)
    output_file = os.path.join(output_dir, output_file)

    if os.path.isfile(output_file):
        print("Masked file already exists, skipping preprocessing.")
        return output_file

    with open(input_file, "r", newline="") as fin, open(output_file, "w", newline="") as fout:
        reader = csv.DictReader(fin)
        writer = csv.writer(fout, lineterminator="\n")
        writer.writerow(["sequence", "id", "masked_index", "target"])

        for row in tqdm(reader, desc="Preprocessing sequences"):
            seq = row["sequence"]
            try:
                seq_id = row["name"]
            except KeyError:
                seq_id = row["id"]
            if max_len is not None:
                seq = seq[:max_len]
            for i in range(len(seq)):
                target = seq[i]
                masked_seq = seq[:i] + "<mask>" + seq[i + 1 :]
                writer.writerow([masked_seq, seq_id, i, target])
    return output_file


class DataCollatorWithPacking(DefaultDataCollator):
    def __init__(self, **kwargs):
        super().__init__(**kwargs)

    def __call__(self, batch):
        # Add position_ids if not present
        if "position_ids" not in batch[0]:
            for item in batch:
                item["position_ids"] = list(range(len(item["input_ids"])))

        # Pack the sequences into a single list
        input_ids_list = [item["input_ids"] for item in batch]
        position_ids_list = [item["position_ids"] for item in batch]
        seqlens = np.array([0] + [len(ids) for ids in input_ids_list])

        packed_batch = {
            "position_ids": np.concatenate(position_ids_list, axis=0),
            "input_ids": np.concatenate(input_ids_list, axis=0),
            "cu_seqlens": np.cumsum(seqlens),
            "max_seqlen": max(seqlens),
        }

        output_batch = super().__call__([packed_batch])
        output_batch["cu_seqlens"] = output_batch["cu_seqlens"].to(torch.int32).squeeze()
        output_batch["masked_index"] = torch.tensor([x["masked_index"] for x in batch], dtype=torch.int32)
        if "id" in batch[0].keys():
            output_batch["id"] = [x["id"] for x in batch]
        if "target" in batch[0].keys():
            output_batch["target"] = [x["target"] for x in batch]

        return output_batch


class PseudoPPLEvaluator:
    """
    Evaluator for pseudo-perplexity evaluation.
    """

    def __init__(
        self,
        packing: bool = True,
        batch_size: int = 128,
        save_steps: int = 1000,
        num_workers: int = 4,
    ):
        self.logger = logging.getLogger(__name__)
        logging.basicConfig(filename=None, level=logging.INFO)

        super().__init__()

        self._name = self.name
        self.packing = packing
        self.batch_size = batch_size
        self.save_steps = save_steps
        self.num_workers = num_workers

    @property
    def name(cls):
        return "pseudo_ppl"

    def _compute_metric(self, model_name: str, output_dir: str):
        parquet_files = [f for f in os.listdir(output_dir) if f.endswith(".parquet")]

        expected_schema = {
            "log_prob": pl.Float32,
            "id": pl.Utf8,
            "masked_index": pl.Int32,
        }

        print(f"Loading {len(parquet_files)} files from {output_dir}")
        df = pl.concat([pl.read_parquet(os.path.join(output_dir, f)).cast(expected_schema) for f in parquet_files])

        ppl = df.group_by("id").mean()

        ppl = ppl.with_columns(
            pl.col("id").map_elements(lambda x: x.split("|")[1] if len(x.split("|")) > 1 else x, return_dtype=pl.Utf8).alias("uniprot_id"),
            ((-pl.col("log_prob")).exp()).alias(f"{model_name}_pseudo_ppl"),
        )

        ppl = ppl.select(["uniprot_id", f"{model_name}_pseudo_ppl"]).to_pandas()

        return ppl

    def __call__(self, model, tokenizer, input_csv: str, max_len: int, model_name: str, data_name: str, output_dir: str) -> Dict[str, Any]:
        """
        Evaluate the pseudo-perplexity.

        Args:
            model: Encoder model to evaluate.

        Returns:
            Dictionary containing the evaluation metrics.
        """

        # Define paths
        output_path = output_dir #os.path.join(output_dir, self.name, model_name, data_name)
        os.makedirs(output_path, exist_ok=True)

        project_config = ProjectConfiguration(
            project_dir=output_path,
            automatic_checkpoint_naming=True,
            total_limit=3,
        )
        accelerator = Accelerator(
            project_config=project_config,
            mixed_precision="fp16",
        )

        with accelerator.main_process_first():
            masked_file = create_masked_dataset(input_file=input_csv, output_dir=output_dir, max_len=max_len)
            print(masked_file)
            dataset = load_dataset("csv", data_files=masked_file)
            
            # Test if dataset is datasetdict
            if isinstance(dataset, dict):
                dataset = dataset["train"]

        # Put model in evaluation mode
        self.logger.info(f"Loading model {model_name}")
        model.eval()

        if self.packing:
            collate_fn = DataCollatorWithPacking()
            dataset.set_transform(
                lambda x: tokenizer(x["sequence"], truncation=False, padding=False, return_tensors=None)
                | {"id": x["id"]}
                | {"masked_index": x["masked_index"]}
                | {"target": tokenizer.convert_tokens_to_ids(x["target"])}
            )
        else:
            dataset.set_transform(
                lambda x: tokenizer(x["sequence"], truncation=False, padding=True, return_tensors="pt", pad_to_multiple_of=8)
                | {"id": x["id"]}
                | {"masked_index": x["masked_index"]}
                | {"target": tokenizer.convert_tokens_to_ids(x["target"])}
            )

        # Dataloader
        dataloader = DataLoader(
            dataset,
            batch_size=self.batch_size,
            collate_fn=collate_fn if self.packing else None,
            shuffle=False,
            prefetch_factor=2,
            num_workers=self.num_workers,
            pin_memory=True,
            persistent_workers=True,
        )

        model, dataloader = accelerator.prepare(model, dataloader)

        full_checkpoint_dir = os.path.join(output_path, "checkpoints")
        if os.path.exists(full_checkpoint_dir):
            # This regular expression was taken from accelerator.load_state()
            it = max(int(re.findall(r"[\/]?([0-9]+)(?=[^\/]*$)", folder)[0]) for folder in os.listdir(full_checkpoint_dir))
            # Remove empty checkpoint folders
            while len(os.listdir(os.path.join(full_checkpoint_dir, f"checkpoint_{it}"))) == 0:
                shutil.rmtree(os.path.join(full_checkpoint_dir, f"checkpoint_{it}"), ignore_errors=True)
                it -= 1

            accelerator.load_state()

            files = glob.glob(os.path.join(output_path, "*_*.parquet"))
            start_batch = max([int(f.split("_")[-1].replace(".parquet", "")) for f in files] + [0])

            print(f"Resuming from batch {start_batch}")
            dataloader = accelerator.skip_first_batches(dataloader, start_batch)
        else:
            start_batch = 0

        # Temporary dict to store data in between dumps
        results = {
            "log_prob": [],
            "id": [],
            "masked_index": [],
        }

        torch.backends.cuda.matmul.allow_tf32 = True
        torch.backends.cudnn.allow_tf32 = True

        # Process bar
        total_batches = len(dataloader)
        pbar = tqdm(
            desc=f"Evaluating metric {self.name} on model {model_name}",
            unit="step",
            initial=0,
            total=total_batches,
            disable=(not accelerator.is_main_process),
            smoothing=0.02,
        )

        # Iterate through batches
        for idx, batch in enumerate(dataloader):
            # Update local batch counter
            input_ids, ids, max_seqlen, cu_seqlens, position_ids, attention_mask, masked_index = (
                batch["input_ids"],
                batch["id"],
                batch.get("max_seqlen", None),
                batch.get("cu_seqlens", None),
                batch.get("position_ids", None),
                batch.get("attention_mask", None),
                batch.get("masked_index", None),
            )
            targets = batch["target"]

            # Compute model output
            with torch.no_grad(), accelerator.autocast():
                # Forward pass
                if self.packing:
                    output = model(
                        input_ids,
                        position_ids=position_ids,
                        max_seqlen=max_seqlen,
                        cu_seqlens=cu_seqlens,
                    )
                else:
                    attention_mask = torch.where(attention_mask.to(torch.bool), float(0.0), float("-inf")).to(torch.float16)
                    output = model(
                        input_ids,
                        attention_mask,
                    )

            if self.packing:
                # Find the masked token for each sequence
                logits_flat = output["logits"].view(-1, output["logits"].size(-1))  # [total_length, vocab_size]
                masked_token = (input_ids.view(-1) == tokenizer.mask_token_id).nonzero(as_tuple=True)[0]

                # Get logits at masked positions
                logits = logits_flat[masked_token]

            else:
                # Find the masked token for each sequence
                masked_token = (input_ids == tokenizer.mask_token_id).nonzero(as_tuple=False)

                # Get logits at masked positions
                logits = output["logits"][masked_token[:, 0], masked_token[:, 1]]

            log_probs = torch.log_softmax(logits, dim=-1)
            target_log_probs = log_probs[torch.arange(logits.size(0)), targets]

            results["masked_index"].extend(masked_index.cpu().numpy())
            results["id"].extend(ids)
            results["log_prob"].extend(target_log_probs.cpu().numpy())

            pbar.update(1)

            if idx > 0 and idx % self.save_steps == 0:
                gathered = accelerator.gather_for_metrics(
                    [
                        {
                            "id": id_,
                            "masked_index": idx_,
                            "log_prob": lp_,
                        }
                        for id_, idx_, lp_ in zip(results["id"], results["masked_index"], results["log_prob"])
                    ]
                )

                accelerator.save_state()

                if accelerator.is_main_process:
                    df = pd.DataFrame(gathered)
                    output_file = os.path.join(output_path, f"data_{(idx + start_batch):05d}.parquet")
                    df.to_parquet(output_file, index=False, compression="gzip")
                    self.logger.info(f"Data saved in file {output_file}")

                results = {"log_prob": [], "id": [], "masked_index": []}

        gathered = accelerator.gather_for_metrics(
            [
                {
                    "id": id_,
                    "masked_index": idx_,
                    "log_prob": lp_,
                }
                for id_, idx_, lp_ in zip(results["id"], results["masked_index"], results["log_prob"])
            ]
        )

        if accelerator.is_main_process:
            df = pd.DataFrame(gathered)
            output_file = os.path.join(output_path, f"data_{idx + start_batch:05d}.parquet")
            df.to_parquet(output_file, index=False, compression="gzip")
            self.logger.info(f"Data saved in file {output_file}")

        pbar.close()
        accelerator.end_training()

        with open(os.path.join(output_path, "done.txt"), "w") as f:
            f.write("done")


def load_preprocess_data(input_csv, n_samples=0):
    """
    Loads the base dataset from a CSV, samples if needed,
    and preprocesses columns (name -> entry_name, extracts uniprot_id).
    """
    print(f"Loading base dataset from {input_csv}...")
    df_main = pd.read_csv(input_csv)

    if n_samples > 0:
        print(f"Using a subset of {n_samples} sequences.")
        df_main = df_main.sample(n=n_samples, random_state=42).reset_index(drop=True)

    if "name" in df_main.columns:
        df_main = df_main.rename(columns={"name": "entry_name"})
        
    if "id" in df_main.columns:
        df_main = df_main.rename(columns={"id": "entry_name"})

    # Ensure required columns exist
    if "entry_name" not in df_main.columns or "sequence" not in df_main.columns:
        print("Error: Input CSV must contain 'entry_name' (or 'name', 'id') and 'sequence' columns.")
        exit(1)

    try:
        df_main["uniprot_id"] = df_main["entry_name"].apply(lambda x: x.split("|")[1])
    except IndexError:
        df_main["uniprot_id"] = df_main["entry_name"]
    df_main["sequence"] = df_main["sequence"].astype(str)
    df_main["uniprot_id"] = df_main["uniprot_id"].astype(str)

    print(f"Loaded and preprocessed {len(df_main)} sequences.")
    return df_main


def download_structure(uniprot_id, output_dir):
    """
    Downloads AFv6 PDB from alphafold.ebi.ac.uk, returns status string.
    """
    file_name = f"AF-{uniprot_id}-F1-model_v6.pdb"
    output_path = Path(output_dir) / file_name

    if output_path.exists():
        return (uniprot_id, "skipped")

    try:
        response = requests.get(f"https://alphafold.ebi.ac.uk/files/{file_name}", timeout=60)
        if response.status_code == 200:
            output_path.write_bytes(response.content)
            return (uniprot_id, "success")
        elif response.status_code == 404:
            return (uniprot_id, "not_found")
        else:
            return (uniprot_id, "http_error")
    except requests.RequestException:
        return (uniprot_id, "request_error")
```
