## Supplementary material for "pLM representations unlock metagenomic space beyond homology": Code bundle: config.yaml

```
defaults:
  - _self_
  # For general configurations of metrics, decision metrics, and target columns
  - sequence_classification: sequence_classification
  - token_classification: token_classification
  - sequence_regression: sequence_regression
  # Specific configurations for the task
  - model:
    - esm_150M
  - data_config:
    - stability_prediction
  - grid: mlp_feature_extraction_restricted

head_type: mlp
do_test: true
return_all_results: true
verbose: reduced
name: null
device: cuda

base_folder: $SCRATCH/apperitif/
output_folder: null
```
