## Supplementary material for "pLM representations unlock metagenomic space beyond homology": Code bundle: biomap_research_sequence_classification.yaml

```
# Base configuration for biomap research sequence classification tasks
_target_: apperitif.tasks.sequence_classification.biomap_research.BiomapResearchSequenceClassificationConfig
sequence_column: seq
label_column: label
label_dtype: int
task_type: "sequence_classification"
```
