## Supplementary material for "pLM representations unlock metagenomic space beyond homology": Code bundle: biomap_research_sequence_regression.yaml

```
# Base configuration for biomap research sequence regression tasks
_target_: apperitif.tasks.sequence_regression.biomap_research.BiomapResearchSequenceRegressionConfig
sequence_column: seq
label_column: label
label_dtype: float
task_type: "sequence_regression"
```
