## Supplementary material for "pLM representations unlock metagenomic space beyond homology": Code bundle: biomap_research_token_classification.yaml

```
# Base configuration for biomap research token classification tasks
_target_: apperitif.tasks.token_classification.biomap_research.BiomapResearchTokenClassificationConfig
sequence_column: seq
label_column: label
label_dtype: int
task_type: "token_classification"
```
