## Supplementary material for "pLM representations unlock metagenomic space beyond homology": Code bundle: mlp_feature_extraction_extended.yaml

```
# MLP Feature Extraction Restricted Grid (frozen backbone, smaller search space)
# This grid is used for feature extraction with frozen backbone and restricted hyperparameters

# Override base MLP parameters for feature extraction
finetuning: false
precomputed: true

# Restricted hyperparameter grid for feature extraction
hidden_size: [512, 1024, 2048, 4096]
learning_rate: [5e-3, 1e-3, 5e-4, 1e-4, 5e-5, 1e-5]
num_epochs: [3, 5, 10, 20, 40, 80]
weight_decay: [0, 1e-4, 1e-3, 1e-2]
batch_size: [16, 32, 64, 128, 256]
seeds: [42, 43, 44]
subbenchmark_name: mlp_feature_extraction_extended
```
