## Supplementary material for "pLM representations unlock metagenomic space beyond homology": Code bundle: mlp_feature_extraction_restricted.yaml

```
# MLP Feature Extraction Grid (frozen backbone)
# This grid is used for feature extraction with frozen backbone

# Override base MLP parameters for feature extraction
finetuning: false
precomputed: true

# Hyperparameter grid for feature extraction
hidden_size: [1024, 2048]
learning_rate: [1e-3, 5e-4, 1e-4]
num_epochs: [10, 20, 40]
weight_decay: [1e-4, 1e-3]
batch_size: [32, 64, 128]
seeds: [42, 43, 44]
subbenchmark_name: mlp_feature_extraction_restricted
```
