## Supplementary material for "pLM representations unlock metagenomic space beyond homology": Code bundle: mlp_finetuning_extended.yaml

```
# MLP Finetuning Restricted Grid (unfrozen backbone, extended search space)
# This grid is used for finetuning with unfrozen backbone and extended hyperparameters

# Override base MLP parameters for finetuning
finetuning: true
precomputed: false

# Extended hyperparameter grid for finetuning
hidden_size: [128, 256, 512, 1024]
learning_rate: [1e-3, 5e-4, 1e-4, 5e-5, 1e-5]
num_epochs: [3, 5, 10, 20]
weight_decay: [0, 1e-5, 1e-4, 1e-3]
batch_size: [4, 8, 16, 32]
seeds: [42, 43, 44]
subbenchmark_name: mlp_finetuning_extended
```
