## Supplementary material for "pLM representations unlock metagenomic space beyond homology": Code bundle: mlp_finetuning_restricted.yaml

```
# MLP Finetuning Grid (unfrozen backbone)
# This grid is used for finetuning with unfrozen backbone

# Override base MLP parameters for finetuning
finetuning: true
precomputed: false

# Hyperparameter grid for finetuning
hidden_size: [128, 256, 512]
learning_rate: [1e-4, 5e-5, 1e-5]
num_epochs: [3, 5, 10]
weight_decay: [0, 1e-4, 1e-3]
batch_size: [8, 16, 32]
seeds: [42]
subbenchmark_name: mlp_finetuning_restricted
```
