## Supplementary material for "pLM representations unlock metagenomic space beyond homology": Code bundle: amplify_120M_local.yaml

```
defaults:
  - amplify
  - _self_

model_name: "MILA_U100"
from_hub: false
base_path: "proseqo/ablations"
vocab_path: "/path/to/AMPLIFY/conf/tokenizer/amplify_vocab.txt"
tag: null
attention_mask_type: bool
attention_mask_dtype: bool
num_total_attn_maps: 240
use_packed_sequence: true
```
