## Supplementary material for "pLM representations unlock metagenomic space beyond homology": Code bundle: amplify_350M_packed.yaml

```
defaults:
  - amplify
  - _self_

local_name: amplify_350M_packed
model_name: "${oc.env:HF_ORG,your-hf-org}/AMPLIFY_350M"
attention_mask_type: bool
attention_mask_dtype: bool
num_total_attn_maps: 480
use_packed_sequence: true
hf_kwargs:
  trust_remote_code: true
```
