## Supplementary material for "pLM representations unlock metagenomic space beyond homology": Code bundle: sequence_regression.yaml

```
# Base configuration for sequence regression evaluators
# This file contains the common task-level parameters that apply to all evaluators for sequence regression

# Common task-level parameters
metrics: ["spearmanr", "pearsonr", "mse", "r2"]
decision_metric: "spearmanr"
reduction_method: "mean"
target_column: "labels"
precomputing_batch_size: 32
autocast_precision: "fp16"
```
