## Supplementary material for "pLM representations unlock metagenomic space beyond homology": Code bundle: token_classification.yaml

```
# Base configuration for token classification evaluators
# This file contains the common task-level parameters that apply to all evaluators for token classification

# Common task-level parameters
metrics: ["accuracy", "f1"]
decision_metric: "accuracy"
target_column: "labels"
precomputing_batch_size: 32
autocast_precision: "fp16"
reduction_method: "mean"
```
