## Supplementary material for "pLM representations unlock metagenomic space beyond homology": Code bundle: consensus.sh

```
#!/bin/bash
#SBATCH --job-name=consensus
#SBATCH --cpus-per-task=6
#SBATCH --mem=32G
#SBATCH --time=1-00:00:00
#SBATCH --output=logs/consensus/%x_%j.out
#SBATCH --error=logs/consensus/%x_%j.out
#SBATCH --nodes=1
#SBATCH --ntasks-per-node=1
#SBATCH --gres=gpu:l40s:1
#SBATCH --partition=long

# Usage: sbatch --job-name=<model> consensus.sh
# <model> uses dots (e.g. MILA_ALL_0.50_1m); HF hub name (underscores) is derived automatically.

module load python/3.10 cuda/12.6.0
source $HOME/proseqo/.venv/bin/activate

SLURM_JOB_NAME=MILA_U100_0.0_mila

# facebook/* models are always loaded from the hub
MODEL_FROM_HUB=0
if [[ "$SLURM_JOB_NAME" == facebook/* ]]; then
    MODEL_FROM_HUB=1
fi

# Check if checkpoint path exists for local models
if [ "$MODEL_FROM_HUB" -eq 0 ] && [ ! -d "$SCRATCH/proseqo/ablations/$SLURM_JOB_NAME/model_checkpoints" ]; then
    echo "Checkpoint path not found for $SLURM_JOB_NAME, loading from hub."
    MODEL_FROM_HUB=1
    MODEL_NAME="${HF_ORG:-your-hf-org}/${SLURM_JOB_NAME//./_}"
    echo "Derived HF hub model name: $MODEL_NAME"
else
    MODEL_NAME="$SLURM_JOB_NAME"
fi

# export MASTER_PORT=$(expr 10000 + $(echo -n $SLURM_JOBID | tail -c 4))
# export MASTER_ADDR=$(scontrol show hostnames $SLURM_JOB_NODELIST | head -n 1)

# cmd=(
#     accelerate launch \
#             --config_file=conf/accelerate_ddp.yaml \
#             --machine_rank=\$SLURM_NODEID \
#             --main_process_ip=$MASTER_ADDR \
#             --main_process_port=$MASTER_PORT \
#             --num_machines=1 \
#             --num_processes=1 \
#         scripts/eval_consensus/sequence_recovery.py \
#         --model_name $MODEL_NAME \
#         --config_path $SCRATCH/proseqo/ablations/$SLURM_JOB_NAME/.hydra/config.yaml \
#         --checkpoint_path $SCRATCH/proseqo/ablations/$SLURM_JOB_NAME/model_checkpoints \
#         --vocab_path /path/to/AMPLIFY/conf/tokenizer/amplify_vocab.txt \
#         --batch_size 128 \
#         --data_path $SCRATCH/proseqo/evaluation/human_proteome_masked.csv \
#         --output_path $SCRATCH/proseqo/sequence_recovery/$SLURM_JOB_NAME \
#         --categories_path data \
#         --save_steps 2000 \
# \
#         --packing true \
#         --compute_scores \
#         --skip_predictions
# )

# srun --kill-on-bad-exit=1 bash -c "$(for a in "${cmd[@]}" ; do echo -n \"$a\" "" ; done)"

python scripts/eval_consensus/sequence_recovery.py \
        --model_name $MODEL_NAME \
        --config_path $SCRATCH/proseqo/ablations/$SLURM_JOB_NAME/.hydra/config.yaml \
        --checkpoint_path $SCRATCH/proseqo/ablations/$SLURM_JOB_NAME/model_checkpoints \
        --vocab_path /path/to/AMPLIFY/conf/tokenizer/amplify_vocab.txt \
        --batch_size 128 \
        --data_path $SCRATCH/proseqo/evaluation/human_proteome_masked.csv \
        --output_path $SCRATCH/proseqo/sequence_recovery/$SLURM_JOB_NAME \
        --categories_path data \
        --save_steps 2000 \
        --packing true \
        --compute_scores \
        --skip_predictions
        
```
