## Supplementary material for "pLM representations unlock metagenomic space beyond homology": Code bundle: jacobian.sh

```
#!/bin/bash
#SBATCH --job-name=jacobian
#SBATCH --cpus-per-task=6
#SBATCH --mem=32G
#SBATCH --time=3:00:00
#SBATCH --output=logs/jacobian/%x_%j.out
#SBATCH --error=logs/jacobian/%x_%j.out
#SBATCH --ntasks-per-node=1
#SBATCH --gres=gpu:l40s:1
#SBATCH --partition=long

# Usage: sbatch --job-name=<model> jacobian.sh [split]
# <model> uses dots (e.g. MILA_ALL_0.50_1m); HF hub name (underscores) is derived automatically.

module load python/3.10 cuda/12.6.0
source $HOME/proseqo/.venv/bin/activate

# Derive HF hub name: dots -> underscores (e.g. 0.50 -> 0_50) if model does not start with facebook/
if [[ "$SLURM_JOB_NAME" == facebook/* ]] || [[ "$SLURM_JOB_NAME" == flair-bio/* ]]; then
    MODEL_NAME="$SLURM_JOB_NAME"
else
    MODEL_NAME="${HF_ORG:-your-hf-org}/${SLURM_JOB_NAME//./_}"
fi

split=${1:-casp16}
output_dir=$SCRATCH/proseqo/categorical_jacobian/$SLURM_JOB_NAME
results_path=$output_dir/results_${split}.json

mkdir -p $output_dir

srun --kill-on-bad-exit=1 python \
    scripts/eval_categorical_jacobian/categorical_jacobian.py \
    --model $MODEL_NAME \
    --split $split \
    --batch-size 64 \
    --results $results_path

exit_code=$?

if [ $exit_code -ne 0 ]; then
    echo "Job failed with exit code $exit_code"
    echo $SLURM_JOB_NAME >> $SCRATCH/proseqo/categorical_jacobian/fail.txt
    exit $exit_code
fi

echo "Job completed successfully!"
echo $SLURM_JOB_NAME >> $SCRATCH/proseqo/categorical_jacobian/success.txt
```
