## Supplementary material for "pLM representations unlock metagenomic space beyond homology": Code bundle: kl_consensus.sh

```
#!/bin/bash
#SBATCH --job-name=kl_consensus
#SBATCH --cpus-per-task=6
#SBATCH --mem=32G
#SBATCH --time=06:00:00
#SBATCH --output=logs/kl_consensus/%x_%j.out
#SBATCH --error=logs/kl_consensus/%x_%j.out
#SBATCH --nodes=1
#SBATCH --ntasks-per-node=1
#SBATCH --gres=gpu:l40s:1
#SBATCH --partition=unkillable

# Usage: sbatch --job-name=<model> kl_consensus.sh
# <model> is the base model name (e.g. MILA_U100_0.30, AMPLIFY_benchmark).

module load python/3.10 cuda/12.6.0
source $HOME/AMPLIFY/.venv/bin/activate

# MODEL_BASE="$SLURM_JOB_NAME"
# MODEL_BASE="flair-bio/amplify-120m"  # For testing; comment out for production
MODEL_BASE="AMPLIFY_120M"  # For testing; comment out for production

# Org-qualified names (containing a '/') are loaded from the HF hub.
FROM_HUB=0
if [[ "$MODEL_BASE" == */* ]]; then
    FROM_HUB=1
fi

# Check if local checkpoint exists; fall back to HF hub if not
if [ "$FROM_HUB" -eq 0 ] && [ ! -d "$SCRATCH/proseqo/ablations/$MODEL_BASE/model_checkpoints" ]; then
    echo "Checkpoint path not found for $MODEL_BASE, loading from hub."
    FROM_HUB=1
fi

if [ "$FROM_HUB" -eq 1 ]; then
    if echo "$MODEL_BASE" | grep -qi "facebook"; then
        MODEL_NAME="$MODEL_BASE"
    elif echo "$MODEL_BASE" | grep -qi "flair-bio"; then
        MODEL_NAME="$MODEL_BASE"
    elif echo "$MODEL_BASE" | grep -qi "chandar-lab"; then
        MODEL_NAME="$MODEL_BASE"
    else
        MODEL_NAME="${HF_ORG:-your-hf-org}/${MODEL_BASE//./_}"
    fi
    echo "Loading from HF hub: $MODEL_NAME"
else
    MODEL_NAME="$MODEL_BASE"
    echo "Loading locally: $MODEL_NAME"
fi

OUTPUT_DIR="$SCRATCH/proseqo/kl_consensus/$MODEL_BASE"
mkdir -p "$OUTPUT_DIR"

python scripts/eval_consensus/kl_consensus.py \
    --model_name "$MODEL_NAME" \
    --config_path "$SCRATCH/proseqo/ablations/$MODEL_BASE/.hydra/config.yaml" \
    --checkpoint_path "$SCRATCH/proseqo/ablations/$MODEL_BASE/model_checkpoints" \
    --vocab_path "$HOME/AMPLIFY/conf/tokenizer/amplify_vocab.txt" \
    --proteome_path "$SCRATCH/proseqo/evaluation/human_proteome.csv" \
    --consensus_path "$SCRATCH/proseqo/human_consensus_positions.csv" \
    --dataset_path "$SCRATCH/proseqo/kl_consensus/dataset_cache" \
    --output_path "$OUTPUT_DIR" \
    --batch_size 64 \
    --save_steps 500 \
    --compute_scores
```
