## Supplementary material for "pLM representations unlock metagenomic space beyond homology": Code bundle: ppl.sh

```
#!/bin/bash
#SBATCH --job-name=ppl
#SBATCH --cpus-per-task=6
#SBATCH --mem=32G
#SBATCH --time=6:00:00
#SBATCH --output=logs/ppl/%x_%j.out
#SBATCH --error=logs/ppl/%x_%j.out
#SBATCH --ntasks-per-node=1
#SBATCH --gres=gpu:l40s:1
#SBATCH --partition=long

# Usage: sbatch --job-name=<model> ppl.sh <data_name> [tag]
# <model> uses dots (e.g. MILA_ALL_0.50_1m); HF hub name (underscores) is derived automatically.

module load python/3.10 cuda/12.6.0
source $HOME/proseqo/.venv/bin/activate

# Derive HF hub name: dots -> underscores (e.g. 0.50 -> 0_50)
MODEL_NAME="${SLURM_JOB_NAME//./_}"

data_name=$1

if [ -n "$2" ]; then
    tag="$2"
else
    tag="latest"
fi

data_path=$SCRATCH/proseqo/datasets/uniprotkb_ec_1_14_11_2025_10_20_max_512_masked.csv

export MASTER_PORT=$(expr 10000 + $(echo -n $SLURM_JOBID | tail -c 4))
export MASTER_ADDR=$(scontrol show hostnames $SLURM_JOB_NODELIST | head -n 1)

cmd=(
    accelerate launch \
        --config_file=conf/accelerate_ddp.yaml \
        --machine_rank=\$SLURM_NODEID \
        --main_process_ip=$MASTER_ADDR \
        --main_process_port=$MASTER_PORT \
        --num_machines=1 \
        --num_processes=1 \
    scripts/eval_ppl/pseudo_perplexity.py \
    --model_name $MODEL_NAME \
    --data_name $data_name \
    --config_path $SCRATCH/proseqo/ablations/$SLURM_JOB_NAME/.hydra/config.yaml \
    --checkpoint_path $SCRATCH/proseqo/ablations/$SLURM_JOB_NAME/model_checkpoints \
    --vocab_path /path/to/AMPLIFY/conf/tokenizer/amplify_vocab.txt \
    --batch_size 128 \
    --data_path $data_path \
    --output_path $SCRATCH/proseqo \
    --compute_scores \
    --tag $tag \
)

srun --kill-on-bad-exit=1 bash -c "$(for a in "${cmd[@]}" ; do echo -n \"$a\" "" ; done)"
```
