## Supplementary material for "pLM representations unlock metagenomic space beyond homology": Code bundle: valset.sh

```
#!/bin/bash
#SBATCH --job-name=valset
#SBATCH --cpus-per-task=6
#SBATCH --mem=32G
#SBATCH --time=1:00:00
#SBATCH --output=logs/valset/%x_%j.out
#SBATCH --error=logs/valset/%x_%j.out
#SBATCH --gres=gpu:1

# Usage: sbatch --job-name=<model> valset.sh [model] [--local]
# Pass --local to load from $SCRATCH/proseqo/ablations/ instead of the HF hub.
# facebook/* models are always loaded from the HF hub regardless of this flag.
# Results are always stored under the base model name (no ${HF_ORG:-your-hf-org}/ prefix).

# # Parse --local flag
# FROM_HUB=1
# for arg in "$@"; do
#     case "$arg" in
#         --local) FROM_HUB=0 ;;
#     esac
# done

# If a single model is passed as argument, use it; otherwise iterate over the
# internal list below.
if [ -n "${1:-}" ]; then
    MODELS=("$1")
fi

# Define the models (used when no argument is passed)
MODELS=(
    'MILA_ALL_0.50_1m'
    'MILA_ALL_0.60_1m'
    'MILA_ALL_0.70_1m'
    'MILA_ALL_0.80_1m'
    'MILA_BFD_0.20_1m'
    'MILA_BFD_0.30_1m'
    'MILA_BFD_0.40_1m'
    'MILA_BFD_0.50_1m'
    'MILA_BFD_0.60_1m'
    'MILA_BFD_0.70_1m'
    'MILA_BFD_0.80_1m'
    'MILA_MGNIFY_0.20_1m'
    'MILA_MGNIFY_0.30_1m'
    'MILA_MGNIFY_0.40_1m'
    'MILA_OPT_0.10_sub10_1m'
    'MILA_OPT_0.20_sub20_1m'
    'MILA_OPT_0.30_sub30_1m'
    'MILA_OPT_0.40_sub40_1m'
    'MILA_U100_0.05_1m'
    'MILA_U100_0.10_1m'
    'MILA_U100_0.15_1m'
    'MILA_U100_0.20_1m'
    'MILA_U100_0.25_1m'
    'MILA_U100_0.30_1m'
    'MILA_U100_0.35_1m'
    'MILA_U100_0.40_1m'
    # "${HF_ORG:-your-hf-org}/AMPLIFY_350M"
    # "${HF_ORG:-your-hf-org}/AMPLIFY_120M"
)

cd ${EVAL_ROOT:-.}
module load python/3.10 cuda/12.6.0
source $HOME/AMPLIFY/.venv/bin/activate

max_len=512

for model in "${MODELS[@]}"; do
    CURRENT_MODEL=$model
    echo "Model: $CURRENT_MODEL"

    if [ -f $SCRATCH/proseqo/uniprot_validation/$CURRENT_MODEL/result.json ]; then
        echo "Result file already exists for $CURRENT_MODEL, skipping..."
        continue
    fi

    # facebook/* models are always loaded from the hub
    MODEL_FROM_HUB=0
    if [[ "$CURRENT_MODEL" == facebook/* ]]; then
        MODEL_FROM_HUB=1
    fi

    # Check if checkpoint path exists for local models
    if [ "$MODEL_FROM_HUB" -eq 0 ] && [ ! -d "$SCRATCH/proseqo/ablations/$CURRENT_MODEL/model_checkpoints" ]; then
        echo "Checkpoint path not found for $CURRENT_MODEL, loading from hub."
        MODEL_FROM_HUB=1
        HUB_MODEL="${HF_ORG:-your-hf-org}/${CURRENT_MODEL//./_}"
        echo "Derived HF hub model name: $HUB_MODEL"
    fi

    if [ "$MODEL_FROM_HUB" -eq 1 ]; then
        if [[ "$CURRENT_MODEL" == */* ]]; then
            HUB_MODEL="$CURRENT_MODEL"
        else
            HUB_MODEL="${HF_ORG:-your-hf-org}/$CURRENT_MODEL"
        fi
        # HF repo names use underscores instead of dots (e.g. 0.50 -> 0_50)
        HUB_MODEL="${HUB_MODEL//./_}"
        echo "Running with model from Hugging Face Hub: $HUB_MODEL"
        python scripts/eval_validation/validation_set_eval.py \
            --data_paths $SCRATCH/proseqo/datasets/uniprot.csv \
            --model_name $HUB_MODEL \
            --output_path $SCRATCH/proseqo/uniprot_validation/$CURRENT_MODEL/result.json \
            --token $HF_TOKEN \
            --max_len $max_len \
            --packing 0
    else
        echo "Running with local model: $CURRENT_MODEL"
        python scripts/eval_validation/validation_set_eval.py \
            --data_paths $SCRATCH/proseqo/datasets/uniprot.csv \
            --model_name $CURRENT_MODEL \
            --checkpoint_path $SCRATCH/proseqo/ablations/$CURRENT_MODEL/model_checkpoints \
            --config_path $SCRATCH/proseqo/ablations/$CURRENT_MODEL/.hydra/config.yaml \
            --vocab_path $HOME/AMPLIFY/conf/tokenizer/amplify_vocab.txt \
            --output_path $SCRATCH/proseqo/uniprot_validation/$CURRENT_MODEL/result.json \
            --max_len $max_len \
            --packing 0
    fi
done

exit_code=$?

if [[ "$exit_code" -ne 0 ]]; then
    echo "Job failed with exit code $exit_code"
    echo $CURRENT_MODEL >> $SCRATCH/proseqo/uniprot_validation/fail.txt
    exit $exit_code
fi

echo "Job completed successfully!"
echo $CURRENT_MODEL >> $SCRATCH/proseqo/uniprot_validation/success.txt
exit $exit_code
```
