## Supplementary material for "pLM representations unlock metagenomic space beyond homology": Code bundle: README.md

# Evaluation Jobs

SLURM job scripts for the evaluation tasks. Each script submits **one model on one task**; run it per model to reproduce the corresponding results. Batch orchestration (looping over model lists) is intentionally not included — drive it from your own submission loop.

## Directory structure

```
jobs/
├── evaluation/                 # zero-shot / calibration task jobs
│   ├── ppl.sh                  # Pseudo-perplexity
│   ├── consensus.sh            # Sequence recovery
│   ├── kl_consensus.sh         # Calibration (KL to evolutionary consensus)
│   ├── jacobian.sh             # Categorical Jacobian (contact prediction)
│   └── valset.sh               # Validation-set language modeling
└── xtrimo/                     # supervised property-prediction jobs
    ├── xtrimo_fe.sh            # MLP feature extraction (frozen backbone)
    ├── xtrimo_ft.sh            # MLP finetuning (unfrozen backbone)
    └── mlp_randomized.sh       # Randomized-baseline transfer scores
```

Logs are written to `logs/<task>/`.

## Specifying a model

`<model>` is the base model name:
- Base name only — no `<HF_ORG>/` prefix (e.g. `MILA_ALL_0.50_1m`).
- `facebook/*` models keep their full prefix (e.g. `facebook/esm2_t33_650M_UR50D`).
- Dots (e.g. `0.50`) are fine — HF hub names with underscores (e.g. `0_50`) are derived automatically.

## Hub vs. local loading

Most tasks support two model sources:

| Flag | Behaviour |
|------|-----------|
| *(default, no flag)* | Load from Hugging Face Hub as `<HF_ORG>/<model>` |
| `--local` | Load from `$SCRATCH/proseqo/ablations/<model>/` |

**`facebook/*` models are always loaded from the Hub regardless of this flag.**

The Hub organization is read from the `HF_ORG` environment variable (default: `your-hf-org`).
Set it to the org hosting the released AMPLIFY checkpoints, e.g. `export HF_ORG=chandar-lab`.

### Dot → underscore conversion for HF hub names

HF repository names cannot contain dots. The scripts automatically convert dots to underscores when constructing the hub model name:

- `MILA_ALL_0.50_1m` → `<HF_ORG>/MILA_ALL_0_50_1m` (passed to Python / HF)
- Results are **always stored under the original base name with dots** (e.g. `$SCRATCH/proseqo/.../MILA_ALL_0.50_1m/`)

This conversion applies to: `xtrimo_fe.sh`, `xtrimo_ft.sh`, `valset.sh`, `consensus.sh`, `ppl.sh`, `jacobian.sh`.

## Per-task job scripts

### `xtrimo_fe.sh` — MLP feature extraction

```bash
sbatch --job-name=<task>_<model> jobs/xtrimo/xtrimo_fe.sh <task> <model> [--local]
```

- Runs `scripts/main.py` with `grid=mlp_feature_extraction_extended`.
- Results written to `$SCRATCH/proseqo/apperitif/` under the base model name.
- Success/fail logged to `$SCRATCH/proseqo/apperitif/mlp_feature_extraction/{success,fail}.txt`.

### `xtrimo_ft.sh` — MLP finetuning (unfrozen backbone)

```bash
# As a SLURM array job (one array element per task):
sbatch --job-name=<model> --array=0-16 jobs/xtrimo/xtrimo_ft.sh [--local]

# Or for a single task:
sbatch --job-name=<model> jobs/xtrimo/xtrimo_ft.sh <task> <model> [--local]
```

- Runs `scripts/main.py` with `grid=mlp_finetuning_extended`.
- Results written to `$SCRATCH/proseqo/apperitif/` under the base model name.
- Success/fail logged to `$SCRATCH/proseqo/apperitif/mlp_finetuning_extended/{success,fail}.txt`.

### `mlp_randomized.sh` — Randomized-baseline transfer scores

```bash
sbatch --job-name=<model> jobs/xtrimo/mlp_randomized.sh <task> <model>
```

- Produces the randomly-initialized baseline used for the transfer-learning comparison.

### `ppl.sh` — Pseudo-perplexity

```bash
sbatch --job-name=<model> jobs/evaluation/ppl.sh <data_name> [tag]
```

- Runs `scripts/eval_ppl/pseudo_perplexity.py` via `accelerate launch`.
- Output written to `$SCRATCH/proseqo/pseudo_ppl/<model>/<data_name>/`.

### `consensus.sh` — Sequence recovery

```bash
sbatch --job-name=<model> jobs/evaluation/consensus.sh
```

- Runs `scripts/eval_consensus/sequence_recovery.py` via `accelerate launch`.
- Output written to `$SCRATCH/proseqo/sequence_recovery/<model>/`.

### `kl_consensus.sh` — Calibration

```bash
sbatch --job-name=<model> jobs/evaluation/kl_consensus.sh
```

- Runs `scripts/eval_consensus/kl_consensus.py`.
- Output written under `$SCRATCH/proseqo/kl_consensus/`.

### `jacobian.sh` — Categorical Jacobian

```bash
sbatch --job-name=<model> jobs/evaluation/jacobian.sh [split]
```

- `split` defaults to `casp16`. Override by passing it as the first argument (e.g. `sbatch ... jacobian.sh cameo`).
- Hub-only: always loads from `<HF_ORG>/<model>` (no `--local` support).
- Output written to `$SCRATCH/proseqo/categorical_jacobian/<model>/results_<split>.json`.

### `valset.sh` — Validation set

```bash
sbatch --job-name=<model> jobs/evaluation/valset.sh [model] [--local]
```

- Runs `scripts/eval_validation/validation_set_eval.py`.
- Output written to `$SCRATCH/proseqo/uniprot_validation/<model>/result.json`.
