## Supplementary material for "pLM representations unlock metagenomic space beyond homology": Code bundle: mlp_randomized.sh

```
#!/bin/bash
#SBATCH --job-name=mlp_randomized
#SBATCH --cpus-per-task=6
#SBATCH --mem=32G
#SBATCH --gres=gpu:l40s:1
#SBATCH --time=6:00:00
#SBATCH --output=logs/mlp_randomized/%x_%j.out
#SBATCH --error=logs/mlp_randomized/%x_%j.out
#SBATCH --partition=long

# Usage: sbatch --job-name=<task>_<model>_randomized mlp_randomized.sh <task> <model>
# Or as a SLURM array job (set SLURM_ARRAY_TASK_ID externally).

export HF_HUB_OFFLINE=1

DATA_CONFIGS=(
    "antibiotic_resistance"
    "cloning_clf"
    "enzyme_catalytic_efficiency"
    "fitness_prediction"
    "fluorescence_prediction"
    "fold_prediction"
    "localization_prediction"
    "material_production"
    "metal_ion_binding"
    "optimal_ph"
    "optimal_temperature"
    "peptide_hla_mhc_affinity"
    "solubility_prediction"
    "ssp_q3"
    "ssp_q8"
    "stability_prediction"
    "tcr_pmhc_affinity"
)

MODELS=(
    "esm_35M"
    "esm_150M"
    "esm_650M"
    "amplify_120M_packed"
    "amplify_350M_packed"
)

TOTAL_TASKS=${#DATA_CONFIGS[@]}
TOTAL_MODELS=${#MODELS[@]}
TOTAL_COMBINATIONS=$((TOTAL_TASKS * TOTAL_MODELS))

if [ -n "$1" ] && [ -n "$2" ]; then
    CURRENT_TASK=$1
    CURRENT_MODEL=$2
    echo "Using arguments passed"
else
    TASK_ID=$SLURM_ARRAY_TASK_ID
    MODEL_INDEX=$((TASK_ID % TOTAL_MODELS))
    TASK_INDEX=$((TASK_ID / TOTAL_MODELS))
    CURRENT_TASK=${DATA_CONFIGS[$TASK_INDEX]}
    CURRENT_MODEL=${MODELS[$MODEL_INDEX]}
    echo "Using Job ID: $SLURM_ARRAY_TASK_ID"
fi

echo "Task: $CURRENT_TASK"
echo "Model: $CURRENT_MODEL"

cd ${EVAL_ROOT:-.}
module load python/3.10 cuda/12.6.0
source $HOME/AMPLIFY/.venv/bin/activate

python scripts/main.py \
    data_config=$CURRENT_TASK \
    model=$CURRENT_MODEL \
    grid=mlp_feature_extraction_restricted \
    grid.subbenchmark_name=mlp_feature_extraction \
    head_type=mlp \
    do_test=true \
    return_all_results=true \
    base_folder=$SCRATCH/proseqo/apperitif \
    verbose=true \
    +model.randomize=true \
    +model.randomize_mode=init \
    +precomputing_batch_size=4

exit_code=$?

if [[ "$exit_code" -ne 0 ]]; then
    echo "Job failed with exit code $exit_code"
    echo "${CURRENT_MODEL}_randomized, ${CURRENT_TASK}" >> $SCRATCH/proseqo/apperitif/mlp_feature_extraction/fail.txt
    exit $exit_code
fi

echo "Job completed successfully!"
echo "${CURRENT_MODEL}_randomized, ${CURRENT_TASK}" >> $SCRATCH/proseqo/apperitif/mlp_feature_extraction/success.txt
exit $exit_code
```
