## Supplementary material for "pLM representations unlock metagenomic space beyond homology": Code bundle: xtrimo_fe.sh

```
#!/bin/bash
#SBATCH --job-name=xtrimo_fe
#SBATCH --cpus-per-task=6
#SBATCH --mem=32G
#SBATCH --time=8:00:00
#SBATCH --output=logs/xtrimo_fe/%x_%j.out
#SBATCH --error=logs/xtrimo_fe/%x_%j.out
#SBATCH --partition=lab-chandar
#SBATCH --gres=gpu:1

# Usage: sbatch --job-name=<task>_<model> xtrimo_fe.sh <task> <model> [--local]
# Pass --local to load from $SCRATCH/proseqo/ablations/ instead of the HF hub.
# facebook/* models are always loaded from the HF hub regardless of this flag.
# Results are always stored under the base model name (no ${HF_ORG:-your-hf-org}/ prefix).

# Parse arguments: positional task/model, optional --local flag
FROM_HUB=1
for arg in "$@"; do
    case "$arg" in
        --local) FROM_HUB=0 ;;
    esac
done

DATA_CONFIGS=(
    "antibiotic_resistance"
    "cloning_clf"
    "enzyme_catalytic_efficiency"
    "fitness_prediction"
    "fluorescence_prediction"
    "fold_prediction"
    "localization_prediction"
    "material_production"
    "metal_ion_binding"
    "optimal_ph"
    "optimal_temperature"
    "peptide_hla_mhc_affinity"
    "solubility_prediction"
    "ssp_q3"
    "ssp_q8"
    "stability_prediction"
    "tcr_pmhc_affinity"
)

if [ -n "$1" ] && [ -n "$2" ]; then
    CURRENT_TASK=$1
    CURRENT_MODEL=$2
    echo "Using arguments passed"
else
    echo "ERROR: task and model must be passed as arguments" >&2
    exit 1
fi

echo "Task: $CURRENT_TASK"
echo "Model: $CURRENT_MODEL"

# facebook/* models are always loaded from the hub
if [[ "$CURRENT_MODEL" == facebook/* ]]; then
    FROM_HUB=1
fi

cd ${EVAL_ROOT:-.}
module load python/3.10 cuda/12.6.0
source $HOME/AMPLIFY/.venv/bin/activate

if [ "$FROM_HUB" -eq 1 ]; then
    # For hub runs, prepend ${HF_ORG:-your-hf-org}/ unless the model already has an owner prefix
    if [[ "$CURRENT_MODEL" == */* ]]; then
        HUB_MODEL="$CURRENT_MODEL"
    else
        HUB_MODEL="${HF_ORG:-your-hf-org}/$CURRENT_MODEL"
    fi
    # HF repo names use underscores instead of dots (e.g. 0.50 -> 0_50)
    HUB_MODEL="${HUB_MODEL//./_}"
    echo "Running with model from Hugging Face Hub: $HUB_MODEL"
    python scripts/main.py \
        data_config=$CURRENT_TASK \
        model=$HUB_MODEL \
        grid=mlp_feature_extraction_extended \
        grid.subbenchmark_name=mlp_feature_extraction \
        head_type=mlp \
        do_test=true \
        return_all_results=true \
        base_folder=$SCRATCH/proseqo/apperitif \
        +precomputing_batch_size=4 \
        verbose=true
else
    echo "Running with local model: $CURRENT_MODEL"
    python scripts/main.py \
        model=amplify_120M_local \
        model.vocab_path=$HOME/AMPLIFY/conf/tokenizer/amplify_vocab.txt \
        model.model_name=$CURRENT_MODEL \
        data_config=$CURRENT_TASK \
        model.base_path=$SCRATCH/proseqo/ablations \
        grid=mlp_feature_extraction_extended \
        grid.subbenchmark_name=mlp_feature_extraction \
        head_type=mlp \
        do_test=true \
        return_all_results=true \
        base_folder=$SCRATCH/proseqo/apperitif \
        +precomputing_batch_size=4 \
        verbose=true
fi

exit_code=$?

if [[ "$exit_code" -ne 0 ]]; then
    echo "Job failed with exit code $exit_code"
    echo "$CURRENT_MODEL, $CURRENT_TASK" >> $SCRATCH/proseqo/apperitif/mlp_feature_extraction/fail.txt
    exit $exit_code
fi

echo "Job completed successfully!"
echo "$CURRENT_MODEL, $CURRENT_TASK" >> $SCRATCH/proseqo/apperitif/mlp_feature_extraction/success.txt
exit $exit_code
```
