## Supplementary material for "pLM representations unlock metagenomic space beyond homology": Code bundle: paper_figures.ipynb

```
# %% [markdown]
## Load results

# %% [code]
# Run once to (re-)generate plot_df.csv
# Add --skip-aggregation to skip the upstream aggregation scripts
!python scripts/prepare_plot_df.py

# %% [code]
import pandas as pd
import matplotlib.pyplot as plt
import seaborn as sns
import os
import json

# %% [code]
plot_df = pd.read_csv("data/plot_df.csv")
# Boolean column is read back as bool
plot_df["1m"] = plot_df["1m"].astype(bool)

metric_to_ylabel = {
    "Calibration (NT)": "Avg. calibration",
    "Calibration (T)": "Avg. calibration",
    "Deep mutational scanning": "Avg. Spearman",
    "Categorical jacobian (CASP16)": "ROC-AUC",
    "Property (FE, ext.)": "Avg. test score (*)",
    "Property (FE, rest.)": "Avg. test score (*)",
}

metric_display_names = {
    "Calibration (NT)": "Calibration (NT)",
    "Calibration (T)": "Calibration (T)",
    "Deep mutational scanning": "Deep mutational scanning",
    "Categorical jacobian (CASP16)": "Structure (CASP16)",
    "Property (FE, ext.)": "Property (supervised)",
    "Property (FE, rest.)": "Property (supervised)",
}

# %% [code]
plot_df

# %% [code]
plot_df[plot_df.model_name == "AMPLIFY_350M"]

# %% [code]
models_for_si = [model for model in plot_df.model_name.unique() if( "MILA" in model and not ("low_steps" in model or "0.0_mila" in model))] + ["AMPLIFY_120M", "AMPLIFY_350M", "AMPLIFY_benchmark", "AMPLIFY_consensus"]

tmp = plot_df[["model_name", "1m", "filtering", "KL homologs (mean)", "Calibration (T)", "Calibration (NT)", "Categorical jacobian (CASP16)", "Deep mutational scanning", "Property (FE, ext.)", "Property (FE, rest.)"]]
tmp = tmp[tmp.model_name.isin(models_for_si)]

# Drop rows where any of the metric columns is NA
tmp = tmp.dropna(subset=["KL homologs (mean)", "Calibration (T)", "Calibration (NT)", "Categorical jacobian (CASP16)", "Deep mutational scanning", "Property (FE, ext.)", "Property (FE, rest.)"], how="any")

# Sort models by name
tmp = tmp.sort_values(by="model_name", ascending=False)

# Remove na
tmp.dropna()

tmp.to_csv("data/in_silico_results.csv", index=False)

# %% [code]
tmp

# %% [code]
u100 = 418_829_208
bfd = 2_032_203_748 
mgy = 2_455_358_433

bfd_qt = (100 - 46)
mgy_qt = (100 - 45)
u100_qt = (100 - 16)

u100 = u100 * (u100_qt / 100)
bfd = bfd * (bfd_qt / 100)
mgy = mgy * (mgy_qt / 100)

for sub in [0.03, 0.05, 0.10, 0.20, 0.30, 0.40]:
    count = u100 + bfd * sub + mgy * sub
    print(f"Subsample {sub:.0%}: {count:.0f} total sequences")
    print(f"U100: {u100:.0f} sequences")
    print(f"BFD: {bfd*sub:.0f} sequences")
    print(f"MGNIFY: {mgy*sub:.0f} sequences")

# %% [code]
for model in ["AMPLIFY_consensus", "AMPLIFY_benchmark"]:
    for task in ["Calibration (NT)", "Categorical jacobian (CASP16)", "Deep mutational scanning"]:
        x = tmp[tmp.model_name == model][task].values[0]
        y = tmp[tmp.model_name == "AMPLIFY_350M"][task].values[0]
        rel = (x - y) / abs(y)
        print(f"{model} vs AMPLIFY_350M on {task}: {rel:.2%} relative difference")

# %% [code]
for model in ["AMPLIFY_consensus", "AMPLIFY_benchmark"]:
    for task in ["Calibration (NT)", "Categorical jacobian (CASP16)", "Deep mutational scanning"]:
        x = tmp[tmp.model_name == model][task].values[0]
        y = tmp[tmp.model_name == "AMPLIFY_350M"][task].values[0]
        abs = (x - y)
        print(f"{model} vs AMPLIFY_350M on {task}: {abs:.3f} difference")

# %% [code]
tmp = pd.read_csv("data/in_silico_results.csv")


# %% [code]
tmp2 = plot_df[["model_name", "1m", "filtering"] + [col for col in plot_df.columns if ("(FE," in col or "Pct Completed" in col)]]
tmp2 = tmp2[tmp2.model_name.isin(tmp.model_name.unique())]
tmp2.sort_values(by=["model_name"], inplace=True)

# Add NA values to columns ending in (FE, extended) for rows where the column "Pct Completed Extended (FE)" is 450

for col in tmp2.columns:
    if col.endswith("(FE, Extended)") or col.endswith("(FE, ext.)"):
        tmp2.loc[tmp2["Pct Completed Extended (FE)"] == 450, col] = pd.NA
        
tmp2 = tmp2[['model_name', '1m', 'filtering', 'Antibiotic Resistance (FE, Restricted)',
       'Cloning Clf (FE, Restricted)',
       'Enzyme Catalytic Efficiency (FE, Restricted)',
       'Fitness (FE, Restricted)', 'Fluorescence (FE, Restricted)',
       'Fold (FE, Restricted)', 'Localization (FE, Restricted)',
       'Material Production (FE, Restricted)',
       'Metal Ion Binding (FE, Restricted)', 'Optimal pH (FE, Restricted)',
       'Optimal Temperature (FE, Restricted)',
       'Peptide HLA MHC Affinity (FE, Restricted)',
       'Solubility (FE, Restricted)', 'SSP Q3 (FE, Restricted)',
       'SSP Q8 (FE, Restricted)', 'Stability (FE, Restricted)',
       'TCR pMHC Affinity (FE, Restricted)',
       'Antibiotic Resistance (FE, Extended)', 'Cloning Clf (FE, Extended)',
       'Enzyme Catalytic Efficiency (FE, Extended)', 'Fitness (FE, Extended)',
       'Fluorescence (FE, Extended)', 'Fold (FE, Extended)',
       'Localization (FE, Extended)', 'Material Production (FE, Extended)',
       'Metal Ion Binding (FE, Extended)', 'Optimal pH (FE, Extended)',
       'Optimal Temperature (FE, Extended)',
       'Peptide HLA MHC Affinity (FE, Extended)', 'Solubility (FE, Extended)',
       'SSP Q3 (FE, Extended)', 'SSP Q8 (FE, Extended)',
       'Stability (FE, Extended)', 'TCR pMHC Affinity (FE, Extended)']]

# Sort columns alphabetically, keeping the first three columns in place
tmp2 = tmp2[tmp2.columns[:3].tolist() + sorted(tmp2.columns[3:])]
tmp2.to_csv("data/property_prediction_results.csv", index=False)

# %% [code]
tmp3 = pd.read_csv("$SCRATCH/proseqo/categorical_jacobian/categorical_jacobian.csv")
tmp3["model_name"] = tmp3["model_name"].apply(lambda x: x.replace("MILA_MGNIFY", "MILA_MGnify").replace("AMPLIFY_consensus_extended", "AMPLIFY_consensus"))
tmp3.drop(columns=["repo_name"], inplace=True)
tmp3 = tmp3[tmp3.model_name.isin(tmp.model_name.unique())]
tmp3.to_csv("data/categorical_jacobian_results.csv", index=False)

# %% [code]
# Read json files
dico = {}
for dataset in ["bfd", "mgnify", "uniref"]:
    with open(f"data/kde_boundaries_{dataset}.json", "r") as f:
        tmp = json.load(f)
        dico[dataset] = tmp

# %% [code]
dico

# %% [code]
# Flatten dico in df with columns dataset, boundary, qt, and save as csv
df = []
for dataset, item in dico.items():
    df.append({'dataset': dataset, 'boundary': item["boundary"], 'qt': item["qt"]})
df = pd.DataFrame(df)

# %% [code]
df

# %% [code]
df["red_boundary"] = 1 - df["boundary"]

# %% [code]
df = df[["dataset", "red_boundary", "qt"]]
df.to_csv("data/kde_boundaries.csv", index=False)

# %% [code]


# %% [code]
plt.rcParams.update(
    {
        "font.family": "sans-serif",
        "font.sans-serif": ["Helvetica", "Arial", "DejaVu Sans"],
        "font.size": 10,
        "axes.labelsize": 12,
        "axes.titlesize": 12,
        "xtick.labelsize": 8,
        "ytick.labelsize": 8,
        "figure.dpi": 100,
        "axes.linewidth": 0.8,
        "xtick.major.width": 0.8,
        "ytick.major.width": 0.8,
    }
)

# %% [code]
def create_gridspec_axes(n_items, ncols=4, figsize_per_item=(3.0, 3.0), subplotspec=None, hspace=None, wspace=None, shared_x=False, shared_y=False):
    """Create gridspec and axes for a grid of plots. Returns (fig, axes, standalone)."""
    n_rows = n_items // ncols + (n_items % ncols > 0)
    if subplotspec is None:
        fig = plt.figure(figsize=(figsize_per_item[0] * ncols, figsize_per_item[1] * n_rows))
        gs = fig.add_gridspec(n_rows, ncols, hspace=hspace, wspace=wspace)
        standalone = True
    else:
        gs = subplotspec.subgridspec(n_rows, ncols, hspace=hspace, wspace=wspace)
        fig = subplotspec.get_topmost_subplotspec().get_gridspec().figure
        standalone = False
    axes = np.array([fig.add_subplot(gs[j // ncols, j % ncols]) for j in range(n_items)])
    if shared_x:
        for ax in axes[1:]:
            ax.sharex(axes[0])
    if shared_y:
        for ax in axes[1:]:
            ax.sharey(axes[0])
    
    return fig, axes, standalone


def finalize_plot(fig, standalone, filename=None, tight_layout=True, dpi=300):
    """Finalize a standalone plot: layout, save PDF+PNG, show."""
    if standalone:
        if tight_layout:
            fig.tight_layout()
        if filename:
            fig.savefig(f"paper_figures/{filename}.pdf", format="pdf", bbox_inches="tight", dpi=dpi)
            fig.savefig(f"paper_figures/{filename}.png", format="png", bbox_inches="tight", dpi=dpi)
        plt.show()


def despine(ax, spines=("top", "right")):
    """Remove specified spines from axis."""
    for spine in spines:
        ax.spines[spine].set_visible(False)

# %% [code]
def plot_all_metrics_histogram(
    df,
    models_to_plot,
    colors,
    metrics_to_plot,
    group_order=None,
    filename=None,
    metric_ranges=None,
    alpha=1,
    baseline_model=None,
    to_hatch="UniRef100",
    ncol=3,
    legend_bbox_to_anchor=(0.5, 1.07),
    shift_ylim=True,
    zero_line_proportion=0.8,
    sharey=False,
    legend_ncol=5,
    fontsize=10,
    subplotspec=None,
):
    relative = baseline_model is not None
    n_rows = math.ceil(len(metrics_to_plot) / ncol)
    bar_width = 0.4 if relative else 0.6
    group_gap = 0.8

    if group_order is None:
        group_order = [list(models_to_plot.keys())]

    # Build ordered bar metadata: (x_pos, display_name, color, is_hatched)
    bars, current_x = [], 0
    for group in group_order:
        for name in group:
            if name != baseline_model:
                bars.append((current_x, name, colors.get(name, "gray"), ((to_hatch is not None) and (to_hatch in name))))
                current_x += bar_width
        current_x += group_gap - bar_width

    # Note: sharey not supported in embedded mode
    if subplotspec is None:
        fig, axes = plt.subplots(n_rows, ncol, figsize=(4.0 * ncol, 3 * n_rows), sharey=sharey)
        standalone = True
        axes = axes.flatten()
    else:
        fig, axes, standalone = create_gridspec_axes(
            n_rows * ncol, ncols=ncol, figsize_per_item=(4.0, 3.0), subplotspec=subplotspec
        )

    df_plot = df[df["model_name"].isin(models_to_plot.keys())].copy()
    df_plot["display_name"] = df_plot["model_name"].map(models_to_plot)
    bar_kw = dict(width=bar_width, linewidth=1.2, capsize=3, alpha=alpha, edgecolor="white")

    shift_ylim = (not sharey) and shift_ylim

    for i, metric in enumerate(metrics_to_plot):
        ax = axes[i]
        means = df_plot.groupby("display_name")[metric].mean()
        if relative:
            baseline = means[baseline_model]
            heights = {n: (means.get(n, 0) - baseline) / baseline * 100 for _, n, _, _ in bars}
        else:
            heights = {n: means.get(n, 0) for _, n, _, _ in bars}

        # Plot bars in two passes (hatched then regular)
        for hatched in (True, False):
            subset = [(x, n, c) for x, n, c, h in bars if h == hatched]
            if not subset:
                continue
            xs, names, clrs = zip(*subset)
            extra = {"hatch": "///"} if hatched else {}
            ax.bar(xs, [heights[n] for n in names], color=clrs, **bar_kw, **extra)

        if not sharey and metric_ranges and metric in metric_ranges:
            ax.set_ylim(metric_ranges[metric])

        despine(ax)
        if relative:
            # Remove bottom spine
            ax.spines["bottom"].set_visible(False)
            ax.axhline(0, color="black", linestyle="-", linewidth=1.5)

            # Make sure the zero line is always at the same position across plots and that full bars are visible
            zero_line_proportion = zero_line_proportion
            current_ylim = ax.get_ylim()

            if shift_ylim:
                zero_line_y = current_ylim[0] + zero_line_proportion * (current_ylim[1] - current_ylim[0])
                shift = zero_line_y - 0

                current_ylim = (current_ylim[0] - shift, current_ylim[1] - shift)

            while current_ylim[1] < 1.1 * max(heights.values()) or current_ylim[0] > 1.1 * min(heights.values()):
                if current_ylim[0] == 0:
                    current_ylim = (current_ylim[0] - 1, current_ylim[1])
                if current_ylim[1] == 0:
                    current_ylim = (current_ylim[0], current_ylim[1] + 1)
                current_ylim = (current_ylim[0] * 1.05, current_ylim[1] * 1.05)

            if not sharey:
                ax.set_ylim(*current_ylim)

        elif not metric_ranges and not sharey:
            current_ylim = ax.get_ylim()
            current_ylim = (current_ylim[0], current_ylim[1] * 1.1)
            ax.set_ylim(*current_ylim)

        y_range = ax.get_ylim()[1] - ax.get_ylim()[0]
        for x, name, _, _ in bars:
            h = heights[name]
            if not np.isnan(h):
                y_text = (0.06 * y_range if h < 0 else -0.06 * y_range) if relative else h + 0.05 * y_range
                ax.text(x, y_text, f"{h:.1f}", ha="center", va="center", color="black", fontsize=fontsize)

        ax.set_title(metric_display_names[metric.replace("_", " ")], fontweight="bold", fontsize=fontsize + 2)  # , pad=15)

        ax.set_xticks([])
        ax.yaxis.set_major_locator(MaxNLocator(nbins=5, integer=True))
        ax.yaxis.grid(True, linestyle="--", which="major", color="grey", alpha=0.7)
        ax.set_ylabel(metric_to_ylabel.get(metric, metric), fontsize=fontsize)

        # sns.despine(ax=ax)

    for j in range(len(metrics_to_plot), len(axes)):
        axes[j].axis("off")

    # Legend
    handles = [
        mpatches.Patch(facecolor=c, label=n, edgecolor="white", **({"hatch": "///"} if h else {}))
        for _, n, c, h in bars
    ]
    # Sort by group order and then alphabetically
    order = [item for sublist in group_order for item in sublist]
    handles.sort(key=lambda x: order.index(x.get_label()) if x.get_label() in order else float('inf'))
    if relative:
        handles.insert(0, mlines.Line2D([], [], color="gray", linestyle="--", label=f"{baseline_model} (baseline)"))

    if standalone:
        fig.legend(handles=handles, loc="upper center", frameon=False, ncol=legend_ncol, bbox_to_anchor=legend_bbox_to_anchor, fontsize=fontsize)
    else:
        # Embedded: center legend over the full panel.
        # legend_bbox_to_anchor[1] is treated as a panel-relative y offset (e.g. 1.08 = 8% above panel top).
        # Convert to figure coordinates using the panel's bounding box.
        ss = subplotspec.get_topmost_subplotspec()
        panel_bbox = ss.get_position(fig)
        center_x = (panel_bbox.x0 + panel_bbox.x1) / 2
        # Map panel-relative y (where 1.0 = panel top) into figure coordinates
        panel_height = panel_bbox.y1 - panel_bbox.y0
        legend_y_fig = panel_bbox.y0 + legend_bbox_to_anchor[1] * panel_height
        fig.legend(
            handles=handles,
            loc="upper center",
            frameon=False,
            ncol=legend_ncol,
            bbox_to_anchor=(center_x, legend_y_fig),
            bbox_transform=fig.transFigure,
            fontsize=fontsize,
        )
    if relative:
        ax = axes[0]
        ax.set_ylabel("Relative Difference (%)", fontsize=fontsize)
        for ax in axes[1:]:
            ax.set_ylabel("")
        # fig.text(0.07, 0.5, "Relative Difference (%)", va="center", rotation="vertical", fontsize=10)

    finalize_plot(fig, standalone, filename=filename)

# %% [markdown]
## Scores

# %% [code]
path1 = "$SCRATCH/proseqo/scores"

dataset_folders = {
    "bfd_1M_no_ambig": "BFD",
    "mgnify_1M_no_ambig": "MGnify",
    "uniref100_1M_no_ambig": "UniRef",
}

model_folders = {
    "AMPLIFY_120M_UR50": "AMPLIFY 120M",
    "esm2_flash_35M": "ESM2 35M",
    "esm2_flash_150M": "ESM2 150M",
    "esm2_flash_650M": "ESM2 650M",
    "MILA_BFD": "MILA BFD",
    "MILA_MGNIFY": "MILA MGnify",
    "MILA_MGnify_no_ambig": "MILA MGnify (no ambig)",
    "MILA_UR100_MGnify_no_ambig": "MILA UR100+MGnify",
}

score_cols = ["mean_cosine_sim", "pseudo_ppl"]

frames = []

# Load scores from parquet files
for model_folder, model_name in model_folders.items():
    model_path = Path(path1) / model_folder / "multi-combined-scores"
    if not model_path.exists():
        print(f"Missing: {model_path}")
        continue
    for ds_folder, ds_name in dataset_folders.items():
        ds_path = model_path / ds_folder
        parts = sorted(ds_path.glob("*.parquet")) if ds_path.exists() else []
        if not parts:
            continue
        df = pd.concat(
            [pd.read_parquet(p, columns=score_cols) for p in parts],
            ignore_index=True,
        )
        df["model"] = model_name
        df["dataset"] = ds_name
        frames.append(df)

# Load AMPLIFY 350M scores from CSV files (original scores used in the ridge plot)
amplify_350m_csvs = {
    "BFD": "data/bfd_scores_1M.csv",
    "MGnify": "data/mgnify_scores_1M.csv",
    "UniRef": "data/u100_scores_1M.csv",
}
for ds_name, csv_path in amplify_350m_csvs.items():
    df = pd.read_csv(csv_path)[["mean_cosine_sim"]]
    df["model"] = "AMPLIFY 350M"
    df["dataset"] = ds_name
    frames.append(df)

scores = pd.concat(frames, ignore_index=True)

# KDE boundaries for the ridge plot
with open("data/kde_boundaries_bfd.json") as f:
    kde_boundaries_bfd = json.load(f)
with open("data/kde_boundaries_uniref.json") as f:
    kde_boundaries_uniref = json.load(f)
with open("data/kde_boundaries_mgnify.json") as f:
    kde_boundaries_mgnify = json.load(f)

datasets = {
    "BFD": {
        "scores": scores[(scores["model"] == "AMPLIFY 350M") & (scores["dataset"] == "BFD")].reset_index(drop=True),
        "boundary": kde_boundaries_bfd["boundary"],
        "xlim": (0.3, 1),
        "qt": kde_boundaries_bfd["qt"],
    },
    "MGnify": {
        "scores": scores[(scores["model"] == "AMPLIFY 350M") & (scores["dataset"] == "MGnify")].reset_index(drop=True),
        "boundary": kde_boundaries_mgnify["boundary"],
        "xlim": (0.3, 1),
        "qt": kde_boundaries_mgnify["qt"],
    },
    "UniRef": {
        "scores": scores[(scores["model"] == "AMPLIFY 350M") & (scores["dataset"] == "UniRef")].reset_index(drop=True),
        "boundary": kde_boundaries_uniref["boundary"],
        "xlim": (0, 1),
        "qt": kde_boundaries_uniref["qt"],
    },
}

print(scores.groupby(["model", "dataset"]).size().unstack())

# %% [code]
avg_boundary = np.mean([datasets[ds]["boundary"] for ds in datasets])
print(f"Average boundary across datasets: {avg_boundary:.4f}")

# %% [code]
def plot_red_ridge(datasets, fontsize=10):
    """
    Ridge plot of RED score distributions across datasets.
    Uses seaborn FacetGrid which always creates its own figure.
    """
    dataset_colors = {
        "UniRef": (0.4044212904941176, 0.5346434904470588, 0.9320019126352941),
        "MGnify": (0.8846434386941177, 0.41001709788235297, 0.32250654924705885),
        "BFD": "#FFAC1C",
    }
    dataset_order = ["BFD", "MGnify", "UniRef"]  # bottom to top
    x_boundary = 1 - 0.885

    # Build a combined dataframe for FacetGrid
    ridge_frames = []
    for name in dataset_order:
        df = datasets[name]["scores"][["mean_cosine_sim"]].copy()
        df["RED Score"] = 1 - df["mean_cosine_sim"]
        df["Dataset"] = name
        ridge_frames.append(df)
    ridge_df = pd.concat(ridge_frames, ignore_index=True)
    ridge_df["Dataset"] = pd.Categorical(ridge_df["Dataset"], categories=dataset_order, ordered=True)

    sns.set_style("white", rc={"axes.facecolor": (0, 0, 0, 0)})

    g = sns.FacetGrid(
        ridge_df,
        row="Dataset",
        hue="Dataset",
        palette=dataset_colors,
        aspect=4,
        height=2,
    )

    g.map(
        sns.histplot,
        "RED Score",
        kde=False,
        kde_kws={"gridsize": 100},
        line_kws={"linewidth": 2},
        alpha=1,
        bins=50,
        linewidth=1,
        edgecolor="white",
        binrange=(0, 1),
    )

    g.figure.subplots_adjust(hspace=-0.75)
    g.set_titles("")
    g.set(yticks=[], ylabel="", xlim=(0, 1))
    g.despine(left=True)
    for ax in g.axes.flat:
        ax.spines["bottom"].set_linewidth(1.5)

    ax = g.axes[-1][0]
    ax.spines["left"].set_visible(True)
    ax.spines["left"].set_linewidth(1.5)
    ax.set_xlabel("RED Score", fontsize=fontsize + 2)
    ax.set_ylabel("Count", fontsize=fontsize + 2, color="black")

    # Add perspective: shift upper subplots progressively to the left
    n_rows = len(g.axes)
    x_shift = -0.05
    for i, ax_row in enumerate(g.axes):
        ax = ax_row[0]
        pos = ax.get_position()
        offset = (n_rows - 1 - i) * x_shift
        ax.set_position([pos.x0 - offset, pos.y0, pos.width, pos.height])

    g.figure.canvas.draw()

    floor_top_margin = 0.0
    floor_bottom_margin = 0.0
    x_ticks = np.arange(0.05, 1.0, 0.05)
    ax_top = g.axes[0][0]
    ax_bot = g.axes[-1][0]
    y_top = ax_top.get_position().y0
    y_bot = ax_bot.get_position().y0
    for xt in x_ticks:
        x_t = g.figure.transFigure.inverted().transform(ax_top.transData.transform((xt, 0)))[0]
        x_b = g.figure.transFigure.inverted().transform(ax_bot.transData.transform((xt, 0)))[0]
        dx = x_b - x_t
        dy = y_bot - y_top
        t_top = floor_top_margin / abs(dy)
        t_bot = floor_bottom_margin / abs(dy)
        line = mlines.Line2D(
            [x_t - t_top * dx, x_b + t_bot * dx],
            [y_top - t_top * dy, y_bot + t_bot * dy],
            transform=g.figure.transFigure,
            color="grey",
            linestyle=(0, (10, 5)),
            linewidth=0.5,
            alpha=0.5,
            clip_on=False,
            zorder=0,
        )
        g.figure.add_artist(line)

    x_top = g.figure.transFigure.inverted().transform(ax_top.transData.transform((x_boundary, 0)))[0]
    x_bot = g.figure.transFigure.inverted().transform(ax_bot.transData.transform((x_boundary, 0)))[0]
    top_margin = 0.15
    bottom_margin = 0.05
    dx = x_bot - x_top
    dy = y_bot - y_top
    top_t = top_margin / abs(dy)
    bottom_t = bottom_margin / abs(dy)
    line = mlines.Line2D(
        [x_top - top_t * dx, x_bot + bottom_t * dx],
        [y_top - top_t * dy, y_bot + bottom_t * dy],
        transform=g.figure.transFigure,
        color="black",
        linestyle="--",
        linewidth=1.5,
        alpha=1,
        clip_on=False,
        zorder=0,
    )
    g.figure.add_artist(line)

    g.figure.text(
        x_bot + (bottom_t + 0.05) * dx,
        y_bot + (bottom_t + 0.05) * dy,
        f"KDE boundary",  # = {x_boundary:.2f}
        fontsize=fontsize,
        fontweight="bold",
        color="black",
        ha="center",
        va="center",
    )

    label_offset = 0.0
    # Use the bottom (widest/rightmost) axis as the fixed x anchor for all labels
    fixed_label_x = g.axes[-1][0].get_position().x1 + label_offset
    for ax_row, name in zip(g.axes, dataset_order):
        ax = ax_row[0]
        pos = ax.get_position()
        y_label = pos.y0 + (pos.y1 - pos.y0) * 0.05
        g.figure.text(
            fixed_label_x,
            y_label,
            name,
            fontsize=fontsize + 2,
            fontweight="bold",
            ha="right",
            va="center",
            bbox=dict(facecolor="white", edgecolor="none", pad=3),
        )

    plt.savefig("paper_figures/red_ridge_plot.pdf", format="pdf", bbox_inches="tight", dpi=300)
    plt.savefig("paper_figures/red_ridge_plot.png", format="png", bbox_inches="tight", dpi=300)
    plt.show()


plot_red_ridge(datasets, fontsize=12)

# %% [markdown]
## Baselines

# %% [code]
group_order_baselines = [
    ["UniRef100"],
    ["ALL"],
    [
        "BFD + UniRef100",
        "BFD",
    ],
    [
        "MGnify + UniRef100",
        "MGnify",
    ],
]

models_to_plot_baselines = {
    "MILA_U100_0.0_mila": "UniRef100",
    "MILA_U50_low_steps": "U50",
    "MILA_UR100_BFD_no_ambig": "BFD + UniRef100",
    "MILA_BFD_no_ambig": "BFD",
    "MILA_UR100_MGnify_no_ambig": "MGnify + UniRef100",
    "MILA_MGnify_no_ambig": "MGnify",
    "MILA_UR100_BFD_MGnify_no_ambig": "ALL",
}

ds_colors_baselines = {
    model: color
    for (model, color) in zip(models_to_plot_baselines.values(), sns.color_palette("coolwarm", n_colors=len(models_to_plot_baselines) - 1))
}

ds_colors_baselines = {
    "MGnify": list(ds_colors_baselines.values())[-1],
    "MGnify + UniRef100": list(ds_colors_baselines.values())[-1],
    "BFD": list(ds_colors_baselines.values())[3],
    "BFD + UniRef100": list(ds_colors_baselines.values())[3],
    "UniRef100": list(ds_colors_baselines.values())[0],
    "ALL": sns.color_palette("coolwarm", n_colors=100)[0],
}

metrics_to_plot_baselines = [
    "Calibration (T)",
    "Calibration (NT)",
    "Deep mutational scanning",
    "Categorical jacobian (CASP16)",
    "Property (FE, ext.)",
]

# plot_all_metrics_histogram(
#     plot_df, models_to_plot_baselines, ds_colors_baselines, metrics_to_plot_baselines,
#     group_order_baselines, ncol=4, legend_ncol=6, baseline_model="UniRef100",
#     legend_bbox_to_anchor=(0.5, 1.08), filename="baselines_all_metrics_relative",
# )

# %% [code]
metric_ranges_final = {
    "Calibration (T)": (0, 40),
    "Calibration (NT)": (0, 40),
    "Deep mutational scanning": (35, 42),
    "Categorical jacobian (CASP16)": (15, 38),
    "Property (FE, ext.)": (65, 85),
    "Property (FE, rest.)": (65, 85),
}

colors_final = {
    "esm_650M": "#31a354",
    "esm_150M": "#a1d99b",
    "AMPLIFY$_{v1}$": "#1f77b4",
    "AMPLIFY$_{benchmark}$": "#50C878",
    "AMPLIFY$_{consensus}$": "#FFAC1C",
}

group_order_final = [
    # ["esm_150M", "esm_650M"],
    ["AMPLIFY$_{v1}$"],
    ["AMPLIFY$_{benchmark}$"],
    ["AMPLIFY$_{consensus}$"],
]

models_to_plot_final = {
    "esm_650M": "esm_650M",
    "esm_150M": "esm_150M",
    "AMPLIFY_350M": "AMPLIFY$_{v1}$",
    "AMPLIFY_benchmark": "AMPLIFY$_{benchmark}$",
    "AMPLIFY_consensus": "AMPLIFY$_{consensus}$",
}

# plot_all_metrics_histogram(
#     plot_df, models_to_plot_final, colors_final, metric_ranges_final.keys(),
#     group_order_final, metric_ranges=metric_ranges_final, ncol=4,
#     legend_bbox_to_anchor=(0.5, 1.1), filename="final_results_histogram",
# )

# %% [code]

def plot_baselines(
    df,
    models_to_plot,
    colors,
    metrics_to_plot,
    group_order=None,
    filename=None,
    metric_ranges=None,
    alpha=1,
    baseline_model=None,
    to_hatch="+ UniRef100",
    ncol=3,
    legend_bbox_to_anchor=(0.5, 1.07),
    sharey=False,
    legend_ncol=5,
    fontsize=10,
    subplotspec=None,
):
    n_rows = math.ceil(len(metrics_to_plot) / ncol)
    bar_width = 0.6
    group_gap = 0.8

    if group_order is None:
        group_order = [list(models_to_plot.keys())]

    # Build ordered bar metadata: (x_pos, display_name, color, is_hatched)
    bars, current_x = [], 0
    for group in group_order:
        for name in group:
            bars.append((current_x, name, colors.get(name, "gray"), ((to_hatch is not None) and (to_hatch in name))))
            current_x += bar_width
        current_x += group_gap - bar_width

    if subplotspec is None:
        fig, axes = plt.subplots(n_rows, ncol, figsize=(4.0 * ncol, 3 * n_rows), sharey=sharey)
        standalone = True
        axes = axes.flatten()
    else:
        fig, axes, standalone = create_gridspec_axes(n_rows * ncol, ncols=ncol, figsize_per_item=(4.0, 3.0), subplotspec=subplotspec)

    df_plot = df[df["model_name"].isin(models_to_plot.keys())].copy()
    df_plot["display_name"] = df_plot["model_name"].map(models_to_plot)
    bar_kw = dict(width=bar_width, linewidth=1.2, capsize=3, alpha=alpha, edgecolor="white", zorder=3)

    for i, metric in enumerate(metrics_to_plot):
        ax = axes[i]
        means = df_plot.groupby("display_name")[metric].mean()
        heights = {n: means.get(n, 0) for _, n, _, _ in bars}
        baseline_val = means.get(baseline_model, None) if baseline_model else None

        for hatched in (True, False):
            subset = [(x, n, c) for x, n, c, h in bars if h == hatched]
            if not subset:
                continue
            xs, names, clrs = zip(*subset)
            extra = {"hatch": "///"} if hatched else {}
            ax.bar(xs, [heights[n] for n in names], color=clrs, **bar_kw, **extra)

        if not sharey and metric_ranges and metric in metric_ranges:
            ax.set_ylim(metric_ranges[metric])
        elif not metric_ranges and not sharey:
            lo, hi = ax.get_ylim()
            ax.set_ylim(lo, hi * 1.1)

        despine(ax)

        y_range = ax.get_ylim()[1] - ax.get_ylim()[0]
        for x, name, _, _ in bars:
            h = heights[name]
            if np.isnan(h):
                continue
            if baseline_val is not None and name != baseline_model:
                label = f"{h - baseline_val:+.1f}"
            else:
                label = f"{h:.1f}"
            ax.text(x, h + 0.03 * y_range, label, ha="center", va="center", color="black", fontsize=fontsize)

        ax.set_title(metric_display_names[metric.replace("_", " ")], fontweight="bold", fontsize=fontsize + 2)
        ax.set_xticks([])
        ax.yaxis.set_major_locator(MaxNLocator(nbins=5, integer=True))
        ax.yaxis.grid(True, linestyle="--", which="major", color="grey", alpha=0.7, zorder=0)
        ax.set_ylabel(metric_to_ylabel[metric], fontsize=fontsize)

        # Augment ylim
        ylim = ax.get_ylim()
        ax.set_ylim(ylim[0], ylim[1] + 0.15 * (ylim[1] - ylim[0]))

    for j in range(len(metrics_to_plot), len(axes)):
        axes[j].axis("off")

    handles = [mpatches.Patch(facecolor=c, label=n, edgecolor="white", **({"hatch": "///"} if h else {})) for _, n, c, h in bars]
    order = [item for sublist in group_order for item in sublist]
    handles.sort(key=lambda x: order.index(x.get_label()) if x.get_label() in order else float("inf"))

    if standalone:
        fig.legend(
            handles=handles, loc="upper center", frameon=False, ncol=legend_ncol, bbox_to_anchor=legend_bbox_to_anchor, fontsize=fontsize
        )
    else:
        ss = subplotspec.get_topmost_subplotspec()
        panel_bbox = ss.get_position(fig)
        center_x = (panel_bbox.x0 + panel_bbox.x1) / 2
        panel_height = panel_bbox.y1 - panel_bbox.y0
        legend_y_fig = panel_bbox.y0 + legend_bbox_to_anchor[1] * panel_height
        fig.legend(
            handles=handles,
            loc="upper center",
            frameon=False,
            ncol=legend_ncol,
            bbox_to_anchor=(center_x, legend_y_fig),
            bbox_transform=fig.transFigure,
            fontsize=fontsize,
        )

    finalize_plot(fig, standalone, filename=filename)

plt.rcParams["hatch.linewidth"] = 2.0

metric_ranges = {
    "Calibration (T)": (0, 34),
    "Calibration (NT)": (0, 34),
}

plot_baselines(
    plot_df,
    models_to_plot_baselines,
    ds_colors_baselines,
    metrics_to_plot_baselines,
    group_order_baselines,
    metric_ranges=metric_ranges,
    ncol=5,
    legend_ncol=6,
    baseline_model="UniRef100",
    legend_bbox_to_anchor=(0.5, 0.0),
    subplotspec=None,
    fontsize=10,
    # filename="baselines_all_metrics_relative",
)

# %% [markdown]
## Clustering

# %% [code]
clust_stats = {
    "MILA_U100_clust30": (49_769_155,14_999_675_123,301.4, 49_499),
    "MILA_U100_clust40": (54_339_116,16_904_987_507,311.1, 49_499),
    "MILA_U100_clust50": (65_847_724,21_188_545_095,321.8, 49_499),
    "MILA_U100_clust60": (81_461_435,27_176_245_527,333.6, 49_499),
    "MILA_U100_clust70": (104_962_993,36_075_707_536,343.7, 49_499),
    "MILA_U100_clust80": (141_071_243,50_162_181_521,355.6, 49_499),
    "MILA_U100_clust90": (199_268_886,74_397_981_972,373.4, 49_499),
}

# Number of epochs
num_steps = 100_000
batch_size = 128 * 4 * 8
num_epochs = {}
for clust_name, (num_seqs, num_tokens, seq_len, num_clusters) in clust_stats.items():
    num_epochs[clust_name] = num_steps * batch_size / clust_stats[clust_name][0]

cluster_names_default = list(range(30, 100, 10))
sequence_counts_default = [clust_stats["MILA_U100_clust" + str(name)][0] for name in cluster_names_default]

bsz_default = 128 * 4 * 8
steps_default = [2 * clust_stats["MILA_U100_clust" + str(name)][0] / bsz_default for name in cluster_names_default]

num_epochs_per_clust_default = [num_epochs["MILA_U100_clust" + str(name)] for name in cluster_names_default]

# %% [code]
def plot_clustering_seq_count(
    cluster_names=cluster_names_default,
    sequence_counts=sequence_counts_default,
    subplotspec=None,
):
    """
    Bar chart: number of sequences per clustering threshold.
    If subplotspec is None: creates own figure and calls plt.show().
    Returns the axis.
    """
    if subplotspec is None:
        fig, ax = plt.subplots(figsize=(3, 3))
        standalone = True
    else:
        fig = subplotspec.get_topmost_subplotspec().get_gridspec().figure
        ax = fig.add_subplot(subplotspec)
        standalone = False

    ax.yaxis.grid(True, linestyle="--", which="major", color="grey", alpha=0.7)
    ax.bar(cluster_names, sequence_counts, color="#4a90e2", width=8)
    despine(ax)
    ax.yaxis.set_major_locator(MaxNLocator(nbins=5, integer=True))
    ax.set_xlabel("Clustering", fontsize=10)
    ax.set_ylabel("Sequence count", fontsize=10)

    finalize_plot(fig, standalone, filename="clustering_sequence_count")
    return ax


# plot_clustering_seq_count()

# %% [code]
def plot_clustering_steps(
    cluster_names=cluster_names_default,
    steps=steps_default,
    subplotspec=None,
):
    """
    Bar chart: number of training steps for 2 epochs per clustering threshold.
    If subplotspec is None: creates own figure and calls plt.show().
    Returns the axis.
    """
    if subplotspec is None:
        fig, ax = plt.subplots(figsize=(3, 3))
        standalone = True
    else:
        fig = subplotspec.get_topmost_subplotspec().get_gridspec().figure
        ax = fig.add_subplot(subplotspec)
        standalone = False

    ax.yaxis.grid(True, linestyle="--", which="major", color="grey", alpha=0.7)
    ax.bar(cluster_names, steps, color="#4a90e2", width=8)
    despine(ax)
    ax.yaxis.set_major_locator(MaxNLocator(nbins=5, integer=True))
    ax.ticklabel_format(style="sci", axis="y", scilimits=(0, 0))
    ax.set_xlabel("Clustering", fontsize=10)
    ax.set_ylabel("Number of steps", fontsize=10)

    finalize_plot(fig, standalone, filename="clustering_steps_for_2_epochs")
    return ax


# plot_clustering_steps()

# %% [code]
def plot_clustering_epochs(
    cluster_names=cluster_names_default,
    epoch_counts=num_epochs_per_clust_default,
    subplotspec=None,
    fontsize=8,
):
    """
    Bar chart: number of epochs per clustering threshold (fixed steps).
    If subplotspec is None: creates own figure and calls plt.show().
    Returns the axis.
    """
    if subplotspec is None:
        fig, ax = plt.subplots(figsize=(3, 3))
        standalone = True
    else:
        fig = subplotspec.get_topmost_subplotspec().get_gridspec().figure
        ax = fig.add_subplot(subplotspec)
        standalone = False

    ax.yaxis.grid(True, linestyle="--", which="major", color="grey", alpha=0.7)
    ax.bar(cluster_names, epoch_counts, color="orange", width=8)
    despine(ax)
    ax.yaxis.set_major_locator(MaxNLocator(nbins=5, integer=True))
    ax.set_xlabel("Clustering", fontsize=fontsize)
    ax.set_ylabel("Epochs", fontsize=fontsize)

    ax.spines["bottom"].set_linewidth(2)
    ax.spines["left"].set_visible(False)

    finalize_plot(fig, standalone, filename="clustering_epochs")
    return ax


# plot_clustering_epochs(fontsize=8)

# %% [code]
plot_df.model_name.unique()

# %% [code]
plot_df[plot_df.model_name == "MILA_U100_0.01"]

# %% [code]
clust_models = [
    "MILA_U100_clust30",
    "MILA_U100_clust50",
    "MILA_U100_clust60",
    "MILA_U100_clust70",
    "MILA_U100_clust80",
    "MILA_U100_clust90",
    # "MILA_U100_baseline",
    # "MILA_U100_0.0",
    # "MILA_U100_0.0_mila",
]

metrics_clustering = [
    "Calibration (NT)",
    "Deep mutational scanning",
    "Categorical jacobian (CASP16)",
    "Property (FE, rest.)",
]

model_colors_clustering = {
    model: color
    for model, color in zip(clust_models, ["#247add", "#247add", "#247add", "#4a90e2", "#6fa8f3", "#9ecfff", "#cfe4ff", "#e6f0ff"])
}


def plot_clustering_scatter(
    plot_df,
    clust_models=clust_models,
    metrics=metrics_clustering,
    ncols=4,
    subplotspec=None,
    fontsize=10,
):
    """
    Scatter grid: effect of clustering threshold on each metric (fixed steps).
    If subplotspec is None: creates own figure and calls plt.show().
    Returns list of axes.
    """
    from scipy.interpolate import UnivariateSpline

    tmp = plot_df[plot_df["model_name"].isin(clust_models)].copy()
    tmp["clustering"] = tmp["model_name"].apply(lambda x: int(x.split("clust")[-1]) if "clust" in x else 100)
    tmp.sort_values("clustering", inplace=True)
    x_vals = tmp["clustering"].values
    
    n_metrics = len(metrics)
    fig, axes, standalone = create_gridspec_axes(
        n_metrics, ncols=ncols, figsize_per_item=(3.0, 3.0), subplotspec=subplotspec, hspace=0.5
    )

    for j, metric in enumerate(metrics):
        ax = axes[j]
        heights = tmp[metric].values
        if len(heights) == 0:
            continue
        valid_indices = ~np.isnan(heights)
        heights = heights[valid_indices]
        x_vals_valid = x_vals[valid_indices]

        if len(heights) >= 4:
            spline = UnivariateSpline(x_vals_valid, heights, k=min(3, len(heights) - 1), s=len(heights))
            x_fit = np.linspace(min(x_vals_valid), max(x_vals_valid), 100)
            y_fit = spline(x_fit)
            ax.plot(x_fit, y_fit, color="red", linestyle="--", linewidth=0.8)
            residuals = heights - spline(x_vals_valid)
            std_residuals = np.std(residuals)
            ax.fill_between(x_fit, y_fit - std_residuals, y_fit + std_residuals, color="#ffe5e5", alpha=1)

        sns.despine(ax=ax)
        ax.set_title(metric_display_names.get(metric, metric), fontsize=fontsize + 2, pad=15, fontweight="bold")
        ax.yaxis.set_major_locator(MaxNLocator(nbins=5, integer=True))
        ax.yaxis.grid(True, linestyle="--", which="major", color="grey", alpha=0.7)
        ax.scatter(x_vals_valid, heights, alpha=1, color=ds_colors_baselines["UniRef100"], s=20, linewidth=0.5)
        ax.set_ylabel(metric_to_ylabel.get(metric, metric), fontsize=fontsize)

    axes[0].set_xlabel("Clustering", fontsize=fontsize)

    if standalone:
        plt.subplots_adjust(left=0.05)
    finalize_plot(fig, standalone, filename="all_clustering_scatter", dpi=400)
    return axes

plot_clustering_scatter(plot_df)

# %% [code]
clust_models_fixed = [
    "MILA_U100_clust30_fixed_epochs",
    "MILA_U100_clust40_fixed_epochs",
    "MILA_U100_clust50_fixed_epochs",
    "MILA_U100_clust60_fixed_epochs",
    "MILA_U100_clust70_fixed_epochs",
    "MILA_U100_clust80_fixed_epochs",
    "MILA_U100_clust90_fixed_epochs",
]

metrics_clustering = [
    "Calibration (NT)",
    "Deep mutational scanning",
    "Categorical jacobian (CASP16)",
    "Property (FE, rest.)",
]

model_colors_clustering_fixed = {
    model: color
    for model, color in zip(clust_models_fixed, ["#247add", "#247add", "#247add", "#4a90e2", "#6fa8f3", "#9ecfff", "#cfe4ff", "#e6f0ff"])
}


def plot_clustering_scatter_fixed_epochs(
    plot_df,
    clust_models=clust_models_fixed,
    metrics=metrics_clustering,
    model_colors=model_colors_clustering_fixed,
    ncols=4,
    subplotspec=None,
):
    """
    Scatter grid: effect of clustering threshold (fixed epochs) on each metric.
    If subplotspec is None: creates own figure and calls plt.show().
    Returns list of axes.
    """
    tmp = plot_df[plot_df["model_name"].isin(clust_models)].copy()
    tmp["clustering"] = tmp["model_name"].apply(lambda x: int(x.split("clust")[-1].split("_")[0]) if "clust" in x else 100)
    tmp.sort_values("clustering", inplace=True)
    x_vals = tmp["clustering"].values

    n_metrics = len(metrics)
    fig, axes, standalone = create_gridspec_axes(
        n_metrics, ncols=ncols, figsize_per_item=(3.0, 3.0), subplotspec=subplotspec, hspace=0.5
    )

    for j, metric in enumerate(metrics):
        ax = axes[j]
        heights = tmp[metric].values
        if len(heights) == 0:
            continue
        valid_indices = ~np.isnan(heights)
        heights = heights[valid_indices]
        x_vals_valid = x_vals[valid_indices]

        ax.scatter(x_vals_valid, heights, alpha=1, color=[model_colors[model] for model in tmp["model_name"].iloc[valid_indices]], s=20, linewidth=0.5)

        if len(heights) >= 3:
            coeffs = np.polyfit(x_vals_valid, heights, deg=2)
            poly_func = np.poly1d(coeffs)
            x_fit = np.linspace(min(x_vals_valid), max(x_vals_valid), 100)
            y_fit = poly_func(x_fit)
            ax.plot(x_fit, y_fit, color="red", linestyle="--", linewidth=0.8)
            residuals = heights - poly_func(x_vals_valid)
            std_residuals = np.std(residuals)
            ax.fill_between(x_fit, y_fit - std_residuals, y_fit + std_residuals, color="red", alpha=0.1)

        sns.despine(ax=ax)
        ax.set_title(metric.replace("_", " "), fontsize=10, pad=15, fontweight="bold")
        ax.yaxis.set_major_locator(MaxNLocator(nbins=5, integer=True))

    # fig.text(0.5, -0.02, "Clustering", ha="center", fontsize=10)

    if standalone:
        plt.subplots_adjust(left=0.05)
    finalize_plot(fig, standalone, filename="all_clustering_scatter_fixed_epochs", dpi=400)
    return axes


# plot_clustering_scatter_fixed_epochs(plot_df)

# %% [markdown]
## Filtering

# %% [code]
with open("data/kde_boundaries_bfd.json", "r") as f:
    kde_boundaries_bfd = json.load(f)

with open("data/kde_boundaries_uniref.json", "r") as f:
    kde_boundaries_uniref = json.load(f)

with open("data/kde_boundaries_mgnify.json", "r") as f:
    kde_boundaries_mgnify = json.load(f)

dataset_boundaries = {
    "U100": {
        "boundary": kde_boundaries_uniref["boundary"],
        "qt": kde_boundaries_uniref["qt"],
    },
    "BFD": {
        "boundary": kde_boundaries_bfd["boundary"],
        "qt": kde_boundaries_bfd["qt"],
    },
    "MGnify": {
        "boundary": kde_boundaries_mgnify["boundary"],
        "qt": kde_boundaries_mgnify["qt"],
    },
}

# %% [markdown]
### 100k steps

# %% [code]
filt_to_plot_100k = {
    "U100": [
        # "MILA_U100_0.0",
        # "MILA_U100_0.0_mila",
        "MILA_U100_0.01",
        "MILA_U100_0.05",
        "MILA_U100_0.10",
        "MILA_U100_0.15",
        "MILA_U100_0.20",
        "MILA_U100_0.25",
        "MILA_U100_0.30",
        "MILA_U100_0.40",
        "MILA_U100_0.50",
        "MILA_U100_0.60",
    ],
    "BFD": [
        "MILA_BFD",
        "MILA_BFD_0.10",
        "MILA_BFD_0.20",
        "MILA_BFD_0.30",
        "MILA_BFD_0.40",
        "MILA_BFD_0.50",
        "MILA_BFD_0.60",
        "MILA_BFD_0.70",
        "MILA_BFD_0.80",
        "MILA_BFD_0.90",
    ],
    "MGnify": [
        "MILA_MGnify",
        "MILA_MGnify_0.10",
        "MILA_MGnify_0.20",
        "MILA_MGnify_0.30",
        "MILA_MGnify_0.40",
        "MILA_MGnify_0.50",
        "MILA_MGnify_0.60",
        "MILA_MGnify_0.70",
        "MILA_MGnify_0.80",
    ],
}

unique_datasets_100k = ["U100", "BFD", "MGnify"]

ds_colors_100k = {
    "U100": ds_colors_baselines["UniRef100"],
    "MGnify": ds_colors_baselines["MGnify"],
    "BFD": ds_colors_baselines["BFD"],
}

metrics_to_plot_100k = [
    "Calibration (T)",
    "Calibration (NT)",
    "Deep mutational scanning",
    "Categorical jacobian (CASP16)",
    "Property (FE, ext.)",
]

tmp_100k = plot_df.copy()
tmp_100k = tmp_100k[tmp_100k["model_name"].isin([m for sublist in filt_to_plot_100k.values() for m in sublist])]

# Compute relative score change compared to baseline for each metric
for metric in metrics_to_plot_100k:
    tmp_100k[metric + "_relative"] = 0.0
    for dataset_name in filt_to_plot_100k.keys():
        baseline_mask = (tmp_100k["model_name"].str.contains(dataset_name)) & (
            tmp_100k["filtering"] == tmp_100k[tmp_100k["dataset"] == dataset_name]["filtering"].min()
        )
        baseline_value = tmp_100k[baseline_mask][metric].values[0]

        mask = tmp_100k["dataset"] == dataset_name
        tmp_100k.loc[mask, metric + "_relative"] = ((tmp_100k.loc[mask, metric] - baseline_value) / baseline_value) * 100

red_scores_100k = {}
for ds in ["MGnify", "BFD", "U100"]:
    scores = pd.read_csv(f"data/{ds.lower()}_scores_1M.csv")
    red_scores_100k[ds] = 1 - scores["mean_cosine_sim"].values

# %% [code]
def plot_filtering_100k(
    tmp,
    red_scores,
    ds_colors,
    unique_datasets,
    metrics_to_plot=metrics_to_plot_100k,
    kde_boundary=0.11,
    ncols=4,
    subplotspec=None,
    fontsize=8,
    plot_titles=True,
    legend_bbox_to_anchor=(1.2, 1.1),
    legend_ncol=1,
    subtitle_bbox=(0.5, -0.02),
):
    """
    Scatter grid: effect of 100k-step filtering on each metric (relative improvement).
    If subplotspec is None: creates own figure and calls plt.show().
    Returns list of axes.
    """
    n_metrics = len(metrics_to_plot)

    fig, axes, standalone = create_gridspec_axes(
        n_metrics, ncols=ncols, figsize_per_item=(3.0, 3.0), subplotspec=subplotspec
    )

    for j, metric in enumerate(metrics_to_plot):
        ax = axes[j]
        for i, dataset_name in enumerate(unique_datasets):
            df_ds = tmp[tmp["dataset"] == dataset_name].copy()
            df_ds = df_ds.dropna(subset=[metric])
            if df_ds.empty:
                continue
            df_ds.sort_values(by="filtering", inplace=True)
            percentages = df_ds["filtering"].values
            x_vals = np.array([np.quantile(red_scores[dataset_name], p) for p in percentages])
            heights = df_ds[metric + "_relative"].values
            color = ds_colors.get(dataset_name, "#333333")
            ax.plot(x_vals, heights, alpha=1, color=color)

        ax.spines["bottom"].set_visible(False)
        sns.despine(ax=ax)
        ax.axhline(y=0, color="black", linestyle="-", linewidth=0.5)
        ax.axvline(x=kde_boundary, color="grey", linestyle=":", alpha=1, linewidth=1)
        if plot_titles:
            ax.set_title(metric_display_names[metric.replace("_", " ")], fontweight="bold", fontsize=fontsize + 2)
            
        # Set a minimum y-axis range
        y_min, y_max = ax.get_ylim()
        if y_max - y_min < 20:
            center = (y_max + y_min) / 2
            ax.set_ylim(center - 10, center + 10)
        
        # Set ticks fontsize
        ax.tick_params(axis='both', which='major', labelsize=fontsize)
            
        if j == 0:
            ax.set_ylabel("Relative difference (%)", fontsize=fontsize)
        ax.set_xlabel("Filtering threshold", fontsize=fontsize)

    if standalone:
        plt.subplots_adjust(left=0.05)
    else:
        ss = subplotspec.get_topmost_subplotspec()
        panel_bbox = ss.get_position(fig)
        center_x = (panel_bbox.x0 + panel_bbox.x1) / 2
        panel_height = panel_bbox.y1 - panel_bbox.y0
        legend_y_fig = panel_bbox.y0 + legend_bbox_to_anchor[1] * panel_height
        handles = [mlines.Line2D([0], [0], color=ds_colors[ds], label=ds) for ds in unique_datasets]
        handles.append(mlines.Line2D([0], [0], color="grey", linestyle="--", label="KDE Bound."))
        labels = unique_datasets + ["KDE Bound."]
        fig.legend(
            handles=handles,
            loc="upper center",
            frameon=False,
            ncol=legend_ncol,
            bbox_to_anchor=(center_x, legend_y_fig),
            bbox_transform=fig.transFigure,
            fontsize=fontsize,
        )
    finalize_plot(fig, standalone, filename="all_filtering_scatter", dpi=400)
    return axes


plot_filtering_100k(tmp_100k, red_scores_100k, ds_colors_100k, unique_datasets_100k, metrics_to_plot_100k, ncols=5)

# %% [code]
ds_colors_100k

# %% [code]
ds_colors_baselines

# %% [code]
import statsmodels.api as sm

def plot_filtering_100k_absolute(
    tmp,
    red_scores,
    ds_colors,
    unique_datasets,
    metrics_to_plot=metrics_to_plot_100k,
    kde_boundary=0.11,
    ncols=4,
    subplotspec=None,
    fontsize=8,
    plot_titles=True,
    subtitle_bbox=(0.5, -0.02),
):
    """
    Scatter grid: effect of 100k-step filtering on each metric (relative improvement).
    If subplotspec is None: creates own figure and calls plt.show().
    Returns list of axes.
    """
    n_metrics = len(metrics_to_plot)

    fig, axes, standalone = create_gridspec_axes(n_metrics, ncols=ncols, figsize_per_item=(3.0, 3.0), subplotspec=subplotspec, shared_y=False)

    for j, metric in enumerate(metrics_to_plot):
        ax = axes[j]
        for i, dataset_name in enumerate(unique_datasets):
            df_ds = tmp[tmp["dataset"] == dataset_name].copy()
            df_ds = df_ds.dropna(subset=[metric])
            if df_ds.empty:
                continue
            df_ds.sort_values(by="filtering", inplace=True)
            percentages = df_ds["filtering"].values
            x_vals = percentages # np.array([np.quantile(red_scores[dataset_name], p) for p in percentages])
            heights = df_ds[metric].values
            color = ds_colors.get(dataset_name, "#333333")
            # ax.plot(x_vals, heights, alpha=1, color=color)
            ax.scatter(x_vals, heights, alpha=0.5, color=color, s=10, edgecolors="none")

            # Lowess smoothing with seaborn.regplot
            if len(x_vals) >= 3:
                sns.regplot(
                    x=x_vals,
                    y=heights,
                    # logistic=True,
                    lowess=True,
                    scatter=False,
                    ax=ax,
                    color=color,
                    line_kws={"linestyle": "-", "linewidth": 1},
                    ci=80,
                )

        ax.spines["bottom"].set_visible(False)
        sns.despine(ax=ax)
        # ax.axhline(y=0, color="black", linestyle="-", linewidth=0.5)
        ax.axvline(x=kde_boundary, color="grey", linestyle=":", alpha=1, linewidth=1)
        if plot_titles:
            ax.set_title(metric_display_names[metric.replace("_", " ")], fontweight="bold", fontsize=fontsize + 2)

        # Set ticks fontsize
        ax.tick_params(axis="both", which="major", labelsize=fontsize)
        ax.set_ylabel(metric_to_ylabel.get(metric, metric), fontsize=fontsize)

        # Minimum y range of 10
        y_min, y_max = ax.get_ylim()
        if y_max - y_min < 12:
            center = (y_max + y_min) / 2
            ax.set_ylim(center - 6, center + 6)

        if j == ncols - 1:
            handles = [mlines.Line2D([0], [0], color=ds_colors[ds], label=ds) for ds in unique_datasets]
            handles.append(mlines.Line2D([0], [0], color="grey", linestyle="--", label="KDE Bound."))
            labels = unique_datasets + ["KDE Bound."]
            ax.legend(handles, labels, loc="upper right", fontsize=fontsize, frameon=False, bbox_to_anchor=(1.2, 1))
        # if j == 0:
        #     ax.set_ylabel("Relative difference (%)", fontsize=fontsize)
        ax.set_xlabel("Filtering threshold", fontsize=fontsize)

    if standalone:
        plt.subplots_adjust(left=0.05)
    # finalize_plot(fig, standalone, filename="all_filtering_scatter", dpi=400)
    return axes


# plot_filtering_100k_absolute(tmp_100k, red_scores_100k, ds_colors_100k, unique_datasets_100k, metrics_to_plot_100k)

# %% [markdown]
### 1M steps

# %% [code]
filt_to_plot_1m = {
    "U100": [
        "MILA_U100_0.0",
        "MILA_U100_0.05_1m",
        "MILA_U100_0.10_1m",
        "MILA_U100_0.15_1m",
        "MILA_U100_0.20_1m",
        "MILA_U100_0.25_1m",
        "MILA_U100_0.30_1m",
        "MILA_U100_0.35_1m",
        "MILA_U100_0.40_1m",
    ],
    "BFD": [
        "MILA_BFD_0.20_1m",
        "MILA_BFD_0.30_1m",
        "MILA_BFD_0.40_1m",
        "MILA_BFD_0.50_1m",
        "MILA_BFD_0.60_1m",
        "MILA_BFD_0.70_1m",
        "MILA_BFD_0.80_1m",
    ],
    "MGnify": [
        "MILA_MGnify_0.10_1m",
        "MILA_MGnify_0.20_1m",
        "MILA_MGnify_0.30_1m",
        "MILA_MGnify_0.40_1m",
    ],
}

unique_datasets_1m = ["U100", "BFD", "MGnify"]

ds_colors_1m = {
    "U100": "#19769e",
    "MGnify": "#50C878",
    "BFD": "#FFAC1C",
}

metrics_to_plot_1m = [
    "Calibration (NT)",
    "Deep mutational scanning",
    # "Contact prediction (avg. long P@L)",
    "Categorical jacobian (CASP16)",
    "Property (FE, rest.)",
]

tmp_1m = plot_df.copy()
tmp_1m = tmp_1m[tmp_1m["model_name"].isin([m for sublist in filt_to_plot_1m.values() for m in sublist])]

# Compute relative score change compared to baseline for each metric
for metric in metrics_to_plot_1m:
    tmp_1m[metric + "_relative"] = 0.0
    for dataset_name in filt_to_plot_1m.keys():
        baseline_mask = (tmp_1m["model_name"].str.contains(dataset_name)) & (
            tmp_1m["filtering"] == tmp_1m[tmp_1m["dataset"] == dataset_name]["filtering"].min()
        )
        baseline_value = tmp_1m[baseline_mask][metric].values[0]

        mask = tmp_1m["dataset"] == dataset_name
        tmp_1m.loc[mask, metric + "_relative"] = ((tmp_1m.loc[mask, metric] - baseline_value) / baseline_value) * 100

red_scores_1m = {}
for ds in ["MGnify", "BFD", "U100"]:
    scores = pd.read_csv(f"data/{ds.lower()}_scores_1M.csv")
    red_scores_1m[ds] = 1 - scores["mean_cosine_sim"].values

# %% [code]
def plot_filtering_1m(
    tmp,
    red_scores,
    ds_colors,
    unique_datasets,
    metrics_to_plot=metrics_to_plot_1m,
    kde_boundary=0.11,
    ncols=4,
    subplotspec=None,
):
    """
    Line grid: effect of 1M-step filtering on each metric (relative improvement).
    If subplotspec is None: creates own figure and calls plt.show().
    Returns list of axes.
    """
    n_metrics = len(metrics_to_plot)

    fig, axes, standalone = create_gridspec_axes(
        n_metrics, ncols=ncols, figsize_per_item=(3.0, 3.0), subplotspec=subplotspec
    )

    for j, metric in enumerate(metrics_to_plot):
        ax = axes[j]
        for i, dataset_name in enumerate(unique_datasets):
            df_ds = tmp[tmp["dataset"] == dataset_name].copy()
            df_ds = df_ds.dropna(subset=[metric])
            if df_ds.empty:
                continue
            df_ds.sort_values(by="filtering", inplace=True)
            percentages = df_ds["filtering"].values
            x_vals = np.array([np.quantile(red_scores[dataset_name], p) for p in percentages])
            x_vals[0] = 0.0
            heights = df_ds[metric + "_relative"].values
            color = ds_colors.get(dataset_name, "#333333")
            ax.plot(x_vals, heights, color=color, linewidth=2, alpha=1, zorder=1, label=dataset_name)

        sns.despine(ax=ax)
        ax.axhline(y=0, color="black", linestyle="-", linewidth=0.5)
        ax.axvline(x=kde_boundary, color="grey", linestyle=":", alpha=1, linewidth=1)
        ax.set_title((lambda s: s[0].upper() + s[1:])(metric.replace("_", " ")), fontsize=10, pad=15, fontweight="bold")
        ax.yaxis.set_major_locator(LinearLocator(numticks=5))
        ax.yaxis.set_major_formatter(FormatStrFormatter("%.0f"))

        if j == ncols - 1:
            handles = [mlines.Line2D([0], [0], color=ds_colors[ds], label=ds) for ds in unique_datasets]
            handles.append(mlines.Line2D([0], [0], color="grey", linestyle="--", label="KDE Bound."))
            labels = unique_datasets + ["KDE Bound."]
            ax.legend(handles, labels, loc="upper right", fontsize=8, frameon=False)

    fig.text(-0.02, 0.5, "Relative improvement (%)", va="center", rotation="vertical", fontsize=10)
    fig.text(0.5, -0.02, "Minimal RED score, post-filtering", ha="center", fontsize=10)

    if standalone:
        plt.subplots_adjust(left=0.05)
    finalize_plot(fig, standalone, filename="all_filtering_scatter", dpi=400)
    return axes


# plot_filtering_1m(tmp, metrics_to_plot_1m, red_scores, ds_colors, unique_datasets)

# %% [markdown]
## Subsampling

# %% [code]
opt_models = [model for model in plot_df["model_name"] if "MILA_OPT" in model and not "1m" in model]

dataset_sizes = {
    "UniRef100": 418_829_208,
    "BFD": 2_032_203_748,
    "MGnify": 2_455_358_433,
}

filtering_percentages = {name: datasets[name.replace("UniRef100", "UniRef")]["qt"] / 100 for name in dataset_sizes.keys()}

post_filtering_sizes = {
    ds: int(size * (1 - filtering_percentages[ds])) for ds, size in dataset_sizes.items()
}

ds_colors_pies = {
    "UniRef100": "#49a9d3",
    "MGnify": "#50C878",
    "BFD": "#FFAC1C",
}

# %% [code]
red_models = [model for model in plot_df["model_name"] if "MILA_RED" in model and not "1m" in model]
models_to_plot_subsampling = {model: model.replace("_0.88", "").replace("MILA_RED_", "") for model in red_models}

models_to_plot_subsampling["MILA_U100_0.0"] = "UniRef100"
group_order_subsampling = [list(models_to_plot_subsampling.values())]
ds_colors_subsampling = {
    model: color
    for model, color in zip(models_to_plot_subsampling.values(), sns.color_palette("flare", n_colors=len(models_to_plot_subsampling)))
}

metrics_to_plot_subsampling = [
    "Calibration (NT)",
    "Deep mutational scanning",
    "Categorical jacobian (CASP16)",
    "Property (FE, rest.)",
]

# %% [code]
# Plot stacked bar chart of dataset sizes throughout subsampling, with subsampling of BFD and MGnify varying from 0.03% to 40% (after filtering)
def plot_dataset_composition_bars_normalized(
    post_filtering_sizes=post_filtering_sizes,
    models_to_plot=models_to_plot_subsampling,
    ds_colors=ds_colors_pies,
    subplotspec=None,
    fontsize=8,
    label_color="black",
):
    """Stacked bar chart showing dataset proportions for each subsampling level."""
    if subplotspec is None:
        fig, ax = plt.subplots(figsize=(4, 4))
        standalone = True
    else:
        gs = subplotspec.subgridspec(1, 1)
        fig = subplotspec.get_topmost_subplotspec().get_gridspec().figure
        ax = fig.add_subplot(gs[0, 0])
        standalone = False

    uniref_size = post_filtering_sizes["UniRef100"]
    bfd_filtered = post_filtering_sizes["BFD"]
    mgnify_filtered = post_filtering_sizes["MGnify"]

    labels = []
    uniref_fracs, bfd_fracs, mgnify_fracs = [], [], []

    for model, display_label in models_to_plot.items():
        if display_label == "UniRef100":
            continue
        pct = float(display_label)
        bfd = bfd_filtered * pct
        mgnify = mgnify_filtered * pct
        uniref = uniref_size

        total = uniref + bfd + mgnify
        uniref_fracs.append(uniref / total)
        bfd_fracs.append(bfd / total)
        mgnify_fracs.append(mgnify / total)
        labels.append(display_label)

    x = range(len(labels))
    ax.bar(x, uniref_fracs, label="UniRef100", color=ds_colors["UniRef100"], linewidth=0, edgecolor=None)
    ax.bar(x, bfd_fracs, bottom=uniref_fracs, label="BFD", color=ds_colors["BFD"], linewidth=0, edgecolor=None)
    ax.bar(
        x,
        mgnify_fracs,
        bottom=[u + b for u, b in zip(uniref_fracs, bfd_fracs)],
        label="MGnify",
        color=ds_colors["MGnify"],
        edgecolor=None,
        linewidth=0,
    )
    
    mid = len(labels) // 2
    label_data = [
        ("UniRef100", uniref_fracs[mid] / 2),
        ("BFD", uniref_fracs[mid] + bfd_fracs[mid] / 2),
        ("MGnify", uniref_fracs[mid] + bfd_fracs[mid] + mgnify_fracs[mid] / 2),
    ]
    for name, y_center in label_data:
        txt = ax.text(mid, y_center, name, ha="center", va="center", fontsize=fontsize - 2,
                rotation=90, color=label_color, fontweight="bold")
        txt.set_path_effects([pe.withStroke(linewidth=1, foreground='black')])


    sns.despine(ax=ax)
    # ax.spines["bottom"].set_visible(False)
    ax.spines["left"].set_visible(False)
    ax.tick_params(axis='both', which='major', labelsize=fontsize - 2)
    ax.set_xticks(x)
    ax.set_xticklabels(labels)
    ax.set_yticklabels([])
    ax.set_ylabel("Dataset composition", fontsize=fontsize)
    ax.spines["bottom"].set_linewidth(2)
    # ax.legend(loc="center left", frameon=False, ncol=1, fontsize=fontsize, bbox_to_anchor=(1.05, 0.5),)
    ax.set_ylim(0, 1)
    ax.yaxis.grid(True, linestyle="--", which="major", color="grey", alpha=0.7)

    finalize_plot(fig, standalone, filename="dataset_composition_pie_charts")

    return ax

plot_dataset_composition_bars_normalized(
    post_filtering_sizes=post_filtering_sizes,
    models_to_plot=models_to_plot_subsampling,
    ds_colors=ds_colors_pies,
    label_color="white",
)

# %% [code]
# Plot stacked bar chart of dataset sizes throughout subsampling, with subsampling of BFD and MGnify varying from 0.03% to 40% (after filtering)
def plot_dataset_composition_bars(
    post_filtering_sizes=post_filtering_sizes,
    models_to_plot=models_to_plot_subsampling,
    ds_colors=ds_colors_pies,
    subplotspec=None,
    fontsize=8,
    label_color="black",
):
    """Stacked bar chart showing absolute dataset sizes for each subsampling level."""
    if subplotspec is None:
        fig, ax = plt.subplots(figsize=(4, 4))
        standalone = True
    else:
        gs = subplotspec.subgridspec(1, 1)
        fig = subplotspec.get_topmost_subplotspec().get_gridspec().figure
        ax = fig.add_subplot(gs[0, 0])
        standalone = False

    uniref_size = post_filtering_sizes["UniRef100"]
    bfd_filtered = post_filtering_sizes["BFD"]
    mgnify_filtered = post_filtering_sizes["MGnify"]

    labels = []
    uniref_counts, bfd_counts, mgnify_counts = [], [], []

    for model, display_label in models_to_plot.items():
        if display_label == "UniRef100":
            continue
        pct = float(display_label)
        uniref_counts.append(uniref_size)
        bfd_counts.append(bfd_filtered * pct)
        mgnify_counts.append(mgnify_filtered * pct)
        labels.append(display_label)

    x = range(len(labels))
    ax.bar(x, uniref_counts, label="UniRef100", color=ds_colors["UniRef100"], linewidth=0, edgecolor=None)
    ax.bar(x, bfd_counts, bottom=uniref_counts, label="BFD", color=ds_colors["BFD"], linewidth=0, edgecolor=None)
    ax.bar(
        x,
        mgnify_counts,
        bottom=[u + b for u, b in zip(uniref_counts, bfd_counts)],
        label="MGnify",
        color=ds_colors["MGnify"],
        edgecolor=None,
        linewidth=0,
    )

    mid = len(labels) - 1
    label_data = [
        ("U100", uniref_counts[mid] / 2),
        ("BFD", uniref_counts[mid] + bfd_counts[mid] / 2),
        ("MGnify", uniref_counts[mid] + bfd_counts[mid] + mgnify_counts[mid] / 2),
    ]
    for name, y_center in label_data:
        txt = ax.text(mid, y_center, name, ha="center", va="center", fontsize=fontsize - 2,
                rotation=90, color=label_color, fontweight="bold")
        txt.set_path_effects([pe.withStroke(linewidth=1, foreground='black')])

    sns.despine(ax=ax)
    ax.spines["left"].set_visible(True)
    ax.tick_params(axis='both', which='major', labelsize=fontsize - 2)
    ax.set_xticks(x)
    ax.set_xticklabels(labels)
    ax.set_ylabel("Number of sequences", fontsize=fontsize)
    ax.spines["bottom"].set_linewidth(2)
    ax.yaxis.grid(True, linestyle="--", which="major", color="grey", alpha=0.7)
    ax.yaxis.set_major_formatter(plt.matplotlib.ticker.FuncFormatter(lambda val, _: f"{val/1e9:.1f}B"))

    finalize_plot(fig, standalone, filename="dataset_composition_bars")

    return ax

plot_dataset_composition_bars(
    post_filtering_sizes=post_filtering_sizes,
    models_to_plot=models_to_plot_subsampling,
    ds_colors=ds_colors_pies,
    label_color="white",
)


# %% [code]
post_filtering_sizes

# %% [code]

def plot_dataset_composition_pies(
    dataset_sizes=dataset_sizes,
    post_filtering_sizes=post_filtering_sizes,
    ds_colors=ds_colors_pies,
    subsampling_percentages=(0.5, 0.2, 0.1),
    subplotspec=None,
):
    """
    2x2 grid of pie charts showing dataset composition at different subsampling levels.
    If subplotspec is None: creates own figure and calls plt.show().
    Returns list of axes.
    """
    if subplotspec is None:
        fig, axes = plt.subplots(2, 2, figsize=(4, 4))
        standalone = True
    else:
        gs = subplotspec.subgridspec(2, 2)
        fig = subplotspec.get_topmost_subplotspec().get_gridspec().figure
        axes = np.array([[fig.add_subplot(gs[r, c]) for c in range(2)] for r in range(2)])
        standalone = False

    axes = axes.flatten()

    ax = axes[0]
    ax.pie(dataset_sizes.values(), autopct=None, startangle=90, colors=[ds_colors[ds] for ds in dataset_sizes.keys()], textprops={"fontsize": 8})
    ax.set_title("ALL", fontsize=10)

    for i, subsampling_percentage in enumerate(subsampling_percentages):
        ax = axes[i + 1]
        subsampled_sizes = {"UniRef100": post_filtering_sizes["UniRef100"]} | {
            ds: int(size * subsampling_percentage)
            for ds, size in post_filtering_sizes.items()
            if ds != "UniRef100"
        }
        ax.pie(subsampled_sizes.values(), autopct=None, startangle=90, colors=[ds_colors[ds] for ds in subsampled_sizes.keys()], textprops={"fontsize": 8})
        ax.set_title(f"OPT-{int(subsampling_percentage * 100)}%", fontsize=10)

    handles = [plt.Rectangle((0, 0), 1, 1, color=color) for color in ds_colors.values()]
    fig.legend(handles, ds_colors.keys(), loc="lower center", ncol=3, fontsize=10, bbox_to_anchor=(0.5, -0.02), frameon=False)

    finalize_plot(fig, standalone, filename="dataset_composition_pie_charts")
    return list(axes)


# plot_dataset_composition_pies()

# %% [markdown]
## Randomization

# %% [code]
models_to_plot = {
    "AMPLIFY_120M": "AMPLIFY 120M",
    "AMPLIFY_350M": "AMPLIFY 350M",
    "esm_35M": "ESM 35M",
    "esm_150M": "ESM 150M",
    "esm_650M": "ESM 650M",
}

prop = pd.read_csv("$SCRATCH/proseqo/apperitif/mlp_feature_extraction_aggregated.csv")

task_counts = prop.groupby("model")["task"].nunique().reset_index()
expected_task_count = 17
valid_models = task_counts[task_counts["task"] == expected_task_count]["model"]
prop = prop[prop["model"].isin(valid_models)]
prop[f"best_test_score_extended"] = prop.apply(
    lambda row: row["best_test_f1_extended"] if not pd.isna(row["best_test_f1_extended"]) else row["best_test_spearmanr_extended"], axis=1
)
prop[f"best_test_score_restricted"] = prop.apply(
    lambda row: row["best_test_f1_restricted"] if not pd.isna(row["best_test_f1_restricted"]) else row["best_test_spearmanr_restricted"], axis=1
)

_esm_names = {
    "esm2_t12_35M_UR50D": "esm_35M",
    "esm2_t30_150M_UR50D": "esm_150M",
    "esm2_t33_650M_UR50D": "esm_650M",
    "esm_flash_150M": "esm_150M",
    "esm_flash_650M": "esm_650M",
}
prop["model_name"] = (
    prop["model"].apply(
        lambda x: x.replace("MGNIFY", "MGnify").replace("U50", "UR50").replace("amplify_", "AMPLIFY_").replace("_packed", "")
    )
).apply(lambda x: _esm_names.get(x, x))

randomized = prop[prop["model_name"].str.contains("init") | prop["model_name"].isin(list(models_to_plot.keys()))].copy()
randomized = randomized[["model_name", "task", "best_test_score_restricted"]]

# Compute differences between models and their "init" counterparts in dataframe
diffs = []
for model in models_to_plot.keys():
    model_scores = randomized[randomized["model_name"] == model][["task", "best_test_score_restricted"]].set_index("task")
    init_model = model + "_init"
    if init_model in randomized["model_name"].values:
        init_scores = randomized[randomized["model_name"] == init_model][["task", "best_test_score_restricted"]].set_index("task")
        diff = (model_scores - init_scores).reset_index()
        diff["model_name"] = model
        diffs.append(diff)
differences = pd.concat(diffs, ignore_index=True)
avg_transfer = differences.groupby("task").describe()[["best_test_score_restricted"]].reset_index()

delta_pt = randomized[randomized["model_name"].str.contains("init")].copy()
delta_pt = delta_pt.groupby("task").describe().reset_index()
delta_pt.columns = ["task", "count", "mean", "std", "min", "25%", "50%", "75%", "max"]
avg_transfer.columns = ["task", "count", "mean", "std", "min", "25%", "50%", "75%", "max"]
delta_pt = delta_pt.merge(avg_transfer, on="task", suffixes=("", "_diff")).sort_values("mean_diff", ascending=False).reset_index(drop=True)
delta_pt["task"] = delta_pt["task"].apply(
    lambda x: x.replace("biomap_research_", "")
    .replace("_", " ")
    .title()
    .replace("Ssp", "SSP")
    .replace("Tcr Pmhc", "TCR pMHC")
    .replace("Hla Mhc", "HLA MHC")
    .replace("Prediction", "")
    .strip()
)

# Identify 1m/100k model pairs in plot_df
scaling_pairs = []
for model in plot_df[plot_df["1m"]]["model_name"].unique():
    model_100k = model.replace("_1m", "")
    if model_100k in plot_df["model_name"].values:
        scaling_pairs.append((model_100k, model))

# Per-task (FE, rest.) columns
task_cols = [c for c in plot_df.columns if "(FE, Restricted)" in c]

# Compute per-task differences (1m - 100k) for each model pair
scaling_diffs = []
for model_100k, model_1m in scaling_pairs:
    row_100k = plot_df[plot_df["model_name"] == model_100k][task_cols].iloc[0]
    row_1m = plot_df[plot_df["model_name"] == model_1m][task_cols].iloc[0]
    diff = (row_1m - row_100k).rename("diff").reset_index()
    diff.columns = ["task", "diff"]
    diff["model"] = model_100k
    scaling_diffs.append(diff)

scaling_differences = pd.concat(scaling_diffs, ignore_index=True).dropna()

# Average 100k score per task (x-axis baseline)
scores_100k = plot_df[plot_df["model_name"].isin([p[0] for p in scaling_pairs])][["model_name"] + task_cols]
avg_100k = scores_100k[task_cols].mean().rename("mean_100k").reset_index()
avg_100k.columns = ["task", "mean_100k"]

scores_1m = plot_df[plot_df["model_name"].isin([p[1] for p in scaling_pairs])][["model_name"] + task_cols]
avg_1m = scores_1m[task_cols].mean().rename("mean_1m").reset_index()
avg_1m.columns = ["task", "mean_1m"]

# Average and std of differences per task
avg_diff = scaling_differences.groupby("task")["diff"].agg(mean_diff="mean", std_diff="std").reset_index()

# Merge and clean task names
scaling_tmp = avg_100k.merge(avg_diff, on="task").sort_values("mean_diff", ascending=False).reset_index(drop=True)
scaling_tmp = scaling_tmp.merge(avg_1m, on="task")
scaling_tmp["task"] = scaling_tmp["task"].apply(
    lambda x: x.replace(" (FE, rest.)", "").replace("Tcr Pmhc", "TCR pMHC").replace("Hla Mhc", "HLA MHC")
)

print(f"Number of model pairs: {len(scaling_pairs)}")
print(f"Number of tasks: {len(scaling_tmp)}")

# %% [code]
tasks_to_label_rand = [
    "Stability",
    "Fold",
    "SSP Q8",
    "SSP Q3",
    "Localization",
    "Fluorescence",
    "Optimal Ph",
    "Antibiotic Resistance",
    "TCR pMHC Affinity",
]


def plot_randomization_scatter_bar(
    delta_pt,
    tasks_to_label=tasks_to_label_rand,
    fontsize=10,
    subplotspec=None,
):
    """
    Scatter (randomized score vs delta) + horizontal bar chart of per-task improvements.
    If subplotspec is None: creates own figure and calls plt.show().
    Returns (ax_scatter, ax_bar).
    """
    norm = plt.Normalize(delta_pt["mean_diff"].min(), delta_pt["mean_diff"].max())
    bar_delta_pt = delta_pt.sort_values("mean_diff", ascending=True)
    bar_colors = plt.cm.coolwarm_r(norm(bar_delta_pt["mean_diff"]))

    if subplotspec is None:
        fig, ax = plt.subplots(1, 1, figsize=(6, 6)) #, gridspec_kw={"width_ratios": [1, 1]})
        standalone = True
    else:
        gs = subplotspec.subgridspec(1, 1) #, width_ratios=[1, 1])
        fig = subplotspec.get_topmost_subplotspec().get_gridspec().figure
        ax = fig.add_subplot(gs[0, 0])
        # ax_bar = fig.add_subplot(gs[0, 1])
        standalone = False

    # --- Left: scatter ---
    ax.grid(True, linestyle="--", alpha=0.5)
    sc = ax.scatter(delta_pt["mean"], delta_pt["mean_diff"], cmap="coolwarm_r", c=delta_pt["mean_diff"], norm=norm, s=100, alpha=1)

    slope, intercept, r_value, p_value, std_err = linregress(delta_pt["mean"], delta_pt["mean_diff"])
    x_vals = np.array(ax.get_xlim())
    y_vals = intercept + slope * x_vals
    ax.plot(x_vals, y_vals, color="black", linestyle="-", linewidth=1, alpha=0.3)
    ax.text(x_vals[-1] + 0.02, y_vals[-1] - 0.02, f"R² = {r_value**2:.2f}", fontsize=fontsize, bbox=dict(facecolor="white", alpha=0.5, edgecolor="black", boxstyle="round,pad=0.5"))

    for i, row in delta_pt.iterrows():
        if row["task"] in tasks_to_label and row["task"] not in ("Optimal Ph", "Fluorescence"):
            ax.text(row["mean"] + 0.03, row["mean_diff"] - 0.001, row["task"], fontsize=fontsize, ha="left", va="center")
        elif row["task"] == "Optimal Ph":
            ax.text(row["mean"], row["mean_diff"] - 0.02, row["task"], fontsize=fontsize, ha="center", va="top")
        elif row["task"] == "Fluorescence":
            ax.text(row["mean"], row["mean_diff"] + 0.02, row["task"], fontsize=fontsize, ha="center", va="bottom")

    ax.axhline(y=0, color="black", linewidth=0.5, alpha=1)
    despine(ax, spines=("top", "right", "bottom"))
    ax.set_xlabel("Randomized test score", fontsize=fontsize)
    ax.set_ylabel(r"$\Delta$ (Randomized - 100k)", fontsize=fontsize)
    ax.tick_params(axis="both", labelsize=fontsize)

    # # --- Right: bar ---
    # ax_bar.barh(bar_delta_pt["task"], bar_delta_pt["mean_diff"], xerr=bar_delta_pt["std_diff"], color=bar_colors, edgecolor="none", error_kw={"ecolor": "black", "capsize": 0, "linewidth": 0.8})
    # ax_bar.axvline(x=0, color="black", linewidth=0.5, alpha=1)
    # ax_bar.set_xlabel(r"$\Delta$ Test score", fontsize=fontsize)
    # despine(ax_bar, spines=("top", "right", "left"))
    # ax_bar.yaxis.set_label_position("right")
    # ax_bar.yaxis.tick_right()
    # ax_bar.tick_params(axis="x", labelsize=fontsize)
    # ax_bar.tick_params(axis="y", labelsize=fontsize, length=0)
    # ax_bar.grid(True, axis="x", linestyle="--", alpha=0.5)

    finalize_plot(fig, standalone, filename="test_score_increase_vs_baseline")
    return ax #, ax_bar


plot_randomization_scatter_bar(delta_pt)

# %% [code]
def _clean_task_name(name):
    """Normalize task names for merging across dataframes."""
    return name.replace(" (FE, Restricted)", "").replace("Optimal Ph", "Optimal pH")

# %% [code]
def plot_stacked_performance(
    delta_pt,
    fontsize=10,
    subplotspec=None,
    figsize=(10, 5)
):
    """
    Vertical stacked bar chart: for each task, a grey bar shows the
    randomized (init) performance (delta_pt["mean"] in [0,1] -> scaled to %).
    A hatched transparent bar shows the delta to 1M (scaling_tmp["mean_1m"]
    already in %). When delta > 0 it stacks above; when delta < 0 it hangs below.
    Tasks are sorted by randomized performance descending.
    """
    if subplotspec is None:
        fig, ax = plt.subplots(figsize=figsize)
        standalone = True
    else:
        fig = subplotspec.get_topmost_subplotspec().get_gridspec().figure
        ax = fig.add_subplot(subplotspec)
        standalone = False

    x = np.arange(len(delta_pt))
    bar_w = 0.6

    delta_pt.sort_values("mean", ascending=False, inplace=True)

    # Grey bar: randomized baseline
    ax.bar(
        x,
        delta_pt["mean"].values,
        bar_w,
        label="Randomized",
        color="lightgrey",
        edgecolor="none",
    )

    ax.plot(
        x,
        delta_pt["mean"].values,
        color="black",
        linewidth=1.5,
        linestyle="none",
        marker="_",
        markersize=bar_w * 40,  # 72 pts per inch approx
        markeredgewidth=1.5,
    )

    # Hatched transparent bar: always drawn from min(mean_pct, mean_1m) with height=|delta|
    bar_bottom = np.minimum(delta_pt["mean"].values, delta_pt["mean"].values + delta_pt["mean_diff"].values)
    bar_height = np.abs(delta_pt["mean_diff"].values)

    norm = TwoSlopeNorm(vmin=delta_pt["mean_diff"].min(), vcenter=0.2, vmax=delta_pt["mean_diff"].max())
    bars = ax.bar(
        x,
        bar_height,
        bar_w,
        bottom=bar_bottom,
        label=r"$\Delta$ (Pre-trained $-$ Random)",
        facecolor="lightblue",
        edgecolor="none",
        linewidth=0.8,
    )
    for bar, val in zip(bars, delta_pt["mean_diff"]):
        bar.set_facecolor(plt.cm.coolwarm_r(norm(val)))

    ax.errorbar(
        x,
        delta_pt["mean"].values + delta_pt["mean_diff"].values,
        yerr=delta_pt["std_diff"].values,
        fmt="none",
        ecolor="black",
        elinewidth=1.0,
        capsize=0,
    )

    ax.set_xticks(x)
    ax.set_xticklabels(delta_pt["task"], fontsize=fontsize - 1, rotation=45, ha="right")
    ax.set_ylabel("Test score (%)", fontsize=fontsize)
    ax.legend(frameon=False, fontsize=fontsize - 1, loc="upper right")
    ax.grid(True, axis="y", linestyle="--", alpha=0.5)
    despine(ax, spines=("top", "right"))
    ax.tick_params(axis="x", length=0)
    ax.tick_params(axis="y", labelsize=fontsize - 1)
    ax.spines["bottom"].set_linewidth(2)

    # finalize_plot(fig, standalone, filename="stacked_performance")
    return ax

plot_stacked_performance(delta_pt, figsize=(6, 4))

# %% [code]
delta_pt[["task", "mean", "mean_diff"]].sort_values("mean_diff")

# %% [code]
task_dict = {
    "Stability": "Stability",
    "Fold": "Fold",
    "Fluorescence": "Fluorescence",
    "SSP Q8": "SSP Q8",
    "Optimal Ph": "Opt. Ph",
    "Localization": "Localization",
    "SSP Q3": "SSP Q3",
    "Optimal Temperature": "Opt. Temp.",
    "Enzyme Catalytic Efficiency": "Enzyme Cat. Eff.",
    "Fitness": "Fitness",
    "Metal Ion Binding": "Metal Ion Bind.",
    "Solubility": "Solubility",
    "Cloning Clf": "Cloning Clf",
    "Material Production": "Mat. Prod.",
    "Peptide HLA MHC Affinity": "Peptide HLA MHC",
    "TCR pMHC Affinity": "TCR pMHC",
    "Antibiotic Resistance": "Antibiotic Res.",
}

def plot_stacked_performance_h(
    delta_pt,
    fontsize=10,
    subplotspec=None,
    figsize=(5, 6),
):
    if subplotspec is None:
        fig, ax = plt.subplots(figsize=figsize)
        standalone = True
    else:
        fig = subplotspec.get_topmost_subplotspec().get_gridspec().figure
        ax = fig.add_subplot(subplotspec)
        standalone = False

    delta_pt.sort_values("mean", ascending=True, inplace=True)

    y = np.arange(len(delta_pt))
    bar_w = 0.6

    # Grey bar: randomized baseline
    ax.barh(y, delta_pt["mean"].values, bar_w, label="Randomized", color="lightgrey", edgecolor="none")

    # Dark line on right edge of randomized baseline bar
    ax.plot(
        delta_pt["mean"].values,
        y,
        color="black",
        linewidth=1.5,
        linestyle="none",
        marker="|",
        markersize=bar_w * 20,
        markeredgewidth=1.5,
    )

    # Colored bar: delta
    bar_left = np.minimum(delta_pt["mean"].values, delta_pt["mean"].values + delta_pt["mean_diff"].values)
    bar_width = np.abs(delta_pt["mean_diff"].values)

    norm = TwoSlopeNorm(vmin=delta_pt["mean_diff"].min(), vcenter=0.2, vmax=delta_pt["mean_diff"].max())
    bars = ax.barh(
        y,
        bar_width,
        bar_w,
        left=bar_left,
        label=None,
        facecolor="lightblue",
        edgecolor="none",
        linewidth=0.8,
    )
    for bar, val in zip(bars, delta_pt["mean_diff"]):
        bar.set_facecolor(plt.cm.coolwarm_r(norm(val)))

    ax.errorbar(
        delta_pt["mean"].values + delta_pt["mean_diff"].values,
        y,
        xerr=delta_pt["std_diff"].values,
        fmt="none",
        ecolor="black",
        elinewidth=1.0,
        capsize=0,
    )

    # Add colorbar for delta
    sm = plt.cm.ScalarMappable(cmap="coolwarm_r", norm=norm)
    sm.set_array([])
    cbar = fig.colorbar(sm, ax=ax, orientation="horizontal", pad=0.07) #, fraction=0.02, pad=0.02)
    cbar.set_label(r"$\Delta$ (Pre-trained $-$ Random)", fontsize=fontsize)
    cbar.ax.tick_params(labelsize=fontsize - 1)
    cbar.outline.set_edgecolor(None)

    # Add dark horizontal line at x=0 on the colorbar to indicate the zero point
    cbar.ax.axhline(y=0, color="black", linewidth=1.5)

    ax.set_yticks(y)
    ax.set_yticklabels(delta_pt["task"].map(task_dict), fontsize=fontsize - 1)
    # ax.set_xlabel("Test score", fontsize=fontsize)
    # ax.legend(frameon=False, fontsize=fontsize, bbox_to_anchor=(0.5, -0.2), loc="lower center")
    ax.grid(True, axis="x", linestyle="--", alpha=0.5)
    despine(ax, spines=("top", "right"))
    ax.tick_params(axis="y", length=0)
    ax.tick_params(axis="x", labelsize=fontsize - 1)
    ax.spines["bottom"].set_linewidth(2)
    # ax.spines["left"].set_visible(False)

    # finalize_plot(fig, standalone, filename="stacked_performance_h")
    plt.savefig("paper_figures/stacked_performance_h.png", dpi=400, bbox_inches="tight")
    return ax

plot_stacked_performance_h(delta_pt, figsize=(5, 6), fontsize=12)

# %% [code]
def plot_transfer_dotplot(
    delta_pt,
    scaling_tmp,
    fontsize=10,
    subplotspec=None,
):
    """
    Dot plot with CIs: for each task, show randomized, 100k, and 1M scores
    as dots with error bars on a shared x-axis. All on 0-100 scale.
    A bar chart of delta (pre-trained vs randomized) is shown to the right,
    aligned task-wise (no y-labels on the bar chart).
    """
    rows = []
    for _, r in delta_pt.iterrows():
        key = _clean_task_name(r["task"])
        sc_row = scaling_tmp[scaling_tmp["task"].apply(_clean_task_name) == key]
        if sc_row.empty:
            continue
        sc_row = sc_row.iloc[0]
        rows.append({
            "task": r["task"],
            "randomized": r["mean"] * 100,
            "randomized_std": r["std"] * 100,
            "score_100k": sc_row["mean_100k"],
            "score_1m": sc_row["mean_1m"],
            "mean_diff": r["mean_diff"],
            "std_diff": r["std_diff"],
        })
    df = pd.DataFrame(rows).sort_values("randomized", ascending=True).reset_index(drop=True)

    # --- Create axes: dotplot (left) + barplot (right) ---
    if subplotspec is None:
        fig, (ax, ax_bar) = plt.subplots(
            1, 2, figsize=(12, 6), gridspec_kw={"width_ratios": [3, 1], "wspace": 0.6},
        )
        standalone = True
    else:
        gs = subplotspec.subgridspec(1, 2, width_ratios=[3, 1], wspace=0.4)
        fig = subplotspec.get_topmost_subplotspec().get_gridspec().figure
        ax = fig.add_subplot(gs[0, 0])
        ax_bar = fig.add_subplot(gs[0, 1])
        standalone = False

    y = np.arange(len(df))
    conditions = [
        ("randomized", "Randomized", "#999999", "o"),
        ("score_100k", "Pre-trained (100k)", "#6495ED", "s"),
        ("score_1m", "Pre-trained (1M)", "#2363DB", "D"),
    ]

    for col, label, color, marker in conditions:
        xerr = df["randomized_std"].values if col == "randomized" else None
        ax.errorbar(
            df[col].values, y,
            xerr=xerr, fmt=marker, color=color, markersize=6,
            capsize=2, linewidth=0.8, label=label, alpha=0.9,
        )

    # Connect dots per task with thin lines
    for i, (_, row) in enumerate(df.iterrows()):
        ax.plot(
            [row["randomized"], row["score_100k"], row["score_1m"]],
            [i, i, i],
            color="#cccccc", linewidth=0.6, zorder=0,
        )

    ax.set_yticks(y)
    ax.set_yticklabels(df["task"], fontsize=fontsize)
    ax.set_xlabel("Test score", fontsize=fontsize)
    ax.legend(frameon=False, fontsize=fontsize, loc="upper right")
    ax.grid(True, axis="x", linestyle="--", alpha=0.5)
    despine(ax, spines=("top", "right", "left"))
    ax.tick_params(axis="y", length=0)
    # Put task labels on the right size of the plot, aligned with the bars
    ax.yaxis.set_label_position("right")
    ax.yaxis.tick_right()
    ax.tick_params(axis="x", labelsize=fontsize)
    ax.spines["bottom"].set_linewidth(2)

    df["mean_diff"] = df["mean_diff"] * 100
    df["std_diff"] = df["std_diff"] * 100

    # --- Right: bar chart of delta, aligned by task ---
    # TODO: Move to the right to avoid superposition with the task labels
    norm = plt.Normalize(df["mean_diff"].min(), df["mean_diff"].max())
    bar_colors = plt.cm.coolwarm_r(norm(df["mean_diff"]))

    ax_bar.barh(
        y, df["mean_diff"], xerr=df["std_diff"],
        color=bar_colors, edgecolor="none",
        error_kw={"ecolor": "black", "capsize": 0, "linewidth": 0.8},
    )
    ax_bar.axvline(x=0, color="black", linewidth=0.5, alpha=1)
    ax_bar.set_xlabel(r"$\Delta$ (Randomized - 100k)", fontsize=fontsize)
    despine(ax_bar, spines=("top", "right", "left"))
    ax_bar.set_yticks(y)
    ax_bar.set_yticklabels([])  # no labels — aligned with dotplot
    ax_bar.tick_params(axis="x", labelsize=fontsize)
    ax_bar.tick_params(axis="y", length=0)
    ax_bar.grid(True, axis="x", linestyle="--", alpha=0.5)
    ax_bar.spines["bottom"].set_linewidth(2)

    # Sync y-limits so rows stay aligned
    for a in (ax, ax_bar):
        a.set_ylim(-0.5, len(df) - 0.5)
        a.invert_yaxis()

    finalize_plot(fig, standalone, filename="transfer_dotplot")
    return ax, ax_bar

# plot_transfer_dotplot(delta_pt, scaling_tmp, fontsize=8)

# %% [markdown]
## Scaling training steps

# %% [code]
metrics_to_plot_scaling = [
    "Calibration (T)",
    "Calibration (NT)",
    "Deep mutational scanning",
    "Categorical jacobian (CASP16)",
    "Property (FE, rest.)",
]

# Identify 1m models (ex MILA_ALL_0.60_1m) and their 100k equivalent (ex MILA_ALL_0.60)
model_pairs = []
for model in plot_df[(plot_df["1m"]) & (plot_df["dataset"] != "OPT") & ~((plot_df["dataset"] == "U100") & (plot_df["filtering"] == 0.35))].model_name.unique():
    model_100k = model.replace("_1m", "")
    if model_100k in plot_df.model_name.unique():
        model_pairs.append((model_100k, model))

rows = []
for metric in metrics_to_plot_scaling:
    for m1, m2 in model_pairs:
        score_100k = plot_df[plot_df["model_name"] == m1][metric].values[0]
        score_1m = plot_df[plot_df["model_name"] == m2][metric].values[0]
        if not pd.isna(score_100k) and not pd.isna(score_1m):
            rows.append(
                {
                    "metric": metric,
                    "dataset": plot_df[plot_df["model_name"] == m1]["dataset"].values[0],
                    "filtering": plot_df[plot_df["model_name"] == m1]["filtering"].values[0],
                    "score_100k": score_100k,
                    "score_1m": score_1m,
                    "relative_improvement": (score_1m - score_100k) / abs(score_100k) * 100,
                }
            )

data = pd.DataFrame(rows)
data = data.groupby("metric").filter(lambda g: len(g) >= 2)

ds_colors = {"U100": "#648FFF", "MGnify": "#FE6100", "BFD": "#FFB000", "ALL": "#DC267F"}  # "OPT": "#800080",

# %% [code]
def plot_scaling_grid(
    data,
    ds_colors,
    ncols=5,
    subplotspec=None,
    fontsize=8,
    legend_bbox_to_anchor=(0.5, 0.05),
    plot_legend=True,
):
    """
    Scatter grid: 100k vs 1M score for each metric.
    If subplotspec is None: creates own figure and calls plt.show().
    Returns list of axes.
    """
    metrics = data["metric"].unique()
    n_metrics = len(metrics)

    fig, axes, standalone = create_gridspec_axes(n_metrics, ncols=ncols, figsize_per_item=(3.0, 3.0), subplotspec=subplotspec)

    for i, metric in enumerate(metrics):
        subset = data[data["metric"] == metric]
        ax = axes[i]
        colors = [ds_colors.get(ds.replace("U100", "UniRef100"), "#333333") for ds in subset["dataset"]]
        # sizes = [100 * (1 - f) for f in subset["filtering"]]
        ax.scatter(subset["score_100k"], subset["score_1m"], color=colors, alpha=1, s=80, edgecolor="black", linewidth=0.5)  # s=sizes,
        min_val = min(subset["score_100k"].min(), subset["score_1m"].min()) * 0.98
        max_val = max(subset["score_100k"].max(), subset["score_1m"].max()) * 1.02
        ax.plot([min_val, max_val], [min_val, max_val], color="black", linestyle="-", linewidth=0.8, alpha=1)
        ax.set_title(metric_display_names.get(metric, metric), fontsize=fontsize + 2, fontweight="bold")
        # ax.set_xlim(min_val, max_val)
        # ax.set_ylim(min_val, max_val)
        ax.yaxis.set_major_locator(MaxNLocator(nbins=5, integer=True))
        despine(ax)
        if i % ncols == 0:
            ax.set_ylabel("1M score", fontsize=fontsize)
            ax.set_xlabel("100k score", fontsize=fontsize)

        ax.yaxis.grid(True, linestyle="--", which="major", color="grey", alpha=0.7)

    for j in range(i + 1, len(axes)):
        axes[j].set_visible(False)

    handles = [
        plt.Line2D(
            [0],
            [0],
            marker="o",
            color="w",
            label=ds.replace("U100", "UniRef100"),
            alpha=1,
            markeredgecolor="black",
            markeredgewidth=0.5,
            markerfacecolor=color,
            markersize=8,
        )
        for ds, color in ds_colors.items()
    ]

    if plot_legend:
        fig.legend(handles=handles, loc="lower center", bbox_to_anchor=legend_bbox_to_anchor, fontsize=fontsize, frameon=False, ncol=len(handles))

    finalize_plot(fig, standalone, filename="1m_vs_100k_comparison")
    return axes


plot_scaling_grid(data, ds_colors)


def plot_scaling_dumbbell(
    data,
    ds_colors,
    ds_order=[
        "UniRef100",
        "ALL",
        "BFD",
        "MGnify",
    ],
    ncols=5,
    subplotspec=None,
    fontsize=8,
    legend_bbox_to_anchor=(1.25, 0.5),
    plot_legend=True,
):
    """
    Dumbbell plot: 100k vs 1M score for each metric, grouped by dataset (averaged ± std over filtering).
    Each panel = one metric. Y-axis = datasets. Hollow dot = 100k, filled dot = 1M, connected by line.
    """
    metrics = data["metric"].unique()
    n_metrics = len(metrics)

    agg = (
        data.groupby(["metric", "dataset"])[["score_100k", "score_1m"]]
        .agg(["mean", "std"])
        .reset_index()
    )
    agg.columns = ["metric", "dataset", "mean_100k", "std_100k", "mean_1m", "std_1m"]
    agg["dataset"] = agg["dataset"].replace({"U100": "UniRef100"})

    # Sort agg by ds_order
    agg["dataset"] = pd.Categorical(agg["dataset"], categories=ds_order, ordered=True)
    agg = agg.sort_values("dataset", ascending=False).reset_index(drop=True)

    fig, axes, standalone = create_gridspec_axes(
        n_metrics, ncols=ncols, figsize_per_item=(3.0, 2.2), subplotspec=subplotspec,
        hspace=0.6, wspace=0.4,
    )

    for i, metric in enumerate(metrics):
        ax = axes[i]
        subset = agg[agg["metric"] == metric]
        y = np.arange(len(subset))

        # connecting line
        for j, row in subset.iterrows():
            yi = list(subset.index).index(j)
            ax.plot(
                [row["mean_100k"], row["mean_1m"]], [yi, yi],
                color="#cccccc", linewidth=1.2, zorder=0,
            )

        # 100k: hollow dot
        ax.errorbar(
            subset["mean_100k"], y,
            xerr=subset["std_100k"].fillna(0),
            fmt="o", markersize=6,
            markerfacecolor="white", markeredgewidth=1.2,
            capsize=2, linewidth=0.8, zorder=2,
            color="#666666", ecolor="#aaaaaa",
        )
        # Color the hollow dots individually
        for j, (yi, row) in enumerate(zip(y, subset.itertuples())):
            ax.plot(
                row.mean_100k, yi,
                marker="o", markersize=6,
                markerfacecolor="white",
                markeredgecolor=ds_colors.get(row.dataset, "#333333"),
                markeredgewidth=1.5,
                zorder=3,
            )

        # 1M: filled dot
        for j, (yi, row) in enumerate(zip(y, subset.itertuples())):
            ax.errorbar(
                row.mean_1m, yi,
                xerr=row.std_1m if not np.isnan(row.std_1m) else 0,
                fmt="D", markersize=6,
                markerfacecolor=ds_colors.get(row.dataset, "#333333"),
                markeredgecolor=ds_colors.get(row.dataset, "#333333"),
                markeredgewidth=0.5,
                capsize=2, linewidth=0.8, zorder=4,
                color=ds_colors.get(row.dataset, "#333333"),
                ecolor=ds_colors.get(row.dataset, "#333333"),
            )

        ax.set_yticks(y)
        ax.set_yticklabels(subset["dataset"] if i == 0 else [], fontsize=fontsize)
        ax.set_title(metric_display_names.get(metric, metric), fontsize=fontsize + 1, fontweight="bold")
        ax.tick_params(axis="y", length=0)
        ax.grid(True, axis="x", linestyle="--", alpha=0.5, color="grey")
        despine(ax, spines=("top", "right", "left"))
        ax.tick_params(axis="x", labelsize=fontsize)

        if plot_legend and i == len(metrics) - 1:
            handles = [
                plt.Line2D([0], [0], marker="o", color="w", markerfacecolor="white",
                        markeredgecolor="#666666", markeredgewidth=1.5, markersize=8, label="100k"),
                plt.Line2D([0], [0], marker="D", color="w", markerfacecolor="#666666",
                        markeredgecolor="#666666", markersize=8, label="1M"),
            ]
            ax.legend(handles=handles, loc="center right", bbox_to_anchor=legend_bbox_to_anchor,
                    fontsize=fontsize, frameon=False, ncol=1)

    for j in range(i + 1, len(axes)):
        axes[j].set_visible(False)

    finalize_plot(fig, standalone, filename="1m_vs_100k_dumbbell")
    return axes


plot_scaling_dumbbell(data, ds_colors)

# %% [code]
ds_colors

# %% [code]
# Plot relative improvements as a bar chart per metric
def plot_relative_improvements(
    data,
    ds_colors,
    ncols=5,
    subplotspec=None,
    fontsize=8,
    legend_bbox_to_anchor=(1.05, 0.5),
    plot_legend=True,
    plot_titles=True,
):
    """
    Bar chart grid: relative improvement (delta / 100k) for each metric.
    If subplotspec is None: creates own figure and calls plt.show().
    Returns list of axes.
    """
    metrics = data["metric"].unique()
    n_metrics = len(metrics)

    fig, axes, standalone = create_gridspec_axes(
        n_metrics, ncols=ncols, figsize_per_item=(3.0, 3.0), subplotspec=subplotspec, shared_y=True
    )

    dataset_order = ["MGnify", "BFD", "U100", "ALL"]
    # Order data
    data["dataset"] = pd.Categorical(data["dataset"], categories=dataset_order, ordered=True)
    data.sort_values("dataset", inplace=True)

    for i, metric in enumerate(metrics):
        subset = data[data["metric"] == metric]
        colors = [ds_colors.get(ds.replace("U100", "UniRef100"), "#333333") for ds in subset["dataset"]]
        ax = axes[i]
        ax.bar(range(len(subset)), subset["relative_improvement"], color=colors, alpha=1, edgecolor="black", linewidth=0.5)
        if plot_titles:
            ax.set_title(metric_display_names.get(metric, metric), fontsize=fontsize + 2, fontweight="bold")
        ax.set_xticklabels([])
        despine(ax)
        if i % ncols == 0:
            ax.set_ylabel("Relative improvement", fontsize=fontsize)
        ax.axhline(0, color="black", linestyle="-", linewidth=0.8)
        ax.yaxis.grid(True, linestyle="--", which="major", color="grey", alpha=0.7)
        ax.spines["bottom"].set_visible(False)

    for j in range(i + 1, len(axes)):
        axes[j].set_visible(False)

    handles = [
        plt.Line2D(
            [0],
            [0],
            marker="o",
            color="w",
            label=ds.replace("U100", "UniRef100"),
            alpha=1,
            markeredgecolor="black",
            markeredgewidth=0.5,
            markerfacecolor=color,
            markersize=8,
        )
        for ds, color in ds_colors.items()
    ]

    if plot_legend:
        fig.legend(handles=handles, loc="lower center", bbox_to_anchor=legend_bbox_to_anchor, fontsize=fontsize, frameon=False, ncol=len(ds_colors))

    finalize_plot(fig, standalone, filename="relative_improvements")
    return axes


# plot_relative_improvements(data, ds_colors)

# %% [code]
tmp = data[data["filtering"].isin([0.2, 0.3, 0.4])]

# %% [code]
def plot_relative_improvements_scatter(
    data,
    ds_colors,
    ncols=5,
    subplotspec=None,
    fontsize=8,
    legend_bbox_to_anchor=(1.05, 0.5),
):
    """
    Scatter plot grid: relative improvement (delta / 100k) for each metric,
    with x axis representing filtering level.
    If subplotspec is None: creates own figure and calls plt.show().
    Returns list of axes.
    """
    metrics = data["metric"].unique()
    n_metrics = len(metrics)

    fig, axes, standalone = create_gridspec_axes(
        n_metrics, ncols=ncols, figsize_per_item=(3.0, 3.0), subplotspec=subplotspec, shared_y=True
    )

    for i, metric in enumerate(metrics):
        subset = data[data["metric"] == metric]
        ax = axes[i]
        for _, row in subset.iterrows():
            color = ds_colors.get(row["dataset"], "#333333")
            ax.scatter(row["filtering"], row["relative_improvement"], color=color, alpha=0.7, edgecolors="black", linewidths=0.5, s=40)
        ax.set_title(metric_display_names.get(metric, metric), fontsize=fontsize + 2, fontweight="bold")
        ax.set_xlabel("Filtering", fontsize=fontsize)
        despine(ax)
        if i % ncols == 0:
            ax.set_ylabel("Relative improvement", fontsize=fontsize)
        ax.axhline(0, color="black", linestyle="-", linewidth=0.8)
        ax.yaxis.grid(True, linestyle="--", which="major", color="grey", alpha=0.7)

    for j in range(i + 1, len(axes)):
        axes[j].set_visible(False)

    handles = [
        plt.Line2D(
            [0],
            [0],
            marker="o",
            color="w",
            label=ds.replace("U100", "UniRef100"),
            alpha=0.7,
            markeredgecolor="black",
            markeredgewidth=0.5,
            markerfacecolor=color,
            markersize=8,
        )
        for ds, color in ds_colors.items()
    ]

    fig.legend(handles=handles, loc="center right", bbox_to_anchor=legend_bbox_to_anchor, fontsize=fontsize, frameon=False, ncol=1)

    finalize_plot(fig, standalone, filename="relative_improvements_scatter")
    return axes

# plot_relative_improvements_scatter(data, ds_colors)

# %% [code]
tasks_to_label_scaling = [
    "Stability",
    "Fold",
    "SSP Q8",
    "SSP Q3",
    "Localization",
    "Fluorescence",
    "Optimal pH",
    "Antibiotic Resistance",
    "TCR pMHC Affinity",
    "Peptide HLA MHC Affinity",
]


def plot_scaling_scatter_bar(
    scaling_tmp,
    tasks_to_label=tasks_to_label_scaling,
    fontsize=12,
    subplotspec=None,
):
    """
    Scatter (100k score vs delta) + horizontal bar of per-task 1M-100k improvements.
    If subplotspec is None: creates own figure and calls plt.show().
    Returns (ax_scatter, ax_bar).
    """
    norm_sc = plt.Normalize(scaling_tmp["mean_diff"].min(), scaling_tmp["mean_diff"].max())
    bar_sc = scaling_tmp.sort_values("mean_diff", ascending=True)
    bar_colors_sc = plt.cm.coolwarm_r(norm_sc(bar_sc["mean_diff"]))

    if subplotspec is None:
        fig, (ax, ax_bar) = plt.subplots(1, 2, figsize=(12, 6), gridspec_kw={"width_ratios": [2, 1]})
        standalone = True
    else:
        gs = subplotspec.subgridspec(1, 2, width_ratios=[2, 1])
        fig = subplotspec.get_topmost_subplotspec().get_gridspec().figure
        ax = fig.add_subplot(gs[0, 0])
        ax_bar = fig.add_subplot(gs[0, 1])
        standalone = False

    ax.grid(True, linestyle="--", alpha=0.5)
    sc = ax.scatter(scaling_tmp["mean_100k"], scaling_tmp["mean_diff"], cmap="coolwarm_r", c=scaling_tmp["mean_diff"], norm=norm_sc, s=100, alpha=1)

    for i, row in scaling_tmp.iterrows():
        if row["task"] in tasks_to_label and row["task"] not in ("Optimal pH", "Fluorescence"):
            ax.text(row["mean_100k"] + 1.5, row["mean_diff"] - 0.005, row["task"], fontsize=fontsize, ha="left", va="center")
        elif row["task"] in ("Optimal pH", "Fluorescence"):
            ax.text(row["mean_100k"], row["mean_diff"] - 0.05, row["task"], fontsize=fontsize, ha="center", va="top")

    ax.axhline(y=0, color="black", linewidth=0.5, alpha=1)
    despine(ax, spines=("top", "right", "bottom"))
    ax.set_xlabel("100k test score", fontsize=fontsize)
    ax.set_ylabel(r"$\Delta$ Test score (1M $-$ 100k)", fontsize=fontsize)
    ax.tick_params(axis="both", labelsize=fontsize)

    ax_bar.barh(bar_sc["task"], bar_sc["mean_diff"], xerr=bar_sc["std_diff"], color=bar_colors_sc, edgecolor="none", error_kw={"ecolor": "black", "capsize": 0, "linewidth": 0.8})
    ax_bar.axvline(x=0, color="black", linewidth=0.5, alpha=1)
    ax_bar.set_xlabel(r"$\Delta$ Test score (1M $-$ 100k)", fontsize=fontsize)
    despine(ax_bar, spines=("top", "right", "left"))
    ax_bar.yaxis.set_label_position("right")
    ax_bar.yaxis.tick_right()
    ax_bar.tick_params(axis="x", labelsize=fontsize)
    ax_bar.tick_params(axis="y", labelsize=fontsize, length=0)
    ax_bar.grid(True, axis="x", linestyle="--", alpha=0.5)

    finalize_plot(fig, standalone, filename="test_score_increase_1m_vs_100k")
    return ax, ax_bar


# plot_scaling_scatter_bar(scaling_tmp)

# %% [code]
tasks_to_label_merged = [
    "Stability",
    "Fold",
    "SSP Q8",
    "SSP Q3",
    "Localization",
    "Fluorescence",
    "Optimal pH",
    "Antibiotic Resistance",
    "TCR pMHC Affinity",
    "Peptide HLA MHC Affinity",
]

scaling_tmp["task"] = scaling_tmp["task"].apply(lambda x: x.replace(" (FE, Restricted)", ""))
merged = delta_pt.merge(scaling_tmp[["task", "mean_diff"]], on="task", suffixes=("_vs_random", "_1m_vs_100k"))
merged["mean_diff_vs_random"] = merged["mean_diff_vs_random"] * 100
merged.to_csv("delta_randomized_vs_delta_scaling.csv", index=False)


def plot_randomization_vs_scaling(
    merged,
    tasks_to_label=tasks_to_label_merged,
    subplotspec=None,
):
    """
    Scatter: delta(1M-100k) vs delta(pre-trained - randomized), colored by randomized score.
    If subplotspec is None: creates own figure and calls plt.show().
    Returns the axis.
    """
    if subplotspec is None:
        fig, ax = plt.subplots(figsize=(8, 6))
        standalone = True
    else:
        fig = subplotspec.get_topmost_subplotspec().get_gridspec().figure
        ax = fig.add_subplot(subplotspec)
        standalone = False

    norm = plt.Normalize(merged["mean"].min(), merged["mean"].max())
    ax.scatter(merged["mean_diff_1m_vs_100k"], merged["mean_diff_vs_random"], cmap="viridis_r", c=merged["mean"], s=50, alpha=1, edgecolor=None, label="task")

    cbar = plt.colorbar(plt.cm.ScalarMappable(norm=norm, cmap="viridis_r"), ax=ax)
    cbar.set_label("Randomized test score", fontsize=12)

    ax.xaxis.set_major_formatter(FormatStrFormatter("%.1f"))
    ax.yaxis.set_major_formatter(FormatStrFormatter("%.0f"))
    ax.tick_params(axis="x", which="major", length=0)
    ax.tick_params(axis="y", which="major", length=0)
    despine(ax, spines=("top", "right", "bottom", "left"))
    ax.axhline(y=0, color="black", linewidth=0.5, alpha=1)
    ax.axvline(x=0, color="black", linewidth=0.5, alpha=1)
    ax.set_xlabel(r"$\Delta$ Test score (1M $-$ 100k)", fontsize=12)
    ax.set_ylabel(r"$\Delta$ Test score (Pre-trained $-$ Randomized)", fontsize=12)

    for i, row in merged.iterrows():
        if row["task"] == "TCR pMHC Affinity":
            ax.text(row["mean_diff_1m_vs_100k"], row["mean_diff_vs_random"] - 1.5, row["task"], fontsize=10, ha="center", va="top")
        elif row["task"] in ("Stability", "Localization"):
            ax.text(row["mean_diff_1m_vs_100k"] - 0.09, row["mean_diff_vs_random"], row["task"], fontsize=10, ha="right", va="center")
        elif row["task"] == "Peptide HLA MHC Affinity":
            ax.text(row["mean_diff_1m_vs_100k"] - 0.3, row["mean_diff_vs_random"] - 1.5, row["task"], fontsize=10, ha="left", va="top")
        elif row["task"] in tasks_to_label:
            ax.text(row["mean_diff_1m_vs_100k"] + 0.09, row["mean_diff_vs_random"], row["task"], fontsize=10, ha="left", va="center")

    ax.grid(True, linestyle="--", alpha=0.5)

    finalize_plot(fig, standalone, filename="delta_randomized_vs_delta_scaling", tight_layout=False)
    return ax


# plot_randomization_vs_scaling(merged)

# %% [markdown]
## Consensus fidelity

# %% [code]
consensus = pd.read_csv("$SCRATCH/proseqo/sequence_recovery/consensus_fidelity.csv")
for threshold in [70, 90, 96]:
    consensus[f"cat_{threshold}_difference"] = consensus[f"cat_{threshold}_non_target"] - consensus[f"cat_{threshold}_target"]

local_df = consensus.merge(plot_df[["model_name", "1m", "filtering", "dataset"]], on="model_name", how="left")

# %% [code]
models_to_plot_consensus = {
    "MILA_U100_0.0": "UniRef100",
    "MILA_BFD": "BFD",
    "MILA_MGnify": "MGnify",
    "MILA_UR100_BFD_MGnify_no_ambig": "ALL",
    "MILA_OPT_0.10_sub10": "OPT 10%",
}

model_colors_consensus = {
    "MILA_U100_0.0": "#247add",
    "MILA_U100_clust50": "#49a9d3",
    "MILA_BFD": "#FFAC1C",
    "MILA_MGnify": "#50C878",
    "MILA_UR100_BFD_MGnify_no_ambig": "#19769e",
    "MILA_OPT_0.20_sub20": "#800080",
    "MILA_OPT_0.10_sub10": "#9C3A9C",
}


def plot_consensus_baselines(
    local_df,
    models_to_plot=models_to_plot_consensus,
    model_colors=model_colors_consensus,
    categories=("cat_70", "cat_90", "cat_96"),
    subplotspec=None,
):
    """
    Bar grid: consensus fidelity (target vs non-target) for baseline models.
    If subplotspec is None: creates own figure and calls plt.show().
    Returns list of axes.
    """
    n_cols = len(categories)
    baseline_df = local_df[local_df["model_name"].isin(models_to_plot)].copy()

    if subplotspec is None:
        fig = plt.figure(figsize=(6 * n_cols, 4))
        gs = fig.add_gridspec(1, n_cols)
        standalone = True
    else:
        gs = subplotspec.subgridspec(1, n_cols)
        fig = subplotspec.get_topmost_subplotspec().get_gridspec().figure
        standalone = False

    axes = [fig.add_subplot(gs[0, i]) for i in range(n_cols)]

    for i, cat in enumerate(categories):
        ax = axes[i]
        baseline_df.sort_values(f"{cat}_target", inplace=True, ascending=False)
        bar_width = 1
        gap_between_groups = 0.5

        for j, model in enumerate(models_to_plot):
            model_df = baseline_df[baseline_df["model_name"] == model]
            if model_df.empty:
                continue
            target_score = model_df[f"{cat}_target"].values[0]
            non_target_score = model_df[f"{cat}_non_target"].values[0]
            ax.bar(j * (bar_width + gap_between_groups + 1) - bar_width / 2, target_score, width=bar_width, color=model_colors.get(model, "#333333"), edgecolor="white", alpha=0.7)
            ax.bar(j * (bar_width + gap_between_groups + 1) + bar_width / 2, non_target_score, width=bar_width, color=model_colors.get(model, "#333333"), edgecolor="white", alpha=0.7, hatch="///")

        ax.set_xticks([j * (bar_width + gap_between_groups + 1) for j in range(len(models_to_plot))])
        ax.set_xticklabels(models_to_plot.values(), fontsize=10)
        ax.set_title(f"{cat.split('_')[1]}% Consensus fidelity", fontsize=12, fontweight="bold")
        if i == 0:
            ax.set_ylabel("Accuracy (%)", fontsize=10)
        despine(ax)

        if i == 0:
            handles = [
                mpatches.Patch(facecolor="white", edgecolor="black", label="Target"),
                mpatches.Patch(facecolor="white", edgecolor="black", label="Non-target", hatch="///"),
            ]
            ax.legend(handles=handles, loc="upper center", fontsize=10, frameon=False)

    if standalone:
        plt.subplots_adjust(left=0.05)
    finalize_plot(fig, standalone)
    return axes


plot_consensus_baselines(local_df)

# %% [code]
models_to_plot_consensus_clust = {
    "MILA_U100_clust30": "UniRef30",
    "MILA_U100_clust40": "UniRef40",
    "MILA_U100_clust50": "UniRef50",
    "MILA_U100_clust60": "UniRef60",
    "MILA_U100_clust70": "UniRef70",
    "MILA_U100_clust80": "UniRef80",
    "MILA_U100_clust90": "UniRef90",
}

model_colors_consensus_clust = {
    "MILA_U100_clust30": "#1f77b4",
    "MILA_U100_clust40": "#2ca02c",
    "MILA_U100_clust50": "#ff7f0e",
    "MILA_U100_clust60": "#d62728",
    "MILA_U100_clust70": "#9467bd",
    "MILA_U100_clust80": "#8c564b",
    "MILA_U100_clust90": "#e377c2",
}


def plot_consensus_clustering(
    local_df,
    models_to_plot=models_to_plot_consensus_clust,
    model_colors=model_colors_consensus_clust,
    categories=("cat_70", "cat_90", "cat_96"),
    subplotspec=None,
):
    """
    Line+bar grid: consensus fidelity vs clustering threshold.
    If subplotspec is None: creates own figure and calls plt.show().
    Returns list of axes.
    """
    n_cols = len(categories)
    baseline_df = local_df[local_df["model_name"].isin(models_to_plot)].copy()
    baseline_df.sort_values("model_name", inplace=True, ascending=False)

    if subplotspec is None:
        fig = plt.figure(figsize=(4 * n_cols, 4))
        gs = fig.add_gridspec(1, n_cols)
        standalone = True
    else:
        gs = subplotspec.subgridspec(1, n_cols)
        fig = subplotspec.get_topmost_subplotspec().get_gridspec().figure
        standalone = False

    axes = [fig.add_subplot(gs[0, i]) for i in range(n_cols)]

    for i, cat in enumerate(categories):
        ax = axes[i]
        ax2 = ax.twinx()
        bar_width = 8

        for j, model in enumerate(models_to_plot):
            model_df = baseline_df[baseline_df["model_name"] == model]
            if model_df.empty:
                continue
            target_score = model_df[f"{cat}_target"].values[0]
            non_target_score = model_df[f"{cat}_non_target"].values[0]
            ax2.bar(int(model.split("_clust")[-1]), non_target_score - target_score, width=bar_width, edgecolor="white", alpha=0.5, color="#cfe4ff")

        ax.plot(baseline_df["model_name"].apply(lambda x: int(x.split("_clust")[-1])), baseline_df[f"{cat}_target"], marker="o", color="#247add", alpha=0.7, label="Target")
        ax.plot(baseline_df["model_name"].apply(lambda x: int(x.split("_clust")[-1])), baseline_df[f"{cat}_non_target"], marker="o", color="#F98B03", alpha=0.7, label="Non-target")
        ax2.axhline(y=0, color="black", linewidth=0.5, alpha=1)
        ax2.set_ylim(-16, 8)

        if i == 2:
            ax2.set_ylabel("Difference", fontsize=10)

        ax.set_title(f"{cat.split('_')[1]}% Consensus fidelity", fontsize=12, fontweight="bold")
        if i == 0:
            ax.set_ylabel("Accuracy (%)", fontsize=10)
            ax.legend(loc="upper center", fontsize=10, frameon=False)
        if i == 1:
            ax.set_xlabel("Clustering threshold", fontsize=10)
        despine(ax)

    if standalone:
        plt.subplots_adjust(left=0.05)
    finalize_plot(fig, standalone)
    return axes

plot_consensus_clustering(local_df)

# %% [markdown]
## Figure panels

Each cell below assembles a multi-panel publication figure.
Edit the `MOSAIC` string and `FIGSIZE` at the top of each cell to reposition panels.

# %% [code]
from matplotlib.transforms import ScaledTranslation


def add_part_labels(axd, offset_x=-8, offset_y=8, fontsize=12):
    """Add bold A/B/C part labels (journal style) to axes in a mosaic dict."""
    for label, ax in axd.items():
        trans = ax.transAxes + ScaledTranslation(offset_x / 72, offset_y / 72, ax.figure.dpi_scale_trans)
        ax.text(0, 1, label, transform=trans, fontsize=fontsize, fontweight="bold", va="top", ha="right")

# %% [code]
from PIL import Image
import matplotlib.pyplot as plt
import matplotlib.image as mpimg


def add_image_panel(ax, filepath):
    """Display an image file in a matplotlib axes."""
    img = mpimg.imread(filepath)
    ax.imshow(img)
    ax.margins(0)
    ax.axis("off")
    ax.set_anchor("W")

# %% [markdown]
### Figure 1

# %% [code]
# ── Layout ────────────────────────────────────────────────────────────────────
FIGSIZE = (18, 14)
MOSAIC = """
AABBBB
CCDDDD
EEEEEE
"""
# ──────────────────────────────────────────────────────────────────────────────

fig = plt.figure(figsize=FIGSIZE)
axd = fig.subplot_mosaic(MOSAIC, gridspec_kw={"hspace": 0, "wspace": 0, "height_ratios": [0.9, 0.9, 1], "width_ratios": [1, 1, 1, 1, 1, 1]})
for ax in axd.values():
    ax.set_axis_off()

# Manual panels loaded from file
# add_image_panel(axd["C"], "paper_figures/panel_1c.png")
# add_image_panel(axd["E"], "paper_figures/panel_1e.png")

add_part_labels(axd, offset_x=-10, offset_y=10, fontsize=12)
# plt.savefig("paper_figures/panel_1.png", bbox_inches="tight", dpi=300)
# plt.savefig("paper_figures/panel_1.pdf", bbox_inches="tight", dpi=300)
plt.show()

# %% [markdown]
### Figure 2

# %% [code]
group_order_baselines = [['UniRef100'],
 ['BFD', 'BFD + UniRef100'],
 ['MGnify', 'MGnify + UniRef100'],
 ['ALL'],]

# %% [code]
# ── Layout ────────────────────────────────────────────────────────────────────
FIGSIZE = (18, 18)
MOSAIC = """
AABBBBB
CCDDDEE
FFFFFFF
GGGGGGG
"""
# ──────────────────────────────────────────────────────────────────────────────

fig = plt.figure(figsize=FIGSIZE)
axd = fig.subplot_mosaic(
    MOSAIC, gridspec_kw={"hspace": 0.4, "wspace": 0.2, "height_ratios": [1.4, 1.2, 1, 1], "width_ratios": [1.2, 1.2, 0.8, 0.8, 0.8, 1, 1]}
)
for ax in axd.values():
    ax.set_axis_off()

# add_image_panel(axd["A"], "paper_figures/fig2b-pipeline-v2.png")
# add_image_panel(axd["B"], "paper_figures/U100_RED_vs_PPL.png")
# add_image_panel(axd["C"], "paper_figures/proseqo-mcs_umap.png")
# add_image_panel(axd["D"], "paper_figures/cosine_sim_ridge_plot.png")

ds_colors_100k["BFD"] = "#FFAC1C"
ds_colors_baselines["BFD"] = "#FFAC1C"
ds_colors_baselines["BFD + UniRef100"] = "#FFAC1C"

plot_baselines(
    plot_df,
    models_to_plot_baselines,
    ds_colors_baselines,
    metrics_to_plot_baselines,
    group_order_baselines,
    ncol=5,
    legend_ncol=6,
    baseline_model="UniRef100",
    legend_bbox_to_anchor=(0.5, 0.0),
    filename="baselines_all_metrics_relative",
    subplotspec=axd["F"].get_subplotspec(),
    fontsize=10,
)

plot_filtering_100k(tmp_100k, red_scores_100k, ds_colors_100k, unique_datasets_100k, ncols=5, subtitle_bbox=(0.5, 0.08), fontsize=10, plot_titles=True,
                    legend_bbox_to_anchor=(0.5, -0.2),
                    legend_ncol=5,
                    subplotspec=axd["G"].get_subplotspec())


add_part_labels(axd, offset_x=-20, offset_y=20, fontsize=12)
# plt.savefig("paper_figures/panel_filtering.png", bbox_inches="tight", dpi=300)
# plt.savefig("paper_figures/panel_filtering.pdf", bbox_inches="tight", dpi=300)
plt.show()

# %% [code]
# ── Layout ────────────────────────────────────────────────────────────────────
FIGSIZE = (18, 18)
MOSAIC = """
AABBBBB
CCDDDEE
FFFFFFF
GGGGGGG
"""
# ──────────────────────────────────────────────────────────────────────────────

fig = plt.figure(figsize=FIGSIZE)
axd = fig.subplot_mosaic(
    MOSAIC, gridspec_kw={"hspace": 0.4, "wspace": 0.2, "height_ratios": [1.4, 1.2, 1, 1], "width_ratios": [1.2, 1.2, 0.8, 0.8, 0.8, 1, 1]}
)
for ax in axd.values():
    ax.set_axis_off()

# add_image_panel(axd["A"], "paper_figures/fig2b-pipeline-v2.png")
# add_image_panel(axd["B"], "paper_figures/U100_RED_vs_PPL.png")
# add_image_panel(axd["C"], "paper_figures/proseqo-mcs_umap.png")
# add_image_panel(axd["D"], "paper_figures/cosine_sim_ridge_plot.png")

ds_colors_100k["BFD"] = "#FFAC1C"
ds_colors_baselines["BFD"] = "#FFAC1C"
ds_colors_baselines["BFD + UniRef100"] = "#FFAC1C"

plot_baselines(
    plot_df,
    models_to_plot_baselines,
    ds_colors_baselines,
    metrics_to_plot_baselines,
    group_order_baselines,
    ncol=5,
    legend_ncol=6,
    baseline_model="UniRef100",
    legend_bbox_to_anchor=(0.5, 0.0),
    filename="baselines_all_metrics_relative",
    subplotspec=axd["F"].get_subplotspec(),
    fontsize=10,
)

plot_filtering_100k(tmp_100k, red_scores_100k, ds_colors_100k, unique_datasets_100k, ncols=5, subtitle_bbox=(0.5, 0.08), fontsize=10, plot_titles=True,
                    legend_bbox_to_anchor=(0.5, -0.2),
                    legend_ncol=5,
                    subplotspec=axd["G"].get_subplotspec())


add_part_labels(axd, offset_x=-20, offset_y=20, fontsize=12)
# plt.savefig("paper_figures/panel_filtering.png", bbox_inches="tight", dpi=300)
# plt.savefig("paper_figures/panel_filtering.pdf", bbox_inches="tight", dpi=300)
plt.show()

# %% [markdown]
### Figure 3

# %% [code]
ds_colors_subsampling

# %% [code]
# ── Layout ────────────────────────────────────────────────────────────────────
FIGSIZE = (18, 14)
MOSAIC = """
BBBBBB
CCCCCC
DDDDDE
FFFFFG
"""
# ──────────────────────────────────────────────────────────────────────────────

tmp = data[(data["dataset"] != "BFD") | (data["filtering"].isin([0.2, 0.3, 0.4]))]
# tmp = data[(data["dataset"] != "U100") | (data["filtering"].isin([0.2, 0.3, 0.4]))]
tmp = tmp[(tmp["dataset"] != "ALL") | (tmp["filtering"].isin([0.6, 0.7, 0.8]))]

metric_order = [
    "Calibration (T)",
    "Calibration (NT)",
    "Deep mutational scanning",
    "Categorical jacobian (CASP16)",
    "Property (FE, rest.)",
]

tmp["metric"] = pd.Categorical(tmp["metric"], categories=metric_order, ordered=True)
tmp.sort_values("metric", inplace=True)

ds_colors_subsampling = {
    model: color
    for (model, color) in zip(
        models_to_plot_subsampling.values(), sns.color_palette("coolwarm", n_colors=len(models_to_plot_subsampling) - 1)
    )
}

ds_colors = {
    "MGnify": list(ds_colors_subsampling.values())[-1],
    "BFD": list(ds_colors_subsampling.values())[3],
    "UniRef100": list(ds_colors_subsampling.values())[0],
    "ALL": sns.color_palette("coolwarm", n_colors=100)[0],
}

fig = plt.figure(figsize=FIGSIZE, constrained_layout=False)
axd = fig.subplot_mosaic(MOSAIC, gridspec_kw={"hspace": 0.4, "wspace": 0.4, "width_ratios": [1.2, 1, 1, 1, 1, 1],})#,  "height_ratios": [1, 1, 1]})
for ax in axd.values():
    ax.set_axis_off()

# plot_stacked_performance_h(delta_pt, fontsize=10, subplotspec=axd["A"].get_subplotspec())
plot_scaling_grid(tmp, ds_colors, fontsize=10, plot_legend=True, subplotspec=axd["B"].get_subplotspec(), legend_bbox_to_anchor=(0.5, 0.69))
plot_relative_improvements(tmp, ds_colors, subplotspec=axd["C"].get_subplotspec(), plot_legend=True, fontsize=10, legend_bbox_to_anchor=(0.5, 0.49), plot_titles=True,)

plot_dataset_composition_bars(
    post_filtering_sizes=post_filtering_sizes,
    models_to_plot=models_to_plot_subsampling,
    ds_colors=ds_colors,
    subplotspec=axd["E"].get_subplotspec(),
    fontsize=10,
    label_color="white",
)

ds_colors_subsampling = {
    model: color
    for (model, color) in zip(
        models_to_plot_subsampling.values(), sns.color_palette("Reds", n_colors=len(models_to_plot_subsampling) - 1)
    )
}

plot_all_metrics_histogram(
    plot_df,
    models_to_plot_subsampling,
    ds_colors_subsampling,
    metrics_to_plot_subsampling,
    group_order_subsampling,
    legend_bbox_to_anchor=(0.5, -0.01),
    baseline_model="UniRef100",
    shift_ylim=True,
    ncol=4,
    zero_line_proportion=0.5,
    sharey=False,
    legend_ncol=7,
    fontsize=10,
    subplotspec=axd["D"].get_subplotspec(),
)

plot_clustering_scatter(plot_df, subplotspec=axd["F"].get_subplotspec(), fontsize=10)
plot_clustering_epochs(subplotspec=axd["G"].get_subplotspec(), fontsize=10)

add_part_labels(axd, offset_x=-20, offset_y=20, fontsize=12)
plt.savefig("paper_figures/panel_scaling.pdf", bbox_inches="tight", dpi=300)
plt.savefig("paper_figures/panel_scaling.png", bbox_inches="tight", dpi=300)
plt.show()

# %% [markdown]
### Figure 4

# %% [code]
# ── Layout ────────────────────────────────────────────────────────────────────
FIGSIZE = (18, 9)
MOSAIC = """
ABBDD
ABBDD
CCCDD
CCCDD
"""
# ──────────────────────────────────────────────────────────────────────────────

fig = plt.figure(figsize=FIGSIZE)
axd = fig.subplot_mosaic(MOSAIC, gridspec_kw={"hspace": 0, "wspace": 0, "height_ratios": [1, 1, 0.8, 0.8], "width_ratios": [1, 1, 1, 1, 1]})
for ax in axd.values():
    ax.set_axis_off()

add_part_labels(axd, offset_x=-20, offset_y=20, fontsize=12)
plt.savefig("paper_figures/panel_final.png", bbox_inches="tight", dpi=300)
plt.savefig("paper_figures/panel_final.pdf", bbox_inches="tight", dpi=300)
plt.show()

# %% [code]
```
