## Supplementary material for "pLM representations unlock metagenomic space beyond homology": Code bundle: test_kl_consensus.py

```
"""
Unit tests for scripts/eval_consensus/kl_consensus.py

Run with:  pytest tests/eval_consensus/test_kl_consensus.py -v
"""

import sys
import types
from pathlib import Path
from unittest.mock import MagicMock, patch

import numpy as np
import pandas as pd
import pytest
import torch

# ---------------------------------------------------------------------------
# Make the script importable without the full amplify package
# ---------------------------------------------------------------------------
# Stub out the amplify import so we can import the module without the package.
amplify_stub = types.ModuleType("amplify")
amplify_model_stub = types.ModuleType("amplify.model")
amplify_model_stub.AMPLIFY = MagicMock()
amplify_stub.model = amplify_model_stub
sys.modules.setdefault("amplify", amplify_stub)
sys.modules.setdefault("amplify.model", amplify_model_stub)

SCRIPT = Path(__file__).parents[2] / "scripts" / "eval_consensus" / "kl_consensus.py"
import importlib.util

spec = importlib.util.spec_from_file_location("kl_consensus", SCRIPT)
kl_mod = importlib.util.module_from_spec(spec)
spec.loader.exec_module(kl_mod)

AA_COLS = kl_mod.AA_COLS
TRUNCATION_LEN = kl_mod.TRUNCATION_LEN
build_dataset = kl_mod.build_dataset
get_aa_token_ids = kl_mod.get_aa_token_ids
softmax_over_aa = kl_mod.softmax_over_aa
kl_divergence = kl_mod.kl_divergence


# ===========================================================================
# softmax_over_aa
# ===========================================================================

class TestSoftmaxOverAa:
    def _mock_aa_ids(self, n_vocab=50):
        """Map each AA to a unique token id."""
        return list(range(n_vocab - 20, n_vocab))  # last 20 tokens

    def test_output_sums_to_one(self):
        np.random.seed(0)
        logits = np.random.randn(50)
        aa_ids = self._mock_aa_ids(50)
        probs = softmax_over_aa(logits, aa_ids)
        assert probs.shape == (20,)
        assert abs(probs.sum() - 1.0) < 1e-6

    def test_all_equal_logits_uniform(self):
        logits = np.zeros(50)
        aa_ids = self._mock_aa_ids(50)
        probs = softmax_over_aa(logits, aa_ids)
        np.testing.assert_allclose(probs, np.ones(20) / 20, atol=1e-6)

    def test_argmax_matches_highest_logit(self):
        logits = np.zeros(50)
        aa_ids = self._mock_aa_ids(50)
        best_aa_idx = 5
        logits[aa_ids[best_aa_idx]] = 100.0  # dominate
        probs = softmax_over_aa(logits, aa_ids)
        assert np.argmax(probs) == best_aa_idx

    def test_non_aa_logits_ignored(self):
        """Logits outside aa_ids should not affect the result."""
        logits_a = np.zeros(50)
        logits_b = np.zeros(50)
        logits_b[0] = 1000.0  # non-AA token index
        aa_ids = self._mock_aa_ids(50)
        np.testing.assert_allclose(
            softmax_over_aa(logits_a, aa_ids),
            softmax_over_aa(logits_b, aa_ids),
            atol=1e-6,
        )

    def test_numerical_stability_large_logits(self):
        """Should not produce inf or nan with very large logits."""
        logits = np.full(50, 1e6)
        aa_ids = self._mock_aa_ids(50)
        probs = softmax_over_aa(logits, aa_ids)
        assert np.all(np.isfinite(probs))
        assert abs(probs.sum() - 1.0) < 1e-6


# ===========================================================================
# kl_divergence
# ===========================================================================

class TestKlDivergence:
    def test_identical_distributions_zero(self):
        p = np.array([0.5, 0.3, 0.2])
        assert kl_divergence(p, p) == pytest.approx(0.0, abs=1e-8)

    def test_non_negative(self):
        np.random.seed(42)
        for _ in range(20):
            p = np.random.dirichlet(np.ones(20))
            q = np.random.dirichlet(np.ones(20))
            assert kl_divergence(p, q) >= 0.0

    def test_asymmetric(self):
        # KL(p||q) != KL(q||p) in general; use distributions with different shapes
        p = np.array([0.7, 0.2, 0.1])
        q = np.array([0.1, 0.5, 0.4])
        assert kl_divergence(p, q) != pytest.approx(kl_divergence(q, p), abs=1e-6)

    def test_known_value(self):
        # KL(p||q) for p=[1,0], q=[0.5,0.5] = log(2) ≈ 0.693
        p = np.array([1.0, 0.0])
        q = np.array([0.5, 0.5])
        expected = np.log(2)
        assert kl_divergence(p, q) == pytest.approx(expected, abs=1e-5)

    def test_zero_mass_in_p_clipped(self):
        """Zero entries in p should not cause -inf (clipped to eps)."""
        p = np.array([1.0, 0.0, 0.0])
        q = np.array([0.5, 0.25, 0.25])
        result = kl_divergence(p, q)
        assert np.isfinite(result)

    def test_returns_float(self):
        p = np.ones(20) / 20
        q = np.ones(20) / 20
        assert isinstance(kl_divergence(p, q), float)


# ===========================================================================
# build_dataset
# ===========================================================================

class TestBuildDataset:
    def _make_proteome(self, tmp_path, sequences: dict[str, str]) -> str:
        df = pd.DataFrame({"id": list(sequences.keys()), "sequence": list(sequences.values())})
        path = str(tmp_path / "proteome.csv")
        df.to_csv(path, index=False)
        return path

    def _make_consensus(self, tmp_path, rows: list[dict]) -> str:
        df = pd.DataFrame(rows)
        path = str(tmp_path / "consensus.csv")
        df.to_csv(path, index=False)
        return path

    def _make_consensus_row(self, seq_name, pos, wt_aa, freqs: dict | None = None) -> dict:
        row = {"sequence_name": seq_name, "position": pos, "wt_aa": wt_aa}
        base = {aa: 0.0 for aa in AA_COLS}
        if freqs:
            base.update(freqs)
        row.update(base)
        return row

    def test_basic_row_included(self, tmp_path):
        seq = "A" * 10
        proteome = self._make_proteome(tmp_path, {"seq1": seq})
        consensus = self._make_consensus(tmp_path, [
            self._make_consensus_row("seq1", 3, "A", {"A": 1.0})
        ])
        ds = build_dataset(proteome, consensus)
        assert len(ds) == 1
        row = ds[0]
        assert row["id"] == "seq1"
        assert row["position"] == 3
        assert row["local_position"] == 3
        assert row["wt_aa"] == "A"
        assert row["consensus_aa"] == "A"  # highest freq AA

    def test_mask_replaces_target_position(self, tmp_path):
        seq = "ACDEFGHIKL"
        proteome = self._make_proteome(tmp_path, {"seq1": seq})
        consensus = self._make_consensus(tmp_path, [
            self._make_consensus_row("seq1", 2, "D", {"D": 1.0})
        ])
        ds = build_dataset(proteome, consensus)
        masked_seq = ds[0]["sequence"]
        # position 2 should be masked
        chars = masked_seq.split("<mask>")
        assert chars[0] == seq[:2]
        assert chars[1] == seq[3:]

    def test_missing_sequence_skipped(self, tmp_path):
        proteome = self._make_proteome(tmp_path, {"seq1": "ACDEF"})
        consensus = self._make_consensus(tmp_path, [
            self._make_consensus_row("seq_missing", 0, "A", {"A": 1.0})
        ])
        ds = build_dataset(proteome, consensus)
        assert len(ds) == 0

    def test_out_of_bounds_position_skipped(self, tmp_path):
        seq = "ACDEF"
        proteome = self._make_proteome(tmp_path, {"seq1": seq})
        consensus = self._make_consensus(tmp_path, [
            self._make_consensus_row("seq1", 999, "A", {"A": 1.0})
        ])
        ds = build_dataset(proteome, consensus)
        assert len(ds) == 0

    def test_truncation_long_sequence(self, tmp_path):
        seq = "A" * 1000
        pos = 500
        proteome = self._make_proteome(tmp_path, {"seq1": seq})
        consensus = self._make_consensus(tmp_path, [
            self._make_consensus_row("seq1", pos, "A", {"A": 1.0})
        ])
        ds = build_dataset(proteome, consensus)
        row = ds[0]
        # The returned sequence (with <mask>) should be based on a TRUNCATION_LEN window
        # local_position must be within the truncated window
        assert 0 <= row["local_position"] < TRUNCATION_LEN
        # The masked sequence length (excluding <mask> tag itself) equals TRUNCATION_LEN
        seq_no_mask = row["sequence"].replace("<mask>", "X")  # replace to count positions
        assert len(seq_no_mask) == TRUNCATION_LEN

    def test_truncation_preserves_local_position(self, tmp_path):
        seq = "A" * 1000
        pos = 500
        proteome = self._make_proteome(tmp_path, {"seq1": seq})
        consensus = self._make_consensus(tmp_path, [
            self._make_consensus_row("seq1", pos, "A", {"A": 1.0})
        ])
        ds = build_dataset(proteome, consensus)
        row = ds[0]
        # Reconstruct: the chars before <mask> should have length == local_position
        before_mask = row["sequence"].split("<mask>")[0]
        assert len(before_mask) == row["local_position"]

    def test_consensus_aa_is_argmax(self, tmp_path):
        seq = "ACDEF"
        proteome = self._make_proteome(tmp_path, {"seq1": seq})
        freqs = {aa: 0.0 for aa in AA_COLS}
        freqs["K"] = 0.8
        freqs["A"] = 0.2
        consensus = self._make_consensus(tmp_path, [
            self._make_consensus_row("seq1", 0, "A", freqs)
        ])
        ds = build_dataset(proteome, consensus)
        assert ds[0]["consensus_aa"] == "K"

    def test_aa_frequencies_stored(self, tmp_path):
        seq = "ACDEF"
        proteome = self._make_proteome(tmp_path, {"seq1": seq})
        freqs = {aa: float(i) for i, aa in enumerate(AA_COLS)}
        consensus = self._make_consensus(tmp_path, [
            self._make_consensus_row("seq1", 0, "A", freqs)
        ])
        ds = build_dataset(proteome, consensus)
        for aa in AA_COLS:
            assert ds[0][aa] == pytest.approx(freqs[aa])


# ===========================================================================
# collate_fn (tested via a mock tokenizer + manual batch)
# ===========================================================================

class TestCollateFn:
    """
    Test the collate_fn logic without a real tokenizer.
    We replicate the closure as it is built in evaluate().
    """

    def _make_collate_fn(self, pad_token_id: int):
        input_ids_pad = pad_token_id

        def collate_fn(batch):
            result = {}
            for key in batch[0]:
                vals = [b[key] for b in batch]
                if key == "input_ids":
                    tensors = [torch.tensor(v) for v in vals]
                    result[key] = torch.nn.utils.rnn.pad_sequence(
                        tensors, batch_first=True, padding_value=input_ids_pad
                    )
                elif key == "attention_mask":
                    tensors = [torch.tensor(v) for v in vals]
                    result[key] = torch.nn.utils.rnn.pad_sequence(
                        tensors, batch_first=True, padding_value=0
                    )
                elif isinstance(vals[0], (int, float)):
                    result[key] = torch.tensor(vals)
                else:
                    result[key] = vals
            return result

        return collate_fn

    def test_variable_length_input_ids_padded(self):
        fn = self._make_collate_fn(pad_token_id=1)
        batch = [
            {"input_ids": [2, 5, 6], "attention_mask": [1, 1, 1]},
            {"input_ids": [2, 7, 8, 9, 10], "attention_mask": [1, 1, 1, 1, 1]},
        ]
        out = fn(batch)
        assert out["input_ids"].shape == (2, 5)
        # short sequence padded with pad_token_id=1
        assert out["input_ids"][0, 3].item() == 1
        assert out["input_ids"][0, 4].item() == 1
        # long sequence untouched
        assert out["input_ids"][1, 4].item() == 10

    def test_attention_mask_padded_with_zero(self):
        fn = self._make_collate_fn(pad_token_id=1)
        batch = [
            {"input_ids": [2, 5], "attention_mask": [1, 1]},
            {"input_ids": [2, 5, 6, 7], "attention_mask": [1, 1, 1, 1]},
        ]
        out = fn(batch)
        assert out["attention_mask"].shape == (2, 4)
        assert out["attention_mask"][0, 2].item() == 0
        assert out["attention_mask"][0, 3].item() == 0

    def test_pad_token_id_used_not_zero(self):
        """input_ids must be padded with the tokenizer's pad_token_id, not 0."""
        PAD = 99
        fn = self._make_collate_fn(pad_token_id=PAD)
        batch = [
            {"input_ids": [2, 3], "attention_mask": [1, 1]},
            {"input_ids": [2, 3, 4, 5], "attention_mask": [1, 1, 1, 1]},
        ]
        out = fn(batch)
        assert out["input_ids"][0, 2].item() == PAD
        assert out["input_ids"][0, 3].item() == PAD

    def test_scalar_fields_become_tensors(self):
        fn = self._make_collate_fn(pad_token_id=0)
        batch = [
            {"input_ids": [1, 2], "attention_mask": [1, 1], "position": 5, "local_position": 2},
            {"input_ids": [1, 2, 3], "attention_mask": [1, 1, 1], "position": 10, "local_position": 4},
        ]
        out = fn(batch)
        assert isinstance(out["position"], torch.Tensor)
        assert out["position"].tolist() == [5, 10]
        assert out["local_position"].tolist() == [2, 4]

    def test_string_fields_stay_as_list(self):
        fn = self._make_collate_fn(pad_token_id=0)
        batch = [
            {"input_ids": [1], "attention_mask": [1], "id": "seq1", "wt_aa": "A"},
            {"input_ids": [1, 2], "attention_mask": [1, 1], "id": "seq2", "wt_aa": "C"},
        ]
        out = fn(batch)
        assert out["id"] == ["seq1", "seq2"]
        assert out["wt_aa"] == ["A", "C"]

    def test_float_aa_freq_columns_become_tensors(self):
        fn = self._make_collate_fn(pad_token_id=0)
        batch = [
            {"input_ids": [1], "attention_mask": [1], "A": 0.5},
            {"input_ids": [1, 2], "attention_mask": [1, 1], "A": 0.2},
        ]
        out = fn(batch)
        assert isinstance(out["A"], torch.Tensor)
        assert out["A"].tolist() == pytest.approx([0.5, 0.2])

    def test_output_shapes_consistent(self):
        fn = self._make_collate_fn(pad_token_id=0)
        batch = [
            {"input_ids": [1, 2, 3], "attention_mask": [1, 1, 1]},
            {"input_ids": [1, 2], "attention_mask": [1, 1]},
            {"input_ids": [1, 2, 3, 4, 5], "attention_mask": [1, 1, 1, 1, 1]},
        ]
        out = fn(batch)
        assert out["input_ids"].shape == out["attention_mask"].shape
        assert out["input_ids"].shape == (3, 5)
```
