## Supplementary material for "pLM representations unlock metagenomic space beyond homology": Code bundle: pyproject.toml

```
[build-system]
requires = ["setuptools >= 61.0"]
build-backend = "setuptools.build_meta"

[project]
name = "apperitif"
version = "0.0.0"
description = "A pipeline for protein Language Model evaluation"
requires-python = ">= 3.10"
dependencies = [
    "torch==2.5.1",
    "numpy==1.26.*",
    "transformers==4.49.0",
    "datasets>=3.1.0",
    "xformers==0.0.28.*",
    "flash-attn==2.7.*",
    "peft",
    "polars",
    "pandas",
    "pyarrow",
    "scikit-learn",
    "hydra-core==1.3.*",
    "accelerate",
    "scipy",
    "matplotlib",
    "tqdm",
    "pyyaml",
]
authors = [
    { name = "David Heurtel-Depeiges", email = "" },
    { name = "Lola Le Breton", email = "" },
    { name = "Quentin Fournier", email = "" },
]
maintainers = [
    { name = "David Heurtel-Depeiges", email = "" },
    { name = "Lola Le Breton", email = "" },
    { name = "Quentin Fournier", email = "" },
]
license = { text = "MIT License" }
keywords = ["Machine Learning", "Large Language Models", "Protein Language Models", "Evaluation"]

[project.optional-dependencies]
dev = ["sphinx", "black", "piccolo-theme", "pytest", "pytest-cov", "pytest-clarity", "xeger"]
```
