## Supplementary material for "pLM representations unlock metagenomic space beyond homology": Code bundle: aggregate_apperitif.py

```
import argparse
import ast
import json
import itertools
from pathlib import Path

import pandas as pd
import yaml


HP_KEYS = ["hidden_size", "learning_rate", "num_epochs", "weight_decay", "batch_size"]


def load_grid(yaml_path):
    with open(yaml_path) as f:
        cfg = yaml.safe_load(f)
    vals = [[float(x) for x in cfg[k]] for k in HP_KEYS]
    return set(itertools.product(*vals))


def parse_hp(k):
    return tuple(float(x) for x in ast.literal_eval(k)) if isinstance(k, str) else tuple(float(x) for x in k)


def best_for_grid(all_results, grid):
    best_val, best_entry = -float("inf"), None
    for k, v in all_results.items():
        hp = parse_hp(k)
        if hp not in grid:
            continue
        if v["validation_score"] > best_val:
            best_val = v["validation_score"]
            best_entry = (hp, v)
    return best_entry


def process_results(results_path, grids):
    with open(results_path) as f:
        data = json.load(f)

    all_results = data["all_results"]
    hp_keys = data.get("hyperparameter_keys", HP_KEYS)
    test_metrics = list(next(iter(all_results.values()))["test_scores"].keys())

    # Completion percentages
    row = {}
    if grids is not None:
        for gname, grid in grids.items():
            found = sum(1 for k in all_results if parse_hp(k) in grid)
            row[f"pct_completed_{gname}"] = round(100 * found / len(grid), 2)

        # Best per grid
        for gname, grid in grids.items():
            entry = best_for_grid(all_results, grid)
            if entry is None:
                for m in test_metrics:
                    row[f"best_test_{m}_{gname}"] = None
                row[f"best_val_score_{gname}"] = None
                row[f"best_val_std_{gname}"] = None
                for k in hp_keys:
                    row[f"best_{k}_{gname}"] = None
                continue
            hp, v = entry
            row[f"best_val_score_{gname}"] = v["validation_score"]
            row[f"best_val_std_{gname}"] = v.get("validation_std", None)
            for m in test_metrics:
                row[f"best_test_{m}_{gname}"] = v["test_scores"][m]
            for i, k in enumerate(hp_keys):
                row[f"best_{k}_{gname}"] = hp[i]

        best_grid = max(grids.keys(), key=lambda g: row[f"best_val_score_{g}"] if row[f"best_val_score_{g}"] is not None else -float("inf"))

        row["best_val_score"] = row[f"best_val_score_{best_grid}"]
        row["best_val_std"] = row[f"best_val_std_{best_grid}"]
        for m in test_metrics:
            row[f"best_test_{m}"] = row[f"best_test_{m}_{best_grid}"]

    else:
        row["num_completed"] = sum(1 for k in all_results if parse_hp(k) is not None)

        # Best overall
        best_val, best_entry = -float("inf"), None
        for k, v in all_results.items():
            # print(k, v)
            if parse_hp(k) is None:
                continue
            if v["validation_score"] > best_val:
                best_val = v["validation_score"]
                best_entry = (k, v)
        if best_entry is not None:
            k, v = best_entry
            row["best_val_score"] = v["validation_score"]
            row["best_val_std"] = v.get("validation_std", None)
            for m in test_metrics:
                row[f"best_test_{m}"] = v["test_scores"][m]
            for i, key in enumerate(hp_keys):
                # print(k, parse_hp(k))
                row[f"best_{key}"] = parse_hp(k)[i]
        else:
            row["best_val_score"] = None
            row["best_val_std"] = None
            for m in test_metrics:
                row[f"best_test_{m}"] = None
            for k in hp_keys:
                row[f"best_{k}"] = None

    return row


def main():
    parser = argparse.ArgumentParser()
    parser.add_argument("base_path", type=str)
    parser.add_argument("--output", "-o", type=str, default="aggregated_results.csv")
    parser.add_argument(
        "--grid-configs",
        nargs="+",
        default=[
            "conf/grid/mlp_feature_extraction_extended.yaml",
            "conf/grid/mlp_feature_extraction_restricted.yaml",
        ],
    )
    parser.add_argument("--no-grid", action="store_true")
    args = parser.parse_args()

    grids = None
    if not args.no_grid:
        grids = {}
        for path in args.grid_configs:
            name = Path(path).stem.split("mlp_feature_extraction_")[-1]
            grids[name] = load_grid(path)

    print(grids)

    base = Path(args.base_path)
    rows = []
    for task_dir in sorted(base.iterdir()):
        if not task_dir.is_dir():
            continue
        task = task_dir.name
        for model_dir in sorted(task_dir.iterdir()):
            if not model_dir.is_dir():
                continue
            results_file = model_dir / "results.json"
            if not results_file.exists():
                print(f"File not found: {results_file}")
                continue
            row = process_results(results_file, grids)
            row["task"] = task
            row["model"] = model_dir.name
            rows.append(row)

    df = pd.DataFrame(rows)
    cols = ["task", "model"] + [c for c in df.columns if c not in ("task", "model")]
    df = df[cols]
    df.to_csv(args.output, index=False)
    print(f"Saved {len(df)} rows to {args.output}")


if __name__ == "__main__":
    main()
```
