## Supplementary material for "pLM representations unlock metagenomic space beyond homology": Code bundle: random_sequences.py

```
import os
import pandas as pd
import random
from argparse import ArgumentParser

if __name__ == "__main__":
    parser = ArgumentParser()
    parser.add_argument("--input_file", type=str, required=True)
    parser.add_argument("--output_path", default=None, type=str, required=False)
    parser.add_argument("--output_name", default="random_512", type=str, required=False)

    args = parser.parse_args()

    if args.output_path is None:
        args.output_path = os.path.dirname(os.path.abspath(args.input_file))

    df = pd.read_csv(args.input_file)
    df = df[df["id"].str.endswith("_0")]
    df = df.sample(n=1000, random_state=42).reset_index(drop=True)

    # Function to shuffle a sequence while keeping the first 'M' (if present)
    def shuffle(seq):
        if not seq:
            return seq
        if seq[0] == "M":
            first = "M"
            rest = list(seq[1:])
        else:
            first = ""
            rest = list(seq)
        random.shuffle(rest)
        return first + "".join(rest)

    df["sequence"] = df["sequence"].apply(shuffle)

    df["id"] = df["id"].apply(lambda x: x.split("_0")[0] + "_rand_0")

    df.to_csv(os.path.join(args.output_path, f"{args.output_name}.csv"), index=False)
```
