## Supplementary material for "pLM representations unlock metagenomic space beyond homology": Code bundle: run_hhblits.py

```
import pandas as pd
import os

human_df = pd.read_csv("UP000005640_9606.csv")
human_df = human_df.sample(frac=1).reset_index(drop=True)

for i, row in human_df.iterrows():
    print(row)
    name = row["name"].split()[0].split("|")[1]
    print(name)
    print(row.sequence)

    bfd_path = "$SCRATCH/proseqo/datasets/bfd/download/bfd_metaclust_clu_complete_id30_c90_final_seq.sorted_opt"

    if not os.path.exists(f"./hhblits_99.50.50/fasta/{name}.fasta"):
        with open(f"./hhblits_99.50.50/fasta/{name}.fasta", "wt") as ofile:
            ofile.write(f">{name}\n{row.sequence}\n")
        os.system(
            f"$HOME/hh-suite/build/bin/hhblits -i ./hhblits_99.50.50/fasta/{name}.fasta -o ./hhblits_99.50.50/results/{name}.result -cpu 40 -d {bfd_path} -oa3m ./hhblits_99.50.50/{name}.align.a3m -id 99 -cov 50 -qid 50"
        )

        with open(f"./hhblits_99.x.x/fasta/{name}.fasta", "wt") as ofile:
            ofile.write(f">{name}\n{row.sequence}\n")
        os.system(
            f"$HOME/hh-suite/build/bin/hhblits -i ./hhblits_99.x.x/fasta/{name}.fasta -o ./hhblits_99.x.x/results/{name}.result -cpu 40 -d {bfd_path} -oa3m ./hhblits_99.x.x/{name}.align.a3m -id 99"
        )
```
