## Supplementary material for "pLM representations unlock metagenomic space beyond homology": Code bundle: aggregate_results.py

```
import pandas as pd
import json
import argparse
from pathlib import Path
import sys


def main():
    parser = argparse.ArgumentParser(description="Aggregate results_*.json files into a DataFrame.")
    parser.add_argument("root_dir", type=str, help="Path to the root directory containing results")
    parser.add_argument("-o", "--output", type=str, default="results.csv", help="Output CSV filename (default: results.csv)")
    args = parser.parse_args()

    root_path = Path(args.root_dir)
    if not root_path.exists():
        sys.exit(f"Error: Directory '{root_path}' not found.")

    files = list(root_path.rglob("results_*.json"))

    if not files:
        sys.exit("No result files found.")

    records = []

    for file_path in files:
        try:
            with open(file_path, "r") as f:
                content = json.load(f)

            # Path structure: {root}/{repo_parts...}/{model_name}/results_{split}.json
            rel_path = file_path.relative_to(root_path)
            parts = rel_path.parts  # e.g. ('<org>', 'MILA_BFD', 'results_casp16.json')

            model_name = parts[-2] if len(parts) >= 2 else None
            repo_name = "/".join(parts[:-2]) if len(parts) >= 3 else None

            entry = {
                "repo_name": repo_name,
                "model_name": model_name,
                # Prefer the model field from JSON if present, else reconstruct from path
                "model": content.get("model") or (f"{repo_name}/{model_name}" if repo_name else model_name),
                "split": content.get("split"),
            }

            # Flatten summary metrics as top-level columns
            summary = content.get("summary", {})
            entry.update(summary)

            # Compute std from per_sequence
            per_seq = content.get("per_sequence", [])
            if per_seq:
                metric_keys = [k for k in per_seq[0].keys() if k not in ("seq_idx", "length")]
                for k in metric_keys:
                    vals = [s[k] for s in per_seq if k in s]
                    entry[f"{k}_std"] = pd.Series(vals).std()

            records.append(entry)

        except Exception as e:
            print(f"[Warn] Skipping {file_path}: {e}", file=sys.stderr)

    df = pd.DataFrame(records)

    df.to_csv(args.output, index=False)
    print(f"Success. {len(df)} records saved to {args.output}")


if __name__ == "__main__":
    main()
```
