## Supplementary material for "pLM representations unlock metagenomic space beyond homology": Code bundle: categorical_jacobian.py

```
import argparse
import json
import os
import torch
import numpy as np
import torch.nn.functional as F
from tqdm import tqdm
from datasets import load_dataset
from sklearn.metrics import roc_auc_score
from transformers import AutoTokenizer, AutoModel, AutoModelForMaskedLM


def get_logits(model, input_ids):
    out = model(input_ids)
    if hasattr(out, "logits"):
        return out.logits
    return out


def get_categorical_jacobian(model, tokenizer, sequence, batch_size=32, device="cuda"):
    """
    Categorical Jacobian J[i,a,j,b] = log P(x_i=a | x_j=b) - log P(x_i=a | wt)
    as defined in "Protein language models learn evolutionary statistics of
    interacting sequence motifs" (Sgarbossa et al.).

    Returns jacobian of shape (L, 20, L, 20).
    """
    model.eval()
    L = len(sequence)

    inputs = tokenizer(sequence, return_tensors="pt").to(device)
    input_ids = inputs["input_ids"]

    if len(input_ids.shape) == 1:
        input_ids = input_ids.unsqueeze(0)  # [1, L+2] with cls/eos tokens

    standard_aa = "ARNDCQEGHILKMFPSTWYV"
    vocab_ids = torch.tensor(tokenizer.convert_tokens_to_ids(list(standard_aa)), device=device)
    num_aa = len(vocab_ids)  # 20

    # Determine offset: some tokenizers prepend a BOS token, others don't
    token_offset = input_ids.shape[1] - L  # total extra tokens; BOS adds 1 at start if present
    has_bos = (
        token_offset > 0
        and input_ids[0, 0].item() == tokenizer.cls_token_id
        or token_offset > 0
        and tokenizer.bos_token_id is not None
        and input_ids[0, 0].item() == tokenizer.bos_token_id
    )
    seq_offset = 1 if has_bos else 0

    with torch.no_grad():
        wt_logits = get_logits(model, input_ids)[0, seq_offset : seq_offset + L, :]  # [L, V]
        wt_log_probs = F.log_softmax(wt_logits, dim=-1)[:, vocab_ids]  # [L, 20]

    # Build all L*20 mutant sequences
    batch_input_ids = input_ids.repeat(L * num_aa, 1)
    mutation_positions = torch.arange(seq_offset, seq_offset + L, device=device).repeat_interleave(num_aa)
    mutation_tokens = vocab_ids.repeat(L)
    row_indices = torch.arange(L * num_aa, device=device)
    batch_input_ids[row_indices, mutation_positions] = mutation_tokens

    jacobian = torch.zeros((L * num_aa, L, num_aa), device=device, dtype=torch.float32)

    dataset = torch.utils.data.TensorDataset(batch_input_ids)
    loader = torch.utils.data.DataLoader(dataset, batch_size=batch_size, shuffle=False)

    offset = 0
    with torch.no_grad():
        for (b_ids,) in tqdm(loader, desc=f"Jacobian L={L}", leave=False):
            b_logits = get_logits(model, b_ids)[:, seq_offset : seq_offset + L, :]  # [B, L, V]
            b_log_probs = F.log_softmax(b_logits, dim=-1)[:, :, vocab_ids]  # [B, L, 20]
            B = b_ids.shape[0]
            jacobian[offset : offset + B] = b_log_probs - wt_log_probs.unsqueeze(0)
            offset += B

    return jacobian.view(L, num_aa, L, num_aa)


def jacobian_to_contacts(jacobian):
    """
    Converts Jacobian (L, 20, L, 20) to a contact map (L, L) via:
    4-pass centering -> symmetrization -> Frobenius norm -> APC correction.
    """
    # 4-pass sequential centering (subtract mean over each dimension in order)
    j = jacobian
    for i in range(4):
        j = j - j.mean(i, keepdim=True)

    # Symmetrize
    j = (j + j.permute(2, 3, 0, 1)) / 2.0

    # Frobenius norm over amino acid axes
    contacts = torch.sqrt((j**2).sum(dim=(1, 3)))  # [L, L]

    # Zero diagonal before APC
    contacts.fill_diagonal_(0)

    # APC
    a1 = contacts.sum(dim=0, keepdim=True)
    a2 = contacts.sum(dim=1, keepdim=True)
    contacts = contacts - (a1 * a2) / contacts.sum()

    contacts.fill_diagonal_(0)
    return contacts


def pairs_to_matrix(pairs, L):
    """Converts a list of [i, j] contact pairs to a symmetric binary matrix."""
    mat = np.zeros((L, L), dtype=np.float32)
    for i, j in pairs:
        mat[i, j] = 1.0
        mat[j, i] = 1.0
    return mat


def compute_metrics(contact_map, true_contacts, L, minsep=6):
    """
    Computes P@L, P@L/2, P@L/5, ESM-style AUC (mean binned precision), and ROC-AUC
    on long-range contacts (sep >= minsep), following the ESM implementation.

    contact_map: (L, L) numpy array of predicted scores
    true_contacts: (L, L) binary numpy array (negative values treated as invalid)
    """
    predictions = torch.from_numpy(contact_map).unsqueeze(0)  # [1, L, L]
    targets = torch.from_numpy(true_contacts).unsqueeze(0)  # [1, L, L]

    seqlen_range = torch.arange(L)
    sep = seqlen_range.unsqueeze(0) - seqlen_range.unsqueeze(1)  # [L, L]
    valid_mask = (sep >= minsep) & (targets[0] >= 0)  # [L, L]

    # Mask invalid positions to -inf so they sink in ranking
    predictions = predictions.masked_fill(~valid_mask.unsqueeze(0), float("-inf"))

    # Upper triangle with minsep offset (avoids double-counting and short-range pairs)
    x_ind, y_ind = np.triu_indices(L, minsep)
    scores = predictions[0, x_ind, y_ind].numpy()  # 1D
    labels = targets[0, x_ind, y_ind].numpy()  # 1D

    if labels.sum() == 0:
        return None

    # ESM-style binned precision: top L*0.1, L*0.2, ..., L*1.0
    indices = np.argsort(scores)[::-1][:L]
    topk_targets = labels[indices]
    if len(topk_targets) < L:
        topk_targets = np.pad(topk_targets, (0, L - len(topk_targets)))
    cumulative_dist = topk_targets.cumsum()
    gather_indices = (np.arange(0.1, 1.1, 0.1) * L).astype(int) - 1
    gather_indices = np.clip(gather_indices, 0, len(cumulative_dist) - 1)
    binned_precisions = cumulative_dist[gather_indices] / (gather_indices + 1)

    metrics = {
        "P@L": float(binned_precisions[9]),
        "P@L/2": float(binned_precisions[4]),
        "P@L/5": float(binned_precisions[1]),
        "AUC": float(binned_precisions.mean()),
        "ROC-AUC": float(roc_auc_score(labels, scores)),
    }
    return metrics


def load_model(model_name, device, hf_token=None):
    tokenizer = AutoTokenizer.from_pretrained(model_name, trust_remote_code=True, token=hf_token)
    # AMPLIFY-family models expose a custom encoder via trust_remote_code (AutoModel);
    # ESM and other standard MLMs use AutoModelForMaskedLM.
    if "esm" not in model_name.lower():
        model = AutoModel.from_pretrained(model_name, trust_remote_code=True, token=hf_token)
    else:
        model = AutoModelForMaskedLM.from_pretrained(model_name, trust_remote_code=True, token=hf_token)
    return model.to(device), tokenizer


def main():
    parser = argparse.ArgumentParser(description="Categorical Jacobian contact prediction on CASP16 dataset.")
    parser.add_argument("--model", default="facebook/esm2_t12_35M_UR50D")
    parser.add_argument("--split", default="casp16", choices=["train", "valid", "test", "casp16"])
    parser.add_argument("--batch-size", type=int, default=16)
    parser.add_argument("--max-len", type=int, default=1024)
    parser.add_argument("--output-dir", default=None, help="If set, save per-sequence contact maps as .npy files.")
    parser.add_argument("--results", default="results.json", help="Path to save aggregated metrics JSON.")
    parser.add_argument("--token", default=None, help="HuggingFace access token.")
    args = parser.parse_args()

    device = "cuda" if torch.cuda.is_available() else "cpu"
    model, tokenizer = load_model(args.model, device, hf_token=args.token)

    ds = load_dataset(
        "davidhd/contact_prediction_binary_casp16",
        split=args.split,
        token=args.token,
    )

    if args.output_dir:
        os.makedirs(args.output_dir, exist_ok=True)

    all_metrics = []

    for i, sample in enumerate(tqdm(ds, desc="Sequences")):
        seq = sample["seq"]
        L = len(seq)

        if L > args.max_len:
            print(f"Skipping sequence {i} (len={L} > {args.max_len})")
            continue

        jacobian = get_categorical_jacobian(model, tokenizer, seq, args.batch_size, device)
        contact_map = jacobian_to_contacts(jacobian).cpu().numpy()

        true_contacts = pairs_to_matrix(sample["label"], L)
        metrics = compute_metrics(contact_map, true_contacts, L)

        if metrics is not None:
            metrics["seq_idx"] = i
            metrics["length"] = L
            all_metrics.append(metrics)

        if args.output_dir:
            np.save(f"{args.output_dir}/contacts_{i:04d}.npy", contact_map)

    # Aggregate
    keys = ["P@L", "P@L/2", "P@L/5", "AUC", "ROC-AUC"]
    summary = {k: float(np.mean([m[k] for m in all_metrics])) for k in keys}
    summary["n_sequences"] = len(all_metrics)

    print("\nResults:")
    for k, v in summary.items():
        print(f"  {k}: {v:.4f}" if isinstance(v, float) else f"  {k}: {v}")

    output = {"model": args.model, "split": args.split, "summary": summary, "per_sequence": all_metrics}
    os.makedirs(os.path.dirname(os.path.abspath(args.results)), exist_ok=True)
    with open(args.results, "w") as f:
        json.dump(output, f, indent=2)
    print(f"Saved results to {args.results}")


if __name__ == "__main__":
    main()
```
