## Supplementary material for "pLM representations unlock metagenomic space beyond homology": Code bundle: aggregate_kl_consensus.py

```
"""Aggregate scores.json files from kl_consensus eval into a single CSV."""

import json
import sys
import argparse
from pathlib import Path

import pandas as pd


def main():
    parser = argparse.ArgumentParser(description=__doc__)
    parser.add_argument("root_dir", type=str, help="Root directory containing per-model result folders")
    parser.add_argument("-o", "--output", type=str, default="kl_consensus_results.csv", help="Output CSV path (default: kl_consensus_results.csv)")
    args = parser.parse_args()

    root = Path(args.root_dir)
    if not root.exists():
        sys.exit(f"Error: '{root}' not found.")

    score_files = sorted(root.rglob("scores.json"))
    if not score_files:
        sys.exit("No scores.json files found.")

    records = []
    for path in score_files:
        try:
            with open(path) as f:
                scores = json.load(f)
            model_name = path.parent.relative_to(root).as_posix()
            records.append({"model": model_name, **scores})
        except Exception as e:
            print(f"[warn] Skipping {path}: {e}", file=sys.stderr)

    df = pd.DataFrame(records)
    cols = ["model"] + [c for c in df.columns if c != "model"]
    df = df[cols].sort_values("model").reset_index(drop=True)

    df.to_csv(args.output, index=False)
    print(f"{len(df)} records saved to {args.output}")


if __name__ == "__main__":
    main()
```
