## Supplementary material for "pLM representations unlock metagenomic space beyond homology": Code bundle: aggregate_results.py

```
import pandas as pd
import json
import argparse
from pathlib import Path
import sys


def parse_path_parts(relative_path):
    """
    Parses a relative path into (repo_name, model_name, max_length).
    Logic assumes:
    - Deepest folder matching 'len' or digits is max_length.
    - Deepest remaining folder is model_name.
    - Anything above model_name is repo_name.
    """
    parts = list(relative_path.parts)
    repo = None
    model = None
    length = None

    if not parts:
        return repo, model, length

    # 1. Identify max_length (optional, usually at the end)
    if len(parts) > 0:
        last = parts[-1]
        # Heuristic: if last folder contains "len" or is purely numeric
        if "len" in last.lower() or last.replace("_", "").isdigit():
            length = parts.pop()

    # 2. Identify model_name (Required, usually the last remaining)
    if len(parts) > 0:
        model = parts.pop()

    # 3. Identify repo_name (Optional, whatever is left)
    if len(parts) > 0:
        repo = "/".join(parts)

    return repo, model, length


def main():
    parser = argparse.ArgumentParser(description="Aggregate result.json files into a DataFrame.")
    parser.add_argument("root_dir", type=str, help="Path to the root directory containing results")
    parser.add_argument("-o", "--output", type=str, default="results.csv", help="Output CSV filename (default: results.csv)")
    args = parser.parse_args()

    root_path = Path(args.root_dir)
    if not root_path.exists():
        sys.exit(f"Error: Directory '{root_path}' not found.")

    # Find all result files (matches score.json or scores.json)
    files = list(root_path.rglob("score*.json"))

    if not files:
        sys.exit("No result files found.")

    records = []

    for file_path in files:
        try:
            # Parse JSON
            with open(file_path, "r") as f:
                content = json.load(f)

            # Flatten if not a dict
            if not isinstance(content, dict):
                content = {"data": content}

            # Parse Path Structure
            rel_path = file_path.parent.relative_to(root_path)
            repo, model, length = parse_path_parts(rel_path)

            entry = {"repo_name": repo, "model_name": model, "max_length": length}
            entry.update(content)
            records.append(entry)

        except Exception as e:
            # Low verbosity: only print errors affecting data integrity
            print(f"[Warn] Skipping {file_path}: {e}", file=sys.stderr)

    # Create DataFrame
    df = pd.DataFrame(records)

    # Reorder columns: Meta -> Data
    cols = list(df.columns)
    meta = ["repo_name", "model_name", "max_length"]
    # Keep only meta cols that actually exist in the data
    meta = [c for c in meta if c in cols]
    others = [c for c in cols if c not in meta]

    df = df[meta + others]

    # Save
    df.to_csv(args.output, index=False)
    print(f"Success. {len(df)} records saved to {args.output}")


if __name__ == "__main__":
    main()
```
