## Supplementary material for "pLM representations unlock metagenomic space beyond homology": Code bundle: extract_consensus_positions.py

```
"""
Extract positions with consensus amino acids from HHblits a3m alignment files.

A consensus position is defined as a position where at least one amino acid
has a frequency >= CONSENSUS_THRESHOLD (default: 0.70).

Output columns:
  sequence_name, position, wt_aa, num_homologs, A, C, D, E, F, G, H, I, K, L, M, N, P, Q, R, S, T, V, W, Y
"""

import argparse
from pathlib import Path
import pandas as pd

AA_COLS = list("ACDEFGHIKLMNPQRSTVWY")
AA_SET = set(AA_COLS)


def parse_a3m(a3m_file: Path):
    """
    Parse an a3m file. Returns (query_seq, homolog_seqs) where:
      - query_seq: str, the first sequence (no gaps)
      - homolog_seqs: list of str, subsequent sequences (uppercase only, gaps removed from insertion columns)
    """
    sequences = []
    current_seq = []
    with open(a3m_file) as f:
        for line in f:
            line = line.rstrip()
            if line.startswith(">"):
                if current_seq:
                    sequences.append("".join(current_seq))
                current_seq = []
            else:
                # In a3m format, lowercase = insertion columns (skip for MSA columns)
                current_seq.append(line)
    if current_seq:
        sequences.append("".join(current_seq))

    if not sequences:
        return None, []

    # Query sequence is the first one; strip insertions (lowercase letters)
    def strip_insertions(seq):
        return "".join(c for c in seq if not c.islower())

    query_seq = strip_insertions(sequences[0])
    homolog_seqs = [strip_insertions(s) for s in sequences[1:]]
    return query_seq, homolog_seqs


def compute_profile(query_seq: str, homolog_seqs: list) -> pd.DataFrame:
    """
    Compute per-position AA frequency profile from MSA columns.
    Only counts non-gap, standard AA characters. Query gaps are excluded.
    Returns a DataFrame with columns: res_ix, AA, num_homologs, A..Y
    """
    query_len = len(query_seq)
    num_homologs = len(homolog_seqs)

    records = []
    query_pos = 0  # 0-based index in query (non-gap positions)

    for col_ix in range(query_len):
        query_aa = query_seq[col_ix]
        if query_aa == "-":
            continue  # skip gap positions in query

        # Collect AAs from homologs at this column (exclude gaps and non-standard)
        counts = {aa: 0 for aa in AA_COLS}
        total = 0
        for hseq in homolog_seqs:
            if col_ix >= len(hseq):
                continue
            aa = hseq[col_ix].upper()
            if aa in AA_SET:
                counts[aa] += 1
                total += 1

        # Also count the query itself if it's a standard AA
        if query_aa.upper() in AA_SET:
            counts[query_aa.upper()] += 1
            total += 1

        if total == 0:
            query_pos += 1
            continue

        freqs = {aa: counts[aa] / total for aa in AA_COLS}
        record = {"res_ix": query_pos, "AA": query_aa.upper() if query_aa != "-" else "-"}
        record.update(freqs)
        records.append(record)
        query_pos += 1

    df = pd.DataFrame(records, columns=["res_ix", "AA"] + AA_COLS)
    return df, num_homologs


def extract_consensus_positions(hhblits_dir: Path, threshold: float) -> pd.DataFrame:
    records = []

    a3m_files = sorted(hhblits_dir.glob("*.a3m"))
    for a3m_file in a3m_files:
        seq_name = a3m_file.stem
        if seq_name.endswith(".align"):
            seq_name = seq_name[: -len(".align")]

        query_seq, homolog_seqs = parse_a3m(a3m_file)
        if query_seq is None:
            continue

        df, num_homologs = compute_profile(query_seq, homolog_seqs)

        # Keep only rows where any AA has frequency >= threshold
        mask = (df[AA_COLS] >= threshold).any(axis=1)
        consensus_rows = df[mask]

        for _, row in consensus_rows.iterrows():
            record = {
                "sequence_name": seq_name,
                "position": int(row["res_ix"]),
                "wt_aa": row["AA"],
                "num_homologs": num_homologs,
            }
            for aa in AA_COLS:
                record[aa] = row[aa]
            records.append(record)

    result = pd.DataFrame(records, columns=["sequence_name", "position", "wt_aa", "num_homologs"] + AA_COLS)
    return result


def main():
    parser = argparse.ArgumentParser(description=__doc__)
    parser.add_argument(
        "--hhblits_dir",
        type=Path,
        required=True,
        help="Directory containing HHblits a3m alignment files",
    )
    parser.add_argument(
        "--threshold",
        type=float,
        default=0.70,
        help="Minimum frequency to consider a position as consensus (default: 0.70)",
    )
    parser.add_argument(
        "--output",
        type=Path,
        default=Path("consensus_positions.csv"),
        help="Output CSV path (default: consensus_positions.csv)",
    )
    args = parser.parse_args()

    print(f"Scanning {args.hhblits_dir} with threshold={args.threshold} ...")
    df = extract_consensus_positions(args.hhblits_dir, args.threshold)
    print(f"Found {len(df)} consensus positions across {df['sequence_name'].nunique()} sequences.")

    df.to_csv(args.output, index=False)
    print(f"Saved to {args.output}")


if __name__ == "__main__":
    main()
```
