## Supplementary material for "pLM representations unlock metagenomic space beyond homology": Code bundle: kl_consensus.py

```
"""
Evaluate model logits against homolog AA frequency profiles using KL divergence.

For each consensus position (AA frequency >= threshold in homologs), the WT residue
is masked and the model predicts a distribution over the 20 standard AAs.
KL(homolog_freq || model_softmax) is computed per position.

Output parquet columns per row (one per sequence×position):
  sequence_name, position, wt_aa, consensus_aa, pred_aa, kl
"""

import os
import re
import json
import logging
from argparse import ArgumentParser
from pathlib import Path

import numpy as np
import pandas as pd
import torch
from tqdm import tqdm
from datasets import Dataset
from transformers import AutoTokenizer, AutoModel, AutoModelForMaskedLM
from torch.utils.data import DataLoader

from amplify.model import AMPLIFY

AA_COLS = list("ACDEFGHIKLMNPQRSTVWY")
TRUNCATION_LEN = 512
logger = logging.getLogger(__name__)
logging.basicConfig(level=logging.INFO)


# ---------------------------------------------------------------------------
# Dataset preparation
# ---------------------------------------------------------------------------


def build_dataset(proteome_path: str, consensus_path: str) -> Dataset:
    """
    Build a flat dataset of (masked_sequence, metadata) rows, one per consensus position.
    Each sequence is truncated to TRUNCATION_LEN with the masked position centred.
    """
    proteome = pd.read_csv(proteome_path).set_index("id")
    consensus = pd.read_csv(consensus_path)  # sequence_name, position, wt_aa, A..Y

    rows = []
    missing = 0
    skipped_non_standard = 0
    for _, cons_row in consensus.iterrows():
        seq_name = cons_row["sequence_name"]
        pos = int(cons_row["position"])

        if seq_name not in proteome.index:
            missing += 1
            continue

        full_seq = proteome.loc[seq_name, "sequence"]
        seq_len = len(full_seq)

        if pos >= seq_len:
            continue

        if cons_row["wt_aa"] not in AA_COLS:
            skipped_non_standard += 1
            continue

        # Determine the consensus AA (highest frequency in profile)
        freqs = cons_row[AA_COLS].values.astype(float)
        consensus_aa = AA_COLS[int(np.argmax(freqs))]

        # Truncate window centred on pos
        if seq_len > TRUNCATION_LEN:
            half = TRUNCATION_LEN // 2
            start = max(0, pos - half)
            end = start + TRUNCATION_LEN
            if end > seq_len:
                end = seq_len
                start = max(0, end - TRUNCATION_LEN)

            local_pos = pos - start  # position within the truncated window

        else:
            start = 0
            end = seq_len
            local_pos = pos

        rows.append(
            {
                "sequence": full_seq[start:end],
                "id": seq_name,
                "position": pos,
                "local_position": local_pos,
                "wt_aa": cons_row["wt_aa"],
                "consensus_aa": consensus_aa,
                **{aa: float(cons_row[aa]) for aa in AA_COLS},
            }
        )

    if missing:
        logger.warning(f"{missing} sequence names from consensus file not found in proteome.")
    if skipped_non_standard:
        logger.info(f"Skipped {skipped_non_standard} consensus positions with non-standard wt_aa.")

    logger.info(f"Built dataset with {len(rows)} consensus positions.")
    return Dataset.from_list(rows)


# ---------------------------------------------------------------------------
# KL computation helpers
# ---------------------------------------------------------------------------


def get_aa_token_ids(tokenizer) -> list[int]:
    """Return token IDs for the 20 standard AAs, in AA_COLS order."""
    ids = tokenizer.convert_tokens_to_ids(AA_COLS)
    assert all(i != tokenizer.unk_token_id for i in ids), "Some AAs mapped to <unk> — check vocab."
    return ids


def softmax_over_aa(logits_np: np.ndarray, aa_token_ids: list[int]) -> np.ndarray:
    """Extract logits for 20 AAs and apply softmax (re-normalised over 20 AAs only)."""
    aa_logits = logits_np[aa_token_ids]
    aa_logits = aa_logits - aa_logits.max()  # numerical stability
    exp = np.exp(aa_logits)
    return exp / exp.sum()


def kl_divergence(p: np.ndarray, q: np.ndarray, eps: float = 1e-10) -> float:
    """KL(p || q) — p is the homolog frequency distribution."""
    p = np.clip(p, eps, 1.0)
    q = np.clip(q, eps, 1.0)
    return float(np.sum(p * np.log(p / q)))


# ---------------------------------------------------------------------------
# Main evaluator
# ---------------------------------------------------------------------------


def evaluate(args):
    os.makedirs(args.output_path, exist_ok=True)

    # ---- Resumability: find last saved parquet ----
    parquet_pattern = re.compile(r"data_(\d+)\.parquet")
    existing = [int(m.group(1)) for f in os.listdir(args.output_path) if (m := parquet_pattern.match(f))]
    start_idx = (max(existing) + 1) if existing else 0
    if start_idx > 0:
        logger.info(f"Resuming from batch {start_idx}")

    if not args.skip_predictions:
        # ---- Load model ----
        from_hub = (
            "esm" in args.model_name
            or "AMPLIFY" in args.model_name
            or args.model_name.lower().startswith("<HF_ORG>/")
            or args.model_name.lower().startswith("flair-bio/")
            or args.model_name.startswith("chandar-lab/")
            or args.model_name.startswith("facebook/")
        )
        if from_hub:
            tokenizer = AutoTokenizer.from_pretrained(args.model_name, trust_remote_code=True, token=args.token)
            if "AMPLIFY" in args.model_name or args.model_name.lower().startswith("<HF_ORG>/"):
                model = AutoModel.from_pretrained(args.model_name, trust_remote_code=True, token=args.token)
            else:
                model = AutoModelForMaskedLM.from_pretrained(args.model_name, trust_remote_code=True, token=args.token)
        else:
            model, tokenizer = AMPLIFY.load(
                checkpoint_path=args.checkpoint_path,
                config_path=args.config_path,
                vocab_path=args.vocab_path,
                tag=args.tag,
            )
            if args.tag is not None:
                args.model_name = args.model_name + "_" + args.tag

        aa_token_ids = get_aa_token_ids(tokenizer)

        probe_ids = tokenizer("A", add_special_tokens=True)["input_ids"]
        bos_offset = 1 if tokenizer.bos_token_id is not None and probe_ids[0] == tokenizer.bos_token_id else 0
        n_special = len(probe_ids) - 1
        logger.info(f"Tokenizer probe: bos_offset={bos_offset}, n_special_tokens={n_special}")

        device = torch.device("cuda" if torch.cuda.is_available() else "cpu")
        model = model.to(device)
        model.eval()

        # ---- Build or load dataset ----
        if args.dataset_path and os.path.isdir(args.dataset_path):
            logger.info(f"Loading pre-built dataset from {args.dataset_path}")
            dataset = Dataset.load_from_disk(args.dataset_path)
        else:
            logger.info("Building dataset from proteome and consensus files...")
            dataset = build_dataset(args.proteome_path, args.consensus_path)
            if args.dataset_path:
                dataset.save_to_disk(args.dataset_path)
                logger.info(f"Saved dataset to {args.dataset_path}")

        max_length = TRUNCATION_LEN + n_special
        mask_token_id = tokenizer.mask_token_id
        assert mask_token_id is not None, "Tokenizer has no mask_token_id."

        def tokenize(batch):
            enc = tokenizer(batch["sequence"], truncation=True, max_length=max_length, padding=False, return_tensors=None)
            mask_token_indices = []
            for i, (seq, lp) in enumerate(zip(batch["sequence"], batch["local_position"])):
                target_pid = bos_offset + lp
                if "position_ids" in enc:
                    pids = enc["position_ids"][i]
                    matches = [j for j, p in enumerate(pids) if p == target_pid]
                    assert len(matches) == 1, (
                        f"Expected exactly one token with position_id={target_pid}, got {matches} "
                        f"(seq len {len(seq)}, lp {lp}, pids={pids})"
                    )
                    mask_idx = matches[0]
                else:
                    mask_idx = target_pid
                    assert mask_idx < len(enc["input_ids"][i]) - (n_special - bos_offset), (
                        f"mask_idx {mask_idx} out of range for tokenized length {len(enc['input_ids'][i])} "
                        f"(seq len {len(seq)}, lp {lp}, n_special {n_special})"
                    )
                enc["input_ids"][i][mask_idx] = mask_token_id
                mask_token_indices.append(mask_idx)
            enc["id"] = batch["id"]
            enc["position"] = batch["position"]
            enc["mask_token_index"] = mask_token_indices
            enc["wt_aa"] = batch["wt_aa"]
            enc["consensus_aa"] = batch["consensus_aa"]
            for aa in AA_COLS:
                enc[aa] = batch[aa]
            return enc

        dataset.set_transform(tokenize)

        input_ids_pad = tokenizer.pad_token_id

        def collate_fn(batch):
            result = {}
            for key in batch[0]:
                vals = [b[key] for b in batch]
                if key == "input_ids":
                    tensors = [torch.tensor(v) for v in vals]
                    result[key] = torch.nn.utils.rnn.pad_sequence(tensors, batch_first=True, padding_value=input_ids_pad)
                elif key == "attention_mask":
                    tensors = [torch.tensor(v) for v in vals]
                    result[key] = torch.nn.utils.rnn.pad_sequence(tensors, batch_first=True, padding_value=0)
                elif key == "position_ids":
                    tensors = [torch.tensor(v) for v in vals]
                    result[key] = torch.nn.utils.rnn.pad_sequence(tensors, batch_first=True, padding_value=0)
                elif isinstance(vals[0], (int, float)):
                    result[key] = torch.tensor(vals)
                else:
                    result[key] = vals
            return result

        dataloader = DataLoader(
            dataset,
            batch_size=args.batch_size,
            shuffle=False,
            collate_fn=collate_fn,
            num_workers=4,
            pin_memory=True,
            prefetch_factor=2,
            persistent_workers=True,
        )

        total_batches = len(dataloader)
        torch.backends.cuda.matmul.allow_tf32 = True
        torch.backends.cudnn.allow_tf32 = True

        buffer = []

        with tqdm(desc=f"KL consensus eval — {args.model_name}", total=total_batches) as pbar:
            for idx, batch in enumerate(dataloader):
                if idx < start_idx:
                    pbar.update(1)
                    continue

                input_ids = batch["input_ids"].to(device)
                attention_mask = batch.get("attention_mask", None)
                if attention_mask is not None:
                    attention_mask = attention_mask.to(device)
                position_ids = batch.get("position_ids", None)
                if position_ids is not None:
                    position_ids = position_ids.to(device)
                mask_token_indices = batch["mask_token_index"]

                with torch.no_grad():
                    try:
                        if position_ids is not None:
                            output = model(input_ids=input_ids, attention_mask=attention_mask, position_ids=position_ids)
                        else:
                            output = model(input_ids=input_ids, attention_mask=attention_mask)
                    except TypeError:
                        output = model(src=input_ids, pad_mask=attention_mask.bool())

                logits = output["logits"]  # [B, seq_len, vocab]

                for i in range(logits.size(0)):
                    mask_idx = int(mask_token_indices[i].item())
                    logit_vec = logits[i, mask_idx].float().cpu().numpy()

                    model_probs = softmax_over_aa(logit_vec, aa_token_ids)
                    pred_aa = AA_COLS[int(np.argmax(model_probs))]

                    wt_aa = batch["wt_aa"][i]
                    consensus_aa = batch["consensus_aa"][i]

                    homolog_freq = np.array([float(batch[aa][i].item()) for aa in AA_COLS])
                    homolog_freq = homolog_freq / homolog_freq.sum()

                    kl = kl_divergence(homolog_freq, model_probs)

                    buffer.append(
                        {
                            "sequence_name": batch["id"][i],
                            "position": int(batch["position"][i].item()),
                            "wt_aa": wt_aa,
                            "consensus_aa": consensus_aa,
                            "pred_aa": pred_aa,
                            "kl": kl,
                        }
                    )

                pbar.update(1)

                flush = (idx > 0 and idx % args.save_steps == 0) or (idx == total_batches - 1)
                if flush:
                    out_file = os.path.join(args.output_path, f"data_{idx:05d}.parquet")
                    pd.DataFrame(buffer).to_parquet(out_file, index=False, compression="gzip")
                    logger.info(f"Saved {out_file}")
                    buffer = []

        with open(os.path.join(args.output_path, "done.txt"), "w") as f:
            f.write("done")

    # ---- Aggregate scores ----
    if args.compute_scores:
        parquet_files = sorted(f for f in os.listdir(args.output_path) if f.startswith("data") and f.endswith(".parquet"))
        df = pd.concat([pd.read_parquet(os.path.join(args.output_path, f)) for f in parquet_files], ignore_index=True)

        wt_is_consensus = df[df["wt_aa"] == df["consensus_aa"]]
        wt_is_not_consensus = df[df["wt_aa"] != df["consensus_aa"]]

        scores = {
            "mean_kl": float(df["kl"].mean()),
            "std_kl": float(df["kl"].std()),
            "median_kl": float(df["kl"].median()),
            # Breakdown by whether WT == consensus
            "mean_kl_wt_is_consensus": float(wt_is_consensus["kl"].mean()),
            "mean_kl_wt_not_consensus": float(wt_is_not_consensus["kl"].mean()),
            # Recovery rates
            "pred_wt_consensus": float((wt_is_consensus["pred_aa"] == wt_is_consensus["wt_aa"]).mean()),
            "pred_other_t": float((wt_is_consensus["pred_aa"] != wt_is_consensus["wt_aa"]).mean()),
            "pred_wt_nt": float((wt_is_not_consensus["pred_aa"] == wt_is_not_consensus["wt_aa"]).mean()),
            "pred_consensus_nt": float((wt_is_not_consensus["pred_aa"] == wt_is_not_consensus["consensus_aa"]).mean()),
            "pred_other_nt": float(
                (
                    (wt_is_not_consensus["pred_aa"] != wt_is_not_consensus["wt_aa"])
                    & (wt_is_not_consensus["pred_aa"] != wt_is_not_consensus["consensus_aa"])
                ).mean()
            ),
            # Counts
            "n_wt_is_consensus": len(wt_is_consensus),
            "n_wt_is_not_consensus": len(wt_is_not_consensus),
            "n_positions": len(df),
        }

        out_scores = os.path.join(args.output_path, "scores.json")
        with open(out_scores, "w") as f:
            json.dump(scores, f, indent=2)
        print(f"Scores saved to {out_scores}")
        print(json.dumps(scores, indent=2))


# ---------------------------------------------------------------------------
# Entry point
# ---------------------------------------------------------------------------

if __name__ == "__main__":
    parser = ArgumentParser(description=__doc__)

    # Model
    parser.add_argument("--model_name", type=str, required=True)
    parser.add_argument("--config_path", type=str, required=False)
    parser.add_argument("--vocab_path", type=str, required=False)
    parser.add_argument("--checkpoint_path", type=str, required=False)
    parser.add_argument("--tag", type=str, required=False)
    parser.add_argument("--token", type=str, default=None)
    parser.add_argument("--batch_size", type=int, default=64)
    parser.add_argument("--bos_token", action="store_true", help="Set if the tokenizer prepends a BOS token (offsets position index by 1).")

    # Data
    parser.add_argument("--proteome_path", type=str, required=False, help="CSV with columns: id, sequence")
    parser.add_argument("--consensus_path", type=str, required=False, help="CSV output of extract_consensus_positions.py")
    parser.add_argument(
        "--dataset_path",
        type=str,
        default=None,
        help="Path to a pre-built HuggingFace Dataset on disk. If it exists, it is loaded directly "
        "(proteome_path and consensus_path are not needed). If it does not exist and this path "
        "is provided, the built dataset is saved here for future reuse.",
    )

    # Output / control
    parser.add_argument("--output_path", type=str, required=True)
    parser.add_argument("--save_steps", type=int, default=500)
    parser.add_argument("--skip_predictions", action="store_true")
    parser.add_argument("--compute_scores", action="store_true")

    args = parser.parse_args()
    evaluate(args)
```
