## Supplementary material for "pLM representations unlock metagenomic space beyond homology": Code bundle: sequence_recovery.py

```
from argparse import ArgumentParser
import os

# HuggingFace
from datasets import load_dataset
from transformers import AutoTokenizer, AutoModel, AutoModelForMaskedLM

import polars as pl
import json

# Ours
from apperitif.evaluation import SequenceRecoveryEvaluator
from amplify.model import AMPLIFY


if __name__ == "__main__":

    parser = ArgumentParser()
    # Model
    parser.add_argument("--model_name", type=str, help="", required=True)
    parser.add_argument("--config_path", type=str, help="", required=False)
    parser.add_argument("--vocab_path", type=str, help="", required=False)
    parser.add_argument("--checkpoint_path", type=str, help="", required=False)
    parser.add_argument("--tag", type=str, help="", required=False)
    parser.add_argument("--batch_size", type=int, default=128, help="", required=False)
    parser.add_argument("--packing", type=bool, default=True, help="", required=False)
    parser.add_argument("--token", default=None, type=str, help="Token to use if the model is private on the hub.")

    # Dataset
    parser.add_argument("--data_path", type=str, help="", required=False)

    # Log
    parser.add_argument("--output_path", type=str, help="", required=True)
    parser.add_argument("--save_steps", type=int, default=10000, help="", required=False)

    # What to run
    parser.add_argument("--skip_predictions", action="store_true", help="")
    parser.add_argument("--compute_scores", action="store_true", help="")
    parser.add_argument("--categories_path", type=str, help="", required=False)

    args = parser.parse_args()

    from_hub = (
        "esm" in args.model_name
        or "AMPLIFY" in args.model_name
        or args.model_name.lower().startswith("<HF_ORG>/")
        or args.model_name.startswith("chandar-lab/")
        or args.model_name.startswith("facebook/")
    )

    if from_hub:
        tokenizer = AutoTokenizer.from_pretrained(args.model_name, trust_remote_code=True, token=args.token)
        if "AMPLIFY" in args.model_name or args.model_name.lower().startswith("<HF_ORG>/"):
            model = AutoModel.from_pretrained(args.model_name, trust_remote_code=True, token=args.token)
        else:
            model = AutoModelForMaskedLM.from_pretrained(args.model_name, trust_remote_code=True, token=args.token)
    else:
        model, tokenizer = AMPLIFY.load(
            checkpoint_path=args.checkpoint_path, config_path=args.config_path, vocab_path=args.vocab_path, tag=args.tag
        )

        if args.tag is not None:
            args.model_name = args.model_name + "_" + args.tag

    evaluator = SequenceRecoveryEvaluator(batch_size=args.batch_size, save_steps=args.save_steps, packing=args.packing)

    if not args.skip_predictions:
        dataset = load_dataset("csv", data_files=args.data_path, split="train")

        evaluator(
            model=model,
            tokenizer=tokenizer,
            dataset=dataset,
            model_name=args.model_name,
            output_path=args.output_path,
        )

    if args.compute_scores:
        if not args.categories_path:
            raise ValueError("Path to categories parquet files needs to be provided to compute scores.")
        cat_70_target = pl.read_parquet(os.path.join(args.categories_path, "cat_70_target.parquet"))
        cat_70_non_target = pl.read_parquet(os.path.join(args.categories_path, "cat_70_non_target.parquet"))
        cat_90_target = pl.read_parquet(os.path.join(args.categories_path, "cat_90_target.parquet"))
        cat_90_non_target = pl.read_parquet(os.path.join(args.categories_path, "cat_90_non_target.parquet"))
        cat_96_target = pl.read_parquet(os.path.join(args.categories_path, "cat_96_target.parquet"))
        cat_96_non_target = pl.read_parquet(os.path.join(args.categories_path, "cat_96_non_target.parquet"))

        homologs_dict = {
            "cat_70_target": cat_70_target,
            "cat_70_non_target": cat_70_non_target,
            "cat_90_target": cat_90_target,
            "cat_90_non_target": cat_90_non_target,
            "cat_96_target": cat_96_target,
            "cat_96_non_target": cat_96_non_target,
        }

        scores = evaluator._compute_metric(
            tokenizer=tokenizer,
            homologs_dict=homologs_dict,
            model_name=args.model_name,
            output_path=args.output_path,
        )

        with open(os.path.join(args.output_path, "scores.json"), "w") as f:
            json.dump(scores, f)
```
