## Supplementary material for "pLM representations unlock metagenomic space beyond homology": Code bundle: create_ppl_sets.py

```
import csv
from argparse import ArgumentParser


if __name__ == "__main__":
    parser = ArgumentParser()
    parser.add_argument("--input_file", type=str, help="", required=True)
    parser.add_argument("--chunk_size", type=int, default=512, help="", required=False)
    parser.add_argument("--max_seq_len", type=int, default=None, help="", required=False)

    args = parser.parse_args()

    if args.max_seq_len is not None:
        output_file_512 = args.input_file.replace(".csv", f"_max_{args.max_seq_len}_512.csv")
        output_file_masked = args.input_file.replace(".csv", f"_max_{args.max_seq_len}_masked.csv")
    else:
        output_file_512 = args.input_file.replace(".csv", f"_512.csv")
        output_file_masked = args.input_file.replace(".csv", f"_masked.csv")

    with open(args.input_file, "r") as fin, open(output_file_512, "w", newline="") as fout:
        reader = csv.DictReader(fin)
        writer = csv.writer(fout, lineterminator="\n")
        writer.writerow(["id", "sequence", "length"])

        for row in reader:
            full_id = (row["name"].split("|")[1] if "|" in row["name"] else row["name"]).strip()
            sequence = row["sequence"]
            if args.max_seq_len and len(sequence) > args.max_seq_len:
                continue
            for i in range(0, len(sequence), args.chunk_size):
                chunk = sequence[i : i + args.chunk_size]
                chunk_id = f"{full_id}_{i // args.chunk_size}"
                if len(chunk) > 5:
                    writer.writerow([chunk_id, chunk, len(chunk)])

    with open(output_file_512, "r", newline="") as fin, open(output_file_masked, "w", newline="") as fout:
        reader = csv.DictReader(fin)
        writer = csv.writer(fout, lineterminator="\n")
        writer.writerow(["sequence", "id", "masked_index", "target"])

        for row in reader:
            seq = row["sequence"]
            seq_id = row["id"]
            for i in range(len(seq)):
                target = seq[i]
                masked_seq = seq[:i] + "<mask>" + seq[i + 1 :]
                writer.writerow([masked_seq, seq_id, i, target])
```
