## Supplementary material for "pLM representations unlock metagenomic space beyond homology": Code bundle: pseudo_perplexity.py

```
import os
import torch
import csv
import math
import argparse

# HuggingFace
from datasets import load_dataset
from argparse import ArgumentParser
from transformers import AutoTokenizer, AutoModel, AutoModelForMaskedLM

# Ours
from apperitif.evaluation import PseudoPPLEvaluator
from amplify.model import AMPLIFY


def compute_pseudo_perplexity(log_probs):
    return math.exp(-sum(log_probs) / len(log_probs))


if __name__ == "__main__":

    parser = ArgumentParser()
    # Model
    parser.add_argument("--model_name", type=str, help="", required=True)
    parser.add_argument("--data_name", type=str, help="", required=True)
    parser.add_argument("--config_path", type=str, help="", required=False)
    parser.add_argument("--vocab_path", type=str, help="", required=False)
    parser.add_argument("--checkpoint_path", type=str, help="", required=False)
    parser.add_argument("--tag", type=str, help="", required=False)
    parser.add_argument("--batch_size", type=int, default=128, help="", required=False)
    parser.add_argument("--not_packing", action="store_true", help="")

    # Dataset
    parser.add_argument("--data_path", type=str, help="", required=False)

    # Log
    parser.add_argument("--output_path", type=str, help="", required=True)
    parser.add_argument("--save_steps", type=int, default=10000, help="", required=False)

    # What to run
    parser.add_argument("--skip_predictions", action="store_true", help="")
    parser.add_argument("--compute_scores", action="store_true", help="")

    args = parser.parse_args()

    # Org-qualified ids (e.g. "<org>/AMPLIFY_350M", "facebook/esm2_...") load from the HF hub;
    # bare base names load from a local checkpoint directory.
    from_hub = "/" in args.model_name or "esm" in args.model_name.lower()
    evaluator = PseudoPPLEvaluator(batch_size=args.batch_size, packing=(not args.not_packing))

    args.tag = None if args.tag == "latest" else args.tag

    if not args.skip_predictions:
        if from_hub:
            tokenizer = AutoTokenizer.from_pretrained(args.model_name, trust_remote_code=True)
            if "esm" not in args.model_name.lower():
                model = AutoModel.from_pretrained(args.model_name, trust_remote_code=True)
            else:
                model = AutoModelForMaskedLM.from_pretrained(args.model_name, trust_remote_code=True)
        else:
            model, tokenizer = AMPLIFY.load(
                checkpoint_path=args.checkpoint_path, config_path=args.config_path, vocab_path=args.vocab_path, tag=args.tag
            )

            if args.tag is not None:
                args.model_name = args.model_name + "_" + args.tag

        device = torch.device("cuda" if torch.cuda.is_available() else "cpu")
        model.to(device)

        dataset = load_dataset("csv", data_files=args.data_path, split="train")

        evaluator(
            model=model,
            tokenizer=tokenizer,
            dataset=dataset,
            model_name=args.model_name,
            data_name=args.data_name,
            output_path=args.output_path,
        )

    if args.compute_scores:
        output_csv = os.path.join(args.output_path, "pseudo_ppl", args.model_name, args.data_name, f"ppl_{args.data_name}_aggregated.csv")

        ppl = evaluator._compute_metric(
            model_name=args.model_name,
            data_name=args.data_name,
            output_path=args.output_path,
        )

        ppl.write_csv(os.path.join(args.output_path, evaluator.name, args.model_name, args.data_name, "ppl.csv"))
```
