## Supplementary material for "pLM representations unlock metagenomic space beyond homology": Code bundle: validation_set_eval.py

```
import os
import yaml
import json
import math
from tqdm import tqdm
from typing import Tuple
from argparse import ArgumentParser, Namespace
import torch
from torch.nn import CrossEntropyLoss
from torch.utils.data import DataLoader

from datasets import load_dataset
from transformers import AutoTokenizer, AutoModel, AutoModelForMaskedLM

from amplify.dataset import get_dataloader
from amplify.dataset.dataloader import CustomDataCollator
from amplify.model import AMPLIFY


def evaluate(
    model: torch.nn.Module,
    dataloader: torch.utils.data.DataLoader,
    loss_fn: torch.nn.modules.loss._Loss,
    vocab_size: int,
    device: torch.device,
    packing: bool,
    from_hub: bool,
    additive_mask: bool = False,
) -> Tuple[int, float, int]:
    """Evaluate the model on the dataloader provided.

    Args:
        model (torch.nn.Module): Model.
        dataloader (torch.utils.data.DataLoader): Dataloader.
        loss_fn (torch.nn.modules.loss._Loss): Loss function.
        vocab_size (int): Total number of tokens in the vocabulary.

    Returns:
        Tuple[int,float,int]: Number of predictions, sum of per-token losses and sum of correct predictions.
    """
    sum_val_loss, num_val_correct, num_val_pred = 0, 0, 0
    with torch.no_grad(), torch.amp.autocast("cuda"):
        for batch in tqdm(dataloader, total=len(dataloader), desc="Evaluating"):
            input_ids, labels, max_seqlen, cu_seqlens, attention_mask, position_ids = (
                batch["input_ids"].to(device),
                batch["labels"].to(device),
                batch.get("max_seqlen", None),
                batch.get("cu_seqlens", None),
                batch.get("attention_mask", None),
                batch.get("position_ids", None),
            )

            if max_seqlen is not None:
                max_seqlen = max_seqlen.to(device)
            if cu_seqlens is not None:
                cu_seqlens = cu_seqlens.squeeze().to(device)
            if attention_mask is not None:
                attention_mask = attention_mask.to(device)
            if position_ids is not None:
                position_ids = position_ids.to(device)

            if packing and from_hub:
                outputs = model(
                    input_ids=input_ids,
                    position_ids=position_ids,
                    max_seqlen=max_seqlen,
                    cu_seqlens=cu_seqlens,
                    attention_mask=attention_mask,
                )
            elif packing:
                outputs = model(
                    input_ids=input_ids,
                    position_ids=position_ids,
                    max_seqlen=max_seqlen,
                    cu_seqlens=cu_seqlens,
                    attention_mask=attention_mask,
                )
            else:
                if additive_mask:
                    attention_mask = torch.where(attention_mask, float(0.0), float("-inf")).type(torch.float16)
                try:
                    outputs = model(input_ids=input_ids, attention_mask=attention_mask)
                except TypeError:
                    outputs = model(src=input_ids, pad_mask=attention_mask)

            logits = outputs.logits

            # Loss over unmasked targets
            loss = loss_fn(logits.view(-1, vocab_size), labels.view(-1))
            valid = labels != -100

            num_preds_batch = valid.sum().item()
            num_val_pred += num_preds_batch
            sum_val_loss += loss.item() * num_preds_batch

            # Accuracy over unmasked targets only
            preds = torch.argmax(logits, dim=-1)
            num_val_correct += (preds[valid] == labels[valid]).sum().item()

    return num_val_pred, sum_val_loss, num_val_correct


def run(args: Namespace):
    try:
        num_workers = len(os.sched_getaffinity(0))
    except AttributeError:
        num_workers = os.cpu_count() or 1

    device = torch.device("cuda" if torch.cuda.is_available() else "cpu")

    packing = bool(args.packing)  # (not "esm" in args.model_name)

    print("packing:", packing)

    # Load model and tokenizer
    from_hub = (
        "esm" in args.model_name
        or "AMPLIFY" in args.model_name
        or args.model_name.lower().startswith("<HF_ORG>/")
        or args.model_name.startswith("chandar-lab/")
        or args.model_name.startswith("facebook/")
    )
    if from_hub:
        tokenizer = AutoTokenizer.from_pretrained(args.model_name, trust_remote_code=True, token=args.token)
        if "AMPLIFY" in args.model_name or args.model_name.lower().startswith("<HF_ORG>/"):
            model = AutoModel.from_pretrained(args.model_name, trust_remote_code=True, token=args.token)
        else:
            model = AutoModelForMaskedLM.from_pretrained(args.model_name, trust_remote_code=True, token=args.token)
    else:
        model, tokenizer = AMPLIFY.load(args.checkpoint_path, args.config_path, args.vocab_path)
    model.to(device)
    model.eval()

    collator = CustomDataCollator(
        tokenizer=tokenizer,
        mlm=True,
        mlm_probability=float(args.mask_probability),
        return_tensors="pt",
        pack_sequences=packing,
    )
    dataset = load_dataset(
        "csv",
        data_files=args.data_paths,
        keep_in_memory=False,
        num_proc=num_workers,
        split="train",
    )

    def transform(inputs):
        return tokenizer(
            inputs["sequence"],
            max_length=args.max_len,
            padding=False,
            truncation=True,
            return_tensors=None,
        )

    dataset.set_transform(transform)

    dataloader = DataLoader(
        dataset=dataset,
        batch_size=args.batch_size,
        shuffle=False,
        collate_fn=collator,
        num_workers=num_workers,
        prefetch_factor=4 if (num_workers > 1) else None,
        pin_memory=(device.type == "cuda"),
        persistent_workers=(num_workers > 1),
    )

    # Loss function
    val_loss_fn = CrossEntropyLoss(reduction="mean", ignore_index=-100)

    num_val_pred, sum_val_loss, num_val_correct = evaluate(
        model,
        dataloader,
        val_loss_fn,
        tokenizer.vocab_size,
        packing=packing,
        device=device,
        from_hub=from_hub,
        additive_mask=args.additive_mask,
    )

    # Metrics
    metrics = {
        "val_loss": (sum_val_loss / num_val_pred) if num_val_pred > 0 else float("nan"),
        "val_perplexity": math.exp(sum_val_loss / num_val_pred) if num_val_pred > 0 else float("inf"),
        "val_accuracy": (num_val_correct / num_val_pred) if num_val_pred > 0 else float("nan"),
        "num_tokens": num_val_pred,
    }

    # Save metrics in json
    os.makedirs(os.path.dirname(args.output_path), exist_ok=True)
    with open(args.output_path, "w") as f:
        json.dump(metrics, f, indent=4)


if __name__ == "__main__":
    parser = ArgumentParser()
    parser.add_argument("--data_paths", nargs="+", required=True, help="Path to the csv file or files to evaluate on.")
    parser.add_argument("--token", type=str, help="HF token.")
    parser.add_argument("--model_name", type=str, required=True, help="Name of the model.")
    parser.add_argument("--checkpoint_path", type=str, help="Path to the local model checkpoint (if not using --model_name).")
    parser.add_argument("--config_path", type=str, help="Path to the model config file.")
    parser.add_argument("--vocab_path", type=str, help="Path to the tokenizer vocabulary file.")
    parser.add_argument("--output_path", type=str, required=True, help="Path to the output json file.")

    parser.add_argument("--mask_probability", type=float, required=False, default=0.15)
    parser.add_argument("--batch_size", type=int, required=False, default=16)
    parser.add_argument("--max_len", type=int, required=False, default=512)
    parser.add_argument("--packing", type=int, required=False, default=1, help="Wether or not to pack sequences. 0 for False, 1 for True.")
    parser.add_argument("--additive_mask", action="store_true", help="Wether or not to use additive mask instead of boolean mask.")

    args = parser.parse_args()
    run(args)
```
