## Supplementary material for "pLM representations unlock metagenomic space beyond homology": Code bundle: main.py

```
"""
Main script for running AMPLIFY evaluation tasks.
This script dynamically constructs evaluator configurations based on the task type.
"""

import logging
from pathlib import Path
import hydra
from omegaconf import DictConfig, OmegaConf
from apperitif.tasks.AbsTask import TaskConfig
from apperitif.utils.config_utils import resolve_evaluator_target
from apperitif.tasks import Task

logging.basicConfig(level=logging.INFO)
logger = logging.getLogger(__name__)


def construct_evaluator_config(cfg: DictConfig) -> DictConfig:
    """
    Construct the evaluator configuration based on the task type from data_config.

    Args:
        cfg: The main configuration object

    Returns:
        DictConfig: The constructed evaluator configuration
    """
    # Get the task type from data_config
    task_type = cfg.data_config.task_type
    logger.info(f"Task type detected: {task_type}")

    # Load the base evaluator config
    base_evaluator_cfg = cfg.get(task_type, None)
    if base_evaluator_cfg is None:
        logger.error(f"Unknown task type: {task_type}. Please check your configuration.")
        raise ValueError(f"Unknown task type: {task_type}")

    # Load the head type config
    head_type = cfg.head_type
    logger.info(f"Using head type: {head_type}")
    head_type_cfg = {"head_type": head_type}

    # Load the grid config
    grid_cfg = cfg.grid

    # Convert to regular dict to avoid struct mode issues, then merge
    base_dict = OmegaConf.to_container(base_evaluator_cfg, resolve=True)
    head_type_dict = head_type_cfg  # Already a dict
    grid_dict = OmegaConf.to_container(grid_cfg, resolve=True)

    # Merge all configurations
    merged_dict = {**base_dict, **head_type_dict, **grid_dict}
    merged_dict["_target_"] = resolve_evaluator_target(
        task_type=task_type,
        head_type=head_type,
    )
    evaluator_config = OmegaConf.create(merged_dict)
    return evaluator_config


@hydra.main(version_base=None, config_path="../conf", config_name="config")
def main(cfg: DictConfig):
    """
    Main function for running AMPLIFY evaluation tasks.

    Args:
        cfg: Hydra configuration object
    """
    logger.info("Starting AMPLIFY evaluation...")
    logger.info("=" * 60)

    # Log the initial configuration
    logger.info("Initial configuration loaded:")
    # logger.info(f"  Name: {cfg.name}")
    # logger.info(f"  Device: {cfg.device}")
    # logger.info(f"  Head type: {cfg.head_type}")
    # logger.info(f"  Grid: {cfg.grid}")
    # logger.info(f"  Do test: {cfg.do_test}")
    # logger.info(f"  Return all results: {cfg.return_all_results}")

    # Instantiate the data config
    logger.info("=" * 60)
    logger.info("Instantiating data configuration...")
    data_config = hydra.utils.instantiate(cfg.data_config)
    logger.info(f"✓ Data config instantiated: {type(data_config).__name__}")
    logger.info(f"  Task type: {data_config.task_type}")
    logger.info(f"  Dataset name: {data_config.dataset_name}")

    # Instantiate the model config
    logger.info("=" * 60)
    logger.info("Instantiating model configuration...")
    model_config = hydra.utils.instantiate(cfg.model)
    logger.info(f"Model config instantiated: {type(model_config).__name__}")
    logger.info(f"  Model name: {model_config.model_name}")
    logger.info(f"  Model type: {model_config.model_type}")

    # Construct the evaluator configuration
    logger.info("=" * 60)
    logger.info("Constructing evaluator configuration...")

    # Extract subbenchmark_name from grid config and remove it
    subbenchmark_name = cfg.grid.get("subbenchmark_name", "default")

    # Create a new grid config without subbenchmark_name
    grid_dict = OmegaConf.to_container(cfg.grid, resolve=True)
    if "subbenchmark_name" in grid_dict:
        del grid_dict["subbenchmark_name"]
    cleaned_grid_cfg = OmegaConf.create(grid_dict)
    cfg.grid = cleaned_grid_cfg
    evaluator_config = construct_evaluator_config(cfg)
    evaluator_config = hydra.utils.instantiate(evaluator_config)
    if "precomputing_batch_size" in cfg:
        evaluator_config.precomputing_batch_size = cfg.precomputing_batch_size  # Override precomputing batch size if specified
    logger.info(f"Evaluator config instantiated: {type(evaluator_config).__name__}")

    # Resolving the task name
    if cfg.name is None:
        # Use the task type from data_config as the default name
        name = data_config.dataset_name.lower()
        name = name.replace("/", "_")
        name = name.replace("-", "_")
        cfg.name = name
        logger.info(f"Task name resolved to: {cfg.name}")

    # Resolving the output folder
    if cfg.output_folder is None:
        model_name = model_config.local_name if model_config.local_name is not None else model_config.model_name.replace("/", "_")
        if model_config.randomize:
            if model_config.randomize_mode != "distribution":
                model_name += f"_{model_config.randomize_mode}"
            else:
                model_name += "_randomized"

        print("model_name", model_name)
        # subbenchmark_name corresponds to the grid type (finetuning, feature_extraction, restricted or not)
        cfg.output_folder = Path(cfg.base_folder) / subbenchmark_name / cfg.name / model_name
        cfg.output_folder.mkdir(parents=True, exist_ok=True)  # Ensure the output folder exists
        logger.info(f"Output folder resolved to: {cfg.output_folder}")

    # Create the TaskConfig object
    logger.info("=" * 60)
    logger.info("Creating TaskConfig object...")

    task_config = TaskConfig(
        data_config=data_config,
        evaluator_config=evaluator_config,
        model_config=model_config,
        name=cfg.name,
        device=cfg.device,
        output_folder=cfg.output_folder,
        return_all_results=cfg.return_all_results,
        do_test=cfg.do_test,
        verbose=cfg.verbose,  # Use the verbose setting from the config
        accumulation_steps=cfg.accumulation_steps if "accumulation_steps" in cfg else None,  # Use the accumulation steps from the config
    )
    # Instantiate the actual task using the from_config method
    logger.info("Instantiating task using Task.from_config...")

    task = Task.from_config(task_config)
    logger.info(f"✓ Task instantiated: {type(task).__name__}")

    logger.info("=" * 60)
    logger.info("Configuration setup completed successfully!")
    logger.info(f"TaskConfig summary:")
    logger.info(f"  Data config type: {type(task_config.data_config).__name__}")
    logger.info(f"  Model config type: {type(task_config.model_config).__name__}")
    logger.info(f"  Evaluator head type: {task_config.evaluator_config.head_type}")
    logger.info(f"  Task type: {task_config.data_config.task_type}")
    logger.info(f"  Task name: {task_config.name}")
    logger.info(f"  Device: {task_config.device}")
    logger.info(f"  Output folder: {task_config.output_folder}")
    logger.info(f"  Task instance: {type(task).__name__}")

    task.evaluate()


if __name__ == "__main__":
    main()
```
