## Supplementary material for "pLM representations unlock metagenomic space beyond homology": Code bundle: prepare_plot_df.py

```
"""
Aggregate evaluation results and build plot_df.csv for the paper figures notebook.

Usage:
    python prepare_plot_df.py [--output OUTPUT]

Runs upstream aggregation scripts, loads all result CSVs, merges them into a
single dataframe and saves it as a CSV.
"""
import argparse
import json
import subprocess
import sys
from pathlib import Path

import numpy as np
import pandas as pd

SCRATCH = "$SCRATCH/proseqo"
REPO = "."
DEFAULT_OUTPUT = Path("notebooks/data/plot_df.csv")

# ── Aggregation scripts ───────────────────────────────────────────────────────

AGGREGATE_SCRIPTS = [
    (
        f"{REPO}/scripts/eval_consensus/aggregate_results.py",
        f"{SCRATCH}/sequence_recovery",
        f"{SCRATCH}/sequence_recovery/consensus_fidelity.csv",
    ),
    (
        f"{REPO}/scripts/eval_categorical_jacobian/aggregate_results.py",
        f"{SCRATCH}/categorical_jacobian/",
        f"{SCRATCH}/categorical_jacobian/categorical_jacobian.csv",
    ),
    (
        f"{REPO}/scripts/eval_validation/aggregate_results.py",
        f"{SCRATCH}/uniprot_validation",
        f"{SCRATCH}/uniprot_validation/uniprot_validation.csv",
    ),
    (
        f"{REPO}/scripts/eval_consensus/aggregate_kl_consensus.py",
        f"{SCRATCH}/kl_consensus",
        f"{SCRATCH}/kl_consensus/kl_consensus_results.csv",
    ),
]


def run_aggregation_scripts():
    for script, input_dir, output_path in AGGREGATE_SCRIPTS:
        print(f"Running {Path(script).name} ...")
        result = subprocess.run(
            [sys.executable, script, input_dir, "-o", output_path],
            capture_output=True, text=True,
        )
        if result.returncode != 0:
            print(f"  WARNING: {Path(script).name} exited with code {result.returncode}")
            print(result.stderr[-500:] if result.stderr else "")
        else:
            print(f"  OK -> {output_path}")


# ── Helper functions (previously in notebook cell 4) ─────────────────────────

ESM_NAMES = {
    "esm2_t12_35M_UR50D": "esm_35M",
    "esm2_t30_150M_UR50D": "esm_150M",
    "esm2_t33_650M_UR50D": "esm_650M",
    "esm_flash_150M": "esm_150M",
    "esm_flash_650M": "esm_650M",
}


def format_model_name(model_name):
    model_name = (
        model_name
        .replace("MGNIFY", "MGnify")
        .replace("U50", "UR50")
        .replace("amplify_", "AMPLIFY_")
        .replace("_packed", "")
        .replace("_0_", "_0.")
    )
    return ESM_NAMES.get(model_name, model_name)


def load_contact_results(base_path):
    records = []
    for results_file in Path(base_path).rglob("results.json"):
        model_dir = results_file.parent
        with open(results_file) as f:
            data = json.load(f)
        best_hp, best_result = max(
            data["all_results"].items(), key=lambda x: x[1]["validation_score"]
        )
        record = {"model_name": model_dir.name}
        for split in ["test_scores", "casp16_scores"]:
            prefix = split.replace("_scores", "")
            for k, v in best_result.get(split, {}).items():
                record[f"{prefix}_{k}"] = v
        records.append(record)

    contact = pd.DataFrame(records)

    for prefix, keyword in [("test", "P@L"), ("test", "long_P@L"), ("test", "AUC"), ("casp", "P@L"), ("casp", "AUC")]:
        cols = [c for c in contact.columns if c.startswith(f"{prefix}_") and keyword in c and "avg" not in c]
        contact[f"{prefix}_avg_{keyword}"] = contact[cols].mean(axis=1)

    return contact


def prepare_xtrimo(input_df, prefix):
    df = input_df.copy()
    for suffix in ["_restricted", "_extended", ""]:
        f1_col = f"best_test_f1{suffix}"
        sp_col = f"best_test_spearmanr{suffix}"
        if f1_col in df.columns or sp_col in df.columns:
            df[f"{prefix}{suffix}"] = df.apply(
                lambda row: (
                    row[f1_col] if not pd.isna(row.get(f1_col, float("nan"))) else row.get(sp_col, float("nan"))
                ),
                axis=1,
            )
    df["model_name"] = df["model"].apply(format_model_name)
    df = df.groupby(["model_name", "task"]).max()
    df.reset_index(inplace=True)

    avg = df.groupby("model_name").mean(numeric_only=True).reset_index().copy()
    pct_col = [c for c in avg.columns if "pct_completed" in c or "num_completed" in c]
    avg.rename(columns={c: f"{prefix}_{c}" for c in pct_col}, inplace=True)
    avg = avg[["model_name"] + [c for c in avg.columns if c.startswith(prefix)]]

    tasks = df.pivot(
        index="model_name",
        columns="task",
        values=[f"{prefix}{s}" for s in ["_restricted", "_extended", ""] if f"{prefix}{s}" in df.columns],
    )
    tasks.columns = ["_".join(col).replace("biomap_research_", "").strip() for col in tasks.columns.values]
    tasks = tasks.reset_index()

    for suffix in ["_restricted", "_extended", ""]:
        q3 = f"{prefix}{suffix}_ssp_q3"
        q8 = f"{prefix}{suffix}_ssp_q8"
        if q3 in tasks.columns and q8 in tasks.columns:
            tasks[f"{prefix}{suffix}_ssp"] = tasks[[q3, q8]].mean(axis=1)

    tasks = tasks[["model_name"] + [c for c in tasks.columns if c.startswith(f"{prefix}_")]]
    return avg.merge(tasks, on="model_name", how="outer")


# ── Loading and merging (previously in notebook cell 6) ──────────────────────

def build_plot_df():
    print("Loading results ...")

    contact = load_contact_results(
        "$SCRATCH/proseqo/apperitif/"
        "mlp_feature_extraction/davidhd_contact_prediction_binary_casp16"
    )
    contact["model_name"] = contact["model_name"].apply(format_model_name)

    jacobian = pd.read_csv(f"{SCRATCH}/categorical_jacobian/categorical_jacobian.csv")
    jacobian["model_name"] = jacobian["model_name"].apply(format_model_name)

    val = pd.read_csv(f"{SCRATCH}/uniprot_validation/uniprot_validation.csv")
    val = val[val.max_length != "max_len_2048"]
    val = val[["model_name", "val_accuracy"]].groupby("model_name").mean(numeric_only=True).reset_index()
    val["model_name"] = val["model_name"].str.replace("_0_", "_0.")
    val["model_name"] = val["model_name"].apply(format_model_name)

    consensus = pd.read_csv(f"{SCRATCH}/sequence_recovery/consensus_fidelity.csv")
    consensus = consensus[[
        "model_name",
        "cat_70_target", "cat_70_non_target",
        "cat_90_target", "cat_90_non_target",
        "cat_96_target", "cat_96_non_target",
    ]]
    consensus["model_name"] = consensus["model_name"].apply(format_model_name)

    kl_consensus = pd.read_csv(f"{SCRATCH}/kl_consensus/kl_consensus_results.csv")
    kl_consensus["model_name"] = kl_consensus["model"].apply(format_model_name)

    prop = pd.read_csv(f"{SCRATCH}/apperitif/mlp_feature_extraction_aggregated.csv")
    fe = prepare_xtrimo(prop, "fe")

    prop2 = pd.read_csv(f"{SCRATCH}/apperitif/mlp_finetuning_extended_aggregated.csv")
    ft = prepare_xtrimo(prop2, "ft")

    dms = pd.read_csv(
        f"{SCRATCH}/proteingym/results/performance/all_models/Spearman/"
        "Summary_performance_DMS_substitutions_Spearman.csv"
    )
    dms = dms[["Model_name", "Average_Spearman", "Bootstrap_standard_error_Spearman"]]
    dms.columns = ["model", "avg_spearman", "bootstrap_standard_error_spearman"]
    dms["model_name"] = (
        dms["model"]
        .apply(lambda x: x
            .replace("AMPLIFY_2_", "")
            .replace("120M_MILA", "MILA")
            .replace("MGNIFY", "MGnify")
            .replace("AMPLIFY_120M_hf_mm", "AMPLIFY_120M")
            .replace("AMPLIFY_350M_hf_mm", "AMPLIFY_350M")
            .replace("350M_consensus", "AMPLIFY_consensus")
            .replace("120M_cluster", "cluster")
            .replace("120M_amplify_opt_", "amplify_opt_")
            .replace("_0_", "_0.")
        )
        .apply(lambda x: ESM_NAMES.get(x, x))
    )
    dms.sort_values("model_name", inplace=True)

    # Normalise consensus model name across all sources
    for ds in [fe, ft, consensus, val, dms, jacobian, kl_consensus]:
        if "AMPLIFY_consensus_extended" in ds["model_name"].values:
            ds["model_name"] = ds["model_name"].replace("AMPLIFY_consensus", "AMPLIFY_consensus_short")
            ds["model_name"] = ds["model_name"].replace("AMPLIFY_consensus_extended", "AMPLIFY_consensus")

    df = val.merge(consensus, on="model_name", how="outer")
    df = df.merge(contact, on="model_name", how="outer")
    df = df.merge(fe, on="model_name", how="outer")
    df = df.merge(ft, on="model_name", how="outer")
    df = df.merge(dms, on="model_name", how="outer")
    df = df.merge(jacobian, on="model_name", how="outer")
    df = df.merge(kl_consensus, on="model_name", how="outer")

    parts = df["model_name"].str.split("_")
    df["1m"] = parts.str[-1] == "1m"
    df["dataset"] = parts.str[1].fillna("")

    def get_filter(x):
        for part in str(x).split("_"):
            if part.replace(".", "", 1).isdigit():
                val = float(part)
                if 0 < val < 1:
                    return val
        return 0

    df["filtering"] = df["model_name"].apply(get_filter)
    df = df[~df["model_name"].isna()]

    # ── Metric renaming map ───────────────────────────────────────────────────
    metrics_to_plot = {
        "val_accuracy": "Sequence recovery",
        "cat_70_non_target": "Consensus fidelity",
        "fe_extended": "Property (FE, ext.)",
        "fe_restricted": "Property (FE, rest.)",
        "fe": "Property (FE)",
        "ft": "Property (FT)",
        "avg_spearman": "Deep mutational scanning",
        "test_avg_P@L": "Contact prediction (avg. P@L)",
        "test_long_P@L": "Contact prediction (long P@L)",
        "test_long_P@L2": "Contact prediction (long P@L2)",
        "test_long_P@L5": "Contact prediction (long P@L5)",
        "test_avg_long_P@L": "Contact prediction (avg. long P@L)",
        "AUC": "Categorical jacobian (CASP16)",
        "mean_kl": "KL homologs (mean)",
        "std_kl": "KL homologs (std)",
        "mean_kl_wt_is_consensus": "KL homologs (T)",
        "mean_kl_wt_not_consensus": "KL homologs (NT)",
        "pred_wt_consensus": "Consensus prediction (T)",
        "pred_consensus_nt": "Consensus prediction (NT)",
        "pred_wt_nt": "WT prediction (NT)",
    }
    for col in list(ft.columns) + list(fe.columns):
        if (col.startswith("fe") or col.startswith("ft")) and col not in metrics_to_plot:
            tmp = col.replace("_prediction", "").replace("_", " ").title()
            items = tmp.split(" ")
            if len(items) == 1:
                continue
            elif items[1] in ["Restricted", "Extended"]:
                f, grid = items[0], items[1]
                task = " ".join(items[2:])
            else:
                f, grid = items[0], None
                task = " ".join(items[1:])
            metric_name = f"{task} ({f.upper()}, {grid})" if grid else f"{task} ({f.upper()})"
            metric_name = (
                metric_name
                .replace("Ssp", "SSP")
                .replace("Ph", "pH")
                .replace("Tcr Pmhc", "TCR pMHC")
                .replace("Hla Mhc", "HLA MHC")
            )
            metrics_to_plot[col] = metric_name

    for k, v in metrics_to_plot.items():
        if k in df.columns:
            df[v] = df[k] * (1 if v == "Consensus fidelity" or "KL" in v else 100)

    plot_df = df[["model_name", "1m", "filtering", "dataset"] + list(metrics_to_plot.values())].copy()
    plot_df["Calibration"] = 100 * np.exp(-plot_df["KL homologs (mean)"])
    plot_df["Calibration (T)"] = 100 * np.exp(-plot_df["KL homologs (T)"])
    plot_df["Calibration (NT)"] = 100 * np.exp(-plot_df["KL homologs (NT)"])

    return plot_df


# ── Entry point ───────────────────────────────────────────────────────────────

def main():
    parser = argparse.ArgumentParser(description=__doc__)
    parser.add_argument("--output", default=str(DEFAULT_OUTPUT), help="Output CSV path")
    parser.add_argument("--skip-aggregation", action="store_true",
                        help="Skip running upstream aggregation scripts")
    args = parser.parse_args()

    if not args.skip_aggregation:
        run_aggregation_scripts()

    plot_df = build_plot_df()

    output = Path(args.output)
    output.parent.mkdir(parents=True, exist_ok=True)
    plot_df.to_csv(output, index=False)
    print(f"\nSaved plot_df ({len(plot_df)} rows, {len(plot_df.columns)} columns) -> {output}")


if __name__ == "__main__":
    main()
```
