## Supplementary material for "pLM representations unlock metagenomic space beyond homology": Code bundle: token_umap.py

```
import csv
import os
from argparse import ArgumentParser

# from cuml import UMAP
from umap import UMAP
import pandas as pd
import numpy as np
import joblib
from tqdm import tqdm
import time
import pickle


def load_df(score_path, sequences_path):
    df = []
    for file in os.listdir(score_path):
        if file.endswith(".parquet"):
            df.append(pd.read_parquet(os.path.join(score_path, file)))

    output = pd.concat(df)

    with open(sequences_path, "rt") as f:
        output_sequences = pd.read_csv(f)

    output = output.merge(output_sequences, left_on="name", right_on="name", how="inner")
    output["length"] = output["sequence"].apply(lambda x: len(x))

    return output


def project_df(df, reducer, embedding_dim=2048):
    output = {}

    for _, row in df.iterrows():
        token_embbedings = []
        for i in range(embedding_dim):
            token_emb = row.get(f"embedding_{i}")
            if token_emb is not None:
                token_embbedings.append(token_emb)
        if token_embbedings:
            token_projections = reducer.transform(token_embbedings)
            output[row.get("name")] = {
                "pseudo_ppl": row.get("pseudo_ppl"),
                "mean_cosine_sim": row.get("mean_cosine_sim"),
                "std_cosine_sim": row.get("std_cosine_sim"),
                "token_projections": token_projections,
            }

    return output


def compute_umap(args):
    df = load_df(
        args.score_path,
        args.sequences_path,
    )

    # Collect all token embeddings and metadata
    all_embeddings = []
    text_residue = []
    positions = []
    ids = []

    for idx, row in tqdm(df.iterrows(), total=len(df)):
        token_emb = row.get(args.column_key)[1:-1]  # We skip the BOS and EOS token
        sequence = list(row["sequence"][: len(token_emb)])
        all_embeddings.extend(token_emb)
        text_residue.extend(sequence)
        positions.extend(list(range(len(token_emb))))
        ids.extend([row["name"]] * len(token_emb))

    idx = np.random.choice(len(all_embeddings), size=args.n_samples, replace=False)
    X_sampled = np.array(all_embeddings)[idx]

    matched_residue = np.array(text_residue)[idx]
    matched_position = np.array(positions)[idx]
    matched_id = np.array(ids)[idx]
    np.save(os.path.join(args.output_path, "matched_residue.npy"), matched_residue)
    np.save(os.path.join(args.output_path, "matched_position.npy"), matched_position)
    np.save(os.path.join(args.output_path, "matched_id.npy"), matched_id)

    del text_residue, all_embeddings, matched_residue

    # Fit UMAP
    start_time = time.time()
    umap_model = UMAP(n_neighbors=15, min_dist=0.1, metric="cosine", random_state=42)
    X_umap = umap_model.fit_transform(X_sampled)

    end_time = time.time()
    print(f"Project the data in 2D with UMAP ({end_time - start_time:.2f}s)")

    joblib.dump(umap_model, os.path.join(args.output_path, "umap_reducer.joblib"))
    np.save(os.path.join(args.output_path, "X_umap.npy"), X_umap)


def reduce_embeddings(reducer):
    reducer = joblib.load("$SCRATCH/proseqo/visuals/umap_reducer.joblib")

    df = load_df(
        args.score_path,
        args.sequences_path,
    )

    projections = project_df(df, reducer)

    with open(f"$SCRATCH/proseqo/visuals/{args.output_name}.pkl", "wb") as f:
        pickle.dump(projections, f)


if __name__ == "__main__":

    args = ArgumentParser()
    args.add_argument(
        "--score_path", type=str, default="$SCRATCH/proseqo/scores/Embedding-scores/uniref100_train_10k"
    )
    args.add_argument(
        "--sequences_path",
        type=str,
        default="$SCRATCH/proseqo/datasets/uniref100_train_10k.csv",
    )
    args.add_argument("--output_path")
    args.add_argument("--n_samples", type=int, default=1000000)
    args.add_argument("--column_key", type=str, default="token_embeddings_per_layer")
    args.add_argument("--output_name", type=str, default="token_umap_projections")
    args = args.parse_args()

    os.makedirs(args.output_path, exist_ok=True)

    compute_umap(args)
```
