## Supplementary material for "pLM representations unlock metagenomic space beyond homology": Code bundle: __init__.py

```
__all__ = [
    "CustomCollatorForSeqClassificationWithPacking",
    "DataCollatorForSeqClassification",
    "DataCollatorWithPacking",
]
from .Collator import (
    CustomCollatorForSeqClassificationWithPacking,
    DataCollatorForSeqTasks,
    DataCollatorWithPacking,
)
```
