## Supplementary material for "pLM representations unlock metagenomic space beyond homology": Code bundle: Collator.py

```
import torch
import numpy as np

from abc import ABC, abstractmethod

from transformers import PreTrainedTokenizerFast, DataCollatorForLanguageModeling, DefaultDataCollator


class CustomCollator(ABC):
    """
    Custom collate function for the DataLoader. It takes a list of samples and returns a batch.
    """

    def __init__(
        self,
        sequence_column: str,
        tokenizer: PreTrainedTokenizerFast,
        max_length: int,
    ):
        self.sequence_column = sequence_column
        self.tokenizer = tokenizer
        self.max_length = max_length

    @abstractmethod
    def __call__(self, inputs):
        pass


class DataCollatorWithPacking(DefaultDataCollator):
    def __init__(self, **kwargs):
        super().__init__(**kwargs)

    def __call__(self, batch):
        # Add position_ids if not present
        if "position_ids" not in batch[0]:
            for item in batch:
                item["position_ids"] = list(range(len(item["input_ids"])))

        # Pack the sequences into a single list
        input_ids_list = [item["input_ids"] for item in batch]
        position_ids_list = [item["position_ids"] for item in batch]
        seqlens = np.array([0] + [len(ids) for ids in input_ids_list])

        packed_batch = {
            "position_ids": np.concatenate(position_ids_list, axis=0),
            "input_ids": np.concatenate(input_ids_list, axis=0),
            "cu_seqlens": np.cumsum(seqlens),
            "max_seqlen": max(seqlens),
        }

        output_batch = super().__call__([packed_batch])
        output_batch["cu_seqlens"] = output_batch["cu_seqlens"].to(torch.int32).squeeze()
        output_batch["masked_index"] = torch.tensor([x["masked_index"] for x in batch], dtype=torch.int32)
        if "id" in batch[0].keys():
            output_batch["id"] = [x["id"] for x in batch]
        if "target" in batch[0].keys():
            output_batch["target"] = [x["target"] for x in batch]

        return output_batch


# class CustomCollatorForSeqRecovery(CustomCollator):

#     def __call__(self, inputs):
#         input_ids, position_ids, targets = torch.tensor([]), torch.tensor([]), torch.tensor([])
#         seqlens = [0]

#         # Tokenize the inputs
#         features = self.tokenizer(
#             [input[self.sequence_column] for input in inputs],
#             padding=False,
#             truncation=False,
#             max_length=self.max_length,
#             return_tensors=None,
#             remove_ambiguous=True,
#         )

#         for i in range(len(features["input_ids"])):
#             # Sequence length
#             seqlen = len(features["input_ids"][i])

#             # Repeat the tensors for the desired batch size
#             repeated_input_ids = torch.tensor([features["input_ids"][i]] * seqlen).to(torch.long)
#             repeated_position_ids = torch.tensor([features["position_ids"][i]] * seqlen).to(torch.long)

#             # Apply masking
#             mask = torch.eye(seqlen, dtype=torch.bool)

#             masked_input_ids = torch.where(mask, self.tokenizer.mask_token_id, repeated_input_ids)
#             repeated_target = torch.where(mask, repeated_input_ids, -100)

#             # Concatenate features
#             input_ids = torch.cat((input_ids, masked_input_ids.flatten()), dim=0)
#             position_ids = torch.cat((position_ids, repeated_position_ids.flatten()), dim=0)
#             targets = torch.cat((targets, repeated_target.flatten()), dim=0)
#             seqlens.extend([seqlen] * seqlen)

#         return {
#             "input_ids": input_ids.to(torch.long).unsqueeze(0),
#             "targets": targets.to(torch.long).unsqueeze(0),
#             "position_ids": position_ids.to(torch.long).unsqueeze(0),
#             "cu_seqlens": torch.tensor(np.cumsum(seqlens), dtype=torch.int32),
#             "max_seqlen": max(seqlens),
#             "names": [input[self.name_column] for input in inputs],
#         }


# class CustomCollatorWithoutPacking(CustomCollator): TODO decide if it's still needed

#     def __call__(self, inputs):
#         input_ids= []

#         # Tokenize the inputs
#         padding = len(inputs) > 1
#         features = self.tokenizer(
#             [input[self.sequence_column] for input in inputs],
#             padding=padding,
#             truncation=True,
#             max_length=self.max_length,
#             pad_to_multiple_of=8,
#             return_tensors="pt",
#             remove_ambiguous=True,
#         )

#         # Repeat the input sequences
#         input_ids = features["input_ids"].repeat_interleave(self.repeat_size, dim=0)
#         attention_mask = features["attention_mask"].repeat_interleave(self.repeat_size, dim=0).to(torch.bool)

#         # Apply random masking for each row except the first of each sample
#         mask = (torch.rand(input_ids.shape, dtype=torch.float32) < self.mask_probability) & attention_mask
#         for i in range(len(features["input_ids"])):
#             mask[i * self.repeat_size] = 0

#         masked_input_ids = torch.where(mask, self.tokenizer.mask_token_id, input_ids)

#         return {
#             "input_ids": masked_input_ids,
#             "attention_mask": attention_mask if padding else None,
#             "names": [input[self.name_column] for input in inputs],
#         }


class CustomCollatorForSeqClassificationWithPacking(CustomCollator):
    """
    Custom collate function for the DataLoader in a sequence classification task with packing.
    It takes a list of samples and returns a packed batch of sequences with labels.
    """

    def __init__(
        self,
        sequence_column: str,
        tokenizer: PreTrainedTokenizerFast,
        max_length: int,
        label_column: str,
        device: torch.device = None,
        label_dtype: torch.dtype = torch.long,
    ):
        super().__init__(sequence_column, tokenizer, max_length)
        self.label_column = label_column
        self.device = device
        self.label_dtype = label_dtype

    def __call__(self, inputs):
        input_ids = []
        position_ids = []
        seqlens = [0]
        labels = []

        # Tokenize the inputs
        features = self.tokenizer(
            [input[self.sequence_column] for input in inputs],
            padding=False,
            truncation=True,
            max_length=self.max_length,
            return_tensors=None,
            remove_ambiguous=True,
        )

        for i in range(len(features["input_ids"])):
            input_id = torch.tensor(features["input_ids"][i]).to(torch.long)
            position_id = torch.arange(len(input_id)).to(torch.long)

            # Sequence length
            seqlen = len(input_id)

            # Append features
            input_ids.append(input_id)
            position_ids.append(position_id)
            seqlens.append(seqlen)

            # Append label
            labels.append(inputs[i][self.label_column])

        input_ids = torch.cat(input_ids, dim=0)
        position_ids = torch.cat(position_ids, dim=0)

        if self.device:
            input_ids = input_ids.to(self.device)
            position_ids = position_ids.to(self.device)
            labels = torch.tensor(labels, dtype=self.label_dtype).to(self.device)
            cu_seqlens = torch.tensor(np.cumsum(seqlens), dtype=torch.int32).to(self.device)
        else:
            labels = torch.tensor(labels, dtype=self.label_dtype)
            cu_seqlens = torch.tensor(np.cumsum(seqlens), dtype=torch.int32)

        return {
            "input_ids": input_ids.unsqueeze(0),
            "position_ids": position_ids.unsqueeze(0),
            "cu_seqlens": cu_seqlens,
            "max_seqlen": max(seqlens),
            "labels": labels,
        }


# TODO truncation!! ?? --> We have no random truncation, so we know each sequence starts with the BOS token.
# TODO shape of labels. to avoid squeezes and everything.
# TODO shape of labels and input_ids and others for token level tasks
# TODO have some more beautiful code LOL (factorize some parts in classes)


class DataCollatorForSeqTasks(DataCollatorForLanguageModeling):
    def __init__(
        self,
        tokenizer: PreTrainedTokenizerFast,
        pack_sequences=False,
        device=None,
        sequence_column: str = "seq",
        max_length: int = 2048,
        label_column: str = "label",
        label_dtype: torch.dtype = torch.long,
        attention_mask_type: str = "bool",
        attention_mask_dtype: torch.dtype = torch.bool,
        token_level_task: bool = False,
        structure_task: bool = False,
        sentinel_value: int | float | None = -100,
        **kwargs,
    ):
        super().__init__(tokenizer=tokenizer, mlm_probability=0, mlm=False, **kwargs)  # Disable Masking for downstream tasks
        self.tokenizer = tokenizer
        self.pack_sequences = pack_sequences
        self.device = device
        self.sequence_column = sequence_column
        self.max_length = min(max_length, self.tokenizer.model_max_length)
        self.label_column = label_column
        self.label_dtype = label_dtype
        self.token_level_task = token_level_task
        self.structure_task = structure_task
        self.sentinel_value = sentinel_value
        self.attention_mask_type = attention_mask_type
        self.attention_mask_dtype = attention_mask_dtype

    def __call__(self, batch):
        if self.pack_sequences:
            # Tokenize the inputs
            input_ids_list = []
            position_ids_list = []
            seqlens = [0]
            labels = []

            for item in batch:
                features = self.tokenizer(
                    item[self.sequence_column],
                    padding=False,
                    truncation=True,
                    max_length=self.max_length,
                    return_tensors=None,
                )
                input_ids = features["input_ids"]
                position_ids = list(range(len(input_ids)))

                input_ids_list.append(input_ids)
                position_ids_list.append(position_ids)
                seqlens.append(len(input_ids))
                labels.append(item[self.label_column])

            if self.token_level_task:
                # Add the sentinel value to start and end of each label sequence if BOS token is present
                for i in range(len(labels)):
                    if input_ids_list[i][0] == self.tokenizer.bos_token_id:
                        labels[i] = [self.sentinel_value] + labels[i] + [self.sentinel_value]
                    # Align labels with positions (truncate/pad as needed)
                    labels[i] = [labels[i][pos] for pos in position_ids_list[i]]
                # Concatenate labels for token-level tasks
                labels = np.concatenate(labels, axis=0)
            if self.structure_task:
                # labels[0] contains a list of 2-lists.
                # Each 2-lists contains the positions of two residues in contact with each other.
                # We will create a nested tensor with tensors of variable size containing the contacts
                contacts = []
                for i in range(len(labels)):
                    contact = torch.tensor(labels[i], dtype=torch.int) + 1  # We add 1 to account for the BOS token in the input_ids
                    # With no random truncation, the BOS is always here, so we can safely add 1s
                    contacts.append(contact)
                labels = torch.nested.nested_tensor(contacts, dtype=torch.int)

            packed_batch = {
                "position_ids": torch.tensor(np.concatenate(position_ids_list, axis=0), dtype=torch.long).unsqueeze(0),
                "input_ids": torch.tensor(np.concatenate(input_ids_list, axis=0), dtype=torch.long).unsqueeze(0),
                "cu_seqlens": torch.tensor(np.cumsum(seqlens), dtype=torch.int32).unsqueeze(0),
                "max_seqlen": max(seqlens),
                "labels": torch.tensor(labels, dtype=self.label_dtype),
            }

            # batch = super().__call__([packed_batch])
            batch = packed_batch
            batch["cu_seqlens"] = batch["cu_seqlens"].to(torch.int32).squeeze()
        else:
            # Tokenize the inputs
            labels = [item.pop(self.label_column) for item in batch]
            # print([len(item[self.sequence_column]) for item in batch])
            for item in batch:
                features = self.tokenizer(
                    item.pop(self.sequence_column),
                    padding=True,
                    pad_to_multiple_of=8,
                    truncation=True,
                    max_length=self.max_length,
                    return_tensors=None,
                )
                item["input_ids"] = features["input_ids"]
                item["attention_mask"] = features["attention_mask"]
            # print([len(item["input_ids"]) for item in batch])
            batch = super().__call__(batch)
            batch["attention_mask"] = batch["attention_mask"].to(torch.bool)
            if self.attention_mask_type == "bool":
                pass
            elif self.attention_mask_type == "additive":
                batch["attention_mask"] = torch.where(batch["attention_mask"], float(0.0), float("-inf")).to(self.attention_mask_dtype)
            elif self.attention_mask_type == "multiplicative":
                batch["attention_mask"] = torch.where(batch["attention_mask"], 1, 0).to(self.attention_mask_dtype)
            else:
                raise ValueError(f"Unknown attention mask type: {self.attention_mask_type}")
            if self.token_level_task:
                # Add the sentinel value to start and end of each label sequence
                for i in range(len(labels)):
                    labels[i] = [self.sentinel_value] + labels[i] + [self.sentinel_value]
                    # Truncate the labels to the max length
                    labels[i] = labels[i][: len(batch["input_ids"][i])]
                # Concatenate labels for token-level tasks
                labels = np.concatenate(labels, axis=0)
                batch["labels"] = torch.tensor(labels, dtype=self.label_dtype)
            elif self.structure_task:
                # labels[0] contains a list of 2-lists.
                # Each 2-lists contains the positions of two residues in contact with each other.
                # We will create a nested tensor with tensors of variable size containing the contacts
                contacts = []
                for i in range(len(labels)):
                    contact = torch.tensor(labels[i], dtype=torch.int) + 1  # We add 1 to account for the BOS token in the input_ids
                    # With no random truncation, the BOS is always here, so we can safely add 1s
                    contacts.append(contact)
                labels = torch.nested.nested_tensor(contacts, dtype=torch.int)
                batch["labels"] = labels
            else:
                batch["labels"] = torch.tensor(labels, dtype=self.label_dtype)

        if self.device:
            for key in batch:
                if isinstance(batch[key], torch.Tensor):
                    batch[key] = batch[key].to(self.device)

        return batch
```
