## Supplementary material for "pLM representations unlock metagenomic space beyond homology": Code bundle: Tokenizer.py

```
import torch
from typing import List, Optional, Union, Dict
from torch import Tensor
import copy

from itertools import compress

# HuggingFace
from tokenizers import Tokenizer
from transformers import PreTrainedTokenizerFast, BatchEncoding
from tokenizers.models import WordPiece
from tokenizers.pre_tokenizers import Split


class ProteinTokenizer(PreTrainedTokenizerFast):

    def __init__(
        self,
        vocab: dict,
        pad_token_id: int,
        mask_token_id: int,
        bos_token_id: int,
        eos_token_id: int,
        unk_token_id: int,
        model_max_length: int,
        other_special_token_ids: Optional[List[int]] = None,
        **kwargs,
    ):
        """Vocabulary comprising the amino acids, and the special tokens <unk>, <bos>, <eos>, <pad> and <mask>.

        Args:
            vocab_path (str): Path to the vocabulary file to load.
            pad_token_id (int): <PAD> token index.
            mask_token_id (int): <MASK> token index.
            bos_token_id (int): <BOS> token index.
            eos_token_id (int): <EOS> token index.
            unk_token_id (int): <UNK> token index.
            other_special_token_ids (Optional[List[int]]): List of additional special tokens.
        """
        # Create vocabulary with special tokens
        token_to_id = dict()
        id_to_token = dict()

        for token, token_id in vocab.items():
            token = token.strip()
            token_to_id[token] = token_id
            id_to_token[token_id] = token

        # Define tokenizer and model
        tokenizer_object = Tokenizer(WordPiece(vocab=token_to_id, unk_token=id_to_token.get(unk_token_id)))

        # Pretokenize by splitting every character
        tokenizer_object.pre_tokenizer = Split("", behavior="removed")

        super().__init__(
            vocab=vocab,
            model_max_length=model_max_length,
            padding_side="right",
            truncation_side="right",
            pad_token_id=pad_token_id,
            pad_token=id_to_token.get(pad_token_id),
            mask_token_id=mask_token_id,
            mask_token=id_to_token.get(mask_token_id),
            bos_token_id=bos_token_id,
            bos_token=id_to_token.get(bos_token_id),
            eos_token_id=eos_token_id,
            eos_token=id_to_token.get(eos_token_id),
            unk_token_id=unk_token_id,
            unk_token=id_to_token.get(unk_token_id),
            other_special_token_ids=other_special_token_ids,
            model_input_names=["input_ids", "attention_mask", "special_tokens_mask"],
            tokenizer_object=tokenizer_object,
        )

        if other_special_token_ids is not None:
            self.add_special_tokens(
                {"additional_special_tokens": list(id_to_token.get(i) for i in other_special_token_ids)}
            )

        self.key_to_padding = {
            "input_ids": self.pad_token_id,
            "attention_mask": 0,
            "special_tokens_mask": 1,
            "position_ids": 0,
        }
        self.key_to_dtype = {
            "input_ids": torch.long,
            "attention_mask": torch.bool,
            "special_tokens_mask": torch.bool,
            "position_ids": torch.int,
        }

    def truncate(
        self,
        encoded_inputs: Dict[str, List[int]],
        max_length: Optional[int] = None,
        random_truncate: bool = True,
    ) -> Dict[str, List[List[int]]]:
        """
        Randomly truncate sequences in encoded inputs to the specified maximum length.

        Args:
            encoded_inputs (BatchEncoding): Tokenized inputs with keys like 'input_ids' as tensors.
            max_length (Optional[int]): Maximum length for truncation. Defaults to model's max length if None.
            random_truncate (bool): Whether to randomly truncate sequences.

        Returns:
            Dict[str, List[List[int]]]: Randomly truncated tokenized inputs.
        """

        for i, sequence in enumerate(encoded_inputs["input_ids"]):
            if len(sequence) > max_length:
                if random_truncate:
                    offset = torch.randint(0, len(sequence) - max_length + 1, (1,)).item()
                else:
                    offset = 0
                for key in encoded_inputs:
                    encoded_inputs[key][i] = encoded_inputs[key][i][offset : offset + max_length]

        # add option for different random truncate

        return encoded_inputs

    def remove_ambiguous(self, encoded_inputs: Dict[str, List[int]]) -> Dict[str, List[List[int]]]:
        """
        Remove ambiguous amino acids from the input sequences.

        Args:
            encoded_inputs (BatchEncoding): Tokenized inputs with keys like 'input_ids' as tensors.

        Returns:
            Dict[str, List[List[int]]]: Tokenized inputs without ambiguous amino acids.
        """

        for i, sequence in enumerate(encoded_inputs["input_ids"]):
            mask = [token_id != self.unk_token_id for token_id in sequence]
            for key in encoded_inputs:
                encoded_inputs[key][i] = list(compress(encoded_inputs[key][i], mask))
        return encoded_inputs

    def _pad(
        self,
        encoded_inputs: Dict[str, List[List[int]]],
        padding: Union[bool, str] = True,
        max_length: Optional[int] = None,
        pad_to_multiple_of: int = 8,
        **kwargs,
    ) -> Dict[str, List[List[int]]]:

        if isinstance(encoded_inputs, list):
            tmp = dict()
            for key in encoded_inputs[0]:
                tmp[key] = [encoded_inputs[i][key] for i in range(len(encoded_inputs))]
            encoded_inputs = tmp

        if max_length is None:
            max_length = self.model_max_length

        sequence_lengths = [len(sequence) for sequence in encoded_inputs["input_ids"]]
        if padding == "longest" or padding == True:
            max_length = min(max_length, max(sequence_lengths))

        if pad_to_multiple_of is not None and (max_length % pad_to_multiple_of != 0):
            max_length = ((max_length // pad_to_multiple_of) + 1) * pad_to_multiple_of

        for i, seq_len in enumerate(sequence_lengths):
            if seq_len < max_length:
                for key in encoded_inputs:
                    encoded_inputs[key][i] = encoded_inputs[key][i] + [self.key_to_padding[key]] * (
                        max_length - seq_len
                    )

        return encoded_inputs

    def pad(
        self,
        encoded_inputs: Dict[str, List[List[int]]],
        padding: Union[bool, str] = True,
        max_length: Optional[int] = None,
        pad_to_multiple_of: int = 8,
        return_tensors: str = "pt",
        **kwargs,
    ) -> Dict[str, List[List[int]]]:

        encoded_inputs = self._pad(
            encoded_inputs,
            padding,
            max_length,
            pad_to_multiple_of,
            **kwargs,
        )

        if return_tensors is not None:
            return BatchEncoding(encoded_inputs, tensor_type=return_tensors)

        return encoded_inputs

    def __call__(
        self,
        text: str | List[str],
        max_length: Optional[int] = None,
        padding: Union[bool, str] = False,
        truncation: bool = False,
        random_truncate: bool = False,
        remove_ambiguous: bool = False,
        return_special_tokens_mask: bool = True,
        return_tensors: str = None,
        add_special_tokens: bool = True,
        **kwargs,
    ) -> Dict[str, Tensor]:

        if isinstance(text, str):
            encoded_inputs = self.__call__(
                [text],
                max_length,
                padding,
                truncation,
                random_truncate,
                remove_ambiguous,
                return_special_tokens_mask,
                return_tensors,
            )
            for key in encoded_inputs:
                encoded_inputs[key] = encoded_inputs[key][0]
            return encoded_inputs

        # Tokenize without truncation or padding
        encoded_inputs = super().__call__(
            text,
            padding=False,
            truncation=False,
            verbose=False,
            return_special_tokens_mask=return_special_tokens_mask,
            **kwargs,
        )

        if max_length is None:
            max_length = self.model_max_length

        # Add special tokens
        if add_special_tokens:
            encoded_inputs["input_ids"] = [
                [self.bos_token_id] + seq + [self.eos_token_id] for seq in encoded_inputs["input_ids"]
            ]
            encoded_inputs["attention_mask"] = [[1, 1] + seq for seq in encoded_inputs["attention_mask"]]
            encoded_inputs["special_tokens_mask"] = [[1] + seq + [1] for seq in encoded_inputs["special_tokens_mask"]]

        # Truncate
        if truncation:
            encoded_inputs = self.truncate(
                encoded_inputs,
                max_length=max_length,  # Need to account for the BOS and EOS tokens
                random_truncate=random_truncate,
            )

        ## NOTE: Moved this to after truncation to avoid the offset when random truncation is used
        # Track original position indexes
        encoded_inputs["position_ids"] = [list(range(len(seq))) for seq in encoded_inputs["input_ids"]]

        # Remove ambiguous amino acids
        if remove_ambiguous:
            encoded_inputs = self.remove_ambiguous(encoded_inputs)

        # Add padding
        if padding:
            encoded_inputs = self._pad(encoded_inputs, max_length=max_length, return_tensors=return_tensors)

        if return_tensors is not None:
            return BatchEncoding(encoded_inputs, tensor_type=return_tensors)

        return encoded_inputs
```
