## Supplementary material for "pLM representations unlock metagenomic space beyond homology": Code bundle: TokenEmbeddingsDataset.py

```
from typing import List, Tuple, Union, Optional
import torch
from torch.utils.data import Dataset


class TokenEmbeddingsDataset(Dataset):
    """
    Dataset for precomputed token-level embeddings.

    Each item returns a tensor of shape (seq_len_i, hidden_size) for embeddings
    and a tensor of shape (seq_len_i,) for labels.
    """

    def __init__(self, embeddings_list: List[torch.Tensor], labels_list: List[torch.Tensor], tokenizer=None):
        """
        Args:
            embeddings_list: List of tensors, each with shape (seq_len_i, hidden_size)
            labels_list: List of tensors, each with shape (seq_len_i,)
            tokenizer: Optional tokenizer for compatibility
        """
        assert len(embeddings_list) == len(labels_list), (
            f"Number of embeddings ({len(embeddings_list)}) must match " f"number of labels ({len(labels_list)})"
        )

        self.embeddings_list = embeddings_list
        self.labels_list = labels_list
        self.tokenizer = tokenizer

        # Validate that all embeddings have the same hidden size
        if len(embeddings_list) > 0:
            hidden_size = embeddings_list[0].shape[-1]
            for i, emb in enumerate(embeddings_list):
                assert emb.shape[-1] == hidden_size, (
                    f"All embeddings must have the same hidden size. "
                    f"Expected {hidden_size}, got {emb.shape[-1]} at index {i}"
                )
                assert emb.shape[0] == labels_list[i].shape[0], (
                    f"Sequence length mismatch at index {i}. "
                    f"Embeddings shape: {emb.shape}, Labels shape: {labels_list[i].shape}"
                )

    def __len__(self) -> int:
        """Return the number of sequences in the dataset."""
        return len(self.embeddings_list)

    def __getitem__(self, idx: int) -> Tuple[torch.Tensor, torch.Tensor]:
        """
        Return the embedding tensor and labels tensor for sequence at index idx.

        Args:
            idx: Index of the sequence

        Returns:
            Tuple of (embedding_tensor, labels_tensor) where:
            - embedding_tensor has shape (seq_len_i, hidden_size)
            - labels_tensor has shape (seq_len_i,)
        """
        return self.embeddings_list[idx], self.labels_list[idx]


def token_embeddings_collate_fn(batch: List[Tuple[torch.Tensor, torch.Tensor]]) -> dict:
    """
    Collate function for TokenEmbeddingsDataset.

    Concatenates variable-length sequence embeddings and labels into batched tensors.

    Args:
        batch: List of (embedding_tensor, labels_tensor) tuples

    Returns:
        Dictionary containing:
        - 'embeddings': Concatenated tensor of shape (total_tokens, hidden_size)
        - 'labels': Concatenated tensor of shape (total_tokens,)
        - 'sequence_lengths': List of sequence lengths for reconstruction
    """
    if not batch:
        return {"embeddings": torch.empty(0, 0), "labels": torch.empty(0, dtype=torch.long), "sequence_lengths": []}

    embeddings_list = []
    labels_list = []
    sequence_lengths = []

    for embedding_tensor, labels_tensor in batch:
        embeddings_list.append(embedding_tensor)
        labels_list.append(labels_tensor)
        sequence_lengths.append(embedding_tensor.shape[0])

    # Concatenate all embeddings and labels
    concatenated_embeddings = torch.cat(embeddings_list, dim=0)
    concatenated_labels = torch.cat(labels_list, dim=0)

    return {"embeddings": concatenated_embeddings, "labels": concatenated_labels, "sequence_lengths": sequence_lengths}
```
