## Supplementary material for "pLM representations unlock metagenomic space beyond homology": Code bundle: Evaluator.py

```
import os
import json
from abc import ABC, abstractmethod
from typing import Any
import logging
import numpy as np
import torch
from dataclasses import dataclass
from pathlib import Path
from datetime import datetime

from apperitif.models.Encoder import Encoder
from apperitif.utils.evaluator_utils import EvaluatorResult, convert_to_json_serializable

# class EvaluatorConfig


@dataclass
class EvaluatorConfig:
    """
    Base configuration class for evaluators.
    This can be extended by specific evaluator configurations.
    """

    # Add common evaluator configuration parameters here if needed
    pass


class Evaluator(ABC):
    """
    Abstract class for evaluators.
    """

    def __init__(self):  # config
        super().__init__()
        # self.config = config
        self._name = self.name
        self.logger = logging.getLogger(__name__)
        logging.basicConfig(filename=None, level=logging.INFO)
        # Seeding?

    @property
    @abstractmethod
    def name(cls):
        """Class name that must be defined in subclasses"""
        pass

    @abstractmethod
    def __call__(self, model: Encoder, **kwargs) -> EvaluatorResult:
        """Evaluate the model on a specific task (classification, regression, ...) for a given set of metrics,
        a specific data setting (train, test, val), and a specific set of hyperparameters.

        Returns:
            EvaluatorResult: Standardized result object with metrics, hyperparameters, and metadata
        """

        pass

    def save_results_to_json(
        self,
        all_scores: dict,
        output_folder: str,
        task_name: str,
        task_description: str,
        filename: str = "results.json",
    ) -> str:
        """Save evaluation results to a JSON file in the output folder.

        Args:
            results: Dictionary object containing the evaluation results
            filename: Name of the JSON file to save (default: "results.json")

        Returns:
            str: Path to the saved JSON file

        Raises:
            ValueError: If output_folder is not configured
        """
        # Create output directory if it doesn't exist
        output_path = Path(output_folder)
        output_path.mkdir(parents=True, exist_ok=True)

        # Ensure filename has .json extension
        if not filename.endswith(".json"):
            filename += ".json"

        file_path = output_path / filename

        results_dict = {
            "all_results": all_scores,
            "task_name": task_name,
            "task_description": task_description,
            "timestamp": datetime.now().isoformat(),
        }

        if os.path.exists(file_path):
            self.logger.warning(f"File {file_path} already exists. Aggregating all results and overwriting.")

            # If the file already exists, load existing results and merge
            with open(file_path, "rt") as f:
                existing_results = json.load(f)

            # Merge existing results with new results
            results_dict["all_results"].update(existing_results.get("all_results", {}))

            if "validation_score" in existing_results and existing_results["validation_score"] > results_dict.get("validation_score", 0):
                self.logger.warning(
                    f"Existing validation score {existing_results['validation_score']} is higher than new score "
                    f"{results_dict.get('validation_score', 0)}. Keeping the existing score."
                )
                for key, value in existing_results.items():
                    if key != "all_results" and key != "timestamp":
                        results_dict[key] = value

        # Save to JSON file
        results_dict["all_results"] = {str(k): convert_to_json_serializable(v) for k, v in results_dict["all_results"].items()}
        with open(file_path, "w", encoding="utf-8") as f:
            json.dump(results_dict, f, indent=2, ensure_ascii=False)

        self.logger.info(f"Results saved to: {file_path}")
        return str(file_path)
```
