## Supplementary material for "pLM representations unlock metagenomic space beyond homology": Code bundle: SequenceClassificationEvaluator.py

```
import os
import ast
import json
import logging
from typing import List, Dict, Tuple, Any, Callable, Optional
import tqdm
from dataclasses import dataclass, field

import numpy as np
import torch
from sklearn.metrics import accuracy_score, f1_score, roc_auc_score

from torch.utils.data import DataLoader, Dataset

from apperitif.evaluation.Evaluator import Evaluator, EvaluatorConfig
from apperitif.models.Encoder import SequenceEncoder
from apperitif.models.regressor import Regressor, MLP
from apperitif.utils.deterministic import set_seed
from apperitif.utils.embedding_utils import get_embeddings_and_labels
from apperitif.utils.optimization import get_linear_scheduler_with_warmup, EarlyStopping
from apperitif.utils.dtype_utils import get_torch_dtype
from apperitif.utils.evaluator_utils import EvaluatorResult, create_standardized_result


@dataclass
class SequenceClassificationEvaluatorConfig(EvaluatorConfig):
    head_type: str = "base"  # Discriminator field for Hydra
    metrics: List[str] = field(default_factory=lambda: ["accuracy", "f1", "roc_auc"])
    decision_metric: Optional[str] = None
    num_labels: Optional[int] = None
    reduction_method: str = "mean"
    target_column: str = "labels"
    precomputing_batch_size: int = 128
    autocast_precision: str = "fp16"  # Changed to string for Hydra compatibility


@dataclass
class MLPSequenceClassificationEvaluatorConfig(SequenceClassificationEvaluatorConfig):
    head_type: str = "mlp"  # Discriminator field for Hydra
    hidden_size: int | list[int] = 512
    learning_rate: float | list[float] = 1e-3
    num_epochs: int | list[int] = 2
    weight_decay: float | list[float] = 0
    batch_size: int | list[int] = 32
    seeds: list[int] = field(default_factory=lambda: [0, 1, 2, 3, 4])
    finetuning: bool = False
    precomputed: bool = True  # Moved to config


class SequenceClassificationEvaluator(Evaluator):
    """
    Base evaluator for sequence classification tasks.
    """

    def __init__(
        self,
        config: SequenceClassificationEvaluatorConfig,
        datasets: Dict[str, Dataset],
        collate_fn: Callable,
    ):
        super().__init__()
        self.logger = logging.getLogger(__name__)
        logging.basicConfig(filename=None, level=logging.INFO)

        self.config = config
        self.datasets = datasets
        self.collate_fn = collate_fn

        # Extract config values
        self.metrics = config.metrics
        assert all(metric in ["accuracy", "f1", "roc_auc"] for metric in self.metrics), "Invalid metric(s)."
        assert len(self.metrics) > 0, "At least one metric must be provided."

        if config.decision_metric is None:
            self.decision_metric = self.metrics[0]
        else:
            assert config.decision_metric in self.metrics, f"Invalid decision metric: {config.decision_metric}"
            self.decision_metric = config.decision_metric

        self.num_labels = config.num_labels
        assert self.num_labels > 1, "Number of labels must be greater than 1."
        self.reduction_method = config.reduction_method
        self.target_column = config.target_column
        self.precomputing_batch_size = config.precomputing_batch_size
        self.autocast_precision = get_torch_dtype(config.autocast_precision)
        self.head_type = config.head_type  # For standardized output

    @property
    def name(cls):
        return "sequence_classification"

    def _compute_metric(self, labels: np.ndarray, preds: np.ndarray, metric: str) -> float:
        """
        Compute the value of the given metric.

        Args:
            labels: True labels.
            preds: Predicted probabilities or logits.
            metric: Metric to compute.

        Returns:
            Value of the metric.
        """
        if metric == "accuracy":
            return accuracy_score(labels, np.argmax(preds, axis=1))
        elif metric == "f1":
            return f1_score(labels, np.argmax(preds, axis=1), average="weighted")
        elif metric == "roc_auc":
            if self.num_labels == 2:
                return roc_auc_score(labels, preds[:, 1])
            else:
                return roc_auc_score(labels, preds, multi_class="ovo", labels=np.arange(self.num_labels))
        else:
            raise ValueError(f"Invalid metric: {metric}")

    def __call__(self, model: SequenceEncoder, **kwargs) -> EvaluatorResult:
        """
        Evaluate the model on a sequence classification task.

        Args:
            model: Encoder model to evaluate.

        Returns:
            Dictionary containing the evaluation metrics on the test set.
        """
        raise NotImplementedError


class MLPSequenceClassificationEvaluator(SequenceClassificationEvaluator):
    def __init__(
        self,
        config: MLPSequenceClassificationEvaluatorConfig,
        datasets: Dict[str, Dataset],
        collate_fn: Callable,
    ):
        super().__init__(config, datasets, collate_fn)
        self.hidden_size = config.hidden_size
        self.learning_rate = config.learning_rate
        self.num_epochs = config.num_epochs
        self.weight_decay = config.weight_decay
        self.batch_size = config.batch_size
        self.finetuning = config.finetuning  # Whether to fine-tune the trunk or not
        self.precomputed = config.precomputed  # Whether to use precomputed embeddings
        assert not self.precomputed if self.finetuning else True, "Cannot finetune with precomputed embeddings."

        if isinstance(self.hidden_size, int):
            self.hidden_size = [self.hidden_size]
        if isinstance(self.learning_rate, float):
            self.learning_rate = [self.learning_rate]
        if isinstance(self.num_epochs, int):
            self.num_epochs = [self.num_epochs]
        if isinstance(self.weight_decay, float):
            self.weight_decay = [self.weight_decay]
        if isinstance(self.batch_size, int):
            self.batch_size = [self.batch_size]

        self.seeds = config.seeds
        set_seed(self.seeds[0])

        self.early_stopping = EarlyStopping(patience=16, mode="max", disable=False)

    @property
    def name(cls):
        return "mlp_classification"

    def _evaluate_mlp_precomputed(
        self, mlp: torch.nn.Module, embedding_loader: DataLoader, device: str = "cuda", do_roc_auc: bool = False
    ) -> Dict[str, float]:
        mlp.eval()
        all_logits = []
        all_labels = []
        with torch.no_grad():
            for batch in embedding_loader:
                embeddings, batch_labels = batch
                embeddings = embeddings.to(device)
                logits = mlp(embeddings)
                all_logits.append(logits.float().cpu().numpy())
                all_labels.append(batch_labels.long().cpu().numpy())
        all_logits = np.concatenate(all_logits, axis=0)
        all_labels = np.concatenate(all_labels, axis=0)
        all_probas = torch.nn.functional.softmax(torch.tensor(all_logits), dim=1).numpy()

        scores = {}
        for metric in self.metrics:
            if metric == "roc_auc" and not do_roc_auc:
                scores[metric] = -1
            else:
                scores[metric] = self._compute_metric(all_labels, all_probas, metric)
        return scores

    def _train_mlp_precomputed(
        self,
        mlp: torch.nn.Module,
        train_loader: DataLoader,
        validation_loader: DataLoader,
        test_loader: DataLoader,
        optimizer: torch.optim.Optimizer,
        scheduler: torch.optim.lr_scheduler.LRScheduler | None,
        criterion: Callable,
        num_epochs: int,
        device: str = "cuda",
        do_test: bool = True,
        verbose: bool | str = "reduced",
        accumulation_steps: int = 1,
    ) -> Tuple[float, Dict[str, float]]:
        disable = (not verbose) or verbose == "reduced"
        pbar = tqdm.tqdm(total=(num_epochs) * len(train_loader) // accumulation_steps, desc="Training MLP", disable=disable)
        best_val_score = -1
        loader_len = len(train_loader)
        should_stop = False
        completed_steps = 0

        with torch.autocast(device_type=device, dtype=torch.float32, enabled=True):
            for n in range(num_epochs):
                mlp.train()
                optimizer.zero_grad()
                for idx, batch in enumerate(train_loader):
                    embeddings, labels = batch
                    embeddings, labels = embeddings.to(device), labels.to(device).long()
                    preds = mlp(embeddings)
                    loss = criterion(preds, labels)
                    loss = loss / accumulation_steps  # Normalize loss for accumulation
                    loss.backward()

                    if (idx + 1) % accumulation_steps == 0 or (idx + 1 == loader_len):
                        optimizer.step()
                        optimizer.zero_grad()
                        if scheduler is not None:
                            scheduler.step()

                        completed_steps += 1
                        pbar.update(1)

                    if ((idx + 1) % (loader_len // 3) == 0) or (idx + 1 == loader_len):
                        val_scores = self._evaluate_mlp_precomputed(mlp, validation_loader, device=device, do_roc_auc=False)
                        val_score = val_scores[self.decision_metric]
                        pbar.set_postfix({"val_score": val_score})
                        if val_score > best_val_score:
                            best_val_score = val_score
                            best_state_dict = {k: v.cpu() for k, v in mlp.state_dict().items()}
                        should_stop = self.early_stopping(val_score)
                        if should_stop:
                            break
                        mlp.train()
                if should_stop:
                    break
            pbar.close()

        if do_test:
            mlp.load_state_dict({k: v.to(device) for k, v in best_state_dict.items()})
            test_scores = self._evaluate_mlp_precomputed(mlp, test_loader, device=device, do_roc_auc=True)
        else:
            test_scores = {metric: -1 for metric in self.metrics}
        return best_val_score, test_scores, completed_steps

    def _evaluate_classifier(
        self, classifier: Regressor, dataloader: DataLoader, device: str = "cuda", do_roc_auc: bool = False
    ) -> Dict[str, float]:
        classifier.eval()
        all_logits = []
        all_labels = []
        with torch.no_grad():
            for batch in dataloader:
                labels = batch.pop(self.target_column).to(device).long().squeeze(0)
                logits = classifier(batch, frozen_backbone=True)
                all_logits.append(logits.cpu().numpy())
                all_labels.append(labels.cpu().numpy().reshape(-1))

        all_logits = np.concatenate(all_logits, axis=0)
        all_labels = np.concatenate(all_labels, axis=0)
        all_probas = torch.nn.functional.softmax(torch.tensor(all_logits), dim=1).numpy()

        scores = {}
        for metric in self.metrics:
            if metric == "roc_auc" and not do_roc_auc:
                scores[metric] = -1
            else:
                scores[metric] = self._compute_metric(all_labels, all_probas, metric)
        return scores

    def _train_classifier(
        self,
        classifier: Regressor,
        train_loader: DataLoader,
        validation_loader: DataLoader,
        test_loader: DataLoader,
        optimizer: torch.optim.Optimizer,
        scheduler: torch.optim.lr_scheduler.LRScheduler | None,
        criterion: Callable,
        num_epochs: int,
        finetuning: bool = False,
        device: str = "cuda",
        do_test: bool = True,
        verbose: bool | str = "reduced",
        accumulation_steps: int = 1,
    ) -> Tuple[float, Dict[str, float]]:
        disable_pbar = (not verbose) or verbose == "reduced"
        pbar = tqdm.tqdm(total=(num_epochs) * len(train_loader) // accumulation_steps, desc="Training Classifier", disable=disable_pbar)
        best_val_score = -1
        loader_len = len(train_loader)
        should_stop = False
        completed_steps = 0

        for n in range(num_epochs):
            frozen_backbone = not finetuning or n == 0
            classifier.train()
            for idx, batch in enumerate(train_loader):
                optimizer.zero_grad()
                labels = batch.pop(self.target_column).to(device).long()
                preds = classifier(batch, frozen_backbone=frozen_backbone)
                loss = criterion(preds, labels.reshape(-1)) / accumulation_steps  # Normalize loss for accumulation
                loss.backward()

                if (idx + 1) % accumulation_steps == 0 or (idx + 1 == loader_len):
                    optimizer.step()
                    optimizer.zero_grad()
                    if scheduler is not None:
                        scheduler.step()

                    completed_steps += 1
                    pbar.update(1)

                if ((idx + 1) % (loader_len // 3) == 0) or (idx + 1 == loader_len):
                    val_scores = self._evaluate_classifier(classifier, validation_loader, device=device, do_roc_auc=False)
                    val_score = val_scores[self.decision_metric]
                    pbar.set_postfix({"val_score": val_score})
                    if val_score > best_val_score:
                        best_val_score = val_score
                        best_state_dict = {k: v.cpu() for k, v in classifier.state_dict().items()}
                    should_stop = self.early_stopping(val_score)
                    if should_stop:
                        break
            if should_stop:
                break

        pbar.close()

        if do_test:
            classifier.load_state_dict({k: v.to(device) for k, v in best_state_dict.items()})
            test_scores = self._evaluate_classifier(classifier, test_loader, device=device, do_roc_auc=True)
        else:
            test_scores = {metric: -1 for metric in self.metrics}
        return best_val_score, test_scores, completed_steps

    def __call__(
        self,
        model: SequenceEncoder,
        model_kwargs: Dict[str, Any] = {},
        device: str = "cuda",
        return_all_scores: bool = False,
        do_test: bool = True,
        verbose: bool | str = "reduced",
        accumulation_steps: int = None,
        output_folder: str = None,
        task_name: Optional[str] = None,
        task_description: Optional[str] = None,
    ) -> EvaluatorResult:
        if self.precomputed:
            precomputing_dataloaders = {
                name: DataLoader(
                    ds,
                    batch_size=self.precomputing_batch_size,
                    collate_fn=self.collate_fn,
                    shuffle=False,
                )
                for name, ds in self.datasets.items()
                if name in ["train", "validation", "test"]
            }

            train_embeddings, train_labels = get_embeddings_and_labels(
                model,
                precomputing_dataloaders["train"],
                self.target_column,
                self.reduction_method,
                model_kwargs,
                self.autocast_precision,
            )
            val_embeddings, val_labels = get_embeddings_and_labels(
                model,
                precomputing_dataloaders["validation"],
                self.target_column,
                self.reduction_method,
                model_kwargs,
                self.autocast_precision,
            )
            if do_test and "test" in precomputing_dataloaders:
                test_embeddings, test_labels = get_embeddings_and_labels(
                    model,
                    precomputing_dataloaders["test"],
                    self.target_column,
                    self.reduction_method,
                    model_kwargs,
                    self.autocast_precision,
                )

            train_dataset = torch.utils.data.TensorDataset(torch.tensor(train_embeddings), torch.tensor(train_labels))
            validation_dataset = torch.utils.data.TensorDataset(torch.tensor(val_embeddings), torch.tensor(val_labels))
            if do_test and "test" in precomputing_dataloaders:
                test_dataset = torch.utils.data.TensorDataset(torch.tensor(test_embeddings), torch.tensor(test_labels))
            train_collate_fn = None
        else:
            train_dataset, validation_dataset = self.datasets["train"], self.datasets["validation"]
            test_dataset = self.datasets.get("test") if do_test else None
            train_collate_fn = self.collate_fn

        all_scores = {}
        hyperparameters = [
            (hs, lr, ne, wd, bs)
            for bs in self.batch_size
            for hs in self.hidden_size
            for lr in self.learning_rate
            for ne in self.num_epochs
            for wd in self.weight_decay
        ]

        # Load previously computed hyperparameters if available
        if output_folder is not None and os.path.exists(os.path.join(output_folder, "results.json")):
            file_path = os.path.join(output_folder, "results.json")
            with open(file_path, "rt") as f:
                results_dict = json.load(f)
            all_scores = results_dict["all_results"]
            hyperparameters_done = [ast.literal_eval(s) for s in list(results_dict["all_results"].keys())]

            print("Complete HP grid:", hyperparameters)
            hyperparameters = set(hyperparameters) - set(hyperparameters_done)

            print("Remaining HP grid:", hyperparameters)
            self.logger.info(
                f"Found {len(hyperparameters_done)} previously computed hyperparameters. Only {len(hyperparameters)} remaining to compute."
            )

            if len(hyperparameters) == 0:
                self.logger.info("All hyperparameters already computed. Finishing job.")
                return None

        global_pbar = tqdm.tqdm(
            total=len(hyperparameters),
            desc="Hyperparameter search",
            disable=(not verbose) or (verbose != "reduced"),
            smoothing=0,
        )
        for hparam in hyperparameters:
            hidden_size, learning_rate, num_epochs, weight_decay, batch_size = hparam
            all_scores[hparam] = []

            if accumulation_steps is None or batch_size % accumulation_steps != 0:
                accumulation_steps = max(1, batch_size // 8)

            print("Hyperparameters:", hparam)
            print("Accumulation steps:", accumulation_steps)

            train_loader = DataLoader(
                train_dataset,
                batch_size=batch_size // accumulation_steps,  # Adjust batch size for accumulation steps
                shuffle=True,
                collate_fn=train_collate_fn,
            )
            validation_loader = DataLoader(
                validation_dataset,
                batch_size=batch_size,
                shuffle=False,
                collate_fn=train_collate_fn,
            )
            if do_test and test_dataset is not None:
                test_loader = DataLoader(test_dataset, batch_size=batch_size, shuffle=False, collate_fn=train_collate_fn)
            else:
                test_loader = None

            for seed in self.seeds:
                print("Seed:", seed)
                set_seed(seed)
                if self.precomputed:
                    mlp = MLP(model.hidden_size, hidden_size, self.num_labels).to(device)
                    optimizer = torch.optim.AdamW(mlp.parameters(), lr=learning_rate, weight_decay=weight_decay, eps=1e-4)
                elif not self.finetuning:
                    model.eval()
                    classifier = Regressor(
                        trunk=model,
                        hidden_size=hidden_size,
                        output_size=self.num_labels,
                        reduction_method=self.reduction_method,
                        trunk_autocast_precision=self.autocast_precision,
                        head_precision=torch.float32,
                    ).to(device)
                    optimizer = torch.optim.AdamW(classifier.head.parameters(), lr=learning_rate, weight_decay=weight_decay, eps=1e-4)
                else:  # Finetuning
                    model.train()
                    classifier = Regressor(
                        trunk=model,
                        hidden_size=hidden_size,
                        output_size=self.num_labels,
                        reduction_method=self.reduction_method,
                        trunk_autocast_precision=self.autocast_precision,
                        head_precision=torch.float32,
                    ).to(device)
                    optimizer = torch.optim.AdamW(classifier.parameters(), lr=learning_rate, weight_decay=weight_decay, eps=1e-4)

                scheduler = get_linear_scheduler_with_warmup(
                    optimizer,
                    num_warmup_steps=int(0.1 * num_epochs * len(train_loader)),
                    num_training_steps=num_epochs * len(train_loader),
                )
                criterion = torch.nn.CrossEntropyLoss()
                self.early_stopping.reset()

                if self.precomputed:
                    val_score, test_scores, completed_steps = self._train_mlp_precomputed(
                        mlp,
                        train_loader,
                        validation_loader,
                        test_loader,
                        optimizer,
                        scheduler,
                        criterion,
                        num_epochs,
                        device,
                        do_test,
                        verbose=verbose,
                        accumulation_steps=accumulation_steps,
                    )
                else:  # Handles both finetuning and head-only training
                    val_score, test_scores, completed_steps = self._train_classifier(
                        classifier,
                        train_loader,
                        validation_loader,
                        test_loader,
                        optimizer,
                        scheduler,
                        criterion,
                        num_epochs,
                        self.finetuning,
                        device,
                        do_test,
                        verbose=verbose,
                        accumulation_steps=accumulation_steps,
                    )
                all_scores[hparam].append((val_score, test_scores, completed_steps))

            mean_val_score = np.mean([s[0] for s in all_scores[hparam]])
            std_val_score = np.std([s[0] for s in all_scores[hparam]])
            mean_test_score = {metric: np.mean([s[1][metric] for s in all_scores[hparam] if s[1] != -1]) for metric in self.metrics}
            std_test_score = {metric: np.std([s[1][metric] for s in all_scores[hparam] if s[1] != -1]) for metric in self.metrics}
            completed_steps = min([s[2] for s in all_scores[hparam]])
            all_scores[hparam] = {
                "validation_score": mean_val_score,
                "validation_std": std_val_score,
                "test_scores": mean_test_score,
                "test_std": std_test_score,
                "min_completed_steps": completed_steps,
            }
            global_pbar.update(1)

            # Checkpoint
            self.save_results_to_json(
                all_scores=all_scores,
                output_folder=output_folder,
                task_name=task_name,
                task_description=task_description,
            )

        global_pbar.close()

        best_hyperparameters = max(all_scores, key=lambda x: all_scores[x]["validation_score"])
        best_val_score = all_scores[best_hyperparameters].get("validation_score", 0)
        best_test_scores = all_scores[best_hyperparameters].get("test_scores", {})
        best_std_val_score = all_scores[best_hyperparameters].get("validation_std", 0)
        best_std_test_scores = all_scores[best_hyperparameters].get("test_std", {})
        best_completed_steps = all_scores[best_hyperparameters].get("min_completed_steps", -1)

        self.logger.info(f"Best hyperparameters: {best_hyperparameters}")
        self.logger.info(f"Best validation score: {best_val_score:.4f} +- {best_std_val_score:.4f}")
        self.logger.info(f"Best test scores: {best_test_scores} +- {best_std_test_scores}")
        self.logger.info(f"Best completed steps: {best_completed_steps}")

        # Convert std_test_scores format (remove "std_" prefix)
        std_metrics = {k.replace("std_", ""): v for k, v in best_std_test_scores.items()}

        return create_standardized_result(
            metrics=best_test_scores,
            evaluator_type=self.head_type,
            best_hyperparameters=best_hyperparameters,
            std_metrics=std_metrics,
            all_results=all_scores,
            validation_score=best_val_score,
            validation_std=best_std_val_score,
            model_name=getattr(model, "name_or_path", ""),
            dataset_name=getattr(self, "dataset_name", ""),
        )
```
