## Supplementary material for "pLM representations unlock metagenomic space beyond homology": Code bundle: SequenceRecoveryEvaluator.py

```
import os
import re
from tqdm import tqdm
import torch
import logging
import numpy as np
import pandas as pd
import polars as pl

from typing import List, Dict, Any

from datasets import Dataset
from accelerate import Accelerator
from accelerate.utils import ProjectConfiguration

from torch.utils.data import DataLoader

from apperitif.evaluation.Evaluator import Evaluator
from apperitif.collator import DataCollatorWithPacking


class SequenceRecoveryEvaluator(Evaluator):
    """
    Evaluator for sequence recovery tasks.

    Args:
        metrics: List of metrics to compute (default: ["accuracy", "f1", "roc_auc"]).
    """

    def __init__(
        self,
        packing: bool = True,
        batch_size: int = 128,
        save_steps: int = 1000,
        num_workers: int = 4,
        metrics: List[str] = ["accuracy", "f1", "roc_auc"],
        decision_metric: str | None = None,
    ):
        self.logger = logging.getLogger(__name__)
        logging.basicConfig(filename=None, level=logging.INFO)

        super().__init__()

        self.packing = packing
        self.batch_size = batch_size
        self.save_steps = save_steps
        self.num_workers = num_workers
        self.metrics = metrics
        assert all(metric in ["accuracy", "f1", "roc_auc"] for metric in self.metrics), "Invalid metric(s)."
        assert len(self.metrics) > 0, "At least one metric must be provided."

        if decision_metric is None:
            self.decision_metric = self.metrics[0]
        else:
            assert decision_metric in self.metrics, f"Invalid decision metric: {decision_metric}"
            self.decision_metric = decision_metric

    @property
    def name(cls):
        return "sequence_recovery"

    def _compute_metric(self, tokenizer, homologs_dict: dict, model_name: str, output_path: str):
        parquet_files = [f for f in os.listdir(output_path) if f.startswith("data") and f.endswith(".parquet")]

        expected_schema = {
            "logits": pl.List(pl.Float64),
            "id": pl.Utf8,
            "masked_index": pl.Int64,
        }

        print(f"Loading {len(parquet_files)} files from {output_path}")
        df = pl.concat([pl.read_parquet(os.path.join(output_path, f)).cast(expected_schema) for f in parquet_files])

        print("Decoding predicted residues")
        df = df.with_columns(
            [
                pl.col("logits").map_elements(lambda logits: tokenizer.decode(np.argmax(logits)), return_dtype=pl.Utf8).alias("predicted"),
                pl.col("masked_index").cast(pl.Int64).alias("res_ix"),
            ]
        )

        print("Parsing sequence identifiers")
        split_df = df["id"].str.split_exact("_", 1)
        df = df.with_columns([split_df.struct.field("field_0").alias("id"), split_df.struct.field("field_1").alias("number")])

        output_file = os.path.join(output_path, f"predictions.parquet")
        print(f"Saving full output to {output_file}")
        # df.write_parquet(output_file, compression="gzip")

        print("Computing accuracy per category")
        scores = {
            key: (
                cat.join(df, on=["id", "res_ix", "number"], how="inner")
                .with_columns((pl.col("predicted") == pl.col("most_common")).alias("correct"))
                .select((pl.col("correct").sum() / pl.len() * 100).alias("accuracy"))
                .item()
            )
            for key, cat in homologs_dict.items()
        }

        return scores

    def __call__(self, model, tokenizer, dataset: Dataset, model_name: str, output_path: str) -> Dict[str, Any]:
        """
        Evaluate the model on a non-target sequence recovery task.

        Args:
            model: Encoder model to evaluate.

        Returns:
            Dictionary containing the evaluation metrics on the test set.
        """
        # Define paths
        os.makedirs(output_path, exist_ok=True)

        project_config = ProjectConfiguration(
            project_dir=output_path,
            automatic_checkpoint_naming=False,
        )
        accelerator = Accelerator(
            project_config=project_config,
            mixed_precision="fp16",
        )

        # Put model in evaluation mode
        self.logger.info(f"Loading model {model_name}")
        model.eval()

        if self.packing:
            collate_fn = DataCollatorWithPacking()
            dataset.set_transform(
                lambda x: tokenizer(x["sequence"], truncation=False, padding=False, return_tensors=None)
                | {"id": x["id"]}
                | {"masked_index": x["masked_index"]}
            )
        else:
            dataset.set_transform(
                lambda x: tokenizer(x["sequence"], truncation=False, padding=True, return_tensors="pt", pad_to_multiple_of=8)
                | {"id": x["id"]}
                | {"masked_index": x["masked_index"]}
            )

        # Dataloader
        dataloader = DataLoader(
            dataset,
            batch_size=self.batch_size,
            collate_fn=collate_fn if self.packing else None,
            shuffle=False,
            prefetch_factor=2,
            num_workers=self.num_workers,
            pin_memory=True,
            persistent_workers=True,
        )

        model, dataloader = accelerator.prepare(model, dataloader)

        # Process bar
        total_batches = len(dataloader)
        pbar = tqdm(
            desc=f"Evaluating metric {self.name} on model {model_name}",
            unit="step",
            initial=0,
            total=total_batches,
            disable=(not accelerator.is_main_process),
            smoothing=0.02,
        )

        # Temporary dict to store data in between dumps
        results = {
            "logits": [],
            "id": [],
            "masked_index": [],
        }

        torch.backends.cuda.matmul.allow_tf32 = True
        torch.backends.cudnn.allow_tf32 = True

        start_idx = 0
        output_files = os.listdir(output_path)
        if len(output_files) > 0:
            # Find the maximum index from files named data_<int>.parquet
            parquet_indices = [
                int(re.match(r"data_(\d+)\.parquet", f).group(1)) for f in output_files if re.match(r"data_(\d+)\.parquet", f)
            ]
            latest = max(parquet_indices) if parquet_indices else 0
            if latest > 0:
                start_idx = latest + 1
                self.logger.info(f"Resuming from index {start_idx}")

        # Iterate through batches
        for idx, batch in enumerate(dataloader):
            # Skip already processed batches
            if idx < start_idx:
                pbar.update(1)
                continue

            # Update local batch counter
            input_ids, ids, max_seqlen, cu_seqlens, position_ids, attention_mask, masked_index = (
                batch["input_ids"],
                batch["id"],
                batch.get("max_seqlen", None),
                batch.get("cu_seqlens", None),
                batch.get("position_ids", None),
                batch.get("attention_mask", None),
                batch.get("masked_index", None),
            )

            # Compute model output
            with torch.no_grad(), accelerator.autocast():
                # Forward pass
                if self.packing:
                    output = model(
                        input_ids,
                        position_ids=position_ids,
                        max_seqlen=max_seqlen,
                        cu_seqlens=cu_seqlens,
                    )
                else:
                    if not "esm" in model_name:
                        attention_mask = torch.where(attention_mask.to(torch.bool), float(0.0), float("-inf")).to(torch.float16)
                    output = model(
                        input_ids=input_ids,
                        attention_mask=attention_mask,
                    )

            if self.packing:
                # Find the masked token for each sequence
                logits_flat = output["logits"].view(-1, output["logits"].size(-1))  # [total_length, vocab_size]
                masked_token = (input_ids.view(-1) == tokenizer.mask_token_id).nonzero(as_tuple=True)[0]

                # Get logits at masked positions
                logits = logits_flat[masked_token]

            else:
                # Find the masked token for each sequence
                masked_token = (input_ids == tokenizer.mask_token_id).nonzero(as_tuple=False)

                # Get logits at masked positions
                logits = output["logits"][masked_token[:, 0], masked_token[:, 1]]

            results["masked_index"].extend(masked_index.cpu().numpy())
            results["logits"].extend([logit.cpu().numpy() for logit in logits])
            results["id"].extend(ids)

            pbar.update(1)

            if (idx > 0 and idx % self.save_steps == 0) or idx == total_batches - 1:
                print(idx, self.save_steps)
                gathered = accelerator.gather_for_metrics(
                    [
                        {"id": id_, "masked_index": idx_, "logits": logit_}
                        for id_, idx_, logit_ in zip(results["id"], results["masked_index"], results["logits"])
                    ]
                )

                if accelerator.is_main_process:
                    df = pd.DataFrame(gathered)
                    output_file = os.path.join(output_path, f"data_{idx:05d}.parquet")
                    df.to_parquet(output_file, index=False, compression="gzip")
                    self.logger.info(f"Data saved in file {output_file}")

                results = {"logits": [], "id": [], "masked_index": []}

        pbar.close()
        accelerator.end_training()

        with open(os.path.join(output_path, "done.txt"), "w") as f:
            f.write("done")
```
