## Supplementary material for "pLM representations unlock metagenomic space beyond homology": Code bundle: SequenceRegressionEvaluator.py

```
import os
import ast
import json
import logging
from typing import List, Dict, Tuple, Any, Callable, Optional
import tqdm
from dataclasses import dataclass, field

import numpy as np
import torch
from sklearn.metrics import mean_squared_error, r2_score
from scipy.stats import spearmanr, pearsonr

from torch.utils.data import DataLoader, Dataset

from apperitif.evaluation.Evaluator import Evaluator, EvaluatorConfig
from apperitif.models.Encoder import SequenceEncoder
from apperitif.models.regressor import MLP, Regressor
from apperitif.utils.deterministic import set_seed
from apperitif.utils.embedding_utils import get_embeddings_and_labels
from apperitif.utils.optimization import get_linear_scheduler_with_warmup, EarlyStopping
from apperitif.utils.dtype_utils import get_torch_dtype
from apperitif.utils.evaluator_utils import EvaluatorResult, create_standardized_result


@dataclass
class SequenceRegressionEvaluatorConfig(EvaluatorConfig):
    head_type: str = "base"  # Discriminator field for Hydra
    metrics: List[str] = field(default_factory=lambda: ["spearmanr", "pearsonr", "mse", "r2"])
    decision_metric: Optional[str] = None
    reduction_method: str = "mean"
    target_column: str = "labels"
    precomputing_batch_size: int = 128
    autocast_precision: str = "fp16"  # Changed to string for Hydra compatibility


@dataclass
class MLPSequenceRegressionEvaluatorConfig(SequenceRegressionEvaluatorConfig):
    head_type: str = "mlp"  # Discriminator field for Hydra
    hidden_size: int | list[int] = 512
    learning_rate: float | list[float] = 1e-3
    num_epochs: int | list[int] = 2
    weight_decay: float | list[float] = 0
    batch_size: int | list[int] = 32
    seeds: list[int] = field(default_factory=lambda: [0, 1, 2, 3, 4])
    finetuning: bool = False
    precomputed: bool = True  # Moved to config


class SequenceRegressionEvaluator(Evaluator):
    def __init__(
        self,
        config: SequenceRegressionEvaluatorConfig,
        datasets: Dict[str, Dataset],
        collate_fn: Callable,
    ):
        super().__init__()
        self.logger = logging.getLogger(__name__)
        logging.basicConfig(filename=None, level=logging.INFO)

        self.config = config
        self.datasets = datasets
        self.collate_fn = collate_fn

        # Extract config values
        self.metrics = config.metrics
        assert all(metric in ["mse", "r2", "pearsonr", "spearmanr"] for metric in self.metrics), "Invalid metric(s)."

        if config.decision_metric is None:
            self.decision_metric = self.metrics[0]
        else:
            assert config.decision_metric in self.metrics, f"Invalid decision metric: {config.decision_metric}"
            self.decision_metric = config.decision_metric

        self.reduction_method = config.reduction_method
        self.target_column = config.target_column
        self.precomputing_batch_size = config.precomputing_batch_size
        self.autocast_precision = get_torch_dtype(config.autocast_precision)
        self.head_type = config.head_type  # For standardized output
        self.precomputed = config.precomputed  # Now an attribute
        self.head_type = config.head_type  # For standardized output

    @property
    def name(cls):
        return "sequence_regression"

    def _compute_metric(self, labels: np.ndarray, preds: np.ndarray, metric: str) -> float:
        if metric == "mse":
            return mean_squared_error(labels, preds)
        elif metric == "r2":
            return r2_score(labels, preds)
        elif metric == "pearsonr":
            return pearsonr(labels, preds)[0]
        elif metric == "spearmanr":
            return spearmanr(labels, preds)[0]
        else:
            raise ValueError(f"Invalid metric: {metric}")

    def __call__(self, model: SequenceEncoder, **kwargs) -> EvaluatorResult:
        """
        Evaluate the model on a sequence classification task.

        Args:
            model: Encoder model to evaluate.

        Returns:
            Dictionary containing the evaluation metrics on the test set.
        """
        raise NotImplementedError


class MLPSequenceRegressionEvaluator(SequenceRegressionEvaluator):
    """
    Evaluator for MLP regression.

    Args:
        config: MLPSequenceRegressionEvaluatorConfig containing all configuration parameters
        datasets: Dictionary containing the train, validation, and test datasets
        collate_fn: Function to collate batches
    """

    def __init__(
        self,
        config: MLPSequenceRegressionEvaluatorConfig,
        datasets: Dict[str, Dataset],
        collate_fn: Callable,
    ):
        super().__init__(config, datasets, collate_fn)

        # Extract MLP-specific config values
        self.hidden_size = config.hidden_size
        self.learning_rate = config.learning_rate
        self.num_epochs = config.num_epochs
        self.weight_decay = config.weight_decay
        self.batch_size = config.batch_size
        self.seeds = config.seeds
        self.finetuning = config.finetuning

        # Validation: ensure precomputed and finetuning settings are compatible
        assert not self.precomputed if self.finetuning else True, (
            "Either precomputed embeddings or finetuning must be enabled. "
            "We cannot finetune the trunk while also using precomputed embeddings."
        )

        # Ensure all hyperparameters are lists for grid search

        if isinstance(self.hidden_size, int):
            self.hidden_size = [self.hidden_size]
        if isinstance(self.learning_rate, float):
            self.learning_rate = [self.learning_rate]
        if isinstance(self.num_epochs, int):
            self.num_epochs = [self.num_epochs]
        if isinstance(self.weight_decay, float):
            self.weight_decay = [self.weight_decay]
        if isinstance(self.batch_size, int):
            self.batch_size = [self.batch_size]

        set_seed(self.seeds[0])  # Set the seed for reproducibility

        self.early_stopping = EarlyStopping(
            patience=16, mode="max", disable=False
        )  # Early stopping over 4 epoch (4 epochs, 4 evaluations per epoch)

    @property
    def name(cls):
        return "mlp_regression"

    def _evaluate_mlp_precomputed(self, mlp: torch.nn.Module, embedding_loader: DataLoader, device: str = "cuda") -> Dict[str, float]:
        mlp.eval()
        all_preds = []
        all_labels = []
        with torch.no_grad():
            for batch in embedding_loader:
                embeddings, batch_labels = batch
                embeddings = embeddings.to(device)
                preds = mlp(embeddings).squeeze(-1)  # Only squeeze the last dimension
                # Ensure preds is always 1D
                if preds.dim() == 0:  # If scalar, make it 1D
                    preds = preds.unsqueeze(0)
                # Float conversion needed for numpy conversion if autocast_precision is bfloat16
                all_preds.append(preds.float().cpu().numpy())
                all_labels.append(batch_labels.float().cpu().numpy())
        all_preds = np.concatenate(all_preds, axis=0)
        all_labels = np.concatenate(all_labels, axis=0)
        # Compute the evaluation metrics
        scores = {}
        for metric in self.metrics:
            scores[metric] = self._compute_metric(all_labels, all_preds, metric)
        return scores

    def _train_mlp_precomputed(
        self,
        mlp: torch.nn.Module,
        train_loader: DataLoader,
        validation_loader: DataLoader,
        test_loader: DataLoader,
        optimizer: torch.optim.Optimizer,
        scheduler: torch.optim.lr_scheduler.LRScheduler | None = None,
        criterion: Callable = torch.nn.MSELoss(),
        num_epochs: int = 2,
        label_dtype: torch.dtype = torch.float32,
        device: str = "cuda",
        do_test: bool = True,
        verbose: bool | str = "reduced",
        accumulation_steps: int = 1,
    ) -> Tuple[float, Dict[str, float]]:
        disable = (not verbose) or verbose == "reduced"
        pbar = tqdm.tqdm(
            total=(num_epochs) * len(train_loader) // accumulation_steps,
            desc=f"Training MLP",
            disable=disable,
        )
        best_val_score = -1
        loader_len = len(train_loader)
        should_stop = False
        completed_steps = 0

        with torch.autocast(device_type=device, dtype=torch.float32, enabled=True):
            for n in range(num_epochs):
                mlp.train()
                optimizer.zero_grad()
                for idx, batch in enumerate(train_loader):
                    embeddings, labels = batch
                    embeddings, labels = embeddings.to(device), labels.to(device).to(label_dtype)
                    preds = mlp(embeddings).squeeze(-1)  # Only squeeze the last dimension
                    # Ensure preds has the same shape as labels
                    if preds.dim() == 0:  # If scalar, make it 1D
                        preds = preds.unsqueeze(0)
                    loss = criterion(preds, labels)
                    loss = loss / accumulation_steps  # Normalize loss for accumulation
                    loss.backward()

                    if (idx + 1) % accumulation_steps == 0 or (idx + 1 == loader_len):
                        optimizer.step()
                        optimizer.zero_grad()
                        if scheduler is not None:
                            scheduler.step()

                        completed_steps += 1
                        pbar.update(1)

                    if ((idx + 1) % (loader_len // 3) == 0) or (idx + 1 == loader_len):
                        val_score = self._evaluate_mlp_precomputed(mlp, validation_loader, device=device)[self.decision_metric]
                        pbar.set_postfix({"val_score": val_score})
                        if val_score > best_val_score:
                            best_val_score = val_score
                            # Save the best state_dict on CPU for use on test set
                            best_state_dict = {k: v.cpu() for k, v in mlp.state_dict().items()}
                        should_stop = self.early_stopping(val_score)
                        if should_stop:
                            break
                        mlp.train()
                if should_stop:
                    break
            pbar.close()

        if do_test:
            # Evaluate the MLP classifier on the test set
            mlp.load_state_dict(best_state_dict)
            test_scores = self._evaluate_mlp_precomputed(mlp, test_loader, device=device)
        else:
            test_scores = {metric: -1 for metric in self.metrics}
        return best_val_score, test_scores, completed_steps

    def _evaluate_regressor(self, regressor: Regressor, dataloader: DataLoader, device: str = "cuda") -> Dict[str, float]:
        """
        Evaluate the regressor on the given dataloader.

        Args:
            regressor: Regressor model to evaluate.
            dataloader: DataLoader for the dataset.
            device: Device to use for evaluation (default: "cuda").

        Returns:
            Dictionary containing the evaluation metrics.
        """
        regressor.eval()
        all_preds = []
        all_labels = []
        with torch.no_grad():
            for batch in dataloader:
                labels = batch.pop(self.target_column).to(device).squeeze(0)
                preds = regressor(batch, frozen_backbone=True).squeeze(-1)  # Only squeeze the last dimension
                # Ensure preds is always 1D
                if preds.dim() == 0:  # If scalar, make it 1D
                    preds = preds.unsqueeze(0)
                all_preds.append(preds.cpu().numpy())
                all_labels.append(labels.cpu().numpy())
        all_preds = np.concatenate(all_preds, axis=0)
        all_labels = np.concatenate(all_labels, axis=0)

        scores = {}
        for metric in self.metrics:
            scores[metric] = self._compute_metric(all_labels, all_preds, metric)
        return scores

    def _train_regressor(
        self,
        regressor: Regressor,
        train_loader: DataLoader,
        validation_loader: DataLoader,
        test_loader: DataLoader,
        optimizer: torch.optim.Optimizer,
        scheduler: torch.optim.lr_scheduler.LRScheduler | None = None,
        criterion: Callable = torch.nn.MSELoss(),
        num_epochs: int = 2,
        finetuning: bool = False,
        label_dtype: torch.dtype = torch.float32,
        device: str = "cuda",
        do_test: bool = True,
        verbose: bool | str = "reduced",
        accumulation_steps: int = 1,
    ) -> Tuple[float, Dict[str, float]]:
        """
        Train the regressor on the training set and evaluate on the validation set.

        Args:
            regressor: Regressor model to train.
            train_loader: DataLoader for the training set.
            validation_loader: DataLoader for the validation set.
            test_loader: DataLoader for the test set.
            optimizer: Optimizer for training.
            scheduler: Learning rate scheduler (optional).
            criterion: Loss function (default: MSELoss).
            num_epochs: Number of epochs to train (default: 2).
            label_dtype: Data type of the labels (default: torch.float32).
            device: Device to use for training and evaluation (default: "cuda").
            do_test: Whether to evaluate on the test set (default: True).

        Returns:
            Tuple containing the best validation score and test scores.
        """
        disable_pbar = (not verbose) or verbose == "reduced"
        pbar = tqdm.tqdm(
            total=(num_epochs) * len(train_loader) // accumulation_steps,
            desc=f"Training Regressor",
            disable=disable_pbar,
        )
        best_val_score = -1
        loader_len = len(train_loader)
        completed_steps = 0

        for n in range(num_epochs):
            # Freeze the backbone if not finetuning or if it's the first epoch
            frozen_backbone = not finetuning or n == 0
            regressor.train()
            for idx, batch in enumerate(train_loader):
                optimizer.zero_grad()
                labels = batch.pop(self.target_column).to(device).to(label_dtype).squeeze(0)
                preds = regressor(batch, frozen_backbone=frozen_backbone).squeeze()
                loss = criterion(preds, labels) / accumulation_steps  # Normalize loss for accumulation
                loss.backward()

                if (idx + 1) % accumulation_steps == 0 or (idx + 1 == loader_len):
                    optimizer.step()
                    optimizer.zero_grad()
                    if scheduler is not None:
                        scheduler.step()

                    completed_steps += 1
                    pbar.update(1)

                if ((idx + 1) % (loader_len // 3) == 0) or (idx + 1 == loader_len):
                    val_score = self._evaluate_regressor(regressor, validation_loader, device=device)[self.decision_metric]
                    pbar.set_postfix({"val_score": val_score})
                    if val_score > best_val_score:
                        best_val_score = val_score
                        # Save the best state_dict on CPU for use on test set
                        best_state_dict = {k: v.cpu() for k, v in regressor.state_dict().items()}
                    should_stop = self.early_stopping(val_score)
                    if should_stop:
                        break
            if should_stop:
                pbar.close()
                break
        pbar.close()
        if do_test:
            # Evaluate the regressor on the test set
            regressor.load_state_dict(best_state_dict)
            test_scores = self._evaluate_regressor(regressor, test_loader, device=device)
        else:
            test_scores = {metric: -1 for metric in self.metrics}
        return best_val_score, test_scores, completed_steps

    def __call__(
        self,
        model: SequenceEncoder,
        model_kwargs: Dict[str, Any] = {},
        device: str = "cuda",
        return_all_scores: bool = False,
        do_test: bool = True,
        verbose: bool | str = "reduced",
        accumulation_steps: int = None,
        output_folder: str = None,
        task_name: Optional[str] = None,
        task_description: Optional[str] = None,
    ) -> EvaluatorResult:
        """
        Evaluate the model on the test set using MLP classification.

        Args:
            model: Encoder model to evaluate.
            model_kwargs: Additional arguments to pass to the model.
            device: Device to use for training and evaluation (default: "cuda").
            return_all_scores: Whether to return all scores for hyperparameter selection (default: False).
            do_test: Whether to evaluate on the test set (default: True).
            verbose: Verbosity level for training progress (default: "reduced").

        Returns:
            Dictionary containing the evaluation metrics on the test set.
        """
        if self.precomputed:
            # Build the precomputing dataloaders
            precomputing_dataloaders = {
                "train": DataLoader(
                    self.datasets["train"],
                    batch_size=self.precomputing_batch_size,
                    collate_fn=self.collate_fn,
                    shuffle=False,
                    num_workers=0,
                ),
                "validation": DataLoader(
                    self.datasets["validation"],
                    batch_size=self.precomputing_batch_size,
                    collate_fn=self.collate_fn,
                    shuffle=False,
                    num_workers=0,
                ),
            }
            if do_test:
                precomputing_dataloaders["test"] = DataLoader(
                    self.datasets["test"],
                    batch_size=self.precomputing_batch_size,
                    collate_fn=self.collate_fn,
                    shuffle=False,
                    num_workers=0,
                )

            # Precompute the embeddings (and labels) for the train, validation and test sets
            train_embeddings, train_labels = get_embeddings_and_labels(
                model=model,
                dataloader=precomputing_dataloaders["train"],
                target_column=self.target_column,
                reduction_method=self.reduction_method,
                model_kwargs=model_kwargs,
                autocast_precision=self.autocast_precision,
            )
            val_embeddings, val_labels = get_embeddings_and_labels(
                model=model,
                dataloader=precomputing_dataloaders["validation"],
                target_column=self.target_column,
                reduction_method=self.reduction_method,
                model_kwargs=model_kwargs,
                autocast_precision=self.autocast_precision,
            )
            # If do_test is True, precompute the test embeddings and labels
            if do_test:
                test_embeddings, test_labels = get_embeddings_and_labels(
                    model=model,
                    dataloader=precomputing_dataloaders["test"],
                    target_column=self.target_column,
                    reduction_method=self.reduction_method,
                    model_kwargs=model_kwargs,
                    autocast_precision=self.autocast_precision,
                )
            train_dataset = torch.utils.data.TensorDataset(torch.tensor(train_embeddings), torch.tensor(train_labels))
            validation_dataset = torch.utils.data.TensorDataset(torch.tensor(val_embeddings), torch.tensor(val_labels))
            if do_test:
                test_dataset = torch.utils.data.TensorDataset(torch.tensor(test_embeddings), torch.tensor(test_labels))
            train_collate_fn = None
        else:
            self.logger.info("Using the original datasets and collate_fn for training.")
            # Use the original datasets and collate_fn
            train_dataset = self.datasets["train"]
            validation_dataset = self.datasets["validation"]
            if do_test:
                test_dataset = self.datasets["test"]
            train_collate_fn = self.collate_fn

        all_scores = {}
        hyperparameters = [
            (hidden_size, learning_rate, num_epochs, weight_decay, batch_size)
            for batch_size in self.batch_size
            for hidden_size in self.hidden_size
            for learning_rate in self.learning_rate
            for num_epochs in self.num_epochs
            for weight_decay in self.weight_decay
        ]

        # Load previously computed hyperparameters if available
        if output_folder is not None and os.path.exists(os.path.join(output_folder, "results.json")):
            file_path = os.path.join(output_folder, "results.json")
            with open(file_path, "rt") as f:
                results_dict = json.load(f)
            all_scores = results_dict["all_results"]
            hyperparameters_done = [ast.literal_eval(s) for s in list(results_dict["all_results"].keys())]

            hyperparameters = set(hyperparameters) - set(hyperparameters_done)
            self.logger.info(
                f"Found {len(set(hyperparameters_done))} previously computed hyperparameters. {len(hyperparameters)} remaining to compute."
            )

        global_pbar = tqdm.tqdm(
            total=len(hyperparameters),
            desc="Hyperparameter search",
            disable=(not verbose) or (verbose != "reduced"),
            smoothing=0,  # Use average speed to estimate the time remaining
        )
        for hparam in hyperparameters:
            hidden_size, learning_rate, num_epochs, weight_decay, batch_size = hparam
            all_scores[hparam] = []

            if accumulation_steps is None or batch_size % accumulation_steps != 0:
                accumulation_steps = max(1, batch_size // 8)

            print("Hyperparameters:", hparam)
            print("Accumulation steps:", accumulation_steps)

            train_loader = torch.utils.data.DataLoader(
                train_dataset,
                batch_size=batch_size // accumulation_steps,  # Adjust batch size for accumulation steps
                shuffle=True,
                collate_fn=train_collate_fn,
            )
            validation_loader = torch.utils.data.DataLoader(
                validation_dataset,
                batch_size=batch_size,
                shuffle=False,
                collate_fn=train_collate_fn,
            )
            if do_test:
                test_loader = torch.utils.data.DataLoader(
                    test_dataset,
                    batch_size=batch_size,
                    shuffle=False,
                    collate_fn=train_collate_fn,
                )
            else:
                test_loader = None

            for seed in self.seeds:
                set_seed(seed)

                print("Seed:", seed)

                # TODO clean that up to the criterion
                if self.precomputed:
                    mlp = MLP(model.hidden_size, hidden_size)
                    dtype = torch.float32  # Default dtype for MLP
                    mlp = mlp.to(device, dtype=dtype)
                    optimizer = torch.optim.AdamW(
                        mlp.parameters(),
                        lr=learning_rate,
                        weight_decay=weight_decay,
                        eps=1e-4,
                    )
                elif not self.finetuning:
                    model.eval()  # Ensure the trunk is in evaluation mode to match _get_embeddings_and_labels
                    regressor = Regressor(
                        trunk=model,
                        use_low_rank=False,
                        reduction_method=self.reduction_method,
                        hidden_size=hidden_size,
                        trunk_autocast_precision=self.autocast_precision,  # Trunk weight precision handled when defining the trunk
                        head_precision=torch.float32,  # Default precision for the head
                    ).to(device)
                    dtype = regressor.head_precision  # Use the head's autocast precision as label dtype
                    optimizer = torch.optim.AdamW(
                        regressor.head.parameters(),
                        lr=learning_rate,
                        weight_decay=weight_decay,
                        eps=1e-4,
                    )
                    # Check if we should add requires grad to false for the trunk parameters
                else:
                    # If finetuning, we use the full model
                    model.train()  # Ensure the trunk is in training mode
                    regressor = Regressor(
                        trunk=model,
                        use_low_rank=False,
                        reduction_method=self.reduction_method,
                        hidden_size=hidden_size,
                        trunk_autocast_precision=self.autocast_precision,  # Trunk weight precision handled when defining the trunk
                        head_precision=torch.float32,  # Default precision for the head
                    ).to(device)
                    dtype = regressor.head_precision
                    optimizer = torch.optim.AdamW(  # TODO check the mixed precision thingy
                        regressor.parameters(),
                        lr=learning_rate,
                        weight_decay=weight_decay,
                        eps=1e-4,
                    )

                scheduler = get_linear_scheduler_with_warmup(
                    optimizer,
                    num_warmup_steps=int(0.1 * num_epochs * len(train_loader)),
                    num_training_steps=num_epochs * len(train_loader),
                )
                criterion = torch.nn.MSELoss()
                self.early_stopping.reset()  # Reset early stopping for each hyperparameter configuration

                if self.precomputed:
                    val_score, test_scores, completed_steps = self._train_mlp_precomputed(
                        mlp=mlp,
                        train_loader=train_loader,
                        validation_loader=validation_loader,
                        test_loader=test_loader,
                        optimizer=optimizer,
                        scheduler=scheduler,
                        criterion=criterion,
                        num_epochs=num_epochs,
                        label_dtype=dtype,
                        device=device,
                        do_test=do_test,
                        verbose=verbose,
                        accumulation_steps=accumulation_steps,
                    )
                elif not self.finetuning:
                    val_score, test_scores, completed_steps = self._train_regressor(
                        regressor=regressor,
                        train_loader=train_loader,
                        validation_loader=validation_loader,
                        test_loader=test_loader,
                        optimizer=optimizer,
                        scheduler=scheduler,
                        criterion=criterion,
                        num_epochs=num_epochs,
                        label_dtype=dtype,
                        device=device,
                        do_test=do_test,
                        verbose=verbose,
                        accumulation_steps=accumulation_steps,
                    )
                else:
                    val_score, test_scores, completed_steps = self._train_regressor(
                        regressor=regressor,
                        train_loader=train_loader,
                        validation_loader=validation_loader,
                        test_loader=test_loader,
                        optimizer=optimizer,
                        scheduler=scheduler,
                        criterion=criterion,
                        num_epochs=num_epochs,
                        finetuning=True,  # We are finetuning the trunk
                        label_dtype=dtype,
                        device=device,
                        do_test=do_test,
                        verbose=verbose,
                        accumulation_steps=accumulation_steps,
                    )
                # Store the scores for this hyperparameter and seed
                all_scores[hparam].append((val_score, test_scores, completed_steps))

            # Compute the average score over the seeds
            mean_val_score = np.mean([score[0] for score in all_scores[hparam]])
            std_val_score = np.std([score[0] for score in all_scores[hparam]])
            mean_test_score = {metric: np.mean([score[1][metric] for score in all_scores[hparam]]) for metric in self.metrics}
            std_test_score = {"std_" + metric: np.std([score[1][metric] for score in all_scores[hparam]]) for metric in self.metrics}
            completed_steps = min([s[2] for s in all_scores[hparam]])

            # Replace the list of scores with their aggregate mean
            all_scores[hparam] = {
                "validation_score": mean_val_score,
                "test_scores": mean_test_score,
                "validation_std": std_val_score,
                "test_std": std_test_score,
                "min_completed_steps": completed_steps,
            }
            global_pbar.update(1)

            # Checkpoint
            self.save_results_to_json(
                all_scores=all_scores,
                output_folder=output_folder,
                task_name=task_name,
                task_description=task_description,
            )

        global_pbar.close()

        # Find the best hyperparameter values based on the validation set
        best_hyperparameters = max(all_scores, key=lambda x: all_scores[x]["validation_score"])
        best_val_score = all_scores[best_hyperparameters].get("validation_score", 0)
        best_test_scores = all_scores[best_hyperparameters].get("test_scores", {})
        best_std_val_score = all_scores[best_hyperparameters].get("validation_std", 0)
        best_std_test_scores = all_scores[best_hyperparameters].get("test_std", {})
        best_completed_steps = all_scores[best_hyperparameters].get("min_completed_steps", -1)

        self.logger.info("Best hyperparameters: {}".format(best_hyperparameters))
        self.logger.info("Best validation score: {} +- {}".format(best_val_score, best_std_val_score))
        self.logger.info("Best test scores: {} +- {}".format(best_test_scores, best_std_test_scores))
        self.logger.info(f"Best completed steps: {best_completed_steps}")

        # Convert std_test_scores format (remove "std_" prefix)
        std_metrics = {k.replace("std_", ""): v for k, v in best_std_test_scores.items()}

        return create_standardized_result(
            metrics=best_test_scores,
            evaluator_type=self.head_type,
            best_hyperparameters=best_hyperparameters,
            std_metrics=std_metrics,
            all_results=all_scores,
            validation_score=best_val_score,
            validation_std=best_std_val_score,
            model_name=getattr(model, "name_or_path", ""),
            dataset_name=getattr(self, "dataset_name", ""),
        )
```
