## Supplementary material for "pLM representations unlock metagenomic space beyond homology": Code bundle: TokenClassificationEvaluator.py

```
import os
import ast
import json
from dataclasses import dataclass, field
from typing import List, Dict, Callable, Any, Optional, Union, Tuple
import torch
import torch.nn as nn
from torch.utils.data import DataLoader, Dataset
import numpy as np
import tqdm
from sklearn.metrics import accuracy_score, f1_score, precision_score, recall_score

from apperitif.evaluation.Evaluator import Evaluator, EvaluatorConfig
from apperitif.utils.dtype_utils import get_torch_dtype
from apperitif.utils.evaluator_utils import EvaluatorResult, create_standardized_result
from apperitif.utils.embedding_utils import get_token_embeddings_and_labels
from apperitif.utils.optimization import get_linear_scheduler_with_warmup, EarlyStopping
from apperitif.utils.deterministic import set_seed
from apperitif.data.TokenEmbeddingsDataset import TokenEmbeddingsDataset, token_embeddings_collate_fn
from apperitif.models.Encoder import TokenEncoder
from apperitif.models.regressor import Regressor, MLP
import logging


@dataclass
class TokenClassificationEvaluatorConfig(EvaluatorConfig):
    head_type: str = "base"  # Discriminator field for Hydra
    metrics: List[str] = field(default_factory=lambda: ["accuracy", "f1", "precision", "recall"])
    decision_metric: Optional[str] = None
    num_labels: Optional[int] = None  # Number of labels for token classification, can be set to None if not known beforehand
    reduction_method: str = "none"
    target_column: str = "labels"
    precomputing_batch_size: int = 128
    autocast_precision: str = "fp16"  # Changed to string for Hydra compatibility
    max_length: Optional[int] = None  # for truncating token-level labels


@dataclass
class MLPTokenClassificationEvaluatorConfig(TokenClassificationEvaluatorConfig):
    head_type: str = "mlp"  # Discriminator field for Hydra
    hidden_size: Union[int, List[int]] = 512
    learning_rate: Union[float, List[float]] = 1e-3
    num_epochs: Union[int, List[int]] = 2
    weight_decay: Union[float, List[float]] = 0
    batch_size: Union[int, List[int]] = 32
    seeds: List[int] = field(default_factory=lambda: [0, 1, 2, 3, 4])
    finetuning: bool = False
    precomputed: bool = True  # Moved to config


class TokenClassificationEvaluator(Evaluator):
    """
    Base evaluator for token classification tasks.
    """

    def __init__(
        self,
        config: TokenClassificationEvaluatorConfig,
        datasets: Dict[str, Dataset],
        collate_fn: Callable,
    ):
        super().__init__()
        self.logger = logging.getLogger(__name__)
        logging.basicConfig(filename=None, level=logging.INFO)

        self.config = config
        self.datasets = datasets
        self.collate_fn = collate_fn

        # Extract config values
        self.metrics = config.metrics
        assert all(metric in ["accuracy", "f1", "precision", "recall"] for metric in self.metrics), "Invalid metric(s)."
        assert len(self.metrics) > 0, "At least one metric must be provided."

        if config.decision_metric is None:
            self.decision_metric = self.metrics[0]
        else:
            assert config.decision_metric in self.metrics, f"Invalid decision metric: {config.decision_metric}"
            self.decision_metric = config.decision_metric

        self.num_labels = config.num_labels
        assert self.num_labels is None or self.num_labels > 1, "Number of labels must be greater than 1 or None."
        self.reduction_method = config.reduction_method
        self.target_column = config.target_column
        self.precomputing_batch_size = config.precomputing_batch_size
        self.autocast_precision = get_torch_dtype(config.autocast_precision)
        self.head_type = config.head_type  # For standardized output

    @property
    def name(cls):
        return "token_classification"

    def _compute_metric(self, labels: np.ndarray, preds: np.ndarray, metric: str) -> float:
        """
        Compute the value of the given metric for token classification.

        Args:
            labels: True labels.
            preds: Predicted probabilities or logits.
            metric: Metric to compute.

        Returns:
            Value of the metric.
        """
        if metric == "accuracy":
            return accuracy_score(labels, np.argmax(preds, axis=1))
        elif metric == "f1":
            return f1_score(labels, np.argmax(preds, axis=1), average="weighted")
        elif metric == "precision":
            return precision_score(labels, np.argmax(preds, axis=1), average="weighted")
        elif metric == "recall":
            return recall_score(labels, np.argmax(preds, axis=1), average="weighted")
        else:
            raise ValueError(f"Invalid metric: {metric}")

    def __call__(self, model: TokenEncoder, **kwargs) -> EvaluatorResult:
        """
        Evaluate the model on a token classification task.

        Args:
            model: TokenEncoder model to evaluate.

        Returns:
            Dictionary containing the evaluation metrics on the test set.
        """
        raise NotImplementedError


class MLPTokenClassificationEvaluator(TokenClassificationEvaluator):
    def __init__(
        self,
        config: MLPTokenClassificationEvaluatorConfig,
        datasets: Dict[str, Dataset],
        collate_fn: Callable,
    ):
        super().__init__(config, datasets, collate_fn)
        self.hidden_size = config.hidden_size
        self.learning_rate = config.learning_rate
        self.num_epochs = config.num_epochs
        self.weight_decay = config.weight_decay
        self.batch_size = config.batch_size
        self.finetuning = config.finetuning  # Whether to fine-tune the trunk or not
        self.precomputed = config.precomputed  # Whether to use precomputed embeddings
        assert not self.precomputed if self.finetuning else True, "Cannot finetune with precomputed embeddings."

        if isinstance(self.hidden_size, int):
            self.hidden_size = [self.hidden_size]
        if isinstance(self.learning_rate, float):
            self.learning_rate = [self.learning_rate]
        if isinstance(self.num_epochs, int):
            self.num_epochs = [self.num_epochs]
        if isinstance(self.weight_decay, float):
            self.weight_decay = [self.weight_decay]
        if isinstance(self.batch_size, int):
            self.batch_size = [self.batch_size]

        self.seeds = config.seeds
        set_seed(self.seeds[0])

        self.early_stopping = EarlyStopping(patience=16, mode="max", disable=False)

    @property
    def name(cls):
        return "mlp_token_classification"

    def _evaluate_mlp_precomputed(self, mlp: torch.nn.Module, embedding_loader: DataLoader, device: str = "cuda") -> Dict[str, float]:
        mlp.eval()
        all_logits = []
        all_labels = []
        with torch.no_grad():
            for batch in embedding_loader:
                embeddings = batch["embeddings"].to(device)
                batch_labels = batch["labels"]
                logits = mlp(embeddings)
                all_logits.append(logits.float().cpu().numpy())
                all_labels.append(batch_labels.long().cpu().numpy())

        all_logits = np.concatenate(all_logits, axis=0)
        all_labels = np.concatenate(all_labels, axis=0)
        all_probas = torch.nn.functional.softmax(torch.tensor(all_logits), dim=1).numpy()

        scores = {}
        for metric in self.metrics:
            scores[metric] = self._compute_metric(all_labels, all_probas, metric)
        return scores

    def _train_mlp_precomputed(
        self,
        mlp: torch.nn.Module,
        train_loader: DataLoader,
        validation_loader: DataLoader,
        test_loader: DataLoader,
        optimizer: torch.optim.Optimizer,
        scheduler: torch.optim.lr_scheduler.LRScheduler | None,
        criterion: Callable,
        num_epochs: int,
        device: str = "cuda",
        do_test: bool = True,
        verbose: Union[bool, str] = "reduced",
        accumulation_steps: int = 1,
    ) -> Tuple[float, Dict[str, float]]:
        disable = (not verbose) or verbose == "reduced"
        pbar = tqdm.tqdm(total=(num_epochs) * len(train_loader) // accumulation_steps, desc="Training MLP", disable=disable)
        best_val_score = -1
        loader_len = len(train_loader)
        should_stop = False
        completed_steps = 0

        with torch.autocast(device_type=device, dtype=torch.float32, enabled=True):
            for n in range(num_epochs):
                mlp.train()
                for idx, batch in enumerate(train_loader):
                    optimizer.zero_grad()
                    embeddings = batch["embeddings"].to(device)
                    labels = batch["labels"].to(device).long()
                    preds = mlp(embeddings)
                    loss = criterion(preds, labels) / accumulation_steps  # Normalize loss for accumulation
                    loss.backward()

                    if (idx + 1) % accumulation_steps == 0 or (idx + 1 == loader_len):
                        optimizer.step()
                        optimizer.zero_grad()
                        if scheduler is not None:
                            scheduler.step()

                        completed_steps += 1
                        pbar.update(1)

                    if ((idx + 1) % (loader_len // 3) == 0) or (idx + 1 == loader_len):
                        val_scores = self._evaluate_mlp_precomputed(mlp, validation_loader, device=device)
                        val_score = val_scores[self.decision_metric]
                        pbar.set_postfix({"val_score": val_score})
                        if val_score > best_val_score:
                            best_val_score = val_score
                            best_state_dict = {k: v.cpu() for k, v in mlp.state_dict().items()}
                        should_stop = self.early_stopping(val_score)
                        if should_stop:
                            break
                        mlp.train()
                if should_stop:
                    break
            pbar.close()

        if do_test:
            mlp.load_state_dict({k: v.to(device) for k, v in best_state_dict.items()})
            test_scores = self._evaluate_mlp_precomputed(mlp, test_loader, device=device)
        else:
            test_scores = {metric: -1 for metric in self.metrics}
        return best_val_score, test_scores, completed_steps

    def _evaluate_classifier(self, classifier: Regressor, dataloader: DataLoader, device: str = "cuda") -> Dict[str, float]:
        """
        Evaluate a classifier (regressor) on token classification data.

        Args:
            classifier: The regressor model to evaluate
            dataloader: DataLoader for evaluation data
            device: Device to run on

        Returns:
            Dictionary of metric scores
        """
        classifier.eval()
        all_logits = []
        all_labels = []

        with torch.no_grad():
            for batch in dataloader:
                # Extract labels and move to device
                labels = batch.pop(self.target_column).to(device).long()

                # Get predictions from classifier
                # The classifier will handle the token encoder internally
                logits = classifier(batch, frozen_backbone=True)

                # Flatten logits and labels for token-level evaluation
                # logits: [batch_size, seq_len, num_labels]
                # labels: [batch_size, seq_len]
                # batch_size, seq_len, num_labels = logits.shape
                # logits_flat = logits.view(-1, num_labels)  # [batch_size * seq_len, num_labels]
                # labels_flat = labels.view(-1)  # [batch_size * seq_len]

                # Filter out -100 labels (padding/special tokens)
                valid_mask = labels != -100
                if valid_mask.any():
                    valid_logits = logits[valid_mask]
                    valid_labels = labels[valid_mask]

                    all_logits.append(valid_logits.cpu().numpy())
                    all_labels.append(valid_labels.cpu().numpy())

        if not all_logits:
            # No valid tokens found
            return {metric: 0.0 for metric in self.metrics}

        all_logits = np.concatenate(all_logits, axis=0)
        all_labels = np.concatenate(all_labels, axis=0)
        all_probas = torch.nn.functional.softmax(torch.tensor(all_logits), dim=1).numpy()

        scores = {}
        for metric in self.metrics:
            scores[metric] = self._compute_metric(all_labels, all_probas, metric)
        return scores

    def _train_classifier(
        self,
        classifier: Regressor,
        train_loader: DataLoader,
        validation_loader: DataLoader,
        test_loader: DataLoader,
        optimizer: torch.optim.Optimizer,
        scheduler: torch.optim.lr_scheduler.LRScheduler | None,
        criterion: Callable,
        num_epochs: int,
        finetuning: bool = False,
        device: str = "cuda",
        do_test: bool = True,
        verbose: Union[bool, str] = "reduced",
        accumulation_steps: int = 1,
    ) -> Tuple[float, Dict[str, float]]:
        """
        Train a classifier (regressor) on token classification data.

        Args:
            classifier: The regressor model to train
            train_loader: DataLoader for training data
            validation_loader: DataLoader for validation data
            test_loader: DataLoader for test data
            optimizer: Optimizer for training
            scheduler: Learning rate scheduler
            criterion: Loss function
            num_epochs: Number of training epochs
            finetuning: Whether to finetune the backbone
            device: Device to run on
            do_test: Whether to evaluate on test set

        Returns:
            Tuple of (best_val_score, test_scores)
        """
        disable_pbar = (not verbose) or verbose == "reduced"
        pbar = tqdm.tqdm(total=(num_epochs) * len(train_loader) // accumulation_steps, desc="Training Classifier", disable=disable_pbar)

        best_val_score = -1
        best_state_dict = None
        loader_len = len(train_loader)
        should_stop = False
        completed_steps = 0

        with torch.autocast(device_type=device, dtype=torch.float32, enabled=True):
            for epoch in range(num_epochs):
                classifier.train()
                frozen_backbone = not finetuning or epoch == 0
                for idx, batch in enumerate(train_loader):
                    optimizer.zero_grad()

                    # Extract labels and move to device
                    labels = batch.pop(self.target_column).to(device).long()

                    # Get predictions from classifier
                    logits = classifier(batch, frozen_backbone=frozen_backbone)

                    # Compute loss (CrossEntropyLoss will ignore -100 labels)
                    loss = criterion(logits, labels) / accumulation_steps
                    loss.backward()

                    if (idx + 1) % accumulation_steps == 0 or (idx + 1 == loader_len):
                        optimizer.step()
                        optimizer.zero_grad()
                        if scheduler is not None:
                            scheduler.step()

                        completed_steps += 1
                        pbar.update(1)

                    # Evaluate on validation set periodically
                    if ((idx + 1) % (loader_len // 3) == 0) or (idx + 1 == loader_len):
                        val_scores = self._evaluate_classifier(classifier, validation_loader, device=device)
                        val_score = val_scores[self.decision_metric]
                        pbar.set_postfix({"val_score": val_score})

                        if val_score > best_val_score:
                            best_val_score = val_score
                            best_state_dict = {k: v.cpu() for k, v in classifier.state_dict().items()}

                        should_stop = self.early_stopping(val_score)
                        if should_stop:
                            break
                        classifier.train()

                if should_stop:
                    break

            pbar.close()

        # Evaluate on test set with best model
        if do_test and test_loader is not None:
            classifier.load_state_dict({k: v.to(device) for k, v in best_state_dict.items()})
            test_scores = self._evaluate_classifier(classifier, test_loader, device=device)
        else:
            test_scores = {metric: -1 for metric in self.metrics}

        return best_val_score, test_scores, completed_steps

    def __call__(
        self,
        model: TokenEncoder,
        model_kwargs: Dict[str, Any] = None,
        return_all_scores: bool = False,
        do_test: bool = True,
        verbose: Union[bool, str] = "reduced",
        accumulation_steps: int = None,
        output_folder: str = None,
        task_name: Optional[str] = None,
        task_description: Optional[str] = None,
    ) -> EvaluatorResult:
        """
        Evaluate the model on a token classification task using MLP.

        Args:
            model: TokenEncoder model to evaluate.
            model_kwargs: Additional keyword arguments to pass to the model.
            return_all_scores: Whether to return all scores or just the best.
            do_test: Whether to evaluate on the test set.

        Returns:
            EvaluatorResult containing the evaluation metrics.
        """
        device = next(model.parameters()).device.type

        if self.precomputed:
            # Extract embeddings once and create datasets
            dataloaders = {
                name: DataLoader(
                    ds,
                    batch_size=self.precomputing_batch_size,
                    collate_fn=self.collate_fn,
                    shuffle=False,
                )
                for name, ds in self.datasets.items()
            }

            # Get token embeddings and labels
            train_data = get_token_embeddings_and_labels(
                model, dataloaders["train"], self.target_column, device, model_kwargs, self.autocast_precision
            )
            validation_data = get_token_embeddings_and_labels(
                model, dataloaders["validation"], self.target_column, device, model_kwargs, self.autocast_precision
            )

            if do_test and "test" in dataloaders:
                test_data = get_token_embeddings_and_labels(
                    model, dataloaders["test"], self.target_column, device, model_kwargs, self.autocast_precision
                )
            else:
                test_data = []

            # Create embedding datasets
            train_embeddings = [item["embeddings"] for item in train_data]
            train_labels = [item["labels"] for item in train_data]
            train_dataset = TokenEmbeddingsDataset(train_embeddings, train_labels)

            validation_embeddings = [item["embeddings"] for item in validation_data]
            validation_labels = [item["labels"] for item in validation_data]
            validation_dataset = TokenEmbeddingsDataset(validation_embeddings, validation_labels)

            if test_data:
                test_embeddings = [item["embeddings"] for item in test_data]
                test_labels = [item["labels"] for item in test_data]
                test_dataset = TokenEmbeddingsDataset(test_embeddings, test_labels)
            else:
                test_dataset = None

            # Determine input dimension and number of labels
            input_dim = train_embeddings[0].shape[-1]
            if self.num_labels is None:
                all_labels = torch.cat(train_labels)
                self.num_labels = len(torch.unique(all_labels))

            train_collate_fn = token_embeddings_collate_fn
        else:
            # Online evaluation - use original datasets
            train_dataset = self.datasets["train"]
            validation_dataset = self.datasets["validation"]
            test_dataset = self.datasets.get("test") if do_test else None
            train_collate_fn = self.collate_fn

            # Determine number of labels if not provided
            if self.num_labels is None:
                # Get a sample batch to determine num_labels
                sample_dataloader = DataLoader(train_dataset, batch_size=1, collate_fn=self.collate_fn, shuffle=False)
                sample_batch = next(iter(sample_dataloader))
                sample_labels = sample_batch[self.target_column]
                # Remove -100 values and get unique labels
                valid_labels = sample_labels[sample_labels != -100]
                self.num_labels = len(torch.unique(valid_labels))

        # Create hyperparameter grid
        hyperparameters = [
            (hs, lr, ne, wd, bs)
            for bs in self.batch_size
            for hs in self.hidden_size
            for lr in self.learning_rate
            for ne in self.num_epochs
            for wd in self.weight_decay
        ]

        # Grid search over hyperparameters
        all_scores = {}

        # Load previously computed hyperparameters if available
        if output_folder is not None and os.path.exists(os.path.join(output_folder, "results.json")):
            file_path = os.path.join(output_folder, "results.json")
            with open(file_path, "rt") as f:
                results_dict = json.load(f)
            all_scores = results_dict["all_results"]
            hyperparameters_done = [ast.literal_eval(s) for s in list(results_dict["all_results"].keys())]

            hyperparameters = set(hyperparameters) - set(hyperparameters_done)
            self.logger.info(
                f"Found {len(hyperparameters_done)} previously computed hyperparameters. Only {len(hyperparameters)} remaining to compute."
            )

        global_pbar = tqdm.tqdm(
            total=len(hyperparameters),
            desc="Hyperparameter search",
            disable=(not verbose) or (verbose != "reduced"),
            smoothing=0,
        )

        for hparam in hyperparameters:
            hidden_size, learning_rate, num_epochs, weight_decay, batch_size = hparam
            all_scores[hparam] = []

            if accumulation_steps is None or batch_size % accumulation_steps != 0:
                accumulation_steps = max(1, batch_size // 8)

            print("Hyperparameters:", hparam)
            print("Accumulation steps:", accumulation_steps)

            # Create data loaders
            train_loader = DataLoader(train_dataset, batch_size=batch_size // accumulation_steps, shuffle=True, collate_fn=train_collate_fn)
            validation_loader = DataLoader(validation_dataset, batch_size=batch_size, shuffle=False, collate_fn=train_collate_fn)
            if do_test and test_dataset is not None:
                test_loader = DataLoader(test_dataset, batch_size=batch_size, shuffle=False, collate_fn=train_collate_fn)
            else:
                test_loader = None

            for seed in self.seeds:
                set_seed(seed)

                if self.precomputed:
                    # Create MLP for precomputed embeddings
                    mlp = MLP(input_dim, hidden_size, self.num_labels).to(device)
                    optimizer = torch.optim.AdamW(mlp.parameters(), lr=learning_rate, weight_decay=weight_decay, eps=1e-4)
                elif not self.finetuning:
                    # Create regressor with frozen backbone
                    model.eval()
                    classifier = Regressor(
                        trunk=model,
                        hidden_size=hidden_size,
                        output_size=self.num_labels,
                        reduction_method=self.reduction_method,
                        trunk_autocast_precision=self.autocast_precision,
                        head_precision=torch.float32,
                    ).to(device)
                    optimizer = torch.optim.AdamW(classifier.head.parameters(), lr=learning_rate, weight_decay=weight_decay, eps=1e-4)
                else:  # Finetuning
                    model.train()
                    classifier = Regressor(
                        trunk=model,
                        hidden_size=hidden_size,
                        output_size=self.num_labels,
                        reduction_method=self.reduction_method,
                        trunk_autocast_precision=self.autocast_precision,
                        head_precision=torch.float32,
                    ).to(device)
                    optimizer = torch.optim.AdamW(classifier.parameters(), lr=learning_rate, weight_decay=weight_decay, eps=1e-4)

                scheduler = get_linear_scheduler_with_warmup(
                    optimizer,
                    num_warmup_steps=int(0.1 * num_epochs * len(train_loader)),
                    num_training_steps=num_epochs * len(train_loader),
                )
                criterion = nn.CrossEntropyLoss(ignore_index=-100)  # Ignore -100 labels
                self.early_stopping.reset()

                # Train and evaluate
                if self.precomputed:
                    val_score, test_scores, completed_steps = self._train_mlp_precomputed(
                        mlp,
                        train_loader,
                        validation_loader,
                        test_loader,
                        optimizer,
                        scheduler,
                        criterion,
                        num_epochs,
                        device,
                        do_test,
                        verbose=verbose,
                        accumulation_steps=accumulation_steps,
                    )
                else:  # Handles both finetuning and head-only training
                    val_score, test_scores, completed_steps = self._train_classifier(
                        classifier,
                        train_loader,
                        validation_loader,
                        test_loader,
                        optimizer,
                        scheduler,
                        criterion,
                        num_epochs,
                        self.finetuning,
                        device,
                        do_test,
                        verbose=verbose,
                        accumulation_steps=accumulation_steps,
                    )
                all_scores[hparam].append((val_score, test_scores, completed_steps))

            # Compute mean and std for this hyperparameter configuration
            mean_val_score = np.mean([s[0] for s in all_scores[hparam]])
            std_val_score = np.std([s[0] for s in all_scores[hparam]])
            mean_test_scores = {
                metric: np.mean([s[1][metric] for s in all_scores[hparam] if s[1][metric] != -1]) for metric in self.metrics
            }
            std_test_scores = {
                "std_" + metric: np.std([s[1][metric] for s in all_scores[hparam] if s[1][metric] != -1]) for metric in self.metrics
            }
            completed_steps = min([s[2] for s in all_scores[hparam]])
            all_scores[hparam] = {
                "validation_score": mean_val_score,
                "test_scores": mean_test_scores,
                "validation_std": std_val_score,
                "test_std": std_test_scores,
                "min_completed_steps": completed_steps,
            }
            global_pbar.update(1)

            # Checkpoint
            self.save_results_to_json(
                all_scores=all_scores,
                output_folder=output_folder,
                task_name=task_name,
                task_description=task_description,
            )

        global_pbar.close()

        # Find best hyperparameters
        best_hyperparameters = max(all_scores, key=lambda x: all_scores[x]["validation_score"])

        best_val_score = all_scores[best_hyperparameters].get("validation_score", 0)
        best_test_scores = all_scores[best_hyperparameters].get("test_scores", {})
        best_std_val_score = all_scores[best_hyperparameters].get("validation_std", 0)
        best_std_test_scores = all_scores[best_hyperparameters].get("test_std", {})
        best_completed_steps = all_scores[best_hyperparameters].get("min_completed_steps", -1)

        self.logger.info(f"Best hyperparameters: {best_hyperparameters}")
        self.logger.info(f"Best validation score: {best_val_score:.4f} +- {best_std_val_score:.4f}")
        self.logger.info(f"Best test scores: {best_test_scores} +- {best_std_test_scores}")
        self.logger.info(f"Best completed steps: {best_completed_steps}")

        # Create result
        result = create_standardized_result(
            metrics=best_test_scores,
            evaluator_type=self.head_type,
            best_hyperparameters=best_hyperparameters,
            std_metrics=best_std_test_scores,
            all_results=all_scores if return_all_scores else None,
            validation_score=best_val_score,
            validation_std=best_std_val_score,
            dataset_name=getattr(self, "dataset_name", "unknown"),
        )

        self.logger.info(f"Created EvaluatorResult: metrics={result.metrics}, evaluator_name={result.evaluator_name}")
        return result
```
