## Supplementary material for "pLM representations unlock metagenomic space beyond homology": Code bundle: Amplify.py

```
from typing import Any, Dict, Optional, Union, List
from dataclasses import dataclass, field

import numpy as np
import os
import torch
from torch import Tensor
from transformers import AutoModel, AutoTokenizer, AutoConfig

from .Encoder import (
    SequenceEncoder,
    TokenEncoder,
    EncoderConfig,
)
from apperitif.collator import DataCollatorForSeqTasks

from apperitif.utils.dtype_utils import get_torch_dtype
from amplify.model import AMPLIFY


@dataclass
class AmplifyConfig(EncoderConfig):
    # Amplify-specific configurations only
    # Note: model_name and model_type are inherited from ModelConfig
    # Note: hf_kwargs, use_hidden_layer, etc. are inherited from EncoderConfig
    local_name: str = None  # Local name if different than model_name
    model_type: str = "amplify"  # Default model type for AMPLIFY models
    model_name: str = "chandar-lab/AMPLIFY_120M"  # AMPLIFY model name on the Hugging Face Hub
    attention_mask_type: str = "additive"
    hf_kwargs: Dict[str, Any] = field(default_factory=lambda: {"trust_remote_code": True})
    max_length: Optional[int] = 2048  # Default max length for padding/truncation
    attention_mask_dtype: str = "fp16"  # Default attention mask type for AMPLIFY models (half precision needed for inference)
    use_attn_maps: bool = True
    num_total_attn_maps: Optional[int] = 240  # Total number of attention maps of AMPLIFY 120M
    from_hub: bool = True
    base_path: str = None
    vocab_path: str = None
    tag: str = None
    randomize: bool = False
    randomize_mode: str = "distribution"


class AmplifyModelMixin:
    def __init__(self, config: AmplifyConfig) -> None:
        print("config.from_hub", config.from_hub)
        if config.from_hub:
            print(config.model_name, config.hf_kwargs)
            self.model = AutoModel.from_pretrained(config.model_name, **config.hf_kwargs)
            self.tokenizer = AutoTokenizer.from_pretrained(config.model_name, **config.hf_kwargs)
        else:
            checkpoint_path = os.path.join(config.base_path, config.model_name, "model_checkpoints")
            config_path = os.path.join(config.base_path, config.model_name, ".hydra/config.yaml")
            self.model, self.tokenizer = AMPLIFY.load(
                checkpoint_path=checkpoint_path, config_path=config_path, vocab_path=config.vocab_path, tag=config.tag
            )
        if config.randomize:
            # Create a copy of the architecture
            model_config = AutoConfig.from_pretrained(config.model_name, **config.hf_kwargs)
            randomized_model = AutoModel.from_config(model_config)

            with torch.no_grad():
                if config.randomize_mode == "distribution":
                    for (name_pretrained, param_pretrained), (name_random, param_random) in zip(
                        self.model.named_parameters(), randomized_model.named_parameters()
                    ):
                        if param_pretrained.shape != param_random.shape:
                            raise ValueError(f"Shape mismatch for {name_pretrained}")

                        # Get stats from pretrained weights
                        tensor = param_pretrained.data
                        mean = tensor.mean()
                        std = tensor.std()

                        # Re-sample from the same distribution
                        param_random.data.copy_(torch.normal(mean=mean, std=std, size=tensor.shape).to(tensor.device))

                else:
                    if hasattr(randomized_model, "_init_weights"):
                        randomized_model.apply(randomized_model._init_weights)
                    elif hasattr(randomized_model, "init_weights"):
                        randomized_model.apply(randomized_model.init_weights)
                    else:
                        raise AttributeError("Model does not have an init_weights or _init_weights method.")

                diffs = []
                for (name_pretrained, p_pretrained), (name_random, p_random) in zip(
                    self.model.named_parameters(), randomized_model.named_parameters()
                ):
                    if p_pretrained.norm() == 0:
                        continue
                    rel_diff = (p_random - p_pretrained).norm() / p_pretrained.norm()
                    diffs.append(rel_diff.item())
                avg_diff = sum(diffs) / len(diffs)
                print(f"Average relative norm difference: {avg_diff:.4f}")

                self.model = randomized_model

        self.attention_mask_type = config.attention_mask_type
        self.attention_mask_dtype = get_torch_dtype(config.attention_mask_dtype)

        # For layer selection in the trunk forward pass
        if self.use_packed_sequence:
            self.total_num_layers = self.model.config.num_hidden_layers
        # Correct the hidden size initialized at 1 in the super class
        self.hidden_size = self.model.config.hidden_size * self.use_hidden_layer


class AmplifySequenceEncoder(SequenceEncoder, AmplifyModelMixin):
    def __init__(self, config: AmplifyConfig, **kwargs: Any):
        super().__init__(config, **kwargs)

        # Initialize model and tokenizer via AmplifyModelMixin
        AmplifyModelMixin.__init__(self, config)

    def forward(
        self,
        batch,
        reduction_method: str = "mean",
    ):
        if self.use_packed_sequence:
            # TODO implement a warning about the number of hidden_states returned
            input_ids, max_seqlen, cu_seqlens, attention_mask, position_ids = (
                batch["input_ids"],
                batch.get("max_seqlen", None),
                batch.get("cu_seqlens", None),
                batch.get("attention_mask", None),
                batch.get("position_ids", None),
            )
            if self.config.from_hub:
                output = self.model(
                    src=input_ids,
                    position_ids=position_ids,
                    max_seqlen=max_seqlen,
                    cu_seqlens=cu_seqlens,
                    pad_mask=attention_mask,
                    output_hidden_states=self.total_num_layers - self.use_hidden_layer,  # Only need the last layers
                )
            else:
                output = self.model(
                    input_ids=input_ids,
                    position_ids=position_ids,
                    max_seqlen=max_seqlen,
                    cu_seqlens=cu_seqlens,
                    attention_mask=attention_mask,
                    output_hidden_states=self.total_num_layers - self.use_hidden_layer,  # Only need the last layers
                )
            output["hidden_states"] = torch.concatenate(output["hidden_states"], dim=-1)
            mean_embeddings = []
            for seq_start, seq_end in zip(cu_seqlens[:-1], cu_seqlens[1:]):
                mean_embeddings.append(self._get_embeddings(output, None, seq_start, seq_end, reduction_method))
        else:
            output = self.model(**batch, output_hidden_states=True)
            mean_embeddings = []
            output["hidden_states"] = torch.cat(
                output["hidden_states"][-self.use_hidden_layer :],  # Default AMPLIFY implementation is all or nothing for hidden states
                dim=-1,
            )
            # TODO try with the torch.isin function to get all special tokens at once? Or maybe not if we want to keep BOS/EOS?
            for i in range(batch["input_ids"].shape[0]):
                if self.attention_mask_type == "additive":
                    seq_mask = batch["attention_mask"][i] == 0
                elif self.attention_mask_type == "multiplicative":
                    seq_mask = batch["attention_mask"][i] == 1
                elif self.attention_mask_type == "bool":
                    seq_mask = batch["attention_mask"][i].bool()  # TODO WTF? ALso we don't have bool masks in AMPLIFY for now. Remove this?
                mean_embeddings.append(
                    self._get_embeddings(output=output, seq_mask=seq_mask, seq_start=i, reduction_method=reduction_method)
                )

        return torch.stack(mean_embeddings)

    def get_collate_fn(
        self,
        sequence_column: str = "seq",
        label_column: str = "label",
        label_dtype: torch.dtype = torch.long,
        device=None,
        **kwargs,
    ):
        return DataCollatorForSeqTasks(
            tokenizer=self.tokenizer,
            pack_sequences=self.use_packed_sequence,
            device=device,
            sequence_column=sequence_column,
            max_length=self.max_length,
            label_column=label_column,
            label_dtype=label_dtype,
            token_level_task=False,
            attention_mask_type=self.attention_mask_type,
            attention_mask_dtype=self.attention_mask_dtype,
            **kwargs,
        )


# DO not forget to remove the special tokens in token level tasks.


class AmplifyTokenEncoder(TokenEncoder):
    def __init__(self, config: AmplifyConfig):
        super(AmplifyTokenEncoder, self).__init__(config)

        # Initialize model and tokenizer via AmplifyModelMixin
        AmplifyModelMixin.__init__(self, config)

        self.padding_token_id = getattr(config, "padding_token_id", 0)

        # Initialize special tokens IDs for masking
        self.special_tokens_ids = []
        if hasattr(self.tokenizer, "pad_token_id") and self.tokenizer.pad_token_id is not None:
            self.special_tokens_ids.append(self.tokenizer.pad_token_id)
        if hasattr(self.tokenizer, "bos_token_id") and self.tokenizer.bos_token_id is not None:
            self.special_tokens_ids.append(self.tokenizer.bos_token_id)
        if hasattr(self.tokenizer, "eos_token_id") and self.tokenizer.eos_token_id is not None:
            self.special_tokens_ids.append(self.tokenizer.eos_token_id)
        if hasattr(self.tokenizer, "unk_token_id") and self.tokenizer.unk_token_id is not None:
            self.special_tokens_ids.append(self.tokenizer.unk_token_id)
        if hasattr(self.tokenizer, "mask_token_id") and self.tokenizer.mask_token_id is not None:
            self.special_tokens_ids.append(self.tokenizer.mask_token_id)
        self.special_tokens_ids = torch.tensor(self.special_tokens_ids, dtype=torch.long)

    def forward(
        self,
        batch: Dict[str, Tensor],
        reduction_method: str = None,  # Shape: (batch_size, seq_len) (With padding)
        return_individual_sequences: bool = False,
    ) -> Union[Tensor, List[Tensor]]:  # Shape: (num_standard_tokens_in_batch, hidden_size) OR List[(seq_len_i, hidden_size)]
        if self.use_packed_sequence:
            # TODO implement a warning about the number of hidden_states returned
            input_ids, max_seqlen, cu_seqlens, attention_mask, position_ids = (
                batch["input_ids"],
                batch.get("max_seqlen", None),
                batch.get("cu_seqlens", None),
                batch.get("attention_mask", None),
                batch.get("position_ids", None),
            )
            if self.config.from_hub:
                output = self.model(
                    src=input_ids,
                    position_ids=position_ids,
                    max_seqlen=max_seqlen,
                    cu_seqlens=cu_seqlens,
                    pad_mask=attention_mask,
                    output_hidden_states=self.total_num_layers - self.use_hidden_layer,  # Only need the last layers
                )
            else:
                output = self.model(
                    input_ids=input_ids,
                    position_ids=position_ids,
                    max_seqlen=max_seqlen,
                    cu_seqlens=cu_seqlens,
                    attention_mask=attention_mask,
                    output_hidden_states=self.total_num_layers - self.use_hidden_layer,  # Only need the last layers
                )

            # TODO remove there is no padding
            output["hidden_states"] = torch.concatenate(output["hidden_states"], dim=-1)
            global_seq_mask = (
                (batch["input_ids"] != self.padding_token_id).squeeze(0).bool()
            )  # Shape: (total_num_tokens_in_batch,)# TODO replace with attention mask?
            embeddings = []
            for seq_start, seq_end in zip(cu_seqlens[:-1], cu_seqlens[1:]):
                seq_mask = global_seq_mask[seq_start:seq_end]
                temp_output = self._get_embeddings(output, seq_mask, seq_start, seq_end)
                embeddings.append(temp_output)
        else:
            output = self.model(**batch, output_hidden_states=True)
            output["hidden_states"] = torch.cat(
                output["hidden_states"][-self.use_hidden_layer :],  # Default AMPLIFY implementation is all or nothing for hidden states
                dim=-1,
            )
            embeddings = []
            for i in range(batch["input_ids"].shape[0]):
                if self.attention_mask_type == "additive":
                    seq_mask = batch["attention_mask"][i] == 0
                elif self.attention_mask_type == "multiplicative":
                    seq_mask = batch["attention_mask"][i] == 1
                elif self.attention_mask_type == "bool":
                    seq_mask = batch["attention_mask"][i].bool()
                else:
                    raise ValueError(f"Unknown attention mask type: {self.attention_mask_type}")
                embeddings.append(
                    self._get_embeddings(
                        output,
                        seq_mask,
                        seq_start=i,
                    )
                )

        if return_individual_sequences:
            return embeddings  # List[(seq_len_i, hidden_size)]
        else:
            out = torch.concatenate(embeddings, dim=0)
            return out

    def get_collate_fn(
        self,
        sequence_column: str = "seq",
        label_column: str = "label",
        label_dtype: torch.dtype = torch.long,
        device=None,
        **kwargs,
    ):
        return DataCollatorForSeqTasks(
            tokenizer=self.tokenizer,
            pack_sequences=self.use_packed_sequence,
            device=device,
            sequence_column=sequence_column,
            max_length=self.max_length,
            label_column=label_column,
            label_dtype=label_dtype,
            token_level_task=True,
            attention_mask_type=self.attention_mask_type,
            attention_mask_dtype=self.attention_mask_dtype,
            **kwargs,
        )
```
