## Supplementary material for "pLM representations unlock metagenomic space beyond homology": Code bundle: Encoder.py

```
from typing import Any, Dict, Union
import torch
from torch import Tensor
from dataclasses import dataclass, field
from typing import Optional

from transformers import PreTrainedModel

# TODO we need to use configs to pass the model configuration and everything else


@dataclass
class EncoderConfig:
    model_type: Optional[str] = None  # Type of model, e.g., "esm", "amplify"
    model_name: Optional[str] = None  # Name of the model, e.g., "facebook/esm2_t12_35M_UR50D"
    hf_kwargs: Optional[Dict[str, Any]] = field(default_factory=dict)  # Hugging Face model kwargs to load from the Hub
    local_kwargs: Optional[Dict[str, Any]] = field(default_factory=dict)  # Local model kwargs to load locally
    max_length: Optional[int] = 1_000_000_000  # Maximum sequence length for padding/truncation PLACEHOLDER
    use_hidden_layer: int = 1
    normalize_embeddings: bool = False
    use_packed_sequence: bool = False
    use_attn_maps: bool = False  # Whether to use attention maps
    num_total_attn_maps: Optional[int] = None  # Total number of attention maps


class Encoder(torch.nn.Module):
    def __init__(self, config: EncoderConfig, **kwargs: Any) -> None:
        super().__init__(**kwargs)
        self.config = config
        self.model_name = getattr(config, "model_name", None)
        self.model_type = getattr(config, "model_type", None)
        self.use_hidden_layer = getattr(config, "use_hidden_layer", 1)
        self.normalize_embeddings = getattr(config, "normalize_embeddings", True)
        self.use_packed_sequence = getattr(config, "use_packed_sequence", False)
        self.max_length = getattr(config, "max_length", 1_000_000_000)
        self.hidden_size = getattr(config, "hidden_size", 1)  # Placeholder, will be set by the model

    def forward(  # PreTrainedModel inherits from torch.nn.Module so Encoder is callable
        self, batch: Dict[str, Tensor], **kwargs: Any
    ) -> Dict[str, Any]:
        pass


class SequenceEncoder(Encoder):
    def __init__(self, config: EncoderConfig, **kwargs: Any) -> None:
        super().__init__(config, **kwargs)

    def _get_embeddings(
        self,
        output,
        seq_mask: Tensor = None,
        seq_start: int = None,
        seq_end: int = None,
        reduction_method: str = "mean",
    ):  # TODO this could be fused with the appending and be one function for the whole batch
        if self.use_packed_sequence:
            embeddings = output["hidden_states"][0][seq_start:seq_end]
        else:
            embeddings = output["hidden_states"][seq_start]
            if seq_mask is not None:
                embeddings = embeddings[seq_mask]

        if self.normalize_embeddings:
            embeddings = torch.nn.functional.normalize(embeddings, p=2, dim=-1)

        if reduction_method == "mean":
            mean_embeddings = embeddings.mean(dim=0)
            return mean_embeddings
        if reduction_method == "sum":
            sum_embeddings = embeddings.sum(dim=0)
            return sum_embeddings
        if reduction_method == "first" or reduction_method == "cls":
            return embeddings[0]

    def forward(
        self,
        batch: Dict[str, Tensor],  # Shape: (batch_size, seq_len) (With padding)
        reduction_method: str = "mean",
        **kwargs: Any,
    ) -> Tensor:  # Shape: (batch_size, hidden_size)
        pass


class TokenEncoder(Encoder):
    def __init__(self, config: EncoderConfig, **kwargs: Any) -> None:
        super().__init__(config, **kwargs)
        # TODO normalize embeddings?

    def _get_embeddings(
        self,
        output,
        seq_mask: Tensor = None,  # Should exclude special tokens, not just padding
        seq_start: int = None,
        seq_end: int = None,
    ):
        # If packed nothing to do (appart from concatenating?)
        # If unpacked, remove the padding and pack together the different sequences.
        if self.use_packed_sequence:
            # Just to remove special tokens. Which would not be padding in the case of a packed implementation
            embeddings = output["hidden_states"][0][seq_start:seq_end][seq_mask]
        else:
            embeddings = output["hidden_states"][seq_start]
            if seq_mask is not None:
                embeddings = embeddings[seq_mask]
        if self.normalize_embeddings:
            embeddings = torch.nn.functional.normalize(embeddings, p=2, dim=-1)

        return embeddings  # Shape: (seq_len, hidden_size) (Exclude any padding and special tokens)

    def forward(
        self, batch: Dict[str, Tensor], **kwargs: Any  # Shape: (batch_size, seq_len) (With padding)
    ) -> Tensor:  # Shape: (batch_size, seq_len, hidden_size) (With padding)
        pass
```
