## Supplementary material for "pLM representations unlock metagenomic space beyond homology": Code bundle: ESM.py

```
from .Encoder import (
    SequenceEncoder,
    TokenEncoder,
    EncoderConfig,
)
import numpy as np
from dataclasses import dataclass, field
from typing import Any, Dict, Optional, Union, List

from transformers import AutoModel, AutoTokenizer, AutoConfig

import torch
from apperitif.collator import DataCollatorForSeqTasks


@dataclass
class ESMConfig(EncoderConfig):
    # ESM-specific configurations only
    # Note: model_name and model_type are inherited from ModelConfig
    # Note: hf_kwargs, use_hidden_layer, etc. are inherited from EncoderConfig
    local_name: str = None  # Local name if different than model_name
    model_type: str = "esm"
    model_name: str = "facebook/esm2_t12_35M_UR50D"  # Default ESM model
    hf_kwargs: Dict[str, Any] = field(default_factory=dict)  # Hugging
    max_length: Optional[int] = 2048  # Maximum sequence length for padding/truncation
    use_attn_maps: bool = True  # Whether to use attention maps
    num_total_attn_maps: Optional[int] = 120  # Total number of attention maps
    randomize: bool = False
    randomize_mode: str = "distribution"


class ESMSequenceEncoder(SequenceEncoder):
    def __init__(self, config: ESMConfig):
        super(ESMSequenceEncoder, self).__init__(config)
        self.model = AutoModel.from_pretrained(config.model_name, **config.hf_kwargs)
        self.tokenizer = AutoTokenizer.from_pretrained(config.model_name, **config.hf_kwargs)
        self.use_hidden_layer = getattr(config, "use_hidden_layer", 1)
        self.name_or_path = config.model_name

        # Correct the hidden size initialized at 1 in the super class
        self.hidden_size = self.model.config.hidden_size * self.use_hidden_layer

        if config.randomize:
            # Create a copy of the architecture
            model_config = AutoConfig.from_pretrained(config.model_name, **config.hf_kwargs)
            randomized_model = AutoModel.from_config(model_config)

            with torch.no_grad():
                if config.randomize_mode == "distribution":
                    for (name_pretrained, param_pretrained), (name_random, param_random) in zip(
                        self.model.named_parameters(), randomized_model.named_parameters()
                    ):
                        if param_pretrained.shape != param_random.shape:
                            raise ValueError(f"Shape mismatch for {name_pretrained}")

                        # Get stats from pretrained weights
                        tensor = param_pretrained.data
                        mean = tensor.mean()
                        std = tensor.std()

                        # Re-sample from the same distribution
                        param_random.data.copy_(torch.normal(mean=mean, std=std, size=tensor.shape).to(tensor.device))

                else:
                    if hasattr(randomized_model, "_init_weights"):
                        randomized_model.apply(randomized_model._init_weights)
                    elif hasattr(randomized_model, "init_weights"):
                        randomized_model.apply(randomized_model.init_weights)
                    else:
                        raise AttributeError("Model does not have an init_weights or _init_weights method.")

                diffs = []
                for (name_pretrained, p_pretrained), (name_random, p_random) in zip(
                    self.model.named_parameters(), randomized_model.named_parameters()
                ):
                    if p_pretrained.norm() == 0:
                        continue
                    rel_diff = (p_random - p_pretrained).norm() / p_pretrained.norm()
                    diffs.append(rel_diff.item())
                avg_diff = sum(diffs) / len(diffs)
                print(f"Average relative norm difference: {avg_diff:.4f}")

                self.model = randomized_model

    def forward(
        self,
        batch,
        reduction_method: str = "mean",
    ):
        input_ids, attention_mask = (
            batch["input_ids"],
            batch.get("attention_mask", None),
        )
        output = self.model(
            input_ids,
            attention_mask=attention_mask,
            output_hidden_states=True,
        )
        output["hidden_states"] = torch.concatenate(output["hidden_states"][-self.use_hidden_layer :], dim=-1)
        mean_embeddings = []
        for i in range(len(input_ids)):
            seq_mask = batch["attention_mask"][i] == 1
            mean_embeddings.append(self._get_embeddings(output, seq_mask, seq_start=i, seq_end=None, reduction_method=reduction_method))

        return torch.stack(mean_embeddings)

    def get_collate_fn(
        self,
        sequence_column: str = "seq",
        label_column: str = "label",
        label_dtype: torch.dtype = torch.long,
        device=None,
        **kwargs,
    ):
        return DataCollatorForSeqTasks(
            tokenizer=self.tokenizer,
            pack_sequences=self.use_packed_sequence,
            device=device,
            sequence_column=sequence_column,
            max_length=self.max_length,
            label_column=label_column,
            label_dtype=label_dtype,
            token_level_task=False,
            **kwargs,
        )


class ESMTokenEncoder(TokenEncoder):
    def __init__(self, config: ESMConfig):
        super(ESMTokenEncoder, self).__init__(config)
        self.model = AutoModel.from_pretrained(config.model_name, **config.hf_kwargs)
        self.tokenizer = AutoTokenizer.from_pretrained(config.model_name, **config.hf_kwargs)
        self.use_hidden_layer = getattr(config, "use_hidden_layer", 1)
        self.name_or_path = config.model_name
        # Correct the hidden size initialized at 1 in the super class
        self.hidden_size = self.model.config.hidden_size * self.use_hidden_layer

        if config.randomize:
            # Create a copy of the architecture
            model_config = AutoConfig.from_pretrained(config.model_name, **config.hf_kwargs)
            randomized_model = AutoModel.from_config(model_config)

            with torch.no_grad():
                if config.randomize_mode == "distribution":
                    for (name_pretrained, param_pretrained), (name_random, param_random) in zip(
                        self.model.named_parameters(), randomized_model.named_parameters()
                    ):
                        if param_pretrained.shape != param_random.shape:
                            raise ValueError(f"Shape mismatch for {name_pretrained}")

                        # Get stats from pretrained weights
                        tensor = param_pretrained.data
                        mean = tensor.mean()
                        std = tensor.std()

                        # Re-sample from the same distribution
                        param_random.data.copy_(torch.normal(mean=mean, std=std, size=tensor.shape).to(tensor.device))

                else:
                    if hasattr(randomized_model, "_init_weights"):
                        randomized_model.apply(randomized_model._init_weights)
                    elif hasattr(randomized_model, "init_weights"):
                        randomized_model.apply(randomized_model.init_weights)
                    else:
                        raise AttributeError("Model does not have an init_weights or _init_weights method.")

                diffs = []
                for (name_pretrained, p_pretrained), (name_random, p_random) in zip(
                    self.model.named_parameters(), randomized_model.named_parameters()
                ):
                    if p_pretrained.norm() == 0:
                        continue
                    rel_diff = (p_random - p_pretrained).norm() / p_pretrained.norm()
                    diffs.append(rel_diff.item())
                avg_diff = sum(diffs) / len(diffs)
                print(f"Average relative norm difference: {avg_diff:.4f}")

                self.model = randomized_model

    def forward(
        self,
        batch,
        reduction_method: str = None,  # Not used in this encoder, but kept for compatibility
        return_individual_sequences: bool = False,
    ) -> Union[torch.Tensor, List[torch.Tensor]]:
        input_ids, attention_mask = (
            batch["input_ids"],
            batch.get("attention_mask", None),
        )
        output = self.model(
            input_ids,
            attention_mask=attention_mask,
            output_hidden_states=True,
        )
        output["hidden_states"] = torch.concatenate(output["hidden_states"][-self.use_hidden_layer :], dim=-1)

        embeddings = []
        for i in range(len(input_ids)):
            seq_mask = batch["attention_mask"][i] == 1
            embeddings.append(self._get_embeddings(output, seq_mask, seq_start=i, seq_end=None))

        if return_individual_sequences:
            return embeddings  # List[(seq_len_i, hidden_size)]
        else:
            out = torch.concatenate(embeddings, dim=0)
            return out

    def get_collate_fn(
        self,
        sequence_column: str = "seq",
        label_column: str = "label",
        label_dtype: torch.dtype = torch.long,
        device=None,
        **kwargs,
    ):
        return DataCollatorForSeqTasks(
            tokenizer=self.tokenizer,
            pack_sequences=self.use_packed_sequence,
            device=device,
            sequence_column=sequence_column,
            max_length=self.max_length,
            label_column=label_column,
            label_dtype=label_dtype,
            token_level_task=True,
            **kwargs,
        )
```
