## Supplementary material for "pLM representations unlock metagenomic space beyond homology": Code bundle: regressor.py

```
from typing import Any, Dict
import torch
from torch import Tensor

from apperitif.models.Encoder import SequenceEncoder, TokenEncoder


class MLP(torch.nn.Module):
    def __init__(self, input_size, hidden_size, output_size=1, activation="gelu"):
        super(MLP, self).__init__()
        self.fc1 = torch.nn.Linear(input_size, hidden_size)
        self.fc2 = torch.nn.Linear(hidden_size, output_size)
        if activation == "relu":
            self.act = torch.nn.ReLU()
        elif activation == "gelu":
            self.act = torch.nn.GELU()
        else:
            raise ValueError(f"Unsupported activation function: {activation}")

    def forward(self, x):  # TODO optimization
        x = self.fc1(x)
        x = self.act(x)
        x = self.fc2(x)
        return x


class Regressor(torch.nn.Module):
    def __init__(
        self,
        trunk: SequenceEncoder | TokenEncoder,  # Potentially Encoder instead of SequenceEncoder (TokenEncoder would be covered too)
        use_low_rank: bool = False,
        reduction_method: str = "mean",
        hidden_size: int = 512,
        output_size: int = 1,
        head_precision: torch.dtype = torch.float32,
        trunk_autocast_precision: torch.dtype = torch.float16,
    ):
        super(Regressor, self).__init__()
        self.trunk = trunk
        self.head_precision = head_precision
        self.head = MLP(trunk.hidden_size, hidden_size, output_size).to(self.head_precision)
        self.trunk_autocast_precision = trunk_autocast_precision

        self.use_low_rank = use_low_rank
        if self.use_low_rank:
            raise NotImplementedError("Low-rank regression is not implemented yet.")
        self.reduction_method = reduction_method
        assert self.reduction_method in [
            "mean",
            "sum",
            "cls",
        ], f"{self.reduction_method} is an invalid reduction method. Must be one of ['mean', 'sum', 'cls']"

    def forward(
        self,
        batch: Dict[str, Tensor],
        frozen_backbone=False,
        device_type: str = "cuda",
        **kwargs: Any,
    ) -> Tensor:
        """
        Forward pass of the regressor.

        Args:
            batch: Dictionary containing the input data.
            **kwargs: Additional arguments for the trunk model.

        Returns:
            Tensor: Output of the regressor.
        """
        if frozen_backbone:
            self.trunk.eval()  # To have similar behavior as the precomputed case
            with (
                torch.no_grad(),
                torch.autocast(
                    device_type=device_type,
                    dtype=self.trunk_autocast_precision,
                ),
            ):
                embeddings = self.trunk(batch, reduction_method=self.reduction_method, **kwargs)
        else:
            self.trunk.train()
            with (
                torch.autocast(
                    device_type=device_type,
                    dtype=self.trunk_autocast_precision,
                ),
            ):
                embeddings = self.trunk(batch, reduction_method=self.reduction_method, **kwargs)

        embeddings = embeddings.to(self.head_precision)
        return self.head(embeddings)
```
