## Supplementary material for "pLM representations unlock metagenomic space beyond homology": Code bundle: AbsTask.py

```
import os
import ast
from abc import ABC, abstractmethod
import json
import logging
from pathlib import Path
from datetime import datetime
from apperitif.models import Encoder
from apperitif.models.Encoder import EncoderConfig
from apperitif.evaluation.Evaluator import Evaluator, EvaluatorConfig
from apperitif.evaluation.SequenceClassificationEvaluator import SequenceClassificationEvaluatorConfig
from apperitif.evaluation.SequenceRegressionEvaluator import SequenceRegressionEvaluatorConfig
from apperitif.utils.evaluator_utils import EvaluatorResult
from dataclasses import dataclass, field
from typing import Any, Dict, Optional, Union


@dataclass
class DataConfig:
    name: str = "default_data"


@dataclass
class TaskConfig:
    # Data config: passed to get_data (can be empty for now)
    data_config: Union[DataConfig, Dict[str, Any]] = field(default_factory=dict)
    # Evaluator config: now uses proper config classes instead of dict
    evaluator_config: Union[
        EvaluatorConfig,
        Dict[str, Any],
    ] = field(default_factory=dict)
    # Model config: now uses proper config classes instead of dict
    model_config: Union[EncoderConfig, Dict[str, Any]] = field(default_factory=dict)
    # Optionally, add a name or description for the task
    name: Optional[str] = None
    description: Optional[str] = None
    device: str = "cuda"  # Default device, can be overridden in subclasses

    # Output folder for results and artifacts
    output_folder: str = "apperitif"

    # Options for detailed results
    return_all_results: bool = False  # Whether to return all hyperparameter grid search results
    do_test: bool = True  # Whether to run the test set evaluation.
    verbose: Union[bool, str] = "reduced"  # Verbosity level for training progress, can be True/False or "reduced"
    accumulation_steps: int = None  # Gradient accumulation steps for training, default is 1 (no accumulation)


class AbsTask(ABC):
    """
    Abstract class for tasks.

    Prepares datasets and determines evaluation metrics, methods and hyperparameters.
    """

    def __init__(self, config: TaskConfig):
        super().__init__()
        self.logger = logging.getLogger(__name__)
        logging.basicConfig(filename=None, level=logging.INFO)
        self.config = config
        self._name = self.name
        self.device = config.device

    @property
    @abstractmethod
    def name(cls):
        """Class name that must be defined in subclasses"""
        pass

    @abstractmethod
    def evaluate(self, **kwargs) -> EvaluatorResult:
        """Evaluate the model for a given task.

        Return standardized EvaluatorResult object with metrics, hyperparameters, and metadata.
        The return_all_results option in config determines whether detailed grid search results are included.

        Args:
            kwargs: Additional arguments for evaluation.

        Returns:
            EvaluatorResult: Standardized result object with metrics and hyperparameter information
        """
        pass

    def aggregate_results_in_json(self, filename: str = "results.json") -> str:
        """Aggregate evaluation results to a JSON file in the output folder.

        Args:
            filename: Name of the JSON file to save (default: "results.json")

        Returns:
            str: Path to the saved JSON file

        Raises:
            ValueError: If output_folder is not configured
        """
        if not self.config.output_folder:
            raise ValueError("output_folder must be configured in TaskConfig to save results")

        # Create output directory if it doesn't exist
        output_path = Path(self.config.output_folder)
        output_path.mkdir(parents=True, exist_ok=True)

        # Ensure filename has .json extension
        if not filename.endswith(".json"):
            filename += ".json"

        file_path = output_path / filename

        if not os.path.exists(file_path):
            raise ValueError(f"File {file_path} does not exists. Cannot aggregate all results.")

        # If the file already exists, load existing results and merge
        with open(file_path, "rt") as f:
            results_dict = json.load(f)

        # Add task-specific metadata
        results_dict.update(
            {
                "task_name": self.config.name or self.name,
                "task_description": self.config.description,
                "timestamp": datetime.now().isoformat(),
                "device": self.config.device,
            }
        )

        best_hyperparameters = max(results_dict["all_results"], key=lambda x: results_dict["all_results"][x].get("validation_score", 0))
        best_results = results_dict["all_results"][best_hyperparameters]

        results_dict.update(
            {
                "best_hyperparameters": ast.literal_eval(best_hyperparameters),
                "validation_score": best_results["validation_score"],
                "metrics": best_results["test_scores"],
                "std_metrics": best_results["test_std"],
            }
        )

        # Save to JSON file
        results_path = output_path / "results.json"
        with open(results_path, "w", encoding="utf-8") as f:
            json.dump(results_dict, f, indent=2, ensure_ascii=False)

        self.logger.info(f"Results saved to: {results_path}")


class Task(AbsTask):
    @classmethod
    def from_config(cls, config: TaskConfig):
        """
        Factory method to create a Task instance from a TaskConfig object.
        Dynamically resolves the correct task class based on the data_config type.

        Args:
            config: TaskConfig object containing all necessary configurations

        Returns:
            Task instance with the provided configuration
        """
        # Import here to avoid circular imports
        from apperitif.tasks.sequence_classification.SequenceClassificationTask import (
            SequenceClassificationTask,
            SequenceClassificationDataConfig,
        )
        from apperitif.tasks.sequence_regression.SequenceRegressionTask import (
            SequenceRegressionTask,
            RegressionDataConfig,
        )
        from apperitif.tasks.token_classification.TokenClassificationTask import (
            TokenClassificationTask,
            TokenClassificationDataConfig,
        )

        data_config = config.data_config

        # Map data config classes to task classes
        config_to_task_mapping = {
            # Base task classes
            SequenceClassificationDataConfig: SequenceClassificationTask,
            RegressionDataConfig: SequenceRegressionTask,
            TokenClassificationDataConfig: TokenClassificationTask,
        }

        # Find the appropriate task class based on data_config type
        data_config_type = type(data_config)

        # Check direct mapping first
        if data_config_type in config_to_task_mapping:
            task_class = config_to_task_mapping[data_config_type]
            return task_class.from_config(config)  # Will further resolve to specific task class

        # If no direct mapping, check inheritance
        for config_class, task_class in config_to_task_mapping.items():
            if isinstance(data_config, config_class):
                return task_class.from_config(config)  # Will further resolve to specific task class

        # If no match found, raise an error
        raise ValueError(
            f"No task class found for data config type: {data_config_type}. " f"Supported types: {list(config_to_task_mapping.keys())}"
        )


# def create_task_from_hydra_config(
#     model_config,
#     evaluator_config,
#     data_config,
#     task_name: Optional[str] = None,
#     device: str = "cuda",
#     return_all_results: bool = False,
#     output_folder: Optional[str] = None,
# ):
#     """
#     Factory function to create a task from Hydra configs.

#     This function resolves anonymous configs to proper config classes
#     and then creates the appropriate task instance.

#     Args:
#         model_config: Anonymous model config (dict or DictConfig)
#         evaluator_config: Anonymous evaluator config (dict or DictConfig)
#         data_config: Anonymous data config (dict or DictConfig)
#         task_name: Optional task name
#         device: Device to use for computation
#         return_all_results: Whether to return all hyperparameter grid search results
#         output_folder: Optional output folder for results and artifacts
#     Returns:
#         Properly configured task instance
#     """
#     from apperitif.utils.config_utils import resolve_all_configs

#     # Resolve all configs to proper config classes
#     resolved_model_config, resolved_evaluator_config, resolved_data_config = resolve_all_configs(
#         model_config, evaluator_config, data_config
#     )

#     # Create TaskConfig with resolved configs
#     task_config = TaskConfig(
#         model_config=resolved_model_config,
#         evaluator_config=resolved_evaluator_config,
#         data_config=resolved_data_config,
#         name=task_name,
#         device=device,
#         return_all_results=return_all_results,
#         output_folder=output_folder,
#     )

#     # Map data config types to task classes
#     from apperitif.tasks.sequence_classification.SequenceClassificationTask import SequenceClassificationTask
#     from apperitif.tasks.sequence_regression.SequenceRegressionTask import SequenceRegressionTask
#     from apperitif.tasks.token_classification.TokenClassificationTask import TokenClassificationTask
#     from apperitif.tasks.sequence_classification.biomap_research import BiomapResearchSequenceClassificationTask
#     from apperitif.tasks.sequence_regression.biomap_research import BiomapResearchSequenceRegressionTask
#     from apperitif.tasks.token_classification.biomap_research import BiomapResearchTokenClassificationTask

#     task_type = resolved_data_config.task_type

#     TASK_CLASS_MAPPING = {
#         "sequence_classification": SequenceClassificationTask,
#         "sequence_regression": SequenceRegressionTask,
#         "token_classification": TokenClassificationTask,
#         "biomap_research_sequence_classification": BiomapResearchSequenceClassificationTask,
#         "biomap_research_sequence_regression": BiomapResearchSequenceRegressionTask,
#         "biomap_research_token_classification": BiomapResearchTokenClassificationTask,
#     }

#     if task_type not in TASK_CLASS_MAPPING:
#         raise ValueError(f"Unknown task_type: {task_type}. Available types: {list(TASK_CLASS_MAPPING.keys())}")

#     task_class = TASK_CLASS_MAPPING[task_type]
#     return task_class(task_config)
```
