## Supplementary material for "pLM representations unlock metagenomic space beyond homology": Code bundle: __init__.py

```
from .biomap_research import BiomapResearchSequenceClassificationConfig, BiomapResearchSequenceClassificationTask

__all__ = [
    "BiomapResearchSequenceClassificationConfig",
    "BiomapResearchSequenceClassificationTask",
]
```
