## Supplementary material for "pLM representations unlock metagenomic space beyond homology": Code bundle: SequenceClassificationTask.py

```
from apperitif.tasks.AbsTask import AbsTask, TaskConfig, DataConfig
from apperitif.evaluation.SequenceClassificationEvaluator import (
    MLPSequenceClassificationEvaluator,
    MLPSequenceClassificationEvaluatorConfig,
)
from apperitif.models.Amplify import AmplifySequenceEncoder, AmplifyConfig
from apperitif.models.ESM import ESMSequenceEncoder, ESMConfig
from apperitif.models.Encoder import EncoderConfig
from apperitif.utils.evaluator_utils import EvaluatorResult
from transformers import AutoTokenizer
from typing import Dict, Any, Tuple, Union
from dataclasses import dataclass
from apperitif.utils.dtype_utils import get_torch_dtype


@dataclass
class SequenceClassificationDataConfig(DataConfig):
    """
    Configuration for sequence classification datasets.
    This can include paths, preprocessing options, etc.
    """

    task_type: str = "sequence_classification"  # Discriminator field for Hydra
    sequence_column: str = "seq"  # Column name for sequences
    label_column: str = "label"  # Column name for labels
    label_dtype: str = "int"  # Data type for labels (e.g., "int", "float")
    num_labels: int = 2  # Number of labels for classification tasks


class SequenceClassificationTask(AbsTask):
    """
    Abstract task for sequence classification using structured config objects.
    """

    @classmethod
    def from_config(cls, config: TaskConfig):
        """
        Factory method to create a SequenceClassificationTask instance from a TaskConfig object.
        Dynamically resolves the correct task class based on the data_config type.

        Args:
            config: TaskConfig object containing all necessary configurations

        Returns:
            SequenceClassificationTask instance with the provided configuration
        """
        # Import biomap research specific classes
        from apperitif.tasks.sequence_classification.biomap_research import (
            BiomapResearchSequenceClassificationTask,
            BiomapResearchSequenceClassificationConfig,
        )

        data_config = config.data_config

        # Map data config classes to task classes for sequence classification
        config_to_task_mapping = {
            SequenceClassificationDataConfig: cls,  # Base class
            BiomapResearchSequenceClassificationConfig: BiomapResearchSequenceClassificationTask,
        }

        # Find the appropriate task class based on data_config type
        data_config_type = type(data_config)

        # Check direct mapping first
        if data_config_type in config_to_task_mapping:
            task_class = config_to_task_mapping[data_config_type]
            return task_class(config)

        # If no direct mapping, check inheritance
        for config_class, task_class in config_to_task_mapping.items():
            if isinstance(data_config, config_class):
                return task_class(config)

        # If no match found, raise an error
        raise ValueError(
            f"No sequence classification task class found for data config type: {data_config_type}. "
            f"Supported types: {list(config_to_task_mapping.keys())}"
        )

    @property
    def name(self):
        return "sequence_classification"

    def get_data(self):
        """To be implemented in child classes: returns datasets."""
        raise NotImplementedError

    def get_encoder(self) -> Tuple[Any, Any]:
        """
        Get encoder and collate function from the task's model config.

        Returns:
            Tuple of (encoder, collate_fn)
        """
        model_config = self.config.model_config
        data_config = self.config.data_config

        # Ensure we have a structured config object
        if not isinstance(model_config, (ESMConfig, AmplifyConfig, EncoderConfig)):
            raise ValueError(f"Expected structured config object, got {type(model_config)}")

        # Create encoder based on config type
        if isinstance(model_config, ESMConfig):
            encoder = ESMSequenceEncoder(config=model_config)
        elif isinstance(model_config, AmplifyConfig):
            encoder = AmplifySequenceEncoder(config=model_config)
        else:
            # Generic SequenceEncoderConfig - need to determine type from model_type
            if model_config.model_type.lower() == "esm":
                encoder = ESMSequenceEncoder(config=model_config)
            elif model_config.model_type.lower() == "amplify":
                encoder = AmplifySequenceEncoder(config=model_config)
            else:
                raise ValueError(f"Unknown model_type: {model_config.model_type}")

        # Create collate function with reasonable defaults
        collate_fn = encoder.get_collate_fn(
            sequence_column=getattr(data_config, "sequence_column", "seq"),
            label_column=getattr(data_config, "label_column", "label"),
            label_dtype=get_torch_dtype(getattr(data_config, "label_dtype", int)),
            device=self.device,
        )

        return encoder, collate_fn

    def get_evaluator(self, datasets: Dict[str, Any], collate_fn: Any) -> Any:
        """
        Get evaluator from the task's evaluator config.

        Args:
            datasets: Dictionary of datasets (train, validation, test)
            collate_fn: Collate function for the datasets

        Returns:
            Configured evaluator instance
        """
        evaluator_config = self.config.evaluator_config
        data_config = self.config.data_config

        evaluator_config.num_labels = getattr(data_config, "num_labels", 2)  # Default to binary classification

        # Use head_type to determine the evaluator class
        head_type = getattr(evaluator_config, "head_type", None)
        if head_type is None:
            raise ValueError(f"evaluator_config must have a 'head_type' attribute, got {type(evaluator_config)}")

        # Create the appropriate config class based on head_type
        if head_type == "mlp":
            # Convert to MLPSequenceClassificationEvaluatorConfig if needed
            if not isinstance(evaluator_config, MLPSequenceClassificationEvaluatorConfig):
                evaluator_config = MLPSequenceClassificationEvaluatorConfig(**evaluator_config)
            return MLPSequenceClassificationEvaluator(config=evaluator_config, datasets=datasets, collate_fn=collate_fn)
        else:
            raise ValueError(f"Unsupported head_type: {head_type}. Supported types: mlp")

    def evaluate(self, **kwargs) -> EvaluatorResult:
        """
        Evaluate the model for this task.

        Args:
            **kwargs: Additional arguments passed to the evaluator

        Returns:
            EvaluatorResult: Standardized evaluation results with hyperparameter information
        """
        # Get datasets
        datasets = self.get_data()

        # Get encoder and collate function
        encoder, collate_fn = self.get_encoder()

        encoder = encoder.to(self.device)

        # Get evaluator
        evaluator = self.get_evaluator(datasets, collate_fn)

        # Merge config options with kwargs
        eval_kwargs = {
            "return_all_scores": self.config.return_all_results,
            "do_test": self.config.do_test,
            "verbose": self.config.verbose,
            "accumulation_steps": self.config.accumulation_steps,
            "task_name": self.config.name,
            "task_description": self.config.description,
            "output_folder": self.config.output_folder,
            **kwargs,
        }
        
        # Run evaluation
        evaluator(encoder, **eval_kwargs)

        # Save results to JSON if output folder is configured
        if self.config.output_folder:
            self.aggregate_results_in_json()
```
