## Supplementary material for "pLM representations unlock metagenomic space beyond homology": Code bundle: __init__.py

```
from .biomap_research import BiomapResearchSequenceRegressionConfig, BiomapResearchSequenceRegressionTask

__all__ = [
    "BiomapResearchSequenceRegressionConfig",
    "BiomapResearchSequenceRegressionTask",
]
```
