## Supplementary material for "pLM representations unlock metagenomic space beyond homology": Code bundle: biomap_research.py

```
from apperitif.tasks.sequence_regression.SequenceRegressionTask import (
    SequenceRegressionTask,
    RegressionDataConfig,
)
from dataclasses import dataclass
from datasets import load_dataset
from typing import Optional


@dataclass
class BiomapResearchSequenceRegressionConfig(RegressionDataConfig):
    """
    Configuration for the Biomap Research sequence regression task.
    This can include paths, preprocessing options, etc.
    """

    task_type: str = "biomap_research_sequence_regression"  # Discriminator field for Hydra
    dataset_name: str = "biomap-research"
    split_train: bool = False
    split_seed: Optional[int] = 42


class BiomapResearchSequenceRegressionTask(SequenceRegressionTask):
    """
    Task for sequence regression using the Biomap Research dataset.
    Inherits from SequenceRegressionTask and uses structured config objects.
    """

    @property
    def name(self):
        return "biomap_research_sequence_regression"

    def get_data(self):
        """To be implemented in child classes: returns datasets."""
        dataset = load_dataset(self.config.data_config.dataset_name)
        # If the label column is not of type float, convert it
        if self.config.data_config.label_column not in dataset["train"].features:
            raise ValueError(f"Label column '{self.config.data_config.label_column}' not found in dataset.")
        if dataset["train"].features[self.config.data_config.label_column].dtype != "float":
            dataset = dataset.map(
                lambda x: {self.config.data_config.label_column: float(x[self.config.data_config.label_column])},
                remove_columns=[self.config.data_config.label_column],
            )
        if self.config.data_config.split_train:
            # Generate the valid split from the train split
            train_dataset = dataset["train"].train_test_split(test_size=0.1, seed=self.config.data_config.split_seed)
            dataset["valid"] = train_dataset["test"]
            dataset["train"] = train_dataset["train"]
        dataset["validation"] = dataset.pop("valid")  # Will fail if no valid split was created
        return dataset
```
