## Supplementary material for "pLM representations unlock metagenomic space beyond homology": Code bundle: __init__.py

```
from .biomap_research import BiomapResearchTokenClassificationConfig, BiomapResearchTokenClassificationTask

__all__ = [
    "BiomapResearchTokenClassificationConfig",
    "BiomapResearchTokenClassificationTask",
]
```
