## Supplementary material for "pLM representations unlock metagenomic space beyond homology": Code bundle: biomap_research.py

```
from apperitif.tasks.token_classification.TokenClassificationTask import (
    TokenClassificationTask,
    TokenClassificationDataConfig,
)
from dataclasses import dataclass
from datasets import load_dataset
from typing import Optional


@dataclass
class BiomapResearchTokenClassificationConfig(TokenClassificationDataConfig):
    """
    Configuration for the Biomap Research token classification task.
    This can include paths, preprocessing options, etc.
    """

    task_type: str = "biomap_research_token_classification"  # Discriminator field for Hydra
    dataset_name: str = "biomap-research"
    split_train: bool = False
    split_seed: Optional[int] = 42
    label_column = "label"  # Default label column, can be overridden in specific tasks
    sequence_column = "seq"


class BiomapResearchTokenClassificationTask(TokenClassificationTask):
    """
    Task for token classification using the Biomap Research dataset.
    Inherits from TokenClassificationTask and uses structured config objects.
    """

    @property
    def name(self):
        return self.config.data_config.dataset_name

    def get_data(self):
        """To be implemented in child classes: returns datasets."""
        dataset = load_dataset(self.config.data_config.dataset_name)
        if self.config.data_config.split_train:
            # Generate the valid split from the train split
            train_dataset = dataset["train"].train_test_split(test_size=0.1, seed=self.config.data_config.split_seed)
            dataset["valid"] = train_dataset["test"]
            dataset["train"] = train_dataset["train"]
        dataset["validation"] = dataset.pop("valid")  # Will fail if no valid split was created
        return dataset
```
