## Supplementary material for "pLM representations unlock metagenomic space beyond homology": Code bundle: config_utils.py

```
"""
Utility functions for handling Hydra config mapping.
These functions help map anonymous config objects to specific config classes
based on discriminator fields like model_type, evaluator_type, and task_type.
"""

from typing import Any, Dict, Union
from omegaconf import DictConfig


def resolve_evaluator_target(
    task_type: str = "sequence_classification",
    head_type: str = "mlp",
) -> str:
    """
    Resolve the _target_ field for the evaluator based on task and head type.
    Args:
        task_type: The type of task (e.g., "sequence_classification", "token_classification", etc.)
        head_type: The type of head to use (currently only "mlp").
    Returns:
        The fully qualified class path for the evaluator.
    """
    # Mapping of task types to evaluator classes
    task_evaluator_mapping = {
        "sequence_classification": {
            "mlp": "apperitif.evaluation.SequenceClassificationEvaluator.MLPSequenceClassificationEvaluatorConfig",
        },
        "sequence_regression": {
            "mlp": "apperitif.evaluation.SequenceRegressionEvaluator.MLPSequenceRegressionEvaluatorConfig",
        },
        "token_classification": {
            "mlp": "apperitif.evaluation.TokenClassificationEvaluator.MLPTokenClassificationEvaluatorConfig",
        },
    }

    # Resolve the evaluator class based on task and head type
    if task_type not in task_evaluator_mapping:
        raise ValueError(f"Unknown task_type: {task_type}. Available types: {list(task_evaluator_mapping.keys())}")
    if head_type not in task_evaluator_mapping[task_type]:
        raise ValueError(
            f"Unknown head_type '{head_type}' for task '{task_type}'. "
            f"Available: {list(task_evaluator_mapping[task_type].keys())}"
        )
    evaluator_class_name = task_evaluator_mapping[task_type][head_type]
    return evaluator_class_name
```
