## Supplementary material for "pLM representations unlock metagenomic space beyond homology": Code bundle: deterministic.py

```
import torch
import numpy as np
import random


def set_seed(seed: int):
    """
    Set the seed for all random number generators to ensure reproducibility.
    Args:
        seed (int): The seed value to set.
    """
    torch.manual_seed(seed)
    torch.cuda.manual_seed_all(seed)
    np.random.seed(seed)
    random.seed(seed)
    # On or Off this does not seem to affect reproducibility
    torch.backends.cudnn.deterministic = True
    torch.backends.cudnn.benchmark = False
```
