## Supplementary material for "pLM representations unlock metagenomic space beyond homology": Code bundle: dtype_utils.py

```
"""
Utility functions for handling data types, especially for Hydra compatibility.
"""

import torch


def get_torch_dtype(precision_str: str) -> torch.dtype:
    """
    Convert precision string to torch dtype for Hydra compatibility.

    Args:
        precision_str: String representation of the precision.
                      Supported values: "fp16", "float16", "bf16", "bfloat16",
                      "fp32", "float32", "fp64", "float64"

    Returns:
        Corresponding torch.dtype

    Raises:
        ValueError: If precision_str is not supported
    """
    precision_map = {
        "fp16": torch.float16,
        "float16": torch.float16,
        "bf16": torch.bfloat16,
        "bfloat16": torch.bfloat16,
        "fp32": torch.float32,
        "float32": torch.float32,
        "fp64": torch.float64,
        "float64": torch.float64,
        "float": torch.float32,  # Alias for float32
        "double": torch.float64,  # Alias for float64
        "int": torch.int32,  # Alias for int32
        "int32": torch.int32,
        "int64": torch.int64,  # Alias for int64
        "long": torch.int64,  # Alias for int64
        "bool": torch.bool,  # Alias for boolean
    }

    if precision_str not in precision_map:
        raise ValueError(f"Unsupported precision: {precision_str}. Supported: {list(precision_map.keys())}")

    return precision_map[precision_str]
```
