## Supplementary material for "pLM representations unlock metagenomic space beyond homology": Code bundle: embedding_utils.py

```
from typing import Dict, Tuple, Any, List
import tqdm

import numpy as np
import torch

from torch.utils.data import DataLoader

from apperitif.models.Encoder import SequenceEncoder, TokenEncoder

# TODO add a token embeddings equivalent


def get_embeddings_and_labels(
    model: SequenceEncoder,
    dataloader: DataLoader,
    target_column: str,
    reduction_method: str,
    model_kwargs: Dict[str, Any],
    autocast_precision: torch.dtype = torch.float16,
) -> Tuple[np.ndarray, np.ndarray]:
    """
    Get sequence embeddings and labels for the given dataloader using the specified model.
    This function computes the embeddings for each token in each sequence in the dataloader, collapses them using the specified reduction method, and returns the embeddings along with their corresponding labels.

    Args:
        model: Encoder model to use.
        dataloader: Dataloader to use.
        reduction_method: Method to reduce the hidden states to a single vector. Can be "mean", "sum", or "cls".
        target_column: The column in the batch that contains the labels.
        model_kwargs: Additional keyword arguments to pass to the model.

    Returns:
        Tuple containing the embeddings and labels.
    """
    model.eval()
    embeddings, labels = [], []
    device = next(model.parameters()).device
    with (
        torch.no_grad(),
        torch.autocast(device_type=device.type, dtype=autocast_precision),
    ):  # Use autocast for mixed precision
        for batch in tqdm.tqdm(dataloader, desc="Computing embeddings"):
            labels.append(batch.pop(target_column).cpu().numpy().reshape(-1))
            embeddings.append(model(batch, reduction_method=reduction_method, **model_kwargs).cpu().numpy())
    embeddings = np.concatenate(embeddings, axis=0)
    labels = np.concatenate(labels, axis=0)
    return embeddings, labels


def get_token_embeddings_and_labels(
    model: TokenEncoder,
    dataloader: DataLoader,
    labels_key: str = "labels",
    device: str = "cuda",
    model_kwargs: Dict[str, Any] = None,
    autocast_precision: torch.dtype = torch.float16,
) -> List[Dict[str, torch.Tensor]]:
    """
    Get token-level embeddings and labels for the given dataloader using the specified model.

    The TokenEncoder models handle special tokens and attention masking internally, so this function
    simply extracts embeddings using return_individual_sequences=True and matches them with labels
    that have -100 values removed (standard practice for token classification).

    Args:
        model: TokenEncoder model to use.
        dataloader: Dataloader to use.
        labels_key: The key in the batch that contains the token-level labels.
        device: Device to run inference on.
        model_kwargs: Additional keyword arguments to pass to the model.
        autocast_precision: Precision for autocast.

    Returns:
        List of dictionaries, each containing:
        - 'embeddings': tensor of shape (seq_len_i, hidden_size)
        - 'labels': tensor of shape (seq_len_i,) with -100 values removed
    """
    if model_kwargs is None:
        model_kwargs = {}

    model.eval()
    results = []
    device_obj = torch.device(device)

    with torch.no_grad(), torch.autocast(device_type=device_obj.type, dtype=autocast_precision):
        for batch in tqdm.tqdm(dataloader, desc="Computing token embeddings"):
            # Extract labels before passing to model (make a copy to avoid modifying original)
            batch_labels = batch.pop(labels_key)
            # Has shape (sum_seq_lens, )

            # Get individual sequence embeddings (already filtered by model)
            individual_embeddings = model(batch, return_individual_sequences=True, **model_kwargs)
            # Process labels for each sequence
            seq_start = 0
            for i, embedding_tensor in enumerate(individual_embeddings):
                # Get labels for this sequence and remove -100 values (padding/special tokens)
                labels = batch_labels[seq_start : seq_start + embedding_tensor.shape[0]]
                seq_start += embedding_tensor.shape[0]
                assert (
                    labels.shape[0] == embedding_tensor.shape[0]
                ), f"Labels shape {labels.shape} does not match embedding shape {embedding_tensor.shape}"
                results.append(
                    {
                        "embeddings": embedding_tensor.cpu(),
                        "labels": labels.cpu(),
                    }
                )

    return results
```
