## Supplementary material for "pLM representations unlock metagenomic space beyond homology": Code bundle: evaluator_utils.py

```
"""
Standardized evaluator output format and key mapping utilities.
"""

from dataclasses import dataclass, field
from typing import Dict, Any, List, Tuple, NamedTuple, Optional, Union
import json
import numpy as np


def convert_to_json_serializable(obj):
    """
    Convert NumPy and PyTorch types to JSON-serializable Python types.
    
    Args:
        obj: The object to convert
        
    Returns:
        JSON-serializable version of the object
    """
    if isinstance(obj, np.ndarray):
        return obj.tolist()
    elif isinstance(obj, (np.integer, np.int32, np.int64)):
        return int(obj)
    elif isinstance(obj, (np.floating, np.float32, np.float64)):
        return float(obj)
    elif isinstance(obj, dict):
        return {key: convert_to_json_serializable(value) for key, value in obj.items()}
    elif isinstance(obj, (list, tuple)):
        return [convert_to_json_serializable(item) for item in obj]
    else:
        # Try to handle PyTorch tensors if they exist
        try:
            if hasattr(obj, 'cpu') and hasattr(obj, 'numpy'):
                return convert_to_json_serializable(obj.cpu().numpy())
        except:
            pass
        return obj


@dataclass
class EvaluatorResult:
    """
    Standardized result format for all evaluators.
    """

    # Main results for each metric
    metrics: Dict[str, float] = field(default_factory=dict)

    # Standard deviation for each metric (if applicable)
    std_metrics: Dict[str, float] = field(default_factory=dict)

    # Best hyperparameters that achieved these results
    best_hyperparameters: Optional[Tuple] = None

    # Key mapping explaining what each position in best_hyperparameters tuple represents
    hyperparameter_keys: Optional[List[str]] = None

    # All results for different hyperparameter combinations (optional, for detailed analysis)
    all_results: Optional[Dict[Tuple, Dict[str, Any]]] = None

    # Validation scores (if available)
    validation_score: Optional[float] = None
    validation_std: Optional[float] = None

    # Additional metadata
    evaluator_name: str = ""
    model_name: str = ""
    dataset_name: str = ""

    def get_hyperparameter_mapping(self) -> Dict[str, Any]:
        """
        Get a human-readable mapping of hyperparameters.

        Returns:
            Dictionary mapping hyperparameter names to their values
        """
        if self.best_hyperparameters is None or self.hyperparameter_keys is None:
            return {}

        if len(self.best_hyperparameters) != len(self.hyperparameter_keys):
            raise ValueError(
                f"Mismatch between hyperparameters ({len(self.best_hyperparameters)}) and keys ({len(self.hyperparameter_keys)})"
            )

        return dict(zip(self.hyperparameter_keys, self.best_hyperparameters))

    def to_dict(self) -> Dict[str, Any]:
        """Convert to dictionary for easy serialization."""
        result = {
            "metrics": convert_to_json_serializable(self.metrics),
            "std_metrics": convert_to_json_serializable(self.std_metrics),
            "evaluator_name": self.evaluator_name,
            "model_name": self.model_name,
            "dataset_name": self.dataset_name,
        }

        if self.best_hyperparameters is not None:
            result["best_hyperparameters"] = convert_to_json_serializable(self.best_hyperparameters)
            result["hyperparameter_keys"] = self.hyperparameter_keys
            result["hyperparameter_mapping"] = convert_to_json_serializable(self.get_hyperparameter_mapping())

        if self.validation_score is not None:
            result["validation_score"] = convert_to_json_serializable(self.validation_score)
            result["validation_std"] = convert_to_json_serializable(self.validation_std)

        if self.all_results is not None:
            # Convert tuple keys to strings for JSON serialization
            result["all_results"] = {str(k): convert_to_json_serializable(v) for k, v in self.all_results.items()}

        return result

    def to_json(self, indent: int = 2) -> str:
        """Convert to JSON string."""
        return json.dumps(self.to_dict(), indent=indent)

    def __str__(self) -> str:
        """Human-readable string representation."""
        lines = [f"EvaluatorResult - {self.evaluator_name}"]

        if self.model_name:
            lines.append(f"Model: {self.model_name}")
        if self.dataset_name:
            lines.append(f"Dataset: {self.dataset_name}")

        lines.append("Metrics:")
        for metric, value in self.metrics.items():
            std_val = self.std_metrics.get(metric, None)
            if std_val is not None:
                lines.append(f"  {metric}: {value:.4f} ± {std_val:.4f}")
            else:
                lines.append(f"  {metric}: {value:.4f}")

        if self.best_hyperparameters is not None:
            lines.append("Best Hyperparameters:")
            mapping = self.get_hyperparameter_mapping()
            for key, value in mapping.items():
                lines.append(f"  {key}: {value}")

        if self.validation_score is not None:
            if self.validation_std is not None:
                lines.append(f"Validation Score: {self.validation_score:.4f} ± {self.validation_std:.4f}")
            else:
                lines.append(f"Validation Score: {self.validation_score:.4f}")

        return "\\n".join(lines)


# Standard hyperparameter key definitions for different evaluator types
HYPERPARAMETER_KEYS = {
    "mlp": ["hidden_size", "learning_rate", "num_epochs", "weight_decay", "batch_size"],  # Generic MLP
}


def get_hyperparameter_keys(evaluator_type: str) -> List[str]:
    """
    Get the standard hyperparameter keys for a given evaluator type.

    Args:
        evaluator_type: The type of evaluator (should match evaluator_type field in config)

    Returns:
        List of hyperparameter key names in order
    """
    return HYPERPARAMETER_KEYS.get(evaluator_type, [])


def create_standardized_result(
    metrics: Dict[str, float],
    evaluator_type: str,
    best_hyperparameters: Optional[Tuple] = None,
    std_metrics: Optional[Dict[str, float]] = None,
    all_results: Optional[Dict[Tuple, Dict[str, Any]]] = None,
    validation_score: Optional[float] = None,
    validation_std: Optional[float] = None,
    model_name: str = "",
    dataset_name: str = "",
) -> EvaluatorResult:
    """
    Create a standardized result object.

    Args:
        metrics: Dictionary of metric names to values
        evaluator_type: Type of evaluator (for determining hyperparameter keys)
        best_hyperparameters: Tuple of best hyperparameter values
        std_metrics: Standard deviations for metrics
        all_results: All results for different hyperparameter combinations
        validation_score: Best validation score
        validation_std: Standard deviation of validation score
        model_name: Name of the model used
        dataset_name: Name of the dataset used

    Returns:
        Standardized EvaluatorResult object
    """
    hyperparameter_keys = get_hyperparameter_keys(evaluator_type)

    return EvaluatorResult(
        metrics=metrics,
        std_metrics=std_metrics or {},
        best_hyperparameters=best_hyperparameters,
        hyperparameter_keys=hyperparameter_keys if best_hyperparameters else None,
        all_results=all_results,
        validation_score=validation_score,
        validation_std=validation_std,
        evaluator_name=evaluator_type,
        model_name=model_name,
        dataset_name=dataset_name,
    )
```
