## Supplementary material for "pLM representations unlock metagenomic space beyond homology": Code bundle: optimization.py

```
import torch
from torch.optim import Optimizer


def get_linear_scheduler_with_warmup(
    optimizer: Optimizer,
    num_warmup_steps: int,
    num_training_steps: int,
):
    """
    Create a linear scheduler with warmup.

    Args:
        optimizer: The optimizer to schedule.
        num_warmup_steps (int): Number of steps for the warmup phase.
        num_training_steps (int): Total number of training steps.
        last_epoch (int, optional): The index of the last epoch. Default is -1.

    Returns:
        torch.optim.lr_scheduler.LinearLR: A linear learning rate scheduler with warmup.
    """
    scheduler_warmup = torch.optim.lr_scheduler.LinearLR(optimizer, start_factor=3e-2, end_factor=1, total_iters=num_warmup_steps)
    scheduler_decay = torch.optim.lr_scheduler.LinearLR(optimizer, start_factor=1, end_factor=0, total_iters=num_training_steps - num_warmup_steps)
    scheduler = torch.optim.lr_scheduler.SequentialLR(optimizer, [scheduler_warmup, scheduler_decay], milestones=[num_warmup_steps])
    return scheduler

class EarlyStopping:
    def __init__(self, patience=5, mode='min', disable=False):
        """
        Args:
            patience (int): Number of validation checks with no improvement before stopping.
            mode (str): 'min' for loss, 'max' for accuracy or other metrics.
        """
        self.patience = patience
        self.mode = mode
        self.best_score = None
        self.counter = 0
        if self.patience <= 0:
            self.disable = True
        else:
            self.disable = False
        self.disable = disable
            

    def __call__(self, current_score):
        if self.disable:
            should_stop = False
            return should_stop

        if self.best_score is None:
            self.best_score = current_score
            should_stop = False
        elif (
            (self.mode == 'min' and current_score < self.best_score) or
            (self.mode == 'max' and current_score > self.best_score)
        ):
            self.best_score = current_score
            self.counter = 0
            should_stop = False
        else:
            self.counter += 1
            if self.counter >= self.patience:
                should_stop = True
            else:
                should_stop = False

        return should_stop

    def reset(self):
        """
        Reset the early stopping counter and best score.
        """
        self.best_score = None
        self.counter = 0
```
