## Supplementary material for "pLM representations unlock metagenomic space beyond homology": Code bundle: accelerate_deepspeed_zero2.yaml

```
compute_environment: LOCAL_MACHINE
debug: false 
distributed_type: DEEPSPEED
deepspeed_config:   
  deepspeed_multinode_launcher: standard
  offload_optimizer_device: none
  offload_param_device: none
  zero3_init_flag: false
  zero_stage: 2
dynamo_backend: 'no'
downcast_bf16: 'no'
gpu_ids: all
machine_rank: 0
main_training_function: main
mixed_precision: 'no'
num_machines: 1
num_processes: 2
rdzv_backend: static
same_network: true
tpu_env: []
tpu_use_cluster: false
tpu_use_sudo: false
use_cpu: false
```
