## Supplementary material for "pLM representations unlock metagenomic space beyond homology": Code bundle: config.yaml

```
defaults:
  - _self_
  - filtering_method: cosine_sim

hydra:
  run:
    dir: outputs/${now:%Y-%m-%d-%H-%M-%S}

data:
  output_path: ${oc.env:DATA_ROOT,./data}/datasets/uniref/filtered/
  output_name: uniref100_train_no_ambig

# Directory of score shards produced by scripts/score.py.
source:
  path: ${oc.env:OUTPUT_ROOT,./outputs}/scores/sequence-scores/uniref100/
  files_list: null

# The sequence CSVs to filter, matched against the retained ids.
sequence_datasets:
  - ${oc.env:DATA_ROOT,./data}/datasets/uniref/train/uniref100_train_no_ambig.csv

# Fraction of the filtered set to keep per epoch, or null to keep everything.
# The paper's mixtures subsample the metagenomic sources at 0.03-0.4.
subsample: null
num_epochs_to_subsample: 1
```
