## Supplementary material for "pLM representations unlock metagenomic space beyond homology": Code bundle: pseudo_ppl.yaml

```
name: Pseudo PPL quantile filtering

use_cols : 
  - 'pseudo_ppl'

filter: quantile

quantiles : # TODO check quantiles
  - 0.01
  - 0.02
  - 0.05
  - 0.1
  - 0.2
  - 0.5
  - 0.9
  - 0.95
  - 0.98
  - 0.99

thresholds: null

larger_is_better: False
```
