## Supplementary material for "pLM representations unlock metagenomic space beyond homology": Code bundle: bfd100.yaml

```
train_no_ambig: False
train_max_seq_id: 1.0
validation_n: 10000
deduplicate_cov_mode: 0
deduplicate_fident: 0.9
deduplicate_sensitivity: 7.5
deduplicate_max_accept: 1000000
deduplicate_max_seqs: 1000000
deduplicate_max_rejected: 1000000
deduplicate_min_ungapped_score: 15
deduplicate_diag_score: 1
```
