## Supplementary material for "pLM representations unlock metagenomic space beyond homology": Code bundle: config.yaml

```
defaults:
  - _self_
  - uniref: uniref100
  - oas: oas90
  - mgnify: mgnify100
  - bfd: bfd100
  - uniprot: uniprot

hydra:
  run:
    dir: outputs/${now:%Y-%m-%d-%H-%M-%S}

# Valid names are the keys built in scripts/process.py:
# uniref50, uniref90, uniref100, oas, bfd, mgnify, uniprot, or "all".
datasets: ["uniref100","oas","bfd","mgnify"]

# Datasets are downloaded and processed under <base_path>/datasets/.
base_path: ${oc.env:DATA_ROOT,./data}
val_dataset_path: ${oc.env:DATA_ROOT,./data}/datasets/combined_validation.fasta.gz

# test_mode restricts each source to a small subset (e.g. 1 of 25 MGnify chunks),
# which is what you want for a first run. Set to False for the full databases.
test_mode: True
num_proc: -1
uniprot_version: "2024_05"
  
process:
  tmp_path: null
  download: False
  preprocess: False
  create_validation: False
  deduplicate: False
  chunk_for_deduplication: True
  max_sequences_per_chunk: 1000000
  clean_tmp: False
  clean_download: False
```
