## Supplementary material for "pLM representations unlock metagenomic space beyond homology": Code bundle: config.yaml

```
defaults:
  - _self_
  - model: 350M
  - dataset: uniref100

num_proc: 8
max_samples_per_file: 10000

# Root directory for the score shards. Scores are written to
# <output_path>/<scorer name>/<dataset name>/data_*.parquet
output_path: ${oc.env:OUTPUT_ROOT,./outputs}/scores

wandb:
  name: null
  project: proseqo
  entity: null
  tags: []
  dir: outputs/${now:%Y-%m-%d-%H-%M-%S}/wandb
  mode: disabled
  log_interval: 100

hydra:
  run:
    dir: outputs/${now:%Y-%m-%d-%H-%M-%S}

accelerate:
  max_checkpoints: 3

tokenizer:
  pretrained_model_name_or_path: chandar-lab/AMPLIFY_350M
  trust_remote_code: True

datacollator:
  packed: True
  pad_to_multiple_of: 8
  padding: True
  pin_memory: True
  num_workers: 8
  mask_probability: 0.15
  batch_size: 64
  samples_per_batch: 4
```
