## Supplementary material for "pLM representations unlock metagenomic space beyond homology": Code bundle: 350M.yaml

```
# AMPLIFY 350M: the model used to compute RED scores for dataset filtering.
# Authentication, if ever needed, is read from the HF_TOKEN environment variable.
trunk:
  pretrained_model_name_or_path: chandar-lab/AMPLIFY_350M
  trust_remote_code: True
  max_length: 2048
```
