## Supplementary material for "pLM representations unlock metagenomic space beyond homology": Code bundle: 350M_consensus.yaml

```
# AMPLIFY-C (AMPLIFY-consensus): the specialized model used for wet-lab candidate
# selection, embedding-based retrieval and expression analysis.
# Authentication, if ever needed, is read from the HF_TOKEN environment variable.
trunk:
  pretrained_model_name_or_path: ${oc.env:HF_ORG,flair-bio}/AMPLIFY-C
  trust_remote_code: True
  max_length: 2048
```
