## Supplementary material for "pLM representations unlock metagenomic space beyond homology": Code bundle: cosine_sim.yaml

```
# Architecture is rebuilt from the hydra config saved by the training run, and the
# weights are loaded from each DeepSpeed checkpoint under checkpoints_dir.
# Used by scripts/cosine_similarity_score.py to track RED across pretraining.
config_path: ${oc.env:OUTPUT_ROOT,./outputs}/ablations/AMPLIFY-C/.hydra/config.yaml
vocab_path: null  # set to override the vocab_path in config_path, e.g. conf/tokenizer/amplify_vocab.txt
max_length: 2048

# Directory containing DeepSpeed checkpoint subdirectories (one per training step).
checkpoints_dir: ${oc.env:OUTPUT_ROOT,./outputs}/ablations/AMPLIFY-C/model_checkpoints
```
