## Supplementary material for "pLM representations unlock metagenomic space beyond homology": Code bundle: cosine_similarity_score.sh

```
#!/bin/bash
# Sweep RED across the checkpoints of a training run, to track how residue
# embedding diversity evolves over pretraining.
#
#SBATCH --job-name=cosine_similarity_score
#SBATCH --time=1-00:00:00
#SBATCH --error=logs/%x_%j.txt
#SBATCH --output=logs/%x_%j.txt
#SBATCH --mem=32G
#SBATCH --gpus-per-task=1
#SBATCH --cpus-per-task=6
#SBATCH --nodes=1
#SBATCH --ntasks=1

set -euo pipefail

: "${DATA_ROOT:?set DATA_ROOT to the directory holding datasets/}"
: "${OUTPUT_ROOT:?set OUTPUT_ROOT to the directory holding the training run}"

export MASTER_PORT=$((10000 + ${SLURM_JOBID: -4}))
export MASTER_ADDR=$(scontrol show hostnames "$SLURM_JOB_NODELIST" | head -n 1)
export OMP_NUM_THREADS=${SLURM_CPUS_PER_TASK:-6}

# source .venv/bin/activate

MODEL_NAME=AMPLIFY-C
RUN_DIR="$OUTPUT_ROOT/ablations/$MODEL_NAME"
DATASET=bfd

accelerate launch \
    --config_file=config/accelerate_default.yaml \
    --machine_rank="${SLURM_NODEID:-0}" \
    --num_cpu_threads_per_process="${SLURM_CPUS_PER_TASK:-6}" \
    --main_process_ip="$MASTER_ADDR" \
    --main_process_port="$MASTER_PORT" \
    --num_processes=$(( ${SLURM_JOB_NUM_NODES:-1} * ${SLURM_GPUS_ON_NODE:-1} )) \
    --num_machines="${SLURM_JOB_NUM_NODES:-1}" \
    --mixed_precision=bf16 \
    scripts/cosine_similarity_score.py \
    num_proc="${SLURM_CPUS_PER_TASK:-6}" \
    output_path="$OUTPUT_ROOT/scores/cosine_similarity/$MODEL_NAME" \
    model=cosine_sim \
    "model.config_path=$RUN_DIR/.hydra/config.yaml" \
    "model.checkpoints_dir=$RUN_DIR/model_checkpoints" \
    dataset=$DATASET \
    max_samples_per_file=100000 \
    datacollator.batch_size=64 \
    datacollator.samples_per_batch=64 \
    datacollator.packed=true

# Merge the per-checkpoint shards into a single dataframe:
# python scripts/build_cosine_sim_dataframe.py \
#     --scores_dir "$OUTPUT_ROOT/scores/cosine_similarity/$MODEL_NAME/cosine-similarity" \
#     --dataset_name $DATASET \
#     --output "$OUTPUT_ROOT/scores/cosine_similarity/$MODEL_NAME/merged.parquet"
```
