## Supplementary material for "pLM representations unlock metagenomic space beyond homology": Code bundle: fasta_to_csv.sh

```
#!/bin/bash
# Convert a FASTA file to the two-column (name, sequence) CSV the scorer reads.
#
# Usage: sbatch jobs/fasta_to_csv.sh <input.fasta[.gz]> <output.csv>
#
#SBATCH --job-name=fasta_to_csv
#SBATCH --time=02:00:00
#SBATCH --error=logs/%x_%j.txt
#SBATCH --output=logs/%x_%j.txt
#SBATCH --cpus-per-task=24
#SBATCH --mem=64G

set -euo pipefail

: "${DATA_ROOT:?set DATA_ROOT to the directory holding datasets/}"

INPUT="${1:-$DATA_ROOT/datasets/uniref/train/uniref100_train_no_ambig.fasta.gz}"
OUTPUT="${2:-$DATA_ROOT/datasets/uniref/train/uniref100_train_no_ambig.csv}"

# source .venv/bin/activate

python scripts/fasta_to_csv.py --input_file "$INPUT" --output_file "$OUTPUT"

# Very large CSVs can be split into chunks for parallel scoring:
# python scripts/chunk.py --input_file "$OUTPUT" --output_dir "$(dirname "$OUTPUT")/chunks"
```
