## Supplementary material for "pLM representations unlock metagenomic space beyond homology": Code bundle: filter.sh

```
#!/bin/bash
# Filter a dataset by RED quantile, and optionally subsample the survivors.
#
# QUANTILE is the proportion of sequences to drop, lowest RED first. The paper
# uses 0.05-0.1 for UniRef100 and 0.2-0.5 for the metagenomic sources; see the
# README for the per-model settings of AMPLIFY-B and AMPLIFY-C.
#
#SBATCH --job-name=filter
#SBATCH --time=04:00:00
#SBATCH --error=logs/%x_%j.txt
#SBATCH --output=logs/%x_%j.txt
#SBATCH --cpus-per-task=16
#SBATCH --mem=128G

set -euo pipefail

: "${DATA_ROOT:?set DATA_ROOT to the directory holding datasets/}"
: "${OUTPUT_ROOT:?set OUTPUT_ROOT to the directory holding scores/}"

QUANTILE="${1:-0.1}"

# source .venv/bin/activate

python scripts/filter.py \
    filtering_method=cosine_sim \
    "source.path=$OUTPUT_ROOT/scores/sequence-scores/uniref100/" \
    "sequence_datasets=[$DATA_ROOT/datasets/uniref/train/uniref100_train_no_ambig.csv]" \
    "data.output_path=$DATA_ROOT/datasets/uniref/filtered/" \
    "data.output_name=uniref100_$QUANTILE" \
    "filtering_method.quantiles=[$QUANTILE]"

# To build a mixture, filter each source then subsample the metagenomic ones.
# subsample is the fraction kept, resampled independently for each epoch:
#     subsample=0.1 num_epochs_to_subsample=5
#
# source.path also accepts a list, to score-filter several shards at once:
#     "source.path=[$OUTPUT_ROOT/scores/sequence-scores/bfd_0,$OUTPUT_ROOT/scores/sequence-scores/bfd_1]"
```
