## Supplementary material for "pLM representations unlock metagenomic space beyond homology": Code bundle: process.sh

```
#!/bin/bash
# Download and preprocess a source database.
#
# Add your cluster's partition/account directives as needed.
#SBATCH --job-name=process
#SBATCH --time=2-00:00:00
#SBATCH --error=logs/%x_%j.txt
#SBATCH --output=logs/%x_%j.txt
#SBATCH --cpus-per-task=16
#SBATCH --mem=32G
#SBATCH --nodes=1
#SBATCH --ntasks-per-node=1

set -euo pipefail

: "${DATA_ROOT:?set DATA_ROOT to the directory that should hold datasets/}"

# source .venv/bin/activate

# aria2c (downloads) and mmseqs (deduplication) must be on PATH; see the README.
command -v aria2c >/dev/null || { echo "aria2c not found on PATH"; exit 1; }
command -v mmseqs >/dev/null || { echo "mmseqs not found on PATH"; exit 1; }

# test_mode=true restricts the download to a small subset. Set it to false for
# the full database; BFD and MGnify are multi-terabyte downloads.
python scripts/process.py \
    datasets=\["bfd"\] \
    base_path="$DATA_ROOT" \
    val_dataset_path="$DATA_ROOT/datasets/combined_validation.fasta.gz" \
    test_mode=true \
    process.download=true \
    process.preprocess=true \
    process.create_validation=false \
    process.deduplicate=false
```
