## Supplementary material for "pLM representations unlock metagenomic space beyond homology": Code bundle: README.md

# Example SLURM jobs

One script per pipeline stage, in the order they are normally run:

| Script | Stage |
| --- | --- |
| `process.sh` | download, preprocess and deduplicate a source database |
| `fasta_to_csv.sh` | convert a FASTA (optionally gzipped) to the CSV the scorer reads |
| `score.sh` | compute RED, pseudo-perplexity and embeddings with a pretrained pLM |
| `cosine_similarity_score.sh` | sweep RED across the checkpoints of a training run |
| `filter.sh` | filter by RED quantile and optionally subsample |

These are examples, not a turnkey pipeline. Every script expects two environment
variables and leaves cluster-specific choices to you:

```bash
export DATA_ROOT=/path/to/data      # holds datasets/
export OUTPUT_ROOT=/path/to/outputs # receives scores/ and filtered sets
```

The `#SBATCH` directives are deliberately minimal. Partition names, GPU
constraints and account flags are cluster-specific: add whatever your site
requires. The runs behind the paper used 1 to 8 H100 GPUs per scoring job.
