## Supplementary material for "pLM representations unlock metagenomic space beyond homology": Code bundle: score.sh

```
#!/bin/bash
# Score sequences with a pretrained pLM: RED (as mean_cosine_sim), pseudo-perplexity
# and mean-pooled embeddings, one forward pass per sequence.
#
#SBATCH --job-name=score
#SBATCH --time=1-00:00:00
#SBATCH --error=logs/%x_%j.txt
#SBATCH --output=logs/%x_%j.txt
#SBATCH --mem=32G
#SBATCH --gpus-per-task=1
#SBATCH --cpus-per-task=6
#SBATCH --nodes=1
#SBATCH --ntasks=1

set -euo pipefail

: "${DATA_ROOT:?set DATA_ROOT to the directory holding datasets/}"
: "${OUTPUT_ROOT:?set OUTPUT_ROOT to the directory that should receive scores/}"

# Unique port per job, so concurrent jobs on one node do not collide.
export MASTER_PORT=$((10000 + ${SLURM_JOBID: -4}))
export MASTER_ADDR=$(scontrol show hostnames "$SLURM_JOB_NODELIST" | head -n 1)
export OMP_NUM_THREADS=${SLURM_CPUS_PER_TASK:-6}

# source .venv/bin/activate

accelerate launch \
    --config_file=config/accelerate_default.yaml \
    --machine_rank="${SLURM_NODEID:-0}" \
    --num_cpu_threads_per_process="${SLURM_CPUS_PER_TASK:-6}" \
    --main_process_ip="$MASTER_ADDR" \
    --main_process_port="$MASTER_PORT" \
    --num_processes=$(( ${SLURM_JOB_NUM_NODES:-1} * ${SLURM_GPUS_ON_NODE:-1} )) \
    --num_machines="${SLURM_JOB_NUM_NODES:-1}" \
    --mixed_precision=bf16 \
    scripts/score.py \
    num_proc="${SLURM_CPUS_PER_TASK:-6}" \
    output_path="$OUTPUT_ROOT/scores" \
    model=350M \
    dataset=uniref100 \
    max_samples_per_file=100000 \
    datacollator.batch_size=128 \
    datacollator.samples_per_batch=128 \
    datacollator.packed=true \
    model.trunk.max_length=512

# To emit per-residue instead of mean-pooled embeddings (used for the
# residue-embedding UMAP), append:
#     +token_level=true
```
