## Supplementary material for "pLM representations unlock metagenomic space beyond homology": Code bundle: pyproject.toml

```
[build-system]
requires = ["setuptools >= 61.0"]
build-backend = "setuptools.build_meta"

[project]
name = "proseqo"
version = "0.0.0"
description = "A data preprocessing pipeline for protein language model training data"
requires-python = ">= 3.10"
dependencies = [
    "torch==2.9.0",
    "deepspeed==0.18.0",
    "wandb==0.22.2",
    "hydra-core==1.3.2",
    "numpy==2.2.6",
    "transformers==4.49.0",
    "datasets==4.2.0",
    "polars==1.34.0",
    "pandas",
    "pyarrow",
    "requests",
    "tqdm",
    "accelerate==1.11.0",
    "flash-attn==2.8.3",
    "upfp",
]

# Not installable from PyPI, and therefore not listed above:
#   - aria2, mmseqs2, seqkit and pigz are external binaries; see the README.
#   - `amplify`, imported by proseqo.scores.CosineSimilarity to rebuild a model
#     from a training checkpoint, lives in the training/ directory of this
#     repository. Install it with `pip install -e ../training` if you need the
#     checkpoint sweep; the rest of the pipeline does not require it.

authors = [
    { name = "Lola Le Breton", email = "" },
    { name = "David Heurtel-Depeiges", email = "" },
    { name = "Quentin Fournier", email = "" },
]
maintainers = [
    { name = "Lola Le Breton", email = "" },
    { name = "David Heurtel-Depeiges", email = "" },
    { name = "Quentin Fournier", email = "" },
]
license = { text = "MIT License" }
keywords = []

[project.optional-dependencies]
dev = ["black", "pytest", "pytest-cov", "pytest-clarity", "xeger"]
```
