## Supplementary material for "pLM representations unlock metagenomic space beyond homology": Code bundle: build_cosine_sim_dataframe.py

```
import os
import logging
from argparse import ArgumentParser

import polars as pl

logger = logging.getLogger(__name__)
logging.basicConfig(level=logging.INFO)


def find_checkpoints(scores_dir: str, dataset_name: str) -> list[tuple[int, str]]:
    """Return (step, checkpoint_path) pairs sorted by training step."""
    entries = []
    for name in os.listdir(scores_dir):
        checkpoint_dir = os.path.join(scores_dir, name)
        dataset_dir = os.path.join(checkpoint_dir, dataset_name)
        if not os.path.isdir(dataset_dir):
            continue
        try:
            step = int(name)
        except ValueError:
            logger.warning(f"Skipping non-numeric checkpoint directory: {name}")
            continue
        if not any(f.endswith(".parquet") for f in os.listdir(dataset_dir)):
            logger.warning(f"No parquet files found in {dataset_dir}, skipping.")
            continue
        entries.append((step, dataset_dir))

    if not entries:
        raise ValueError(f"No valid checkpoint directories found under {scores_dir} for dataset '{dataset_name}'")

    return sorted(entries, key=lambda x: x[0])


def load_checkpoint(dataset_dir: str, step: int) -> pl.LazyFrame:
    """Lazily load all shards for one checkpoint and rename the score column."""
    pattern = os.path.join(dataset_dir, "data_*.parquet")
    return (
        pl.scan_parquet(pattern)
        .select(["names", "mean_cosine_sim"])
        .rename({"mean_cosine_sim": f"step_{step}"})
    )


def build_dataframe(scores_dir: str, dataset_name: str) -> pl.DataFrame:
    checkpoints = find_checkpoints(scores_dir, dataset_name)
    logger.info(f"Found {len(checkpoints)} checkpoint(s): {[s for s, _ in checkpoints]}")

    # Build from the first checkpoint, then left-join all subsequent ones
    base_step, base_dir = checkpoints[0]
    df = load_checkpoint(base_dir, base_step)

    for step, dataset_dir in checkpoints[1:]:
        df = df.join(load_checkpoint(dataset_dir, step), on="names", how="left")

    logger.info("Collecting dataframe...")
    return df.collect()


def main(args):
    df = build_dataframe(args.scores_dir, args.dataset_name)

    n_samples, n_cols = df.shape
    n_checkpoints = n_cols - 1  # subtract the 'name' column
    logger.info(f"Merged dataframe: {n_samples} samples × {n_checkpoints} checkpoints")

    os.makedirs(os.path.dirname(os.path.abspath(args.output)), exist_ok=True)

    if args.output.endswith(".csv"):
        df.write_csv(args.output)
    else:
        df.write_parquet(args.output, compression="zstd")

    logger.info(f"Saved to {args.output}")


if __name__ == "__main__":
    parser = ArgumentParser(
        description="Merge per-checkpoint cosine similarity scores into a single dataframe."
    )
    parser.add_argument(
        "--scores_dir",
        type=str,
        required=True,
        help="Root cosine-similarity scores directory, e.g. $SCRATCH/proseqo/scores/AMPLIFY-C/cosine-similarity",
    )
    parser.add_argument(
        "--dataset_name",
        type=str,
        required=True,
        help="Dataset subdirectory name, e.g. mgnify_1M_no_ambig",
    )
    parser.add_argument(
        "--output",
        type=str,
        required=True,
        help="Output file path (.parquet or .csv)",
    )

    main(parser.parse_args())
```
