## Supplementary material for "pLM representations unlock metagenomic space beyond homology": Code bundle: chunk.py

```
from proseqo.utils import chunk_csv_file
from argparse import ArgumentParser


def main():
    parser = ArgumentParser()
    parser.add_argument("--input_file", type=str)
    parser.add_argument("--output_dir", type=str)
    args = parser.parse_args()

    chunk_csv_file(args.input_file, max_sequences_per_chunk=60000000, output_dir=args.output_dir)


if __name__ == "__main__":
    main()
```
