## Supplementary material for "pLM representations unlock metagenomic space beyond homology": Code bundle: cosine_similarity_score.py

```
import os
import logging
import hydra
from omegaconf import DictConfig

# HuggingFace
from datasets import load_dataset

# Ours
from proseqo.scores import CosineSimilarity

logger = logging.getLogger(__name__)


def _find_checkpoints(checkpoints_dir: str) -> list[str]:
    """Return sorted list of DeepSpeed checkpoint subdirectories found in checkpoints_dir."""
    if not os.path.isdir(checkpoints_dir):
        raise ValueError(f"checkpoints_dir does not exist or is not a directory: {checkpoints_dir}")

    paths = sorted(
        os.path.join(checkpoints_dir, name)
        for name in os.listdir(checkpoints_dir)
        if os.path.isdir(os.path.join(checkpoints_dir, name)) and name != "latest"
    )

    if not paths:
        raise ValueError(f"No checkpoint subdirectories found in: {checkpoints_dir}")

    return paths


@hydra.main(config_path="../config/score", config_name="config", version_base="1.1")
def main(cfg: DictConfig):
    logging.basicConfig(filename=None, level=logging.INFO)

    # model.checkpoint_path can be set to score a single checkpoint (used by the array job).
    # Otherwise, all checkpoints under model.checkpoints_dir are scored sequentially.
    single = cfg.model.get("checkpoint_path", None)
    if single is not None:
        checkpoints = [single]
        logger.info(f"Scoring single checkpoint: {single}")
    else:
        checkpoints_dir = cfg.model.checkpoints_dir
        checkpoints = _find_checkpoints(checkpoints_dir)
        logger.info(f"Found {len(checkpoints)} checkpoint(s) in {checkpoints_dir}")

    scorer = CosineSimilarity(cfg)

    dataset = load_dataset(
        "csv",
        data_files=cfg.dataset.path,
        keep_in_memory=False,
        split="train",
        streaming=False,
        num_proc=cfg.num_proc,
    )

    for checkpoint_path in checkpoints:
        checkpoint_name = os.path.basename(checkpoint_path)
        logger.info(f"Scoring with checkpoint: {checkpoint_name}")
        scorer.score(
            dataset,
            dataset_name=cfg.dataset.name,
            output_path=cfg.output_path,
            checkpoint_path=checkpoint_path,
        )
        logger.info(f"Done with checkpoint: {checkpoint_name}")


if __name__ == "__main__":
    main()
```
