## Supplementary material for "pLM representations unlock metagenomic space beyond homology": Code bundle: extract_scores_without_embeddings.py

```
"""Strip the embedding columns out of score shards.

Scoring writes the mean-pooled embedding alongside the scalar scores, which makes
the shards two orders of magnitude larger than they need to be for filtering.
This produces score-only copies, cheap to load when all you need is RED or
perplexity.
"""

import os
import logging
import pyarrow.parquet as pq
from argparse import ArgumentParser

logger = logging.getLogger(__name__)
logging.basicConfig(level=logging.INFO)


def main(args):
    os.makedirs(args.output_path, exist_ok=True)

    columns_to_drop = None
    for file in sorted(os.listdir(args.score_path)):
        if file.endswith(".parquet"):
            file_path = os.path.join(args.score_path, file)
            logger.info(f"Extracting scores from file {file_path}")

            # Load parquet file
            df = pq.read_table(file_path)

            # Drop the embedding columns, whether stored as one list column
            # ("embeddings", "token_embeddings_per_layer") or as one column per
            # dimension ("embedding_0", "embedding_1", ...).
            if columns_to_drop is None:
                columns_to_drop = [col for col in df.column_names if "embedding" in col]
                logger.info(f"Dropping columns: {columns_to_drop}")
            df = df.drop(columns_to_drop)

            # Save to parquet
            pq.write_table(df, os.path.join(args.output_path, file))


if __name__ == "__main__":
    args = ArgumentParser(description="Write score shards without their embedding columns.")
    args.add_argument("--score_path", type=str, required=True, help="Directory of score shards")
    args.add_argument("--output_path", type=str, required=True, help="Directory for the stripped copies")
    args = args.parse_args()

    main(args)
```
