## Supplementary material for "pLM representations unlock metagenomic space beyond homology": Code bundle: fasta_to_csv.py

```
from proseqo.utils import fasta_to_csv
from argparse import ArgumentParser


def main():
    parser = ArgumentParser()
    parser.add_argument("--input_file", type=str)
    parser.add_argument("--output_file", type=str)
    args = parser.parse_args()

    fasta_to_csv(args.input_file, args.output_file)


if __name__ == "__main__":
    main()
```
