## Supplementary material for "pLM representations unlock metagenomic space beyond homology": Code bundle: filter.py

```
import pandas as pd
import os
import random
import pyarrow as pa
import pyarrow.dataset as ds
import pyarrow.compute as pc
import pyarrow.parquet as pq
import pyarrow.csv as pcsv
from omegaconf import DictConfig, OmegaConf, ListConfig
import logging
import hydra
from abc import ABC, abstractmethod

logger = logging.getLogger(__name__)


def compute_quantiles(df: pd.DataFrame, metric: str, quantiles: list):
    quantiles = df[metric].quantile(quantiles)
    return quantiles


class Filter(ABC):
    """
    Abstract class for filtering.

    Filter the dataframe based on some criteria and saves the resulting sequence_ids (with the scores) in a file.
    """

    def __init__(self, use_cols: list, logger: logging.Logger = logger):
        self.use_cols = use_cols
        self.logger = logger

    @abstractmethod
    def __call__(self, df: pd.DataFrame, **kwargs) -> None:
        pass


class QuantileFilter(Filter):
    """
    Filter based on a list of quantiles.
    """

    def __init__(self, use_cols: list, quantiles: list, output_paths: list, larger_is_better: bool = True, logger: logging.Logger = logger):
        super().__init__(use_cols, logger)
        assert len(self.use_cols) == 1, "Only one score is allowed"
        self.quantiles = quantiles
        self.output_paths = output_paths
        assert len(self.output_paths) == len(self.quantiles), "Number of output paths must be equal to the number of quantiles"
        self.larger_is_better = larger_is_better

    def __call__(self, table: pa.Table):
        score_col = self.use_cols[0]

        self.logger.info("Computing quantiles")
        # Compute the quantiles of the table
        if not self.larger_is_better:
            quantiles = [1 - q for q in self.quantiles]
        else:
            quantiles = self.quantiles

        quantiles_values = pc.quantile(pc.cast(table.column(score_col), pa.float64()), quantiles)
        quantiles_values = [q.as_py() for q in quantiles_values]

        self.logger.info("Quantiles values: " + str(quantiles_values))
        for i, threshold in enumerate(quantiles_values):
            # get output_file extension
            self.logger.info("Filtering with threshold: " + str(threshold))
            if not self.larger_is_better:
                filtered_table = table.filter(pc.less_equal(pc.cast(table.column(score_col), pa.float64()), threshold))
            else:
                filtered_table = table.filter(pc.greater_equal(pc.cast(table.column(score_col), pa.float64()), threshold))

            _, ext = os.path.splitext(self.output_paths[i])
            if ext == ".parquet":
                pq.write_table(filtered_table, self.output_paths[i], compression="snappy")
            elif ext == ".csv":
                pcsv.write_csv(filtered_table, self.output_paths[i])
            else:
                raise ValueError("Output file extension not supported")


class ThresholdsFilter(Filter):
    """
    Filter based on a list of thresholds.
    """

    def __init__(
        self, use_cols: list, thresholds: list, output_paths: list, larger_is_better: bool = True, logger: logging.Logger = logger
    ):
        super().__init__(use_cols, logger)
        assert len(self.use_cols) == 1, "Only one score is allowed"
        self.thresholds = thresholds
        self.output_paths = output_paths
        assert len(self.output_paths) == len(self.thresholds), "Number of output paths must be equal to the number of thresholds"
        self.larger_is_better = larger_is_better

    def __call__(self, table: pa.Table):
        score_col = self.use_cols[0]

        for i, threshold in enumerate(self.thresholds):
            # get output_file extension
            if not self.larger_is_better:
                filtered_table = table.filter(pc.less_equal(pc.cast(table.column(score_col), pa.float64()), threshold))
            else:
                filtered_table = table.filter(pc.greater_equal(pc.cast(table.column(score_col), pa.float64()), threshold))

            _, ext = os.path.splitext(self.output_paths[i])
            if ext == ".parquet":
                pq.write_table(filtered_table, self.output_paths[i], compression="snappy")
            elif ext == ".csv":
                pcsv.write_csv(filtered_table, self.output_paths[i])
            else:
                raise ValueError("Output file extension not supported")


def get_filter(cfg: DictConfig, base_output_path: str, output_name: str, logger: logging.Logger = logger) -> Filter:
    """
    Get the filter based on the configuration.
    """
    if cfg.filtering_method.filter == "quantile":
        output_paths = [os.path.join(base_output_path, output_name + "_info" + ".parquet") for q in cfg.filtering_method.quantiles]

        return (
            QuantileFilter(
                use_cols=cfg.filtering_method.use_cols,
                quantiles=cfg.filtering_method.quantiles,
                larger_is_better=cfg.filtering_method.larger_is_better,
                output_paths=output_paths,
                logger=logger,
            ),
            output_paths,
        )

    elif cfg.filtering_method.filter == "threshold":
        output_paths = [os.path.join(base_output_path, output_name + "_info" + ".parquet") for t in cfg.filtering_method.thresholds]

        return (
            ThresholdsFilter(
                use_cols=cfg.filtering_method.use_cols,
                thresholds=cfg.filtering_method.thresholds,
                larger_is_better=cfg.filtering_method.larger_is_better,
                output_paths=output_paths,
                logger=logger,
            ),
            output_paths,
        )


@hydra.main(config_path="../config/filter", config_name="config", version_base="1.1")
def main(cfg: DictConfig):
    logging.basicConfig(filename=None, level=logging.INFO)
    os.makedirs(cfg.data.output_path, exist_ok=True)

    logger.info("Configuration :" + str(OmegaConf.to_object(cfg)))
    logger.info("Starting filtering")
    logger.info("Reading Scores")

    use_cols = list(cfg.filtering_method.use_cols)

    ## Load dataset as PyArrow dataset (avoids Pandas overhead)
    if cfg.source.path is not None:
        if isinstance(cfg.source.path, ListConfig):
            path = OmegaConf.to_container(cfg.source.path, resolve=True)  # convert to python list
            # Get the schema from the first path
            first_dataset = ds.dataset(path[0], format="parquet")
            schema = first_dataset.schema
            # Changes the schema to use large_string for "names" column
            # Create new list of fields, replacing only 'names' with large_string
            new_fields = [pa.field("names", pa.large_string()) if f.name == "names" else f for f in schema]

            # Construct new schema
            new_schema = pa.schema(new_fields)
            dataset = ds.dataset([ds.dataset(p, format="parquet", schema=new_schema) for p in path])
        else:
            print("Loading dataset from path: " + cfg.source.path)
            dataset = ds.dataset(cfg.source.path, format="parquet")
    elif cfg.source.files_list is not None:
        dataset = ds.dataset(cfg.source.files_list, format="parquet")
    else:
        raise ValueError("No source defined")

    table = dataset.to_table(columns=["names"] + use_cols)

    logger.info("Table shape: " + str(table.shape))
    logger.info("Scores loaded")
    logger.info("Filtering method: " + cfg.filtering_method.name)

    # Get the filter
    filter, filtered_ids_paths = get_filter(cfg, cfg.data.output_path, cfg.data.output_name, logger)

    # Save filtered sequence ids to filtered_ids_paths
    logger.info("Applying filter and saving filtered ids to <dataset>_info_<quantile>.parquet files")
    filter(table)

    logger.info("Matching with the sequence dataset")

    # Open array of filtered sequence ids
    filtered_ids = [ds.dataset(output_path, format="parquet").to_table(columns=["names"]).column(0) for output_path in filtered_ids_paths]

    # Iterate over chunks of the dataset and write filtered sequences to csv
    for i, dataset in enumerate(cfg.sequence_datasets):
        logger.info(f"Processing dataset {i}, {dataset}")
        table = pcsv.read_csv(
            dataset, convert_options=pcsv.ConvertOptions(column_types={"name": pa.large_string(), "sequence": pa.large_string()})
        )
        # Iterate over filtering quantiles or thresholds
        for j, (ids_list, id_path) in enumerate(zip(filtered_ids, filtered_ids_paths)):
            output_dir = os.path.dirname(id_path)
            output_file = os.path.basename(id_path)
            output_file = (
                output_file.replace("info", "sequences").replace(".parquet", f"_{i}.csv")
                if len(cfg.sequence_datasets) > 1
                else output_file.replace("info", "sequences").replace(".parquet", f".csv")
            )
            filtered_table = table.filter(pc.is_in(table.column("name"), ids_list))
            if "subsample" in cfg and cfg.subsample is not None and filtered_table.num_rows > 0:
                num_epochs_to_subsample = cfg.get("num_epochs_to_subsample", 1)
                for epoch in range(num_epochs_to_subsample):
                    logger.info(
                        f"Subsampling the sequences for dataset {i}, epoch {epoch}, subsample rate {cfg.subsample} from {filtered_table.num_rows} to {int(cfg.subsample * filtered_table.num_rows)}"
                    )
                    random.seed(epoch)
                    tmp = filtered_table.take(random.sample(range(filtered_table.num_rows), k=int(cfg.subsample * filtered_table.num_rows)))
                    os.makedirs(os.path.join(output_dir, f"epoch_{epoch}"), exist_ok=True)
                    pcsv.write_csv(
                        tmp,
                        os.path.join(output_dir, f"epoch_{epoch}", output_file),
                        write_options=pcsv.WriteOptions(include_header=True, quoting_style="none"),
                    )

            else:
                pcsv.write_csv(
                    filtered_table,
                    os.path.join(output_dir, output_file),
                    write_options=pcsv.WriteOptions(include_header=True, quoting_style="none"),
                )

    logger.info("Done")


if __name__ == "__main__":
    main()
```
