## Supplementary material for "pLM representations unlock metagenomic space beyond homology": Code bundle: filter_bfd_headers.py

```
import polars as pl
import os
import argparse
import gzip
from tqdm import tqdm

if __name__ == "__main__":
    # Parse command-line arguments
    parser = argparse.ArgumentParser()
    parser.add_argument("--input_path", type=str, default="scores/bfd/multi-combined-scores/bfd")
    parser.add_argument("--output_path", type=str, default="scores/bfd/multi-combined-scores/bfd")
    parser.add_argument("--sequences_path", type=str, default="bfd_v2_headers.sorted.txt.gz")
    args = parser.parse_args()

    with gzip.open(args.sequences_path, "rt") as f:
        sequences_to_filter = [line.strip() for line in f]

    filter_df = pl.DataFrame({"names": sequences_to_filter})
    del sequences_to_filter
    print("sequences loaded")

    for file in tqdm(os.listdir(args.input_path), desc="Filtering chunks", unit_scale=True):
        if file.endswith(".parquet"):
            scores = pl.read_parquet(os.path.join(args.input_path, file))

            # If you read parquet in chunks
            scores = pl.read_parquet("scores.parquet")
            filtered = scores.join(filter_df, on="names", how="inner").sink_parquet(
                os.path.join(args.output_path, file.replace(".parquet", ".filtered.parquet")), maintain_order=False
            )
```
