## Supplementary material for "pLM representations unlock metagenomic space beyond homology": Code bundle: README.md

# pLM representations unlock metagenomic space beyond homology

This repository holds the code accompanying the paper *pLM representations unlock metagenomic space beyond homology*.

 Metagenomic sequencing has uncovered billions of proteins from uncultured microorganisms, vastly expanding the known protein space. Yet most remain functionally inaccessible because existing annotation methods depend on close homologs or accurate structure predictions. Here, we show that protein language models (pLMs) can unlock this diversity only when their training data are appropriately curated. We introduce Residue Embedding Diversity (RED), a metric for protein quality assessment orders of magnitude cheaper than likelihood, and a calibration task that measures model alignment with natural evolutionary distributions. We discover a fundamental trade-off between evolutionary calibration and structural modeling, establishing training data composition as a primary determinant of pLM behavior. Finally, we successfully retrieve diverse enzyme candidates from billions of metagenomic sequences and validate their expression in vivo.

## Incoming release

Note that these repositories are currently undergoing a large code refacto and will be released soon as updated packages.

## Layout

| Directory | Role |
| --- | --- |
| [`processing/`](processing/) | Builds the pretraining datasets: downloads and normalizes UniRef, BFD, MGnify and OAS, scores every sequence with RED, filters on that score, and assembles the mixtures. |
| [`training/`](training/) | Pretrains the models. A fork of the public [`chandar-lab/AMPLIFY`](https://github.com/chandar-lab/AMPLIFY) codebase. |
| [`evaluation/`](evaluation/) | In-silico benchmarks: the categorical-Jacobian contact task (CASP16), the calibration task, and the supervised xTrimo property-prediction suite. `notebooks/paper_figures.ipynb` produces the paper figures. See the official ProteinGym repository for DMS evaluation code. |
| [`enzymes/`](enzymes/) | The wet-lab discovery campaign: EC 1.14.11 candidate filtering and MMR selection, metagenomic retrieval from BFD/MGnify, and expression analysis. |

## Install

Each directory is a standalone package:

```bash
cd <directory>
python -m venv env && source env/bin/activate
pip install -e .
```

Several steps additionally require external binaries or manually-installed tools (MMseqs2, hh-suite, SignalP, ProteinMPNN, ESM-IF, OpenFold). Each directory's README lists what it needs.

## Data

The source databases (UniRef, BFD, MGnify, OAS, ProteinGym, CASP16, xTrimo) are public and are individually downloaded by the pipelines.

## License

MIT — see [LICENSE](LICENSE). `training/` and `evaluation/` additionally carry the upstream AMPLIFY MIT license for fork attribution.
